# Correction of a Factor VIII genomic inversion with designer-recombinases

**DOI:** 10.1101/2020.11.02.328013

**Authors:** Felix Lansing, Liliya Mukhametzyanova, Teresa Rojo-Romanos, Kentaro Iwasawa, Masaki Kimura, Maciej Paszkowski-Rogacz, Janet Karpinski, Tobias Grass, Jan Sonntag, Paul Martin Schneider, Ceren Günes, Jenna Hoersten, Lukas Theo Schmitt, Natalia Rodriguez-Muela, Ralf Knöfler, Takanori Takebe, Frank Buchholz

## Abstract

Despite advances in nuclease-based genome editing technologies, correcting human disease-causing genomic inversions remains a challenge. Here, we describe the potential use of a recombinase-based system to correct a 140 kb genomic inversion of the F8 gene, which is frequently found in patients diagnosed with severe Hemophilia A. Employing substrate-linked directed molecular evolution, we developed a fused heterodimeric recombinase system (RecF8) achieving 30% inversion of the target sequence in human tissue culture cells. Transient RecF8 treatment of endothelial cells, differentiated from patient derived induced pluripotent stem cells (iPSCs) of a hemophilic donor, resulted in prominent correction of the inversion and restored Factor VIII mRNA expression. Our data suggests that designer-recombinases may represent efficient and specific means towards treatment of monogenic diseases caused by large gene inversions.

## Introduction

Today, patients with monogenic diseases are primarily treated with palliative care. However, within the past decades, disease-causing mutations have been identified in over 8,000 monogenic diseases allowing for the development of cure strategies that target the underlying genetic defects ^1,2^. Nuclease-based genome editing approaches can be used to correct the mutations for many of these genetic diseases ^3–5^. Nevertheless, genetic disorders caused by large genomic alterations, such as gene inversions, remain a challenge. Progress has been reported to correct gene inversions with nuclease-based approaches ^6,7^ but there are several drawbacks hampering the applied use of nucleases for these genetic alterations.

In order to repair DNA inversions with nucleases, two cuts have to be introduced into the genome, releasing the DNA fragment and subsequently relying on the fragment reinserting in the opposite orientation. This reversion is caused passively by the cell’s DNA repair machinery and not actively by the nuclease itself. Therefore, editing efficiencies rely on an effective DNA repair machinery, which is most active in mitotic cells and typically inefficient in postmitotic cells. Most certainly a nuclease-based treatment to correct inversions will require a cell-based therapy, where the correction and screening of the patient-derived cells would have to be performed *ex vivo* in mitotic cells ^7^. Additionally, it is largely unpredictable how the cell is going to repair the introduced lesions, providing a risk for sequence alterations such as small insertions/deletions (indels), thereby increasing the likelihood of unwanted mutations after the genome editing process. Furthermore, nuclease-based edited clones may also carry other undesirable small and large genomic rearrangements ^5,8,9^.

An alternative genome engineering tool with the potential to invert DNA in mitotic and postmitotic cells are site-specific recombinases such as the tyrosine recombinase Cre/loxP system ^10,11^. Cre can precisely invert genomic sequences in model organisms when the loxP target sites are aligned on the same chromosome in an inverted orientation ^12^. To overcome the native target preference of Cre, the methods of directed molecular evolution can be applied to develop novel Cre-type recombinases to recognize and actively recombine desired target sites ^13,14^, with first examples of evolved Cre-type recombinases demonstrating their broader potential for therapeutic applications ^15,16^.

We therefore reasoned that designer-recombinases could be suitable to correct genomic inversions involved in monogenic diseases. To test this hypothesis, we decided to evolve recombinases targeting sequences that are implicated in a disease-causing inversion of the F8 gene. The resulting disease is a severe form of a blood clotting defect (Hemophilia A) where no Factor VIII activity is detectable. This inversion is caused by an intrachromosomal recombination event of two homologous regions found on the X-chromosome. One repeat is located in intron 1 (int1h-1) of the F8 gene and the other is located 140 kb upstream (int1h-2), outside the F8 gene. This inversion displaces exon 1 roughly 140 kb upstream of the F8 gene and fully disrupts Factor VIII expression ^17,18^. Restoring 2-5% of the Factor VIII activity is thought to be sufficient to alleviate severe Hemophilia A symptoms ^19^, indicating that methods able to correct a moderate number of cells in the body could potentially result in clinical benefits.

## Results

### Identification of loxF8 target site and directed evolution of F8 recombinases

The int1h homologous regions are nearly perfect inverted repeats of 1,041 bp harboring only one mismatch. Therefore, almost any target site for site specific recombinases within these homologous regions would theoretically qualify as an inversion substrate for the recombination reaction. Typical target sites for Cre-type recombinases consist of 13 bp inverted repeats flanking an 8 bp spacer sequence ^10^. The ideal target sequence for Cre-type recombinases would consist of a perfect inverted repeat left and right from the spacer. This perfect sequence pattern does not exist within the int1h-repeats. However, evolved Cre-type recombinases can be coaxed to recombine target sites with some asymmetry ^15,16^. Therefore, the int1h sequences were searched for target sites with tolerable asymmetry between possible half-sites. Systematically searching through the 1,041 bp int1h sequences revealed 82 potential target sites (Fig. 1a, Supplementary Fig. 1, Supplementary Table 1). We chose the candidate sequence with the highest score (hereafter referred to as loxF8) of this list as a target for the evolution of Cre-type recombinases. The loxF8 site has 6 asymmetric positions and is located between base pairs 76-109 in the int1h regions (Fig. 1a).

**Figure 1:**
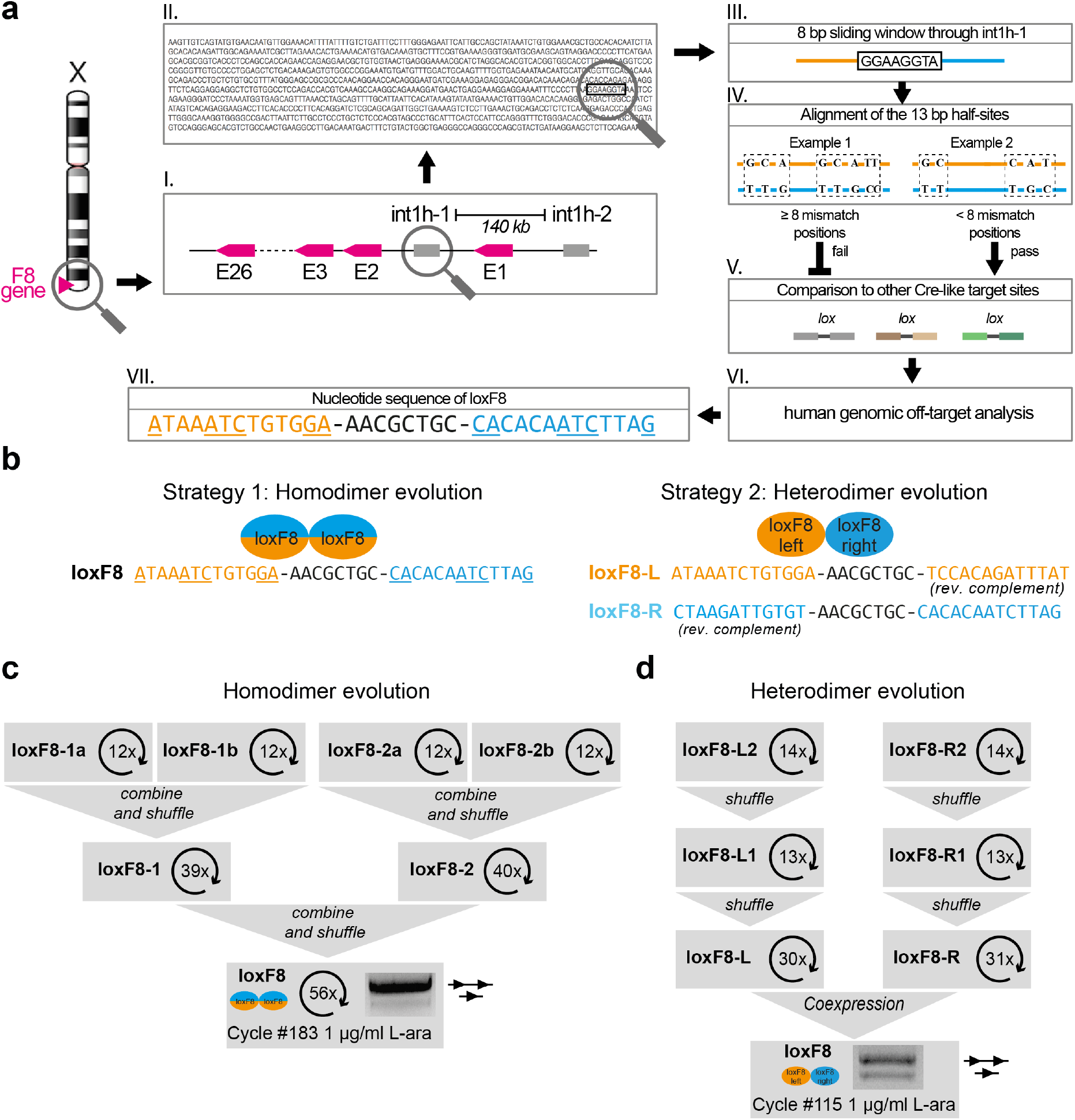
Directed evolution of site specific recombinases for the loxF8 target site. a) Schematic overview of the target site identification in the inverted repeats (int1h) surrounding the F8 gene. For each 8 bp sequence in int1h-1 the surrounding 13 bp were compared. Every 13 bp sequence pair with less than 8 mismatch positions was analyzed for similarity to previous target sites targeted by evolved Cre-type recombinases. After off-target analysis the loxF8 target site was nominated. Underlined nucleotides display asymmetric positions. The nucleotide sequence of the left (orange) and the right (blue) half sites are indicated. b) Strategies to evolve either a homodimer or a heterodimer to target the loxF8 site. The monomer targets the left, as well as the right half-site sequence (illustrated by hybrid orange-blue ellipses). The monomers of the heterodimer recognize either the left (loxF8-L, orange) or the right (loxF8-R, blue) half-site of the loxF8 target site. c) Schematic overview of the evolution of a monomer for the loxF8 target site. Evolutions for the subsites loxF8-1a, loxF8-1b, loxF8-2a and loxF8-2b were stepwise collapsed via loxF8-1 and loxF8-2 to finally obtain a recombinase library active on the loxF8 target site. The numbers indicate the evolutions cycles needed for each subsite. In total 183 evolution cycles were needed to acquire an active library that recombines loxF8 at low recombinase induction levels (1 μg/ml L-ara). The gel of the final library is shown where the upper band represents the unrecombined plasmid (illustrated by a line with two triangles) and the lower band showing the recombined plasmid (a line with one triangle). d) Schematic overview of the evolution of two monomers that can be combined as heterodimers targeting the loxF8 site. Independent evolutions were performed on the subsites loxF8-L2, loxF8-L1, loxF8-L, loxF8-R2, loxF8-R1 and loxF8-R. Libraries with activity on the loxF8-L (orange) and loxF8-R (blue) were co-expressed to recombine loxF8. In total 115 evolution cycles were needed to acquire an active library that recombines loxF8 at low recombinase induction levels (1 μg/ml L-ara). The gel of the final library is shown where the upper band represents the unrecombined plasmid (illustrated by a line with two triangles) and the lower band showing the recombined plasmid (a line with one triangle).

Selecting an asymmetric target site provides the opportunity to compare two different evolution strategies. A single recombinase can be evolved to recognize both 13 bp half-sites or two recombinases can be evolved in parallel for each 13 bp half-site (Fig. 1b). Combining the two recombinases allows to form a functional heterodimer capable of recombining the asymmetric site ^20^. To compare the two approaches, we performed two parallel substrate-linked directed evolution (SLiDE) ^13^ experiments in *E. coli* to either generate single or dual-recombinases that recombine loxF8 (Fig. 1b). For the single recombinase, 6 subsites and 183 rounds of SLiDE were required to obtain recombinase libraries with activity on the loxF8 target site (Fig. 1c, Supplementary Fig. 1).

In order to evolve the dual-recombinases to cooperatively recombine the loxF8 site as heterodimers, two separate evolutions were initiated. Two recombinase libraries were evolved for either the left (loxF8-L) or the right (loxF8-R) half-site of the loxF8 site (Fig. 1b). SLiDE evolution through 3 subsites each (Supplementary Fig. 1) and 115 rounds of directed evolution generated two libraries with activity on loxF8-L or loxF8-R, respectively. Hence, 68 evolution cycles less were required for the dual-recombinase approach. Importantly, only upon co-expression of both recombinase libraries, efficient recombination occurred on the asymmetric loxF8 target site (Fig. 1d), demonstrating that the participation of both monomers was required to recombine loxF8. Because the recombination activity was higher for the heterodimer recombinases than the monomers, we continued with the heterodimer libraries for further experiments (Fig. 1c and d).

### Selection of a heterodimer recombinase

In order to select a candidate heterodimer recombinase, we screened for recombinases that were well tolerated in human cells. The loxF8-L and loxF8-R libraries were expressed for 21 days in HeLa cells utilizing a doxycycline inducible lentiviral system (Fig. 2a). Recombinases were subsequently retrieved from genomic DNA and their activity was confirmed in *E. coli* (Supplementary Fig. 2). Next, 96 single heterodimer clones were analyzed by PCR for their activity on the loxF8 site (Fig. 2b). The heterodimer clone (D7) showed the best activity on the loxF8 site and was chosen as a candidate for detailed examination. D7 showed an induction dependent recombination activity on the loxF8 target site with nearly full recombination at 100 μg/ml L-arabinose (Fig. 2b).

Plotting the amino acid differences found in D7 onto a co-crystal structure of Cre bound to loxP ^21^, showed that the majority of the altered amino acid reside in proximity to DNA (Fig. 2c), consistent with their role in influencing DNA-binding specificity ^10,22^. The overall comparison of the amino acid sequence of the two D7 monomers to Cre revealed 46 changes (13%) in the left monomer and 39 changes (11%) in the right monomer (Fig. 2d), suggesting that considerable changes in the protein sequence were necessary to achieve high activity on the new target site. Interestingly, some of the common changes have previously been found in evolved Cre-type recombinases (Y77H, S108G, I166V, A175S, N317T and I320S) (Supplementary Fig. 2), indicating that these residues might be beneficial for evolved Cre-type recombinases in general, or that these changes are potentially important for the stability of the enzymes, as suggested by a previous study ^23^. For both monomers most changes were acquired in helix J (5-7 changes), suggesting that this helix plays a pivotal role for DNA-binding specificity. Indeed, this helix interacts with the DNA major grove near the base pairs 10 and 11 and is important for binding of Cre to loxP ^22^. Interestingly, several amino acid substitutions in helix J differ between the left and the right monomer further supporting the idea that this helix is crucial for DNA-binding selectivity (Fig. 2d). Altogether, the mutational data provide a starting point for a more rational understanding for DNA-binding specificity of designer-recombinases.

**Figure 2:**
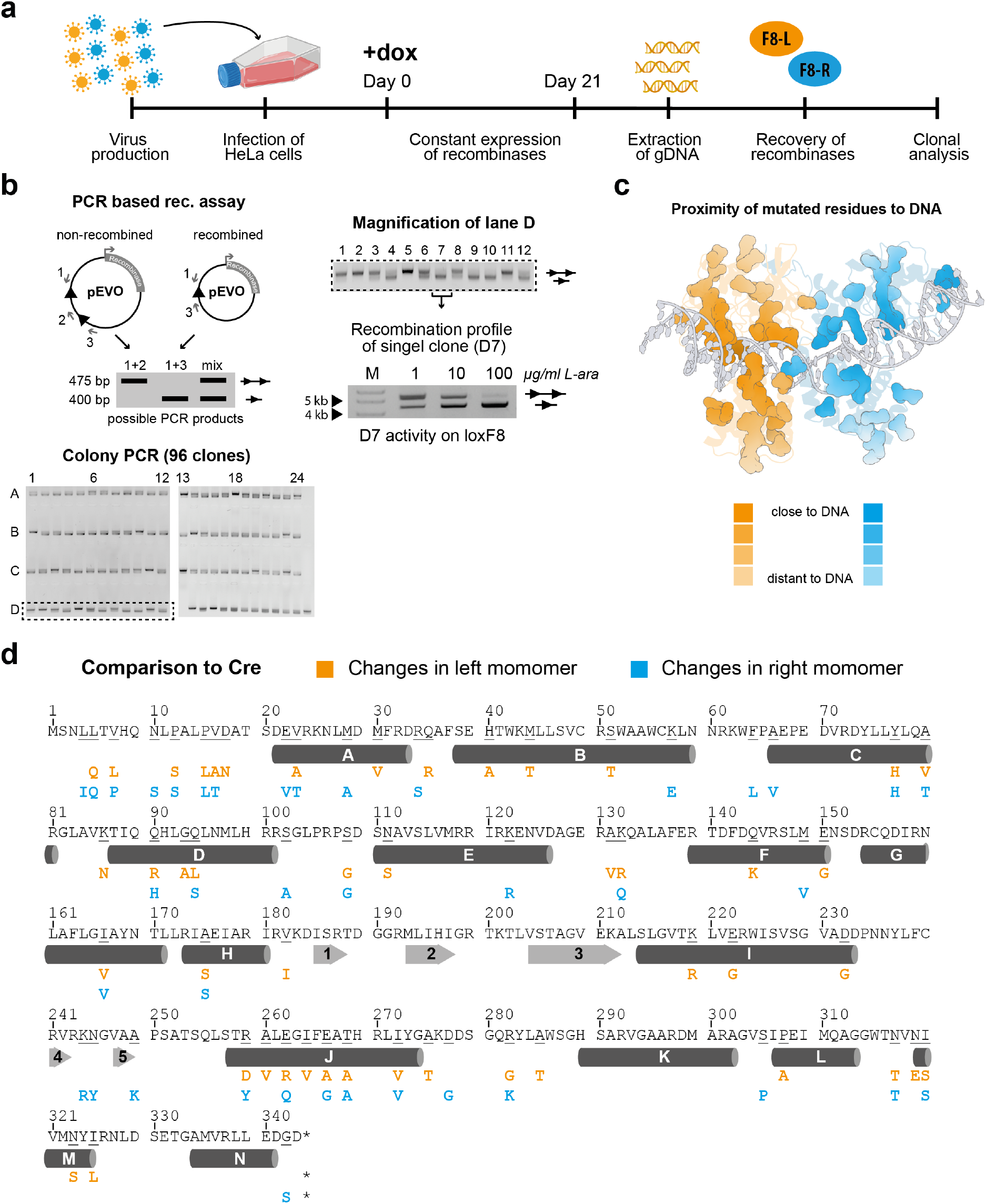
Identification of a heterodimer clone with robust activity on the loxF8 site. a) Schematic overview of the tolerance screen of recombinase clones in human cells. Both recombinases libraries were integrated and expressed for 21 days in HeLa cells under constant recombinase expression. Recombinase libraries were retrieved from genomic DNA. b) The PCR based recombination assay is schematically shown with the possible amplicons if non-recombined pEVO plasmid (475 bp), recombined pEVO plasmid (400 bp) or a mix of both plasmids is used as a template for the PCR. Grey numbered arrows show the primer binding sites and black triangles show the loxF8 target site in the pEVO vector. Ninety-six heterodimer clones were expressed in E. coli and analyzed with this assay. A magnification of lane D is presented, harboring the best clone identified (lane D clone 7). D7 was validated in a plasmid-based assay at indicated L-arabinose concentrations. The marker (M) bands at 4 kb and 5 kb are indicated with unrecombined and recombined plasmids indicated by lines with one, or two triangles, respectively. c) Mapping of mutations to a Cre protein structure bound to loxP. Residues that have changed during the evolution in the D7 heterodimer are highlighted with the intensities of the color representing their proximity to DNA (dark = close, faint = far). Changes found in the left monomer are shown in orange and changes found in the right monomer are shown in blue. The protein structure was colored and adapted using Protein Imager (3dproteinimaging.com) ^52^. d) Mutation analysis of the D7 monomers (one-letter code). The amino acid sequence of Cre recombinase is shown as a reference. The orange letters represent changes found in the left monomer and the blue letters represent changes found in the right monomer. Secondary structure elements are indicated, with alpha-helices displayed as cylinders with letters and beta-sheets represented as numbered arrows. Asterisks denote stop codons. Parts of the figure were created with BioRender.com.

### D7 recombines loxF8 in human cells

In order to test the activity of the D7 heterodimer in human cells, a HEK293T^loxF8^ reporter cell line was generated (Supplementary Fig. 3), which expresses the fluorescent mCherry protein after successful recombination of the loxF8 target sites (Fig. 3a). HEK293T^loxF8^ cells were co-transfected with mRNA coding for the recombinases together with tagBFP mRNA (Fig. 3b, Supplementary Fig. 3), to evaluate whether mRNA delivery is a suitable method to express recombinases in human cells. Only cells transfected with both monomers showed expression of mCherry, demonstrating that heterodimer formation is required to achieve recombination (Fig. 3 c). Analysis of the FACS data showed that the heterodimer recombined ~76% of transfected HEK293T^loxF8^ reporter cells (Fig. 3d, Supplementary Fig. 3), signifying the robust recombination efficiency achieved by mRNA-mediated D7 delivery.

**Figure 3:**
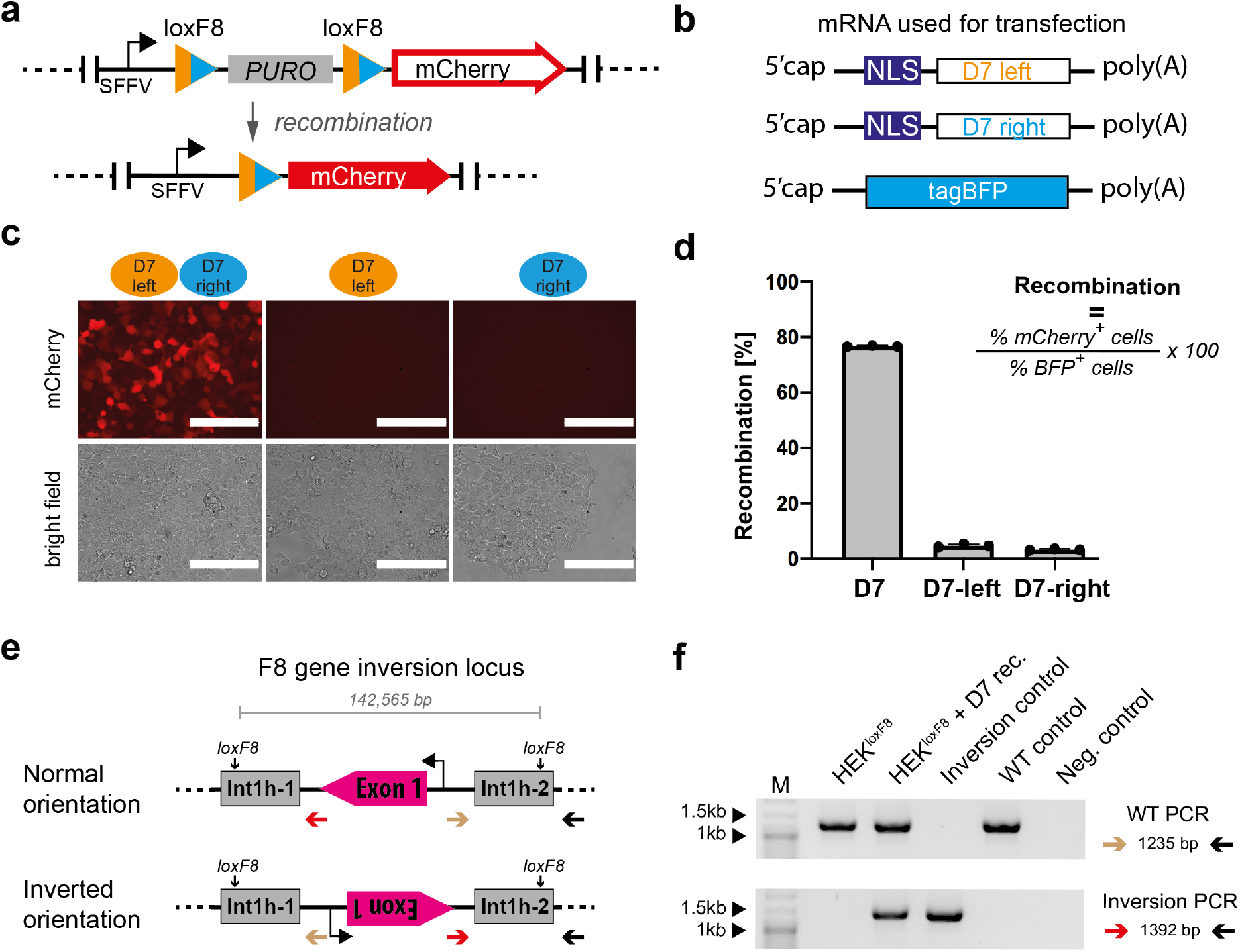
Activity of the D7 heterodimer in human cells. a) Schematic overview of the integrated reporter construct in HEK293T cells. Site-specific recombination excises the spleen focus forming virus (SFFV) promoter-driven puromycin resistance gene (PURO), which leads to expression of the red-fluorescent mCherry gene. b) Depiction of the mRNAs that were transfected for transient expression of the recombinase dimer. tagBFP mRNA was used to monitor transfection efficiencies. NLS, nuclear localization signal. c) Fluorescent and brightfield images of transfected HEK293T^loxF8^ reporter cells with indicated recombinases. Note that mCherry expression is only visible in cells receiving both monomers. 200 μm white scale bars are indicated. d) Quantification of the recombination efficiencies 48 h after transfection of HEK293T^loxF8^ reporter cells with indicated recombinases, analyzed by FACS (n=3, biological replicates are shown as dots). Error bars represent standard deviation of the mean (SD). Recombination efficiencies were calculated based on the presented formula. e) Schematic overview of a fraction of the F8 gene displaying the PCR primers used to detect the orientation of the loxF8 locus. Exons are displayed in magenta and the repeated regions int1h-1 and int1h-2 are shown in grey. Primer binding sites are indicated with arrows. The transcription start site is depicted by a black arrow. f) Gel image of PCR products generated using indicated primer combinations to detect the orientation of the loxF8 locus with and without treatment with the D7 heterodimer. Band sizes of the marker lane (M) are indicated.

Because HEK293T^loxF8^ cells carry the F8 gene in the wild type orientation, recombinase expression could potentially invert the locus into the disease orientation. To test for this possibility, genomic DNA was extracted from transfected cells and analyzed for the conceivable genomic inversion of the loxF8 locus. Two PCR assays were designed at the inversion sites to detect the orientation of the DNA fragment between the endogenous loxF8 target sites (Fig. 3e). Genomic DNA form a male donor with the F8 int1h inversion was used as a positive control. Indeed, only the HEK293T^loxF8^ cells expressing the D7 heterodimer showed the inversion specific band of the expected size (Fig. 3f). Sequencing of the PCR product confirmed that expression of the D7 recombinase led to the int1h inversion of the loxF8 locus in these cells at nucleotide precision. We conclude that a heterodimer of evolved recombinases can invert a 140 kb target sequence found in the human genome.

### Identification of potential human genomic off-targets

Because the D7 heterodimer consists of two recombinases, which can form either a heterodimer or two different homodimers, the amount of potential recognition sequences is increased and could result in the increased chance of unintended recombination at off-target sites. To identify potential off-targets, we bioinformatically analyzed the human genome for sequences with similarity to loxF8, loxF8-L and loxF8-R (Supplementary Fig. 4, Supplementary Table 2) ^16,24^. Thirteen predicted human off-target sites were cloned into the pEVO reporter vector and tested in *E. Coli*. Eleven of these predicted human off-target-sites were not recombined by the D7 heterodimer, showing that D7 has high target-site selectivity. However, expression of the recombinase dimer revealed that one asymmetric (HG2) and one symmetric (HG2L) off-target sequence were indeed recombined by D7, albeit at lower efficiency than the loxF8 site (Fig. 4a).

**Figure 4:**
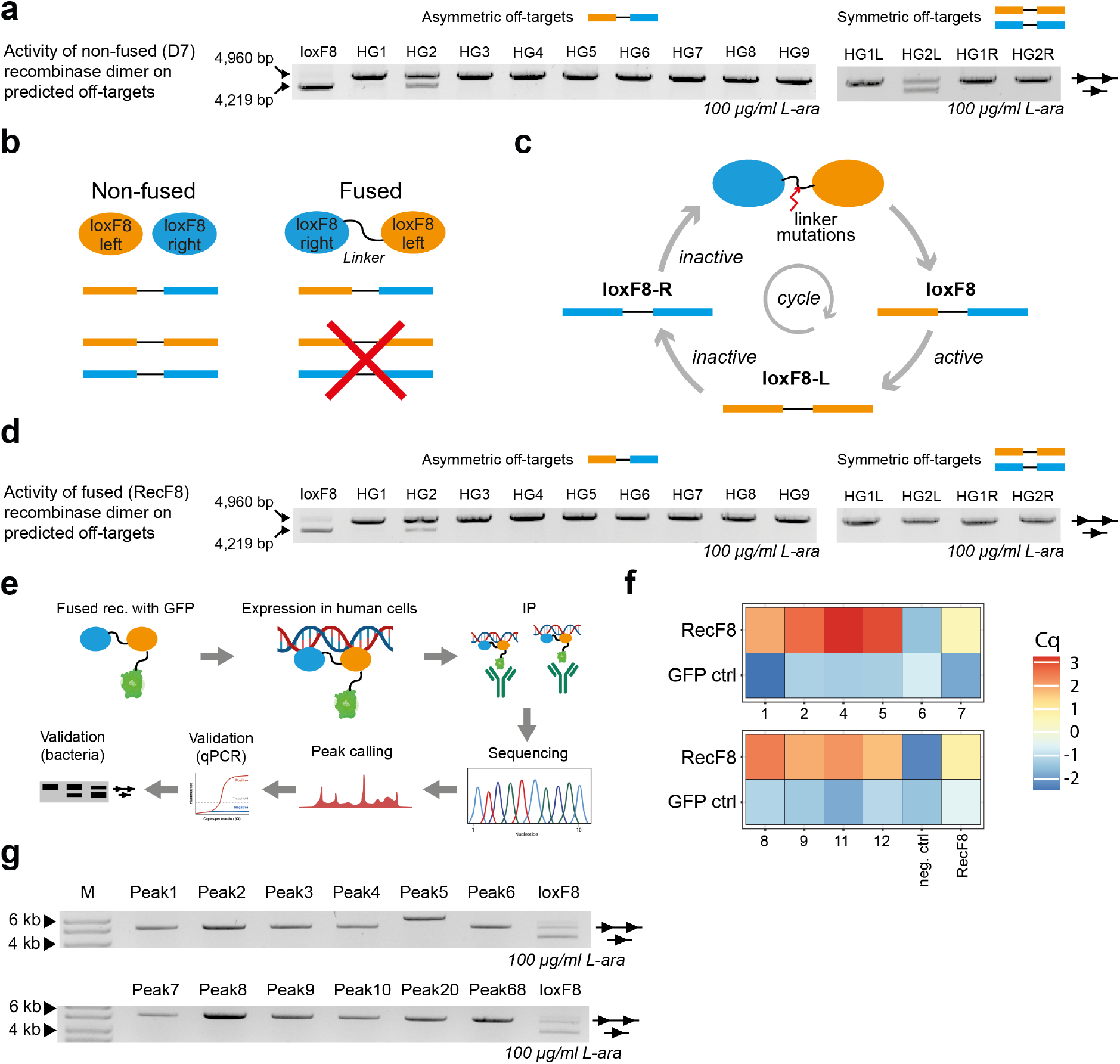
Off-target analysis of F8 recombinases. a) Activity of the D7 heterodimer on predicted asymmetric (nine) and symmetric (four) human loxF8-like sites. Recombination activities of D7 expressed at 100 μg/ml L-arabinose induction are shown. The upper band represents non-recombined plasmid DNA (line with two triangles, 4,960 bp) whereas the lower band shows the recombined form of the plasmid (line with one triangle, 4,219 bp). Recombination-level of the loxF8 site is shown as control. b) Schematic overview of physical linking of recombinases and their predicted activity on symmetric target sites. c) Schematic presentation of the linker selection. Active clones on the loxF8 target site are selected against activity on the symmetric target sites loxF8-L and loxF8-R. The red arrow indicates mutagenesis of the linker region that was subjected to directed evolution. d) Activity of the linked RecF8 dimer on predicted human asymmetric (nine) and symmetric (four) loxF8-like off-target-sites. Recombination activities of RecF8 expressed at 100 μg/ml L-arabinose induction are shown. e) Pipeline of the ChIP-seq experimental procedure to identify RecF8 binding sites in the human genome. f) qPCR-based validation of ten putative binding sites identified by ChIP-seq. The RecF8 sample is compared to the GFP control sample. Enrichment is displayed by a heatmap of the difference of the Cq in the input sample and in the IP sample for each peak. Depletion is displayed in blue and enrichment is displayed in red. g) Bacterial plasmid-based RecF8 activity assay of the ten RecF8-binding sequences. Ninety-five bp surrounding the peak summit of twelve peaks were cloned as an excision substrate in the reporter vector. Recombination-level of the loxF8 site is shown as control. The upper band represents non-recombined substrate (line with two triangles), whereas the lower band shows the recombined plasmid (line with one triangle). The marker lane (M) for 4 kb and 6 kb is shown with arrows. Parts of the figure were created with BioRender.com.

### Linking of recombinase monomers reduces off-target recombination

To reduce the chance of recombination at off-target sites we focused on constraining the monomers from homodimerization. To achieve this goal, we physically fused the D7 monomers to enforce the desired heterodimer assembly (Fig. 4b). To fuse the monomers, we began with a common linker motif of eight glycine-glycine-serine (GGS) repeats ^25^. Indeed, the fused D7 heterodimer recombined the loxF8 target site, although its activity was approximately 5 times reduced compared to the non-fused D7 dimer (Supplementary Fig. 4), presumably because the linker prevented the individual monomers to assemble optimally on the DNA. In order to enhance the activity of the fused monomers, we designed a linker library that was evolved for 10 cycles of SLiDE to find the most active linker variants (Fig. 4c, Supplementary Fig. 4). The final library showed robust activity on the loxF8 target site at low arabinose induction, indicating that variants with improved activity had been selected (Supplementary Fig. 4). At this stage, twelve single clones were analyzed for their activity on loxF8, loxF8-L and loxF8-R. One clone (from here on referred to as RecF8) (Supplementary Fig. 4) was selected, because it showed similar activity on the loxF8 target site as D7, without recombining loxF8-L and loxF8-R, thus abolishing activity on the symmetric target sites (Supplementary Fig. 4). Next, RecF8 was tested on the predicted human off-targets. In contrast to D7, RecF8 showed no observable activity on the symmetric HG2L off-target (Fig. 4d), signifying its improved properties. Interestingly, the off-target activity on the asymmetric HG2 off-target site was also reduced (Fig. 4d), indicating that the linking might also have positive effects on asymmetric off-target activity. We conclude that by fusing the two monomers to limit unwanted homotetramer formation, RecF8 shows improvements in specificity without the loss of activity at the on-target site.

### An experimental off-target assay suggests high target site selectivity for RecF8

Bioinformatic off-target predictions are limited to the depth of the input data consequently producing an incomplete off-target prediction profile for the desired recombinase. To increase the safety profile and enhance the classification of potential off-targets for designer-recombinases, we sought of an experimental method to nominate potential off-targets. Because DNA binding is a prerequisite for successful recombination, we developed a ChIP-Seq-based assay to identify putative RecF8 off-targets in the human genome (Fig. 4e). Eighty-five high confidence RecF8 binding sites were identified with this method (Supplementary Table 3). Twelve hits were subjected to validation by qPCR. For ten out of twelve binding sites efficient primers could be designed and for nine out of these ten sites we were able to confirm the RecF8-mediated DNA enrichment by qPCR (Fig. 4f). Hence, we were able to identify DNA regions that are occupied by RecF8 in the human genome, thereby representing putative off-target sites. RecF8 activity on these potential off-target sites was experimentally tested by cloning the genomic sequences as excision substrates into our bacterial reporter vector. Induction of RecF8 expression at high L-arabinose levels did not result in any detectable recombination at the sites (Fig. 4g), indicating that these sequences, while bound, are not a recombination substrate for RecF8.

Because D7 and RecF8 were both active on the HG2 off-target site in a plasmid-based assay in *E. coli*, we finally investigated whether this sequence is actually altered in human tissue culture cells upon RecF8 expression. Noteworthy, no HG2 recombination activity on the human sites could be detected after expressing RecF8 at high levels with a sensitive PCR-based assay (Supplementary Fig. 4), suggesting that RecF8-mediated alteration at this sequence does not occur in the human genome, at least not at frequencies detectable by PCR. We conclude that RecF8 is fairly specific with minimal off-target activity detectable in a bacterial reporter assay.

### RecF8 expression corrects the F8 exon 1 inversion

Due to the promising characteristics of RecF8, we next tested its activity in mammalian cells. Expression of RecF8 in human cells resulted in similar recombination efficiencies in the HEK293T^loxF8^ reporter cell line compared to the D7 clone and the inversion of the loxF8 locus was also detectable by PCR on genomic DNA (Supplementary Fig. 5). In order to quantify the genomic inversion efficiencies, a qPCR-based assay was developed. Both recombinases (D7 and RecF8) showed around 25-30% inversion of the loxF8 locus on genomic DNA 48 h after mRNA transfection, demonstrating robust and rapid inversion kinetics of the 140 kb genomic DNA fragment (Fig. 5a). In order to investigate whether RecF8 can correct the disease-causing exon 1 inversion in patient derived cells, iPSCs were generated from blood cells of a hemophiliac donor. The iPSCs were characterized in detail and protocols for efficient mRNA transfections were established (Fig. 5b and Supplementary Fig. 5). Forty-eight hours post transfection with RecF8 mRNA, genomic DNA was isolated and analyzed for the inversion of the loxF8 locus via PCR. Only iPSCs (patient derived and wild-type (WT) control) treated with RecF8 showed the expected PCR bands, indicating that the loxF8 locus had been inverted (Fig. 5c). Sequencing of the PCR bands revealed that the inversion in the F8 iPSCs resulted in the same sequence as it is found in non-treated WT iPSCs. This result demonstrates that the RecF8-mediated inversion brings the exon 1 of the F8 gene back into the right orientation and thereby corrects the genetic defect in patient derived cells at nucleotide precision. Conversely, the same sequence found in patient derived iPSCs was identified in WT iPSCs after treating them with RecF8. This result validates that the genomic inversion can be induced in both directions in a patient and WT genetic backgrounds.

**Figure 5:**
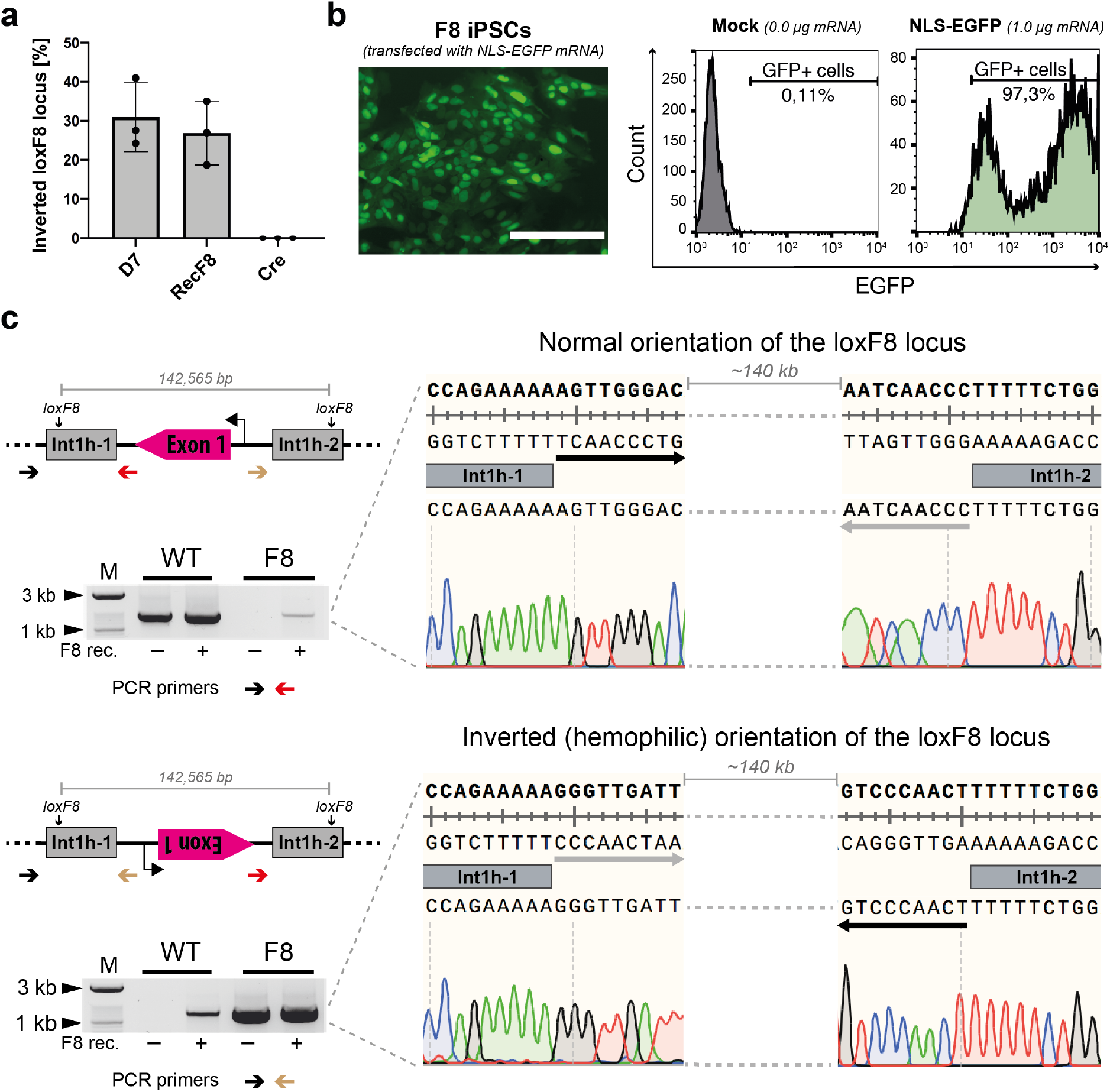
RecF8 induces the genomic inversion of the loxF8 locus in mammalian cells. a) qPCR-based quantification of the genomic inversion efficiencies at the loxF8 locus after treatment with indicated recombinases in HEK293T cells (n=3, biological replicates are shown as dots). Error bars represent standard deviation of the mean (SD). The quantification of the inversion was extrapolated from a standard of defined ratios of WT and F8 patient iPSCs genomic DNA (1%, 5%, 10%, 25% and 50% hemophilic orientation of the loxF8 locus, n=3). b) Transfection of patient derived induced pluripotent stem cells (iPSCs) with NLS-EGFP mRNA. A representative microscopy image of 48 h post transfection with 1 μg of NLS-EGFP mRNA is shown. White bar indicates 200 μm scale bar. A FACS histogram of mock and NLS-EGFP transfected iPSCs 48 h post transfection is shown to the right. NLS, nuclear localization sequence. c) PCR-based detection of loxF8 locus inversion in WT and patient derived iPSCs after RecF8 treatment. Marker (M) lanes at 1 kb and 3 kb are indicated. Sequencing reads of the PCR product confirming the corrected orientation of the DNA sequence after RecF8 treatment are displayed. The black and grey arrows indicate the sequence orientation between the int1h-1 and int1h-2 repeats of the loxF8 locus.

### RecF8-mediated Factor VIII expression is restored in patient derived endothelial cells

It has been suggested that endothelial cells (ECs) of the liver are the main producer cells for Factor VIII in the human body ^26,27^. In order to test if F8 gene expression can be restored in this cell type *in vitro*, we differentiated patient derived iPSCs into endothelial cells (ECs) and delivered RecF8 in form of mRNA into the cells (Fig. 6a and b). Forty-eight hours post transfection, isolated genomic DNA was analyzed for the inversion using a PCR-based assay (Fig. 6c). Only the patient derived ECs treated with RecF8 mRNA showed the PCR bands characteristic for the WT control (Fig. 6c). Quantification of the genomic DNA inversion in ECs treated with the RecF8 mRNA revealed that around ~9% of the loxF8 locus had been reverted back to the WT orientation (Fig. 6d). To examine whether the corrected cells express Factor VIII, we isolated mRNA from non-treated and RecF8-treated cells and performed RT-PCR assays. No amplicon could be generated from the untreated ECs, consistent with the inability of these cells to produce Factor VIII mRNA. In contrast, the Factor VIII transcript was readily detected after treating the ECs with different amounts of RecF8 mRNA (Fig. 6e). Sequencing the transcript revealed the correct boundary of exons 1 and 2 demonstrating a precise reversion, transcription and splicing of the F8 gene (Fig. 6f). Further sequenced exons indicated the presence of the full-length Factor VIII transcript (Supplementary Fig. 6). These results demonstrate that the evolved RecF8 recombinase can be used to specifically invert the 140 kb DNA fragment between int1h-1 and int1h-2 and restore the expression of the F8 gene in patient derived ECs.

**Figure 6:**
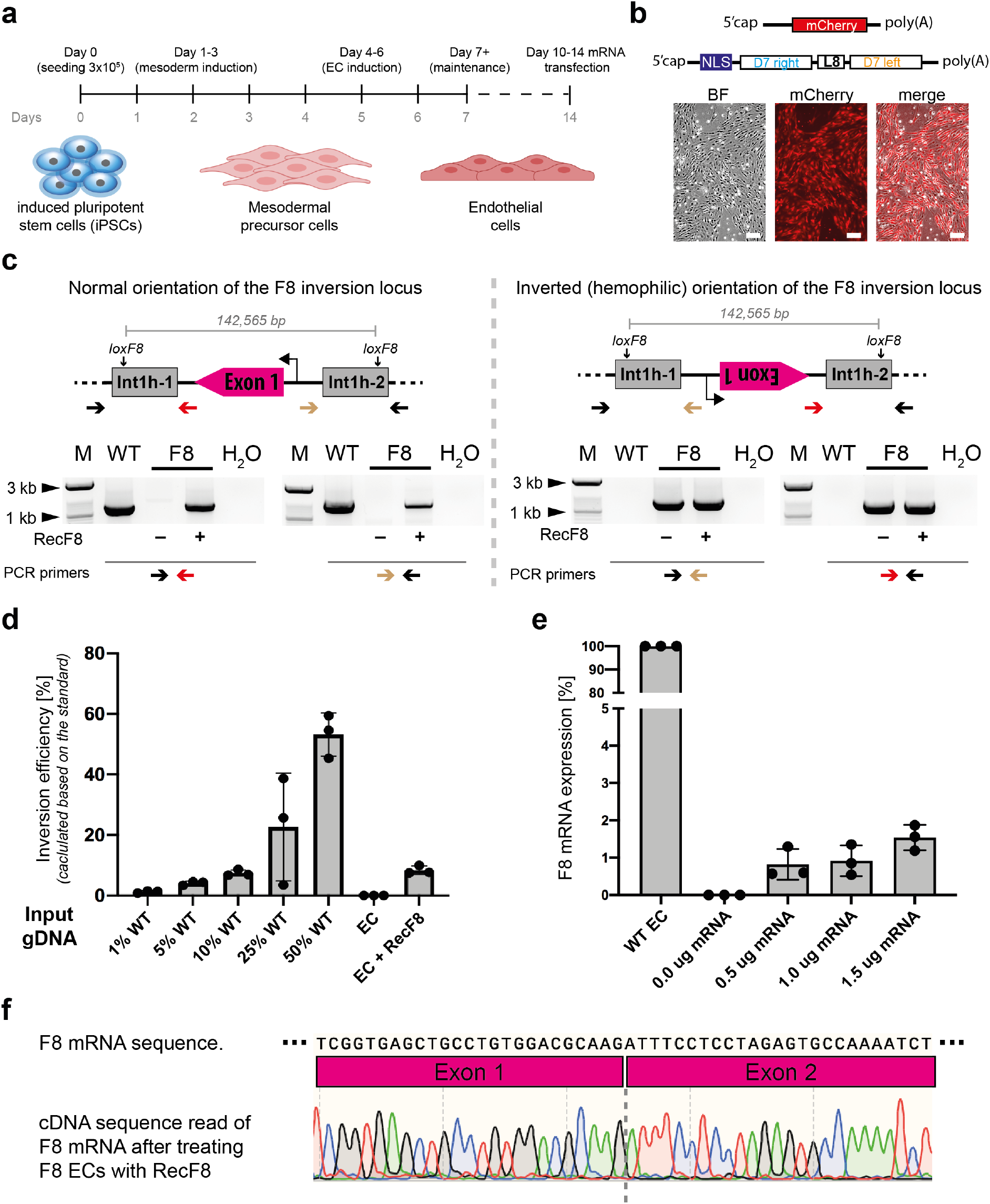
Functional correction of the inverted F8 gene in differentiated endothelial cells. a) Schematic overview of the EC differentiation protocol. Important steps are highlighted on the timeline. b) Efficiency of transfection of mCherry mRNA to differentiated endothelial cells. Brightfield (BF) image, mCherry image and the merged image a shown. Images were taken 48 h post transfection. White bars indicated 50 μm scale bar. c) RecF8 expression corrects the int1h inversion. The arrangement of different primers around the first and second loxF8 site to detect the orientation of the full 140 kb fragment before and after RecF8 treatment are shown in the top panels. The lower panels show gels of PCR products obtained on genomic DNA from patient specific ECs with and without treatment with RecF8. The combination of the primers used for every PCR gel picture is shown below. d) qPCR-based quantification of the genomic inversion efficiencies in ECs after RecF8 treatment. The quantification of the inversion was extrapolated from a standard of defined ratios of WT and F8 patient iPSCs genomic DNA (1%, 5%, 10%, 25% and 50% normal orientation of the loxF8 locus, n=3, replicates are shown as dots). Error bars represent standard deviation of the mean (SD). e) qPCR-based quantification of Factor VIII mRNA transcript (amplification of exon1-exon2 boundary transcript). The relative expression of the Factor VIII mRNA was measured after treating patient specific ECs with indicated amounts of RecF8 mRNA (n=3, replicates are shown as dots). Error bars represent standard deviation of the mean (SD). Factor VIII mRNA expression in untreated WT ECs was used for normalization. The Factor VIII mRNA expression was normalized against ß-actin expression. f) Sequencing read of Factor VIII cDNA of RecF8 treated patient-derived ECs. Note that the exon1-exon2 boundary can only occur, if the inversion is corrected to the WT orientation, the gene is transcribed and the pre-mRNA spliced correctly. Parts of the figure were created with BioRender.com.

## Discussion

Genome editing tools are rapidly being developed for therapeutic applications to devise novel curative strategies for monogenic diseases ^28^. Gene inversions are a particularly challenging genetic alteration to correct with the most commonly used nuclease-based genome editing tools ^6,7^. We explored the potential of site specific recombinases (SSRs) to correct a large gene inversion frequently found in Hemophilia A. SSRs have previously been shown to catalyze large genomic inversions and their target specificity can be altered through directed molecular evolution ^13,14,29,30^. Another advantage of SSRs is their independence of the host cell DNA repair machinery and other cofactors. This allows these enzymes to work efficiently *in vivo* in most cell types, including postmitotic cells ^10^.

To obtain SSRs that recognize a sequence in the inverted int1h repeats surrounding the F8 gene, we applied the well-established substrate-linked directed evolution approach ^13,15,16,20^ and obtained Cre-type enzymes with activity on the desired sites. We acquired both single and heterodimeric Cre-type recombinase systems with robust activity on the loxF8 target sites, demonstrating multiple strategies to successfully bypass the native symmetric preference of Cre. The heterodimeric system required less evolution cycles to develop a novel recombinase of comparable efficiency, suggesting that this method might be a more economical approach for asymmetric target sites. Additionally, developing a heterodimeric recombinase system consisting of evolved monomers with different sequence specificities provides greater flexibility when selecting potential target sites because the asymmetries in each half-site is no longer a limitation ^20^. However, using two evolved monomers instead of one increase potential off-target activity. Each monomer can be combined into a homotetrameric complex to recombine potential symmetric off-targets or into a heterotetrametric complex to recombine asymmetric off-targets. We mitigated this problem by linking the two monomers forming a heterodimer (RecF8). The fusion of the two monomers eliminated observed off-target recombination of a symmetric target site and reduced the effect on an asymmetric off-target site that the non-linked recombinase actively recombined. We assume that the linking of the two monomers sterically obstructed synapse formation and allowed only activity on asymmetric target sites where the linked monomers could bind simultaneously. We believe that this approach should be generally applicable and expedite the process to generate novel, evolved hetero-specific recombinases with desirable properties in the future.

Single-dose mRNA-mediated expression of RecF8 resulted in up to 30% inversion of the loxF8 locus in HEK293T cells and around 9% in postmitotic iPSC-derived endothelial cells. As the limit for SSR-mediated inversion is 50%, these results demonstrate the potential of SSR as an inversion genome editing tool. Because such percentages would be sufficient to change a severe Hemophilia A phenotype (<1% Factor VIII activity) into a moderate (2-5% Factor VIII activity) or mild (5-30% Factor VIII activity) phenotype in patients ^19^, we believe that RecF8 constitutes a promising lead-recombinase for therapeutic exploitation. However, conclusive results of how much inversion of the F8 gene can be reached *in vivo* will first have to be evaluated in an F8 inversion animal model. Ideally, in such an animal model, the recombinase would be transiently delivered to the cells of interest for example with the use of lipid nanoparticles packaged mRNA or through adeno-associated virus (AAV)-mediated delivery ^31,32^. We envision that such delivery methods could potentially reach correction levels that lead to therapeutically relevant Factor VIII expression in the organism. These studies should also reveal if RecF8 expression is tolerated and does not cause unwanted site-effects. If necessary, RecF8 specificity could be further improved utilizing established methods ^33–35^. Overall, our work introduces designer-recombinases as a promising genome-editing tool to correct human diseases caused by genomic inversions. Importantly, numerous other human disease have already been identified to be caused by genomic inversions, including the intron 22 inversion in Hemophilia A ^19^, Sotos Syndrome ^36^ or Hunter Syndrome ^37^, with additional disease-causing inversions likely waiting to be discovered ^38,39^. All of these could potentially benefit from a genome editing strategy employing designer-recombinases.

## Online Material and methods

Primer lists and cycling programs can be found in Supplementary Table 4 and 5.

### Target site identification

In order to nominate a suitable target site, we first identified a sequence of the inverted repeat int1h, by aligning the first intron of the F8 gene against a reverse-complement sequence of 200 kb DNA fragment located upstream of the F8 transcription start site (reference genome assembly hg38), using EMBOSS Water as an alignment tool ^40^. In the following step, using a python script, the inverted repeat (1,041 bp) was scanned for all occurrences of potential target sites, defined as palindromic repeats of 13 bp (half-sites) separated by 8 bp spacer sequence. The half-sites were allowed to have up to 7 positions of asymmetry. A set of 82 target sites fulfilling this criterion was sorted by a larger count of mismatches between one of the half-sites (left or right) and any of the half-sites recognized by previously evolved Cre-type recombinase libraries ^15,16^. As the final target site, we picked the sequence with the highest score from the sorted list and named it loxF8.

In order to avoid potential off-targeting, we scanned the human genome for any occurrences of the selected target site, allowing up to one mismatch in each half-site and any sequence in the spacer region. A search for genomic sequences with the highest resemblance towards loxF8 was performed using an exhaustive short sequence aligner PatMaN, allowing up to 8 mismatches in both half-sites in total ^41^.

### Plasmid construction

Evolution vectors with the different target sites for SLiDE were cloned as described previously ^20^.

A DNA fragment (synthesized by Twist Bioscience) coding for TRE3G-EF1a-Tet-ON^®^ 3G-P2A-eGFP was inserted into the lentiviral backbone of the plentiSAMv2, a gift from Feng Zhang, (Addgene plasmid #75112; http://n2t.net/addgene:75112; RRID:Addgene_75112) ^42^ utilizing NheI and KpnI restriction enzymes (NEB). The resulting plasmid (plentiX) can be used to clone recombinases under the control of a doxycycline inducible promoter employing BsrGI and XbaI restriction enzymes (NEB). In order to fuse the recombinases to EGFP the plasmid was modified and a TRE3G-NLS-eGFP-EF1a-Tet-ON^®^ 3G-P2A-PURO was inserted between the LTRs. The resulting plasmid was used to exchange EGFP with mCherry or tagBFP.

The plentiCRISPR v2, a gift from Feng Zhang, (Addgene plasmid #52961; http://n2t.net/addgene:52961; RRID:Addgene_52961) ^43^ was used to generate the loxF8 reporter plasmid containing a SFFVpromoter-loxF8-PURO-loxF8-mCherry cassette between the LTRs.

### Substrate linked directed evolution (SLiDE)

Recombinases were evolved using substrate linked directed evolution as described previously ^13,15,16,20^. A schematic overview of the experimental procedure is depicted in Supplementary Fig. 6. Recombinase libraries were evolved stepwise on different subsites to obtain active variants on the final target sites (loxF8, loxF8-L and loxF8-R).

### Clonal analysis of recombinases

Recombinase activity was either analyzed with a plasmid-based assay or a PCR-based assay as previously described ^20^. Briefly, recombination of the respective target sites on the evolution plasmid leads to the excision of a fragment from the plasmid. The resulting size difference is an indication for recombinase activity and can be detected by a restriction digest or by PCR.

### Cell culture

HEK293T (ATCC) and HeLa (MPI-CBG, Dresden) were cultured in DMEM, Dulbecco’s modified Eagle’s medium (Gibco) with 10% fetal bovine (tetracycline free) serum and 1% Penicillin-Streptomycin (10 000 U/ml, ThermoFisher).

### Fluorescent activated cell analysis

HEK293T, HeLa or iPSCs were washed once with PBS and then detached using Trypsin (Gibco) or Accutase (Sigma). Cells were resuspended in FACS buffer (PBS with 2.5mM EDTA and 1% BSA) and analyzed with the MACSQuant^®^ VYB Flow Cytometer (Miltenyi) or the BD FACSCanto™ II Cell Analyzer (BD). Analysis of the data was performed using FlowJo™ 10 (BD).

### Long term expression of recombinase libraries

Recombinase libraries were cut out of the evolution vectors via BsrGI and XbaI restriction sites, purified from an agarose gel and subsequently ligated to the plentiX viral vector. HeLa (MPI-CBG Dresden) cells were transduced with lentiviral particles generated from the lentiX-loxF8-L and lentiX-loxF8-R libraries. Transduced HeLa cells will express EGFP and recombinase expression can be induced upon administration of doxycycline. Cells were transduced with a MOI of one to enrich for clones harboring only one recombinase integration per cell. Recombinase expression was induced for 21 days with 100 ng/ml doxyclcline and the medium was renewed every other day. Next, genomic DNA was isolated using the QIAamp DNA Blood Mini Kit (Qiagen) and recombinases were retrieved by PCR using the high-fidelity Herculase II Phusion DNA polymerase (Agilent). Retrieved recombinase libraries were ligated into the evolution vectors and the activity of single clones and libraries was assessed by PCR or by a digest as previously described ^20^.

### Generation of the HEK293T^loxF8^ cell line

HEK293T (ATCC) cells were transduced with lentiviral particles generated from the SFFVpromoter-loxF8-PURO-loxF8-mCherry plasmid. The cells were exposed to 2 μg/ml puromycin selection 48 h after transduction for 7 days. Genomic DNA was isolated from the surviving cells and a reporter specific PCR was performed. The PCR fragment was sequenced to confirm that the reporter construct was integrated in the genome.

### In vitro transcription

Recombinase, eGFP, tagBFP and mCherry mRNA was produced by in vitro transcription (IVT) using the HiScribe™ T7 ARCA mRNA Kit (NEB) and purified using the Monarch^©^ RNA Cleanup Kit (NEB). The DNA templates for the IVT were generated by PCR using the lentiviral plasmids with eGFP (primers 1+2), mCherry (primerss 3+4), tagBFP (primers 5+6). Recombinase DNA templates for IVT were generated by PCR using the pEVO vectors as templates with D7 (primer 7+8 and 9+10) or RecF8 (primers 11+12). Cycling program 3 was used for the mRNA transcription and poly(A) tailing reaction.

### mRNA transfection

IVT produced mRNA was transfected using Lipofectamine™ MessengerMAX™ Transfection Reagent (ThermoFisher). HEK293T^loxF8^ cells were transfected in a 12-well format, seeded at a density of 250,000 cells/well and transfected 24 h after seeding. For each well 250 ng of mRNA (200 ng recombinase mRNA and 50 ng tagBFP mRNA) and 2 μl Lipofectamine™ MessengerMAX™ was mixed with 50 μl Opti-MEM I Reduced Serum Medium (ThermoFisher, prewarmed at RT). After 5 minutes of incubation at RT the mRNA and Lipofectamine™ MessengerMAX™ mixtures were combined, shortly vortexed and incubated for 10 minutes. The transfection mixture was directly added to the cells without changing the medium. Cells were analyzed 48 h post transfection by FACS and fluorescent microscopy.

iPSCs were cultured using StemFit basic 02 (AJINOMOTO) and iMatrix-511 silk laminin coating (NIPPI) as described elsewhere ^44^. iPSCs were transfected as previously described ^45^.

### Detection of the int1h inversion by PCR on genomic DNA

Genomic DNA of HEK293T^loxF8^, iPSCs and ECs was isolated 48 h post transfection using the QIAamp DNA Blood Mini Kit (Qiagen). Two sets of primers were designed to detect the orientation of the 140 kb DNA fragment between the two loxF8 target sites. Primers (15 and 18) are located outside the fragment and do not change upon recombination. Primers (16 and 17) bind inside the 140 kb fragment and will change their orientation upon recombination. Primer pairs 15+16 and 17+18 were used to detect the WT orientation of the 140 kb fragment. Primer pairs 15+17 and 16+18 were used to detect the inverted variant. Independent of the orientation and the primer combinations, cycling program 1 was used for the PCR.

### Linker evolution

The linkers of eight GGS repeats were synthesized by Sigma-Aldrich as annealing compatible oligonucleotides. The designed linker library contained the core 12 amino acids coded by the degenerate codon RVM that were flanked by two GGS repeats from each side, and the sequence was synthesized (Biomers). In both cases, the linker sequences were inserted in the pEVO plasmid via XhoI and BsrGI, between the two recombinase monomers. The linker selection was performed following the SLiDE protocol, with the exception of using the high-fidelity Herculase II Phusion DNA polymerase (Agilent) in order to select the heterodimer fused with a linker with the best properties without introducing new mutations in the recombinase sequences. By varying selection of active and inactive recombinase heterodimers on the loxF8 and symmetric sites (loxF8-L and loxF8-R), respectively, the counter selection was performed.

### Inversion quantification

Quantification of the inversion efficiencies was performed with a qPCR-based assay. In order to detect the WT orientation primer pair 17+18 was used together with a TaqMan specific probe for this amplicon. To detect the inversion orientation primer pair 16+18 was used. For both reactions cycling program 2 was used. A standard curve of 1%, 5%, 10%, 25% and 50% inversion was generated by mixing genomic DNA of WT iPSCs and F8 iPSCs at appropriate rations. The standard curve was used to extrapolate the inversion efficiency of the genomic DNA samples. As the standard curve was generated using genomic DNA of male iPSCs (one X-chromosome), the calculated inversion efficiencies for the HEK293T cells (female, two X-chromosomes) were divided by two.

### ChIP-Seq and qPCR validation of potential binding sites for RecF8

RecF8 recombinase was fused with EGFP and cloned in a modified version of the tetracycline inducible plentiX vector. Hela cells were infected with the lentivirus and selected with 2 μg/ml puromycin 48 h after transduction for 7 days.

Cells were grown in 10 cm dishes and the expression of RecF8 or EGFP (control) was induced for 24h with 100ng/mL doxycycline. The cell lines were crosslinked with 1% formaldehyde for 10 min at room temperature and further processed following the manufacturer’s protocol for High Cell number using the kit TruChip Chromatin Shearing Kit (Covaris). Chromatin shearing was performed using a Covaris M220 sonicator. 1% of the sheared chromatin was separated for qPCR validation (input sample) and the rest was used for immunoprecipitation.

Sonicated chromatin was immunoprecipitated using a goat GFP-antibody (MPI-CBG antibody facility) and Protein G sepharose beads (Protein G Sepharose^®^ 4 Fast Flow, GE Healthcare). Eluates were reverse crosslinked followed by RNA and protein digestion.

Sequencing libraries were prepared using NEBNext^®^ Ultra™ DNA Library Prep Kit for Illumina^®^ from 17-75 ng of ChIP DNA with 15 PCR amplification cycle and size selection using AMPure XP beads. Paired-end sequencing was performed on an Illumina HiSeq 2000, aiming for approximately 25 million pairs of sequencing reads per sample, with each read being 76 bp long. Sequencing reads were aligned to a human reference genome assembly GRCh38.p12 ^46^ using STAR aligner ^47^ tuned for ChIP-Seq analysis pipeline (disabled intron detection, 400 bp of maximum gap between read mates, up to 50 reported alignments of multi-mapper reads). Peak calling was performed with Genrich (https://github.com/jsh58/Genrich), using the ENCODE blacklist (v2) ^48^ for filtering out problematic regions. All steps involving manipulations and comparisons of genomic intervals were done using BEDTools ^49^. Visualizations of the ChIP-Seq pile-up signals were generated with the USCS Genome Browser ^50^ (USCS genome browser tracks are available upon request), directly from BAM files sorted by samtools command line tool ^51^.

Ten out of twelve peaks that were identified by ChIP-Seq were additionally tested by qPCR for recombinase binding (Primers 49-78). qPCR analysis comparing immunoprecipitated samples and input samples was performed using SYBR green MasterMix (Thermo Scientific ABsolute qPCR SYBR Green Mix). Twelve peak-sequences were tested in a plasmid-based assay for recombination. A DNA insert (95 bp) around each peak was generated by PCR (primer 23-46) using cycling program 4 and cloned in the pEVO vector via BglII digestion and ligation. After expression of the recombinase dimer at 100 μg/ml L-arabinose the recombination of the peak sequence was analyzed by the resulting size difference after a potential recombination event.

### HG2 off-target translocation detection

Primers were designed around the HG2 off-target (chromosome 15, primer 19 and 20) site and its potential recombination target site HG2-1 (chromosome 7, primer 21 and 22). PCR products generated from these combinations were sequenced and the presence of the off-targets was validated. Next, a combination of primes as depicted in Supplementary Fig. 4 was used for PCR. PCR products of primer combinations 21+20 and 19+22 should reveal a translocation event. PCR program 5 was used.

### IPSCs and endothelial cell culturing, differentiation and transfection

Reprogramming, culturing, and characterization of patient derived iPSCs (F8 iPSCs) were performed at the Stem Cell Engineering Facility of the Center for Molecular and Cellular Bioengineering (CMCB) at TU Dresden using the CytoTune-iPS 2.0 Sendai Reprogramming Kit (Thermo Fisher Scientific, A16517, Waltham, MA, USA) according to the manufacturer’s guidelines. Human iPSCs were cultured in StemFit02 and laminin coating as described elsewhere ^44^. Endothelial cells were differentiated from F8 iPSCs and maintained in a 6-well dish as described elsewhere ^44^. For each well 700 ng mRNA (250 ng of each recombinase mRNA and 200 ng mCherry mRNA) and 4 μl Lipofectamine™ MessengerMAX™ was mixed with 100 μl Opti-MEM I Reduced Serum Medium (ThermoFisher, prewarmed at RT) and 4 μl Lipofectamine™ MessengerMAX™. The mRNA and Lipofectamine™ MessengerMAX™ mixtures were combined and incubated 15 minutes at RT. Afterwards the transfection mixture was added to the ECs. Four hours after transfection the EC maintenance medium was renewed and the cells were analyzed 48 h post transfection.

### Factor VIII qPCR

RNA was isolated using the RNeasy mini kit (Qiagen). Reverse transcription was carried out using the High Capacity cDNA Reverse Transcription Kit (Applied Biosystems) according to the manufacturer’s protocol. qPCR was carried out using the TaqMan gene-expression master mix (Applied Biosystems) on a QuantStudio 5 Real-Time PCR System (Thermo). In order to detect the right boundary of exon 1 and exon 2 after recombinase mediated correction of the genomic inversion, specific primers (47 and 48) in exon 1 and exon 2 were used together with the universal probe #27 of the Roche Universal ProbeLibrary (Roche). F8 mRNA expression was normalized to ß-actin expression. Probe information for each target gene were obtained from the Universal ProbeLibrary Assay Design Center (https://qpcr.probefinder.com/organ-ism.jsp).

## Acknowledgements

This work was supported, in part, by the European Union [ERC 742133, H2020 UPGRADE 825825] the BMBF GO-Bio [031B0633], the Else Kröner Fresenius Stiftung and the Bayer Hemophilia Awards Program (BHAP). Furthermore, we thank the DRESDEN-concept Genome Center at the CMCB from the TU Dresden for performing the sequencing of the ChIP samples, the Flow Cytometry Core Facility and Stem Cell Engineering Core Facility of the CMCB Technology Platform at TU Dresden for their excellent support.

## Contributions

F.L., J.S., P.M.S. and J.K. performed the directed evolution of the new recombinases. L.M., F.L. and T.R-R. designed, planned performed the linker development. T.R-R. developed and performed the recombinase ChIP-Seq. experiment. F.L., M.K. and K.I. performed the iPSCs and the EC differentiation work in the US. F.L., T.G. and N.R-M. worked with the patient derived iPSCs in Germany. J.H. performed structure-based analyses and evaluated data. M.P-R. and L.T.S. contributed to the bioinformatic analysis and the target site identification. M.P-R. processed the ChIP-Seq. data. R.K. organized and transferred the patient specific cells. C.G. reprogrammed patient specific cells to iPSCs. F.B. and T.T. designed the study, directed the study, evaluated data and wrote the manuscript. F.L. analyzed the data and wrote the manuscript.

## Data availability

The datasets generated during and/or analyzed during the current study are not publicly available due to the current filing process of the patent but are available from the corresponding author F.B. on reasonable request.

## Code availability

The data for this project are confidential, but may be obtained with Data Use Agreements with the Technical University of Dresden, Germany. Researchers interested in access to the data may contact F.B. at. It can take some months to negotiate data use agreements and gain access to the data. The author will assist with any reasonable replication attempts for two years following publication.

---

**Supplementary Figure 1:**
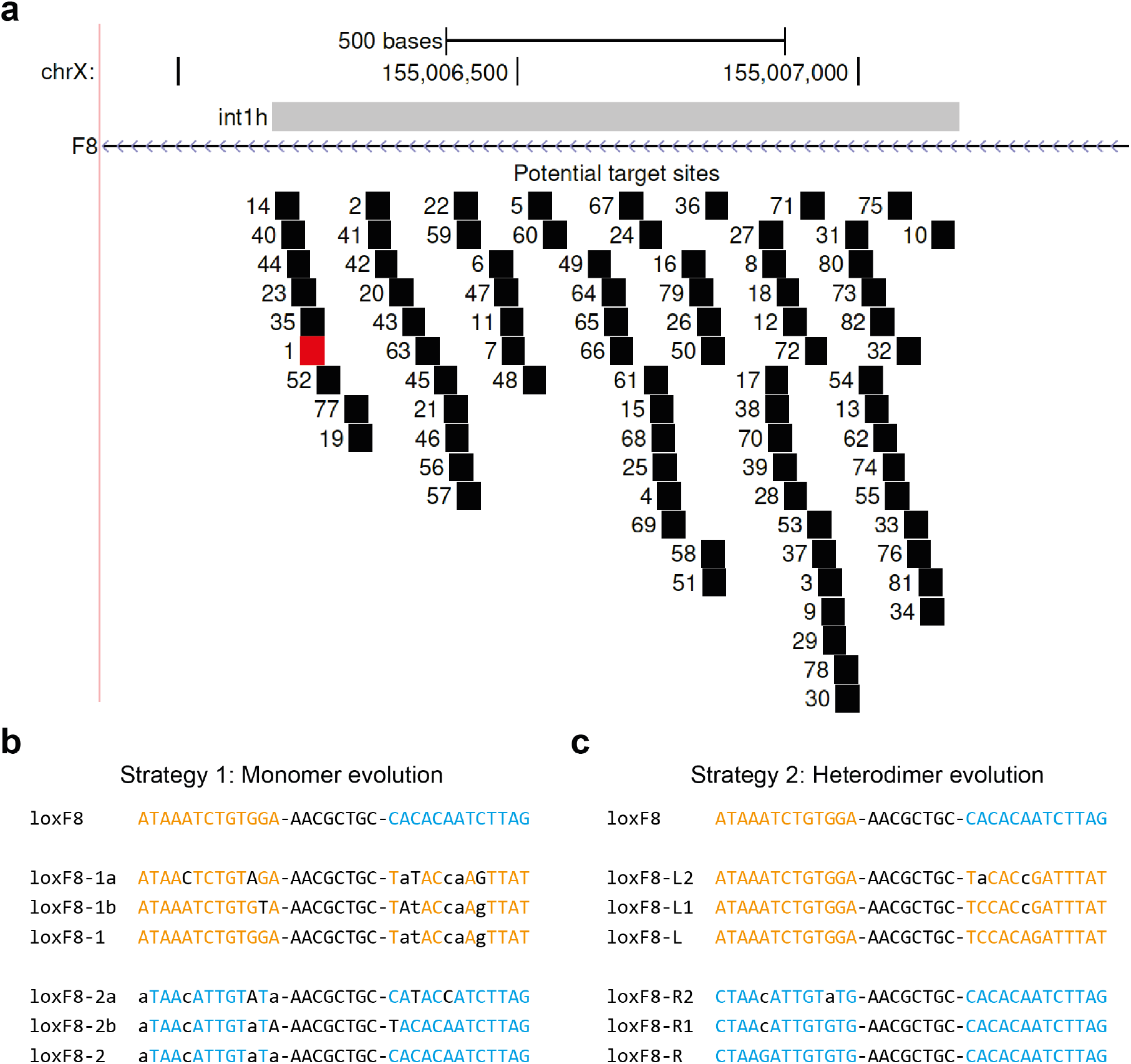
loxF8 target site identification and subsites used in SLiDE (homodimer and heterodimer evolution). a) Genome browser view of int1h-1 repeat (grey) on the X-chromosome with potential unique 34 bp recombinase target sites (black boxes). The target site that was used to evolve recombinases is marked in red. In total 82 potential target sites were nominated and sorted based on the number of mismatches and similarity to previously addressed target sites. b) loxF8 target site and the evolution subsites that were used for the homodimer evolution. c) loxF8 target site and the evolution subsites that were used for the heterodimer evolution. b) and c) The left half sites are depicted in orange and the right half sites in blue. Differences compared to the loxF8 site or its half-sites are presented with black letters. Lower case letters show asymmetry in the evolution targets.

**Supplementary Figure 2:**
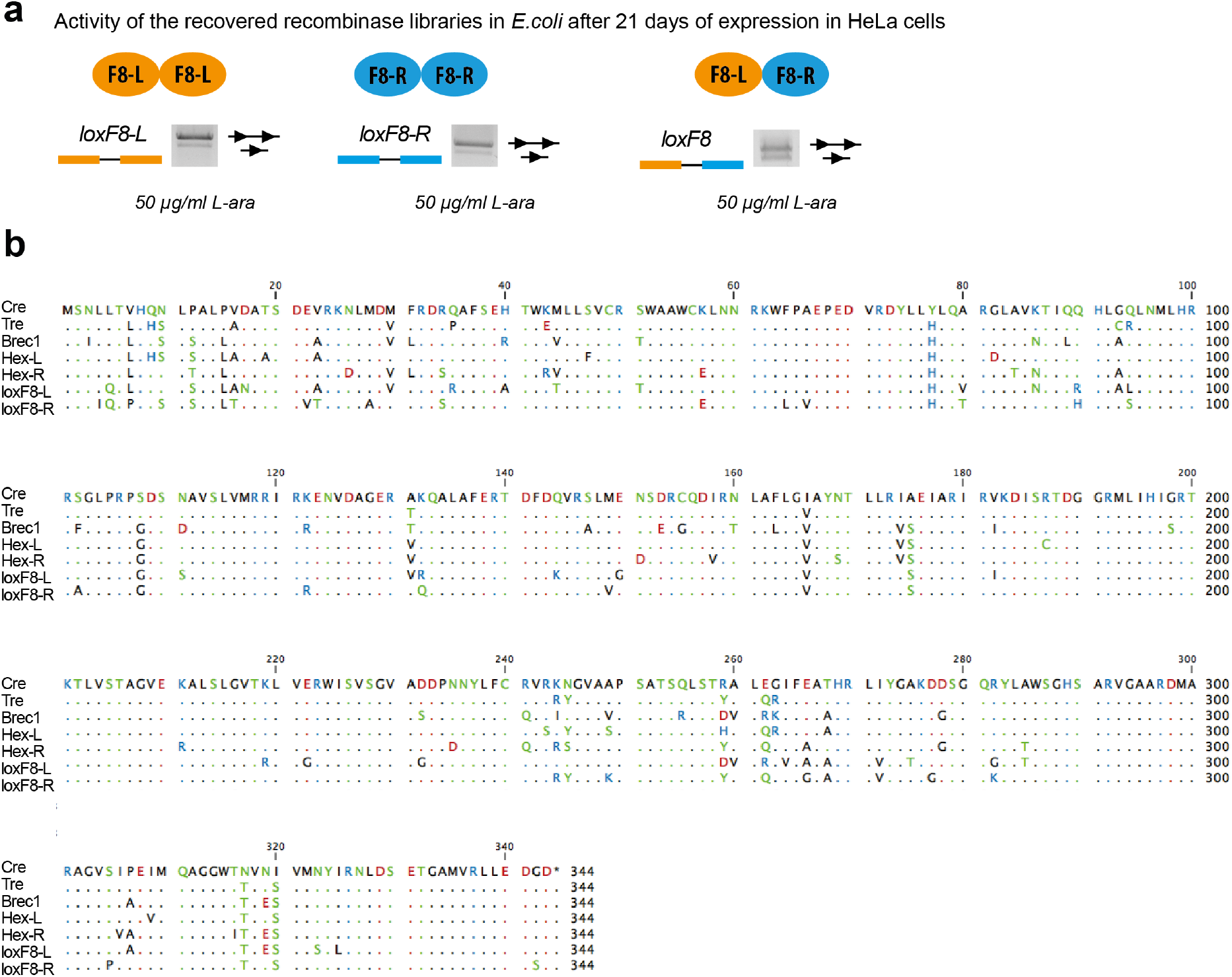
a) Activity of retrieved recombinases after tolerance screen in human cells (HeLa). The retrieved recombinases show activity both as monomers and dimers on the respective target sites. The upper band represents non-recombined substrate whereas the lower band shows recombined substrate at indicated L-arabinose concentrations. b) Amino acid alignment of selected Cre-type recombinases compared to Cre. The alignment was performed using the CLC Genomics Workbench (https://wwww.qiagenbioinformatics.com). Residues were colored based on polarity. The same amino acids found in the reference sequence and the alignment are represented as dots.

**Supplementary Figure 3:**
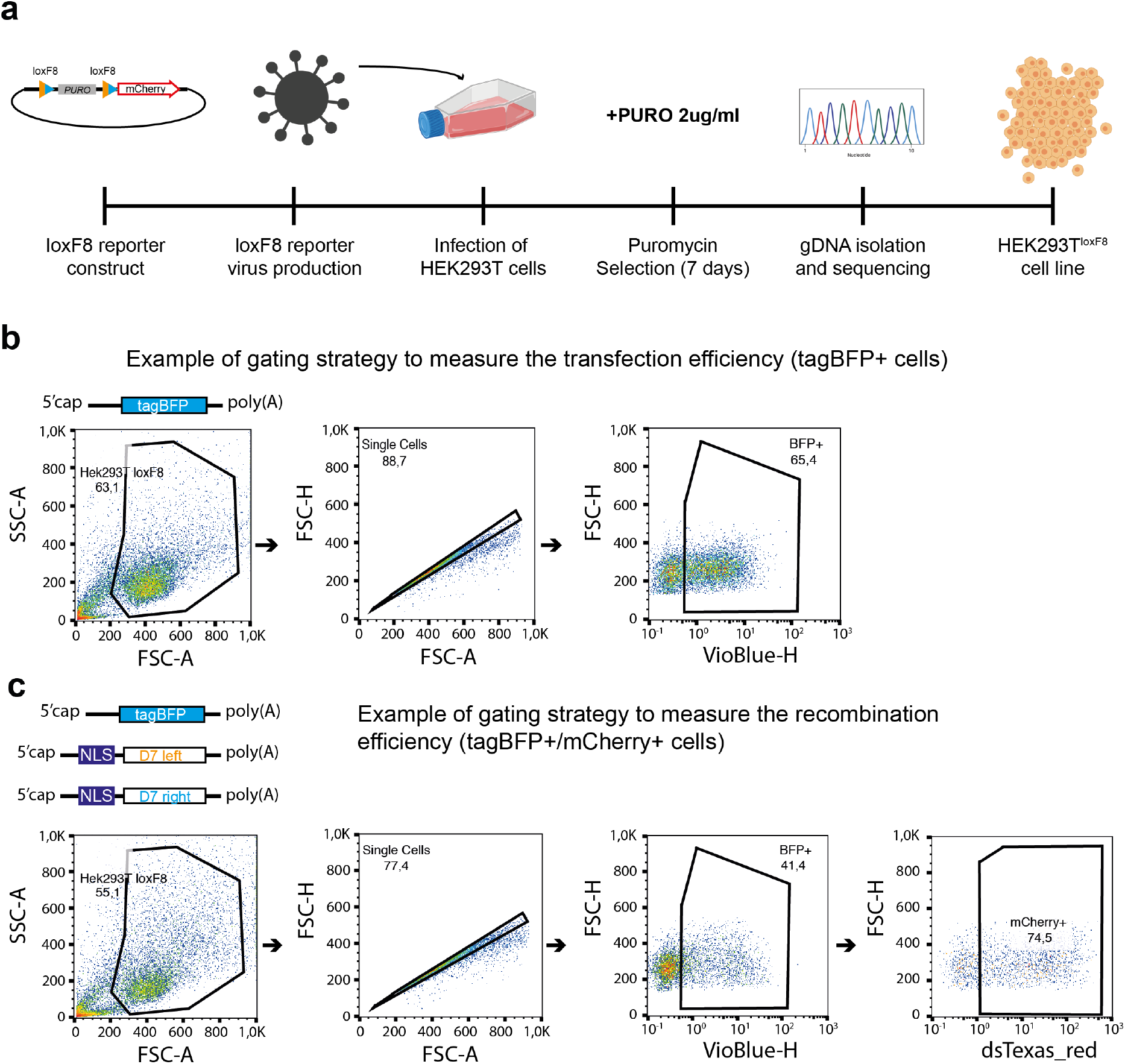
Strategy to evaluate recombinases in mammalian cells. a) Schematic representation of the generation of the HEK293T^loxF8^ reporter cell line. Important steps are shown on the time-line. b) Representative FACS plot and gating strategy to evaluate the transfection efficiency measured by tagBFP+ cells (transfection of tagBFP mRNA only). c) Representative FACS plot and gating strategy of HEK293T^loxF8^ reporter cells that were transfected with tagBFP and indicated recombinase mRNAs. Upon successful site-specific recombination, cells express mCherry. tagBFP+ and mCherry+ were used to calculate the recombination efficiencies (see formula in Fig. 3). Parts of the figure are created with BioRender.com.

**Supplementary Figure 4:**
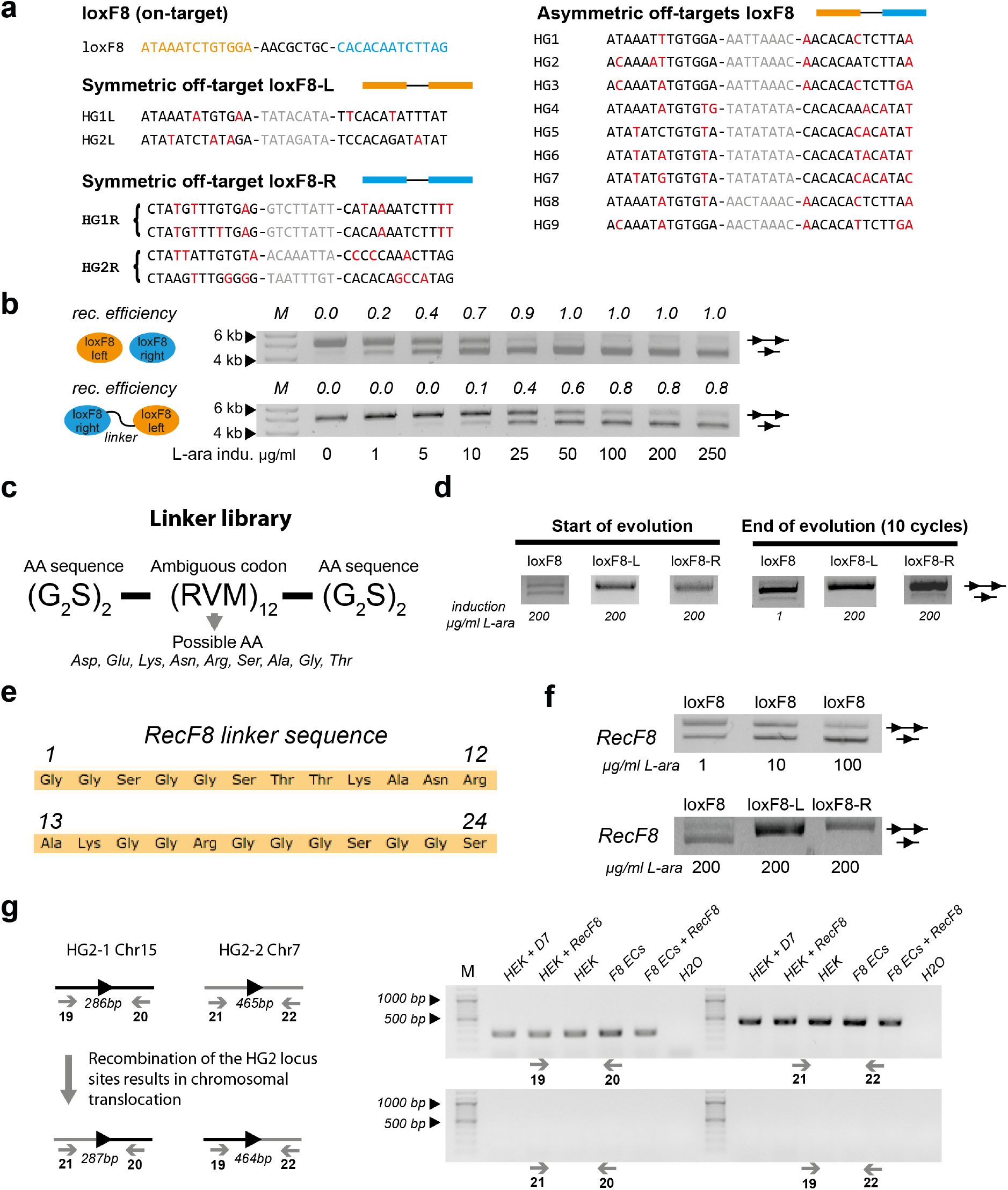
Off-target characterization a) Nucleotide sequences of computationally predicted off-targets with high similarity to loxF8, loxF8-L and loxF8-R. Differences to the respective target sites are marked in red. b) Dose dependent activity of the non-fused and fused (GGS-linker) D7 dimer at different expression levels. The upper band represents non-recombined substrate whereas the lower band shows recombination. L-arabinose induction levels are shown on the bottom. The recombination efficiencies were calculated from the ratios of the intensities of the bands using Fiji ^53^. c) Linker library sequence used to create a pool of differently linked recombinases (the recombinases are always the same). d) Activity of the fused recombinases with the linker library before and after evolution of the linker at indicated L-arabinose induction levels on indicated target sites. e) Linker sequence of RecF8. Amino acids are shown as three letter code with positions indicated. f) Activity of RecF8 on the loxF8 target site at indicated induction levels on displayed target site. The upper band represents non-recombined substrate whereas the lower band shows recombination. g) Design of a PCR assay that would detect RecF8-mediated recombination of the asymmetric HG2 off-target site on chromosomes 7 and 15. Marker (M) lanes at 500 bp and 1000 bp are indicated. Primer binding sites are depicted with grey arrows. The PCR product size is indicated for each PCR primer combination.

**Supplementary Figure 5:**
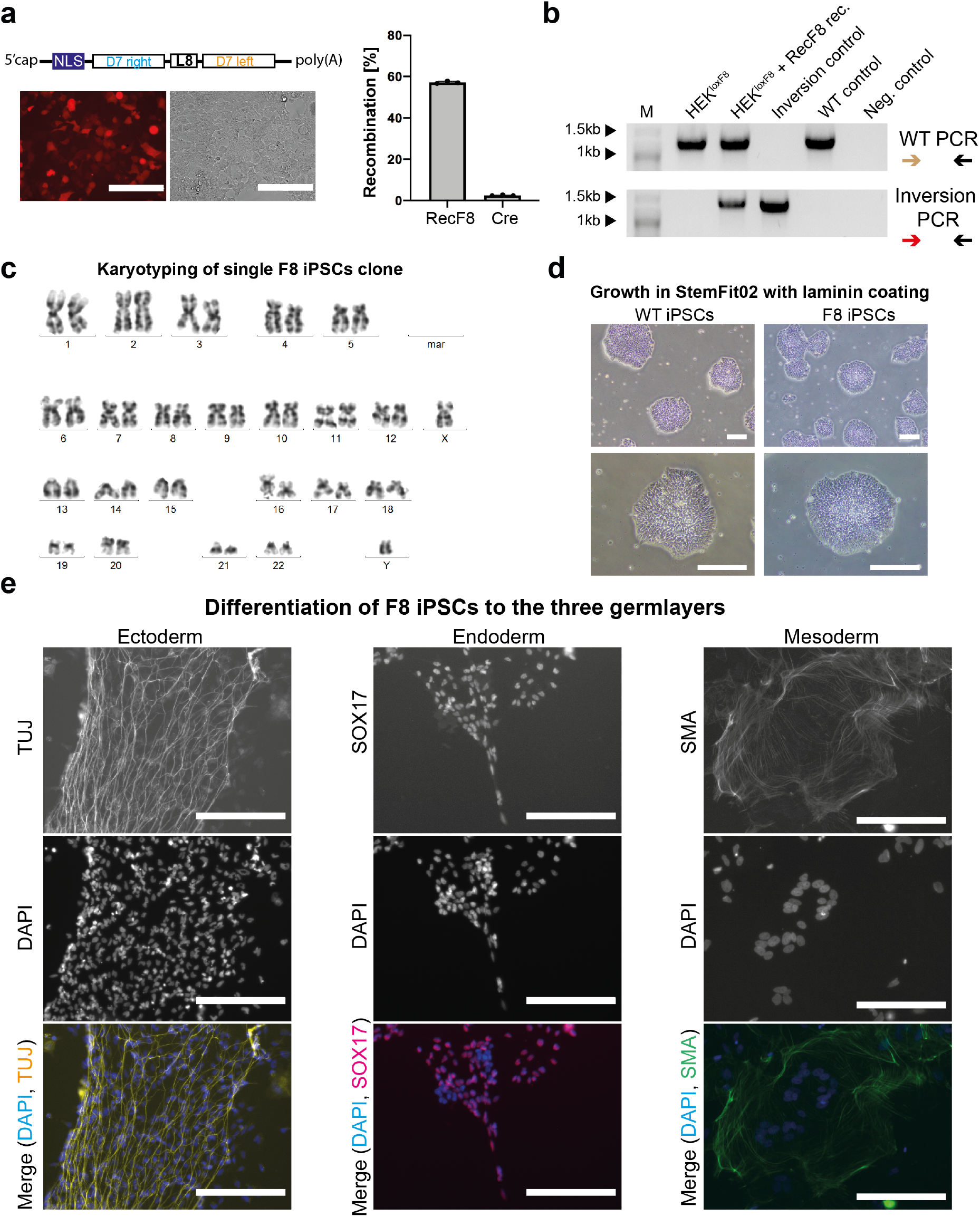
Activity of RecF8 in HEK293T cells and patient derived iPSCs culturing and characterization. a) Transient expression (mRNA) of RecF8 dimer in HEK293T^loxF8^ reporter cell line and quantification of the recombination efficiency (n=3, biological replicates are shown as dots). Error bars represent standard deviation of the mean (SD). White bars indicate 200 μm scale bars. b) PCR-based detection of the loxF8 locus orientation in HEK293T^loxF8^ reporter cells treated with RecF8. Marker (M) lanes at 1 kb and 1.5 kb are indicated. The primer combination used for the PCR is shown next to the gel image. The same primer as shown in Fig. 5 were used. c) Karyotyping of patient derived iPSCs. A normal male karyotype without abnormalities is seen for the patient derived iPSCs. d) Morphological comparison of patient derived iPSCs to WT iPSCs cell line grown in StemFit02 with laminin coating. White bars indicate 200 μm scale bars. e) Differentiation capacity of patient derived iPSCs to ectoderm, endoderm and mesoderm. The ectodermal marker beta-III-tubulin (TUJ) was used to show ectodermal differentiation capacity. The endodermal marker SRY-related HMG-box 17 (SOX17) was used to show endodermal differentiation capacity. The mesodermal marker smooth muscle actin (SMA) was used to show mesodermal differentiation capacity. DAPI was used in all three differentiations to stain the nucleus of the cells. The black and white images of the staining’s as well as the merge images with the respective colors are shown. White bars indicate 200 μm scale bars.

**Supplementary Figure 6:**
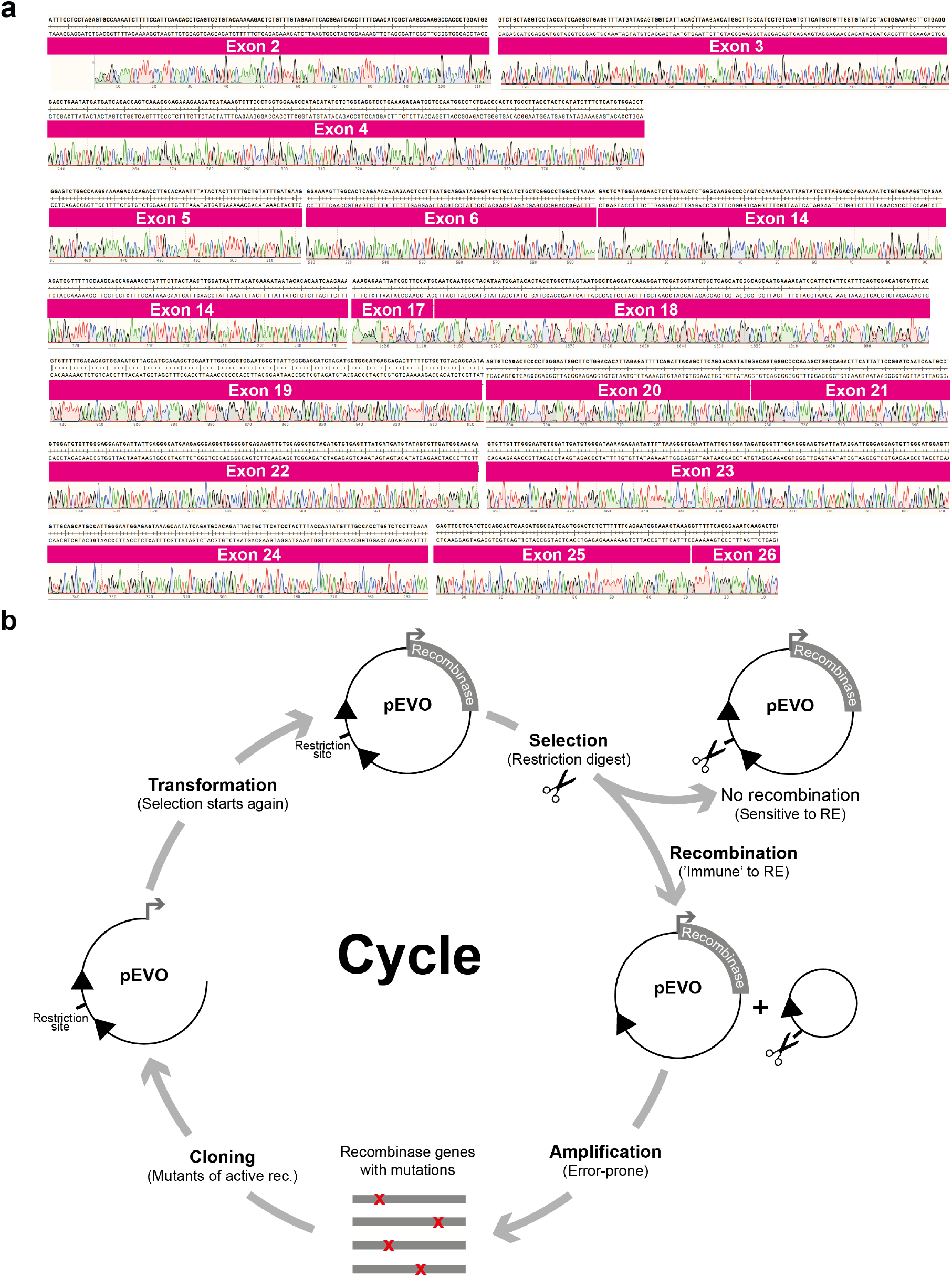
Factor VIII cDNA sequencing reads and graphical scheme for the substrate-linked directed evolution (SLiDE) strategy. a) Sequencing reads of the Factor VIII transcript (cDNA) generated from patient specific ECs treated with RecF8. Exons are marked in magenta and numbered. The sequence and the chromatogram for each read are indicated. b) The evolution cycle starts with recombinase libraries cloned into the pEVO vector (top middle). The recombinase gene is indicated in gray. Two lox-like sites that serve as the substrate for the recombinase are indicated as black triangles. A unique restriction site between the lox-like sites is shown. Expression of the recombinase can be induced by different levels of L-arabinose (typically ranging from 1 μg/ml – 200 μg/ml). The unique restriction site is removed after active recombinases recombine the two lox-like target sites. The resulting pEVO plasmid (only one lox-like site left) will be ‘immune’ the restriction enzyme (RE) recognizing the unique restriction site. Amplifying only the active recombinase by error-prone PCR generates a new library of recombinases (mutations are depicted as red asterisks in the recombinase gene, bottom middle) that is again cloned into the empty pEVO vector with two lox-like target sites to start a new evolution cycle.

## Tables

Supplementary Table 1: 82 potential recombinase target sites (34 bp) found in the int1 repeat surrounding the F8 gene. Genomic coordinates are indicated. The target sites are ranked by the number of mismatches (asymmetry) between the left and right half sites of a given spacer. The target sites are then compared to target sites of previous evolutions and the maximum mismatch count is displayed. The first sequence of this list has the lowest asymmetry and the lowest maximum mismatch count and was used in this study to evolve recombinases. It is called loxF8 (highlighted in yellow).

Supplementary Table 2: Potential off-targets with similarity towards loxF8, loxF8-L and loxF8-R. The chromosomal coordinates and sequence (left half-site, right half-site and spacer) of each identified site are displayed. For each half-site of a potential off-target site the mismatches are displayed separately. The total mismatches indicate the similarity of the identified site compared to the full loxF8, loxF8-L or loxF8-R. A maximum of 7 mismatches was allowed to identify potential off-targets.

Supplementary Table 3: Putative binding sites identified by recombinase ChIP-seq. 85 potential binding sites are shown with their genomic coordinates and the summit of each peak is displayed.

Supplementary Table 4: Nucleotide sequences of primers used in this study. Top strand primers are marked with a ‘_F’ and bottom strand primers are marked with a ‘_R’. A small description of use of the primer is shown.

Supplementary Table 5: Cycling programs used for different PCRs.

