## Supplementary Table 1 for "Correction of a Factor VIII genomic inversion with designer-recombinases"

| | Potential target site sequence | | | | Genomic coordinates (hg38) | | Left half-site - the most similar library | | | Right half-site - the most similar library | | | |
|---|---|---|---|---|---|---|---|---|---|---|---|---|---|
| # | Left half-site | Spacer | Right half-si | Asymmetry | h-1 (plus stra | int1h-2 (minus strand) | Target site | Half-site sequence | Mismatches | Target site | Half-site sequence | Mismatches | Maximum mismatch count |
| 1 | AATCTGTGGA | AACGCTGC | CACACAATCT | 6 | 5006182-155 | chrX:155148713-155148746 | loxBTR_1B | ATACCACTGTATA | 5 | loxBTR_1C | ATAACATTGCTTA | 5 | 5 |
| 2 | AGGGTGGATG | CGAAGCAG | TAAGGACCCC | 6 | 5006279-155 | chrX:155148616-155148649 | loxBTR_1A | AACACATTGTATA | 6 | loxBTR_2 | AAGGCAAGCTTTA | 5 | 6 |
| 3 | GCAAAGGTGG | GGCCGACT | TAATTCTTGC | 7 | 5006942-155 | chrX:155147953-155147986 | loxBTR_2a+2 | AAGGCAAGGTTTA | 6 | loxBTR_2 | AAGGCAAGCTTTA | 3 | 6 |
| 4 | CCCTTAAGGA | AGGTAAAA | TCCAGAAGGG | 5 | 5006706-155 | chrX:155148189-155148222 | loxBTR_1a | AACCCATTGTATA | 7 | loxBTR_1a | AACCCATTGTATA | 6 | 7 |
| 5 | ACAAAGCAGA | CCCTGCTC | TGTGCGTTTA | 6 | 5006517-155 | chrX:155148378-155148411 | loxBTR_2A | AAGACTAGGTATA | 5 | loxBTR_2B | ATAGCAAGGTATA | 7 | 7 |
| 6 | GAAATGTGAT | GTTTGGAC | TGCAAGTTTT | 7 | 5006459-155 | chrX:155148436-155148469 | loxBTR_2a+2 | AAGGCAAGGTTTA | 7 | loxBTR_1A | AACACATTGTATA | 7 | 7 |
| 7 | TGCAAGTTTT | GGTGAGAA | ATAACAATGC | 7 | 5006477-155 | chrX:155148418-155148451 | loxBTR_2a+2 | AAGGCAAGGTTTA | 7 | loxBTR_1A | AACACATTGTATA | 6 | 7 |
| 8 | CCCTTCACAG | GATCTCGC | AGCAGATTGG | 7 | 5006860-155 | chrX:155148035-155148068 | loxBTR_1a | AACCCATTGTATA | 7 | loxBTR_2 | AAGGCAAGCTTTA | 6 | 7 |
| 9 | GGTGGGGCCG | ACTTAATT | CTTGCCTCCC | 7 | 5006947-155 | chrX:155147948-155147981 | loxBTR_2a | AAGGCTAGGTATA | 7 | loxBTR_2B | ATAGCAAGGTATA | 7 | 7 |
| 10 | GCCCAGCGTA | CTGATAAG | GAAGCTCTTC | 7 | 5007109-155 | chrX:155147786-155147819 | loxBTR_1 | AACCCACTGCTTA | 6 | loxBTR_1C | ATAACATTGCTTA | 7 | 7 |
| 11 | ACTGCAAGTT | TTGGTGAG | AAATAACAAT | 5 | 5006475-155 | chrX:155148420-155148453 | loxBTR_2A | AAGACTAGGTATA | 8 | loxBTR_1b+2 | AACACACTGCTTA | 8 | 8 |
| 12 | AAAAGTCTCC | TTGAAACT | GCAGACCTCT | 5 | 5006891-155 | chrX:155148004-155148037 | loxBTR_2b | ATAACAAGCTTTA | 7 | loxBTR_1b+2 | AACACACTGCTTA | 8 | 8 |
| 13 | AGGGTTTCTG | GGACACCC | GAGAAAGCAC | 5 | 5007011-155 | chrX:155147884-155147917 | loxBTR_1A | AACACATTGTATA | 8 | loxBTR_1 | AACCCACTGCTTA | 8 | 8 |
| 14 | GTCTGATTTC | CTTTGGGA | GAATTCATTG | 6 | 5006145-155 | chrX:155148750-155148783 | loxBTR_2 | AAGGCAAGCTTTA | 8 | loxBTR_2 | AAGGCAAGCTTTA | 8 | 8 |
| 15 | GGAAAATTTC | CCCTTAAG | GAAGGTAAAA | 6 | 5006696-155 | chrX:155148199-155148232 | loxBTR_2 | AAGGCAAGCTTTA | 7 | loxBTR_1C | ATAACATTGCTTA | 8 | 8 |
| 16 | AATGGTGAGC | AGTTTAAA | CCTAGCAGTT | 6 | 5006742-155 | chrX:155148153-155148186 | loxBTR_2A | AAGACTAGGTATA | 8 | loxBTR_1A | AACACATTGTATA | 8 | 8 |
| 17 | TCACAGGATC | TCGCAGCA | GATTGGCTGA | 6 | 5006864-155 | chrX:155148031-155148064 | loxBTR_1B | ATACCACTGTATA | 7 | loxBTR_1B | ATACCACTGTATA | 8 | 8 |
| 18 | AGATTGGCTG | AAAAGTCT | CCTTGAAACT | 6 | 5006881-155 | chrX:155148014-155148047 | loxBTR_1A | AACACATTGTATA | 5 | loxBTR_1A | AACACATTGTATA | 8 | 8 |
| 19 | CATGTGACAA | AGTGCTTT | CCGTGAAAAG | 7 | 5006253-155 | chrX:155148642-155148675 | loxBTR_1 | AACCCACTGCTTA | 7 | loxBTR_1a | AACCCATTGTATA | 8 | 8 |
| 20 | TGAAGCACGC | GGTCACCC | TCCAGCCACC | 7 | 5006312-155 | chrX:155148583-155148616 | loxBTR_2b | ATAACAAGCTTTA | 8 | loxBTR_2 | AAGGCAAGCTTTA | 8 | 8 |
| 21 | ACGGTGGCAC | CTTCCAGC | AGGTCCCCGG | 7 | 5006393-155 | chrX:155148502-155148535 | loxBTR_1A | AACACATTGTATA | 8 | loxBTR_1 | AACCCACTGCTTA | 8 | 8 |
| 22 | AGCAGGTCCC | CGGGGTTG | TGCCCCTGGA | 7 | 5006408-155 | chrX:155148487-155148520 | loxBTR_2A | AAGACTAGGTATA | 8 | loxBTR_2A | AAGACTAGGTATA | 7 | 8 |
| 23 | CATTGCCAGC | TATAAATC | TGTGGAAACG | 7 | 5006168-155 | chrX:155148727-155148760 | loxBTR_2C | ATAACTAGCTTTA | 8 | loxBTR_1a | AACCCATTGTATA | 7 | 8 |
| 24 | ATGAACTGAG | GAAAGGAG | GAAAATTTCC | 7 | 5006679-155 | chrX:155148216-155148249 | loxBTR_2 | AAGGCAAGCTTTA | 8 | loxBTR_2 | AAGGCAAGCTTTA | 5 | 8 |
| 25 | AAATTTCCCC | TTAAGGAA | GGTAAAATCC | 7 | 5006699-155 | chrX:155148196-155148229 | loxBTR_1C | ATAACATTGCTTA | 7 | loxBTR_1A | AACACATTGTATA | 8 | 8 |
| 26 | GCAGTTTTGC | ATTAATTC | ACATAAAGTA | 7 | 5006764-155 | chrX:155148131-155148164 | loxBTR_1B | ATACCACTGTATA | 7 | loxBTR_1C | ATAACATTGCTTA | 8 | 8 |
| 27 | CACCCCTTCA | CAGGATCT | CGCAGCAGAT | 7 | 5006857-155 | chrX:155148038-155148071 | loxBTR_1B | ATACCACTGTATA | 8 | loxBTR_2C | ATAACTAGCTTTA | 8 | 8 |
| 28 | AAAGTCTCCT | TGAAACTG | CAGACCTCTC | 7 | 5006892-155 | chrX:155148003-155148036 | loxBTR_2b | ATAACAAGCTTTA | 8 | loxBTR_2B | ATAGCAAGGTATA | 6 | 8 |
| 29 | GTGGGGCCGA | CTTAATTC | TTGCCTCCCT | 7 | 5006948-155 | chrX:155147947-155147980 | loxBTR_2a | AAGGCTAGGTATA | 6 | loxBTR_1A | AACACATTGTATA | 8 | 8 |
| 30 | GCCTCCCTGC | TCTCCCAC | GTAGCCCTGC | 7 | 5006968-155 | chrX:155147927-155147960 | loxBTR_1C | ATAACATTGCTTA | 8 | loxBTR_2a+2 | AAGGCAAGGTTTA | 5 | 8 |
| 31 | CCACGTAGCC | CTGCATTT | CACTCCATTC | 7 | 5006982-155 | chrX:155147913-155147946 | loxBTR_1 | AACCCACTGCTTA | 8 | loxBTR_2B | ATAGCAAGGTATA | 7 | 8 |
| 32 | TGCCAACTGA | AGGCCTTG | ACAAATGACT | 7 | 5007058-155 | chrX:155147837-155147870 | loxBTR_1 | AACCCACTGCTTA | 8 | loxBTR_2C | ATAACTAGCTTTA | 8 | 8 |
| 33 | GCCTTGACAA | ATGACTTT | CTGTACTGGC | 7 | 5007070-155 | chrX:155147825-155147858 | loxBTR_1A | AACACATTGTATA | 6 | loxBTR_2B | ATAGCAAGGTATA | 8 | 8 |
| 34 | ACTGGCTGAG | GGCCAGGG | CCCAGCGTAC | 7 | 5007092-155 | chrX:155147803-155147836 | loxBTR_2C | ATAACTAGCTTTA | 7 | loxBTR_2 | AAGGCAAGCTTTA | 8 | 8 |
| 35 | AAATCTGTGG | AAACGCTG | CCACACAATC | 4 | 5006181-155 | chrX:155148714-155148747 | loxBTR_1A | AACACATTGTATA | 9 | loxBTR_2 | AAGGCAAGCTTTA | 9 | 9 |
| 36 | TAATTCACAT | AAAGTATA | ATGAAAACTG | 5 | 5006776-155 | chrX:155148119-155148152 | loxBTR_1A | AACACATTGTATA | 9 | loxBTR_1a | AACCCATTGTATA | 8 | 9 |
| 37 | TGAGTTGGGC | AAAGGTGG | GGCCGACTTA | 5 | 5006934-155 | chrX:155147961-155147994 | loxBTR_1 | AACCCACTGCTTA | 9 | loxBTR_1 | AACCCACTGCTTA | 9 | 9 |
| 38 | CACAGGATCT | CGCAGCAG | ATTGGCTGAA | 6 | 5006865-155 | chrX:155148030-155148063 | loxBTR_2B | ATAGCAAGGTATA | 9 | loxBTR_2B | ATAGCAAGGTATA | 9 | 9 |
| 39 | GCAGCAGATT | GGCTGAAA | AGTCTCCTTG | 6 | 5006876-155 | chrX:155148019-155148052 | loxBTR_2B | ATAGCAAGGTATA | 7 | loxBTR_1B | ATACCACTGTATA | 9 | 9 |
| 40 | TCCTTTGGGA | GAATTCAT | TGCCAGCTAT | 7 | 5006153-155 | chrX:155148742-155148775 | loxBTR_1C | ATAACATTGCTTA | 7 | loxBTR_1C | ATAACATTGCTTA | 9 | 9 |
| 41 | GGGTGGATGC | GAAGCAGT | AAGGACCCCC | 7 | 5006280-155 | chrX:155148615-155148648 | loxBTR_2a+2 | AAGGCAAGGTTTA | 7 | loxBTR_2 | AAGGCAAGCTTTA | 9 | 9 |
| 42 | GAAGCAGTAA | GGACCCCC | TTCATGAAGC | 7 | 5006290-155 | chrX:155148605-155148638 | loxBTR_1 | AACCCACTGCTTA | 9 | loxBTR_2a | AAGGCTAGGTATA | 8 | 9 |
| 43 | CCAGCCACCA | GAACCAGA | GGAACGCTGT | 7 | 5006331-155 | chrX:155148564-155148597 | loxBTR_1 | AACCCACTGCTTA | 9 | loxBTR_1A | AACACATTGTATA | 6 | 9 |
| 44 | GAGAATTCAT | TGCCAGCT | ATAAATCTGT | 7 | 5006161-155 | chrX:155148734-155148767 | loxBTR_2 | AAGGCAAGCTTTA | 9 | loxBTR_1A | AACACATTGTATA | 8 | 9 |
| 45 | ATCTAGGCAC | ACGTCACG | GTGGCACCTT | 7 | 5006378-155 | chrX:155148517-155148550 | loxBTR_1A | AACACATTGTATA | 9 | loxBTR_1C | ATAACATTGCTTA | 9 | 9 |
| 46 | CGGTGGCACC | TTCCAGCA | GGTCCCCGGG | 7 | 5006394-155 | chrX:155148501-155148534 | loxBTR_1B | ATACCACTGTATA | 9 | loxBTR_1a | AACCCATTGTATA | 6 | 9 |
| 47 | TGTTTGGACT | GCAAGTTT | TGGTGAGAAA | 7 | 5006468-155 | chrX:155148427-155148460 | loxBTR_1A | AACACATTGTATA | 9 | loxBTR_1B | ATACCACTGTATA | 8 | 9 |
| 48 | GGTTGCAGAC | AAAGCAGA | CCCTGCTCTG | 7 | 5006509-155 | chrX:155148386-155148419 | loxBTR_2C | ATAACTAGCTTTA | 9 | loxBTR_2C | ATAACTAGCTTTA | 9 | 9 |
| 49 | ACAGACACAC | CAGAGAGA | GGTTCTCAGG | 7 | 5006604-155 | chrX:155148291-155148324 | loxBTR_2b | ATAACAAGCTTTA | 8 | loxBTR_1B | ATACCACTGTATA | 9 | 9 |
| 50 | TTGCATTAAT | TCACATAA | AGTATAATGA | 7 | 5006770-155 | chrX:155148125-155148158 | loxBTR_1B | ATACCACTGTATA | 9 | loxBTR_1A | AACACATTGTATA | 7 | 9 |
| 51 | CATTAATTCA | CATAAAGT | ATAATGAAAA | 7 | 5006773-155 | chrX:155148122-155148155 | loxBTR_2C | ATAACTAGCTTTA | 8 | loxBTR_2 | AAGGCAAGCTTTA | 9 | 9 |
| 52 | ATCTTAGCAC | ACAAGATT | GGCAGAAAAT | 7 | 5006206-155 | chrX:155148689-155148722 | loxBTR_1C | ATAACATTGCTTA | 8 | loxBTR_1A | AACACATTGTATA | 9 | 9 |
| 53 | GACCCACTGA | GTTGGGCA | AAGGTGGGGC | 7 | 5006927-155 | chrX:155147968-155148001 | loxBTR_1b | ATAACACTGCTTA | 8 | loxBTR_1 | AACCCACTGCTTA | 9 | 9 |
| 54 | CTCCATTCCA | GGGTTTCT | GGGACACCCG | 7 | 5007002-155 | chrX:155147893-155147926 | loxBTR_1B | ATACCACTGTATA | 8 | loxBTR_1B | ATACCACTGTATA | 9 | 9 |
| 55 | GTCCAGGGAG | CACGTCTG | CCAACTGAAG | 7 | 5007042-155 | chrX:155147853-155147886 | loxBTR_1B | ATACCACTGTATA | 9 | loxBTR_1 | AACCCACTGCTTA | 8 | 9 |
| 56 | CACCTTCCAG | CAGGTCCC | CGGGGTTGTG | 5 | 5006400-155 | chrX:155148495-155148528 | loxBTR_1 | AACCCACTGCTTA | 9 | loxBTR_2 | AAGGCAAGCTTTA | 10 | 10 |
| 57 | GGTCCCCGGG | GTTGTGCC | CCTGGAGCTC | 5 | 5006412-155 | chrX:155148483-155148516 | loxBTR_1b | ATAACACTGCTTA | 10 | loxBTR_1B | ATACCACTGTATA | 9 | 10 |
| 58 | TGCATTAATT | CACATAAA | GTATAATGAA | 5 | 5006771-155 | chrX:155148124-155148157 | loxBTR_1B | ATACCACTGTATA | 9 | loxBTR_1C | ATAACATTGCTTA | 10 | 10 |
| 59 | AGGTCCCCGG | GGTTGTGC | CCCTGGAGCT | 6 | 5006411-155 | chrX:155148484-155148517 | loxBTR_1A | AACACATTGTATA | 9 | loxBTR_1A | AACACATTGTATA | 10 | 10 |
| 60 | GCGTTTATGG | GAGCCGCG | CCCAACAGGA | 6 | 5006538-155 | chrX:155148357-155148390 | loxBTR_1C | ATAACATTGCTTA | 9 | loxBTR_1A | AACACATTGTATA | 10 | 10 |
| 61 | AGGAAAGGAG | GAAAATTT | CCCCTTAAGG | 6 | 5006687-155 | chrX:155148208-155148241 | loxBTR_2A | AAGACTAGGTATA | 10 | loxBTR_1B | ATACCACTGTATA | 10 | 10 |
| 62 | CACCCGAGAA | AGCACGTA | GTCCAGGGAG | 6 | 5007024-155 | chrX:155147871-155147904 | loxBTR_1B | ATACCACTGTATA | 8 | loxBTR_1B | ATACCACTGTATA | 10 | 10 |
| 63 | AACGCTGTGG | TAACTGAG | GGAAAACGCA | 7 | 5006351-155 | chrX:155148544-155148577 | loxBTR_1C | ATAACATTGCTTA | 10 | loxBTR_1B | ATACCACTGTATA | 9 | 10 |
| 64 | TCTCAGGAGG | AGGCTCTG | TGGCCTCCAG | 7 | 5006625-155 | chrX:155148270-155148303 | loxBTR_1B | ATACCACTGTATA | 9 | loxBTR_1 | AACCCACTGCTTA | 10 | 10 |
| 65 | CAGGAGGAGG | CTCTGTGG | CCTCCAGACC | 7 | 5006628-155 | chrX:155148267-155148300 | loxBTR_1B | ATACCACTGTATA | 10 | loxBTR_1B | ATACCACTGTATA | 9 | 10 |
| 66 | GGCTCTGTGG | CCTCCAGA | CCACGTCAAA | 7 | 5006636-155 | chrX:155148259-155148292 | loxBTR_1C | ATAACATTGCTTA | 10 | loxBTR_2C | ATAACTAGCTTTA | 10 | 10 |
| 67 | CAGACCACGT | CAAAGCCA | AGGCAGAAAG | 7 | 5006650-155 | chrX:155148245-155148278 | loxBTR_2C | ATAACTAGCTTTA | 10 | loxBTR_1a | AACCCATTGTATA | 9 | 10 |
| 68 | AAAATTTCCC | CTTAAGGA | AGGTAAAATC | 7 | 5006698-155 | chrX:155148197-155148230 | loxBTR_1A | AACACATTGTATA | 8 | loxBTR_1C | ATAACATTGCTTA | 10 | 10 |
| 69 | AGGAAGGTAA | AATCCAGA | AGGGATCCCT | 7 | 5006712-155 | chrX:155148183-155148216 | loxBTR_1C | ATAACATTGCTTA | 7 | loxBTR_1C | ATAACATTGCTTA | 10 | 10 |
| 70 | GATCTCGCAG | CAGATTGG | CTGAAAAGTC | 7 | 5006870-155 | chrX:155148025-155148058 | loxBTR_2 | AAGGCAAGCTTTA | 9 | loxBTR_1A | AACACATTGTATA | 10 | 10 |
| 71 | CTCTCAAGGA | GACCCACT | GAGTTGGGCA | 7 | 5006917-155 | chrX:155147978-155148011 | loxBTR_1a | AACCCATTGTATA | 10 | loxBTR_1B | ATACCACTGTATA | 10 | 10 |
| 72 | AAGGAGACCC | ACTGAGTT | GGGCAAAGGT | 7 | 5006922-155 | chrX:155147973-155148006 | loxBTR_1A | AACACATTGTATA | 10 | loxBTR_1A | AACACATTGTATA | 8 | 10 |
| 73 | TTCCAGGGTT | TCTGGGAC | ACCCGAGAAA | 7 | 5007007-155 | chrX:155147888-155147921 | loxBTR_1B | ATACCACTGTATA | 9 | loxBTR_1 | AACCCACTGCTTA | 10 | 10 |
| 74 | CACGTAGTCC | AGGGAGCA | CGTCTGCCAA | 7 | 5007036-155 | chrX:155147859-155147892 | loxBTR_1 | AACCCACTGCTTA | 8 | loxBTR_2A | AAGACTAGGTATA | 10 | 10 |
| 75 | AGGGAGCACG | TCTGCCAA | CTGAAGGCCT | 7 | 5007046-155 | chrX:155147849-155147882 | loxBTR_1A | AACACATTGTATA | 9 | loxBTR_1B | ATACCACTGTATA | 10 | 10 |
| 76 | TTGACAAATG | ACTTTCTG | TACTGGCTGA | 7 | 5007073-155 | chrX:155147822-155147855 | loxBTR_1A | AACACATTGTATA | 10 | loxBTR_1 | AACCCACTGCTTA | 8 | 10 |
| 77 | CTGAAAACAT | GTGACAAA | GTGCTTTCCG | 4 | 5006246-155 | chrX:155148649-155148682 | loxBTR_1B | ATACCACTGTATA | 10 | loxBTR_1B | ATACCACTGTATA | 11 | 11 |
| 78 | TTGCCTCCCT | GCTCTCCC | ACGTAGCCCT | 6 | 5006966-155 | chrX:155147929-155147962 | loxBTR_1 | AACCCACTGCTTA | 10 | loxBTR_1A | AACACATTGTATA | 11 | 11 |
| 79 | GTTTAAACCT | AGCAGTTT | TGCATTAATT | 7 | 5006753-155 | chrX:155148142-155148175 | loxBTR_1C | ATAACATTGCTTA | 11 | loxBTR_2A | AAGACTAGGTATA | 8 | 11 |
| 80 | AGCCCTGCAT | TTCACTCC | ATTCCAGGGT | 7 | 5006988-155 | chrX:155147907-155147940 | loxBTR_1b | ATAACACTGCTTA | 11 | loxBTR_1A | AACACATTGTATA | 7 | 11 |
| 81 | TACTGGCTGA | GGGCCAGG | GCCCAGCGTA | 7 | 5007091-155 | chrX:155147804-155147837 | loxBTR_1C | ATAACATTGCTTA | 7 | loxBTR_2 | AAGGCAAGCTTTA | 11 | 11 |
| 82 | TGGGACACCC | GAGAAAGC | ACGTAGTCCA | 7 | 5007019-155 | chrX:155147876-155147909 | loxBTR_2C | ATAACTAGCTTTA | 11 | loxBTR_1 | AACCCACTGCTTA | 12 | 12 |
