## Supplementary Table 2 for "Correction of a Factor VIII genomic inversion with designer-recombinases"

| | | | Matches vs. loxF8L or loxF8R | | | | | | |
|---|---|---|---|---|---|---|---|---|---|
| | | | Left half-site | | Righ half-site | | | | |
| Start position (hg38) | | Sequence on the + strand | Match | Mismatches | Match | Mismatches | Spacer/Rev. comp. spacer | Sequence label | total mismatches |
| chrX | 155148712 | CTAAGATTGTGTG-GCAGCGTT-TCCACAGATTTAT | loxF8R | 0 | loxF8L | 0 | AACGCTGC/GCAGCGTT | loxF8 | 0 |
| chrX | 155006181 | ATAAATCTGTGGA-AACGCTGC-CACACAATCTTAG | loxF8L | 0 | loxF8R | 0 | AACGCTGC/GCAGCGTT | loxF8 | 0 |
| --- | | | | | | | | | |
| chr1 | 114990203 | ACAAATTTGTGGA-AATTAAAC-AACACACTCTTAG | loxF8L | 2 | loxF8R | 2 | AATTAAAC/GTTTAATT | | 4 |
| chr6 | 143914350 | TTAAGAGTGTGTT-GTTTAATT-TCCACAAATTTAT | loxF8R | 3 | loxF8L | 1 | AATTAAAC/GTTTAATT | HG1 | 4 |
| chr8 | 94350077 | ATAAATTTGTGGA-AATTAAAC-AACACACTCTTAA | loxF8L | 1 | loxF8R | 3 | AATTAAAC/GTTTAATT | HG1 | 4 |
| chr9 | 62050082 | TTAAGAATGTGTT-GTTTAATT-TCCACAAATTTTT | loxF8R | 3 | loxF8L | 2 | AATTAAAC/GTTTAATT | | 5 |
| chr15 | 85059144 | TTAAGATTGTGTT-GTTTAATT-TCCACAATTTTGT | loxF8R | 2 | loxF8L | 3 | AATTAAAC/GTTTAATT | HG2 | 5 |
| chr3 | 5776625 | CTAAGAATGTGTT-GTTTAATT-TCTACATATTTGT | loxF8R | 2 | loxF8L | 3 | AATTAAAC/GTTTAATT | | 5 |
| chr16 | 52944118 | ACAAATATGTGGA-AATTAAAC-AACACACTCTTAA | loxF8L | 2 | loxF8R | 3 | AATTAAAC/GTTTAATT | | 5 |
| chr2 | 121309734 | ACAAATATGTGGA-AATTAAAC-AACACACTCTTAA | loxF8L | 2 | loxF8R | 3 | AATTAAAC/GTTTAATT | | 5 |
| chr1 | 152790797 | ACAAATATGTGGA-AATTAAAC-AACACACTCTTAT | loxF8L | 2 | loxF8R | 3 | AATTAAAC/GTTTAATT | | 5 |
| chr3 | 128129270 | ACAAATATGTGGA-AATTAAAC-CATACATTCTTAA | loxF8L | 2 | loxF8R | 3 | AATTAAAC/GTTTAATT | | 5 |
| chr4 | 26706795 | TTAAGACTGTGTT-GTTTAATT-TCCATAGATTTGT | loxF8R | 3 | loxF8L | 2 | AATTAAAC/GTTTAATT | | 5 |
| chr14 | 70624934 | ACAAATGTGTGGA-AATTAAAC-AACACACTCTTAA | loxF8L | 2 | loxF8R | 3 | AATTAAAC/GTTTAATT | | 5 |
| chr1 | 103970289 | TTAAGAGTGTGTT-GTTTAATT-TCCACAAATTTGT | loxF8R | 3 | loxF8L | 2 | AATTAAAC/GTTTAATT | | 5 |
| chr1 | 153080535 | ACAAATTTGTGGA-AATTAAAC-AACACACTCTTAA | loxF8L | 2 | loxF8R | 3 | AATTAAAC/GTTTAATT | | 5 |
| chr3 | 119141128 | AGAAATATGTGGA-AATTAAAC-AACACATTCTTAA | loxF8L | 2 | loxF8R | 3 | AATTAAAC/GTTTAATT | | 5 |
| chr15 | 44752124 | TTAAGAGTGTGTT-GTTTAATT-TTCACATATTTAT | loxF8R | 3 | loxF8L | 2 | AATTAAAC/GTTTAATT | | 5 |
| chr4 | 38300917 | ATAAATATGTGAA-AATTAAAC-AACACACTCTTAA | loxF8L | 2 | loxF8R | 3 | AATTAAAC/GTTTAATT | | 5 |
| chr4 | 136183206 | AAAAATATGTGAA-AATTAAAC-AACACATTCTTAA | loxF8L | 3 | loxF8R | 3 | AATTAAAC/GTTTAATT | | 6 |
| chr9 | 38825655 | TTAAGAATGTGTT-GTTTAATT-TCCACAAATGTTT | loxF8R | 3 | loxF8L | 3 | AATTAAAC/GTTTAATT | | 6 |
| chr2 | 1358617 | CCAAGAGTGTGTT-GTTTAATT-TCCACATTTTTGT | loxF8R | 3 | loxF8L | 3 | AATTAAAC/GTTTAATT | | 6 |
| chr5 | 84711284 | TTAAGACTGTGTT-GTTTAATT-TCCACATTTTTGT | loxF8R | 3 | loxF8L | 3 | AATTAAAC/GTTTAATT | | 6 |
| chr2 | 138408554 | TTAAGAATGTGTT-GTTTAATT-TCCACATTTTTGT | loxF8R | 3 | loxF8L | 3 | AATTAAAC/GTTTAATT | | 6 |
| chr7 | 149338626 | ACAAAATTGTGGA-AATTAAAC-AACACATTCTTAA | loxF8L | 3 | loxF8R | 3 | AATTAAAC/GTTTAATT | | 6 |
| chr1 | 244399353 | TTAAGAGTGTGTT-GTTTAATT-TCCACGTATTTGT | loxF8R | 3 | loxF8L | 3 | AATTAAAC/GTTTAATT | | 6 |
| chrX | 55537486 | ATAAGAGTGTGTT-GTTTAATT-TCTACATATTTGT | loxF8R | 3 | loxF8L | 3 | AATTAAAC/GTTTAATT | | 6 |
| chr15 | 50889382 | ACAAATATGTAGA-AATTAAAC-CACACACTTTTAA | loxF8L | 3 | loxF8R | 3 | AATTAAAC/GTTTAATT | | 6 |
| chr17 | 33998635 | TTAAGAATGTGTT-GTTTAATT-TCCAAATATTTGT | loxF8R | 3 | loxF8L | 3 | AATTAAAC/GTTTAATT | | 6 |
| chr13 | 41165687 | TTAAGAGTGTGTT-GTTTAATT-TCCATACATTTGT | loxF8R | 3 | loxF8L | 3 | AATTAAAC/GTTTAATT | | 6 |
| chr8 | 10749587 | TTAAGAATGTGTT-GTTTAATT-TCCCCAAATTTGT | loxF8R | 3 | loxF8L | 3 | AATTAAAC/GTTTAATT | | 6 |
| chr19 | 49319932 | ACAAATTTGTAGA-AATTAAAC-AACACACTCTTAA | loxF8L | 3 | loxF8R | 3 | AATTAAAC/GTTTAATT | | 6 |
| chr15 | 69775513 | TTAAGAGTGTGTC-GTTTAATT-TTCACAAATTTGT | loxF8R | 3 | loxF8L | 3 | AATTAAAC/GTTTAATT | | 6 |
| chr13 | 72056379 | TTAAGAGTGTGTT-GTTTAATT-GCCACAAATTTGT | loxF8R | 3 | loxF8L | 3 | AATTAAAC/GTTTAATT | | 6 |
| chr14 | 50567750 | TTAAGAGTGTGTT-GTTTAATT-GCCACAAATTTGT | loxF8R | 3 | loxF8L | 3 | AATTAAAC/GTTTAATT | | 6 |
| chrX | 134359020 | ACAAATTTGTGGG-AATTAAAC-CACACATTCCTAA | loxF8L | 3 | loxF8R | 3 | AATTAAAC/GTTTAATT | | 6 |
| chr1 | 168159249 | CCAAGAGTGTGTT-GTTTAATT-TCCACATACTTGT | loxF8R | 3 | loxF8L | 3 | AATTAAAC/GTTTAATT | | 6 |
| chr16 | 47096018 | ACAAGTCTGTGTA-AATTAAAC-TACACACTCTTAC | loxF8L | 3 | loxF8R | 3 | AATTAAAC/GTTTAATT | | 6 |
| chr14 | 51470766 | TTAAGATGGTGTT-GTTTAATT-TCTAGATATTTAT | loxF8R | 3 | loxF8L | 3 | AATTAAAC/GTTTAATT | | 6 |
| chr9 | 73390993 | TTAAGAGTGTGTT-GTTTAATT-TTCACATATTCAT | loxF8R | 3 | loxF8L | 3 | AATTAAAC/GTTTAATT | | 6 |
| chrX | 88000189 | TTAAGAGTGTGTT-GTTTAATT-TCCACATATTTGG | loxF8R | 3 | loxF8L | 3 | AATTAAAC/GTTTAATT | | 6 |
| chr11 | 134897607 | TTAAGACTGTGTT-GTTTAATT-TCCACAAATTTGC | loxF8R | 3 | loxF8L | 3 | AATTAAAC/GTTTAATT | | 6 |
| chr6 | 8157412 | TTAAGAGTATGTT-GTTTAATT-TCCACAGATTTGT | loxF8R | 4 | loxF8L | 1 | AATTAAAC/GTTTAATT | | 5 |
| chr12 | 75229236 | TCAAGAGTGTGTT-GTTTAATT-TCCACATATTTAT | loxF8R | 4 | loxF8L | 1 | AATTAAAC/GTTTAATT | | 5 |
| chr9 | 6155500 | ATAAATTTGTGGA-AATTAAAC-AACAAACTCTTAA | loxF8L | 1 | loxF8R | 4 | AATTAAAC/GTTTAATT | | 5 |
| chr13 | 29134152 | TTAAGAGTGTGCT-GTTTAATT-TCCACACATTTTT | loxF8R | 4 | loxF8L | 2 | AATTAAAC/GTTTAATT | | 6 |
| chr9 | 67538277 | AAAAATTTGTGGA-AATTAAAC-AACACATTCTTTT | loxF8L | 2 | loxF8R | 4 | AATTAAAC/GTTTAATT | | 6 |
| chr1 | 108550349 | AAATATTTGTTGA-GTTTAATT-CACATAATTTTAG | loxF8L | 4 | loxF8R | 2 | AATTAAAC/GTTTAATT | | 6 |
| chr11 | 118041514 | ACAAATATGTGGA-AATTAAAC-AACACACTCCTAA | loxF8L | 2 | loxF8R | 4 | AATTAAAC/GTTTAATT | | 6 |
| chr15 | 87436075 | ACAAATATGTGGA-AATTAAAC-AACACACTCCTAA | loxF8L | 2 | loxF8R | 4 | AATTAAAC/GTTTAATT | | 6 |
| chr18 | 25909452 | TTAGGAGTGTGTT-GTTTAATT-TCCACATATTTGT | loxF8R | 4 | loxF8L | 2 | AATTAAAC/GTTTAATT | | 6 |
| chr3 | 50712734 | TTAGGAGTGTGTT-GTTTAATT-TCCACATATTTGT | loxF8R | 4 | loxF8L | 2 | AATTAAAC/GTTTAATT | | 6 |
| chr5 | 149705405 | TTAGGAGTGTGTT-GTTTAATT-TCCACATATTTGT | loxF8R | 4 | loxF8L | 2 | AATTAAAC/GTTTAATT | | 6 |
| chrX | 103619148 | ACAAATATGTGGA-AATTAAAC-AACACACTCCTAA | loxF8L | 2 | loxF8R | 4 | AATTAAAC/GTTTAATT | | 6 |
| chr7 | 488065 | ACAAATATGTGGA-AATTAAAC-AACACACTCCTAC | loxF8L | 2 | loxF8R | 4 | AATTAAAC/GTTTAATT | | 6 |
| chr16 | 32245509 | AGAAGAGTGTGTT-GTTTAATT-TCCACATATTTGT | loxF8R | 4 | loxF8L | 2 | AATTAAAC/GTTTAATT | | 6 |
| chr10 | 58539987 | ACAAATATGTGGA-AATTAAAC-AACACACTCTTGA | loxF8L | 2 | loxF8R | 4 | AATTAAAC/GTTTAATT | HG3 | 6 |
| chr11 | 61431528 | TCAAGAGTGTGTT-GTTTAATT-TCCACATATTTGT | loxF8R | 4 | loxF8L | 2 | AATTAAAC/GTTTAATT | HG3 | 6 |
| chr16 | 35372381 | ACAAATATGTGGA-AATTAAAC-AACACACTCTTGA | loxF8L | 2 | loxF8R | 4 | AATTAAAC/GTTTAATT | HG3 | 6 |
| chr2 | 201240889 | ACAAATATGTGGA-AATTAAAC-AACACACTCTTGA | loxF8L | 2 | loxF8R | 4 | AATTAAAC/GTTTAATT | HG3 | 6 |
| chr2 | 206778413 | TCAAGAGTGTGTT-GTTTAATT-TCCACATATTTGT | loxF8R | 4 | loxF8L | 2 | AATTAAAC/GTTTAATT | HG3 | 6 |
| chr7 | 72700538 | TTAACAGTGTGTT-GTTTAATT-TCCACATATTTGT | loxF8R | 4 | loxF8L | 2 | AATTAAAC/GTTTAATT | | 6 |
| chr5 | 165509925 | ACAAATATGTGGA-AATTAAAC-AACACATTCCTAA | loxF8L | 2 | loxF8R | 4 | AATTAAAC/GTTTAATT | | 6 |
| chr8 | 78424774 | TTGAGAATGTGTT-GTTTAATT-TCCACATATTTGT | loxF8R | 4 | loxF8L | 2 | AATTAAAC/GTTTAATT | | 6 |
| chr1 | 30591556 | ACAAATATGTGGA-AATTAAAC-AACACATTTTTAA | loxF8L | 2 | loxF8R | 4 | AATTAAAC/GTTTAATT | | 6 |
| chr7 | 9314298 | ACAAATATGTGGA-AATTAAAC-AACAGAATCTTGA | loxF8L | 2 | loxF8R | 4 | AATTAAAC/GTTTAATT | | 6 |
| chr2 | 131163837 | TTAAGAATATGTT-GTTTAATT-TCCACATATTTGT | loxF8R | 4 | loxF8L | 2 | AATTAAAC/GTTTAATT | | 6 |
| chr4 | 109791932 | TTAAGAATATGTT-GTTTAATT-TCCACATATTTGT | loxF8R | 4 | loxF8L | 2 | AATTAAAC/GTTTAATT | | 6 |
| chr20 | 22178731 | ACAAATATGTGGA-AATTAAAC-AATACACTCTTAA | loxF8L | 2 | loxF8R | 4 | AATTAAAC/GTTTAATT | | 6 |
| chrX | 82687505 | CTACCATTGTGCT-GTTTAATT-TCCACATATTTGT | loxF8R | 4 | loxF8L | 2 | AATTAAAC/GTTTAATT | | 6 |
| chr9 | 17109489 | TTAGAAGTGTGTG-GTTTAATT-TCCACATATTTGT | loxF8R | 4 | loxF8L | 2 | AATTAAAC/GTTTAATT | | 6 |
| chr7 | 38485834 | ACAAATGTGTGGA-AATTAAAC-AACACAATTTTGA | loxF8L | 2 | loxF8R | 4 | AATTAAAC/GTTTAATT | | 6 |
| chr1 | 234339254 | ACAAATTTGTGGA-AATTAAAC-AACACACATTTAA | loxF8L | 2 | loxF8L | 4 | AATTAAAC/GTTTAATT | | 6 |
| chr13 | 105650308 | ACAAATTTGTGGA-AATTAAAC-AACATACTCTTAC | loxF8L | 2 | loxF8R | 4 | AATTAAAC/GTTTAATT | | 6 |
| chr1 | 153082571 | ACAAATTTGTGGA-AATTAAAC-AACGCACTCTTAA | loxF8L | 2 | loxF8R | 4 | AATTAAAC/GTTTAATT | | 6 |
| chr7 | 34615794 | ACAAATTTGTGGA-AATTAAAC-AGCACACTCTTAA | loxF8L | 2 | loxF8R | 4 | AATTAAAC/GTTTAATT | | 6 |
| chr8 | 47456539 | ATAAACATGTGGA-AATTAAAC-AACACAGCCTTAT | loxF8L | 2 | loxF8L | 4 | AATTAAAC/GTTTAATT | | 6 |
| chr1 | 43490066 | TTAGGAGTGTGTT-GTTTAATT-TTCACATATTTAT | loxF8R | 4 | loxF8L | 2 | AATTAAAC/GTTTAATT | | 6 |
| chrX | 121005078 | TTAAGAATATGTT-GTTTAATT-TTCACATATTTAT | loxF8R | 4 | loxF8L | 2 | AATTAAAC/GTTTAATT | | 6 |
| chr3 | 6459210 | TTAAGAGTGTGTG-GTTTAATT-TTCTCATACTTAT | loxF8R | 2 | loxF8L | 4 | AATTAAAC/GTTTAATT | | 6 |
| chr10 | 5437963 | ATAAGTATGTGGA-AATTAAAC-AACACATTATTAA | loxF8L | 2 | loxF8R | 4 | AATTAAAC/GTTTAATT | | 6 |
| chr1 | 118722988 | AAAAAAATGTGGA-AATTAAAC-TACACACTGTTAA | loxF8L | 3 | loxF8R | 4 | AATTAAAC/GTTTAATT | | 7 |
| chr2 | 19534805 | AAAAATGTATGAA-AATTAAAC-AACACACTCTTAA | loxF8L | 4 | loxF8R | 3 | AATTAAAC/GTTTAATT | | 7 |

| | | | | | | | | |
|---|---|---|---|---|---|---|---|---|
| chr8 | 71857506 | AAATGATTGTGTT-GTTTAATT-TCCATAAATTTGT | loxF8R | 4 | loxF8L | 3 | AATTAAAC/GTTTAATT | 7 |
| chr5 | 105478755 | AATAATATGTGGA-AATTAAAC-AACAGAATCCTAA | loxF8L | 3 | loxF8R | 4 | AATTAAAC/GTTTAATT | 7 |
| chr5 | 147377202 | TTAAGAGTTTGTT-GTTTAATT-TCCACATTTTTGT | loxF8R | 4 | loxF8L | 3 | AATTAAAC/GTTTAATT | 7 |
| chr17 | 53523927 | TTAAGGGTGTGTT-GTTTAATT-TCCACATGTTTGT | loxF8R | 4 | loxF8L | 3 | AATTAAAC/GTTTAATT | 7 |
| chr3 | 121366027 | TTAAGAGAGTGTT-GTTTAATT-TCCACAAGTTTGT | loxF8R | 4 | loxF8L | 3 | AATTAAAC/GTTTAATT | 7 |
| chr10 | 75316772 | ACAAAGATGTGGA-AATTAAAC-AACACATTCTTCA | loxF8L | 3 | loxF8R | 4 | AATTAAAC/GTTTAATT | 7 |
| chr7 | 92978637 | TTAAGAGTATGTT-GTTTAATT-TCCACTTATTTGT | loxF8R | 4 | loxF8L | 3 | AATTAAAC/GTTTAATT | 7 |
| chr8 | 128731687 | TTAAGAGTGTGTT-GTTTAATT-TCCATGTATTTGT | loxF8R | 3 | loxF8L | 4 | AATTAAAC/GTTTAATT | 7 |
| chr22 | 43785370 | ACAAATACGTGGA-AATTAAAC-AACACACTCCTAA | loxF8L | 3 | loxF8R | 4 | AATTAAAC/GTTTAATT | 7 |
| chrX | 101193331 | TTAGGAGTGTGTT-GTTTAATT-TCCACGTATTTGT | loxF8R | 4 | loxF8L | 3 | AATTAAAC/GTTTAATT | 7 |
| chr1 | 158527121 | ACAAATAGGTGGA-AATTAAAC-AACACACTCATAA | loxF8L | 3 | loxF8R | 4 | AATTAAAC/GTTTAATT | 7 |
| chr5 | 18189162 | CTAGGACTGTGTT-GTTTAATT-TCTATATATTTGT | loxF8R | 3 | loxF8L | 4 | AATTAAAC/GTTTAATT | 7 |
| chr4 | 150954921 | ACAAATATATGGA-AATTAAAC-AACACACTCCTAA | loxF8L | 3 | loxF8R | 4 | AATTAAAC/GTTTAATT | 7 |
| chr20 | 22360494 | ACAAATATATGGA-AATTAAAC-AATACACTCTTAA | loxF8L | 3 | loxF8R | 4 | AATTAAAC/GTTTAATT | 7 |
| chrX | 150722012 | TTAGGAGTGTGTC-GTTTAATT-TCCATATATTTGT | loxF8R | 4 | loxF8L | 3 | AATTAAAC/GTTTAATT | 7 |
| chr12 | 76231002 | TCAAGAGTGTGTA-GTTTAATT-TCCATATATTTGT | loxF8R | 4 | loxF8L | 3 | AATTAAAC/GTTTAATT | 7 |
| chrX | 18816258 | TTAAGATTTTGTT-GTTTAATT-TTCTCATATTTGT | loxF8R | 3 | loxF8L | 4 | AATTAAAC/GTTTAATT | 7 |
| chr2 | 236389755 | TCAAGAGTGTGTT-GTTTAATT-TCCGCATATTTGT | loxF8R | 4 | loxF8L | 3 | AATTAAAC/GTTTAATT | 7 |
| chr1 | 59799784 | TTAAGAGTATGTT-GTTTAATT-TCCGCATATTTGT | loxF8R | 4 | loxF8L | 3 | AATTAAAC/GTTTAATT | 7 |
| chr12 | 43366664 | ACAAATATGCGGA-AATTAAAC-TCCACACTCTTAA | loxF8L | 3 | loxF8L | 4 | AATTAAAC/GTTTAATT | 7 |
| chr21 | 24347619 | TCAATATTGTGTT-GTTTAATT-TCTACATATTTGT | loxF8R | 4 | loxF8L | 3 | AATTAAAC/GTTTAATT | 7 |
| chr15 | 100948735 | ACAAATATGTAGA-AATTAAAC-AACACACTATTAA | loxF8L | 3 | loxF8R | 4 | AATTAAAC/GTTTAATT | 7 |
| chr10 | 103158106 | ACAAATATGTAGA-AATTAAAC-AACACACTCCTAA | loxF8L | 3 | loxF8R | 4 | AATTAAAC/GTTTAATT | 7 |
| chrX | 128069411 | ACAAATATGTAGA-AATTAAAC-AACACACTCCTAA | loxF8L | 3 | loxF8R | 4 | AATTAAAC/GTTTAATT | 7 |
| chr10 | 81209768 | ACAAATATGTAGA-AATTAAAC-AACACACTCTTTA | loxF8L | 3 | loxF8R | 4 | AATTAAAC/GTTTAATT | 7 |
| chr5 | 174783449 | ATAACAGTGTGTT-GTTTAATT-TCTACATATTTGT | loxF8R | 4 | loxF8L | 3 | AATTAAAC/GTTTAATT | 7 |
| chr15 | 20583517 | ACAAATATGTAGA-AATTAAAC-AATACACTCTTAA | loxF8L | 3 | loxF8R | 4 | AATTAAAC/GTTTAATT | 7 |
| chr5 | 114290854 | TTCAGAGTGTGTA-GTTTAATT-TCTACATATTTGT | loxF8R | 4 | loxF8L | 3 | AATTAAAC/GTTTAATT | 7 |
| chr3 | 158125486 | TCAAGAGTGTGTA-GTTTAATT-TCTACATATTTGT | loxF8R | 4 | loxF8L | 3 | AATTAAAC/GTTTAATT | 7 |
| chr1 | 7204426 | GTAAGAATGTGTT-GTTTAATT-TATACATATTTGT | loxF8R | 3 | loxF8L | 4 | AATTAAAC/GTTTAATT | 7 |
| chr4 | 11992901 | ACAAATATGTGAA-AATTAAAC-AACACAATCCTGA | loxF8L | 3 | loxF8R | 4 | AATTAAAC/GTTTAATT | 7 |
| chr12 | 115054828 | CTTTGAGTGTGTT-GTTTAATT-TTCACATATTTGT | loxF8R | 4 | loxF8L | 3 | AATTAAAC/GTTTAATT | 7 |
| chr14 | 21706003 | TTAGGAGTGTGTT-GTTTAATT-TTCACATATTTGT | loxF8R | 4 | loxF8L | 3 | AATTAAAC/GTTTAATT | 7 |
| chr2 | 85703190 | TTAGGAGTGTGTT-GTTTAATT-TTCACATATTTGT | loxF8R | 4 | loxF8L | 3 | AATTAAAC/GTTTAATT | 7 |
| chr4 | 187643337 | ACAAATATGTGAA-AATTAAAC-AACACACTCCTAA | loxF8L | 3 | loxF8R | 4 | AATTAAAC/GTTTAATT | 7 |
| chr20 | 11707729 | ACAAATATGTGAA-AATTAAAC-AACACACTCTTGA | loxF8L | 3 | loxF8R | 4 | AATTAAAC/GTTTAATT | 7 |
| chr6 | 119578723 | ACAAATATGTGAA-AATTAAAC-AACACACTCTTGT | loxF8L | 3 | loxF8R | 4 | AATTAAAC/GTTTAATT | 7 |
| chr21 | 44564996 | ACAAATATGTGAA-AATTAAAC-AGCACACTCTTAT | loxF8L | 3 | loxF8R | 4 | AATTAAAC/GTTTAATT | 7 |
| chr13 | 70690754 | GCAAGACTGTGTC-GTTTAATT-TTCACATATTTGT | loxF8R | 4 | loxF8L | 3 | AATTAAAC/GTTTAATT | 7 |
| chr14 | 103987518 | TTAGGAGTGTGTT-GTTTAATT-TGCACATATTTGT | loxF8R | 4 | loxF8L | 3 | AATTAAAC/GTTTAATT | 7 |
| chr21 | 44633496 | ATAAGTATGTGTT-GTTTAATT-GCCACATATTTGT | loxF8L | 4 | loxF8L | 3 | AATTAAAC/GTTTAATT | 7 |
| chr7 | 19277431 | TTAGGAGTGTGTT-GTTTAATT-TCCAAATATTTGT | loxF8R | 4 | loxF8L | 3 | AATTAAAC/GTTTAATT | 7 |
| chrY | 11803659 | ACAAATATTTGGA-AATTAAAC-AACACACTTTTAA | loxF8L | 3 | loxF8R | 4 | AATTAAAC/GTTTAATT | 7 |
| chr12 | 921005 | ACAAATCAGTAGA-AATTAAAC-ACCACACTCTTAA | loxF8L | 3 | loxF8R | 4 | AATTAAAC/GTTTAATT | 7 |
| chr18 | 72068922 | TTAAGAATGTGTT-GTTTAATT-TCCATGCATTTGT | loxF8R | 3 | loxF8L | 4 | AATTAAAC/GTTTAATT | 7 |
| chr14 | 30963806 | ACAAATTTATGGA-AATTAAAC-AACACACTCAAAG | loxF8L | 3 | loxF8R | 4 | AATTAAAC/GTTTAATT | 7 |
| chr1 | 153047640 | ACAAATTTGTAGA-AATTAAAC-AACAAACTCTTAA | loxF8L | 3 | loxF8R | 4 | AATTAAAC/GTTTAATT | 7 |
| chr1 | 153120648 | ACAAATTTGTAGA-AATTAAAC-AACACACTCTTTA | loxF8L | 3 | loxF8R | 4 | AATTAAAC/GTTTAATT | 7 |
| chr15 | 100273946 | TTAATAGTATGTG-GTTTAATT-TCTACAAATTTGT | loxF8R | 4 | loxF8L | 3 | AATTAAAC/GTTTAATT | 7 |
| chr13 | 71059930 | TTAATATTATGTT-GTTTAATT-TTCACAAATTTGT | loxF8R | 4 | loxF8L | 3 | AATTAAAC/GTTTAATT | 7 |
| chr2 | 237175045 | TTAAGAGTGTGGT-GTTTAATT-TTCACAAATTTGT | loxF8R | 4 | loxF8L | 3 | AATTAAAC/GTTTAATT | 7 |
| chrX | 71337653 | TTTAGAATGTGTT-GTTTAATT-TCCACATACTTGT | loxF8R | 4 | loxF8L | 3 | AATTAAAC/GTTTAATT | 7 |
| chr2 | 172576730 | ATAGGATTGTGTT-GTTTAATT-GCCACATACTTGT | loxF8R | 3 | loxF8L | 4 | AATTAAAC/GTTTAATT | 7 |
| chr11 | 45957495 | ACAAGTGTATGGA-AATTAAAC-AACACATTCTTAA | loxF8L | 4 | loxF8R | 3 | AATTAAAC/GTTTAATT | 7 |
| chr3 | 33525134 | ACAAGTTTGAGGA-AATTAAAC-AACACACTCTTAA | loxF8L | 4 | loxF8R | 3 | AATTAAAC/GTTTAATT | 7 |
| chr18 | 1617285 | ACAATTATGTGGA-AATTAAAC-AACATATTCTTAA | loxF8L | 3 | loxF8R | 4 | AATTAAAC/GTTTAATT | 7 |
| chr11 | 44023913 | TGAAGAGTGTGTT-GTTTAATT-TCCACATATGTGT | loxF8R | 4 | loxF8L | 3 | AATTAAAC/GTTTAATT | 7 |
| chr12 | 23317436 | ACAGATATGTGGA-AATTAAAC-AACACCATCCTAA | loxF8L | 3 | loxF8R | 4 | AATTAAAC/GTTTAATT | 7 |
| chr17 | 10431863 | TTAGGAGTGTGTT-GTTTAATT-TCCACAAATCTGT | loxF8R | 4 | loxF8L | 3 | AATTAAAC/GTTTAATT | 7 |
| chr4 | 1390234 | TTAAGAGTTTGTT-GTTTAATT-TCCACATATATGT | loxF8R | 4 | loxF8L | 3 | AATTAAAC/GTTTAATT | 7 |
| chr3 | 21211427 | TTAAGATCATGTT-GTTTAATT-TCCACATATATGT | loxF8R | 4 | loxF8L | 3 | AATTAAAC/GTTTAATT | 7 |
| chr5 | 450306 | TTAAGAGTATGTT-GTTTAATT-TCCACAAATTGGT | loxF8R | 4 | loxF8L | 3 | AATTAAAC/GTTTAATT | 7 |
| chr13 | 88969170 | ATAAACATGTGAA-AATTAAAC-AAAACATTCTTAA | loxF8L | 3 | loxF8R | 4 | AATTAAAC/GTTTAATT | 7 |
| chr1 | 116094211 | TTGGGATTGTGTT-GTTTAATT-TTTACATATTTAT | loxF8R | 4 | loxF8L | 3 | AATTAAAC/GTTTAATT | 7 |
| chr1 | 28004510 | ATAAATATGTAGG-AATTAAAC-AATACAATCCTAA | loxF8L | 3 | loxF8R | 4 | AATTAAAC/GTTTAATT | 7 |
| chr1 | 190712502 | TTAAAAATGTGTT-GTTTAATT-TTCAAACATTTAT | loxF8R | 4 | loxF8L | 3 | AATTAAAC/GTTTAATT | 7 |
| chr4 | 167752520 | ATGAATATGTGCA-AATTAAAC-AACACCATCCTAA | loxF8L | 3 | loxF8R | 4 | AATTAAAC/GTTTAATT | 7 |
| chr2 | 200380949 | ATTAAACTATGGA-AATTAAAC-CACAGAAACAAAG | loxF8L | 3 | loxF8R | 4 | AATTAAAC/GTTTAATT | 7 |
| chr18 | 13147734 | TTCAGAGTGTGTT-GTTTAATT-TCCACATATTTGG | loxF8R | 4 | loxF8L | 3 | AATTAAAC/GTTTAATT | 7 |
| chr2 | 112132164 | CCAAATATGTGGA-AATTAAAC-CACACAGTTCTAA | loxF8L | 3 | loxF8R | 4 | AATTAAAC/GTTTAATT | 7 |
| chr2 | 156835070 | TTAAAATTGTGTT-GTTTAATT-TCCAAATATTCAG | loxF8R | 3 | loxF8L | 4 | AATTAAAC/GTTTAATT | 7 |
| chr2 | 14139634 | GCAAATATGTGGA-AATTAAAC-TACACATTGTAAG | loxF8L | 3 | loxF8R | 4 | AATTAAAC/GTTTAATT | 7 |
| chr10 | 96891867 | TTAAGAATGTATT-GTTTAATT-TCCACATATTTGC | loxF8R | 4 | loxF8L | 3 | AATTAAAC/GTTTAATT | 7 |
| chr9 | 137123743 | TTAAGAGTATGTT-GTTTAATT-GCCACAGATTTGC | loxF8R | 4 | loxF8L | 3 | AATTAAAC/GTTTAATT | 7 |
| chr1 | 59846774 | TTAGGAATGTGTT-GTTTAATT-TACAAAGATTTAA | loxF8R | 4 | loxF8L | 3 | AATTAAAC/GTTTAATT | 7 |
| chr9 | 113436794 | TTAAGAGTGTGTT-GTTTAATT-TCCATATACTTAA | loxF8R | 3 | loxF8L | 4 | AATTAAAC/GTTTAATT | 7 |
| chr1 | 164353024 | TTGAATATGTGGA-AATTAAAC-AACACACTCATAA | loxF8L | 3 | loxF8R | 4 | AATTAAAC/GTTTAATT | 7 |
| --- | | | | | | | | 0 |
| chr12 | 116225942 | ATAAATATGAGGA-AAAGACTA-ACAACAGATTTAT | loxF8L | 2 | loxF8L | 2 | AAAGACTA/TAGTCTTT | 4 |
| chrX | 130500096 | TTAAAATTTTATG-TAGTCTTT-CCTACAGATTCAT | loxF8R | 4 | loxF8L | 3 | AAAGACTA/TAGTCTTT | 7 |
| --- | | | | | | | | 0 |
| chr3 | 164127645 | ATATATCTGTGCA-CATTTATA-TACACATATTTAT | loxF8L | 2 | loxF8L | 2 | CATTTATA/TATAAATG | 4 |
| chr7 | 138584148 | ATAACTCTGTCCA-CATTTATA-TTCAAAGGTTTAT | loxF8L | 3 | loxF8L | 3 | CATTTATA/TATAAATG | 6 |
| chrX | 45940094 | GTAAGTATGTGGA-TATAAATG-GCAACAGCTTTGT | loxF8L | 3 | loxF8L | 4 | CATTTATA/TATAAATG | 7 |
| chrX | 5302461 | ATAAAAATGTGTA-TATAAATG-TTCACAAACATAT | loxF8L | 3 | loxF8L | 4 | CATTTATA/TATAAATG | 7 |
| chr4 | 164402556 | ATATGTGTGTGTG-TATAAATG-TACACATATATAT | loxF8R | 4 | loxF8L | 3 | CATTTATA/TATAAATG | 7 |
| --- | | | | | | | | 0 |

| | | | | | | | | | |
|---|---|---|---|---|---|---|---|---|---|
| chr2 | 212549011 | ATAAATATGTGAA-TATACATA-TTCACATATTTAT | loxF8L | 2 | loxF8L | 2 | TATACATA/TATGTATA | HG1L | 4 |
| chr2 | 212549087 | ATAAATATGTGAA-TATACATA-TTCACATATTTAT | loxF8L | 2 | loxF8L | 2 | TATACATA/TATGTATA | HG1L | 4 |
| chr2 | 212549049 | ATAAATATGTGAA-TATACATA-TTCACATATTTAT | loxF8L | 2 | loxF8L | 2 | TATACATA/TATGTATA | HG1L | 4 |
| chr2 | 212548973 | ATAAATATGTGAA-TATACATA-TTCACATATTTAT | loxF8L | 2 | loxF8L | 2 | TATACATA/TATGTATA | HG1L | 4 |
| chr2 | 212549125 | ATAAATATGTGAA-TATACATA-TTCATATATTTAT | loxF8L | 2 | loxF8L | 3 | TATACATA/TATGTATA | | 5 |
| chr2 | 212548935 | ATAAATATATGAA-TATACATA-TTCACATATTTAT | loxF8L | 3 | loxF8L | 2 | TATACATA/TATGTATA | | 5 |
| chr7 | 120980619 | TTTAGAATGTGTG-TATACATA-TATACAGATGTAT | loxF8R | 3 | loxF8L | 3 | TATACATA/TATGTATA | | 6 |
| chr1 | 184645593 | ATATATATGTGTA-TATGTATA-TACACATATATAT | loxF8L | 3 | loxF8L | 3 | TATACATA/TATGTATA | | 6 |
| chr15 | 29378724 | ATAAATCTGTCTA-TATACATA-TACATACATATAT | loxF8L | 2 | loxF8L | 4 | TATACATA/TATGTATA | | 6 |
| chrX | 45723880 | ACAGATATGTGTA-TATGTATA-TACACAGATATGT | loxF8L | 4 | loxF8L | 3 | TATACATA/TATGTATA | | 7 |
| chr10 | 99814464 | ACATATGTGTGTA-TATACATA-CACACAAACATAT | loxF8L | 4 | loxF8R | 3 | TATACATA/TATGTATA | | 7 |
| chrX | 46615635 | AGAAATATGTGTA-TATACATA-TACATATATATAT | loxF8L | 3 | loxF8L | 4 | TATACATA/TATGTATA | | 7 |
| chr8 | 29837751 | AGATATGTGTGTA-TATACATA-TGCACACATATAT | loxF8L | 4 | loxF8L | 3 | TATACATA/TATGTATA | | 7 |
| chr3 | 1161520 | ATAAACATGTGTA-TATACATA-TATATACATTTAT | loxF8L | 3 | loxF8L | 4 | TATACATA/TATGTATA | | 7 |
| chr14 | 70353457 | ATAAATATATATA-TATACATA-CACACACACATAG | loxF8L | 4 | loxF8R | 3 | TATACATA/TATGTATA | | 7 |
| chr10 | 7513195 | ATACATGTTTGTA-TATGTATA-TACATATATTTAT | loxF8L | 4 | loxF8L | 3 | TATACATA/TATGTATA | | 7 |
| chr10 | 7513117 | ATACATGTTTGTA-TATGTATA-TACATATATTTAT | loxF8L | 4 | loxF8L | 3 | TATACATA/TATGTATA | | 7 |
| chr9 | 11954267 | ATAAATATGTTTA-TATGTATA-TATGCATATTTAT | loxF8L | 3 | loxF8L | 4 | TATACATA/TATGTATA | | 7 |
| chr6 | 102362134 | ATAAGTGTGTATA-TATGTATA-TATACATATTTAT | loxF8L | 4 | loxF8L | 3 | TATACATA/TATGTATA | | 7 |
| chr4 | 122269167 | ATAAATATGTTTA-TATGTATA-CATACAGTCTATG | loxF8L | 3 | loxF8R | 4 | TATACATA/TATGTATA | | 7 |
| chr8 | 118165577 | ATAAATCTATATA-TATACATA-TATACATATATAT | loxF8L | 3 | loxF8L | 4 | TATACATA/TATGTATA | | 7 |
| chr10 | 108988339 | ATACATGTGTATA-TATACATA-TATACACATTTAT | loxF8L | 4 | loxF8L | 3 | TATACATA/TATGTATA | | 7 |
| chr18 | 72081353 | ATAAATTTGTGTG-TATGTATA-TATACATATATAT | loxF8L | 3 | loxF8L | 4 | TATACATA/TATGTATA | | 7 |
| chr5 | 90067987 | ATATATTTGTGTA-TATACATA-CACATACTGTTAT | loxF8L | 3 | loxF8R | 4 | TATACATA/TATGTATA | | 7 |
| chr14 | 92382133 | ATAAGTGTGTGTA-TATACATA-TATACACATATAT | loxF8L | 3 | loxF8L | 4 | TATACATA/TATGTATA | | 7 |
| chr19 | 31514934 | ATAATAGTGTGTG-TATACATA-TACATATATATAT | loxF8R | 3 | loxF8L | 4 | TATACATA/TATGTATA | | 7 |
| chr9 | 11866153 | ATAATTTTGTGTA-TATACATA-TATACAAATATAT | loxF8L | 3 | loxF8L | 4 | TATACATA/TATGTATA | | 7 |
| chr9 | 11866121 | ATAATTTTGTGTA-TATACATA-TATACAAATATAT | loxF8L | 3 | loxF8L | 4 | TATACATA/TATGTATA | | 7 |
| chr12 | 70027781 | ATATATGTGTGTA-TATACATA-TACACATGTGTAT | loxF8L | 3 | loxF8L | 4 | TATACATA/TATGTATA | | 7 |
| chr7 | 157806341 | ATATATGTGTATA-TATGTATA-TCTACATATGTAT | loxF8L | 4 | loxF8L | 3 | TATACATA/TATGTATA | | 7 |
| chr7 | 77217355 | ATATATATGTGTG-TATGTATA-TGCACATATGTAT | loxF8L | 4 | loxF8L | 3 | TATACATA/TATGTATA | | 7 |
| chr19 | 9477194 | ATACATATGTGCA-TATACATA-TATACATATATAT | loxF8L | 3 | loxF8L | 4 | TATACATA/TATGTATA | | 7 |
| chr2 | 131791252 | ATACATATGTGTA-TATGTATA-TATACACATATAT | loxF8L | 3 | loxF8L | 4 | TATACATA/TATGTATA | | 7 |
| chr2 | 131791284 | ATACATATGTGTA-TATGTATA-TATACACATATAT | loxF8L | 3 | loxF8L | 4 | TATACATA/TATGTATA | | 7 |
| chr2 | 131791220 | ATACATATGTGTA-TATGTATA-TATACACATATAT | loxF8L | 3 | loxF8L | 4 | TATACATA/TATGTATA | | 7 |
| chr6 | 79987564 | ATACATATGTGTA-TATACATA-TATACACATATAT | loxF8L | 3 | loxF8L | 4 | TATACATA/TATGTATA | | 7 |
| chrX | 114508179 | ATATATGTGTGTA-TATACATA-TTCATATATATAT | loxF8L | 3 | loxF8L | 4 | TATACATA/TATGTATA | | 7 |
| chrX | 33010339 | ATATATATGTGTG-TATGTATA-TCCATATATATAT | loxF8L | 4 | loxF8L | 3 | TATACATA/TATGTATA | | 7 |
| chr3 | 141889806 | ATATATGTGTGTA-TATACATA-TACATATATATAT | loxF8L | 3 | loxF8L | 4 | TATACATA/TATGTATA | | 7 |
| chr12 | 44024815 | ATATATGTGTATA-TATGTATA-TCTACATATATAT | loxF8L | 4 | loxF8L | 3 | TATACATA/TATGTATA | | 7 |
| chr16 | 62654301 | ATATATATGTATA-TATACATA-CACACACACATAG | loxF8L | 4 | loxF8R | 3 | TATACATA/TATGTATA | | 7 |
| chr8 | 59182020 | ATATATTTGTGTA-TATGTATA-TATACATATATAT | loxF8L | 3 | loxF8L | 4 | TATACATA/TATGTATA | | 7 |
| chr8 | 136207540 | ATATATGTGTGTA-TATACATA-TATACATATATAT | loxF8L | 3 | loxF8L | 4 | TATACATA/TATGTATA | | 7 |
| chr5 | 142026344 | ATATATATGTATA-TATACATA-TACACATATATAT | loxF8L | 4 | loxF8L | 3 | TATACATA/TATGTATA | | 7 |
| chr5 | 169714954 | ATATATGTGTGCA-TATGTATA-TATACATATATAT | loxF8L | 3 | loxF8L | 4 | TATACATA/TATGTATA | | 7 |
| chr14 | 71068463 | ATATATATGTGCA-TATGTATA-CACACACACATAT | loxF8L | 3 | loxF8R | 4 | TATACATA/TATGTATA | | 7 |
| chr14 | 93158536 | ATATATGTGTGTA-TATACATA-CACACATATATAT | loxF8L | 3 | loxF8L | 4 | TATACATA/TATGTATA | | 7 |
| chr6 | 75834810 | ATATATGTGTATA-TATACATA-CACACAATTCTGG | loxF8L | 4 | loxF8R | 3 | TATACATA/TATGTATA | | 7 |
| chr21 | 23284359 | ATATATGTGTGTA-TATGTATA-TATACACATATAT | loxF8L | 3 | loxF8L | 4 | TATACATA/TATGTATA | | 7 |
| chr3 | 134701385 | ATATATGTGTATA-TATACATA-TACACACATATAT | loxF8L | 4 | loxF8L | 3 | TATACATA/TATGTATA | | 7 |
| chr6 | 8597284 | ATATATGTGTGTA-TATGTATA-CACACACACACAG | loxF8L | 3 | loxF8R | 4 | TATACATA/TATGTATA | | 7 |
| chr3 | 140717669 | TTATATGTGTGTA-TATGTATA-TACACAGACATAT | loxF8L | 4 | loxF8L | 3 | TATACATA/TATGTATA | | 7 |
| chr6 | 57545256 | ATATTTCTGTGTA-TATGTATA-TACATAGATACAT | loxF8L | 3 | loxF8L | 4 | TATACATA/TATGTATA | | 7 |
| chr3 | 136134577 | CTTAAAATGTGTG-TATGTATA-TACACAAGCTAAA | loxF8R | 3 | loxF8R | 4 | TATACATA/TATGTATA | | 7 |
| --- | | | | | | | | | 0 |
| chr10 | 44602701 | ATATATATGTGGA-TATATATA-TCCACATATATAT | loxF8L | 2 | loxF8L | 2 | TATATATA/TATATATA | | 4 |
| chr10 | 44602739 | ATATATATGTGAA-TATATATA-TCCACATATATAT | loxF8L | 3 | loxF8L | 2 | TATATATA/TATATATA | | 5 |
| chr6 | 65440834 | AAAAATATGTGTA-TATATATA-TATATAGATTTAT | loxF8L | 3 | loxF8L | 3 | TATATATA/TATATATA | | 6 |
| chr20 | 9643585 | ATATGTTTGTGTG-TATATATA-CACACATATTTAT | loxF8R | 3 | loxF8L | 3 | TATATATA/TATATATA | HG4 | 6 |
| chr6 | 88920471 | TTAAGGATGTGTG-TATATATA-TTCATACATTTAT | loxF8R | 3 | loxF8L | 3 | TATATATA/TATATATA | | 6 |
| chr17 | 2588188 | ATAGATATGTGTA-TATATATA-CAGACATCCTTAG | loxF8L | 3 | loxF8R | 3 | TATATATA/TATATATA | | 6 |
| chr7 | 23079460 | ATAGATCTGTATA-TATATATA-TACACATATATAT | loxF8L | 3 | loxF8L | 3 | TATATATA/TATATATA | | 6 |
| chr9 | 2589105 | ATATATATATGGA-TATATATA-TCCATATATATAT | loxF8L | 3 | loxF8L | 3 | TATATATA/TATATATA | | 6 |
| chr4 | 166665426 | ATATATATCTGGA-TATATATA-TCCAGATATATAT | loxF8L | 3 | loxF8L | 3 | TATATATA/TATATATA | | 6 |
| chr4 | 110883492 | ATATATATGTGAA-TATATATA-TTCACATATATAT | loxF8L | 3 | loxF8L | 3 | TATATATA/TATATATA | | 6 |
| chr7 | 44043496 | CTATGTGTGTGTG-TATATATA-TACACATATATAT | loxF8R | 3 | loxF8L | 3 | TATATATA/TATATATA | | 6 |
| chr4 | 172177834 | ATATATATGTGTA-TATATATA-CACACATACTAAG | loxF8L | 3 | loxF8R | 3 | TATATATA/TATATATA | | 6 |
| chr4 | 182161710 | ATATATATGTGTA-TATATATA-TACACAAATATAT | loxF8L | 3 | loxF8L | 3 | TATATATA/TATATATA | | 6 |
| chr4 | 182161758 | ATATATATGTGTA-TATATATA-TACACAAATATAT | loxF8L | 3 | loxF8L | 3 | TATATATA/TATATATA | | 6 |
| chr10 | 33170028 | ATATATATGTGTA-TATATATA-TACACACATATAT | loxF8L | 3 | loxF8L | 3 | TATATATA/TATATATA | | 6 |
| chr10 | 33170056 | ATATATATGTGTA-TATATATA-TACACACATATAT | loxF8L | 3 | loxF8L | 3 | TATATATA/TATATATA | | 6 |
| chr10 | 128732573 | ATATATGTGTGTA-TATATATA-TACACATATATAT | loxF8L | 3 | loxF8L | 3 | TATATATA/TATATATA | | 6 |
| chr13 | 67613383 | ATATATATGTGTA-TATATATA-TACACACATATAT | loxF8L | 3 | loxF8L | 3 | TATATATA/TATATATA | | 6 |
| chr18 | 5308628 | ATATATATGTGTA-TATATATA-TACACACATATAT | loxF8L | 3 | loxF8L | 3 | TATATATA/TATATATA | | 6 |
| chr19 | 23761004 | ATATATGTGTGTA-TATATATA-TACACATATATAT | loxF8L | 3 | loxF8L | 3 | TATATATA/TATATATA | | 6 |
| chr3 | 29007533 | ATATATATGTGTA-TATATATA-TACACACATATAT | loxF8L | 3 | loxF8L | 3 | TATATATA/TATATATA | | 6 |
| chr4 | 180049910 | ATATATATGTGTA-TATATATA-TACACACATATAT | loxF8L | 3 | loxF8L | 3 | TATATATA/TATATATA | | 6 |
| chr6 | 48621165 | ATATATATGTGTA-TATATATA-TACACACATATAT | loxF8L | 3 | loxF8L | 3 | TATATATA/TATATATA | | 6 |
| chr7 | 135206754 | ATATATATGTGTA-TATATATA-TACACACATATAT | loxF8L | 3 | loxF8L | 3 | TATATATA/TATATATA | | 6 |
| chr12 | 60908598 | ATATATATGTGTA-TATATATA-TACACATATATAT | loxF8L | 3 | loxF8L | 3 | TATATATA/TATATATA | | 6 |
| chr20 | 53050130 | ATATATATGTGTA-TATATATA-TACACATATATAT | loxF8L | 3 | loxF8L | 3 | TATATATA/TATATATA | | 6 |
| chr3 | 71397025 | ATATATATGTGTA-TATATATA-TACACATATATAT | loxF8L | 3 | loxF8L | 3 | TATATATA/TATATATA | | 6 |
| chr4 | 37738026 | ATATATATGTGTA-TATATATA-TACACATATATAT | loxF8L | 3 | loxF8L | 3 | TATATATA/TATATATA | | 6 |
| chr4 | 37738000 | ATATATATGTGTA-TATATATA-TACACATATATAT | loxF8L | 3 | loxF8L | 3 | TATATATA/TATATATA | | 6 |
| chr7 | 71701832 | ATATATATGTGTA-TATATATA-TACACATATATAT | loxF8L | 3 | loxF8L | 3 | TATATATA/TATATATA | | 6 |
| chr7 | 102338066 | ATATATATGTGTA-TATATATA-TACACATATATAT | loxF8L | 3 | loxF8L | 3 | TATATATA/TATATATA | | 6 |
| chrX | 151742983 | ATATATATGTGTA-TATATATA-TACACATATATAT | loxF8L | 3 | loxF8L | 3 | TATATATA/TATATATA | | 6 |

| | | | | | | | | | |
|---|---|---|---|---|---|---|---|---|---|
| chr4 | 11031005 | ATATATGTGTGCA-TATATATA-TACACATATATAT | loxF8L | 3 | loxF8L | 3 | TATATATA/TATATATA | | 6 |
| chr3 | 78638304 | ATATATGTGTGTA-TATATATA-CACACACACTTAA | loxF8L | 3 | loxF8R | 3 | TATATATA/TATATATA | | 6 |
| chr5 | 130077788 | ATATATGTGTGTA-TATATATA-TACACACATATAT | loxF8L | 3 | loxF8L | 3 | TATATATA/TATATATA | | 6 |
| chr7 | 149133682 | ATATATGTGTGTA-TATATATA-TACACACATATAT | loxF8L | 3 | loxF8L | 3 | TATATATA/TATATATA | | 6 |
| chrX | 54071973 | ATATATGTGTGTA-TATATATA-TACACACATATAT | loxF8L | 3 | loxF8L | 3 | TATATATA/TATATATA | | 6 |
| chrX | 118709434 | ATATATGTGTGTA-TATATATA-TACACACATATAT | loxF8L | 3 | loxF8L | 3 | TATATATA/TATATATA | | 6 |
| chr5 | 62179684 | AAAAGTCTGCAGA-TATATATA-TCCAGATATTTAT | loxF8L | 4 | loxF8L | 2 | TATATATA/TATATATA | | 6 |
| chr13 | 104128435 | ATAAATATGTGTA-TATATATA-TGGACATATATAT | loxF8L | 2 | loxF8L | 4 | TATATATA/TATATATA | | 6 |
| chrX | 33269957 | ATATATATGTATA-TATATATA-ATCACAGATTTAT | loxF8L | 4 | loxF8L | 2 | TATATATA/TATATATA | | 6 |
| chr12 | 78777543 | ATAAGATAGTGTG-TATATATA-TATACATATATAT | loxF8R | 2 | loxF8L | 4 | TATATATA/TATATATA | | 6 |
| chr1 | 216117803 | ATATATATATGTA-TATATATA-TACACAGATATAT | loxF8L | 4 | loxF8L | 2 | TATATATA/TATATATA | | 6 |
| chr22 | 38377188 | ATATATCTGTGTA-TATATATA-CACACACACATAT | loxF8L | 2 | loxF8R | 4 | TATATATA/TATATATA | HG5 | 6 |
| chr7 | 9940394 | ATATGTATGTGTA-TATATATA-TCCACAAATATAT | loxF8L | 4 | loxF8L | 2 | TATATATA/TATATATA | | 6 |
| chr6 | 4895542 | ATATGTGTGTGTG-TATATATA-TACACACATTTTT | loxF8R | 4 | loxF8L | 3 | TATATATA/TATATATA | | 7 |
| chr13 | 85215891 | ATATATATGTGGG-TATATATA-CACACACATTTTT | loxF8L | 3 | loxF8L | 4 | TATATATA/TATATATA | | 7 |
| chrX | 92911453 | ATATATGTGTGTA-TATATATA-TACACACATATTT | loxF8L | 3 | loxF8L | 4 | TATATATA/TATATATA | | 7 |
| chr11 | 58752106 | ACAAATATGTGTA-TATATATA-TACACATACATAT | loxF8L | 3 | loxF8L | 4 | TATATATA/TATATATA | | 7 |
| chr18 | 38684186 | ACAAATATGTGTA-TATATATA-TATACAGGTATAT | loxF8L | 3 | loxF8L | 4 | TATATATA/TATATATA | | 7 |
| chr3 | 2440745 | ATATATGTGTGAA-TATATATA-CACACATATTTGT | loxF8L | 3 | loxF8L | 4 | TATATATA/TATATATA | | 7 |
| chr7 | 94933023 | ACACATCTCTGCA-TATATATA-TACACATATTTAG | loxF8L | 4 | loxF8L | 3 | TATATATA/TATATATA | | 7 |
| chr11 | 32755578 | ATATATGTGTGTA-TATATATA-TCCATATATATGT | loxF8L | 3 | loxF8L | 4 | TATATATA/TATATATA | | 7 |
| chr13 | 27052057 | ACATATATATGGA-TATATATA-TACACACATATAT | loxF8L | 4 | loxF8L | 3 | TATATATA/TATATATA | | 7 |
| chr14 | 59908821 | ACATATATATGGA-TATATATA-TACACATATATAT | loxF8L | 4 | loxF8L | 3 | TATATATA/TATATATA | | 7 |
| chr11 | 89164426 | ATATATGTGTGTA-TATATATA-TACACATATATGT | loxF8L | 3 | loxF8L | 4 | TATATATA/TATATATA | | 7 |
| chr7 | 75511867 | ACATATATGTGTA-TATATATA-TACACACATATAT | loxF8L | 4 | loxF8L | 3 | TATATATA/TATATATA | | 7 |
| chr8 | 125890942 | ACATATATGTGTA-TATATATA-TACACACATATAT | loxF8L | 4 | loxF8L | 3 | TATATATA/TATATATA | | 7 |
| chr13 | 67613311 | ATATATATGTGTA-TATATATA-TACACATATATGT | loxF8L | 3 | loxF8L | 4 | TATATATA/TATATATA | | 7 |
| chr20 | 14702652 | ATATATATGTGTA-TATATATA-TACACATATATGT | loxF8L | 3 | loxF8L | 4 | TATATATA/TATATATA | | 7 |
| chr3 | 46180742 | ACATATATGTGTA-TATATATA-TACACATATATAT | loxF8L | 4 | loxF8L | 3 | TATATATA/TATATATA | | 7 |
| chr4 | 37737974 | ACATATATGTGTA-TATATATA-TACACATATATAT | loxF8L | 4 | loxF8L | 3 | TATATATA/TATATATA | | 7 |
| chr9 | 70408989 | ATATATATGTGTA-TATATATA-TACACATATATGT | loxF8L | 3 | loxF8L | 4 | TATATATA/TATATATA | | 7 |
| chrX | 146043099 | ATACATATGTGTA-TATATATA-TACACATATATGT | loxF8L | 3 | loxF8L | 4 | TATATATA/TATATATA | | 7 |
| chr6 | 43289960 | ATATATGTGTGTA-TATATATA-TACACACATATGT | loxF8L | 3 | loxF8L | 4 | TATATATA/TATATATA | | 7 |
| chr4 | 75187308 | AGATATGTGTGTA-TATATATA-TACACACATATAT | loxF8L | 4 | loxF8L | 3 | TATATATA/TATATATA | | 7 |
| chr7 | 52749477 | ATATATATGTGTG-TATATATA-TACACACTTTTAT | loxF8L | 4 | loxF8L | 3 | TATATATA/TATATATA | | 7 |
| chr7 | 44616606 | ATATATATATGTA-TATATATA-TACACATGTTTAT | loxF8L | 4 | loxF8L | 3 | TATATATA/TATATATA | | 7 |
| chrX | 116283761 | ATAAACCTGTATA-TATATATA-CACACAACTATAT | loxF8L | 3 | loxF8R | 4 | TATATATA/TATATATA | | 7 |
| chr6 | 153054107 | ATATATGTGTATA-TATATATA-TATACAGCTTTAT | loxF8L | 4 | loxF8L | 3 | TATATATA/TATATATA | | 7 |
| chr14 | 38066848 | ATATATATGTATA-TATATATA-TCCATTTATTTAT | loxF8L | 4 | loxF8L | 3 | TATATATA/TATATATA | | 7 |
| chrX | 107782364 | ATAAATATATATA-TATATATA-CCCACACATATAT | loxF8L | 4 | loxF8L | 3 | TATATATA/TATATATA | | 7 |
| chr5 | 117316388 | ATAAATATATATA-TATATATA-TACACATATATAT | loxF8L | 4 | loxF8L | 3 | TATATATA/TATATATA | | 7 |
| chr14 | 83998518 | ATAAATATATGTG-TATATATA-TGCACATATATAT | loxF8L | 4 | loxF8L | 3 | TATATATA/TATATATA | | 7 |
| chr14 | 83998986 | ATAAATATATGTG-TATATATA-TGCACATATATAT | loxF8L | 4 | loxF8L | 3 | TATATATA/TATATATA | | 7 |
| chr14 | 27031795 | ATAAATATGTGTG-TATATATA-CACACACACATAT | loxF8L | 3 | loxF8R | 4 | TATATATA/TATATATA | | 7 |
| chr16 | 10043918 | ATAAATATGTGTG-TATATATA-CACACATATATAT | loxF8L | 3 | loxF8L | 4 | TATATATA/TATATATA | | 7 |
| chr5 | 137329351 | ATAAATCTATTTA-TATATATA-TACACCTATATAT | loxF8L | 3 | loxF8L | 4 | TATATATA/TATATATA | | 7 |
| chr4 | 162324514 | ATATATGTGTATA-TATATATA-TAACCAGATTTAT | loxF8L | 4 | loxF8L | 3 | TATATATA/TATATATA | | 7 |
| chr2 | 35576990 | ATAAATCTGTATC-TATATATA-TACACACACATAT | loxF8L | 3 | loxF8L | 4 | TATATATA/TATATATA | | 7 |
| chr4 | 158255541 | ATATATGTGTGTG-TATATATA-TATACACATTTAT | loxF8L | 4 | loxF8L | 3 | TATATATA/TATATATA | | 7 |
| chr10 | 27790243 | GTATATATGTGCA-TATATATA-TAAACACATTTAT | loxF8L | 4 | loxF8L | 3 | TATATATA/TATATATA | | 7 |
| chr16 | 77409029 | ATAAGTATGTGTA-TATATATA-CACACATATATAT | loxF8L | 3 | loxF8L | 4 | TATATATA/TATATATA | | 7 |
| chr4 | 80608082 | ATATATGTGTGTG-TATATATA-CACACATACTTAT | loxF8L | 4 | loxF8R | 3 | TATATATA/TATATATA | | 7 |
| chr6 | 98457319 | CCTAGTGTGTGTG-TATATATA-TGCACACACTTAT | loxF8R | 4 | loxF8L | 3 | TATATATA/TATATATA | | 7 |
| chr5 | 165395445 | ATATATATATGTA-TATATATA-TACACACACTTAT | loxF8L | 4 | loxF8L | 3 | TATATATA/TATATATA | | 7 |
| chrX | 140412930 | ATACATATGTGTA-TATATATA-TACACATTATTAT | loxF8L | 3 | loxF8L | 4 | TATATATA/TATATATA | | 7 |
| chr21 | 34324961 | ATACAAGTGTGTA-TATATATA-TACACATATATAT | loxF8L | 4 | loxF8L | 3 | TATATATA/TATATATA | | 7 |
| chrX | 136686655 | ATACACATGTGTA-TATATATA-TACACATATATAT | loxF8L | 4 | loxF8L | 3 | TATATATA/TATATATA | | 7 |
| chr6 | 67605102 | ATACACGTGTGTA-TATATATA-TACACACATATAT | loxF8L | 4 | loxF8L | 3 | TATATATA/TATATATA | | 7 |
| chr1 | 215230028 | ATACATATATAGA-TATATATA-CACACAGATATAT | loxF8L | 4 | loxF8L | 3 | TATATATA/TATATATA | | 7 |
| chr3 | 108819788 | ATATATATGTGTA-TATATATA-TCTATATATGTAT | loxF8L | 3 | loxF8L | 4 | TATATATA/TATATATA | | 7 |
| chr4 | 119909898 | ATACATATATGGA-TATATATA-CACACATATATAT | loxF8L | 3 | loxF8L | 4 | TATATATA/TATATATA | | 7 |
| chr6 | 152224948 | ATATATATGTGTA-TATATATA-TACATATATGTAT | loxF8L | 3 | loxF8L | 4 | TATATATA/TATATATA | | 7 |
| chr7 | 147585792 | ATATGTGTGTGTG-TATATATA-TCTACATATGTAT | loxF8R | 4 | loxF8L | 3 | TATATATA/TATATATA | | 7 |
| chr13 | 79607628 | ATATATATGTGTA-TATATATA-TATACATATGTAT | loxF8L | 3 | loxF8L | 4 | TATATATA/TATATATA | | 7 |
| chr9 | 119040426 | ATATGTGTGTGTG-TATATATA-TGCACATATGTAT | loxF8R | 4 | loxF8L | 3 | TATATATA/TATATATA | | 7 |
| chr18 | 52528277 | ATACATATGTGCA-TATATATA-TTCCAAGATATAT | loxF8L | 3 | loxF8L | 4 | TATATATA/TATATATA | | 7 |
| chr5 | 116964430 | ATACATATGTGTA-TATATATA-CACACACACATAT | loxF8L | 3 | loxF8R | 4 | TATATATA/TATATATA | | 7 |
| chr9 | 15642276 | ATATGTGTGTGTG-TATATATA-TACACATATGTAT | loxF8R | 4 | loxF8L | 3 | TATATATA/TATATATA | | 7 |
| chr2 | 110491515 | ATACATATGTGTA-TATATATA-TACACACACATAT | loxF8L | 3 | loxF8L | 4 | TATATATA/TATATATA | | 7 |
| chr2 | 109879957 | ATATGTGTGTGTA-TATATATA-TACACATATGTAT | loxF8L | 4 | loxF8L | 3 | TATATATA/TATATATA | | 7 |
| chrX | 140413021 | ATACATATGTGTA-TATATATA-TACACATATATAC | loxF8L | 3 | loxF8L | 4 | TATATATA/TATATATA | | 7 |
| chr12 | 44882328 | ATATATGTATGTA-TATATATA-TACACATATGTAT | loxF8L | 4 | loxF8L | 3 | TATATATA/TATATATA | | 7 |
| chr8 | 1579320 | ATATATGTGTGTA-TATATATA-GACACATATGTAT | loxF8L | 3 | loxF8L | 4 | TATATATA/TATATATA | | 7 |
| chr15 | 37242963 | ATACATTTGTGTA-TATATATA-CACACATATGTAT | loxF8L | 3 | loxF8L | 4 | TATATATA/TATATATA | | 7 |
| chr3 | 189057807 | ATATATGTGTGTA-TATATATA-CACACATATGTAT | loxF8L | 3 | loxF8L | 4 | TATATATA/TATATATA | | 7 |
| chr16 | 65011006 | ATATATATGTGTA-TATATATA-CACACATATGTAT | loxF8L | 3 | loxF8L | 4 | TATATATA/TATATATA | | 7 |
| chr21 | 29792177 | GTATATGTGTGGA-TATATATA-CACACATATGTAT | loxF8L | 3 | loxF8L | 4 | TATATATA/TATATATA | | 7 |
| chr3 | 103079449 | ATACATCTATGTG-TATATATA-TACACACATATAT | loxF8L | 4 | loxF8L | 3 | TATATATA/TATATATA | | 7 |
| chr13 | 58553809 | ATATAAGTGTGTG-TATATATA-CACACAGATGTAT | loxF8R | 4 | loxF8L | 3 | TATATATA/TATATATA | | 7 |
| chr2 | 175896841 | ATACATGTGTATA-TATATATA-TACACATATATAT | loxF8L | 4 | loxF8L | 3 | TATATATA/TATATATA | | 7 |
| chrX | 15953635 | ATACATGTGTGAA-TATATATA-TCTATATATATAT | loxF8L | 3 | loxF8L | 4 | TATATATA/TATATATA | | 7 |
| chr5 | 57278525 | ATACATGTGTGTA-TATATATA-TACACACACATAT | loxF8L | 3 | loxF8L | 4 | TATATATA/TATATATA | | 7 |
| chr5 | 162179187 | ATACATGTGTGTA-TATATATA-TACACGTATATAT | loxF8L | 3 | loxF8L | 4 | TATATATA/TATATATA | | 7 |
| chrX | 7736952 | ATACATTTGTGTA-TATATATA-TACACACTCATAT | loxF8L | 3 | loxF8R | 4 | TATATATA/TATATATA | | 7 |
| chr6 | 56636389 | ATACGTATGTGTG-TATATATA-TACACACATATAT | loxF8R | 4 | loxF8L | 3 | TATATATA/TATATATA | | 7 |
| chr3 | 113935533 | ATACGTGTGTGTA-TATATATA-TACACATATATAT | loxF8L | 4 | loxF8L | 3 | TATATATA/TATATATA | | 7 |
| chr1 | 182918119 | ATACGTGTGTGTG-TATATATA-TACACATATATAT | loxF8R | 4 | loxF8L | 3 | TATATATA/TATATATA | | 7 |

| | | | | | | | | |
|---|---|---|---|---|---|---|---|---|
| chr9 | 80757831 | ATATATATGTGTA-TATATATA-CACACACACGTAT | loxF8L | 3 | loxF8R | 4 | TATATATA/TATATATA | 7 |
| chr4 | 137135338 | ATAGAACTGTGTA-TATATATA-CACACACACACAG | loxF8L | 3 | loxF8R | 4 | TATATATA/TATATATA | 7 |
| chr2 | 44442297 | ATAGAGTTGTGTG-TATATATA-TACACACATATAT | loxF8R | 4 | loxF8L | 3 | TATATATA/TATATATA | 7 |
| chr3 | 8046418 | ATATATCTATCTA-TATATATA-TCTACATATCTAT | loxF8L | 4 | loxF8L | 3 | TATATATA/TATATATA | 7 |
| chr2 | 58542348 | ATATATATGTGTA-TATATATA-TACACATGTATAT | loxF8L | 3 | loxF8L | 4 | TATATATA/TATATATA | 7 |
| chr15 | 35463775 | ATATACCTATGGA-TATATATA-TATACACATTCAT | loxF8L | 3 | loxF8L | 4 | TATATATA/TATATATA | 7 |
| chr7 | 85279775 | ATATATACGTGCA-TATATATA-CACACACTCATAT | loxF8L | 4 | loxF8R | 3 | TATATATA/TATATATA | 7 |
| chr13 | 90621174 | ATATATACGTGTA-TATATATA-TACACACATATAT | loxF8L | 4 | loxF8L | 3 | TATATATA/TATATATA | 7 |
| chrX | 56058209 | ATATATACGTGTA-TATATATA-TACACACATATAT | loxF8L | 4 | loxF8L | 3 | TATATATA/TATATATA | 7 |
| chr7 | 90461790 | ATATATATGTGTA-TATATATA-TTCATATATATAT | loxF8L | 3 | loxF8L | 4 | TATATATA/TATATATA | 7 |
| chr13 | 67613489 | ATATATATGTGTA-TATATATA-TGCATATATATAT | loxF8L | 3 | loxF8L | 4 | TATATATA/TATATATA | 7 |
| chr8 | 72561766 | ATATATATATGGA-TATATATA-TACATATATATAT | loxF8L | 3 | loxF8L | 4 | TATATATA/TATATATA | 7 |
| chr8 | 113934419 | ATATATATATGGA-TATATATA-TATACACATATAT | loxF8L | 3 | loxF8L | 4 | TATATATA/TATATATA | 7 |
| chrX | 84191523 | ATATATGTGTATA-TATATATA-TCCATATATATAT | loxF8L | 4 | loxF8L | 3 | TATATATA/TATATATA | 7 |
| chrX | 77097298 | ATATATATGTATA-TATATATA-TCCATATATATAT | loxF8L | 4 | loxF8L | 3 | TATATATA/TATATATA | 7 |
| chr6 | 104387492 | ATATATATATGGA-TATATATA-TCCATATATATAA | loxF8L | 3 | loxF8L | 4 | TATATATA/TATATATA | 7 |
| chr14 | 47540543 | ATATATATATGTA-TATATATA-CACACACACATAG | loxF8L | 4 | loxF8R | 3 | TATATATA/TATATATA | 7 |
| chr11 | 111181286 | ATATATGTGTGTA-TATATATA-TACATATATATAT | loxF8L | 3 | loxF8L | 4 | TATATATA/TATATATA | 7 |
| chr21 | 17069163 | ATATATGTGTGTA-TATATATA-TACATATATATAT | loxF8L | 3 | loxF8L | 4 | TATATATA/TATATATA | 7 |
| chr4 | 166346410 | ATATATGTGTGTA-TATATATA-TACATATATATAT | loxF8L | 3 | loxF8L | 4 | TATATATA/TATATATA | 7 |
| chr4 | 166346466 | ATATATGTGTGTA-TATATATA-TACATATATATAT | loxF8L | 3 | loxF8L | 4 | TATATATA/TATATATA | 7 |
| chr4 | 166346438 | ATATATGTGTGTA-TATATATA-TACATATATATAT | loxF8L | 3 | loxF8L | 4 | TATATATA/TATATATA | 7 |
| chr5 | 78315954 | ATATATGTGTGTA-TATATATA-TACATATATATAT | loxF8L | 3 | loxF8L | 4 | TATATATA/TATATATA | 7 |
| chr8 | 34389804 | ATATATGTGTGTA-TATATATA-TACATATATATAT | loxF8L | 3 | loxF8L | 4 | TATATATA/TATATATA | 7 |
| chr8 | 128363242 | ATATATGTGTGTA-TATATATA-TACATATATATAT | loxF8L | 3 | loxF8L | 4 | TATATATA/TATATATA | 7 |
| chr8 | 128363354 | ATATATGTGTGTA-TATATATA-TACATATATATAT | loxF8L | 3 | loxF8L | 4 | TATATATA/TATATATA | 7 |
| chr8 | 128363270 | ATATATGTGTGTA-TATATATA-TACATATATATAT | loxF8L | 3 | loxF8L | 4 | TATATATA/TATATATA | 7 |
| chr8 | 128363298 | ATATATGTGTGTA-TATATATA-TACATATATATAT | loxF8L | 3 | loxF8L | 4 | TATATATA/TATATATA | 7 |
| chr8 | 128363326 | ATATATGTGTGTA-TATATATA-TACATATATATAT | loxF8L | 3 | loxF8L | 4 | TATATATA/TATATATA | 7 |
| chr14 | 27299180 | ATATATATGTGTA-TATATATA-TACATATATATAT | loxF8L | 3 | loxF8L | 4 | TATATATA/TATATATA | 7 |
| chr15 | 88915923 | ATATATATGTGTA-TATATATA-TACATATATATAT | loxF8L | 3 | loxF8L | 4 | TATATATA/TATATATA | 7 |
| chr15 | 88915887 | ATATATATGTGTA-TATATATA-TACATATATATAT | loxF8L | 3 | loxF8L | 4 | TATATATA/TATATATA | 7 |
| chr16 | 5207559 | ATATATATATGTA-TATATATA-TACACATATATAT | loxF8L | 4 | loxF8L | 3 | TATATATA/TATATATA | 7 |
| chr20 | 14702626 | ATATATATATGTA-TATATATA-TACACATATATAT | loxF8L | 4 | loxF8L | 3 | TATATATA/TATATATA | 7 |
| chr20 | 14702602 | ATATATATGTGTA-TATATATA-TACATATATATAT | loxF8L | 3 | loxF8L | 4 | TATATATA/TATATATA | 7 |
| chr21 | 23402118 | ATATATATGTGTA-TATATATA-TACATATATATAT | loxF8L | 3 | loxF8L | 4 | TATATATA/TATATATA | 7 |
| chr22 | 16523170 | ATATATATGTGTA-TATATATA-TACATATATATAT | loxF8L | 3 | loxF8L | 4 | TATATATA/TATATATA | 7 |
| chr2 | 136433086 | ATATATATGTGTA-TATATATA-TACATATATATAT | loxF8L | 3 | loxF8L | 4 | TATATATA/TATATATA | 7 |
| chr2 | 145354179 | ATATATATATGTA-TATATATA-TACACATATATAT | loxF8L | 4 | loxF8L | 3 | TATATATA/TATATATA | 7 |
| chr3 | 114303494 | ATATATATATGTA-TATATATA-TACACATATATAT | loxF8L | 4 | loxF8L | 3 | TATATATA/TATATATA | 7 |
| chr3 | 153301304 | ATATATATGTGTA-TATATATA-TACATATATATAT | loxF8L | 3 | loxF8L | 4 | TATATATA/TATATATA | 7 |
| chr4 | 105284099 | ATATATATATGTA-TATATATA-TACACATATATAT | loxF8L | 4 | loxF8L | 3 | TATATATA/TATATATA | 7 |
| chr4 | 111574844 | ATATATATGTGTA-TATATATA-TACATATATATAT | loxF8L | 3 | loxF8L | 4 | TATATATA/TATATATA | 7 |
| chr5 | 85600254 | ATATATATGTGTA-TATATATA-TACATATATATAT | loxF8L | 3 | loxF8L | 4 | TATATATA/TATATATA | 7 |
| chr6 | 121679782 | ATATATATATGTA-TATATATA-TACACATATATAT | loxF8L | 4 | loxF8L | 3 | TATATATA/TATATATA | 7 |
| chr6 | 115044479 | ATATATATGTGTA-TATATATA-TACATATATATAT | loxF8L | 3 | loxF8L | 4 | TATATATA/TATATATA | 7 |
| chr9 | 12223218 | ATATATATGTGTA-TATATATA-TACATATATATAT | loxF8L | 3 | loxF8L | 4 | TATATATA/TATATATA | 7 |
| chrX | 8594727 | ATATATATGTGTA-TATATATA-TACATATATATAT | loxF8L | 3 | loxF8L | 4 | TATATATA/TATATATA | 7 |
| chr21 | 20586762 | ATATATATATGTA-TATATATA-TACCCAGATATAT | loxF8L | 4 | loxF8L | 3 | TATATATA/TATATATA | 7 |
| chr8 | 68158062 | ATATATATATGTA-TATATATA-TTCACATATATAT | loxF8L | 4 | loxF8L | 3 | TATATATA/TATATATA | 7 |
| chrX | 147509455 | ATATATATGTGCA-TATATATA-TGCGCATATATAT | loxF8L | 3 | loxF8L | 4 | TATATATA/TATATATA | 7 |
| chr14 | 41943955 | ATATATGTGTGTA-TATATATA-TACGCATATATAT | loxF8L | 3 | loxF8L | 4 | TATATATA/TATATATA | 7 |
| chr3 | 190881825 | ATATATATGCGTA-TATATATA-TACACACATATAT | loxF8L | 4 | loxF8L | 3 | TATATATA/TATATATA | 7 |
| chr11 | 32755710 | ATATATATGTGTA-TATATATA-TCTCCATATATAT | loxF8L | 3 | loxF8L | 4 | TATATATA/TATATATA | 7 |
| chr11 | 32755684 | ATATATATGTGTA-TATATATA-TCTCCATATATAT | loxF8L | 3 | loxF8L | 4 | TATATATA/TATATATA | 7 |
| chr11 | 32755632 | ATATATATGTGTA-TATATATA-TCTCCATATATAT | loxF8L | 3 | loxF8L | 4 | TATATATA/TATATATA | 7 |
| chr11 | 32755658 | ATATATATGTGTA-TATATATA-TCTCCATATATAT | loxF8L | 3 | loxF8L | 4 | TATATATA/TATATATA | 7 |
| chr10 | 124657151 | ATATATGTGTGTA-TATATATA-TGTACATATATAT | loxF8L | 3 | loxF8L | 4 | TATATATA/TATATATA | 7 |
| chr12 | 39416676 | ATATATGTGTGTA-TATATATA-TGTACATATATAT | loxF8L | 3 | loxF8L | 4 | TATATATA/TATATATA | 7 |
| chr3 | 21295942 | ATATATGTGTGTA-TATATATA-TGTACATATATAT | loxF8L | 3 | loxF8L | 4 | TATATATA/TATATATA | 7 |
| chr12 | 39416648 | ATATATATGTGTA-TATATATA-TGTACATATATAT | loxF8L | 3 | loxF8L | 4 | TATATATA/TATATATA | 7 |
| chr12 | 39416726 | ATATATATGTGTA-TATATATA-TGTACATATATAT | loxF8L | 3 | loxF8L | 4 | TATATATA/TATATATA | 7 |
| chr12 | 39416752 | ATATATATGTGTA-TATATATA-TGTACATATATAT | loxF8L | 3 | loxF8L | 4 | TATATATA/TATATATA | 7 |
| chr2 | 49779949 | ATATATGTATAGA-TATATATA-TCTACATATATAT | loxF8L | 4 | loxF8L | 3 | TATATATA/TATATATA | 7 |
| chr4 | 184372704 | CTAATGGTGTGTG-TATATATA-TATACATATATAT | loxF8R | 3 | loxF8L | 4 | TATATATA/TATATATA | 7 |
| chr12 | 88655262 | ATATATATGTATA-TATATATA-CACACACCCTAAG | loxF8L | 4 | loxF8R | 3 | TATATATA/TATATATA | 7 |
| chr12 | 86885932 | GTAACAGTGTGTG-TATATATA-TATACATATATAT | loxF8R | 3 | loxF8L | 4 | TATATATA/TATATATA | 7 |
| chrX | 83445880 | ATATATATGTATA-TATATATA-TACACAAATATAT | loxF8L | 4 | loxF8L | 3 | TATATATA/TATATATA | 7 |
| chr10 | 76240835 | ATATATGTGTGTA-TATATATA-TATACATATATAT | loxF8L | 3 | loxF8L | 4 | TATATATA/TATATATA | 7 |
| chr14 | 98969305 | ATATATATGTATA-TATATATA-TACACACATATAT | loxF8L | 4 | loxF8L | 3 | TATATATA/TATATATA | 7 |
| chr15 | 21642164 | ATATATATGTATA-TATATATA-TACACACATATAT | loxF8L | 4 | loxF8L | 3 | TATATATA/TATATATA | 7 |
| chr15 | 95749098 | ATATATGTGTGTA-TATATATA-TATACATATATAT | loxF8L | 3 | loxF8L | 4 | TATATATA/TATATATA | 7 |
| chr2 | 30796133 | ATATATGTGTGTA-TATATATA-TATACATATATAT | loxF8L | 3 | loxF8L | 4 | TATATATA/TATATATA | 7 |
| chr2 | 235284279 | ATATATATGTATA-TATATATA-TACACACATATAT | loxF8L | 4 | loxF8L | 3 | TATATATA/TATATATA | 7 |
| chr3 | 174373922 | ATATATATGTATA-TATATATA-TACACACATATAT | loxF8L | 4 | loxF8L | 3 | TATATATA/TATATATA | 7 |
| chr7 | 49622002 | ATATATATGTATA-TATATATA-TACACACATATAT | loxF8L | 4 | loxF8L | 3 | TATATATA/TATATATA | 7 |
| chr11 | 124586249 | ATATATATGTGTA-TATATATA-TATACATATATAT | loxF8L | 3 | loxF8L | 4 | TATATATA/TATATATA | 7 |
| chr12 | 60908570 | ATATATATGTGTA-TATATATA-TATACATATATAT | loxF8L | 3 | loxF8L | 4 | TATATATA/TATATATA | 7 |
| chr12 | 60908516 | ATATATATGTGTA-TATATATA-TATACATATATAT | loxF8L | 3 | loxF8L | 4 | TATATATA/TATATATA | 7 |
| chr12 | 60908544 | ATATATATGTGTA-TATATATA-TATACATATATAT | loxF8L | 3 | loxF8L | 4 | TATATATA/TATATATA | 7 |
| chr12 | 81462559 | ATATATATGTGTA-TATATATA-TATACATATATAT | loxF8L | 3 | loxF8L | 4 | TATATATA/TATATATA | 7 |
| chr13 | 58617725 | ATATATATGTGTA-TATATATA-TATACATATATAT | loxF8L | 3 | loxF8L | 4 | TATATATA/TATATATA | 7 |
| chr14 | 32510451 | ATATATATGTGTA-TATATATA-TATACATATATAT | loxF8L | 3 | loxF8L | 4 | TATATATA/TATATATA | 7 |
| chr14 | 82600618 | ATATATATGTGTA-TATATATA-TATACATATATAT | loxF8L | 3 | loxF8L | 4 | TATATATA/TATATATA | 7 |
| chr19 | 38974007 | ATATATATGTGTA-TATATATA-TATACATATATAT | loxF8L | 3 | loxF8L | 4 | TATATATA/TATATATA | 7 |
| chr1 | 201901587 | ATATATATGTATA-TATATATA-TACACATATATAT | loxF8L | 4 | loxF8L | 3 | TATATATA/TATATATA | 7 |
| chr21 | 24633406 | ATATATATGTGTA-TATATATA-TATACATATATAT | loxF8L | 3 | loxF8L | 4 | TATATATA/TATATATA | 7 |

| | | | | | | | | | |
|---|---|---|---|---|---|---|---|---|---|
| chr3 | 71397075 | ATATATATGTGTA-TATATATA-TATACATATATAT | loxF8L | 3 | loxF8L | 4 | TATATATA/TATATATA | | 7 |
| chr3 | 153301266 | ATATATATGTGTA-TATATATA-TATACATATATAT | loxF8L | 3 | loxF8L | 4 | TATATATA/TATATATA | | 7 |
| chr4 | 47976314 | ATATATATGTATA-TATATATA-TACATATATATAT | loxF8L | 4 | loxF8L | 3 | TATATATA/TATATATA | | 7 |
| chr4 | 111574870 | ATATATATGTGTA-TATATATA-TATACATATATAT | loxF8L | 3 | loxF8L | 4 | TATATATA/TATATATA | | 7 |
| chr6 | 13003956 | ATATATATGTATA-TATATATA-TACATATATATAT | loxF8L | 4 | loxF8L | 3 | TATATATA/TATATATA | | 7 |
| chr7 | 71701872 | ATATATATGTATA-TATATATA-TACATATATATAT | loxF8L | 4 | loxF8L | 3 | TATATATA/TATATATA | | 7 |
| chr7 | 147083541 | ATATATATGTATA-TATATATA-TACATATATATAT | loxF8L | 4 | loxF8L | 3 | TATATATA/TATATATA | | 7 |
| chr9 | 42735178 | ATATATATGTATA-TATATATA-TACATATATATAT | loxF8L | 4 | loxF8L | 3 | TATATATA/TATATATA | | 7 |
| chr9 | 111219373 | ATATATATGTGTA-TATATATA-TATACATATATAT | loxF8L | 3 | loxF8L | 4 | TATATATA/TATATATA | | 7 |
| chrX | 13191844 | ATATATATGTGTA-TATATATA-TATACATATATAT | loxF8L | 3 | loxF8L | 4 | TATATATA/TATATATA | | 7 |
| chrX | 75272321 | ATATATATGTATA-TATATATA-TACATATATATAT | loxF8L | 4 | loxF8L | 3 | TATATATA/TATATATA | | 7 |
| chr12 | 16093095 | GTGAATATGTGAA-TATATATA-TTCACATATATAT | loxF8L | 4 | loxF8L | 3 | TATATATA/TATATATA | | 7 |
| chr5 | 82108373 | ATATATATGTGCA-TATATATA-CACACAGTATATG | loxF8L | 3 | loxF8R | 4 | TATATATA/TATATATA | | 7 |
| chr14 | 95778045 | TGCATATTGTGTG-TATATATA-TACACATATATAT | loxF8R | 4 | loxF8L | 3 | TATATATA/TATATATA | | 7 |
| chr10 | 31242006 | ATATATATGTGTA-TATATATA-CACACACACATAC | loxF8L | 3 | loxF8R | 4 | TATATATA/TATATATA | | 7 |
| chr2 | 193556185 | GTATGTGTGTGTG-TATATATA-TACACATATATAT | loxF8R | 4 | loxF8L | 3 | TATATATA/TATATATA | | 7 |
| chr6 | 76311798 | ATATATATGTGTA-TATATATA-CACACACACATAC | loxF8L | 3 | loxF8R | 4 | TATATATA/TATATATA | | 7 |
| chr8 | 129064170 | ATATATATGTGTA-TATATATA-CACACACACATAC | loxF8L | 3 | loxF8R | 4 | TATATATA/TATATATA | | 7 |
| chr10 | 60912204 | ATATATATGTGTA-TATATATA-CACACACACATAT | loxF8L | 3 | loxF8R | 4 | TATATATA/TATATATA | | 7 |
| chr12 | 129147161 | ATATGTGTGTGTG-TATATATA-TACACATATATAT | loxF8R | 4 | loxF8L | 3 | TATATATA/TATATATA | | 7 |
| chr1 | 54957268 | ATATGTGTGTGTG-TATATATA-TACACATATATAT | loxF8R | 4 | loxF8L | 3 | TATATATA/TATATATA | | 7 |
| chr3 | 29007263 | ATATGTGTGTGTG-TATATATA-TACACATATATAT | loxF8R | 4 | loxF8L | 3 | TATATATA/TATATATA | | 7 |
| chr3 | 9881783 | ATATATATGTGTA-TATATATA-CACACACACATAT | loxF8L | 3 | loxF8R | 4 | TATATATA/TATATATA | | 7 |
| chr7 | 102338140 | ATATGTGTGTGTG-TATATATA-TACACATATATAT | loxF8R | 4 | loxF8L | 3 | TATATATA/TATATATA | | 7 |
| chr9 | 25518072 | ATATATATGTGTA-TATATATA-CACACACACATAT | loxF8L | 3 | loxF8R | 4 | TATATATA/TATATATA | | 7 |
| chr9 | 80757751 | ATATATATGTGTA-TATATATA-CACACACACATAT | loxF8L | 3 | loxF8R | 4 | TATATATA/TATATATA | | 7 |
| chr15 | 94336734 | ATATATGTGTGTG-TATATATA-TACACATATATAT | loxF8L | 4 | loxF8L | 3 | TATATATA/TATATATA | | 7 |
| chr16 | 46898919 | ATATATATGTGTA-TATATATA-CACACACATATAT | loxF8L | 3 | loxF8L | 4 | TATATATA/TATATATA | | 7 |
| chr16 | 50116404 | ATATATGTGTGTG-TATATATA-TACACATATATAT | loxF8L | 4 | loxF8L | 3 | TATATATA/TATATATA | | 7 |
| chr18 | 5308570 | ATATATATGTGTA-TATATATA-CACACACATATAT | loxF8L | 3 | loxF8L | 4 | TATATATA/TATATATA | | 7 |
| chr18 | 51781560 | ATATATGTGTGTG-TATATATA-TACACATATATAT | loxF8L | 4 | loxF8L | 3 | TATATATA/TATATATA | | 7 |
| chr1 | 146481510 | ATATATATGTGTA-TATATATA-CACACACATATAT | loxF8L | 3 | loxF8L | 4 | TATATATA/TATATATA | | 7 |
| chr2 | 87915593 | ATATATGTGTGTG-TATATATA-TACACATATATAT | loxF8L | 4 | loxF8L | 3 | TATATATA/TATATATA | | 7 |
| chr2 | 177631642 | ATATATGTGTGTG-TATATATA-TACACATATATAT | loxF8L | 4 | loxF8L | 3 | TATATATA/TATATATA | | 7 |
| chr5 | 96787791 | ATATATGTGTGTG-TATATATA-TACACATATATAT | loxF8L | 4 | loxF8L | 3 | TATATATA/TATATATA | | 7 |
| chr6 | 123793941 | ATATATGTGTGTG-TATATATA-TACACATATATAT | loxF8L | 4 | loxF8L | 3 | TATATATA/TATATATA | | 7 |
| chr7 | 102338110 | ATATATGTGTGTG-TATATATA-TACACATATATAT | loxF8L | 4 | loxF8L | 3 | TATATATA/TATATATA | | 7 |
| chrX | 144123504 | ATATATATGTGTA-TATATATA-CACACACATATAT | loxF8L | 3 | loxF8L | 4 | TATATATA/TATATATA | | 7 |
| chr5 | 22819936 | GTATGTATGTGTG-TATATATA-TACACATATATAT | loxF8R | 4 | loxF8L | 3 | TATATATA/TATATATA | | 7 |
| chr9 | 21861312 | ATATATATGTGTA-TATATATA-CACACATACATAC | loxF8L | 3 | loxF8R | 4 | TATATATA/TATATATA | | 7 |
| chr1 | 71943007 | ATATATATGTGTA-TATATATA-CACACATACATAT | loxF8L | 3 | loxF8R | 4 | TATATATA/TATATATA | HG6 | 7 |
| chr1 | 71943065 | ATATATATGTGTA-TATATATA-CACACATACATAT | loxF8L | 3 | loxF8R | 4 | TATATATA/TATATATA | HG6 | 7 |
| chr12 | 72430446 | ATATATATGTGTA-TATATATA-CACACATATATAT | loxF8L | 3 | loxF8L | 4 | TATATATA/TATATATA | | 7 |
| chr13 | 56465829 | ATATATATGTGTA-TATATATA-CACACATATATAT | loxF8L | 3 | loxF8L | 4 | TATATATA/TATATATA | | 7 |
| chr15 | 101894988 | ATATATATGTGTG-TATATATA-TACACATATATAT | loxF8L | 4 | loxF8L | 3 | TATATATA/TATATATA | | 7 |
| chr1 | 667982 | ATATATATGTGTA-TATATATA-CACACATATATAT | loxF8L | 3 | loxF8L | 4 | TATATATA/TATATATA | | 7 |
| chr1 | 432230 | ATATATATGTGTA-TATATATA-CACACATATATAT | loxF8L | 3 | loxF8L | 4 | TATATATA/TATATATA | | 7 |
| chr1 | 54957378 | ATATATATGTGTG-TATATATA-TACACATATATAT | loxF8L | 4 | loxF8L | 3 | TATATATA/TATATATA | | 7 |
| chr1 | 238056300 | ATATATATGTGTG-TATATATA-TACACATATATAT | loxF8L | 4 | loxF8L | 3 | TATATATA/TATATATA | | 7 |
| chr5 | 181386623 | ATATATATGTGTG-TATATATA-TACACATATATAT | loxF8L | 4 | loxF8L | 3 | TATATATA/TATATATA | | 7 |
| chr6 | 88188 | ATATATATGTGTA-TATATATA-CACACATATATAT | loxF8L | 3 | loxF8L | 4 | TATATATA/TATATATA | | 7 |
| chr6 | 89099417 | ATATATATGTGTG-TATATATA-TACACATATATAT | loxF8L | 4 | loxF8L | 3 | TATATATA/TATATATA | | 7 |
| chr6 | 64686890 | ATATATATGTGTA-TATATATA-CACACATATATAT | loxF8L | 3 | loxF8L | 4 | TATATATA/TATATATA | | 7 |
| chr6 | 170662151 | ATATATATGTGTG-TATATATA-TACACATATATAT | loxF8L | 4 | loxF8L | 3 | TATATATA/TATATATA | | 7 |
| chr8 | 145931 | ATATATATGTGTA-TATATATA-CACACATATATAT | loxF8L | 3 | loxF8L | 4 | TATATATA/TATATATA | | 7 |
| chr8 | 66718411 | ATATATATGTGTA-TATATATA-CACACATATATAT | loxF8L | 3 | loxF8L | 4 | TATATATA/TATATATA | | 7 |
| chrX | 95270759 | ATATATATGTGTG-TATATATA-TACACATATATAT | loxF8L | 4 | loxF8L | 3 | TATATATA/TATATATA | | 7 |
| chr16 | 8612727 | ATATATATGTGTA-TATATATA-CACATCATCATAT | loxF8L | 3 | loxF8R | 4 | TATATATA/TATATATA | | 7 |
| chr2 | 166475069 | TTATATTTGTGTA-TATATATA-TACACATATATAT | loxF8L | 4 | loxF8L | 3 | TATATATA/TATATATA | | 7 |
| chr11 | 37404214 | ATATATATGTGTA-TATATATA-TACACACACATAT | loxF8L | 3 | loxF8L | 4 | TATATATA/TATATATA | | 7 |
| chr15 | 80655796 | ATATGTGTGTGTA-TATATATA-TACACATATATAT | loxF8L | 4 | loxF8L | 3 | TATATATA/TATATATA | | 7 |
| chr18 | 5308598 | ATATATATGTGTA-TATATATA-TACACACACATAT | loxF8L | 3 | loxF8L | 4 | TATATATA/TATATATA | | 7 |
| chr18 | 5308540 | ATATATATGTGTA-TATATATA-TACACACACATAT | loxF8L | 3 | loxF8L | 4 | TATATATA/TATATATA | | 7 |
| chr1 | 30926932 | ATATATATGTGTA-TATATATA-TACACACACATAT | loxF8L | 3 | loxF8L | 4 | TATATATA/TATATATA | | 7 |
| chr1 | 49949987 | ATATATATGTGTA-TATATATA-TACACACACATAT | loxF8L | 3 | loxF8L | 4 | TATATATA/TATATATA | | 7 |
| chr2 | 30796189 | ATATGTGTGTGTA-TATATATA-TACACATATATAT | loxF8L | 4 | loxF8L | 3 | TATATATA/TATATATA | | 7 |
| chr4 | 71887089 | ATATATATGTGTA-TATATATA-TACACACACATAT | loxF8L | 3 | loxF8L | 4 | TATATATA/TATATATA | | 7 |
| chr4 | 71887044 | ATATATATGTGTA-TATATATA-TACACACACATAT | loxF8L | 3 | loxF8L | 4 | TATATATA/TATATATA | | 7 |
| chr5 | 81327160 | ATATATATGTGTA-TATATATA-TACACACACATAT | loxF8L | 3 | loxF8L | 4 | TATATATA/TATATATA | | 7 |
| chr6 | 45962175 | ATATGTGTGTGTA-TATATATA-TACACATATATAT | loxF8L | 4 | loxF8L | 3 | TATATATA/TATATATA | | 7 |
| chr9 | 111219405 | ATATGTGTGTGTA-TATATATA-TACACATATATAT | loxF8L | 4 | loxF8L | 3 | TATATATA/TATATATA | | 7 |
| chrX | 49181606 | ATATATATGTGTA-TATATATA-TACACACACATAT | loxF8L | 3 | loxF8L | 4 | TATATATA/TATATATA | | 7 |
| chrX | 143668066 | ATATATATGTGTA-TATATATA-TACACACACATAT | loxF8L | 3 | loxF8L | 4 | TATATATA/TATATATA | | 7 |
| chr13 | 90570738 | ATGTATGTGTGTA-TATATATA-TACACATATATAT | loxF8L | 4 | loxF8L | 3 | TATATATA/TATATATA | | 7 |
| chr3 | 155321983 | ATATATATGTGTA-TATATATA-TACACATACATAT | loxF8L | 3 | loxF8L | 4 | TATATATA/TATATATA | | 7 |
| chrX | 146392057 | ATATATATGTGTA-TATATATA-TACACATACATAT | loxF8L | 3 | loxF8L | 4 | TATATATA/TATATATA | | 7 |
| chrX | 146392027 | ATATATATGTGTA-TATATATA-TACACATACATAT | loxF8L | 3 | loxF8L | 4 | TATATATA/TATATATA | | 7 |
| chr13 | 103929517 | TTATATATGTGTA-TATATATA-TACACATATATAT | loxF8L | 4 | loxF8L | 3 | TATATATA/TATATATA | | 7 |
| chr11 | 68084401 | GTATATATGTGTA-TATATATA-TACACATATATAT | loxF8L | 4 | loxF8L | 3 | TATATATA/TATATATA | | 7 |
| chr11 | 68084337 | ATATATATGTGTA-TATATATA-TACACATATATAC | loxF8L | 3 | loxF8L | 4 | TATATATA/TATATATA | | 7 |
| chr5 | 108772309 | ATATATGTATGTA-TATATATA-TACACATATATAT | loxF8L | 4 | loxF8L | 3 | TATATATA/TATATATA | | 7 |
| chr11 | 124586281 | ATATATATGTGTA-TATATATA-TATACACATATAT | loxF8L | 3 | loxF8L | 4 | TATATATA/TATATATA | | 7 |
| chr11 | 124586283 | ATATATGTGTATA-TATATATA-TACACATATATAT | loxF8L | 4 | loxF8L | 3 | TATATATA/TATATATA | | 7 |
| chr13 | 89273953 | ATATATATGTGTA-TATATATA-TATACACATATAT | loxF8L | 3 | loxF8L | 4 | TATATATA/TATATATA | | 7 |
| chr13 | 89273955 | ATATATGTGTATA-TATATATA-TACACATATATAT | loxF8L | 4 | loxF8L | 3 | TATATATA/TATATATA | | 7 |
| chr18 | 68280094 | ATATATATGTGTA-TATATATA-TATACACATATAT | loxF8L | 3 | loxF8L | 4 | TATATATA/TATATATA | | 7 |
| chr18 | 68280096 | ATATATGTGTATA-TATATATA-TACACATATATAT | loxF8L | 4 | loxF8L | 3 | TATATATA/TATATATA | | 7 |

| | | | | | | | | | |
|---|---|---|---|---|---|---|---|---|---|
| chr18 | 68280058 | ATATATGTGTATA-TATATATA-TACACATATATAT | loxF8L | 4 | loxF8L | 3 | TATATATA/TATATATA | | 7 |
| chr18 | 68280132 | ATATATGTGTATA-TATATATA-TACACATATATAT | loxF8L | 4 | loxF8L | 3 | TATATATA/TATATATA | | 7 |
| chr18 | 68280056 | ATATATATGTGTA-TATATATA-TATACACATATAT | loxF8L | 3 | loxF8L | 4 | TATATATA/TATATATA | | 7 |
| chr1 | 201901649 | ATATATATGTGTA-TATATATA-TATACACATATAT | loxF8L | 3 | loxF8L | 4 | TATATATA/TATATATA | | 7 |
| chr1 | 201901651 | ATATATGTGTATA-TATATATA-TACACATATATAT | loxF8L | 4 | loxF8L | 3 | TATATATA/TATATATA | | 7 |
| chr2 | 117278321 | ATATATATGTGTA-TATATATA-TATACACATATAT | loxF8L | 3 | loxF8L | 4 | TATATATA/TATATATA | | 7 |
| chr2 | 117278323 | ATATATGTGTATA-TATATATA-TACACATATATAT | loxF8L | 4 | loxF8L | 3 | TATATATA/TATATATA | | 7 |
| chr2 | 141314819 | ATATATATGTGTA-TATATATA-TATACACATATAT | loxF8L | 3 | loxF8L | 4 | TATATATA/TATATATA | | 7 |
| chr2 | 141314821 | ATATATGTGTATA-TATATATA-TACACATATATAT | loxF8L | 4 | loxF8L | 3 | TATATATA/TATATATA | | 7 |
| chr3 | 153301340 | ATATATATGTGTA-TATATATA-TATACACATATAT | loxF8L | 3 | loxF8L | 4 | TATATATA/TATATATA | | 7 |
| chr3 | 153301342 | ATATATGTGTATA-TATATATA-TACACATATATAT | loxF8L | 4 | loxF8L | 3 | TATATATA/TATATATA | | 7 |
| chr3 | 153301368 | ATATATATGTGTA-TATATATA-TATACACATATAT | loxF8L | 3 | loxF8L | 4 | TATATATA/TATATATA | | 7 |
| chr3 | 153301370 | ATATATGTGTATA-TATATATA-TACACATATATAT | loxF8L | 4 | loxF8L | 3 | TATATATA/TATATATA | | 7 |
| chr8 | 89455386 | ATATATATGTGTA-TATATATA-TATACACATATAT | loxF8L | 3 | loxF8L | 4 | TATATATA/TATATATA | | 7 |
| chr8 | 89455388 | ATATATGTGTATA-TATATATA-TACACATATATAT | loxF8L | 4 | loxF8L | 3 | TATATATA/TATATATA | | 7 |
| chrX | 112257878 | ATATATATGTGTA-TATATATA-TATACACATATAT | loxF8L | 3 | loxF8L | 4 | TATATATA/TATATATA | | 7 |
| chrX | 112257880 | ATATATGTGTATA-TATATATA-TACACATATATAT | loxF8L | 4 | loxF8L | 3 | TATATATA/TATATATA | | 7 |
| chr7 | 97651085 | ATATATCTATATA-TATATATA-TACACATATATAT | loxF8L | 4 | loxF8L | 3 | TATATATA/TATATATA | | 7 |
| chr12 | 11331864 | ATATATGTGAGCA-TATATATA-TACACATATATAT | loxF8L | 4 | loxF8L | 3 | TATATATA/TATATATA | | 7 |
| chr20 | 31296063 | ATATATATTTGAA-TATATATA-TACACATATATAT | loxF8L | 4 | loxF8L | 3 | TATATATA/TATATATA | | 7 |
| chr20 | 31296091 | ATATATATTTGAA-TATATATA-TACACATATATAT | loxF8L | 4 | loxF8L | 3 | TATATATA/TATATATA | | 7 |
| chr20 | 31296225 | ATATATATTTGAA-TATATATA-TACACATATATAT | loxF8L | 4 | loxF8L | 3 | TATATATA/TATATATA | | 7 |
| chr20 | 31296251 | ATATATATTTGAA-TATATATA-TACACATATATAT | loxF8L | 4 | loxF8L | 3 | TATATATA/TATATATA | | 7 |
| chr20 | 31296169 | ATATATATTTGAA-TATATATA-TACACATATATAT | loxF8L | 4 | loxF8L | 3 | TATATATA/TATATATA | | 7 |
| chr12 | 66852481 | CTAACAAAGTGTG-TATATATA-CACACATATATAT | loxF8R | 3 | loxF8L | 4 | TATATATA/TATATATA | | 7 |
| chr10 | 54779453 | ATATATGTGTGTA-TATATATA-CACACATATATAT | loxF8L | 3 | loxF8L | 4 | TATATATA/TATATATA | | 7 |
| chr13 | 36609205 | ATATATGTGTGTA-TATATATA-CACACATATATAT | loxF8L | 3 | loxF8L | 4 | TATATATA/TATATATA | | 7 |
| chr15 | 53515042 | ATATATATGTGTG-TATATATA-TACACACATATAT | loxF8L | 4 | loxF8L | 3 | TATATATA/TATATATA | | 7 |
| chr15 | 53515044 | ATATATGTGTGTA-TATATATA-CACACATATATAT | loxF8L | 3 | loxF8L | 4 | TATATATA/TATATATA | | 7 |
| chr3 | 29007503 | ATATATATGTGTG-TATATATA-TACACACATATAT | loxF8L | 4 | loxF8L | 3 | TATATATA/TATATATA | | 7 |
| chr3 | 29007505 | ATATATGTGTGTA-TATATATA-CACACATATATAT | loxF8L | 3 | loxF8L | 4 | TATATATA/TATATATA | | 7 |
| chr7 | 31590039 | ATATATATGTGTG-TATATATA-TACACACATATAT | loxF8L | 4 | loxF8L | 3 | TATATATA/TATATATA | | 7 |
| chr7 | 137638467 | ATATATGTGTGTA-TATATATA-CACACATATATAT | loxF8L | 3 | loxF8L | 4 | TATATATA/TATATATA | | 7 |
| chr7 | 137638541 | ATATATATGTGTG-TATATATA-TACACACATATAT | loxF8L | 4 | loxF8L | 3 | TATATATA/TATATATA | | 7 |
| chr10 | 89115137 | ATATATATGTGTT-TATATATA-TACACACATATAT | loxF8L | 4 | loxF8L | 3 | TATATATA/TATATATA | | 7 |
| chr2 | 30601868 | ATCTATCTATAGA-TATATATA-TCTATAGATATAT | loxF8L | 4 | loxF8L | 3 | TATATATA/TATATATA | | 7 |
| chr11 | 71942763 | ATATATCTGTATA-TATATATA-TATATAGATATAT | loxF8L | 3 | loxF8L | 4 | TATATATA/TATATATA | | 7 |
| chr13 | 68285878 | ATATATCTATATA-TATATATA-TGCACACATATAT | loxF8L | 4 | loxF8L | 3 | TATATATA/TATATATA | | 7 |
| chr1 | 69034977 | ATATATGTGTGTG-TATATATA-TACATAGATATAT | loxF8L | 4 | loxF8L | 3 | TATATATA/TATATATA | | 7 |
| chr13 | 19489713 | ATATATCTGTATA-TATATATA-CACACACACATAT | loxF8L | 3 | loxF8R | 4 | TATATATA/TATATATA | | 7 |
| chr10 | 28638369 | ATATATGTATGTA-TATATATA-CACACAGATATAT | loxF8L | 4 | loxF8L | 3 | TATATATA/TATATATA | | 7 |
| chr13 | 86929580 | ATATATCTGTTTA-TATATATA-CACACAAACACAC | loxF8L | 3 | loxF8R | 4 | TATATATA/TATATATA | | 7 |
| chr7 | 84791672 | ATATATGTATGCA-TATATATA-TACACACATATAT | loxF8L | 4 | loxF8L | 3 | TATATATA/TATATATA | | 7 |
| chrX | 5566164 | ATATATGTATGTA-TATATATA-CACATACTATTAG | loxF8L | 4 | loxF8R | 3 | TATATATA/TATATATA | | 7 |
| chr4 | 21200315 | ATATATGTGTGTA-TATATATA-TACATACATATAT | loxF8L | 3 | loxF8L | 4 | TATATATA/TATATATA | | 7 |
| chr4 | 21200343 | ATATATGTGTGTA-TATATATA-TACATACATATAT | loxF8L | 3 | loxF8L | 4 | TATATATA/TATATATA | | 7 |
| chr12 | 11331808 | ATATATGTGAGCA-TATATATA-TACACACATATAT | loxF8L | 4 | loxF8L | 3 | TATATATA/TATATATA | | 7 |
| chr16 | 7912699 | ATATATGTGAGTA-TATATATA-TACACACATATAT | loxF8L | 4 | loxF8L | 3 | TATATATA/TATATATA | | 7 |
| chrX | 147440881 | ATATATGTGTGTA-TATATATA-TGTACACATATAT | loxF8L | 3 | loxF8L | 4 | TATATATA/TATATATA | | 7 |
| chr10 | 128732605 | ATATATGTGTATA-TATATATA-TACACACATATAT | loxF8L | 4 | loxF8L | 3 | TATATATA/TATATATA | | 7 |
| chr11 | 41034464 | ATATATGTGTGTA-TATATATA-TATACACATATAT | loxF8L | 3 | loxF8L | 4 | TATATATA/TATATATA | | 7 |
| chr12 | 63751277 | ATATATGTGTATA-TATATATA-TACACACATATAT | loxF8L | 4 | loxF8L | 3 | TATATATA/TATATATA | | 7 |
| chr12 | 63751245 | ATATATGTGTGTA-TATATATA-TATACACATATAT | loxF8L | 3 | loxF8L | 4 | TATATATA/TATATATA | | 7 |
| chr13 | 72927934 | ATATATGTGTGTA-TATATATA-TATACACATATAT | loxF8L | 3 | loxF8L | 4 | TATATATA/TATATATA | | 7 |
| chr13 | 87193180 | ATATATGTGTATA-TATATATA-TACACACATATAT | loxF8L | 4 | loxF8L | 3 | TATATATA/TATATATA | | 7 |
| chr2 | 75379260 | ATATATGTGTATA-TATATATA-TACACACATATAT | loxF8L | 4 | loxF8L | 3 | TATATATA/TATATATA | | 7 |
| chr3 | 158019574 | ATATATGTGTGTA-TATATATA-TATACACATATAT | loxF8L | 3 | loxF8L | 4 | TATATATA/TATATATA | | 7 |
| chr7 | 30194306 | ATATATGTGTGTA-TATATATA-TATACACATATAT | loxF8L | 3 | loxF8L | 4 | TATATATA/TATATATA | | 7 |
| chrX | 2893817 | ATATATGTGTATA-TATATATA-TACACACATATAT | loxF8L | 4 | loxF8L | 3 | TATATATA/TATATATA | | 7 |
| chr12 | 44470472 | ATATATGTGTGCA-TATATATA-CACACACACATAT | loxF8L | 3 | loxF8R | 4 | TATATATA/TATATATA | | 7 |
| chr14 | 95778169 | ATATATGTGTGCA-TATATATA-TACACACATACAT | loxF8L | 3 | loxF8L | 4 | TATATATA/TATATATA | | 7 |
| chr7 | 52719149 | TATATATTGTGTG-TATATATA-TACACACATATAT | loxF8R | 4 | loxF8L | 3 | TATATATA/TATATATA | | 7 |
| chr12 | 107915398 | ATATATGTGTGTA-TATATATA-CACACACACAGAG | loxF8L | 3 | loxF8R | 4 | TATATATA/TATATATA | | 7 |
| chr1 | 107753471 | GTATGTGTGTGTG-TATATATA-TACACACATATAT | loxF8R | 4 | loxF8L | 3 | TATATATA/TATATATA | HG7 | 7 |
| chr4 | 27402951 | GTATGTGTGTGTG-TATATATA-TACACACATATAT | loxF8R | 4 | loxF8L | 3 | TATATATA/TATATATA | HG7 | 7 |
| chr5 | 174895874 | ATATATGTGTGTA-TATATATA-CACACACACATAC | loxF8L | 3 | loxF8R | 4 | TATATATA/TATATATA | HG7 | 7 |
| chr6 | 120026275 | GTATGTGTGTGTG-TATATATA-TACACACATATAT | loxF8R | 4 | loxF8L | 3 | TATATATA/TATATATA | HG7 | 7 |
| chr6 | 93476680 | GTATGTGTGTGTG-TATATATA-TACACACATATAT | loxF8R | 4 | loxF8L | 3 | TATATATA/TATATATA | HG7 | 7 |
| chrX | 66285884 | GTATGTGTGTGTG-TATATATA-TACACACATATAT | loxF8R | 4 | loxF8L | 3 | TATATATA/TATATATA | HG7 | 7 |
| chr10 | 33169998 | ATATATGTGTGTA-TATATATA-CACACACACATAT | loxF8L | 3 | loxF8R | 4 | TATATATA/TATATATA | | 7 |
| chr4 | 46473488 | ATATGTGTGTGTG-TATATATA-TACACACATATAT | loxF8R | 4 | loxF8L | 3 | TATATATA/TATATATA | | 7 |
| chr9 | 28057510 | ATATATGTGTGTA-TATATATA-CACACACACATAT | loxF8L | 3 | loxF8R | 4 | TATATATA/TATATATA | | 7 |
| chr12 | 11874403 | ATATATGTGTGTA-TATATATA-CACACACATATAT | loxF8L | 3 | loxF8L | 4 | TATATATA/TATATATA | | 7 |
| chr3 | 46180658 | ATATATGTGTGTA-TATATATA-CACACACATATAT | loxF8L | 3 | loxF8L | 4 | TATATATA/TATATATA | | 7 |
| chr4 | 104604579 | ATATATGTGTGTG-TATATATA-TACACACATATAT | loxF8L | 4 | loxF8L | 3 | TATATATA/TATATATA | | 7 |
| chr6 | 56636351 | ATATATGTGTGTG-TATATATA-TACACACATATAT | loxF8L | 4 | loxF8L | 3 | TATATATA/TATATATA | | 7 |
| chr7 | 78070126 | ATATATGTGTGTG-TATATATA-TACACACATATAT | loxF8L | 4 | loxF8L | 3 | TATATATA/TATATATA | | 7 |
| chr7 | 137638505 | ATATATGTGTGTA-TATATATA-CACACACATATAT | loxF8L | 3 | loxF8L | 4 | TATATATA/TATATATA | | 7 |
| chr9 | 80225247 | ATATATGTGTGTA-TATATATA-CACACACATATAT | loxF8L | 3 | loxF8L | 4 | TATATATA/TATATATA | | 7 |
| chr17 | 69463455 | ATATGTATGTGTG-TATATATA-TACACACATATAT | loxF8R | 4 | loxF8L | 3 | TATATATA/TATATATA | | 7 |
| chr2 | 56617368 | ATATGTATGTGTG-TATATATA-TACACACATATAT | loxF8R | 4 | loxF8L | 3 | TATATATA/TATATATA | | 7 |
| chr7 | 31590041 | ATATATGTGTGTA-TATATATA-CACACATATATAG | loxF8L | 3 | loxF8R | 4 | TATATATA/TATATATA | | 7 |
| chr15 | 58679353 | ATATATGTGTGTA-TATATATA-CACACATGCATAC | loxF8L | 3 | loxF8R | 4 | TATATATA/TATATATA | | 7 |
| chr11 | 22821091 | CAGATATTATGTG-TATATATA-TACACACATATAT | loxF8R | 4 | loxF8L | 3 | TATATATA/TATATATA | | 7 |
| chr10 | 54779503 | ATATATGTGTGTA-TATATATA-TACACACACATAT | loxF8L | 3 | loxF8L | 4 | TATATATA/TATATATA | | 7 |
| chr11 | 133499797 | ATATGTGTGTGTA-TATATATA-TACACACATATAT | loxF8L | 4 | loxF8L | 3 | TATATATA/TATATATA | | 7 |
| chr13 | 67613345 | ATATATGTGTGTA-TATATATA-TACACACACATAT | loxF8L | 3 | loxF8L | 4 | TATATATA/TATATATA | | 7 |

| | | | | | | | | |
|---|---|---|---|---|---|---|---|---|
| chr13 | 75066705 | ATATATGTGTGTA-TATATATA-TACACACACATAT | loxF8L | 3 | loxF8L | 4 | TATATATA/TATATATA | 7 |
| chr18 | 5308512 | ATATGTGTGTGTA-TATATATA-TACACACATATAT | loxF8L | 4 | loxF8L | 3 | TATATATA/TATATATA | 7 |
| chr20 | 53558867 | ATATATGTGTGTA-TATATATA-TACACACACATAT | loxF8L | 3 | loxF8L | 4 | TATATATA/TATATATA | 7 |
| chr21 | 13384085 | ATATATGTGTGTA-TATATATA-TACACACACATAT | loxF8L | 3 | loxF8L | 4 | TATATATA/TATATATA | 7 |
| chr7 | 140298610 | ATATATGTGTGTA-TATATATA-TACACACACATAT | loxF8L | 3 | loxF8L | 4 | TATATATA/TATATATA | 7 |
| chrX | 18418691 | ATATATGTGTGTA-TATATATA-TACACACACATAT | loxF8L | 3 | loxF8L | 4 | TATATATA/TATATATA | 7 |
| chr10 | 43479074 | ATGTATGTGTGTA-TATATATA-TACACACATATAT | loxF8L | 4 | loxF8L | 3 | TATATATA/TATATATA | 7 |
| chr1 | 181567928 | ATGTATGTGTGTA-TATATATA-TACACACATATAT | loxF8L | 4 | loxF8L | 3 | TATATATA/TATATATA | 7 |
| chr20 | 2517355 | GTATATGTGTGTA-TATATATA-TACACACATATAT | loxF8L | 4 | loxF8L | 3 | TATATATA/TATATATA | 7 |
| chr14 | 29826662 | GTATATATGTGTA-TATATATA-TACACACATATAT | loxF8L | 4 | loxF8L | 3 | TATATATA/TATATATA | 7 |
| chr2 | 51536800 | GTATATATGTGTA-TATATATA-TACACACATATAT | loxF8L | 4 | loxF8L | 3 | TATATATA/TATATATA | 7 |
| chr5 | 111186373 | ATATATGTGTGTA-TATATATA-TACACATATATAC | loxF8L | 3 | loxF8L | 4 | TATATATA/TATATATA | 7 |
| chr3 | 139802729 | GTACATATGTGTA-TATATATA-TACACACATATAT | loxF8L | 4 | loxF8L | 3 | TATATATA/TATATATA | 7 |
| chr12 | 62831748 | ATATATTCGTGTA-TATATATA-TACACACATATAT | loxF8L | 4 | loxF8L | 3 | TATATATA/TATATATA | 7 |
| chr2 | 22391972 | ATATATGTGTGTA-TATATATA-TATAAAATCTTTG | loxF8L | 3 | loxF8R | 4 | TATATATA/TATATATA | 7 |
| chr16 | 63255386 | ATATATTTGTGCA-TATATATA-TACACATATATAA | loxF8L | 3 | loxF8L | 4 | TATATATA/TATATATA | 7 |
| chrX | 57438136 | ATATATTTGTGTA-TATATATA-TACACACATATAC | loxF8L | 3 | loxF8L | 4 | TATATATA/TATATATA | 7 |
| chr12 | 78876544 | ATATGAGTGTGTG-TATATATA-TACACACATATAC | loxF8R | 3 | loxF8L | 4 | TATATATA/TATATATA | 7 |
| chr1 | 244170358 | GTATATCTGTGTA-TATATATA-CACACATACATAT | loxF8L | 3 | loxF8R | 4 | TATATATA/TATATATA | 7 |
| chr14 | 42606943 | GTATGTGTGTGTG-TATATATA-TACACAGACATAT | loxF8R | 4 | loxF8L | 3 | TATATATA/TATATATA | 7 |
| chr6 | 2337243 | ATATGTGTGTGTG-TATATATA-CACACAATTTTGC | loxF8R | 4 | loxF8R | 3 | TATATATA/TATATATA | 7 |
| chr2 | 115458621 | ATGAGTTTGTGGA-TATATATA-CACACACACATAT | loxF8L | 3 | loxF8R | 4 | TATATATA/TATATATA | 7 |
| chr13 | 37708642 | CTAAAATTATGTA-TATATATA-CACACACACATAC | loxF8R | 3 | loxF8R | 4 | TATATATA/TATATATA | 7 |
| chr4 | 53196029 | CTAGGATAGTGTT-TATATATA-AAAATAATTTTAG | loxF8R | 3 | loxF8R | 4 | TATATATA/TATATATA | 7 |
| chr6 | 4143705 | GTATATGTGTGTA-TATATATA-CACATACTGTTAG | loxF8L | 4 | loxF8R | 3 | TATATATA/TATATATA | 7 |
| --- | | | | | | | | 0 |
| chr3 | 70041583 | TTGAATCTGTGGA-TTCGAGCA-ACCACAGATTTAA | loxF8L | 2 | loxF8L | 2 | TGCTCGAA/TTCGAGCA | 4 |
| chr12 | 97904551 | AGAAAATTGAGGA-TTCGAGCA-TCTACAGATTCTT | loxF8L | 4 | loxF8L | 3 | TGCTCGAA/TTCGAGCA | 7 |
| --- | | | | | | | | 0 |
| chr1 | 160672479 | CTCTGATTTTGTG-AAATCATA-ACCCCAGAATTAT | loxF8R | 3 | loxF8L | 3 | AAATCATA/TATGATTT | 6 |
| chr4 | 45448603 | ATAAATTTGTGGA-TATGATTT-CATAGCATCTTAA | loxF8L | 1 | loxF8R | 4 | AAATCATA/TATGATTT | 5 |
| --- | | | | | | | | 0 |
| chr7 | 55094172 | ATCAATCTGTGGA-AGTGAGTT-TCCACTGACTGAT | loxF8L | 1 | loxF8L | 3 | AACTCACT/AGTGAGTT | 4 |
| chr3 | 118961674 | CTAAGGGTATGGG-AGTGAGTT-CAGACAATAATAG | loxF8R | 4 | loxF8R | 3 | AACTCACT/AGTGAGTT | 7 |
| --- | | | | | | | | 0 |
| chrX | 83165875 | ATAAATGTGTGTT-GTTTACTT-TCCACATATTTGT | loxF8L | 3 | loxF8L | 2 | AAGTAAAC/GTTTACTT | 5 |
| chr9 | 9813018 | ACAAATATGTGGA-AAGTAAAC-AACACACTCTTAA | loxF8L | 2 | loxF8R | 3 | AAGTAAAC/GTTTACTT | 5 |
| chr5 | 180147564 | TTAAGAATGTGTT-GTTTACTT-TCCACAAATTTGT | loxF8R | 3 | loxF8L | 2 | AAGTAAAC/GTTTACTT | 5 |
| chr6 | 34265606 | GCAAATATGTGGA-AAGTAAAC-CACACACTCTTAA | loxF8L | 3 | loxF8R | 2 | AAGTAAAC/GTTTACTT | 5 |
| chrX | 37983849 | ACAAATCTGTGGA-AAGTAAAC-GACACACTCCTAA | loxF8L | 1 | loxF8R | 4 | AAGTAAAC/GTTTACTT | 5 |
| chr5 | 83382581 | ACAAATATGTGGA-AAGTAAAC-AACATACTCTTAA | loxF8L | 2 | loxF8R | 4 | AAGTAAAC/GTTTACTT | 6 |
| chr2 | 32794312 | ACAAAATTATGGA-AAGTAAAC-AACACACTCTTAA | loxF8L | 4 | loxF8R | 3 | AAGTAAAC/GTTTACTT | 7 |
| chr11 | 67297914 | TTAGGAGTGTGTT-GTTTACTT-TCTACATATTTGT | loxF8R | 4 | loxF8L | 3 | AAGTAAAC/GTTTACTT | 7 |
| chr2 | 64650051 | ATCAAATTGTGGG-GTTTACTT-CAGAGAATCCTAG | loxF8L | 4 | loxF8R | 3 | AAGTAAAC/GTTTACTT | 7 |
| chr19 | 39078654 | CTCAGATTGGGTA-AAGTAAAC-CACACAATGGAAT | loxF8R | 3 | loxF8R | 4 | AAGTAAAC/GTTTACTT | 7 |
| chr2 | 189860522 | GTAAAATTGTGTT-GTTTACTT-TCCAAATATTTGG | loxF8R | 3 | loxF8L | 4 | AAGTAAAC/GTTTACTT | 7 |
| --- | | | | | | | | 0 |
| chrX | 135588536 | ATAAATATGTGGA-AATAAAAC-AACACACTCTTAA | loxF8L | 1 | loxF8R | 3 | AATAAAAC/GTTTTATT | 4 |
| chr4 | 62676017 | GTAAGACTGTGTT-GTTTTATT-TCTACATATTTAT | loxF8R | 3 | loxF8L | 2 | AATAAAAC/GTTTTATT | 5 |
| chr9 | 69703604 | TTATGAGTGTGTT-GTTTTATT-TCCACATATTTAT | loxF8R | 4 | loxF8L | 1 | AATAAAAC/GTTTTATT | 5 |
| chr7 | 152199683 | ACAAAATTATGGA-AATAAAAC-CACACAATTATAC | loxF8L | 4 | loxF8R | 3 | AATAAAAC/GTTTTATT | 7 |
| chr1 | 48431636 | ACCAATTTATGGA-AATAAAAC-AACACACTCTTAA | loxF8L | 4 | loxF8R | 3 | AATAAAAC/GTTTTATT | 7 |
| chr7 | 2066102 | ATAAAGGTCTCGA-AATAAAAC-CACACACTCCAAG | loxF8L | 4 | loxF8R | 3 | AATAAAAC/GTTTTATT | 7 |
| chr15 | 89967252 | ATAAATTGGTGGG-AATAAAAC-AACACACTTTTAA | loxF8L | 3 | loxF8R | 4 | AATAAAAC/GTTTTATT | 7 |
| chr2 | 17542065 | GAAAAGCTGGGGA-GTTTTATT-TCTACATATTAAT | loxF8L | 4 | loxF8L | 3 | AATAAAAC/GTTTTATT | 7 |
| --- | | | | | | | | 0 |
| chr1 | 44159649 | TTAAGAGTGTGTT-TTTTAATT-TCCACAGATTTGT | loxF8R | 3 | loxF8L | 1 | AATTAAAA/TTTTAATT | 4 |
| chrX | 84733195 | TTAAGAGTGTGTT-TTTTAATT-TCCACATATTTGT | loxF8R | 3 | loxF8L | 2 | AATTAAAA/TTTTAATT | 5 |
| chr4 | 167219769 | AGAAATCATTGAA-TTTTAATT-TACTCAAATTTAT | loxF8L | 4 | loxF8L | 3 | AATTAAAA/TTTTAATT | 7 |
| chr6 | 51512470 | AGAAATGTGTGCA-TTTTAATT-GCCTCTGATTTAA | loxF8L | 3 | loxF8L | 4 | AATTAAAA/TTTTAATT | 7 |
| chr10 | 113841834 | CTAAATTTGAAGA-AATTAAAA-GCCCCAGCTTTAT | loxF8L | 4 | loxF8L | 3 | AATTAAAA/TTTTAATT | 7 |
| --- | | | | | | | | 0 |
| chr1 | 60311594 | ACAAATCTGTGGA-AATTAAAG-CACATATTCTTAA | loxF8L | 1 | loxF8R | 3 | AATTAAAG/CTTTAATT | 4 |
| chr18 | 22508252 | GAAAGAGTGTGTT-CTTTAATT-TCCACATATTTGT | loxF8R | 4 | loxF8L | 2 | AATTAAAG/CTTTAATT | 6 |
| chr5 | 126398865 | TTAGGAGTGTGTA-CTTTAATT-TCCACATATTTGT | loxF8R | 4 | loxF8L | 2 | AATTAAAG/CTTTAATT | 6 |
| chrX | 145784262 | ACAAATATGTGTA-AATTAAAG-AACACATTCCTGG | loxF8L | 3 | loxF8R | 4 | AATTAAAG/CTTTAATT | 7 |
| chr1 | 102273538 | TTAAGAGCATGTG-CTTTAATT-TCCACATATCTGT | loxF8R | 4 | loxF8L | 3 | AATTAAAG/CTTTAATT | 7 |
| chr21 | 23628294 | TTAAAACTGTGTA-CTTTAATT-TTCAAATATTTAG | loxF8L | 3 | loxF8L | 4 | AATTAAAG/CTTTAATT | 7 |
| --- | | | | | | | | 0 |
| chr18 | 60817110 | TTAAGAATGTGTC-ATTTAATT-TCCACATATTTTT | loxF8R | 3 | loxF8L | 2 | AATTAAAT/ATTTAATT | 5 |
| chr9 | 93877388 | ACAAATATGTGAA-AATTAAAT-AACACAGTCTTAG | loxF8L | 3 | loxF8R | 2 | AATTAAAT/ATTTAATT | 5 |
| chr3 | 116333628 | ACAAATTTGTGGA-AATTAAAT-AACACACTCTTAA | loxF8L | 2 | loxF8R | 3 | AATTAAAT/ATTTAATT | 5 |
| chr7 | 153212209 | TTAAGAGTGTGTT-ATTTAATT-TCCACAAATTTGT | loxF8R | 3 | loxF8L | 2 | AATTAAAT/ATTTAATT | 5 |
| chr6 | 65224271 | ATAAATATGTGAA-AATTAAAT-GACACACTCTTAA | loxF8L | 2 | loxF8R | 3 | AATTAAAT/ATTTAATT | 5 |
| chr9 | 13887007 | AAAAACTTGTGGA-AATTAAAT-GACACAGTCTTAA | loxF8L | 3 | loxF8R | 3 | AATTAAAT/ATTTAATT | 6 |
| chr18 | 44411799 | GGAAATCTGGGGA-ATTTAATT-TACTCAGATTTTT | loxF8L | 3 | loxF8L | 3 | AATTAAAT/ATTTAATT | 6 |
| chr9 | 75165097 | TTAAGAATGTGTA-ATTTAATT-TCCATATATTTGT | loxF8R | 3 | loxF8L | 3 | AATTAAAT/ATTTAATT | 6 |
| chr8 | 30024247 | ACAAATATGTGAA-AATTAAAT-AACACACTCTTAA | loxF8L | 3 | loxF8R | 3 | AATTAAAT/ATTTAATT | 6 |
| chr2 | 128461650 | ATAAACAGGTGGA-AATTAAAT-AACACATTCTTAC | loxF8L | 3 | loxF8R | 3 | AATTAAAT/ATTTAATT | 6 |
| chr16 | 76630302 | TGAGGATTGTGTT-ATTTAATT-TCCACATATTTAT | loxF8R | 4 | loxF8L | 1 | AATTAAAT/ATTTAATT | 5 |
| chr9 | 97816121 | TTAAGAGTATGTT-ATTTAATT-TCCACATATTTTT | loxF8R | 4 | loxF8L | 2 | AATTAAAT/ATTTAATT | 6 |
| chr9 | 83088012 | TTAGGAATGTGTT-ATTTAATT-TCCACATATTTGT | loxF8R | 4 | loxF8L | 2 | AATTAAAT/ATTTAATT | 6 |
| chr9 | 103075551 | ACAAATATGTGGA-AATTAAAT-AACACATTCCTGG | loxF8L | 2 | loxF8R | 4 | AATTAAAT/ATTTAATT | 6 |
| chr1 | 68426718 | TCAAGAGTGTGCG-ATTTAATT-TCCACATATTTGT | loxF8R | 4 | loxF8L | 2 | AATTAAAT/ATTTAATT | 6 |
| chr12 | 30563161 | ACAAATGTATGTA-AATTAAAT-AACACACTCTTAG | loxF8L | 4 | loxF8R | 2 | AATTAAAT/ATTTAATT | 6 |
| chr3 | 171690607 | TTAACAGTGTGTT-ATTTAATT-TCCACAAATTTGT | loxF8R | 4 | loxF8L | 2 | AATTAAAT/ATTTAATT | 6 |
| chr2 | 110709053 | TTAAGAGTGTGCC-ATTTAATT-TCCACAAATTTCT | loxF8R | 4 | loxF8L | 2 | AATTAAAT/ATTTAATT | 6 |

| | | | | | | | | | |
|---|---|---|---|---|---|---|---|---|---|
| chr6 | 74718896 | TTAAGAGTGTATT-ATTTAATT-TCCACAATTTTAT | loxF8R | 4 | loxF8L | 2 | AATTAAAT/ATTTAATT | | 6 |
| chr21 | 38232716 | TTATGAATGTGTT-ATTTAATT-TCCAAATATTTAT | loxF8R | 4 | loxF8L | 2 | AATTAAAT/ATTTAATT | | 6 |
| chr11 | 112443372 | TTAAGAGTGTGCT-ATTTAATT-TCCACAAGTTTGT | loxF8R | 4 | loxF8L | 3 | AATTAAAT/ATTTAATT | | 7 |
| chr7 | 130826910 | TTAGGAGTGTGTT-ATTTAATT-TCCACGTATTTGT | loxF8R | 4 | loxF8L | 3 | AATTAAAT/ATTTAATT | | 7 |
| chrX | 83767766 | TTAACAGTGTATG-ATTTAATT-TCCACGTATTTGT | loxF8R | 4 | loxF8L | 3 | AATTAAAT/ATTTAATT | | 7 |
| chr10 | 106165479 | ACAAATATATGGA-AATTAAAT-GACACACTCCTAA | loxF8L | 3 | loxF8R | 4 | AATTAAAT/ATTTAATT | | 7 |
| chr18 | 22895476 | ACAAATATCTGGA-AATTAAAT-AACACAATTCTAC | loxF8L | 3 | loxF8R | 4 | AATTAAAT/ATTTAATT | | 7 |
| chr11 | 85569484 | TTAAGAGTGTATT-ATTTAATT-TCAACATATTTGT | loxF8R | 4 | loxF8L | 3 | AATTAAAT/ATTTAATT | | 7 |
| chr12 | 71254931 | ACAAATATTTGGA-AATTAAAT-AACACACTCTTGA | loxF8L | 3 | loxF8R | 4 | AATTAAAT/ATTTAATT | | 7 |
| chr13 | 39885843 | ACAAATATTTGGA-AATTAAAT-AACATACTCTTAA | loxF8L | 3 | loxF8R | 4 | AATTAAAT/ATTTAATT | | 7 |
| chr11 | 97828833 | TTAGGAGTGTTTG-ATTTAATT-TCCAAATATTTGT | loxF8R | 4 | loxF8L | 3 | AATTAAAT/ATTTAATT | | 7 |
| chr6 | 77405273 | TTAGGAGTGTGTC-ATTTAATT-TCCACATATATGT | loxF8R | 4 | loxF8L | 3 | AATTAAAT/ATTTAATT | | 7 |
| chr1 | 108417453 | ATAAATGGGTAGA-AATTAAAT-AACACACTCTTGA | loxF8L | 3 | loxF8R | 4 | AATTAAAT/ATTTAATT | | 7 |
| chr1 | 108276891 | TCAAGAGTGTGTT-ATTTAATT-TCTACCCATTTAT | loxF8R | 4 | loxF8L | 3 | AATTAAAT/ATTTAATT | | 7 |
| chr14 | 27875654 | TTAATAGTGTGTC-ATTTAATT-TTCAAACATTTAT | loxF8R | 4 | loxF8L | 3 | AATTAAAT/ATTTAATT | | 7 |
| chrX | 83200258 | ATACATATGCGGA-AATTAAAT-TACACACTATTAA | loxF8L | 3 | loxF8R | 4 | AATTAAAT/ATTTAATT | | 7 |
| chr3 | 27158865 | ATATTTTTGTGTG-ATTTAATT-TTCACAAATATAT | loxF8R | 4 | loxF8L | 3 | AATTAAAT/ATTTAATT | | 7 |
| chr4 | 108469120 | CAAAATATGTGGA-AATTAAAT-AACACAATTCTAA | loxF8L | 3 | loxF8R | 4 | AATTAAAT/ATTTAATT | | 7 |
| chr3 | 136124088 | CAAAATATTTGGA-AATTAAAT-GACACACTCTTAA | loxF8L | 4 | loxF8R | 3 | AATTAAAT/ATTTAATT | | 7 |
| chr12 | 60604638 | TTTAAATTGTGGA-AATTAAAT-AACACACTCTAAG | loxF8L | 4 | loxF8R | 3 | AATTAAAT/ATTTAATT | | 7 |
| chr14 | 49812665 | TTATAACTGTGGT-ATTTAATT-TCCACATACTTAC | loxF8L | 4 | loxF8L | 3 | AATTAAAT/ATTTAATT | | 7 |
| --- | | | | | | | | | 0 |
| chr7 | 98095428 | GTAAGAGTGTGTT-GCTTAATT-TCCACATATTTGT | loxF8R | 3 | loxF8L | 2 | AATTAAGC/GCTTAATT | | 5 |
| chr2 | 161151045 | TTAAGAATGTGTT-GCTTAATT-TGCACATATTTGT | loxF8R | 3 | loxF8L | 3 | AATTAAGC/GCTTAATT | | 6 |
| chr11 | 61939241 | ACAAATATGTGGG-AATTAAGC-AACACACTCTTAA | loxF8L | 3 | loxF8R | 3 | AATTAAGC/GCTTAATT | | 6 |
| chr1 | 107085327 | TTAAGAGTGTGTT-GCTTAATT-TCTACAAATTTGT | loxF8R | 3 | loxF8L | 3 | AATTAAGC/GCTTAATT | | 6 |
| chrX | 154301104 | GTAAGAGTGTGTT-GCTTAATT-TTCACAAATTTGT | loxF8R | 3 | loxF8L | 3 | AATTAAGC/GCTTAATT | | 6 |
| chr3 | 27894636 | ACAAATCTGTGGA-AATTAAGC-AACACACTGTTAA | loxF8L | 1 | loxF8R | 4 | AATTAAGC/GCTTAATT | | 5 |
| chr16 | 21336920 | TTAAGAGTGTTTT-GCTTAATT-TCCACATATTTAT | loxF8R | 4 | loxF8L | 1 | AATTAAGC/GCTTAATT | | 5 |
| chr12 | 21260808 | TTAAGAGTATGTT-GCTTAATT-TCTACAAATTTAT | loxF8R | 4 | loxF8L | 2 | AATTAAGC/GCTTAATT | | 6 |
| chr1 | 219885605 | ATAGATATGTGGA-AATTAAGC-AACACATTCTTGA | loxF8L | 2 | loxF8R | 4 | AATTAAGC/GCTTAATT | | 6 |
| chr6 | 95969294 | GTAAATATGTGGA-AATTAAGC-AACACACTTATAG | loxF8L | 2 | loxF8R | 4 | AATTAAGC/GCTTAATT | | 6 |
| chr11 | 107090794 | ACAAATATGTGAA-AATTAAGC-AACACACTCTTGA | loxF8L | 3 | loxF8R | 4 | AATTAAGC/GCTTAATT | | 7 |
| chr3 | 14748716 | TTAATAATGTGTT-GCTTAATT-TTCACATATTTGT | loxF8R | 4 | loxF8L | 3 | AATTAAGC/GCTTAATT | | 7 |
| chr5 | 29721691 | GTAAAAGTATGTG-GCTTAATT-TCCAAATATTTGT | loxF8R | 4 | loxF8L | 3 | AATTAAGC/GCTTAATT | | 7 |
| chr4 | 21464458 | TCAAGAGTGTGTT-GCTTAATT-TCCACATAATTGT | loxF8R | 4 | loxF8L | 3 | AATTAAGC/GCTTAATT | | 7 |
| chr2 | 86533947 | TTAAGAATGTGTT-GCTTAATT-TCCATACATCTGT | loxF8R | 3 | loxF8L | 4 | AATTAAGC/GCTTAATT | | 7 |
| chr13 | 60979196 | AGAAATATATGGA-AATTAAGC-AACATATTCTTAC | loxF8L | 3 | loxF8R | 4 | AATTAAGC/GCTTAATT | | 7 |
| chr20 | 11595401 | AGAAATGAGTGAA-AATTAAGC-AACACACTCTTAA | loxF8L | 4 | loxF8R | 3 | AATTAAGC/GCTTAATT | | 7 |
| --- | | | | | | | | | 0 |
| chr15 | 26250510 | TTAAGAGTGTGTT-GTGTAATT-TCCACAAATTTGT | loxF8R | 3 | loxF8L | 2 | AATTACAC/GTGTAATT | | 5 |
| chr12 | 98016158 | ATAAATATGTGGA-AATTACAC-AACACACTCCTAA | loxF8L | 1 | loxF8R | 4 | AATTACAC/GTGTAATT | | 5 |
| chr10 | 43244657 | ACAAATATGTGGA-AATTACAC-AACATATTCTTAA | loxF8L | 2 | loxF8R | 4 | AATTACAC/GTGTAATT | | 6 |
| chr15 | 26918269 | ACAAATGTGTGTT-GTGTAATT-TCCACATATGTAT | loxF8L | 4 | loxF8L | 2 | AATTACAC/GTGTAATT | | 6 |
| chrX | 18753714 | TTAAGAGTATGTT-GTGTAATT-TCTACATATTTAT | loxF8R | 4 | loxF8L | 2 | AATTACAC/GTGTAATT | | 6 |
| chr6 | 97479125 | TTAAGTATGTGGT-GTGTAATT-TCTACATATTTTT | loxF8L | 4 | loxF8L | 3 | AATTACAC/GTGTAATT | | 7 |
| chr13 | 26317652 | ACAAAATTGTGGA-AATTACAC-AACACACTCTTTA | loxF8L | 3 | loxF8R | 4 | AATTACAC/GTGTAATT | | 7 |
| chr1 | 173073690 | ACAAATATATGGA-AATTACAC-AACACACTCTAAA | loxF8L | 3 | loxF8R | 4 | AATTACAC/GTGTAATT | | 7 |
| chr11 | 121738785 | TTAAGACTATGTT-GTGTAATT-TCCACAAATGTGT | loxF8R | 4 | loxF8L | 3 | AATTACAC/GTGTAATT | | 7 |
| --- | | | | | | | | | 0 |
| chr8 | 131414092 | ATTAATCTGTGCT-CACACCCA-TTCACAGATTTAT | loxF8L | 3 | loxF8L | 1 | CACACCCA/TGGGTGTG | | 4 |
| chr1 | 158819728 | CTCAGCTTATCTG-CACACCCA-TCCACATATACAT | loxF8R | 4 | loxF8L | 3 | CACACCCA/TGGGTGTG | | 7 |
| --- | | | | | | | | | 0 |
| chr2 | 162585883 | ATAAATGTGTGAA-CTCATAAA-GCCACAGAATGAT | loxF8L | 2 | loxF8L | 3 | CTCATAAA/TTTATGAG | | 5 |
| chr9 | 130828900 | ATAAATCTCTGGA-TTTATGAG-TGCCCAGAATAAT | loxF8L | 1 | loxF8L | 4 | CTCATAAA/TTTATGAG | | 5 |
| --- | | | | | | | | | 0 |
| chr1 | 117384368 | TTAAGAGTGTGTT-GTTTAATA-TCCACATATTTAT | loxF8R | 3 | loxF8L | 1 | GTTTAATA/TATTAAAC | | 4 |
| chr13 | 20015221 | TTAAGAGTGTGTA-GTTTAATA-TCCCCATATTTGT | loxF8R | 3 | loxF8L | 3 | GTTTAATA/TATTAAAC | | 6 |
| chr3 | 121588918 | ACAAATATGTGGA-TATTAAAC-AACACATTCCTAA | loxF8L | 2 | loxF8R | 4 | GTTTAATA/TATTAAAC | | 6 |
| chr6 | 37421773 | TTAAGAGTATGTT-GTTTAATA-TCCACAAATTTGT | loxF8R | 4 | loxF8L | 2 | GTTTAATA/TATTAAAC | | 6 |
| chr5 | 84456236 | AAAAATATCTGGA-TATTAAAC-CACATATTCCTAA | loxF8L | 3 | loxF8R | 4 | GTTTAATA/TATTAAAC | | 7 |
| chr5 | 122678753 | ACACATATGTGGA-TATTAAAC-AACACATTCCTAA | loxF8L | 3 | loxF8R | 4 | GTTTAATA/TATTAAAC | | 7 |
| chr12 | 59794458 | TTAGAAGTGTGTG-GTTTAATA-TCCAAATATTTAG | loxF8R | 4 | loxF8L | 3 | GTTTAATA/TATTAAAC | | 7 |
| chr5 | 111105757 | GAAAGATTTTGTT-TATTAAAC-AACAAAATCTTAC | loxF8R | 4 | loxF8R | 3 | GTTTAATA/TATTAAAC | | 7 |
| chr10 | 79427546 | GCAAATATGTGGA-TATTAAAC-AACACACTCCTAA | loxF8L | 3 | loxF8R | 4 | GTTTAATA/TATTAAAC | | 7 |
| chr2 | 19208149 | TAAAGTCTGTGCA-GTTTAATA-TCACCAGATTTAA | loxF8L | 4 | loxF8L | 3 | GTTTAATA/TATTAAAC | | 7 |
| --- | | | | | | | | | 0 |
| chr5 | 118438609 | ATATATCTATAGA-TATAGATA-TCCACAGATATAT | loxF8L | 3 | loxF8L | 1 | TATAGATA/TATCTATA | HG2L | 4 |
| chr5 | 118438581 | ATATATCTATAGA-TATAGATA-TCCACAGATATAT | loxF8L | 3 | loxF8L | 1 | TATAGATA/TATCTATA | HG2L | 4 |
| chr5 | 118438637 | ATATATCTATAGA-TATAGATA-TCCACAGATATAT | loxF8L | 3 | loxF8L | 1 | TATAGATA/TATCTATA | HG2L | 4 |
| chr5 | 118438665 | ATATATCTATAGA-TATAGATA-TCCACAGATATAT | loxF8L | 3 | loxF8L | 1 | TATAGATA/TATCTATA | HG2L | 4 |
| chr1 | 43911588 | ATATATCTGTAGC-TATAGATA-TCTACAGATATAT | loxF8L | 3 | loxF8L | 2 | TATAGATA/TATCTATA | | 5 |
| chr1 | 43911616 | ATATATCTATAGA-TATCTATA-TCTATAGATATAT | loxF8L | 3 | loxF8L | 3 | TATAGATA/TATCTATA | | 6 |
| chr5 | 118438553 | ATATATCTATAGA-TATAGATA-TCCACAGATATAT | loxF8L | 4 | loxF8L | 1 | TATAGATA/TATCTATA | | 5 |
| chr4 | 67468058 | ATATATCTATAGA-TATAGATA-TCTTTAGATTTTT | loxF8L | 3 | loxF8L | 4 | TATAGATA/TATCTATA | | 7 |
| chr1 | 43911560 | ACATATCTGTAGC-TATAGATA-TCTATAGATATAT | loxF8L | 4 | loxF8L | 3 | TATAGATA/TATCTATA | | 7 |
| chr8 | 32364146 | ATATATGTGTGCA-TATAGATA-TCTACAGATGACT | loxF8L | 3 | loxF8L | 4 | TATAGATA/TATCTATA | | 7 |
| chr6 | 126857379 | ATAAAAGTGTGTA-TATAGATA-TATACACATATAT | loxF8L | 3 | loxF8L | 4 | TATAGATA/TATCTATA | | 7 |
| chr8 | 136992129 | ATACATATGTAGA-TATCTATA-TCCATATATACAT | loxF8L | 3 | loxF8L | 4 | TATAGATA/TATCTATA | | 7 |
| chr8 | 136991877 | ATACATATGTAGA-TATCTATA-TCCATATATACAT | loxF8L | 3 | loxF8L | 4 | TATAGATA/TATCTATA | | 7 |
| chr8 | 136991961 | ATACATATGTAGA-TATCTATA-TCCATATATACAT | loxF8L | 3 | loxF8L | 4 | TATAGATA/TATCTATA | | 7 |
| chr8 | 136991989 | ATACATATGTAGA-TATCTATA-TCCATATATACAT | loxF8L | 3 | loxF8L | 4 | TATAGATA/TATCTATA | | 7 |
| chr8 | 136991933 | ATACATATGTAGA-TATCTATA-TCCATATATACAT | loxF8L | 3 | loxF8L | 4 | TATAGATA/TATCTATA | | 7 |
| chr8 | 136991905 | ATACATATGTAGA-TATCTATA-TCCATATATACAT | loxF8L | 3 | loxF8L | 4 | TATAGATA/TATCTATA | | 7 |
| chr8 | 136992017 | ATACATATGTAGA-TATCTATA-TCCATATATACAT | loxF8L | 3 | loxF8L | 4 | TATAGATA/TATCTATA | | 7 |
| chr8 | 136992101 | ATACATATGTAGA-TATCTATA-TCCATATATACAT | loxF8L | 3 | loxF8L | 4 | TATAGATA/TATCTATA | | 7 |
| chr8 | 136992045 | ATACATATGTAGA-TATCTATA-TCCATATATACAT | loxF8L | 3 | loxF8L | 4 | TATAGATA/TATCTATA | | 7 |

| | | | | | | | | |
|---|---|---|---|---|---|---|---|---|
| chr8 | 136992073 | ATACATATGTAGA-TATCTATA-TCCATATATACAT | loxF8L | 3 | loxF8L | 4 | TATAGATA/TATCTATA | 7 |
| chr1 | 43911504 | ATACATCTATAGA-TATAGATA-TCTATAGACATAT | loxF8L | 3 | loxF8L | 4 | TATAGATA/TATCTATA | 7 |
| chr13 | 19489641 | ATAGATATGTGTA-TATCTATA-TACACAAATACAT | loxF8L | 3 | loxF8L | 4 | TATAGATA/TATCTATA | 7 |
| chrX | 107781751 | ATAGATCTATAGA-TATCTATA-TATATAGATCTAT | loxF8L | 3 | loxF8L | 4 | TATAGATA/TATCTATA | 7 |
| chr1 | 72216406 | ATATATATATGTA-TATCTATA-TACACACATATAT | loxF8L | 4 | loxF8L | 3 | TATAGATA/TATCTATA | 7 |
| chr5 | 118438693 | ATATATCTATAGA-TATAGATA-TCTGTAGATATAT | loxF8L | 3 | loxF8L | 4 | TATAGATA/TATCTATA | 7 |
| chr4 | 20472409 | ATATATCTATATA-TATAGATA-TCTATAGATATAT | loxF8L | 4 | loxF8L | 3 | TATAGATA/TATCTATA | 7 |
| chr5 | 55582128 | ATCCATCTATAGA-TATAGATA-TCTATAGATATAT | loxF8L | 4 | loxF8L | 3 | TATAGATA/TATCTATA | 7 |
| chr1 | 43911644 | ATATATCTGTAGC-TATAGATA-TCTATAGATATAG | loxF8L | 3 | loxF8L | 4 | TATAGATA/TATCTATA | 7 |
| chr1 | 193564111 | ATATATTTGCGTA-TATCTATA-TCTACACATATAT | loxF8L | 4 | loxF8L | 3 | TATAGATA/TATCTATA | 7 |
| chr11 | 49512267 | ATATATGTGTGTA-TATCTATA-CACACACATATAT | loxF8L | 3 | loxF8L | 4 | TATAGATA/TATCTATA | 7 |
| chrX | 34627676 | ATATATGTGTGTA-TATCTATA-TCTATAGATGTAG | loxF8L | 3 | loxF8L | 4 | TATAGATA/TATCTATA | 7 |
| chr1 | 189812026 | CTAAGTGTGTATA-TATAGATA-CACACATACTTAA | loxF8R | 4 | loxF8R | 3 | TATAGATA/TATCTATA | 7 |
| --- | | | | | | | | 0 |
| chr6 | 100272064 | TCAAATTTGTGGA-TTTTTTTT-TCCATACATTTAT | loxF8L | 3 | loxF8L | 2 | AAAAAAAA/TTTTTTTT | 5 |
| chr22 | 38628928 | AGAAATGCTTGGA-AAAAAAAA-TTCACAGTATTAT | loxF8L | 4 | loxF8L | 3 | AAAAAAAA/TTTTTTTT | 7 |
| chrX | 6362101 | ATAAATCTTGGGG-AAAAAAAA-AACACCATTTTAT | loxF8L | 3 | loxF8R | 4 | AAAAAAAA/TTTTTTTT | 7 |
| chr8 | 134704885 | ATAAACCTTTAAA-AAAAAAAA-CAAAAAAACTTAG | loxF8L | 4 | loxF8R | 3 | AAAAAAAA/TTTTTTTT | 7 |
| chr11 | 93438207 | ATAATTTTTTGGA-AAAAAAAA-GCTATAGCTTTAT | loxF8L | 3 | loxF8L | 4 | AAAAAAAA/TTTTTTTT | 7 |
| chr8 | 9564363 | ATAAATATGTTAA-AAAAAAAA-ATCTCAAGCTTAG | loxF8L | 3 | loxF8R | 4 | AAAAAAAA/TTTTTTTT | 7 |
| chr1 | 246393180 | CTGAGACTATGTT-AAAAAAAA-TACACAGAACTAT | loxF8R | 4 | loxF8L | 3 | AAAAAAAA/TTTTTTTT | 7 |
| chr17 | 6620397 | ATTTATCTGAGGA-AAAAAAAA-TACATATATATAT | loxF8L | 3 | loxF8L | 4 | AAAAAAAA/TTTTTTTT | 7 |
| chrY | 3515732 | TGAAATGTGTGGA-TTTTTTTT-TCAACAAATACAT | loxF8L | 3 | loxF8L | 4 | AAAAAAAA/TTTTTTTT | 7 |
| chr15 | 36635369 | ATTTATATGTGGA-TTTTTTTT-CAACTAAACTTAG | loxF8L | 3 | loxF8R | 4 | AAAAAAAA/TTTTTTTT | 7 |
| --- | | | | | | | | 0 |
| chr2 | 142366620 | CTAAATCTGTTGT-AAATTTTT-TCCTCAGTTTTAA | loxF8L | 3 | loxF8L | 3 | AAAAATTT/AAATTTTT | 6 |
| chr11 | 118512109 | AAAAATCAGTTAA-AAATTTTT-TTCACAGCTTTAT | loxF8L | 4 | loxF8L | 2 | AAAAATTT/AAATTTTT | 6 |
| chr7 | 22933854 | TTAATATTCTTTG-AAATTTTT-GACACAATGTTAG | loxF8R | 4 | loxF8R | 2 | AAAAATTT/AAATTTTT | 6 |
| chr4 | 16355536 | ATAAGGCTGTGAG-AAATTTTT-TCAAGTGATTTAT | loxF8L | 4 | loxF8L | 3 | AAAAATTT/AAATTTTT | 7 |
| --- | | | | | | | | 0 |
| chr3 | 3834003 | ATAAGAGTTTGTG-TCTGTTTT-TCCACAGTTTCAG | loxF8R | 3 | loxF8L | 3 | AAAACAGA/TCTGTTTT | 6 |
| chr21 | 27319736 | CTAAGAACTTGTG-TCTGTTTT-CCCACAATTTCAG | loxF8R | 3 | loxF8R | 3 | AAAACAGA/TCTGTTTT | 6 |
| chr4 | 114748104 | CTAAAACTGTGGA-AAAACAGA-CACAAGCTCTTAT | loxF8L | 2 | loxF8R | 4 | AAAACAGA/TCTGTTTT | 6 |
| chr5 | 110208288 | ATAAGAACTTGTG-TCTGTTTT-TCCACAGTTTCAG | loxF8R | 4 | loxF8L | 3 | AAAACAGA/TCTGTTTT | 7 |
| chr6 | 51052669 | CTGAAATTGTGGA-AAAACAGA-CACAAATTCTTAT | loxF8L | 4 | loxF8R | 3 | AAAACAGA/TCTGTTTT | 7 |
| chr12 | 81941552 | ATAAGAGCATGTG-TCTGTTTT-CCCACAATTTCAG | loxF8R | 4 | loxF8R | 3 | AAAACAGA/TCTGTTTT | 7 |
| chr11 | 59276729 | CTAAAATTGTGGA-AAAACAGA-CACATGCTCTTAT | loxF8L | 3 | loxF8R | 4 | AAAACAGA/TCTGTTTT | 7 |
| chr2 | 18878272 | CTGAAATTGTGGA-AAAACAGA-CACACGCTCTTAT | loxF8L | 4 | loxF8R | 3 | AAAACAGA/TCTGTTTT | 7 |
| chrX | 122610000 | CTGAAATTGTGGA-AAAACAGA-CACACGCTCTTAT | loxF8L | 4 | loxF8R | 3 | AAAACAGA/TCTGTTTT | 7 |
| chr8 | 81242550 | ATAAGAGCTTGTG-TCTGTTTT-CCCACAATTTCAG | loxF8R | 4 | loxF8R | 3 | AAAACAGA/TCTGTTTT | 7 |
| chr1 | 204235525 | CTGAAATTGTGTA-AAAACAGA-CACAAGCTCTTAT | loxF8R | 3 | loxF8R | 4 | AAAACAGA/TCTGTTTT | 7 |
| chrY | 21298597 | ATGAAATTGTGGA-AAAACAGA-CACAAGCTCTTAT | loxF8L | 3 | loxF8R | 4 | AAAACAGA/TCTGTTTT | 7 |
| chr1 | 105753791 | CTAAAATTGTGGA-AAAACAGA-CACAAGCTCTTAT | loxF8L | 3 | loxF8R | 4 | AAAACAGA/TCTGTTTT | 7 |
| chr6 | 102871503 | CTAAAATTGTGGA-AAAACAGA-CACAAGCTCTTAT | loxF8L | 3 | loxF8R | 4 | AAAACAGA/TCTGTTTT | 7 |
| chrX | 75909283 | CTAAAATTGTGGA-AAAACAGA-CACAAGCTCTTAT | loxF8L | 3 | loxF8R | 4 | AAAACAGA/TCTGTTTT | 7 |
| chr12 | 79499486 | ATAAGAGCTTGTG-TCTGTTTT-TCCACAGTTTCAG | loxF8R | 4 | loxF8L | 3 | AAAACAGA/TCTGTTTT | 7 |
| chr13 | 69504973 | ATAAGAGCTTGTG-TCTGTTTT-TCCACAGTTTCAG | loxF8R | 4 | loxF8L | 3 | AAAACAGA/TCTGTTTT | 7 |
| chr1 | 100568606 | CTGAAACTGTGGA-AAAACAGA-CACAAGCTCTTAT | loxF8L | 3 | loxF8R | 4 | AAAACAGA/TCTGTTTT | 7 |
| chr6 | 121637428 | ATAAGAGCTTGTG-TCTGTTTT-TCCACAGTTTCAG | loxF8R | 4 | loxF8L | 3 | AAAACAGA/TCTGTTTT | 7 |
| chr12 | 83720350 | CAGAAATTGTGGG-AAAACAGA-CACAAACTCTTAT | loxF8R | 4 | loxF8R | 3 | AAAACAGA/TCTGTTTT | 7 |
| chr4 | 107730431 | ATAAGAGTTTGTG-TCTGTTTT-TCCACAATTATAG | loxF8R | 3 | loxF8L | 4 | AAAACAGA/TCTGTTTT | 7 |
| chr14 | 39894115 | CTGAAATTGTGGA-AAAACAGA-CACAAACTCTTAT | loxF8L | 4 | loxF8R | 3 | AAAACAGA/TCTGTTTT | 7 |
| chr21 | 15577396 | CTGAAATTGTGGA-AAAACAGA-CACAAACTCTTAT | loxF8L | 4 | loxF8R | 3 | AAAACAGA/TCTGTTTT | 7 |
| chr12 | 14299027 | ATAAGAGTTTGTG-TCTGTTTT-TGCACAATTTCAG | loxF8R | 3 | loxF8R | 4 | AAAACAGA/TCTGTTTT | 7 |
| chr19 | 22024478 | ATAAGAGTTTGTG-TCTGTTTT-TGCACAATTTCAG | loxF8R | 3 | loxF8R | 4 | AAAACAGA/TCTGTTTT | 7 |
| chr12 | 84402619 | CTGAAATTGTGGA-AAAACAGA-CACAAGATCTTAT | loxF8L | 4 | loxF8R | 3 | AAAACAGA/TCTGTTTT | 7 |
| chr3 | 174885246 | ATAAGATCTTGTG-TCTGTTTT-TCCACAATTTCAG | loxF8R | 3 | loxF8L | 4 | AAAACAGA/TCTGTTTT | 7 |
| chr3 | 118363580 | CTAAGAGCTTGTG-TCTGTTTT-TCCACAATTTCAG | loxF8R | 3 | loxF8L | 4 | AAAACAGA/TCTGTTTT | 7 |
| chrX | 69870812 | GTAAGAGCTTGTG-TCTGTTTT-TCCACAGTTTCAG | loxF8R | 4 | loxF8L | 3 | AAAACAGA/TCTGTTTT | 7 |
| --- | | | | | | | | 0 |
| chr5 | 154273825 | TTAAAGCAGTGGA-ACACTTTT-CACATAATCTTAT | loxF8L | 3 | loxF8R | 2 | AAAAGTGT/ACACTTTT | 5 |
| chr1 | 180102527 | CTATGATTTTGTT-ACACTTTT-CACCCATTATTGG | loxF8R | 3 | loxF8R | 4 | AAAAGTGT/ACACTTTT | 7 |
| --- | | | | | | | | 0 |
| chr12 | 62318076 | ATTAATATGTGGA-AAAATAAG-CACACACTGTCAG | loxF8L | 2 | loxF8R | 3 | AAAATAAG/CTTATTTT | 5 |
| chr12 | 81138221 | AAAAATATGAGGA-CTTATTTT-TCTGCAAATTAAT | loxF8L | 3 | loxF8L | 4 | AAAATAAG/CTTATTTT | 7 |
| chr3 | 108071284 | TTAAAACTGTGGC-CTTATTTT-TACATGTATTTAT | loxF8L | 3 | loxF8L | 4 | AAAATAAG/CTTATTTT | 7 |
| chr17 | 58154549 | CCAAGATTATCTG-CTTATTTT-TCTTCACATTTAC | loxF8R | 3 | loxF8L | 4 | AAAATAAG/CTTATTTT | 7 |
| --- | | | | | | | | 0 |
| chr2 | 178145023 | CAAAGATTTTGAG-TGAGGTTT-ACCATAGATTTTT | loxF8R | 3 | loxF8L | 3 | AAACCTCA/TGAGGTTT | 6 |
| chr7 | 115653227 | TTTGGATTGTGTT-TGAGGTTT-TCTCCAGATTTAT | loxF8R | 4 | loxF8L | 2 | AAACCTCA/TGAGGTTT | 6 |
| --- | | | | | | | | 0 |
| chr2 | 182071648 | ATAAATCATTTGA-AATTCTTT-TGCACAGATTTGT | loxF8L | 3 | loxF8L | 2 | AAAGAATT/AATTCTTT | 5 |
| chr7 | 127437385 | ATACCTCTGGGGA-AATTCTTT-TCTGCATAGTTAT | loxF8L | 3 | loxF8L | 4 | AAAGAATT/AATTCTTT | 7 |
| --- | | | | | | | | 0 |
| chr2 | 84187684 | ATAAATATTTGTA-AAAGATAA-CACACTATGTTAG | loxF8L | 3 | loxF8R | 2 | AAAGATAA/TTATCTTT | 5 |
| chr2 | 103032336 | ATAATTTTGTTTA-TTATCTTT-TCCCCAGCTTTTT | loxF8L | 4 | loxF8L | 3 | AAAGATAA/TTATCTTT | 7 |
| chr4 | 82025124 | ATATATTTGTCCA-TTATCTTT-TCAACAGATCTTT | loxF8L | 4 | loxF8L | 3 | AAAGATAA/TTATCTTT | 7 |
| --- | | | | | | | | 0 |
| chr14 | 89844369 | CTAAAATTGAATG-AAATAATT-CATACAATTTTAC | loxF8R | 3 | loxF8R | 3 | AAATAATT/AATTATTT | 6 |
| chr9 | 28860097 | AGAAATATGTGGA-AAATAATT-AAGAAAATCTTAC | loxF8L | 2 | loxF8R | 4 | AAATAATT/AATTATTT | 6 |
| chr11 | 39624368 | TTTAATCAGTTGA-AAATAATT-TCCACATTTTTGT | loxF8L | 4 | loxF8L | 3 | AAATAATT/AATTATTT | 7 |
| chr12 | 96525232 | CTTTCATTGTGTG-AATTATTT-TTCACAGTCTTTG | loxF8R | 3 | loxF8R | 4 | AAATAATT/AATTATTT | 7 |
| --- | | | | | | | | 0 |
| chr13 | 67446048 | ATAACTTTGTGGA-ACATATTT-TTGACTGATTTAT | loxF8L | 2 | loxF8L | 3 | AAATATGT/ACATATTT | 5 |
| chr5 | 173060823 | AAACAGCTGTAGA-AAATATGT-TCTACATATTTTT | loxF8L | 4 | loxF8L | 3 | AAATATGT/ACATATTT | 7 |
| chr11 | 15483204 | AAAAATTCTTGGA-ACATATTT-TGCATAGATTTAG | loxF8L | 4 | loxF8L | 3 | AAATATGT/ACATATTT | 7 |
| chr4 | 158737942 | ATAAATCTATCTA-AAATATGT-TCAATATTTTTAT | loxF8L | 3 | loxF8L | 4 | AAATATGT/ACATATTT | 7 |

| | | | | | | | | | |
|---|---|---|---|---|---|---|---|---|---|
| --- | | | | | | | | | 0 |
| chr13 | 67806383 | ATATGCTTGTGTG-TGTATGTT-CACACATATTTAT | loxF8R | 3 | loxF8L | 3 | AACATACA/TGTATGTT | | 6 |
| chr5 | 118544370 | ATAAATATGTAGA-TGTATGTT-TCTATATATATAT | loxF8L | 2 | loxF8L | 4 | AACATACA/TGTATGTT | | 6 |
| chr2 | 8987794 | CCAATTTTGTTTG-AACATACA-CACACACTCTAAT | loxF8R | 4 | loxF8R | 3 | AACATACA/TGTATGTT | | 7 |
| --- | | | | | | | | | 0 |
| chr10 | 49590345 | TTAAGAGTGTGTT-GTTTAGTT-TACACATATTTAT | loxF8R | 3 | loxF8L | 2 | AACTAAAC/GTTTAGTT | HG8 | 5 |
| chr9 | 135831201 | ACAAATACGTGGA-AACTAAAC-AACACATTCTTAA | loxF8L | 3 | loxF8R | 3 | AACTAAAC/GTTTAGTT | | 6 |
| chr1 | 111302514 | TTAAGAGTGTGTT-GTTTAGTT-TCCACAAATTGGT | loxF8R | 3 | loxF8L | 3 | AACTAAAC/GTTTAGTT | | 6 |
| chr3 | 165673739 | TCAAGAGTGTGTT-GTTTAGTT-TCCACATATTTGT | loxF8R | 4 | loxF8L | 2 | AACTAAAC/GTTTAGTT | | 6 |
| chrX | 27566989 | TCAAGAGTGTGTT-GTTTAGTT-TCCACATATTTGT | loxF8R | 4 | loxF8L | 2 | AACTAAAC/GTTTAGTT | | 6 |
| chrX | 154866593 | ACAAATATGTGGA-AACTAAAC-AACACATTCTTGA | loxF8L | 2 | loxF8R | 4 | AACTAAAC/GTTTAGTT | HG9 | 6 |
| chr16 | 59893671 | TCAAGAGTGTGTT-GTTTAGTT-TCCACAAATTTGT | loxF8R | 4 | loxF8L | 2 | AACTAAAC/GTTTAGTT | | 6 |
| chr10 | 92401082 | ACAAATATTTGGA-AACTAAAC-AACACAGGTTTTT | loxF8L | 3 | loxF8L | 4 | AACTAAAC/GTTTAGTT | | 7 |
| chr21 | 44383728 | ACAAAGTTGTGGA-AACTAAAC-GACACTACCTTAC | loxF8L | 3 | loxF8R | 4 | AACTAAAC/GTTTAGTT | | 7 |
| chr13 | 107153030 | TCAAGAGTGTGTT-GTTTAGTT-TCCATATATTTGT | loxF8R | 4 | loxF8L | 3 | AACTAAAC/GTTTAGTT | | 7 |
| chrX | 114502800 | ACAAATATATGGA-AACTAAAC-AACACACTCTTGA | loxF8L | 3 | loxF8R | 4 | AACTAAAC/GTTTAGTT | | 7 |
| chr9 | 91050854 | ACAAATATGTAGA-AACTAAAC-AACACAATATTGA | loxF8L | 3 | loxF8R | 4 | AACTAAAC/GTTTAGTT | | 7 |
| chrX | 127041759 | ATACATGTGTGCA-AACTAAAC-AACACATTCCTAA | loxF8L | 3 | loxF8R | 4 | AACTAAAC/GTTTAGTT | | 7 |
| --- | | | | | | | | | 0 |
| chr6 | 120084236 | AAAAATATGAGGA-AAGTAAGT-CTCACAAGCATAG | loxF8L | 3 | loxF8R | 3 | AAGTAAGT/ACTTACTT | | 6 |
| chr2 | 31034862 | TTAGGAGTGTGTC-ACTTACTT-TCCACATATTTGT | loxF8R | 4 | loxF8L | 2 | AAGTAAGT/ACTTACTT | | 6 |
| --- | | | | | | | | | 0 |
| chr12 | 114453677 | ATAAATGTGTAGA-AATAAAAT-CAGACACCTAAG | loxF8L | 2 | loxF8R | 3 | AATAAAAT/ATTTTATT | | 5 |
| chr2 | 34929593 | CTTAGGTTGTCTG-ATTTTATT-TCTACACCTTTAT | loxF8R | 3 | loxF8L | 3 | AATAAAAT/ATTTTATT | | 6 |
| chr11 | 49017250 | AAAAAATTGTATG-AATAAAAT-CACAAAAGCCTAG | loxF8R | 4 | loxF8R | 3 | AATAAAAT/ATTTTATT | | 7 |
| chr11 | 89818694 | CTAGGCTTTTGTG-ATTTTATT-CATACAATTTTTT | loxF8R | 3 | loxF8R | 4 | AATAAAAT/ATTTTATT | | 7 |
| chr11 | 90025862 | AAAAAATTGTATG-AATAAAAT-CACAAAAGCCTAG | loxF8R | 4 | loxF8R | 3 | AATAAAAT/ATTTTATT | | 7 |
| chr12 | 107738157 | CAAAATCTGTGGT-ATTTTATT-ATCACAACCTGAG | loxF8L | 3 | loxF8R | 4 | AATAAAAT/ATTTTATT | | 7 |
| chr5 | 8350662 | CTCAGTCTGTGGA-ATTTTATT-CTGGCAATCCTAG | loxF8L | 3 | loxF8R | 4 | AATAAAAT/ATTTTATT | | 7 |
| --- | | | | | | | | | 0 |
| chr1 | 73784148 | CTAAGACTGCCTG-AATACAAA-CACAGAATATTAG | loxF8R | 3 | loxF8R | 2 | AATACAAA/TTTGTATT | | 5 |
| chr2 | 192225383 | CCAAGAGTGTTTG-AATACAAA-TCCGCATTTTTAT | loxF8R | 3 | loxF8L | 3 | AATACAAA/TTTGTATT | | 6 |
| chr14 | 83719715 | ATAAATTTGAGGG-AATACAAA-CAATCAAACTTAG | loxF8L | 3 | loxF8R | 3 | AATACAAA/TTTGTATT | | 6 |
| chr5 | 157074110 | GTAAAGCTAAGGA-TTTGTATT-TCCAGAAATTTTT | loxF8L | 4 | loxF8L | 3 | AATACAAA/TTTGTATT | | 7 |
| chr13 | 22459154 | ACAAATCTGTCTA-AATACAAA-TACACAAATCTGT | loxF8L | 3 | loxF8L | 4 | AATACAAA/TTTGTATT | | 7 |
| --- | | | | | | | | | 0 |
| chr20 | 22935066 | TTAAGATTGTATT-GTTTGATT-TCCACAAATTAAT | loxF8R | 3 | loxF8L | 2 | AATCAAAC/GTTTGATT | | 5 |
| chrX | 67848723 | ACAAATATGTGAA-AATCAAAC-AACACAGTCTTAA | loxF8L | 3 | loxF8R | 3 | AATCAAAC/GTTTGATT | | 6 |
| chr8 | 95279581 | TTAAGAGTGTGTT-GTTTGATT-TCCAAAAATTTGT | loxF8R | 3 | loxF8L | 3 | AATCAAAC/GTTTGATT | | 6 |
| chr3 | 135424905 | ACAAATATGTGGA-AATCAAAC-AACACAGTCCTAA | loxF8L | 2 | loxF8R | 4 | AATCAAAC/GTTTGATT | | 6 |
| chr3 | 157995757 | TTAAGAGTCTGTT-GTTTGATT-TCCACATATTTGT | loxF8R | 4 | loxF8L | 2 | AATCAAAC/GTTTGATT | | 6 |
| chr10 | 113863326 | ACAAATTTGTGGA-AATCAAAC-AGCACACTCTTAA | loxF8L | 2 | loxF8R | 4 | AATCAAAC/GTTTGATT | | 6 |
| chr11 | 86883842 | TTAGGAGTGTGTT-GTTTGATT-TCTACATATTTAT | loxF8R | 4 | loxF8L | 2 | AATCAAAC/GTTTGATT | | 6 |
| chr18 | 78843429 | TTAAGAGTATGTT-GTTTGATT-TTCACAAATTTTT | loxF8R | 4 | loxF8L | 3 | AATCAAAC/GTTTGATT | | 7 |
| chr4 | 139666968 | CTCACGTTTTGTG-GTTTGATT-CACACAATCTGTT | loxF8R | 4 | loxF8R | 3 | AATCAAAC/GTTTGATT | | 7 |
| chr15 | 73963846 | CTAAAAGTGTGTT-GTTTGATT-TCCAAACATTTGG | loxF8R | 3 | loxF8L | 4 | AATCAAAC/GTTTGATT | | 7 |
| --- | | | | | | | | | 0 |
| chr18 | 36225803 | TTAAGAGTGTGTT-GTTTCATT-TCCACATATTTGT | loxF8R | 3 | loxF8L | 2 | AATGAAAC/GTTTCATT | | 5 |
| chr19 | 44022644 | ACAAGTATGTGGA-AATGAAAC-AACACATTCTTAA | loxF8L | 3 | loxF8R | 3 | AATGAAAC/GTTTCATT | | 6 |
| chr4 | 80826919 | GCAACTCTGTGGA-AATGAAAC-AACACATTCTTAA | loxF8L | 3 | loxF8R | 3 | AATGAAAC/GTTTCATT | | 6 |
| chrX | 97490673 | TAAAGAGTGTGTT-GTTTCATT-TCCACATATTTGT | loxF8R | 4 | loxF8L | 2 | AATGAAAC/GTTTCATT | | 6 |
| chr3 | 82869094 | ACAAATATGTGAA-AATGAAAC-AACACACACTTAA | loxF8L | 3 | loxF8R | 4 | AATGAAAC/GTTTCATT | | 7 |
| chr10 | 32242115 | TCAAGAGTGTGTT-GTTTCATT-CCCACATATTTGT | loxF8R | 4 | loxF8L | 3 | AATGAAAC/GTTTCATT | | 7 |
| chr11 | 2505245 | TTAAGAGTATGTT-GTTTCATT-TCCATACATTTGT | loxF8R | 4 | loxF8L | 3 | AATGAAAC/GTTTCATT | | 7 |
| --- | | | | | | | | | 0 |
| chr17 | 82045792 | CTAAGAGTGCGTT-GTTACATT-TCCACAAATTTGT | loxF8R | 3 | loxF8L | 2 | AATGTAAC/GTTACATT | | 5 |
| chr5 | 161720803 | ATAAGAGTGTGTT-GTTACATT-TCCACAATTTTGT | loxF8R | 3 | loxF8L | 3 | AATGTAAC/GTTACATT | | 6 |
| chr11 | 100202415 | TTATGATTATGTG-GTTACATT-TACACTATATGAG | loxF8R | 3 | loxF8R | 4 | AATGTAAC/GTTACATT | | 7 |
| --- | | | | | | | | | 0 |
| chr7 | 86540611 | TTAAATATTTGGA-AATTAAGT-CACATAATTTTAA | loxF8L | 3 | loxF8R | 3 | AATTAAGT/ACTTAATT | | 6 |
| chr14 | 58932908 | CTAGGAGCGTGTT-ACTTAATT-TCCACACATTTGT | loxF8R | 4 | loxF8L | 2 | AATTAAGT/ACTTAATT | | 6 |
| --- | | | | | | | | | 0 |
| chr12 | 41753083 | GTAAGAGTGTGTT-GATTAATT-TTCATAAATTTAT | loxF8R | 3 | loxF8L | 3 | AATTAATC/GATTAATT | | 6 |
| chr8 | 42781064 | TTAAGAGTGTGTT-GATTAATT-TCCGCATATTCAT | loxF8R | 3 | loxF8L | 3 | AATTAATC/GATTAATT | | 6 |
| chr15 | 36148996 | AAAAATTTGTGGA-AATTAATC-AACACACCCTTAA | loxF8L | 2 | loxF8R | 4 | AATTAATC/GATTAATT | | 6 |
| chr4 | 89592968 | TTAGGAGTGTGTT-GATTAATT-TCCACATATCTGT | loxF8R | 4 | loxF8L | 3 | AATTAATC/GATTAATT | | 7 |
| --- | | | | | | | | | 0 |
| chr14 | 59047792 | TTAAGAGTGTGTT-AATTAATT-TCCACATATTTTT | loxF8R | 3 | loxF8L | 2 | AATTAATT/AATTAATT | | 5 |
| chr1 | 236324599 | ATAAATAGGTGGA-AATTAATT-AACACAGTCTTAA | loxF8L | 2 | loxF8R | 3 | AATTAATT/AATTAATT | | 5 |
| chr8 | 127966659 | CAATTTTTGTGTG-AATTAATT-TCAACAGACTTAA | loxF8R | 4 | loxF8L | 3 | AATTAATT/AATTAATT | | 7 |
| --- | | | | | | | | | 0 |
| chr5 | 125706394 | ACAAATATGTGGA-AATTATAC-AACCCATTCTTAG | loxF8L | 2 | loxF8R | 3 | AATTATAC/GTATAATT | | 5 |
| chrX | 31672165 | TTAAGAGTGTGTT-GTATAATT-TTCTCATATTTAT | loxF8R | 3 | loxF8L | 3 | AATTATAC/GTATAATT | | 6 |
| chr7 | 157126747 | TTAGGAGTGTGTT-GTATAATT-TCCACATATTTGT | loxF8R | 4 | loxF8L | 2 | AATTATAC/GTATAATT | | 6 |
| chr3 | 56489871 | AGAAGTCTCTGGC-GTATAATT-CACAGAATCTTAT | loxF8L | 4 | loxF8R | 2 | AATTATAC/GTATAATT | | 6 |
| chr7 | 45447673 | AAAAATATTTAGA-AATTATAC-CACACACTCTGAA | loxF8L | 4 | loxF8R | 3 | AATTATAC/GTATAATT | | 7 |
| --- | | | | | | | | | 0 |
| chr14 | 101644518 | AAAAATTTGTGGA-AATTGAAC-AACACACTCTTAA | loxF8L | 2 | loxF8R | 3 | AATTGAAC/GTTCAATT | | 5 |
| chrX | 35873621 | TTAAGAGTGTGTT-GTTCAATT-TCCGCATATTTGT | loxF8R | 3 | loxF8L | 3 | AATTGAAC/GTTCAATT | | 6 |
| chr8 | 40653114 | AAAAATAAGTAGA-AATTGAAC-CACACACTCTTAA | loxF8L | 4 | loxF8R | 2 | AATTGAAC/GTTCAATT | | 6 |
| chr7 | 123884297 | ACAAATATGTGAA-AATTGAAC-AACACACTCCTAA | loxF8L | 3 | loxF8R | 4 | AATTGAAC/GTTCAATT | | 7 |
| chr1 | 68832402 | ACAAATTTGTTGA-AATTGAAC-ATCACACTCTTAA | loxF8L | 3 | loxF8R | 4 | AATTGAAC/GTTCAATT | | 7 |
| --- | | | | | | | | | 0 |
| chr3 | 59845542 | TTAAGATTGTGTT-ATTCAATT-TTCACAAATTTTT | loxF8R | 2 | loxF8L | 3 | AATTGAAT/ATTCAATT | | 5 |
| chrX | 110326378 | TTAGGAGTGTGTT-ATTCAATT-TCCACATATTTGT | loxF8R | 4 | loxF8L | 2 | AATTGAAT/ATTCAATT | | 6 |
| chr4 | 22254605 | ATAAATATGTGAA-AATTGAAT-AATACAGTCTTAA | loxF8L | 2 | loxF8R | 4 | AATTGAAT/ATTCAATT | | 6 |
| chr10 | 51255169 | AATAATTTGTGGA-ATTCAATT-TCTACATATGTTT | loxF8L | 3 | loxF8L | 4 | AATTGAAT/ATTCAATT | | 7 |

| | | | | | | | | | |
|---|---|---|---|---|---|---|---|---|---|
| chr4 | 104049337 | ATAAATATGTTGC-ATTCAATT-TTATCATATTTAT | loxF8L | 3 | loxF8L | 4 | AATTGAAT/ATTCAATT | | 7 |
| --- | | | | | | | | | 0 |
| chr5 | 84649996 | TTAAGATTCTGTG-AATTTAAA-TCCACACAGTAAT | loxF8R | 2 | loxF8L | 3 | AATTTAAA/TTTAAATT | | 5 |
| chr3 | 14220425 | TTAAGAGTGTGTT-TTTAAATT-TCCACATACTTGT | loxF8R | 3 | loxF8L | 3 | AATTTAAA/TTTAAATT | | 6 |
| chr8 | 69711899 | CTAATATTATGAG-TTTAAATT-TGCACATATATAT | loxF8R | 3 | loxF8L | 3 | AATTTAAA/TTTAAATT | | 6 |
| chr13 | 42436876 | TTAAGGTGGTGTT-TTTAAATT-TCCACTTATTTGT | loxF8R | 4 | loxF8L | 3 | AATTTAAA/TTTAAATT | | 7 |
| chr11 | 57041288 | ACGAATATGTGGA-AATTTAAA-AACACACTCCTAA | loxF8L | 3 | loxF8R | 4 | AATTTAAA/TTTAAATT | | 7 |
| chr2 | 46777545 | ATAAATGAATGGA-TTTAAATT-TCCACATGTTGCT | loxF8L | 3 | loxF8L | 4 | AATTTAAA/TTTAAATT | | 7 |
| chr9 | 97867797 | TTAAGAATGAGTT-TTTAAATT-TCTATAGATTCAT | loxF8R | 4 | loxF8L | 3 | AATTTAAA/TTTAAATT | | 7 |
| --- | | | | | | | | | 0 |
| chr12 | 60180283 | CTACGGTTGTGTA-ACAAATTA-TCATCAGATTTAG | loxF8R | 3 | loxF8L | 3 | ACAAATTA/TAATTTGT | | 6 |
| chr10 | 11434104 | CTAAGTTTGGGGG-TAATTTGT-CACACAGCCATAG | loxF8R | 3 | loxF8R | 3 | ACAAATTA/TAATTTGT | HG1R | 6 |
| chr1 | 151245902 | CTATTATTGTGTA-ACAAATTA-CCCCCAAACTTAG | loxF8R | 3 | loxF8R | 3 | ACAAATTA/TAATTTGT | HG2R | 6 |
| chr7 | 95687255 | TTATGCTTTTGTG-TAATTTGT-CCCACAGATATAT | loxF8R | 4 | loxF8L | 2 | ACAAATTA/TAATTTGT | | 6 |
| chr2 | 80566133 | ATATTTCTGTGTA-ACAAATTA-CCCGCAAATTTAA | loxF8L | 3 | loxF8L | 4 | ACAAATTA/TAATTTGT | | 7 |
| chr3 | 134591944 | CTACGGCTGTGTA-ACAAATTA-CCCCCAGATTTAG | loxF8R | 4 | loxF8L | 3 | ACAAATTA/TAATTTGT | | 7 |
| chr22 | 50596205 | CTAAATTTGAGGA-TAATTTGT-TACACAGTAATAG | loxF8L | 3 | loxF8R | 4 | ACAAATTA/TAATTTGT | | 7 |
| chr12 | 128772604 | CTAAATTTGGGAG-TAATTTGT-TACACAATAATAG | loxF8R | 4 | loxF8R | 3 | ACAAATTA/TAATTTGT | | 7 |
| chr1 | 160296685 | CTAAATTTGGGGA-TAATTTGT-TACACAGAAATAG | loxF8L | 3 | loxF8L | 4 | ACAAATTA/TAATTTGT | | 7 |
| chr5 | 58869402 | CTAATTTTGTGGA-TAATTTGT-TACACAGAAATAG | loxF8L | 3 | loxF8L | 4 | ACAAATTA/TAATTTGT | | 7 |
| chr16 | 22953577 | CTCATATTGTTTG-TAATTTGT-TCCCAATATTTAA | loxF8R | 3 | loxF8L | 4 | ACAAATTA/TAATTTGT | | 7 |
| --- | | | | | | | | | 0 |
| chr5 | 13471175 | AAACATCTGTGTA-GTAATTGT-CACTTAATCTTAG | loxF8L | 3 | loxF8R | 2 | ACAATTAC/GTAATTGT | | 5 |
| chr3 | 79877872 | ATATATATATGGA-ACAATTAC-AACAAAATGTTAA | loxF8L | 3 | loxF8R | 4 | ACAATTAC/GTAATTGT | | 7 |
| --- | | | | | | | | | 0 |
| chr21 | 21214825 | ATATATATGTGTA-TATATGGT-TGAACAGAATTAT | loxF8L | 3 | loxF8L | 3 | ACCATATA/TATATGGT | | 6 |
| chr2 | 205027855 | CAAAGATTAGTTG-ACCATATA-TGCACAGGTTTAT | loxF8R | 4 | loxF8L | 2 | ACCATATA/TATATGGT | | 6 |
| chr18 | 36223381 | CAAAGATCATGTA-ACCATATA-TGCATGGATTTAT | loxF8R | 4 | loxF8L | 3 | ACCATATA/TATATGGT | | 7 |
| chr1 | 170957462 | CAAAGATTACTTG-ACCATATA-TTAATAGATTTAT | loxF8R | 4 | loxF8L | 3 | ACCATATA/TATATGGT | | 7 |
| chrX | 109511978 | ATAAATCTTCACA-TATATGGT-CAAAGAATCTTCG | loxF8L | 4 | loxF8R | 3 | ACCATATA/TATATGGT | | 7 |
| chr10 | 64660987 | CGAAGATTATTTG-ACCATATA-TGCATGGATCTAT | loxF8R | 3 | loxF8L | 4 | ACCATATA/TATATGGT | | 7 |
| --- | | | | | | | | | 0 |
| chrX | 93862135 | ATAAATCTGTATT-TTTAAAGT-GCCACAGATTAAT | loxF8L | 3 | loxF8L | 2 | ACTTTAAA/TTTAAAGT | | 5 |
| chr8 | 54117888 | TTAAGATTGTGTT-TTTAAAGT-TCCATATTTTTGT | loxF8R | 2 | loxF8L | 4 | ACTTTAAA/TTTAAAGT | | 6 |
| --- | | | | | | | | | 0 |
| chr5 | 101773510 | ATAAATATGTGAA-TTTATTCT-TCCCCAGAGTTAA | loxF8L | 2 | loxF8L | 3 | AGAATAAA/TTTATTCT | | 5 |
| chr12 | 88618030 | ATAAATATATGAA-AGAATAAA-TGGACAGATTTTG | loxF8L | 3 | loxF8L | 4 | AGAATAAA/TTTATTCT | | 7 |
| chr5 | 62618561 | ATCAAGGGGTGGA-AGAATAAA-CATAAAATATTAG | loxF8L | 4 | loxF8R | 3 | AGAATAAA/TTTATTCT | | 7 |
| --- | | | | | | | | | 0 |
| chrX | 134342277 | ATAAAACTGTTGA-AGAGCAAA-GACAAAATCTTAA | loxF8L | 2 | loxF8R | 3 | AGAGCAAA/TTTGCTCT | | 5 |
| chr13 | 109621698 | ATGGATATGTGGA-TTTGCTCT-TCCATTGATTAGT | loxF8L | 3 | loxF8L | 4 | AGAGCAAA/TTTGCTCT | | 7 |
| chr8 | 63653404 | CAAAGATACTGTG-AGAGCAAA-TACATAAATTTCT | loxF8R | 3 | loxF8L | 4 | AGAGCAAA/TTTGCTCT | | 7 |
| --- | | | | | | | | | 0 |
| chr17 | 60539534 | AAAAGTCTGTGGT-AGAGTTTG-CACAGAATCTTAT | loxF8L | 3 | loxF8R | 2 | AGAGTTTG/CAAACTCT | | 5 |
| chr14 | 78513970 | TTTAATCTGTGGC-CAAACTCT-TGCTCAGATCTAG | loxF8L | 3 | loxF8L | 4 | AGAGTTTG/CAAACTCT | | 7 |
| --- | | | | | | | | | 0 |
| chrY | 11338463 | CTGGGATGGTGTG-AGCCACTG-TGCCCAGACTTAT | loxF8R | 3 | loxF8L | 3 | AGCCACTG/CAGTGGCT | | 6 |
| chr14 | 76160703 | TGAAGACTGGGTG-CAGTGGCT-CAGACATTCTTAG | loxF8R | 4 | loxF8R | 2 | AGCCACTG/CAGTGGCT | | 6 |
| chr22 | 16426812 | CTGGGATGGTGTG-AGCCACTG-TGCTCAGCCTTAT | loxF8R | 3 | loxF8L | 4 | AGCCACTG/CAGTGGCT | | 7 |
| chr10 | 38638860 | CTGGGATGGTGTG-AGCCACTG-TGCCCAGCCTTAT | loxF8R | 3 | loxF8L | 4 | AGCCACTG/CAGTGGCT | | 7 |
| chr10 | 42278897 | ATAAGGCTGGGCA-CAGTGGCT-CACACCATCCCAG | loxF8L | 4 | loxF8R | 3 | AGCCACTG/CAGTGGCT | | 7 |
| chr16 | 32438262 | ATAAGGCTGGGCA-CAGTGGCT-CACACCATCCCAG | loxF8L | 4 | loxF8R | 3 | AGCCACTG/CAGTGGCT | | 7 |
| chr4 | 102899006 | CTGGGATGGTGTG-AGCCACTG-TGCCCAGCCTTAT | loxF8R | 3 | loxF8L | 4 | AGCCACTG/CAGTGGCT | | 7 |
| chr9 | 21387739 | CTAGGCTGGGGTG-CAGTGGCT-CCATCAATCTTAG | loxF8R | 4 | loxF8R | 3 | AGCCACTG/CAGTGGCT | | 7 |
| --- | | | | | | | | | 0 |
| chr3 | 143939194 | ATAAATATGTGGG-AGCTAAAC-CACATAATCTTGA | loxF8L | 2 | loxF8R | 3 | AGCTAAAC/GTTTAGCT | | 5 |
| chr6 | 6426489 | ACAAATCTGTAAG-AGCTAAAC-AACACAGTCTTAA | loxF8L | 4 | loxF8R | 3 | AGCTAAAC/GTTTAGCT | | 7 |
| --- | | | | | | | | | 0 |
| chr11 | 19017963 | CTAACATTTTGTA-AGGAAACT-GCCAGAGATTTAG | loxF8R | 3 | loxF8L | 3 | AGGAAACT/AGTTTCCT | | 6 |
| chr1 | 198660191 | CTAACATTGTATG-AGTTTCCT-TTCACATACTTTT | loxF8R | 2 | loxF8L | 4 | AGGAAACT/AGTTTCCT | | 6 |
| --- | | | | | | | | | 0 |
| chrX | 134349098 | AAAAATATGTGGG-AGTTAAAA-ACCCCAGATATAT | loxF8L | 3 | loxF8L | 3 | AGTTAAAA/TTTTAACT | | 6 |
| chr20 | 11219859 | CAAAGATTGTGTC-TTTTAACT-TACACACATTAAC | loxF8R | 2 | loxF8L | 4 | AGTTAAAA/TTTTAACT | | 6 |
| chr1 | 234009027 | CAAAGTTTGTTTG-TTTTAACT-TCAACATAGTTTT | loxF8R | 3 | loxF8L | 4 | AGTTAAAA/TTTTAACT | | 7 |
| chr7 | 32508424 | TTAATATTGTGTT-TTTTAACT-CACTCCAAATTAG | loxF8R | 3 | loxF8R | 4 | AGTTAAAA/TTTTAACT | | 7 |
| --- | | | | | | | | | 0 |
| chr20 | 20347111 | ACAAATTTGTGGA-AGTTAAAT-AACACATTCTTAA | loxF8L | 2 | loxF8R | 3 | AGTTAAAT/ATTTAACT | | 5 |
| chr1 | 232125404 | TTAAGAGTGTGTC-ATTTAACT-TGCACATATTTGT | loxF8R | 3 | loxF8L | 3 | AGTTAAAT/ATTTAACT | | 6 |
| chr7 | 25182806 | CTTAGCTTTTGTG-ATTTAACT-CACTCCATCTCAG | loxF8R | 3 | loxF8R | 3 | AGTTAAAT/ATTTAACT | | 6 |
| chr5 | 29720948 | TTAATAATGTGTC-ATTTAACT-TCCACATATTTGT | loxF8R | 4 | loxF8L | 2 | AGTTAAAT/ATTTAACT | | 6 |
| --- | | | | | | | | | 0 |
| chrX | 51986752 | TTAAGACTGTGTG-GCTTAACT-TCTACATATTTGT | loxF8R | 2 | loxF8L | 3 | AGTTAAGC/GCTTAACT | | 5 |
| chr5 | 105278878 | CAAAGAATGGGTG-GCTTAACT-AACACGAATTTAT | loxF8R | 3 | loxF8L | 4 | AGTTAAGC/GCTTAACT | | 7 |
| --- | | | | | | | | | 0 |
| chr1 | 176772321 | CTAAGATTTTGTT-TTTGAACT-AACAAAATCCTAG | loxF8R | 2 | loxF8R | 3 | AGTTCAAA/TTTGAACT | | 5 |
| chr6 | 137859314 | ATAATTTTCTCGA-AGTTCAAA-GCCAGAGATGTAT | loxF8L | 4 | loxF8L | 3 | AGTTCAAA/TTTGAACT | | 7 |
| --- | | | | | | | | | 0 |
| chr1 | 163234689 | CTATTATTGTGTA-ATAAAACA-CACAAAAACTTAG | loxF8R | 3 | loxF8R | 2 | ATAAAACA/TGTTTTAT | | 5 |
| chr13 | 44286379 | GGATATCTTTGGA-ATAAAACA-TCCACTGTTTTGT | loxF8L | 4 | loxF8L | 3 | ATAAAACA/TGTTTTAT | | 7 |
| chr6 | 153744975 | ATATATTTTTGCA-ATAAAACA-TTTACTGATTTAT | loxF8L | 4 | loxF8L | 3 | ATAAAACA/TGTTTTAT | | 7 |
| --- | | | | | | | | | 0 |
| chr10 | 51855280 | CTGAGATAGTGTG-TTTTATAT-CAGACAATCTGAT | loxF8R | 2 | loxF8R | 3 | ATATAAAA/TTTTATAT | | 5 |
| chr15 | 48526247 | ATCAATATCTGGA-ATATAAAA-AAAGAATCTCAG | loxF8L | 3 | loxF8R | 4 | ATATAAAA/TTTTATAT | | 7 |
| --- | | | | | | | | | 0 |
| chr14 | 55030250 | ATAAAGCAGTGAA-GAAAATAT-AATACAATCTTAT | loxF8L | 3 | loxF8R | 3 | ATATTTTC/GAAAATAT | | 6 |
| chr13 | 90061426 | ATAAATATATGGA-ATATTTTC-ACAACAGATTCAA | loxF8L | 2 | loxF8L | 4 | ATATTTTC/GAAAATAT | | 6 |
| chr3 | 155922421 | AAATGACTGTGGA-GAAAATAT-TCCACAGATATTC | loxF8L | 4 | loxF8L | 3 | ATATTTTC/GAAAATAT | | 7 |

| | | | | | | | | |
|---|---|---|---|---|---|---|---|---|
| --- | | | | | | | | 0 |
| chr19 | 34714672 | CAAAGCCTGTGTG-ATCCAGCA-AACACAATATTAG | loxF8R | 3 | loxF8R | 2 | ATCCAGCA/TGCTGGAT | 5 |
| chr9 | 6888571 | AAAACTCAGTAGA-TGCTGGAT-CCCACATTCTTTG | loxF8L | 4 | loxF8R | 3 | ATCCAGCA/TGCTGGAT | 7 |
| --- | | | | | | | | 0 |
| chr22 | 18808956 | CTAAGCTTCTGTT-TCATTCAT-TCCACAAATATAT | loxF8R | 3 | loxF8L | 2 | ATGAATGA/TCATTCAT | 5 |
| chr22 | 21243411 | CTAAGCTTCTGTT-TCATTCAT-TCCACAAATATAT | loxF8R | 3 | loxF8L | 2 | ATGAATGA/TCATTCAT | 5 |
| chr1 | 116630565 | CTAAGCTTCTGTT-TCATTCAT-TCCAAAAATATAT | loxF8R | 3 | loxF8L | 3 | ATGAATGA/TCATTCAT | 6 |
| --- | | | | | | | | 0 |
| chr4 | 61969353 | CTAAACTTGTGAG-ATGTAGGC-TCCACAATTGTAT | loxF8R | 3 | loxF8L | 3 | ATGTAGGC/GCCTACAT | 6 |
| chr5 | 106640116 | CAAAGATTGTGAG-GCCTACAT-CTCACAGATGTCT | loxF8R | 2 | loxF8L | 4 | ATGTAGGC/GCCTACAT | 6 |
| --- | | | | | | | | 0 |
| chr1 | 166658748 | ATGATTATGTGGA-TCCTTAAT-TTCACAAATTTCT | loxF8L | 3 | loxF8L | 3 | ATTAAGGA/TCCTTAAT | 6 |
| chr18 | 9442919 | GTAAATCTGGGGA-TCCTTAAT-TGCTCATATTTAC | loxF8L | 2 | loxF8L | 4 | ATTAAGGA/TCCTTAAT | 6 |
| chr7 | 24948647 | AAAAATCTCAGGA-TCCTTAAT-TCTACATCTTTAC | loxF8L | 3 | loxF8L | 4 | ATTAAGGA/TCCTTAAT | 7 |
| chr16 | 22902766 | TTTAGTTTGTGTT-TCCTTAAT-TTTAGAGATTTAT | loxF8R | 4 | loxF8L | 3 | ATTAAGGA/TCCTTAAT | 7 |
| --- | | | | | | | | 0 |
| chr9 | 15954443 | TTAAGAGTGTGTT-GTTTAAAT-TCCACAATTTTGT | loxF8R | 3 | loxF8L | 3 | ATTTAAAC/GTTTAAAT | 6 |
| chr6 | 27223649 | CACAGATTGTGTG-ATTTAAAC-AACAGACATTTAT | loxF8R | 2 | loxF8L | 4 | ATTTAAAC/GTTTAAAT | 6 |
| chr2 | 213761137 | ACAAAACTTTGGA-ATTTAAAC-AATACACTCTTAA | loxF8L | 3 | loxF8R | 4 | ATTTAAAC/GTTTAAAT | 7 |
| chr10 | 32632705 | TTAGGAGTGTGTT-GTTTAAAT-TCCACTTATTTGT | loxF8R | 4 | loxF8L | 3 | ATTTAAAC/GTTTAAAT | 7 |
| chr6 | 43731478 | AGAACTATGTAGA-ATTTAAAC-AACACACTCTTAA | loxF8L | 4 | loxF8R | 3 | ATTTAAAC/GTTTAAAT | 7 |
| chr7 | 136515092 | ATAAATTTCTGTC-ATTTAAAC-CACTCAATCCAAG | loxF8L | 4 | loxF8R | 3 | ATTTAAAC/GTTTAAAT | 7 |
| chr12 | 18476931 | CACAGATTGGGTG-ATTTAAAC-AACAGAAATTTAT | loxF8R | 3 | loxF8L | 4 | ATTTAAAC/GTTTAAAT | 7 |
| chr6 | 81984339 | ATAAATTTCTGTT-GTTTAAAT-CACTCAATCTATG | loxF8L | 4 | loxF8R | 3 | ATTTAAAC/GTTTAAAT | 7 |
| chr9 | 6348634 | TTAAATTTGTAGA-ATTTAAAC-AACATACTCTTAA | loxF8L | 3 | loxF8R | 4 | ATTTAAAC/GTTTAAAT | 7 |
| --- | | | | | | | | 0 |
| chr4 | 34503669 | CAAAGACAGTGTG-GCTTAAAT-AACACAATTTTAT | loxF8R | 3 | loxF8R | 3 | ATTTAAGC/GCTTAAAT | 6 |
| chr9 | 70979716 | CAAAGATTGGGTG-GCTTAAAT-AACAGAAATTTAT | loxF8R | 2 | loxF8L | 4 | ATTTAAGC/GCTTAAAT | 6 |
| chr18 | 57105704 | CATAAATTGGGTG-GCTTAAAT-ACCAGAGATTTCT | loxF8R | 4 | loxF8L | 3 | ATTTAAGC/GCTTAAAT | 7 |
| chr16 | 5357995 | CTCAGACTGGGTG-GCTTAAAT-GACACAACTTTAT | loxF8R | 3 | loxF8L | 4 | ATTTAAGC/GCTTAAAT | 7 |
| chr5 | 16548858 | ATAAATATCTGTT-ATTTAAGC-CACCCAATCTATG | loxF8L | 4 | loxF8R | 3 | ATTTAAGC/GCTTAAAT | 7 |
| chr20 | 59187497 | ATAAATATTTGTC-ATTTAAGC-CACCCAATCTAGG | loxF8L | 4 | loxF8R | 3 | ATTTAAGC/GCTTAAAT | 7 |
| chr11 | 10603927 | ATAAATGTTTGCC-ATTTAAGC-CACCCAGTCTAAG | loxF8L | 4 | loxF8R | 3 | ATTTAAGC/GCTTAAAT | 7 |
| chr20 | 23176825 | CAAAGACTGGGTG-GCTTAAAT-AACAAACATTTAT | loxF8R | 3 | loxF8L | 4 | ATTTAAGC/GCTTAAAT | 7 |
| chr3 | 94399418 | CACAGACTGGGTG-GCTTAAAT-TACAGAAATTTAT | loxF8R | 4 | loxF8L | 3 | ATTTAAGC/GCTTAAAT | 7 |
| chr5 | 120450336 | ATAAATTTCTGTT-ATTTAAGC-CACACAGTCTACG | loxF8L | 4 | loxF8R | 3 | ATTTAAGC/GCTTAAAT | 7 |
| chr6 | 143589051 | CACAGATTGGGTG-GCTTAAAT-AACAGAAATTTAT | loxF8R | 3 | loxF8L | 4 | ATTTAAGC/GCTTAAAT | 7 |
| chr9 | 84176825 | CTCAGACTGTGGG-GCTTAAAT-AACAGAAATTTAT | loxF8R | 3 | loxF8L | 4 | ATTTAAGC/GCTTAAAT | 7 |
| chr17 | 19188959 | ATAAATTTGTGTT-ATTTAAGC-CACAAAGTCTGTG | loxF8L | 3 | loxF8R | 4 | ATTTAAGC/GCTTAAAT | 7 |
| chr17 | 19111673 | CACAGACTTTGTG-GCTTAAAT-AACACAATTTTAT | loxF8R | 4 | loxF8L | 3 | ATTTAAGC/GCTTAAAT | 7 |
| chr17 | 19062858 | ATAAATTTGTGTT-ATTTAAGC-CACAAAGTCTGTG | loxF8L | 3 | loxF8R | 4 | ATTTAAGC/GCTTAAAT | 7 |
| chr3 | 115310429 | CACAGAGTGGGTG-GCTTAAAT-AACACAAATTTAT | loxF8R | 4 | loxF8L | 3 | ATTTAAGC/GCTTAAAT | 7 |
| --- | | | | | | | | 0 |
| chr8 | 49455839 | ATAAATCTGTTCT-TTATAAAT-TACCCAGTCTTAG | loxF8L | 3 | loxF8R | 3 | ATTTATAA/TTATAAAT | 6 |
| chr14 | 104079138 | CTGAGACTGGGTA-ATTTATAA-ACGACAGATTTAT | loxF8R | 4 | loxF8L | 2 | ATTTATAA/TTATAAAT | 6 |
| chr2 | 74904876 | ATAAATCTGTGAC-TTATAAAT-AGCAGAAATTTAT | loxF8L | 2 | loxF8L | 4 | ATTTATAA/TTATAAAT | 6 |
| chr10 | 121816701 | AAAAATCTGTTGT-TTATAAAT-TACCCAGTCTCAG | loxF8L | 3 | loxF8R | 4 | ATTTATAA/TTATAAAT | 7 |
| chr1 | 102546822 | CTGAGATTGTATA-ATTTATAA-TACACAGAAATTT | loxF8R | 3 | loxF8L | 4 | ATTTATAA/TTATAAAT | 7 |
| chr3 | 106622464 | CTAAGACTGGGTA-ATTTATAA-TCAACAGAAATTT | loxF8R | 3 | loxF8L | 4 | ATTTATAA/TTATAAAT | 7 |
| chr2 | 67417248 | CTGAGACTGGGTA-ATTTATAA-TGAACAGATTTGT | loxF8R | 4 | loxF8L | 3 | ATTTATAA/TTATAAAT | 7 |
| chr3 | 114286976 | CTTAGATTGGGTA-ATTTATAA-ACAACAGAATTCT | loxF8R | 3 | loxF8L | 4 | ATTTATAA/TTATAAAT | 7 |
| chr1 | 43720429 | CTAAGACTGGATA-ATTTATAA-AAAACAGATTTAT | loxF8R | 4 | loxF8L | 3 | ATTTATAA/TTATAAAT | 7 |
| chr11 | 112906474 | ATAAATTTGTTGT-TTATAAAT-TACCCAGTCTAAG | loxF8L | 3 | loxF8R | 4 | ATTTATAA/TTATAAAT | 7 |
| chr1 | 187361754 | CACAGACTGGGTG-ATTTATAA-TAAACAAATTTAT | loxF8R | 4 | loxF8L | 3 | ATTTATAA/TTATAAAT | 7 |
| chr11 | 130552325 | ATATTTCTGTTGT-TTATAAAT-TACCCAATCTAAG | loxF8L | 4 | loxF8R | 3 | ATTTATAA/TTATAAAT | 7 |
| --- | | | | | | | | 0 |
| chr4 | 116719938 | GTATATCTCTGGA-TATAAAAT-TTCACAGAGTTAT | loxF8L | 3 | loxF8L | 2 | ATTTTATA/TATAAAAT | 5 |
| chr4 | 44346337 | AAAAATCTGTGAT-ATTTTATA-TCCTCACATTCTT | loxF8L | 3 | loxF8L | 4 | ATTTTATA/TATAAAAT | 7 |
| chr9 | 113750980 | ATAAATTTCTGTG-TATAAAAT-TACTCAGTCTTAG | loxF8L | 4 | loxF8R | 3 | ATTTTATA/TATAAAAT | 7 |
| chr13 | 81836196 | CTAAGTGTGTTTC-ATTTTATA-AGCACAGATTGAT | loxF8R | 4 | loxF8L | 3 | ATTTTATA/TATAAAAT | 7 |
| --- | | | | | | | | 0 |
| chr7 | 117288822 | CTAAGTTTGTATT-TTAAAAAT-AACACAGATTTAT | loxF8R | 3 | loxF8L | 2 | ATTTTTAA/TTAAAAAT | 5 |
| chr8 | 66828880 | ATAGCCCTGAGGA-ATTTTTAA-TCCACAGATTGTT | loxF8L | 4 | loxF8L | 2 | ATTTTTAA/TTAAAAAT | 6 |
| chr10 | 130156108 | CTTAGACTGAGTA-ATTTTTAA-GCAACAGATTTAT | loxF8R | 4 | loxF8L | 2 | ATTTTTAA/TTAAAAAT | 6 |
| chr13 | 83204373 | AAACATATTTGGA-ATTTTTAA-CACATGATCTTAA | loxF8L | 4 | loxF8R | 3 | ATTTTTAA/TTAAAAAT | 7 |
| chr1 | 7972541 | TTATTATTGTGTT-TTAAAAAT-TTCCCAGTTTTAT | loxF8R | 4 | loxF8L | 3 | ATTTTTAA/TTAAAAAT | 7 |
| chr6 | 76658414 | ATTTATATGTGGA-ATTTTTAA-ACGTCAAATTTAT | loxF8L | 3 | loxF8L | 4 | ATTTTTAA/TTAAAAAT | 7 |
| chr1 | 14390643 | TTGAGACTGGGTG-ATTTTTAA-TAAACAAATTTAT | loxF8R | 4 | loxF8L | 3 | ATTTTTAA/TTAAAAAT | 7 |
| chr2 | 240321343 | GTGATTCTGTGTA-TTAAAAAT-GCCACAAATATAT | loxF8L | 4 | loxF8L | 3 | ATTTTTAA/TTAAAAAT | 7 |
| --- | | | | | | | | 0 |
| chr18 | 21464716 | CTGACATTATGTG-CAAAAAAG-CACACAATGTCAC | loxF8R | 3 | loxF8R | 3 | CAAAAAAG/CTTTTTTG | 6 |
| chr15 | 101711220 | ATTCTTCTGGGGA-CAAAAAAG-AAAACAATCTTAG | loxF8L | 4 | loxF8R | 2 | CAAAAAAG/CTTTTTTG | 6 |
| --- | | | | | | | | 0 |
| chr15 | 56632946 | CTAAGTGTGTGTG-CATGTTTG-TACACATATATAT | loxF8R | 2 | loxF8L | 3 | CAAACATG/CATGTTTG | 5 |
| chr19 | 40686013 | GTGAGACTGTGTG-CATGTTTG-TACACAACCGTAG | loxF8R | 3 | loxF8R | 3 | CAAACATG/CATGTTTG | 6 |
| chr20 | 44614267 | ATAAATGTGTGTA-CATGTTTG-TCAATAATTTTAT | loxF8L | 2 | loxF8L | 4 | CAAACATG/CATGTTTG | 6 |
| chr18 | 26502750 | CAAAGATTGTTTT-CAAACATG-CAAACAATGTATG | loxF8R | 3 | loxF8R | 4 | CAAACATG/CATGTTTG | 7 |
| --- | | | | | | | | 0 |
| chr7 | 2697940 | ATATATTTGTGTA-TATTTGTG-TGCACACATGTAT | loxF8L | 3 | loxF8L | 3 | CACAAATA/TATTTGTG | 6 |
| chr8 | 120259362 | CTAAGAATGAGTG-CACAAATA-AAGACAGATTTTT | loxF8R | 2 | loxF8L | 4 | CACAAATA/TATTTGTG | 6 |
| chr10 | 32939831 | ACAGTACTGTGGA-CACAAATA-CACACTACATTAG | loxF8L | 4 | loxF8R | 3 | CACAAATA/TATTTGTG | 7 |
| --- | | | | | | | | 0 |
| chr21 | 16362017 | GTAAATATGTGCA-CACACACA-TGCATACATTTAT | loxF8L | 3 | loxF8L | 3 | CACACACA/TGTGTGTG | 6 |
| chr1 | 56592501 | ATCAAACTGTTGA-TGTGTGTG-TCTACAGGGTTAT | loxF8L | 3 | loxF8L | 3 | CACACACA/TGTGTGTG | 6 |
| chrY | 14599203 | CTATATTTGTGTG-TGTGTGTG-TCCATATATATAT | loxF8R | 3 | loxF8L | 3 | CACACACA/TGTGTGTG | 6 |
| chr4 | 60572863 | ATATATATGTGTA-TGTGTGTG-TGCACATATATAT | loxF8L | 3 | loxF8L | 3 | CACACACA/TGTGTGTG | 6 |
| chr4 | 60572671 | ATATATGTGTGTA-TGTGTGTG-TACACATATATAT | loxF8L | 3 | loxF8L | 3 | CACACACA/TGTGTGTG | 6 |

| | | | | | | | | |
|---|---|---|---|---|---|---|---|---|
| chr4 | 60572699 | ATATATGTGTGTA-TGTGTGTG-TACACATATATAT | loxF8L | 3 | loxF8L | 3 | CACACACA/TGTGTGTG | 6 |
| chr4 | 60572783 | ATATATGTGTGTA-TGTGTGTG-TACACATATATAT | loxF8L | 3 | loxF8L | 3 | CACACACA/TGTGTGTG | 6 |
| chr4 | 60572727 | ATATATGTGTGTA-TGTGTGTG-TACACATATATAT | loxF8L | 3 | loxF8L | 3 | CACACACA/TGTGTGTG | 6 |
| chr4 | 60572755 | ATATATGTGTGTA-TGTGTGTG-TACACATATATAT | loxF8L | 3 | loxF8L | 3 | CACACACA/TGTGTGTG | 6 |
| chr4 | 60572811 | ATATATGTGTGTA-TGTGTGTG-TACACATATATAT | loxF8L | 3 | loxF8L | 3 | CACACACA/TGTGTGTG | 6 |
| chr4 | 60572837 | ATATATATGTGTA-TGTGTGTG-TACACATATATAT | loxF8L | 3 | loxF8L | 3 | CACACACA/TGTGTGTG | 6 |
| chr1 | 40140719 | AAATATGTCTGGA-CACACACA-CACACAATCATAC | loxF8L | 4 | loxF8R | 2 | CACACACA/TGTGTGTG | 6 |
| chr13 | 30227086 | ATAAATATGTGTA-CACACACA-CACACACTCTCTC | loxF8L | 2 | loxF8R | 4 | CACACACA/TGTGTGTG | 6 |
| chr3 | 151797795 | ATAAATATGTGTA-TGTGTGTG-TACATATATGTAT | loxF8L | 2 | loxF8L | 4 | CACACACA/TGTGTGTG | 6 |
| chr4 | 12757540 | ATACATATATGCA-CACACACA-TACACAGATATAT | loxF8L | 4 | loxF8L | 2 | CACACACA/TGTGTGTG | 6 |
| chr1 | 79042150 | CTAGTATTGTGTG-TGTGTGTG-TGCATATATATAT | loxF8R | 2 | loxF8L | 4 | CACACACA/TGTGTGTG | 6 |
| chr1 | 20872104 | AAAAATCTGTGTG-TGTGTGTG-TATACACATATAT | loxF8L | 3 | loxF8L | 4 | CACACACA/TGTGTGTG | 7 |
| chr3 | 59018919 | AAAAATGTGTATA-CACACACA-CACACAATCACAC | loxF8L | 4 | loxF8R | 3 | CACACACA/TGTGTGTG | 7 |
| chr9 | 84615295 | AAAATTTTGTGTG-CACACACA-CACACACACTTAT | loxF8R | 4 | loxF8R | 3 | CACACACA/TGTGTGTG | 7 |
| chr3 | 189717172 | ACAAATATGTACA-CACACACA-CACACAGTGTTTG | loxF8L | 4 | loxF8R | 3 | CACACACA/TGTGTGTG | 7 |
| chr5 | 75286840 | CAAGGTGTGTGTG-TGTGTGTG-TGCACATATTTGT | loxF8R | 4 | loxF8L | 3 | CACACACA/TGTGTGTG | 7 |
| chr1 | 105568046 | AGAAGATTGTGTA-CACACACA-CACACACACATGG | loxF8R | 3 | loxF8R | 4 | CACACACA/TGTGTGTG | 7 |
| chr22 | 15628652 | CTGCGTGTGTGTG-TGTGTGTG-TCCACATATATCT | loxF8R | 4 | loxF8L | 3 | CACACACA/TGTGTGTG | 7 |
| chr3 | 100591084 | ATAAAAATTTGGA-CACACACA-CACACACACGTGG | loxF8L | 3 | loxF8R | 4 | CACACACA/TGTGTGTG | 7 |
| chr10 | 106130132 | ATATATGTGTGCA-TGTGTGTG-TGTATATATTTAT | loxF8L | 3 | loxF8L | 4 | CACACACA/TGTGTGTG | 7 |
| chr12 | 130043140 | ATGAATGTGTGCG-CACACACA-TGCATACATTTAT | loxF8L | 4 | loxF8L | 3 | CACACACA/TGTGTGTG | 7 |
| chr8 | 108025092 | ATAAATGTGTATA-CACACACA-CACACAAGCAAAC | loxF8L | 3 | loxF8R | 4 | CACACACA/TGTGTGTG | 7 |
| chr17 | 52486606 | ATAAATGTGTATA-CACACACA-CACACACACACAG | loxF8L | 3 | loxF8R | 4 | CACACACA/TGTGTGTG | 7 |
| chr8 | 89732962 | ATAAATGTGTGTG-TGTGTGTG-CACCCAATAGCAG | loxF8L | 3 | loxF8R | 4 | CACACACA/TGTGTGTG | 7 |
| chr8 | 125635566 | ATATATATGTATA-CACACACA-CACACAAATTTAT | loxF8L | 4 | loxF8L | 3 | CACACACA/TGTGTGTG | 7 |
| chr11 | 49784779 | ATAAATTTGTGTG-TGTGTGTG-TATACATATGTAT | loxF8L | 3 | loxF8L | 4 | CACACACA/TGTGTGTG | 7 |
| chr11 | 93757117 | ATAAGTATGTGTG-TGTGTGTG-TGCATACATATAT | loxF8R | 3 | loxF8L | 4 | CACACACA/TGTGTGTG | 7 |
| chr4 | 131056602 | ATACATATATGGA-CACACACA-TACATATATATAT | loxF8L | 3 | loxF8L | 4 | CACACACA/TGTGTGTG | 7 |
| chr7 | 53468836 | CTACGTGTGTGTG-TGTGTGTG-TACATATATGTAT | loxF8R | 3 | loxF8L | 4 | CACACACA/TGTGTGTG | 7 |
| chr9 | 21562431 | ATACATATGTGTA-TGTGTGTG-TATACATATGTAT | loxF8L | 3 | loxF8L | 4 | CACACACA/TGTGTGTG | 7 |
| chr2 | 112142761 | GTGTGTTTGTGTG-TGTGTGTG-TCAATAGATGTAT | loxF8R | 4 | loxF8L | 3 | CACACACA/TGTGTGTG | 7 |
| chr18 | 55239492 | TTAAAACAGTGTG-TGTGTGTG-TCCATACATGTAT | loxF8R | 4 | loxF8L | 3 | CACACACA/TGTGTGTG | 7 |
| chr1 | 194524699 | ATATATCTTTACA-CACACACA-TACACATATCTAT | loxF8L | 4 | loxF8L | 3 | CACACACA/TGTGTGTG | 7 |
| chr2 | 105164775 | ATAGATGTATGCA-CACACACA-TACACACATATAT | loxF8L | 4 | loxF8L | 3 | CACACACA/TGTGTGTG | 7 |
| chr11 | 26400847 | ATATATATATAGA-CACACACA-CACACAAACATAT | loxF8L | 4 | loxF8R | 3 | CACACACA/TGTGTGTG | 7 |
| chr12 | 131704353 | ATATATATATAGA-CACACACA-CACACACAATTAG | loxF8L | 4 | loxF8R | 3 | CACACACA/TGTGTGTG | 7 |
| chr2 | 198114788 | CTCTGAATGTGTG-TGTGTGTG-TGCATATATATAT | loxF8R | 3 | loxF8L | 4 | CACACACA/TGTGTGTG | 7 |
| chr7 | 145190405 | CTAAAAAGCTGTG-TGTGTGTG-TCCATATATATAT | loxF8R | 4 | loxF8L | 3 | CACACACA/TGTGTGTG | 7 |
| chr3 | 12130127 | CCAAGTGTGTGTG-TGTGTGTG-TACATATATATAT | loxF8R | 3 | loxF8L | 4 | CACACACA/TGTGTGTG | 7 |
| chr2 | 219380832 | ATATATATGTACA-CACACACA-CACACACACTGAG | loxF8L | 4 | loxF8R | 3 | CACACACA/TGTGTGTG | 7 |
| chr15 | 66509796 | CTATAATTATGTG-TGTGTGTG-GCTACATATATAT | loxF8R | 3 | loxF8L | 4 | CACACACA/TGTGTGTG | 7 |
| chr6 | 77349206 | ATATATCTGTGTG-TGTGTGTG-TATACATATATAT | loxF8L | 3 | loxF8L | 4 | CACACACA/TGTGTGTG | 7 |
| chr13 | 60977771 | ATATATATGTGTA-TGTGTGTG-TATACATATATAT | loxF8L | 3 | loxF8L | 4 | CACACACA/TGTGTGTG | 7 |
| chr13 | 60977797 | ATATATATGTGTA-TGTGTGTG-TATACATATATAT | loxF8L | 3 | loxF8L | 4 | CACACACA/TGTGTGTG | 7 |
| chr13 | 60977875 | ATATATATGTGTA-TGTGTGTG-TATACATATATAT | loxF8L | 3 | loxF8L | 4 | CACACACA/TGTGTGTG | 7 |
| chr13 | 60977823 | ATATATATGTGTA-TGTGTGTG-TATACATATATAT | loxF8L | 3 | loxF8L | 4 | CACACACA/TGTGTGTG | 7 |
| chr13 | 60977849 | ATATATATGTGTA-TGTGTGTG-TATACATATATAT | loxF8L | 3 | loxF8L | 4 | CACACACA/TGTGTGTG | 7 |
| chrX | 148085470 | ATATATATGTGTG-TGTGTGTG-TGCACATATATAT | loxF8L | 4 | loxF8L | 3 | CACACACA/TGTGTGTG | 7 |
| chr18 | 69022278 | ATATATATGTGGG-CACACACA-CACACATATATAT | loxF8L | 3 | loxF8L | 4 | CACACACA/TGTGTGTG | 7 |
| chr6 | 51969760 | ATATATATGTGTA-CACACACA-CACACACATATAT | loxF8L | 3 | loxF8L | 4 | CACACACA/TGTGTGTG | 7 |
| chrX | 67205462 | ATATATCTATATA-CACACACA-TACACACATATAT | loxF8L | 4 | loxF8L | 3 | CACACACA/TGTGTGTG | 7 |
| chr7 | 110853123 | ATATATCTGAGTA-CACACACA-CACACAAACACAT | loxF8L | 3 | loxF8R | 4 | CACACACA/TGTGTGTG | 7 |
| chr11 | 80994829 | ATATATCTGTGTG-TGTGTGTG-TCTACATACATAT | loxF8L | 3 | loxF8L | 4 | CACACACA/TGTGTGTG | 7 |
| chr13 | 71730582 | ATATGAGTGTTTG-CACACACA-TACACACATATAT | loxF8R | 4 | loxF8L | 3 | CACACACA/TGTGTGTG | 7 |
| chr3 | 59588761 | ATATATGTGTGTA-CACACACA-CACACACACATAT | loxF8L | 3 | loxF8R | 4 | CACACACA/TGTGTGTG | 7 |
| chr7 | 35643599 | ATATATGTGTGTA-CACACACA-CACACACACTTTC | loxF8L | 3 | loxF8R | 4 | CACACACA/TGTGTGTG | 7 |
| chr9 | 131202374 | ATATATGTGTGTA-CACACACA-CACACACATATAT | loxF8L | 3 | loxF8L | 4 | CACACACA/TGTGTGTG | 7 |
| chr12 | 14472807 | ATATGAATGTATG-CACACACA-CACACTATATTAA | loxF8R | 4 | loxF8R | 3 | CACACACA/TGTGTGTG | 7 |
| chr4 | 66977495 | TTATATGTGTGTA-CACACACA-CACACACACATAG | loxF8L | 4 | loxF8R | 3 | CACACACA/TGTGTGTG | 7 |
| chr16 | 76919167 | GTGAGAGTGTGTG-TGTGTGTG-TCCCCATATTTCC | loxF8R | 3 | loxF8L | 4 | CACACACA/TGTGTGTG | 7 |
| chr2 | 22088610 | GTAAATCAGTGCA-CACACACA-CACACACACATAC | loxF8L | 3 | loxF8R | 4 | CACACACA/TGTGTGTG | 7 |
| chr3 | 113653725 | TAAAATATGTGTA-CACACACA-CACACATACTTAC | loxF8L | 4 | loxF8R | 3 | CACACACA/TGTGTGTG | 7 |
| --- | | | | | | | | 0 |
| chr11 | 86685872 | ATATATCTGTATA-CACATATA-TACACACATATAT | loxF8L | 3 | loxF8L | 3 | CACATATA/TATATGTG | 6 |
| chr4 | 100178477 | ATATATGTGTGCA-CACATATA-TACACACATATAT | loxF8L | 3 | loxF8L | 3 | CACATATA/TATATGTG | 6 |
| chr4 | 100178449 | ATATATGTGTGCA-CACATATA-TACACACATATAT | loxF8L | 3 | loxF8L | 3 | CACATATA/TATATGTG | 6 |
| chr4 | 100178411 | ATATATGTGTGCA-CACATATA-TACACACATATAT | loxF8L | 3 | loxF8L | 3 | CACATATA/TATATGTG | 6 |
| chrX | 75983758 | ATAAATGTGTGCA-TATATGTG-TGCACACACATAT | loxF8L | 2 | loxF8L | 4 | CACATATA/TATATGTG | 6 |
| chr11 | 81233761 | GTATATATGTGAA-CACATATA-TACACAGATTCAT | loxF8L | 4 | loxF8L | 2 | CACATATA/TATATGTG | 6 |
| chr12 | 102095408 | ATCAACCTGTGTA-TATATGTG-TGCATATATTTTT | loxF8L | 3 | loxF8L | 4 | CACATATA/TATATGTG | 7 |
| chr6 | 46414789 | ACAAATCTGTCTA-TATATGTG-CACACATATATAT | loxF8L | 3 | loxF8L | 4 | CACATATA/TATATGTG | 7 |
| chr4 | 100178437 | ACATATATGTGTA-TATATGTG-TGCACACATATAT | loxF8L | 4 | loxF8L | 3 | CACATATA/TATATGTG | 7 |
| chr4 | 100178505 | ATATATGTGTGCA-CACATATA-TACACACATATGT | loxF8L | 3 | loxF8L | 4 | CACATATA/TATATGTG | 7 |
| chr8 | 47121497 | ATAAATATGTATA-CACATATA-TACACAAATATCT | loxF8L | 3 | loxF8L | 4 | CACATATA/TATATGTG | 7 |
| chr13 | 65749376 | CAAAGTTTGTTTA-CACATATA-TGCACGGGTTTAT | loxF8R | 4 | loxF8L | 3 | CACATATA/TATATGTG | 7 |
| chr11 | 104525169 | ATAAATATATACA-CACATATA-TACACACATATAT | loxF8L | 4 | loxF8L | 3 | CACATATA/TATATGTG | 7 |
| chr2 | 208637167 | ATACATATGTGTA-TATATGTG-TGCATACATATAT | loxF8L | 3 | loxF8L | 4 | CACATATA/TATATGTG | 7 |
| chr6 | 92330380 | ATACATTAGTGCA-CACATATA-TACACAAATTAAT | loxF8L | 4 | loxF8L | 3 | CACATATA/TATATGTG | 7 |
| chr11 | 84628079 | ATATATATGTACA-CACATATA-TACACACATATAT | loxF8L | 4 | loxF8L | 3 | CACATATA/TATATGTG | 7 |
| chr11 | 29460112 | ATATATATGTGTA-CACATATA-TATACATATATAT | loxF8L | 3 | loxF8L | 4 | CACATATA/TATATGTG | 7 |
| chr11 | 29460138 | ATATATATGTGTA-CACATATA-TATACATATATAT | loxF8L | 3 | loxF8L | 4 | CACATATA/TATATGTG | 7 |
| chr2 | 208637307 | ATATATATGTGTA-TATATGTG-TGCATACATATAT | loxF8L | 3 | loxF8L | 4 | CACATATA/TATATGTG | 7 |
| chr2 | 208637475 | ATATATATGTGTA-TATATGTG-TGCATACATATAT | loxF8L | 3 | loxF8L | 4 | CACATATA/TATATGTG | 7 |
| chr2 | 208637503 | ATATATATGTGTA-TATATGTG-TGCATACATATAT | loxF8L | 3 | loxF8L | 4 | CACATATA/TATATGTG | 7 |
| chr2 | 208637251 | ATATATATGTGTA-TATATGTG-TGCATACATATAT | loxF8L | 3 | loxF8L | 4 | CACATATA/TATATGTG | 7 |
| chr2 | 208637223 | ATATATATGTGTA-TATATGTG-TGCATACATATAT | loxF8L | 3 | loxF8L | 4 | CACATATA/TATATGTG | 7 |
| chr2 | 208637195 | ATATATATGTGTA-TATATGTG-TGCATACATATAT | loxF8L | 3 | loxF8L | 4 | CACATATA/TATATGTG | 7 |

| | | | | | | | | |
|---|---|---|---|---|---|---|---|---|
| chr2 | 208637279 | ATATATATGTGTA-TATATGTG-TGCATACATATAT | loxF8L | 3 | loxF8L | 4 | CACATATA/TATATGTG | 7 |
| --- | | | | | | | | 0 |
| chr6 | 147679114 | ATAAATCTATGAA-TATTTATG-TACACATTTTTAT | loxF8L | 2 | loxF8L | 3 | CATAAATA/TATTTATG | 5 |
| chr12 | 24275350 | ATACATGTGTGTA-CATAAATA-CCCACAAACTTGG | loxF8L | 3 | loxF8R | 3 | CATAAATA/TATTTATG | 6 |
| chr1 | 232804339 | GTAGATGTGTGTA-TATTTATG-TACACACATTTGT | loxF8L | 4 | loxF8L | 3 | CATAAATA/TATTTATG | 7 |
| chr13 | 78124215 | ATAAATATAGGGA-CATAAATA-TACACATAGATAT | loxF8L | 3 | loxF8L | 4 | CATAAATA/TATTTATG | 7 |
| chr7 | 20242785 | ATACATATATGTA-CATAAATA-TATACATATTTAT | loxF8L | 4 | loxF8L | 3 | CATAAATA/TATTTATG | 7 |
| chr5 | 118229357 | ATATATCTGTACA-CATAAATA-TAGACATAGTTAT | loxF8L | 3 | loxF8L | 4 | CATAAATA/TATTTATG | 7 |
| chr12 | 129576487 | ATATATCTGTTTA-CATAAATA-TATACACATATAT | loxF8L | 3 | loxF8L | 4 | CATAAATA/TATTTATG | 7 |
| chr12 | 129576527 | ATATATCTGTTTA-CATAAATA-TATACACATATAT | loxF8L | 3 | loxF8L | 4 | CATAAATA/TATTTATG | 7 |
| --- | | | | | | | | 0 |
| chr4 | 179679257 | ATATATGTGTGTA-CATATATG-TACACACATATAT | loxF8L | 3 | loxF8L | 3 | CATATATG/CATATATG | 6 |
| chr4 | 179679201 | ATATATGTGTGTA-CATATATG-TACACACATATAT | loxF8L | 3 | loxF8L | 3 | CATATATG/CATATATG | 6 |
| chr2 | 234177898 | CTAGGAATATGTT-CATATATG-TCCATATATTTAT | loxF8R | 4 | loxF8L | 2 | CATATATG/CATATATG | 6 |
| chrX | 121064451 | ATATATATGTGTA-CATATATG-TATACATATATAT | loxF8L | 3 | loxF8L | 4 | CATATATG/CATATATG | 7 |
| --- | | | | | | | | 0 |
| chr7 | 146938430 | ATACATGTGTGTA-TACATATG-TACACATATATAT | loxF8L | 3 | loxF8L | 3 | CATATGTA/TACATATG | 6 |
| chr9 | 1443139 | ATAAATATATAAA-CATATGTA-TACACATATTTAT | loxF8L | 4 | loxF8L | 2 | CATATGTA/TACATATG | 6 |
| chr7 | 145219292 | CTATGCTTATGTG-CATATGTA-CACACACGCTTGT | loxF8R | 3 | loxF8R | 4 | CATATGTA/TACATATG | 7 |
| chr7 | 41776030 | ATACATATGTATA-CATATGTA-TACACACATATAT | loxF8L | 4 | loxF8L | 3 | CATATGTA/TACATATG | 7 |
| chr7 | 41776018 | ATATATATGTGCA-TACATATG-TATACATATGTAT | loxF8L | 3 | loxF8L | 4 | CATATGTA/TACATATG | 7 |
| chr5 | 84043735 | ATACATGTGTATA-CATATGTA-CACATATTATTAG | loxF8L | 4 | loxF8R | 3 | CATATGTA/TACATATG | 7 |
| chr16 | 48186680 | TTAAATGTATGTA-TACATATG-TCCATATATATAT | loxF8L | 4 | loxF8L | 3 | CATATGTA/TACATATG | 7 |
| chr12 | 19617688 | ATATATATATGTA-CATATGTA-TACACAAATATAT | loxF8L | 4 | loxF8L | 3 | CATATGTA/TACATATG | 7 |
| chr17 | 52093129 | ATATATGTGTAGA-TACATATG-TACACATACATAT | loxF8L | 3 | loxF8L | 4 | CATATGTA/TACATATG | 7 |
| --- | | | | | | | | 0 |
| chrX | 99209834 | CTGAGAATGTGAG-CCCAGATG-TGCACAGATTTAC | loxF8R | 3 | loxF8L | 2 | CATCTGGG/CCCAGATG | 5 |
| chr4 | 136237795 | CTTAGCTTCTGGG-CCCAGATG-CCCTCAGACTTAT | loxF8R | 4 | loxF8L | 3 | CATCTGGG/CCCAGATG | 7 |
| --- | | | | | | | | 0 |
| chr5 | 111873307 | ATAAACTTATGGA-CTGAAAAC-CACACAAACTTAT | loxF8L | 3 | loxF8R | 2 | CTGAAAAC/GTTTTCAG | 5 |
| chr20 | 45575462 | AAAGATCTGTACA-CTGAAAAC-TACAAAATATTAG | loxF8L | 4 | loxF8R | 3 | CTGAAAAC/GTTTTCAG | 7 |
| --- | | | | | | | | 0 |
| chr3 | 106325090 | ACAAATCTGGGGG-CTTAAAAC-AACACAATTTTAG | loxF8L | 3 | loxF8R | 2 | CTTAAAAC/GTTTTAAG | 5 |
| chr5 | 36935679 | ATAAAGCTGGTGA-CTTAAAAC-AACAGAGATTTAT | loxF8L | 3 | loxF8L | 3 | CTTAAAAC/GTTTTAAG | 6 |
| chr5 | 82789660 | ATAAATCTGTGTT-GTTTTAAG-TCACCAAGTTTAT | loxF8L | 2 | loxF8L | 4 | CTTAAAAC/GTTTTAAG | 6 |
| chr9 | 26761009 | ATAAATCTGTGTG-GTTTTAAG-CACTAAATCTGTG | loxF8L | 2 | loxF8R | 4 | CTTAAAAC/GTTTTAAG | 6 |
| chr14 | 51516689 | AAAAAGCTGTGGG-CTTAAAAC-TCAGCTTATTTAT | loxF8L | 3 | loxF8L | 4 | CTTAAAAC/GTTTTAAG | 7 |
| chr16 | 65448757 | ATAAATTTGTGTT-GTTTTAAG-TCATCAGGTTTGT | loxF8L | 3 | loxF8L | 4 | CTTAAAAC/GTTTTAAG | 7 |
| chr1 | 109871760 | ATAAATTTGTGTT-GTTTTAAG-TCACCAAATTTGT | loxF8L | 3 | loxF8L | 4 | CTTAAAAC/GTTTTAAG | 7 |
| chr2 | 164581662 | ATAAATATGTGTT-GTTTTAAG-TCACCAAATTTGT | loxF8L | 3 | loxF8L | 4 | CTTAAAAC/GTTTTAAG | 7 |
| chr2 | 189014000 | ATAAATTTAGTGA-CTTAAAAC-CACACAGATTTCT | loxF8L | 4 | loxF8L | 3 | CTTAAAAC/GTTTTAAG | 7 |
| chr5 | 116357611 | ATAAACCTGGTGG-CTTAAAAC-AACACAAATTTAT | loxF8L | 4 | loxF8L | 3 | CTTAAAAC/GTTTTAAG | 7 |
| chr2 | 11520287 | ATAGATCTATTGG-CTTAAAAC-AGCACATATTTAT | loxF8L | 4 | loxF8L | 3 | CTTAAAAC/GTTTTAAG | 7 |
| chr10 | 93210726 | ATAAATCTCTGTT-GTTTTAAG-CCAACCCATTTAT | loxF8L | 3 | loxF8L | 4 | CTTAAAAC/GTTTTAAG | 7 |
| chr20 | 37185561 | ATAAATGTAGGGG-CTTAAAAC-ACCACCGACTTAT | loxF8L | 4 | loxF8L | 3 | CTTAAAAC/GTTTTAAG | 7 |
| --- | | | | | | | | 0 |
| chr2 | 187472189 | ATAATATTTTGTG-GCTGTGTC-TCCACATTTTTAT | loxF8R | 3 | loxF8L | 2 | GACACAGC/GCTGTGTC | 5 |
| chr2 | 98613667 | ATGAGATTTTGGG-GACACAGC-CAAACAATATCAG | loxF8R | 4 | loxF8R | 3 | GACACAGC/GCTGTGTC | 7 |
| --- | | | | | | | | 0 |
| chr4 | 47017791 | TTGAAGCTGTGGA-GATATTCA-TCCACAGATTGTT | loxF8L | 3 | loxF8L | 2 | GATATTCA/TGAATATC | 5 |
| chr8 | 125067555 | CTAAGACTGTGGT-GATATTCA-TTCAAATATTTTT | loxF8R | 3 | loxF8L | 4 | GATATTCA/TGAATATC | 7 |
| chr8 | 112745615 | ATGAGTTTGTGTG-GATATTCA-TGCAAAAATTTAC | loxF8R | 3 | loxF8L | 4 | GATATTCA/TGAATATC | 7 |
| --- | | | | | | | | 0 |
| chr7 | 50685102 | CTATGATTCTGTG-GATTTAAC-TCCATAAATTTAA | loxF8R | 2 | loxF8L | 3 | GATTTAAC/GTTAAATC | 5 |
| chr1 | 215453324 | ATATATCGGTGGT-GTTAAATC-TCCAGAGATTTTA | loxF8L | 3 | loxF8L | 3 | GATTTAAC/GTTAAATC | 6 |
| chr10 | 114066906 | ATAAATGGGTGCA-GATTTAAC-TTGACACATTAAT | loxF8L | 3 | loxF8L | 4 | GATTTAAC/GTTAAATC | 7 |
| --- | | | | | | | | 0 |
| chr4 | 185024030 | AGAAATCCATGGA-GCACTCCA-CACACATTCTGAG | loxF8L | 3 | loxF8R | 2 | GCACTCCA/TGGAGTGC | 5 |
| chr7 | 41250361 | TTCACATTGTGTG-GCACTCCA-TTCACTCATTTGT | loxF8R | 3 | loxF8L | 4 | GCACTCCA/TGGAGTGC | 7 |
| --- | | | | | | | | 0 |
| chr4 | 21383225 | ATAAATTTCTGGT-GTTTATGC-CACACAGTCTGTG | loxF8L | 3 | loxF8R | 3 | GCATAAAC/GTTTATGC | 6 |
| chr18 | 63905601 | ATACATGTTTGTA-GTTTATGC-CACACAATCTATG | loxF8L | 4 | loxF8R | 2 | GCATAAAC/GTTTATGC | 6 |
| chr1 | 223372771 | CACAGATTGGGTG-GCATAAAC-AACAGACATTTAT | loxF8R | 3 | loxF8L | 4 | GCATAAAC/GTTTATGC | 7 |
| chr5 | 22960989 | CAAAGACTGGGTG-GCATAAAC-AACAAAGATGTAT | loxF8R | 3 | loxF8L | 4 | GCATAAAC/GTTTATGC | 7 |
| --- | | | | | | | | 0 |
| chr3 | 97493306 | CACAGATTGGGTG-GCTTAAAA-GGCACAAATTTAT | loxF8R | 3 | loxF8L | 3 | GCTTAAAA/TTTTAAGC | 6 |
| chrX | 44005178 | CACAAATTTTGTG-GCTTAAAA-AACACAGATTTAT | loxF8R | 4 | loxF8L | 2 | GCTTAAAA/TTTTAAGC | 6 |
| chr8 | 22832476 | ATAAATGTGTGTT-TTTTAAGC-CACCCAGTCTATG | loxF8L | 3 | loxF8R | 4 | GCTTAAAA/TTTTAAGC | 7 |
| chr4 | 112991405 | CACAGATAGGGTG-GCTTAAAA-CCCAGAAATTTAT | loxF8R | 4 | loxF8L | 3 | GCTTAAAA/TTTTAAGC | 7 |
| chr3 | 174493525 | ATAAATTTCTGTT-TTTTAAGC-CACACAGTCTGTG | loxF8L | 4 | loxF8R | 3 | GCTTAAAA/TTTTAAGC | 7 |
| chr15 | 72088160 | ATAAATTTTTGTT-TTTTAAGC-CACTCAATCTGTG | loxF8L | 4 | loxF8R | 3 | GCTTAAAA/TTTTAAGC | 7 |
| chr13 | 27492423 | CTAAATTTCTGTT-TTTTAAGC-CACACAGTCTGTG | loxF8R | 4 | loxF8R | 3 | GCTTAAAA/TTTTAAGC | 7 |
| --- | | | | | | | | 0 |
| chr4 | 54846164 | CACAGATTGGGTG-GCTTAAAC-ACCACAGATGTAT | loxF8R | 3 | loxF8L | 2 | GCTTAAAC/GTTTAAGC | 5 |
| chr11 | 87134079 | CACAGACTGTGTG-GCTTAAAC-AGCAGAGATTTAT | loxF8R | 3 | loxF8L | 3 | GCTTAAAC/GTTTAAGC | 6 |
| chr3 | 34897284 | CAAAGACTGGGTG-GCTTAAAC-ACCAGAAATTTAT | loxF8R | 3 | loxF8L | 3 | GCTTAAAC/GTTTAAGC | 6 |
| chr5 | 170852876 | CACAGACTGTGTG-GCTTAAAC-CACACAAATTTAT | loxF8R | 3 | loxF8L | 3 | GCTTAAAC/GTTTAAGC | 6 |
| chr1 | 208000566 | TTCAGACTGTGTG-GCTTAAAC-AACACAAATTTAT | loxF8R | 3 | loxF8L | 3 | GCTTAAAC/GTTTAAGC | 6 |
| chr17 | 11146773 | CACAGACTGGGTG-GCTTAAAC-ACCAGAGATTTAT | loxF8R | 4 | loxF8L | 2 | GCTTAAAC/GTTTAAGC | 6 |
| chr12 | 41130644 | ATAAATCTCTGTA-GTTTAAGC-CACACAAGCTGTA | loxF8L | 2 | loxF8R | 4 | GCTTAAAC/GTTTAAGC | 6 |
| chr3 | 90037366 | ATAAATGTCTGTT-GTTTAAGC-CACACAGTCTTTG | loxF8L | 4 | loxF8R | 2 | GCTTAAAC/GTTTAAGC | 6 |
| chr9 | 89163771 | CACAGACTGGGTG-GCTTAAAC-TACACACATTTAT | loxF8R | 4 | loxF8L | 2 | GCTTAAAC/GTTTAAGC | 6 |
| chr3 | 110627533 | ATAAATGTTTGTT-GTTTAAGC-CACACAATCTATG | loxF8L | 4 | loxF8R | 2 | GCTTAAAC/GTTTAAGC | 6 |
| chr8 | 55554779 | ATAAATTTCTGTT-GTTTAAGC-CACACAATCATAA | loxF8L | 4 | loxF8R | 2 | GCTTAAAC/GTTTAAGC | 6 |
| chr3 | 60805273 | CAAGGATTGTGTG-GCTTAAAC-AACAGAAATTTAT | loxF8R | 2 | loxF8L | 4 | GCTTAAAC/GTTTAAGC | 6 |
| chr4 | 30392885 | ATACATCTATGTT-GTTTAAGC-CACTCAATCTTTG | loxF8L | 4 | loxF8R | 2 | GCTTAAAC/GTTTAAGC | 6 |
| chr19 | 46575797 | AAAAGATTGGGTG-GCTTAAAC-AACAGACATTTAT | loxF8R | 3 | loxF8L | 4 | GCTTAAAC/GTTTAAGC | 7 |

| | | | | | | | | |
|---|---|---|---|---|---|---|---|---|
| chr8 | 69013922 | AGAAATCTGTGTT-GTTTAAGC-CACCCAGTCTGTG | loxF8L | 3 | loxF8R | 4 | GCTTAAAC/GTTTAAGC | 7 |
| chr1 | 34174735 | ATAAAATTCTGTT-GTTTAAGC-CACACAGTCTGTG | loxF8R | 4 | loxF8R | 3 | GCTTAAAC/GTTTAAGC | 7 |
| chrX | 6549611 | ATAAATATCTGTT-GTTTAAGC-CACACAGCCTTTG | loxF8L | 4 | loxF8R | 3 | GCTTAAAC/GTTTAAGC | 7 |
| chr10 | 99379843 | CATAGACTGTGTG-GCTTAAAC-AACAGATATTTAT | loxF8R | 3 | loxF8L | 4 | GCTTAAAC/GTTTAAGC | 7 |
| chr13 | 46929994 | ATAAATATTTGTT-GTTTAAGC-CACCCAATCTATG | loxF8L | 4 | loxF8R | 3 | GCTTAAAC/GTTTAAGC | 7 |
| chr8 | 26535052 | CATAGATTGGGTG-GCTTAAAC-AACAAATATTTAT | loxF8R | 3 | loxF8L | 4 | GCTTAAAC/GTTTAAGC | 7 |
| chrX | 126508087 | CATAGATTGAGTG-GCTTAAAC-AACAAATATTTAT | loxF8R | 3 | loxF8L | 4 | GCTTAAAC/GTTTAAGC | 7 |
| chr22 | 33401810 | ATAAATCTATGTT-GTTTAAGC-CACCCAGTCTGTG | loxF8L | 3 | loxF8R | 4 | GCTTAAAC/GTTTAAGC | 7 |
| chr7 | 96701517 | ATAAATCTCTGAT-GTTTAAGC-CACCCAGTCTATG | loxF8L | 3 | loxF8R | 4 | GCTTAAAC/GTTTAAGC | 7 |
| chrX | 103144662 | ATAAATCTCTGCT-GTTTAAGC-CACACAGTCTATA | loxF8L | 3 | loxF8R | 4 | GCTTAAAC/GTTTAAGC | 7 |
| chr9 | 129841131 | CACAGACTGGGTG-GCTTAAAC-GACAGAGATTTAT | loxF8R | 4 | loxF8L | 3 | GCTTAAAC/GTTTAAGC | 7 |
| chr19 | 47967419 | CACAGATGGGGTG-GCTTAAAC-GACAGAGATTTAT | loxF8R | 4 | loxF8L | 3 | GCTTAAAC/GTTTAAGC | 7 |
| chr4 | 104877336 | ATAAATCTCTGTT-GTTTAAGC-CACAAACTCTATG | loxF8L | 3 | loxF8R | 4 | GCTTAAAC/GTTTAAGC | 7 |
| chr12 | 114563391 | CACAAACTGTGTG-GCTTAAAC-AACAGAGATTTAT | loxF8R | 4 | loxF8L | 3 | GCTTAAAC/GTTTAAGC | 7 |
| chrX | 69895957 | CACAAACTGTGTG-GCTTAAAC-AACAGAGATTTAT | loxF8R | 4 | loxF8L | 3 | GCTTAAAC/GTTTAAGC | 7 |
| chr2 | 140551118 | ATAAATCTCTGTT-GTTTAAGC-CACCCAGTCTATG | loxF8L | 3 | loxF8R | 4 | GCTTAAAC/GTTTAAGC | 7 |
| chr5 | 107274460 | ATAAATCTCTGTT-GTTTAAGC-CACCCAGTCTGTG | loxF8L | 3 | loxF8R | 4 | GCTTAAAC/GTTTAAGC | 7 |
| chr6 | 82827093 | ATAAATCTCTGTT-GTTTAAGC-CACCCAGTCTGTG | loxF8L | 3 | loxF8R | 4 | GCTTAAAC/GTTTAAGC | 7 |
| chr9 | 107193498 | ATAAATCTCTGTT-GTTTAAGC-CACCCAGTCTGTG | loxF8L | 3 | loxF8R | 4 | GCTTAAAC/GTTTAAGC | 7 |
| chr10 | 72083015 | ATAAATCTCTGTT-GTTTAAGC-CACCCCATCTCTG | loxF8L | 3 | loxF8R | 4 | GCTTAAAC/GTTTAAGC | 7 |
| chr10 | 120984950 | CACAGATAGAGTG-GCTTAAAC-AACAGAGATTTAT | loxF8R | 4 | loxF8L | 3 | GCTTAAAC/GTTTAAGC | 7 |
| chr14 | 95819015 | CACAGATTGCCTG-GCTTAAAC-AACAGAGATTTAT | loxF8R | 4 | loxF8L | 3 | GCTTAAAC/GTTTAAGC | 7 |
| chr6 | 100866809 | ATAAATCTTTGTT-GTTTAAGC-CACCTAATCTGTG | loxF8L | 3 | loxF8R | 4 | GCTTAAAC/GTTTAAGC | 7 |
| chr16 | 85895078 | CATAGCTTGTGTG-GCTTAAAC-AGCAGACATTTAT | loxF8R | 3 | loxF8L | 4 | GCTTAAAC/GTTTAAGC | 7 |
| chr7 | 91115227 | CTTAGACTGGGTG-GCTTAAAC-AGCAGACATTTAT | loxF8R | 3 | loxF8L | 4 | GCTTAAAC/GTTTAAGC | 7 |
| chr2 | 53029114 | ATAAATGTCTGGT-GTTTAAGC-CACCCATTCTATG | loxF8L | 3 | loxF8R | 4 | GCTTAAAC/GTTTAAGC | 7 |
| chr15 | 57360761 | TTAAGACTTTGTG-GCTTAAAC-AACAGACATTTAT | loxF8R | 3 | loxF8L | 4 | GCTTAAAC/GTTTAAGC | 7 |
| chr4 | 14326414 | ATAAATGTCTGTT-GTTTAAGC-CACACAGTCTATG | loxF8L | 4 | loxF8R | 3 | GCTTAAAC/GTTTAAGC | 7 |
| chr2 | 12314900 | CATAGATTGGGTG-GCTTAAAC-AACAGACATTTAT | loxF8R | 3 | loxF8L | 4 | GCTTAAAC/GTTTAAGC | 7 |
| chr8 | 94399771 | CATAGATTGGGTG-GCTTAAAC-AACAGACATTTAT | loxF8R | 3 | loxF8L | 4 | GCTTAAAC/GTTTAAGC | 7 |
| chr18 | 52654419 | CACAGATTGGGTG-GCTTAAAC-AACAGACATTTAT | loxF8R | 3 | loxF8L | 4 | GCTTAAAC/GTTTAAGC | 7 |
| chr1 | 230001277 | CACAGATTGGGTG-GCTTAAAC-AACAGACATTTAT | loxF8R | 3 | loxF8L | 4 | GCTTAAAC/GTTTAAGC | 7 |
| chr5 | 163956811 | ATAAATGTCTGTT-GTTTAAGC-CACCCAATCTGTG | loxF8L | 4 | loxF8R | 3 | GCTTAAAC/GTTTAAGC | 7 |
| chr9 | 114944048 | CTTAGACTGGGTG-GCTTAAAC-AACAGACATTTAT | loxF8R | 3 | loxF8L | 4 | GCTTAAAC/GTTTAAGC | 7 |
| chr1 | 232921996 | CATAGATTGAGTG-GCTTAAAC-AACAGACATTTAT | loxF8R | 3 | loxF8L | 4 | GCTTAAAC/GTTTAAGC | 7 |
| chr4 | 54383832 | ATAAATGTCTGTT-GTTTAAGC-TACACAGTCTAAG | loxF8L | 4 | loxF8R | 3 | GCTTAAAC/GTTTAAGC | 7 |
| chr5 | 168387529 | CATAGATTGGGTG-GCTTAAAC-GTTACACATTTAT | loxF8R | 3 | loxF8L | 4 | GCTTAAAC/GTTTAAGC | 7 |
| chr4 | 189364870 | CACAGACTGGGTG-GCTTAAAC-AACACACATTTAT | loxF8R | 4 | loxF8L | 3 | GCTTAAAC/GTTTAAGC | 7 |
| chrX | 13575569 | ATAAATGTGTGTT-GTTTAAGC-CACCCAGTCTGTG | loxF8L | 3 | loxF8R | 4 | GCTTAAAC/GTTTAAGC | 7 |
| chr8 | 115175861 | ATAAATGTGTGTT-GTTTAAGC-CACTCAGTCTGTG | loxF8L | 3 | loxF8R | 4 | GCTTAAAC/GTTTAAGC | 7 |
| chr4 | 26892236 | CAAAGACTGCGTG-GCTTAAAC-ATCAAACATTTAT | loxF8R | 3 | loxF8L | 4 | GCTTAAAC/GTTTAAGC | 7 |
| chr14 | 85760769 | ATAAATGTTTGCT-GTTTAAGC-CACCCAATCTATG | loxF8L | 4 | loxF8R | 3 | GCTTAAAC/GTTTAAGC | 7 |
| chrX | 128869732 | CGTAGAATGGGTG-GCTTAAAC-ACCAAACATTTAT | loxF8R | 4 | loxF8L | 3 | GCTTAAAC/GTTTAAGC | 7 |
| chr13 | 21115302 | ATAAATGTTTGTT-GTTTAAGC-CACACAGTCTATG | loxF8L | 4 | loxF8R | 3 | GCTTAAAC/GTTTAAGC | 7 |
| chr1 | 170144656 | ATAAATGTTTGTT-GTTTAAGC-CACATAGTCTTTG | loxF8L | 4 | loxF8R | 3 | GCTTAAAC/GTTTAAGC | 7 |
| chr21 | 25628930 | CATAGATTGAGTG-GCTTAAAC-AACAAACATTTAT | loxF8R | 3 | loxF8L | 4 | GCTTAAAC/GTTTAAGC | 7 |
| chr21 | 25629017 | CATAGATTGAGTG-GCTTAAAC-AACAAACATTTAT | loxF8R | 3 | loxF8L | 4 | GCTTAAAC/GTTTAAGC | 7 |
| chrX | 110349270 | CCCAGATTGAGTG-GCTTAAAC-AACAAACATTTAT | loxF8R | 3 | loxF8L | 4 | GCTTAAAC/GTTTAAGC | 7 |
| chr6 | 95390709 | CTAAGACTGAGTA-GCTTAAAC-AACAAACATTTAT | loxF8R | 3 | loxF8L | 4 | GCTTAAAC/GTTTAAGC | 7 |
| chr7 | 47065340 | ATAAATTTATGTT-GTTTAAGC-CACACAGTCTGTG | loxF8L | 4 | loxF8R | 3 | GCTTAAAC/GTTTAAGC | 7 |
| chr5 | 104054931 | ATAAATTTATGTT-GTTTAAGC-CTCACAATCTATG | loxF8L | 4 | loxF8R | 3 | GCTTAAAC/GTTTAAGC | 7 |
| chr13 | 100913832 | ATAAATTTCTGAT-GTTTAAGC-CACCCAATCTGTG | loxF8L | 4 | loxF8R | 3 | GCTTAAAC/GTTTAAGC | 7 |
| chr12 | 42003452 | ATAAATTTCTGTA-GTTTAAGC-CACCCAGTCTGTG | loxF8L | 3 | loxF8R | 4 | GCTTAAAC/GTTTAAGC | 7 |
| chr6 | 152514840 | ATAAATTTCTGTA-GTTTAAGC-CACCCAGTCTGTG | loxF8L | 3 | loxF8R | 4 | GCTTAAAC/GTTTAAGC | 7 |
| chr7 | 113968198 | ATAAATTTCTGTA-GTTTAAGC-CACTCAGACTTAT | loxF8L | 3 | loxF8L | 4 | GCTTAAAC/GTTTAAGC | 7 |
| chr8 | 58551386 | ATAAATTTCTGTC-GTTTAAGC-CACCCAATCTATG | loxF8L | 4 | loxF8R | 3 | GCTTAAAC/GTTTAAGC | 7 |
| chr1 | 175053094 | CAGAGGTTGTGTG-GCTTAAAC-AACAGAAATTTAT | loxF8R | 3 | loxF8L | 4 | GCTTAAAC/GTTTAAGC | 7 |
| chr16 | 6771253 | ATAAATTTCTGTT-GTTTAAGC-CACACAGTCTACG | loxF8L | 4 | loxF8R | 3 | GCTTAAAC/GTTTAAGC | 7 |
| chr10 | 126282034 | CAGAGACTGTGTG-GCTTAAAC-AACAGAAATTTAT | loxF8R | 3 | loxF8L | 4 | GCTTAAAC/GTTTAAGC | 7 |
| chr16 | 54153425 | ATAAATTTCTGTT-GTTTAAGC-CACACAGTCTGTG | loxF8L | 4 | loxF8R | 3 | GCTTAAAC/GTTTAAGC | 7 |
| chr17 | 62510504 | CACAGACTGTGTG-GCTTAAAC-AACAGAAATTTAT | loxF8R | 3 | loxF8L | 4 | GCTTAAAC/GTTTAAGC | 7 |
| chr19 | 38866793 | ATAAATTTCTGTT-GTTTAAGC-CACACAGTCTGTG | loxF8L | 4 | loxF8R | 3 | GCTTAAAC/GTTTAAGC | 7 |
| chr2 | 117492981 | CACAGACTGTGTG-GCTTAAAC-AACAGAAATTTAT | loxF8R | 3 | loxF8L | 4 | GCTTAAAC/GTTTAAGC | 7 |
| chr3 | 73347092 | CACAGACTGTGTG-GCTTAAAC-AACAGAAATTTAT | loxF8R | 3 | loxF8L | 4 | GCTTAAAC/GTTTAAGC | 7 |
| chr7 | 128564414 | CACAGACTGTGTG-GCTTAAAC-AACAGAAATTTAT | loxF8R | 3 | loxF8L | 4 | GCTTAAAC/GTTTAAGC | 7 |
| chr10 | 61042075 | ATAAATTTCTGTT-GTTTAAGC-CACCCAATCAGAG | loxF8L | 4 | loxF8R | 3 | GCTTAAAC/GTTTAAGC | 7 |
| chr13 | 37670180 | ATAAATTTCTGTT-GTTTAAGC-CACCCAATCTATG | loxF8L | 4 | loxF8R | 3 | GCTTAAAC/GTTTAAGC | 7 |
| chr14 | 97051701 | CATAGATTGGGTG-GCTTAAAC-AACAGAAATTTAT | loxF8R | 3 | loxF8L | 4 | GCTTAAAC/GTTTAAGC | 7 |
| chr1 | 179235530 | ATAAATTTCTGTT-GTTTAAGC-CACCCAATCTGTG | loxF8L | 4 | loxF8R | 3 | GCTTAAAC/GTTTAAGC | 7 |
| chr20 | 21014150 | CACAGATTGGGTG-GCTTAAAC-AACAGAAATTTAT | loxF8R | 3 | loxF8L | 4 | GCTTAAAC/GTTTAAGC | 7 |
| chr2 | 207259558 | CACAGATTGGGTG-GCTTAAAC-AACAGAAATTTAT | loxF8R | 3 | loxF8L | 4 | GCTTAAAC/GTTTAAGC | 7 |
| chr5 | 78201147 | ATAAATTTCTGTT-GTTTAAGC-CACCCAATCTGTG | loxF8L | 4 | loxF8R | 3 | GCTTAAAC/GTTTAAGC | 7 |
| chr8 | 57883620 | ATAAATTTCTGTT-GTTTAAGC-CACCCAATCTGTG | loxF8L | 4 | loxF8R | 3 | GCTTAAAC/GTTTAAGC | 7 |
| chr9 | 3018756 | CACAGATTGGGTG-GCTTAAAC-AACAGAAATTTAT | loxF8R | 3 | loxF8L | 4 | GCTTAAAC/GTTTAAGC | 7 |
| chr12 | 33939095 | ATAAATTTCTGTT-GTTTAAGC-CACCCAGTCTAAG | loxF8L | 4 | loxF8R | 3 | GCTTAAAC/GTTTAAGC | 7 |
| chr5 | 54279299 | CTTAGACTGGGTG-GCTTAAAC-AACAGAAATTTAT | loxF8R | 3 | loxF8L | 4 | GCTTAAAC/GTTTAAGC | 7 |
| chr2 | 20429514 | CTGAGACTGGGTG-GCTTAAAC-AACAGAAATTTAT | loxF8R | 3 | loxF8L | 4 | GCTTAAAC/GTTTAAGC | 7 |
| chr22 | 26173660 | CTTAGATAGGGTG-GCTTAAAC-AACAGAAATTTAT | loxF8R | 3 | loxF8L | 4 | GCTTAAAC/GTTTAAGC | 7 |
| chr5 | 101867922 | CACAGATTGAGTG-GCTTAAAC-AACAGAAATTTAT | loxF8R | 3 | loxF8L | 4 | GCTTAAAC/GTTTAAGC | 7 |
| chr7 | 48324129 | CACAGATTGAGTG-GCTTAAAC-AACAGAAATTTAT | loxF8R | 3 | loxF8L | 4 | GCTTAAAC/GTTTAAGC | 7 |
| chr3 | 19286513 | ATAAATTTCTGTT-GTTTAAGC-CACTCAGTCTGAG | loxF8L | 4 | loxF8R | 3 | GCTTAAAC/GTTTAAGC | 7 |
| chr11 | 87382246 | CAAAGACTGAGTG-GCTTAAAC-AACAGAAATTTAT | loxF8R | 3 | loxF8L | 4 | GCTTAAAC/GTTTAAGC | 7 |
| chrX | 69552808 | CAAAGACTGTGTA-GCTTAAAC-AACAGAAATTTAT | loxF8R | 3 | loxF8L | 4 | GCTTAAAC/GTTTAAGC | 7 |
| chr3 | 47561519 | ATAAATTTGTGCT-GTTTAAGC-CACCCAGTCTGTG | loxF8L | 3 | loxF8R | 4 | GCTTAAAC/GTTTAAGC | 7 |
| chr21 | 23811850 | TATAGACTGTGTG-GCTTAAAC-AACACAAATTTAT | loxF8R | 4 | loxF8L | 3 | GCTTAAAC/GTTTAAGC | 7 |
| chr17 | 16248818 | CATAGACTGGGTG-GCTTAAAC-AACACAAATTTAT | loxF8R | 4 | loxF8L | 3 | GCTTAAAC/GTTTAAGC | 7 |
| chr14 | 59798384 | CACAGATTTAGTG-GCTTAAAC-AACACAAATTTAT | loxF8R | 4 | loxF8L | 3 | GCTTAAAC/GTTTAAGC | 7 |

| | | | | | | | | |
|---|---|---|---|---|---|---|---|---|
| chr9 | 115987708 | ATAAATTTGTGTT-GTTTAAGC-CACTAAATCTGTG | loxF8L | 3 | loxF8R | 4 | GCTTAAAC/GTTTAAGC | 7 |
| chr4 | 14231063 | ATAAATTTTTGTT-GTTTAAGC-CACACAGTCTATG | loxF8L | 4 | loxF8R | 3 | GCTTAAAC/GTTTAAGC | 7 |
| chr8 | 29363935 | CACAGACTGTGTG-GCTTAAAC-AACAAAAATTTAT | loxF8R | 3 | loxF8L | 4 | GCTTAAAC/GTTTAAGC | 7 |
| chr3 | 110308508 | CACAGATTGGGTG-GCTTAAAC-AACAAAAATTTAT | loxF8R | 3 | loxF8L | 4 | GCTTAAAC/GTTTAAGC | 7 |
| chr3 | 118390460 | ATAAATTTTTGTT-GTTTAAGC-CACCCAGTCTTTG | loxF8L | 4 | loxF8R | 3 | GCTTAAAC/GTTTAAGC | 7 |
| chr12 | 38215852 | ATAAATTTTTGTT-GTTTAAGC-CACCCATTCTGAG | loxF8L | 4 | loxF8R | 3 | GCTTAAAC/GTTTAAGC | 7 |
| chr7 | 24377068 | ATAACTTTCTGGT-GTTTAAGC-CAAGCAATCTGAG | loxF8L | 4 | loxF8R | 3 | GCTTAAAC/GTTTAAGC | 7 |
| chr20 | 8957887 | CACAGAATGTGTG-GCTTAAAC-AACAGAAACTTAT | loxF8R | 3 | loxF8R | 4 | GCTTAAAC/GTTTAAGC | 7 |
| chr11 | 106905321 | CATAGATTGGGTG-GCTTAAAC-AACAGAAACTTAT | loxF8R | 3 | loxF8R | 4 | GCTTAAAC/GTTTAAGC | 7 |
| chr10 | 123296960 | ATAAGTTTCTGTT-GTTTAAGC-CACTCAATCTATG | loxF8R | 4 | loxF8R | 3 | GCTTAAAC/GTTTAAGC | 7 |
| chr6 | 83470819 | ATAAGTTTGTGTT-GTTTAAGC-CACCCAGTCTGTG | loxF8R | 3 | loxF8R | 4 | GCTTAAAC/GTTTAAGC | 7 |
| chr17 | 62918593 | CATAGACTGTGTG-GCTTAAAC-AACAGAGATGTAT | loxF8R | 3 | loxF8L | 4 | GCTTAAAC/GTTTAAGC | 7 |
| chr5 | 13398154 | ATATATTTGTGGT-GTTTAAGC-CACCCAGTCTAGG | loxF8L | 3 | loxF8R | 4 | GCTTAAAC/GTTTAAGC | 7 |
| --- | | | | | | | | 0 |
| chr2 | 45957213 | CAAAGATTTTTTG-TGTTTTCC-TCCACTGTTTTAT | loxF8R | 3 | loxF8L | 2 | GGAAAACA/TGTTTTCC | 5 |
| chr14 | 44029073 | ACAAATGTGTGAA-TGTTTTCC-CACATACACTAAG | loxF8L | 3 | loxF8R | 4 | GGAAAACA/TGTTTTCC | 7 |
| --- | | | | | | | | 0 |
| chr2 | 80869463 | ATAAATTTGTGTT-GTTTTAAC-CACACAGTTTTTG | loxF8L | 3 | loxF8R | 3 | GTTAAAAC/GTTTTAAC | 6 |
| chr2 | 193039018 | ATAGATATGTGGA-GTTAAAAC-TCCCATGATTAAT | loxF8L | 2 | loxF8L | 4 | GTTAAAAC/GTTTTAAC | 6 |
| chr9 | 83941180 | ATAAATGTCTGTT-GTTTTAAC-CACTAAATTTTAG | loxF8L | 4 | loxF8R | 3 | GTTAAAAC/GTTTTAAC | 7 |
| chr5 | 30068662 | CACAGACTGTGTG-GTTAAAAC-AACAGAAATTTAT | loxF8R | 3 | loxF8L | 4 | GTTAAAAC/GTTTTAAC | 7 |
| chr12 | 77078592 | ATAAATTTCTGTT-GTTAAAAC-CACCCAATCTGTG | loxF8L | 4 | loxF8R | 3 | GTTAAAAC/GTTTTAAC | 7 |
| --- | | | | | | | | 0 |
| chr10 | 47180030 | AAAAGTCTGTGTA-TATATTTA-TACATATATTTAT | loxF8L | 3 | loxF8L | 3 | TAAATATA/TATATTTA | 6 |
| chrX | 140345371 | ATATATGTGTGCA-TAAATATA-TACACATATATAT | loxF8L | 3 | loxF8L | 3 | TAAATATA/TATATTTA | 6 |
| chr21 | 33502034 | TTCAATGTATGGA-TATATTTA-TCCATATATTTAT | loxF8L | 4 | loxF8L | 2 | TAAATATA/TATATTTA | 6 |
| chr15 | 94244057 | ATACATATGTGTA-TATATTTA-TGCACACATATGT | loxF8L | 3 | loxF8L | 4 | TAAATATA/TATATTTA | 7 |
| chr15 | 94243980 | ATACATATGTGTA-TATATTTA-TGCACACATATGT | loxF8L | 3 | loxF8L | 4 | TAAATATA/TATATTTA | 7 |
| chr14 | 82925533 | ATAAAATTCTGAG-TATATTTA-CACACAAATGTAG | loxF8R | 4 | loxF8R | 3 | TAAATATA/TATATTTA | 7 |
| chr3 | 94015986 | ATAAACATGAGAA-TATATTTA-TATACAATCTTAT | loxF8L | 4 | loxF8R | 3 | TAAATATA/TATATTTA | 7 |
| chr11 | 21057474 | TTATGCTTGTTTG-TAAATATA-TCAATATATTTAT | loxF8R | 4 | loxF8L | 3 | TAAATATA/TATATTTA | 7 |
| chr1 | 187718091 | CTAATATTTTATT-TAAATATA-TCTTCATATTTAT | loxF8R | 4 | loxF8L | 3 | TAAATATA/TATATTTA | 7 |
| chr1 | 199518390 | ATAAATATGTACA-TAAATATA-TATACAATTTTAC | loxF8L | 3 | loxF8R | 4 | TAAATATA/TATATTTA | 7 |
| chr20 | 13685563 | ATAAATATGTATA-TAAATATA-CACACATACTTGA | loxF8L | 3 | loxF8R | 4 | TAAATATA/TATATTTA | 7 |
| chr8 | 100052260 | ATAAATGTATATA-TATATTTA-TATACATATTTAT | loxF8L | 4 | loxF8L | 3 | TAAATATA/TATATTTA | 7 |
| chr1 | 95340539 | ATAGATATATGCA-TATATTTA-TAAACATATTTAT | loxF8L | 4 | loxF8L | 3 | TAAATATA/TATATTTA | 7 |
| chr16 | 8103581 | GTAAATATGTTGA-TAAATATA-AACAATGATTTAT | loxF8L | 3 | loxF8L | 4 | TAAATATA/TATATTTA | 7 |
| chrX | 26082270 | ATATATCTATAGA-TATATTTA-TAGATACATTTAT | loxF8L | 3 | loxF8L | 4 | TAAATATA/TATATTTA | 7 |
| chr8 | 137420094 | ATAAATGTGTATA-TATATTTA-TATACATATATAT | loxF8L | 3 | loxF8L | 4 | TAAATATA/TATATTTA | 7 |
| chr15 | 94243860 | ATACATATGTGTA-TATATTTA-TGTACACATATAT | loxF8L | 3 | loxF8L | 4 | TAAATATA/TATATTTA | 7 |
| chr7 | 8279814 | ATATATATGTGTA-TAAATATA-CACACACATATAT | loxF8L | 3 | loxF8L | 4 | TAAATATA/TATATTTA | 7 |
| chr4 | 92583888 | ATATATATGTGTA-TATATTTA-TATACACATATAT | loxF8L | 3 | loxF8L | 4 | TAAATATA/TATATTTA | 7 |
| chr2 | 225018780 | CTAAGATGTCATG-TAAATATA-TATACAGATATAT | loxF8R | 4 | loxF8L | 3 | TAAATATA/TATATTTA | 7 |
| chr18 | 67105524 | ATATATGTGTGTA-TATATTTA-TATACACATATAT | loxF8L | 3 | loxF8L | 4 | TAAATATA/TATATTTA | 7 |
| chr4 | 52733423 | ATATATGTGTGTA-TATATTTA-TACACATATATAC | loxF8L | 3 | loxF8L | 4 | TAAATATA/TATATTTA | 7 |
| chrX | 2166444 | CACACACTGTGTG-TATATTTA-CACACAATGTGTG | loxF8R | 4 | loxF8R | 3 | TAAATATA/TATATTTA | 7 |
| chrY | 2166444 | CACACACTGTGTG-TATATTTA-CACACAATGTGTG | loxF8R | 4 | loxF8R | 3 | TAAATATA/TATATTTA | 7 |
| --- | | | | | | | | 0 |
| chr7 | 110812069 | ATAAACCTATAGA-TACATATA-TCCACATATGTAT | loxF8L | 3 | loxF8L | 2 | TACATATA/TATATGTA | 5 |
| chr10 | 19023211 | ATAAATTTGTATA-TATATGTA-TGCACATATATAT | loxF8L | 3 | loxF8L | 3 | TACATATA/TATATGTA | 6 |
| chr4 | 26734925 | ATATATGTGTGTA-TATATGTA-TACACATATGTAT | loxF8L | 3 | loxF8L | 3 | TACATATA/TATATGTA | 6 |
| chr9 | 35880162 | ATACATATGTGTA-TACATATA-TACACATATGTAT | loxF8L | 3 | loxF8L | 3 | TACATATA/TATATGTA | 6 |
| chr8 | 85566526 | ATATATATGTGTA-TACATATA-TACACACATATAT | loxF8L | 3 | loxF8L | 3 | TACATATA/TATATGTA | 6 |
| chr14 | 81549734 | ATATATATGTGTA-TATATGTA-TACACATATATAT | loxF8L | 3 | loxF8L | 3 | TACATATA/TATATGTA | 6 |
| chr8 | 15232780 | ATATATGTGTGTA-TACATATA-TACACACATATAT | loxF8L | 3 | loxF8L | 3 | TACATATA/TATATGTA | 6 |
| chr5 | 114517608 | AAAAATCTGTGTA-TATATGTA-TGTACATATGTAT | loxF8L | 2 | loxF8L | 4 | TACATATA/TATATGTA | 6 |
| chr4 | 179679287 | ATATATCTGTGTA-TATATGTA-TACACACAGATAT | loxF8L | 2 | loxF8L | 4 | TACATATA/TATATGTA | 6 |
| chr14 | 58240130 | TTAAGAATGTTTG-TATATGTA-TCCAAAGATCATT | loxF8R | 3 | loxF8L | 4 | TACATATA/TATATGTA | 7 |
| chr4 | 32007507 | ACACATCTATGTA-TACATATA-TACACATGTTTAT | loxF8L | 4 | loxF8L | 3 | TACATATA/TATATGTA | 7 |
| chr5 | 142715191 | ACATATATGTGTA-TACATATA-TACACATATATAT | loxF8L | 4 | loxF8L | 3 | TACATATA/TATATGTA | 7 |
| chr12 | 81625126 | ACATATGTGTGTA-TACATATA-TACACACATATAT | loxF8L | 4 | loxF8L | 3 | TACATATA/TATATGTA | 7 |
| chr2 | 203960312 | ATATATCTATGCA-TACATATA-TACACACATATGT | loxF8L | 3 | loxF8L | 4 | TACATATA/TATATGTA | 7 |
| chr3 | 28731842 | ATAAACGTGTGAA-TACATATA-TAGATATATTTAT | loxF8L | 3 | loxF8L | 4 | TACATATA/TATATGTA | 7 |
| chr13 | 105089171 | ATATATATGTGTA-TACATATA-TGTATATATTTAT | loxF8L | 3 | loxF8L | 4 | TACATATA/TATATGTA | 7 |
| chrX | 69168353 | ATAAGTGTATGTA-TACATATA-TATACAGATATAT | loxF8L | 4 | loxF8L | 3 | TACATATA/TATATGTA | 7 |
| chr21 | 22216484 | ATATATGTGTGTA-TATATGTA-TGCATATATGTAT | loxF8L | 3 | loxF8L | 4 | TACATATA/TATATGTA | 7 |
| chr6 | 132015854 | ATACATATATGTA-TACATATA-TACACACATATAT | loxF8L | 4 | loxF8L | 3 | TACATATA/TATATGTA | 7 |
| chr2 | 155412366 | ATATATATGTGTA-TATATGTA-TACATATATGTAT | loxF8L | 3 | loxF8L | 4 | TACATATA/TATATGTA | 7 |
| chr8 | 126870329 | ATATATATGTGTA-TACATATA-TACATATATGTAT | loxF8L | 3 | loxF8L | 4 | TACATATA/TATATGTA | 7 |
| chr19 | 28159667 | ATACATATGTAGA-TACATATA-AACACACATGTAT | loxF8L | 3 | loxF8L | 4 | TACATATA/TATATGTA | 7 |
| chr2 | 191795593 | ATACATATGTATA-TACATATA-TACACACATATAT | loxF8L | 4 | loxF8L | 3 | TACATATA/TATATGTA | 7 |
| chr21 | 26311754 | ATACATATGTGCA-TACATATA-TACACATACATAT | loxF8L | 3 | loxF8L | 4 | TACATATA/TATATGTA | 7 |
| chr18 | 64143545 | CACATATTATGTG-TACATATA-TACACATATGTAT | loxF8R | 4 | loxF8L | 3 | TACATATA/TATATGTA | 7 |
| chr18 | 52497362 | ATACATATGTGTA-TATATGTA-TACACACATATAC | loxF8L | 3 | loxF8L | 4 | TACATATA/TATATGTA | 7 |
| chr18 | 52497392 | ATACATATGTGTA-TATATGTA-TACACACATATAC | loxF8L | 3 | loxF8L | 4 | TACATATA/TATATGTA | 7 |
| chr4 | 101470595 | ATACATATGTGTA-TACATATA-TACATATATATAT | loxF8L | 3 | loxF8L | 4 | TACATATA/TATATGTA | 7 |
| chrX | 116116141 | ATATATGTGTATA-TATATGTA-TACACATATGTAT | loxF8L | 4 | loxF8L | 3 | TACATATA/TATATGTA | 7 |
| chr13 | 76143827 | ATATAGCTGTATA-TACATATA-TACACATATGTAT | loxF8L | 4 | loxF8L | 3 | TACATATA/TATATGTA | 7 |
| chr10 | 93235316 | ATATATATGTGTG-TATATGTA-TACACACATGTAT | loxF8L | 4 | loxF8L | 3 | TACATATA/TATATGTA | 7 |
| chr14 | 42252603 | ATATATGTGTATA-TACATATA-TACACACATGTAT | loxF8L | 4 | loxF8L | 3 | TACATATA/TATATGTA | 7 |
| chr4 | 80864465 | ATACCTATGTGTA-TACATATA-TACACATATATAT | loxF8L | 4 | loxF8L | 3 | TACATATA/TATATGTA | 7 |
| chr5 | 15047078 | ATATATCCATGGA-TACATATA-TATACATATCTAT | loxF8L | 3 | loxF8L | 4 | TACATATA/TATATGTA | 7 |
| chr12 | 39712612 | ATAGATATGTGTA-TATATGTA-TACACATTTATAT | loxF8L | 3 | loxF8L | 4 | TACATATA/TATATGTA | 7 |
| chr13 | 88175344 | ATATATATATGTA-TATATGTA-TACACATATATAT | loxF8L | 4 | loxF8L | 3 | TACATATA/TATATGTA | 7 |
| chr4 | 157830392 | ATATATATATGTA-TACATATA-TACACATATATAT | loxF8L | 4 | loxF8L | 3 | TACATATA/TATATGTA | 7 |
| chr1 | 58730415 | ATATATTTGTGCA-TATATGTA-TACATATATATAT | loxF8L | 3 | loxF8L | 4 | TACATATA/TATATGTA | 7 |
| chr16 | 82116139 | ATATATGTGTGTA-TATATGTA-TTTACATATATAT | loxF8L | 3 | loxF8L | 4 | TACATATA/TATATGTA | 7 |

| | | | | | | | | |
|---|---|---|---|---|---|---|---|---|
| chr11 | 24602285 | ATATATATGTACA-TACATATA-TACACACATATAT | loxF8L | 4 | loxF8L | 3 | TACATATA/TATATGTA | 7 |
| chr5 | 68263168 | ATATATCTATATA-TACATATA-TCTACATATATAT | loxF8L | 4 | loxF8L | 3 | TACATATA/TATATGTA | 7 |
| chr13 | 70778203 | TTTAAATTGTGTG-TACATATA-TATACATATATAT | loxF8R | 3 | loxF8L | 4 | TACATATA/TATATGTA | 7 |
| chr4 | 179679229 | ATATATATGTATA-TATATGTA-TACACACATATAT | loxF8L | 4 | loxF8L | 3 | TACATATA/TATATGTA | 7 |
| chr16 | 17098144 | ATATATATGTGTA-TATATGTA-TATACATATATAT | loxF8L | 3 | loxF8L | 4 | TACATATA/TATATGTA | 7 |
| chr4 | 70583360 | ATATATATGTGTA-TATATGTA-TATACATATATAT | loxF8L | 3 | loxF8L | 4 | TACATATA/TATATGTA | 7 |
| chr5 | 157093086 | ATATATATGTGTA-TACATATA-TATACATATATAT | loxF8L | 3 | loxF8L | 4 | TACATATA/TATATGTA | 7 |
| chr11 | 23895048 | ATATATATGTGCA-TACATATA-CACACATATATAT | loxF8L | 3 | loxF8L | 4 | TACATATA/TATATGTA | 7 |
| chr7 | 43004541 | ATATATTTATGTA-TACATATA-TACACATATATAT | loxF8L | 4 | loxF8L | 3 | TACATATA/TATATGTA | 7 |
| chr18 | 50789787 | ATATATATGTGTA-TATATGTA-TACATACATATAT | loxF8L | 3 | loxF8L | 4 | TACATATA/TATATGTA | 7 |
| chr18 | 50789815 | ATATATATGTGTA-TATATGTA-TACATACATATAT | loxF8L | 3 | loxF8L | 4 | TACATATA/TATATGTA | 7 |
| chr10 | 19973327 | ATATATATGTGTA-TATATGTA-TACTCACATATAT | loxF8L | 3 | loxF8L | 4 | TACATATA/TATATGTA | 7 |
| chr12 | 69041504 | ATATATATGTGTA-TACATATA-TATACACATATAT | loxF8L | 3 | loxF8L | 4 | TACATATA/TATATGTA | 7 |
| chr4 | 11467136 | ATATATGTGTATA-TATATGTA-TACACATATATAT | loxF8L | 4 | loxF8L | 3 | TACATATA/TATATGTA | 7 |
| chr1 | 234995616 | ATATATATGTGTA-TATATGTA-TATATAGATATAT | loxF8L | 3 | loxF8L | 4 | TACATATA/TATATGTA | 7 |
| chr6 | 55986850 | ATATATCTATATA-TACATATA-TACACATATATAT | loxF8L | 4 | loxF8L | 3 | TACATATA/TATATGTA | 7 |
| chr3 | 144966308 | ATATATCCATGGA-TATATGTA-TACACATATATAC | loxF8L | 3 | loxF8L | 4 | TACATATA/TATATGTA | 7 |
| chr4 | 82222988 | ATATGTGTGTGTA-TATATGTA-TACATAGATATAT | loxF8L | 4 | loxF8L | 3 | TACATATA/TATATGTA | 7 |
| chr4 | 179679431 | ATATATGTGTGTA-TATATGTA-TACATACATATAT | loxF8L | 3 | loxF8L | 4 | TACATATA/TATATGTA | 7 |
| chr4 | 179679403 | ATATATGTGTGTA-TATATGTA-TACATACATATAT | loxF8L | 3 | loxF8L | 4 | TACATATA/TATATGTA | 7 |
| chr6 | 64551155 | ATATATGTGTATA-TACATATA-TACACACATATAT | loxF8L | 4 | loxF8L | 3 | TACATATA/TATATGTA | 7 |
| chr9 | 31039478 | ATGAGTGTGTGTA-TACATATA-TACACACATATAT | loxF8L | 4 | loxF8L | 3 | TACATATA/TATATGTA | 7 |
| chr15 | 44140070 | ATATATGTGTGTA-TACATATA-TACACACATATAG | loxF8L | 3 | loxF8L | 4 | TACATATA/TATATGTA | 7 |
| chr6 | 75757037 | GTATATATGTGTA-TATATGTA-TACACACATATAT | loxF8L | 4 | loxF8L | 3 | TACATATA/TATATGTA | 7 |
| chr6 | 75756940 | GTATATATGTGTA-TATATGTA-TACACACATATAT | loxF8L | 4 | loxF8L | 3 | TACATATA/TATATGTA | 7 |
| chr10 | 57150222 | ATCAATCAATGGA-TACATATA-TCAACAGAATGAA | loxF8L | 3 | loxF8L | 4 | TACATATA/TATATGTA | 7 |
| chr3 | 132147151 | GTATATGTGTGTA-TATATGTA-TACACAGATGCAT | loxF8L | 4 | loxF8L | 3 | TACATATA/TATATGTA | 7 |
| --- | | | | | | | | 0 |
| chr1 | 172873333 | ATAAATCTATGTA-TGTATCTA-TCTACATATTTGT | loxF8L | 2 | loxF8L | 3 | TAGATACA/TGTATCTA | 5 |
| chr4 | 36303438 | AAAAATCTGAGGA-TGTATCTA-TCTATATATTTTT | loxF8L | 2 | loxF8L | 4 | TAGATACA/TGTATCTA | 6 |
| chr9 | 91668437 | ATAAGAAGGTGTG-TGTATCTA-TCTATAGAATTTT | loxF8R | 3 | loxF8L | 4 | TAGATACA/TGTATCTA | 7 |
| chrX | 49754269 | TTAAATGCATGGA-TAGATACA-TGCAGATATTTAT | loxF8L | 4 | loxF8L | 3 | TAGATACA/TGTATCTA | 7 |
| chr4 | 30277984 | ATATATGTGTAGA-TAGATACA-TATACACATATAT | loxF8L | 3 | loxF8L | 4 | TAGATACA/TGTATCTA | 7 |
| --- | | | | | | | | 0 |
| chr4 | 55780475 | ATAAACATATGGA-TTTGTATA-CACACAATCATTC | loxF8L | 3 | loxF8R | 3 | TATACAAA/TTTGTATA | 6 |
| chr4 | 146592794 | AGATATTTGTGAA-TTTGTATA-CACATAATCATAG | loxF8L | 4 | loxF8R | 2 | TATACAAA/TTTGTATA | 6 |
| chrY | 15708080 | ATATATCTGTGGG-TATACAAA-CACACATTAATAT | loxF8L | 2 | loxF8R | 4 | TATACAAA/TTTGTATA | 6 |
| chr6 | 151860514 | ATATATGTGTGTA-TATACAAA-CACACAGACATAC | loxF8L | 3 | loxF8R | 4 | TATACAAA/TTTGTATA | 7 |
| --- | | | | | | | | 0 |
| chrX | 44149990 | ATAAATACGTGTA-TATACACA-CACACAAACTTTT | loxF8L | 3 | loxF8R | 3 | TATACACA/TGTGTATA | 6 |
| chr2 | 75869383 | ATATATCTATGTA-TGTGTATA-TCCCCAACTTTAT | loxF8L | 3 | loxF8L | 3 | TATACACA/TGTGTATA | 6 |
| chr10 | 99814066 | ATATATATGTGTA-TGTGTATA-TACACATATATAT | loxF8L | 3 | loxF8L | 3 | TATACACA/TGTGTATA | 6 |
| chr7 | 66236399 | ATATATATATGTA-TATACACA-CACACACTTTTAG | loxF8L | 4 | loxF8R | 2 | TATACACA/TGTGTATA | 6 |
| chr9 | 17356975 | TTATGAGTGTGTG-TGTGTATA-TATACATATTTGT | loxF8R | 3 | loxF8L | 4 | TATACACA/TGTGTATA | 7 |
| chr22 | 35195660 | ACACATATGTGTA-TATACACA-TACACACATATAT | loxF8L | 4 | loxF8L | 3 | TATACACA/TGTGTATA | 7 |
| chrX | 5692950 | ACACATGTGTGTA-TATACACA-TACACACATATAT | loxF8L | 4 | loxF8L | 3 | TATACACA/TGTGTATA | 7 |
| chr9 | 26104796 | ACATATATGTGTA-TATACACA-TACACACATATAT | loxF8L | 4 | loxF8L | 3 | TATACACA/TGTGTATA | 7 |
| chr10 | 17968044 | CTTTGTTTGTGTG-TGTGTATA-TACACACATATGT | loxF8R | 3 | loxF8L | 4 | TATACACA/TGTGTATA | 7 |
| chr4 | 12648826 | ATATATGTGTGTA-TATACACA-TACACACATATGT | loxF8L | 3 | loxF8L | 4 | TATACACA/TGTGTATA | 7 |
| chr15 | 46227786 | AGAAATGTGTGTA-TATACACA-CACACATACATAT | loxF8L | 3 | loxF8R | 4 | TATACACA/TGTGTATA | 7 |
| chr7 | 4571336 | AGATATATGTGTA-TGTGTATA-TATACAGATATAT | loxF8L | 4 | loxF8L | 3 | TATACACA/TGTGTATA | 7 |
| chr3 | 114702437 | ATAAACCTATAAA-TATACACA-TACACAGATGAAT | loxF8L | 4 | loxF8L | 3 | TATACACA/TGTGTATA | 7 |
| chr5 | 120576933 | ATAAATATATGTA-TATACACA-TATACACATATAT | loxF8L | 3 | loxF8L | 4 | TATACACA/TGTGTATA | 7 |
| chr4 | 75495888 | ATTTGTCTGTGTA-TATACACA-CACACATATTTAT | loxF8L | 4 | loxF8L | 3 | TATACACA/TGTGTATA | 7 |
| chrX | 21731695 | ATAAATGTGTATA-TGTGTATA-TGTACACATTAAT | loxF8L | 3 | loxF8L | 4 | TATACACA/TGTGTATA | 7 |
| chr14 | 94188568 | ATATATATGTGTG-TATACACA-CACACACACTTAT | loxF8L | 4 | loxF8R | 3 | TATACACA/TGTGTATA | 7 |
| chr13 | 76480478 | ATACATATGTGCA-TGTGTATA-TACACATACATAT | loxF8L | 3 | loxF8L | 4 | TATACACA/TGTGTATA | 7 |
| chr13 | 76480696 | ATACATATGTGCA-TGTGTATA-TACACATACATAT | loxF8L | 3 | loxF8L | 4 | TATACACA/TGTGTATA | 7 |
| chr13 | 76480748 | ATACATATGTGCA-TGTGTATA-TACACATACATAT | loxF8L | 3 | loxF8L | 4 | TATACACA/TGTGTATA | 7 |
| chr13 | 76480722 | ATACATATGTGCA-TGTGTATA-TACACATACATAT | loxF8L | 3 | loxF8L | 4 | TATACACA/TGTGTATA | 7 |
| chr5 | 20542436 | CAATGTGTGTGTG-TATACACA-TACACATATGTAT | loxF8R | 4 | loxF8L | 3 | TATACACA/TGTGTATA | 7 |
| chr6 | 44330579 | TAAAGAGTGTGTG-TGTGTATA-CACACATATGTAT | loxF8R | 3 | loxF8L | 4 | TATACACA/TGTGTATA | 7 |
| chr4 | 83975351 | ATATATATATGTA-TATACACA-CCCACACATGTAT | loxF8L | 4 | loxF8L | 3 | TATACACA/TGTGTATA | 7 |
| chr9 | 112289772 | CTTATTTTGTGTG-TGTGTATA-TATACATATATAT | loxF8R | 3 | loxF8L | 4 | TATACACA/TGTGTATA | 7 |
| chr9 | 26104891 | ATATATGTGTGTA-TGTGTATA-TATACATATATAT | loxF8L | 3 | loxF8L | 4 | TATACACA/TGTGTATA | 7 |
| chr9 | 26104943 | ATATATATGTGTA-TGTGTATA-TATACATATATAT | loxF8L | 3 | loxF8L | 4 | TATACACA/TGTGTATA | 7 |
| chr9 | 26104917 | ATATATATGTGTA-TGTGTATA-TATACATATATAT | loxF8L | 3 | loxF8L | 4 | TATACACA/TGTGTATA | 7 |
| chr10 | 56226231 | ATATATGTGTATA-TATACACA-TGCACATATATAT | loxF8L | 4 | loxF8L | 3 | TATACACA/TGTGTATA | 7 |
| chr1 | 70692648 | CCACGTGTGTGTG-TGTGTATA-TACACATATATAT | loxF8R | 4 | loxF8L | 3 | TATACACA/TGTGTATA | 7 |
| chr8 | 90233529 | ATATATATGTGTA-TATACACA-CACACACATATAT | loxF8L | 3 | loxF8L | 4 | TATACACA/TGTGTATA | 7 |
| chrX | 8061812 | ATATATGTGTGTG-TGTGTATA-TACACATATATAT | loxF8L | 4 | loxF8L | 3 | TATACACA/TGTGTATA | 7 |
| chr13 | 108699103 | ATATATATGTGTG-TGTGTATA-TACACATATATAT | loxF8L | 4 | loxF8L | 3 | TATACACA/TGTGTATA | 7 |
| chr21 | 21264944 | ATATATATGTGTA-TGTGTATA-TATACACATATAT | loxF8L | 3 | loxF8L | 4 | TATACACA/TGTGTATA | 7 |
| chr4 | 156161766 | ATATATATGTGTA-TGTGTATA-TATACACATATAT | loxF8L | 3 | loxF8L | 4 | TATACACA/TGTGTATA | 7 |
| chr4 | 59779570 | ATATATGTGTGCA-TATACACA-CACACATATATAT | loxF8L | 3 | loxF8L | 4 | TATACACA/TGTGTATA | 7 |
| chr14 | 66372138 | CTAAGAATATGTT-TGTGTATA-TATACACATATAT | loxF8R | 3 | loxF8L | 4 | TATACACA/TGTGTATA | 7 |
| chr12 | 97621887 | ATATATGTGTGTA-TGTGTATA-TATACACATATAT | loxF8L | 3 | loxF8L | 4 | TATACACA/TGTGTATA | 7 |
| chr12 | 97621857 | ATATATGTGTGTA-TGTGTATA-TATACACATATAT | loxF8L | 3 | loxF8L | 4 | TATACACA/TGTGTATA | 7 |
| chr12 | 97621827 | ATATATGTGTGTA-TGTGTATA-TATACACATATAT | loxF8L | 3 | loxF8L | 4 | TATACACA/TGTGTATA | 7 |
| chr6 | 13131162 | ATATATGTGTGTA-TATACACA-CACACACACATAT | loxF8L | 3 | loxF8R | 4 | TATACACA/TGTGTATA | 7 |
| chr3 | 23980756 | ATATATGTGTGTG-TGTGTATA-TACACACATATAT | loxF8L | 4 | loxF8L | 3 | TATACACA/TGTGTATA | 7 |
| chr3 | 73885399 | ATGTATCTGTATA-TATACACA-CACATAATTTTAA | loxF8L | 4 | loxF8R | 3 | TATACACA/TGTGTATA | 7 |
| --- | | | | | | | | 0 |
| chr5 | 175142056 | ATAAAACTAAGGA-TGTATATA-TACACACATATAT | loxF8L | 3 | loxF8L | 3 | TATATACA/TGTATATA | 6 |
| chrY | 17944115 | ATAAATGTGTTTA-TATATACA-CACACAAACATAT | loxF8L | 3 | loxF8R | 3 | TATATACA/TGTATATA | 6 |
| chrY | 17961828 | ATATGTTTGTGTG-TGTATATA-TAAACACATTTAT | loxF8R | 3 | loxF8L | 3 | TATATACA/TGTATATA | 6 |
| chr5 | 15265388 | ATACATATGTGTA-TGTATATA-TACACACATATAT | loxF8L | 3 | loxF8L | 3 | TATATACA/TGTATATA | 6 |

| | | | | | | | | |
|---|---|---|---|---|---|---|---|---|
| chr5 | 15265422 | ATACATATGTGTA-TGTATATA-TACACACATATAT | loxF8L | 3 | loxF8L | 3 | TATATACA/TGTATATA | 6 |
| chr5 | 15265286 | ATACATATGTGTA-TGTATATA-TACACACATATAT | loxF8L | 3 | loxF8L | 3 | TATATACA/TGTATATA | 6 |
| chr5 | 15265354 | ATACATATGTGTA-TGTATATA-TACACACATATAT | loxF8L | 3 | loxF8L | 3 | TATATACA/TGTATATA | 6 |
| chr5 | 15265320 | ATACATATGTGTA-TGTATATA-TACACACATATAT | loxF8L | 3 | loxF8L | 3 | TATATACA/TGTATATA | 6 |
| chr18 | 27670451 | ATATATATGTGTA-TATATACA-TACACACATATAT | loxF8L | 3 | loxF8L | 3 | TATATACA/TGTATATA | 6 |
| chr5 | 160091985 | ATATATGTGTGTA-TATATACA-TACACGGATATAT | loxF8L | 3 | loxF8L | 3 | TATATACA/TGTATATA | 6 |
| chrX | 56745410 | ATACATATATGTA-TATATACA-TACACAGATTTTT | loxF8L | 4 | loxF8L | 2 | TATATACA/TGTATATA | 6 |
| chr10 | 130420690 | ATAAATCTATGCA-TGTATATA-TACACATGTATAT | loxF8L | 2 | loxF8L | 4 | TATATACA/TGTATATA | 6 |
| chr16 | 56711107 | ATAAATGTGTGTA-TATATACA-TATACATATATAT | loxF8L | 2 | loxF8L | 4 | TATATACA/TGTATATA | 6 |
| chr5 | 175783620 | ATAAATGTGTGTA-TGTATATA-TATACATATATAT | loxF8L | 2 | loxF8L | 4 | TATATACA/TGTATATA | 6 |
| chrX | 113930917 | CTATGAATGTGTG-TGTATATA-TGCATACATATAT | loxF8R | 2 | loxF8L | 4 | TATATACA/TGTATATA | 6 |
| chr5 | 62980634 | ATATATGTATGTA-TGTATATA-CACACAATTTTTT | loxF8L | 4 | loxF8R | 3 | TATATACA/TGTATATA | 7 |
| chr17 | 51568133 | ATATATGTGTGTG-TATATACA-TCTACAAATTTTT | loxF8L | 4 | loxF8L | 3 | TATATACA/TGTATATA | 7 |
| chr14 | 93158576 | ATATATATGTGTG-TGTATATA-TACACAGATGTGT | loxF8L | 4 | loxF8L | 3 | TATATACA/TGTATATA | 7 |
| chr8 | 82797849 | ACATATATGTGTA-TATATACA-CACACACACATAG | loxF8L | 4 | loxF8R | 3 | TATATACA/TGTATATA | 7 |
| chr19 | 34902678 | ATATATATGTGCA-TGTATATA-TACACATATATGT | loxF8L | 3 | loxF8L | 4 | TATATACA/TGTATATA | 7 |
| chr12 | 104096799 | AGAAATGTGTGTA-TGTATATA-TACATACATATAT | loxF8L | 3 | loxF8L | 4 | TATATACA/TGTATATA | 7 |
| chr6 | 75834808 | ATATATATGTGTA-TATATACA-TACACACAATTCT | loxF8L | 3 | loxF8L | 4 | TATATACA/TGTATATA | 7 |
| chr12 | 129144991 | AGTAATATGTGTA-TATATACA-CACACATATTTAT | loxF8L | 4 | loxF8L | 3 | TATATACA/TGTATATA | 7 |
| chr11 | 119152684 | CTTGGATTGTGAG-TGTATATA-CACATACTTTTAT | loxF8R | 3 | loxF8R | 4 | TATATACA/TGTATATA | 7 |
| chr12 | 102578666 | ATAAATATATGTA-TATATACA-CACACACACGTAT | loxF8L | 3 | loxF8R | 4 | TATATACA/TGTATATA | 7 |
| chrX | 62965147 | ATATATTTATGTA-TGTATATA-TACATATATTTAT | loxF8L | 4 | loxF8L | 3 | TATATACA/TGTATATA | 7 |
| chr7 | 117212737 | ATAAATATGCATA-TGTATATA-TACACATATGTAT | loxF8L | 4 | loxF8L | 3 | TATATACA/TGTATATA | 7 |
| chr9 | 9195019 | ATATGTGTGTGTG-TGTATATA-TATACATATTTAT | loxF8R | 4 | loxF8L | 3 | TATATACA/TGTATATA | 7 |
| chr10 | 35206271 | ATAAATATGTATA-TATATACA-TGCATATATATAT | loxF8L | 3 | loxF8L | 4 | TATATACA/TGTATATA | 7 |
| chr4 | 31964102 | ATAAATATGTGTG-TGTATATA-TGTACATATATAT | loxF8L | 3 | loxF8L | 4 | TATATACA/TGTATATA | 7 |
| chr11 | 84666445 | ATAAATCTCTTTA-TGTATATA-TCTATATATGTAT | loxF8L | 3 | loxF8L | 4 | TATATACA/TGTATATA | 7 |
| chr2 | 16829522 | ATAAATGTATATA-TGTATATA-TGCACATATATAT | loxF8L | 4 | loxF8L | 3 | TATATACA/TGTATATA | 7 |
| chr2 | 49340966 | ATATATATGTATA-TATATACA-CACACACATTTAT | loxF8L | 4 | loxF8L | 3 | TATATACA/TGTATATA | 7 |
| chrY | 25533498 | ATATGTTTGTATG-TGTATATA-TAAACACATTTAT | loxF8R | 4 | loxF8L | 3 | TATATACA/TGTATATA | 7 |
| chrY | 25553623 | ATAAATGTGTTTA-TATATACA-CATACAAACATAT | loxF8L | 3 | loxF8R | 4 | TATATACA/TGTATATA | 7 |
| chrY | 24116477 | ATATGTTTGTATG-TGTATATA-TAAACACATTTAT | loxF8R | 4 | loxF8L | 3 | TATATACA/TGTATATA | 7 |
| chrY | 24136606 | ATAAATGTGTTTA-TATATACA-CATACAAACATAT | loxF8L | 3 | loxF8R | 4 | TATATACA/TGTATATA | 7 |
| chr17 | 2713618 | ATATATGTGTGTA-TATATACA-CATACATACTTAT | loxF8L | 3 | loxF8R | 4 | TATATACA/TGTATATA | 7 |
| chr17 | 54722114 | ATAAGTGTGTGTG-TGTATATA-TACATACATATAT | loxF8R | 3 | loxF8L | 4 | TATATACA/TGTATATA | 7 |
| chr9 | 17738541 | ATAAGTGTGTGTG-TATATACA-TACATATATATAT | loxF8R | 3 | loxF8L | 4 | TATATACA/TGTATATA | 7 |
| chr12 | 82342616 | ATATATGTGTGTA-TGTATATA-TACATATATGTAT | loxF8L | 3 | loxF8L | 4 | TATATACA/TGTATATA | 7 |
| chr4 | 59899795 | ATATATGTGTGTA-TATATACA-TACATATATGTAT | loxF8L | 3 | loxF8L | 4 | TATATACA/TGTATATA | 7 |
| chr6 | 106139108 | ATATATGTGTGTA-TATATACA-TACATATATGTAT | loxF8L | 3 | loxF8L | 4 | TATATACA/TGTATATA | 7 |
| chr6 | 117923588 | ATACATATGTGTA-TATATACA-TATACATATGTAT | loxF8L | 3 | loxF8L | 4 | TATATACA/TGTATATA | 7 |
| chr10 | 35683225 | ATATATGTGTGTA-TATATACA-CGCACATATGTAT | loxF8L | 3 | loxF8L | 4 | TATATACA/TGTATATA | 7 |
| chr12 | 44882461 | ATACATATGTGTA-TATATACA-TACACACACATAT | loxF8L | 3 | loxF8L | 4 | TATATACA/TGTATATA | 7 |
| chr20 | 35097754 | ATACATATGTGTA-TATATACA-TACACATATATAC | loxF8L | 3 | loxF8L | 4 | TATATACA/TGTATATA | 7 |
| chr2 | 123955487 | ATACATATGTGTA-TATATACA-TACATACATATAT | loxF8L | 3 | loxF8L | 4 | TATATACA/TGTATATA | 7 |
| chr5 | 161776335 | ATATATGTATGTA-TGTATATA-TACACATATGTAT | loxF8L | 4 | loxF8L | 3 | TATATACA/TGTATATA | 7 |
| chr12 | 16065834 | CTAAATGTGTATA-TGTATATA-TACACATATGTAT | loxF8L | 4 | loxF8L | 3 | TATATACA/TGTATATA | 7 |
| chr12 | 7329077 | ATATATATCTGAA-TGTATATA-TACACATATGTAT | loxF8L | 4 | loxF8L | 3 | TATATACA/TGTATATA | 7 |
| chr3 | 66208907 | ATATAAATGTGTA-TATATACA-TACACATATATAT | loxF8L | 4 | loxF8L | 3 | TATATACA/TGTATATA | 7 |
| chr11 | 55849477 | ATATAGCTATAGA-TATATACA-TACACAATTTTAA | loxF8L | 4 | loxF8R | 3 | TATATACA/TGTATATA | 7 |
| chr13 | 65113958 | ATATATATATGGA-TATATACA-CACACATACATAT | loxF8L | 3 | loxF8R | 4 | TATATACA/TGTATATA | 7 |
| chr14 | 41539380 | ATATATATATGTA-TATATACA-CACACAAACTTTA | loxF8L | 4 | loxF8R | 3 | TATATACA/TGTATATA | 7 |
| chr1 | 144541962 | ATATATATATGTA-TATATACA-TACACATATATAT | loxF8L | 4 | loxF8L | 3 | TATATACA/TGTATATA | 7 |
| chr1 | 144541990 | ATATATATATGTA-TATATACA-TACACATATATAT | loxF8L | 4 | loxF8L | 3 | TATATACA/TGTATATA | 7 |
| chr1 | 144542018 | ATATATATATGTA-TATATACA-TACACATATATAT | loxF8L | 4 | loxF8L | 3 | TATATACA/TGTATATA | 7 |
| chr4 | 59952165 | ATATATATGTATA-TATATACA-CACACACACATAG | loxF8L | 4 | loxF8R | 3 | TATATACA/TGTATATA | 7 |
| chr6 | 8299852 | ATATATATGTATA-TATATACA-CACACAGATATAT | loxF8L | 4 | loxF8L | 3 | TATATACA/TGTATATA | 7 |
| chr18 | 65845200 | ATATATATGTATA-TATATACA-TACACACATATAT | loxF8L | 4 | loxF8L | 3 | TATATACA/TGTATATA | 7 |
| chr6 | 63125251 | ATATATATGTATA-TATATACA-TGCACACATATAT | loxF8L | 4 | loxF8L | 3 | TATATACA/TGTATATA | 7 |
| chr4 | 90365358 | ATATATATGTGCA-TATATACA-TATACACATATAT | loxF8L | 3 | loxF8L | 4 | TATATACA/TGTATATA | 7 |
| chr6 | 121509678 | ATATATGTGTATA-TGTATATA-TGCACATATATAT | loxF8L | 4 | loxF8L | 3 | TATATACA/TGTATATA | 7 |
| chr13 | 80478498 | ATATGTGTGTGTG-TGTATATA-TACACATATATAT | loxF8R | 4 | loxF8L | 3 | TATATACA/TGTATATA | 7 |
| chr16 | 31814442 | ATATATATGTGTA-TATATACA-CACACACACATAT | loxF8L | 3 | loxF8R | 4 | TATATACA/TGTATATA | 7 |
| chr5 | 19848928 | ATATATATGTGTA-TATATACA-CACACACACATAT | loxF8L | 3 | loxF8R | 4 | TATATACA/TGTATATA | 7 |
| chr11 | 116793512 | ATATATGTGTGTG-TGTATATA-TACACATATATAT | loxF8L | 4 | loxF8L | 3 | TATATACA/TGTATATA | 7 |
| chr2 | 191141586 | ATATATATGTGTA-TATATACA-CACACACATATAT | loxF8L | 3 | loxF8L | 4 | TATATACA/TGTATATA | 7 |
| chr3 | 28515792 | ATATATGTGTGTG-TGTATATA-TACACATATATAT | loxF8L | 4 | loxF8L | 3 | TATATACA/TGTATATA | 7 |
| chr9 | 32892387 | ATATATATGTGTA-TATATACA-CACACACATATAT | loxF8L | 3 | loxF8L | 4 | TATATACA/TGTATATA | 7 |
| chr1 | 189635220 | ATATATATGTGTA-TATATACA-CACACATATATAT | loxF8L | 3 | loxF8L | 4 | TATATACA/TGTATATA | 7 |
| chr21 | 19773187 | ATATATATGTGTG-TGTATATA-TACACATATATAT | loxF8L | 4 | loxF8L | 3 | TATATACA/TGTATATA | 7 |
| chr22 | 20620208 | ATATATATGTGTG-TGTATATA-TACACATATATAT | loxF8L | 4 | loxF8L | 3 | TATATACA/TGTATATA | 7 |
| chr3 | 182256889 | ATATATATGTGTG-TGTATATA-TACACATATATAT | loxF8L | 4 | loxF8L | 3 | TATATACA/TGTATATA | 7 |
| chr16 | 31814472 | ATATATATGTGTA-TATATACA-CTCACACATATAT | loxF8L | 3 | loxF8L | 4 | TATATACA/TGTATATA | 7 |
| chr2 | 83185141 | GAAAATCTGTTTA-TATATACA-TACACATATATAT | loxF8L | 4 | loxF8L | 3 | TATATACA/TGTATATA | 7 |
| chr2 | 131771016 | ATGTATATGTGTA-TATATACA-TACACATATATAT | loxF8L | 4 | loxF8L | 3 | TATATACA/TGTATATA | 7 |
| chr4 | 93042379 | ATATATATGTGTA-TATATACA-TACATACATATAT | loxF8L | 3 | loxF8L | 4 | TATATACA/TGTATATA | 7 |
| chr19 | 13290730 | ATATATGTGTATA-TATATACA-TACACATATATAT | loxF8L | 4 | loxF8L | 3 | TATATACA/TGTATATA | 7 |
| chrY | 6156818 | ATATATGTGTATA-TGTATATA-TACACATATATAT | loxF8L | 4 | loxF8L | 3 | TATATACA/TGTATATA | 7 |
| chr1 | 72111047 | CTAAATTTTTAGA-TATATACA-TACACATATATAT | loxF8L | 4 | loxF8L | 3 | TATATACA/TGTATATA | 7 |
| chr8 | 113005038 | ATATATATGTGTC-TGTATATA-TACACACATATAT | loxF8L | 4 | loxF8L | 3 | TATATACA/TGTATATA | 7 |
| chr18 | 54105073 | ATATATGTGTGTA-TGTATATA-CACACATATATAT | loxF8L | 3 | loxF8L | 4 | TATATACA/TGTATATA | 7 |
| chr5 | 53137136 | ATATATGTGTGTA-TATATACA-CACACATATATAT | loxF8L | 3 | loxF8L | 4 | TATATACA/TGTATATA | 7 |
| chr8 | 36801667 | ATATATGTGTGTA-TATATACA-CACACATATATAT | loxF8L | 3 | loxF8L | 4 | TATATACA/TGTATATA | 7 |
| chr10 | 43837088 | ATATGAGTGTGTG-TGTATATA-TATATAGATATAT | loxF8R | 3 | loxF8L | 4 | TATATACA/TGTATATA | 7 |
| chr8 | 87862055 | GTATATCTGTGTA-TATATACA-TATATAGATATAT | loxF8L | 3 | loxF8L | 4 | TATATACA/TGTATATA | 7 |
| chr5 | 112243228 | ATATATCTATGTA-TATATACA-CACACACACATAT | loxF8L | 3 | loxF8R | 4 | TATATACA/TGTATATA | 7 |
| chr14 | 38040162 | GTACATGTGTGCA-TATATACA-TATACAGATATAT | loxF8L | 4 | loxF8L | 3 | TATATACA/TGTATATA | 7 |
| chr7 | 79830860 | GTATGTGTGTGTG-TATATACA-CACACAGATATAT | loxF8R | 4 | loxF8L | 3 | TATATACA/TGTATATA | 7 |

| | | | | | | | | |
|---|---|---|---|---|---|---|---|---|
| chr8 | 126662362 | ATATATGTGTGGG-TGTATATA-TACATACATATAT | loxF8L | 3 | loxF8L | 4 | TATATACA/TGTATATA | 7 |
| chr22 | 46940693 | ATATATGTGTGTA-TGTATATA-CACACACACACAG | loxF8L | 3 | loxF8R | 4 | TATATACA/TGTATATA | 7 |
| chr9 | 22808291 | ATATATGTGTGTA-TATATACA-CACACACATAT | loxF8L | 3 | loxF8R | 4 | TATATACA/TGTATATA | 7 |
| chrX | 46847687 | ATATATGTGTGTG-TGTATATA-TACACACATATAT | loxF8L | 4 | loxF8L | 3 | TATATACA/TGTATATA | 7 |
| chr2 | 143471169 | ATGAATATGTGTG-TGTATATA-TACACACATATAT | loxF8L | 4 | loxF8L | 3 | TATATACA/TGTATATA | 7 |
| chr13 | 58083980 | ATGTATGTGTGTA-TATATACA-TACACACATATAT | loxF8L | 4 | loxF8L | 3 | TATATACA/TGTATATA | 7 |
| chr1 | 78532535 | ATATATGTGTGTA-TATATACA-TACACACATATAC | loxF8L | 3 | loxF8L | 4 | TATATACA/TGTATATA | 7 |
| chr14 | 40341642 | ATATATTTCTGAA-TATATACA-CACACATGCTTAC | loxF8L | 4 | loxF8R | 3 | TATATACA/TGTATATA | 7 |
| chr5 | 92430495 | ATGAATATGTACA-TATATACA-TTCACAGACATAT | loxF8L | 4 | loxF8L | 3 | TATATACA/TGTATATA | 7 |
| chr2 | 147176759 | CAAATAATGTGTG-TATATACA-TACACACACATAT | loxF8R | 3 | loxF8L | 4 | TATATACA/TGTATATA | 7 |
| chr2 | 50206838 | CAAACAATGTGTG-TGTATATA-CACACACACACAG | loxF8R | 3 | loxF8R | 4 | TATATACA/TGTATATA | 7 |
| chr15 | 71695458 | TTAATAATGTGTG-TATATACA-CATAAAATATTTG | loxF8R | 3 | loxF8R | 4 | TATATACA/TGTATATA | 7 |
| chr13 | 101380058 | GTATATATGTGGA-TGTATATA-TACACATATATAC | loxF8L | 3 | loxF8L | 4 | TATATACA/TGTATATA | 7 |
| --- | | | | | | | | 0 |
| chr5 | 111160949 | ATACATATGTTGA-TATCTTAA-TCCTCAAAGTTAT | loxF8L | 3 | loxF8L | 3 | TATCTTAA/TTAAGATA | 6 |
| chr12 | 112831695 | CTCAGACTGTGTG-TATCTTAA-AACATTGATTTAT | loxF8R | 2 | loxF8L | 4 | TATCTTAA/TTAAGATA | 6 |
| --- | | | | | | | | 0 |
| chrX | 36553518 | ATAAATATGTAAA-TGTACATA-TACACATATTTAC | loxF8L | 3 | loxF8L | 3 | TATGTACA/TGTACATA | 6 |
| chr7 | 113223394 | TTAATTTTGTGTG-TGTACATA-TAAACATATTTAT | loxF8R | 3 | loxF8L | 3 | TATGTACA/TGTACATA | 6 |
| chr10 | 131836555 | ATAAGTGTGTGTG-TGTACATA-TACACATATATAT | loxF8R | 3 | loxF8L | 3 | TATGTACA/TGTACATA | 6 |
| chr3 | 181906035 | ATATATTTGTGTG-TGTACATA-TCCACCTATTTAT | loxF8L | 4 | loxF8L | 2 | TATGTACA/TGTACATA | 6 |
| chr20 | 19096471 | ATATACATGTGTA-TGTACATA-TATACATATTTAT | loxF8L | 4 | loxF8L | 3 | TATGTACA/TGTACATA | 7 |
| chr1 | 181146215 | ATATATGTGTGCA-TATGTACA-CTCACAGGTGTAT | loxF8L | 3 | loxF8L | 4 | TATGTACA/TGTACATA | 7 |
| chr1 | 79631461 | ATATAAATGTGTA-TGTACATA-TACACATATGTAT | loxF8L | 4 | loxF8L | 3 | TATGTACA/TGTACATA | 7 |
| chr3 | 568270 | ATACATGTGTGTA-TGTACATA-CATACAAGCATAT | loxF8L | 3 | loxF8R | 4 | TATGTACA/TGTACATA | 7 |
| chr15 | 44369247 | ATATATATGTATA-TATGTACA-TATACATTCTTAG | loxF8L | 4 | loxF8R | 3 | TATGTACA/TGTACATA | 7 |
| chr3 | 40164309 | CTAAGTATATATG-TATGTACA-TGCACATATATAT | loxF8R | 4 | loxF8L | 3 | TATGTACA/TGTACATA | 7 |
| chr16 | 25163493 | ATATATATGTGTA-TATGTACA-CATACAATCATTC | loxF8L | 3 | loxF8R | 4 | TATGTACA/TGTACATA | 7 |
| chr10 | 75749445 | ATATATATGTGTA-TATGTACA-TACACACACATAT | loxF8L | 3 | loxF8L | 4 | TATGTACA/TGTACATA | 7 |
| chr7 | 119152469 | ATATATATGTGTA-TATGTACA-TGCACACACATAT | loxF8L | 3 | loxF8L | 4 | TATGTACA/TGTACATA | 7 |
| chr1 | 200816669 | GTATATGTGTGTA-TATGTACA-TGCACACATATAT | loxF8L | 4 | loxF8L | 3 | TATGTACA/TGTACATA | 7 |
| --- | | | | | | | | 0 |
| chrX | 137217572 | CTAAGATTTTATG-TTGTAATA-TCCACTGTTTTAA | loxF8R | 2 | loxF8L | 3 | TATTACAA/TTGTAATA | 5 |
| chr2 | 213210865 | ATAAAACATTGGA-TATTACAA-AATATAATCTTAG | loxF8L | 3 | loxF8R | 3 | TATTACAA/TTGTAATA | 6 |
| chr4 | 54601389 | TTAAAACTTTGGA-TATTACAA-AATAGAATTTTAG | loxF8L | 3 | loxF8R | 4 | TATTACAA/TTGTAATA | 7 |
| chr1 | 72811865 | TTAAAACATTGGA-TATTACAA-AATATAATCTTAG | loxF8L | 4 | loxF8R | 3 | TATTACAA/TTGTAATA | 7 |
| chr11 | 56399890 | TTAAAACATTGGA-TATTACAA-AATAGAATCTTAG | loxF8L | 4 | loxF8R | 3 | TATTACAA/TTGTAATA | 7 |
| chr1 | 184814640 | CTAAGATTCTATT-TTGTAATA-TCCAATGTTTTAA | loxF8R | 3 | loxF8L | 4 | TATTACAA/TTGTAATA | 7 |
| chr3 | 84049394 | CTAAGATTCTATT-TTGTAATA-TCCAATGTTTTAA | loxF8R | 3 | loxF8L | 4 | TATTACAA/TTGTAATA | 7 |
| chr4 | 42678835 | CTAAGATTCTATT-TTGTAATA-TCCAATGTTTTAA | loxF8R | 3 | loxF8L | 4 | TATTACAA/TTGTAATA | 7 |
| chr6 | 36882220 | CTAAGATTCTATT-TTGTAATA-TCCAATGTTTTAA | loxF8R | 3 | loxF8L | 4 | TATTACAA/TTGTAATA | 7 |
| --- | | | | | | | | 0 |
| chr9 | 74781847 | CCACATCTGTGGA-TTCAAATA-ACCACAGATTGAA | loxF8L | 3 | loxF8L | 3 | TATTTGAA/TTCAAATA | 6 |
| chr6 | 164046670 | TTGAATCTTTGGG-TTCAAATA-TCCAAAGATTTCT | loxF8L | 4 | loxF8L | 2 | TATTTGAA/TTCAAATA | 6 |
| --- | | | | | | | | 0 |
| chr7 | 105556538 | CTATGATTGTATC-TGAAAAAA-TTAACTGATTTAT | loxF8R | 3 | loxF8L | 3 | TGAAAAAA/TTTTTTCA | 6 |
| chr6 | 124560708 | CTAAGATTCTATG-TGAAAAAA-TTCACCTTCTTAG | loxF8R | 2 | loxF8R | 4 | TGAAAAAA/TTTTTTCA | 6 |
| chr2 | 210565180 | TAAATATTGTGTG-TGAAAAAA-TCTGCAGTCTTAT | loxF8R | 3 | loxF8L | 4 | TGAAAAAA/TTTTTTCA | 7 |
| chr8 | 53161192 | CCAAGATTTTGTA-TTTTTTCA-GACATAGTCTTTG | loxF8R | 3 | loxF8R | 4 | TGAAAAAA/TTTTTTCA | 7 |
| chr4 | 47997044 | GTGAGATTCTGTC-TGAAAAAA-AACAAAAACTTAG | loxF8R | 4 | loxF8R | 3 | TGAAAAAA/TTTTTTCA | 7 |
| --- | | | | | | | | 0 |
| chr2 | 226862003 | CTAATTTTGTGTG-TGTATGCA-TACACATATATAT | loxF8R | 2 | loxF8L | 3 | TGCATACA/TGTATGCA | 5 |
| chr2 | 137447943 | AAAAATATGAGAA-TGCATACA-GACACATTCATAG | loxF8L | 4 | loxF8R | 3 | TGCATACA/TGTATGCA | 7 |
| chr19 | 53180541 | ATAAAACCATGGG-TGTATGCA-TCTACATATTCAT | loxF8L | 4 | loxF8L | 3 | TGCATACA/TGTATGCA | 7 |
| --- | | | | | | | | 0 |
| chr5 | 109179587 | CCACATCTGTGGA-TTCAACCA-TCCAGAGATTTTT | loxF8L | 3 | loxF8L | 2 | TGGTTGAA/TTCAACCA | 5 |
| chr5 | 134167348 | CTACATCTGTGGA-TTCAACCA-ACCACAGATTAAA | loxF8L | 2 | loxF8L | 3 | TGGTTGAA/TTCAACCA | 5 |
| chr8 | 63184095 | TTCAATCTGTGAA-TGGTTGAA-TCCACAGATGTAG | loxF8L | 3 | loxF8L | 2 | TGGTTGAA/TTCAACCA | 5 |
| chr5 | 125045667 | CCACATCTGTGGA-TTCAACCA-ACCGCAGATTCAT | loxF8L | 3 | loxF8L | 3 | TGGTTGAA/TTCAACCA | 6 |
| chr13 | 49117562 | CCAAATCTATGGA-TTCAACCA-ACCACAGATTGAA | loxF8L | 3 | loxF8L | 3 | TGGTTGAA/TTCAACCA | 6 |
| chr12 | 131716584 | CCACATCTGTGGA-TTCAACCA-ACCACAGATCTAA | loxF8L | 3 | loxF8L | 3 | TGGTTGAA/TTCAACCA | 6 |
| chr9 | 20502484 | CCACATCTGTGGA-TTCAACCA-ACCACAGATTGAA | loxF8L | 3 | loxF8L | 3 | TGGTTGAA/TTCAACCA | 6 |
| chr17 | 63667803 | CTCAATCTGTGGT-TGGTTGAA-TCCACAGATGCAG | loxF8L | 3 | loxF8L | 3 | TGGTTGAA/TTCAACCA | 6 |
| chr8 | 31069420 | CTGCATCTGTGGA-TTCAACCA-ACCACAGATTGAG | loxF8L | 3 | loxF8L | 3 | TGGTTGAA/TTCAACCA | 6 |
| chr3 | 185026137 | CTGCATCTGTGGA-TTCAACCA-ACCACAGATTGAA | loxF8L | 3 | loxF8L | 3 | TGGTTGAA/TTCAACCA | 6 |
| chr12 | 50781354 | TTCAATCTGTGGT-TGGTTGAA-TCCACAGATGTGA | loxF8L | 3 | loxF8L | 3 | TGGTTGAA/TTCAACCA | 6 |
| chr6 | 127488799 | TCACATCTGTGGA-TTCAACCA-ACCACAGATTGAA | loxF8L | 3 | loxF8L | 3 | TGGTTGAA/TTCAACCA | 6 |
| chrX | 104557875 | TTAAATCTCTGGT-TGGTTGAA-TCCACAGATGTCA | loxF8L | 3 | loxF8L | 3 | TGGTTGAA/TTCAACCA | 6 |
| chr8 | 82523326 | TTGCATCTGTGGA-TTCAACCA-ACCACAGATTGAA | loxF8L | 3 | loxF8L | 3 | TGGTTGAA/TTCAACCA | 6 |
| chr8 | 67259126 | TTCAATTTGTGGT-TGGTTGAA-TCCACAGATGTAG | loxF8L | 4 | loxF8L | 2 | TGGTTGAA/TTCAACCA | 6 |
| chr18 | 24468511 | TTCTATCTGTGGT-TGGTTGAA-TCCACAGATGTAG | loxF8L | 4 | loxF8L | 2 | TGGTTGAA/TTCAACCA | 6 |
| chr2 | 52748932 | CTATATCTGTGGA-TTCAACCA-ACCACAGATCAAA | loxF8L | 2 | loxF8L | 4 | TGGTTGAA/TTCAACCA | 6 |
| chr15 | 50486002 | TTCAATCTGTGGA-TGGTTGAA-TTCACAGATGCAG | loxF8L | 2 | loxF8L | 4 | TGGTTGAA/TTCAACCA | 6 |
| chrY | 19566096 | TTTGATCTGTGGT-TGGTTGAA-TCTACAGATGTCT | loxF8L | 4 | loxF8L | 3 | TGGTTGAA/TTCAACCA | 7 |
| chr18 | 55600751 | CAACATCTGTGGA-TTCAACCA-GCCACAGATCAAA | loxF8L | 3 | loxF8L | 4 | TGGTTGAA/TTCAACCA | 7 |
| chrX | 1795835 | TTTCATCTGTGGT-TGGTTGAA-TCCACGGATTTGG | loxF8L | 4 | loxF8L | 3 | TGGTTGAA/TTCAACCA | 7 |
| chrY | 1795835 | TTTCATCTGTGGT-TGGTTGAA-TCCACGGATTTGG | loxF8L | 4 | loxF8L | 3 | TGGTTGAA/TTCAACCA | 7 |
| chr17 | 37031127 | CCACATCAGTGGA-TTCAACCA-ACCACAGATTGAA | loxF8L | 4 | loxF8L | 3 | TGGTTGAA/TTCAACCA | 7 |
| chr1 | 203493070 | CCACATCCGTGGA-TTCAACCA-ACCACAGATTGAA | loxF8L | 4 | loxF8L | 3 | TGGTTGAA/TTCAACCA | 7 |
| chr18 | 11869548 | CCACATCTGCGGA-TTCAACCA-ACCACAGATTGAA | loxF8L | 4 | loxF8L | 3 | TGGTTGAA/TTCAACCA | 7 |
| chr14 | 64672054 | CCACATCTGTGGA-TTCAACCA-ACCACAGATAGAA | loxF8L | 3 | loxF8L | 4 | TGGTTGAA/TTCAACCA | 7 |
| chr6 | 15869273 | CCACATCTGTGGA-TTCAACCA-ACCACAGATCAAA | loxF8L | 3 | loxF8L | 4 | TGGTTGAA/TTCAACCA | 7 |
| chr6 | 111467531 | CCACATCTGTGGA-TTCAACCA-ACCACAGATCAAA | loxF8L | 3 | loxF8L | 4 | TGGTTGAA/TTCAACCA | 7 |
| chr6 | 123777164 | CTTGATCTGTGGT-TGGTTGAA-TCCACAGATGTGG | loxF8L | 4 | loxF8L | 3 | TGGTTGAA/TTCAACCA | 7 |
| chr10 | 32000957 | TTTCATCTGTGGT-TGGTTGAA-TCCACAGATGTGG | loxF8L | 4 | loxF8L | 3 | TGGTTGAA/TTCAACCA | 7 |
| chr17 | 63771169 | CCACATCTGTGGA-TTCAACCA-ACCACAGATGGAA | loxF8L | 3 | loxF8L | 4 | TGGTTGAA/TTCAACCA | 7 |
| chr2 | 205404162 | CCACATCTGTGGA-TTCAACCA-ACCGCAGATTGAA | loxF8L | 3 | loxF8L | 4 | TGGTTGAA/TTCAACCA | 7 |

| | | | | | | | | |
|---|---|---|---|---|---|---|---|---|
| chr6 | 106350383 | CCACATCTGTGGA-TTCAACCA-GCCACAGATAAAA | loxF8L | 3 | loxF8L | 4 | TGGTTGAA/TTCAACCA | 7 |
| chr4 | 156899119 | CCACATTTGTGGA-TTCAACCA-ACCACAGATTGAA | loxF8L | 4 | loxF8L | 3 | TGGTTGAA/TTCAACCA | 7 |
| chr5 | 102273938 | CCATATCTGTGGA-TTCAACCA-ACCACAGATCAAA | loxF8L | 3 | loxF8L | 4 | TGGTTGAA/TTCAACCA | 7 |
| chr6 | 131333508 | TTTGATCTGTGGT-TGGTTGAA-TCCACAGATATGG | loxF8L | 4 | loxF8L | 3 | TGGTTGAA/TTCAACCA | 7 |
| chr7 | 95359273 | TTCAATCTCTGGT-TGGTTGAA-TCCATAGATGTAG | loxF8L | 4 | loxF8L | 3 | TGGTTGAA/TTCAACCA | 7 |
| chr18 | 9655644 | CTACATCTGTGAA-TTCAACCA-ACCATAGATTGAA | loxF8L | 3 | loxF8L | 4 | TGGTTGAA/TTCAACCA | 7 |
| chr5 | 60646528 | CTGCATCTATGGA-TTCAACCA-ACCACAGATTGAA | loxF8L | 4 | loxF8L | 3 | TGGTTGAA/TTCAACCA | 7 |
| chr2 | 127369262 | TTCTATCTGTGGT-TGGTTGAA-TCCACAGATGCAG | loxF8L | 4 | loxF8L | 3 | TGGTTGAA/TTCAACCA | 7 |
| chr4 | 146450148 | CTGCATCTGTGGA-TTCAACCA-ACCACAGATCAAA | loxF8L | 3 | loxF8L | 4 | TGGTTGAA/TTCAACCA | 7 |
| chr8 | 101352019 | CTGCATCTGTGGA-TTCAACCA-ACCACAGATCAAA | loxF8L | 3 | loxF8L | 4 | TGGTTGAA/TTCAACCA | 7 |
| chr9 | 121109292 | TTTAATCCGTGGT-TGGTTGAA-TCCACAGATGCAG | loxF8L | 4 | loxF8L | 3 | TGGTTGAA/TTCAACCA | 7 |
| chr3 | 4739488 | CTGCATCTGTGGA-TTCAACCA-ACTGCAGATTTAA | loxF8L | 3 | loxF8L | 4 | TGGTTGAA/TTCAACCA | 7 |
| chr7 | 141749049 | CTGCATCTGTGGA-TTCAACCA-AGCACAGATTGAA | loxF8L | 3 | loxF8L | 4 | TGGTTGAA/TTCAACCA | 7 |
| chr4 | 123135526 | CTGCATCTTTGGA-TTCAACCA-ACCACAGATTGAA | loxF8L | 4 | loxF8L | 3 | TGGTTGAA/TTCAACCA | 7 |
| chr7 | 114571555 | CTGTATCTGTGGA-TTCAACCA-ACCACAGATCCAA | loxF8L | 3 | loxF8L | 4 | TGGTTGAA/TTCAACCA | 7 |
| chr17 | 18057761 | CTGTATCTGTGGA-TTCAACCA-GCCACAGTTTGAA | loxF8L | 3 | loxF8L | 4 | TGGTTGAA/TTCAACCA | 7 |
| chr7 | 87010799 | TTCAATCTGAGGT-TGGTTGAA-TCCACAGATGTGC | loxF8L | 4 | loxF8L | 3 | TGGTTGAA/TTCAACCA | 7 |
| chr3 | 161061518 | GTACATTTGTGGA-TTCAACCA-ACCACAGATCAAA | loxF8L | 3 | loxF8L | 4 | TGGTTGAA/TTCAACCA | 7 |
| chr3 | 155006867 | TGCCATCTGTGGA-TTCAACCA-ACCACAGATTGAC | loxF8L | 4 | loxF8L | 3 | TGGTTGAA/TTCAACCA | 7 |
| chr19 | 44144868 | TTGCATTTGTGGA-TTCAACCA-ACCACAGATTGAA | loxF8L | 4 | loxF8L | 3 | TGGTTGAA/TTCAACCA | 7 |
| chr8 | 112117554 | TTTAATCTGTAAA-TTCAACCA-ACCACAGATTGAA | loxF8L | 4 | loxF8L | 3 | TGGTTGAA/TTCAACCA | 7 |
| chr9 | 90181630 | TTCCATCTGTGGT-TGGTTGAA-TCCACAGATGCAA | loxF8L | 4 | loxF8L | 3 | TGGTTGAA/TTCAACCA | 7 |
| --- | | | | | | | | 0 |
| chr5 | 58400365 | AGAAATCTGAGAA-TTCAGAAA-TTCTCAGATTTCT | loxF8L | 3 | loxF8L | 3 | TTCAGAAA/TTTCTGAA | 6 |
| chr5 | 174997211 | ATAAAACTGTTGA-TTTCTGAA-AAGAGAATCTCAG | loxF8L | 2 | loxF8R | 4 | TTCAGAAA/TTTCTGAA | 6 |
| chr6 | 145370139 | ATAAATTTGGAGA-TTCAGAAA-ACTACAGACTTGT | loxF8L | 3 | loxF8L | 4 | TTCAGAAA/TTTCTGAA | 7 |
| --- | | | | | | | | 0 |
| chr1 | 188767000 | ATAAATAAGTTGA-AAAAAGCA-CATACAATTTTAA | loxF8L | 3 | loxF8R | 3 | AAAAAGCA/TGCTTTTT | 6 |
| chr21 | 40393675 | ATATATCCGTCTA-TGCTTTTT-TCTACATATATAT | loxF8L | 4 | loxF8L | 3 | AAAAAGCA/TGCTTTTT | 7 |
| --- | | | | | | | | 0 |
| chr19 | 22237925 | ATAAGAGTTTGTG-TGTGTTTT-TCCACAATTTCAT | loxF8R | 3 | loxF8L | 3 | AAAACACA/TGTGTTTT | 6 |
| chrX | 34667918 | ATCAAATTGTGTG-AAAACACA-CACACATTCACAG | loxF8R | 3 | loxF8R | 3 | AAAACACA/TGTGTTTT | 6 |
| chr6 | 67227656 | TTATTATTGTGTG-TGTGTTTT-CCCATAGATATTT | loxF8R | 3 | loxF8L | 4 | AAAACACA/TGTGTTTT | 7 |
| chr17 | 13237143 | ATAAAACTTAGTA-TGTGTTTT-TCTACAACTTTAT | loxF8L | 4 | loxF8L | 3 | AAAACACA/TGTGTTTT | 7 |
| chr4 | 86224749 | ATAAATCTGAGTT-TGTGTTTT-CCCTCAAAATTAG | loxF8L | 3 | loxF8R | 4 | AAAACACA/TGTGTTTT | 7 |
| chr4 | 67436089 | ATAAATCTTTAAA-AAAACACA-CACACAAAAAAAG | loxF8L | 3 | loxF8R | 4 | AAAACACA/TGTGTTTT | 7 |
| chr3 | 87314490 | GTAAAATTATGTG-AAAACACA-CACACATACATAT | loxF8R | 3 | loxF8R | 4 | AAAACACA/TGTGTTTT | 7 |
| --- | | | | | | | | 0 |
| chr11 | 83506887 | ATAAGAGTTTGTG-CCTGTTTT-TCCACAATTTCAT | loxF8R | 3 | loxF8L | 3 | AAAACAGG/CCTGTTTT | 6 |
| chr11 | 56040236 | CTGAAATTGTGGA-AAAACAGG-CACAAACTCTTAT | loxF8L | 4 | loxF8R | 3 | AAAACAGG/CCTGTTTT | 7 |
| chr15 | 76696671 | TTTAGATAGTGTG-AAAACAGG-ACCACAAAGATAT | loxF8R | 3 | loxF8L | 4 | AAAACAGG/CCTGTTTT | 7 |
| --- | | | | | | | | 0 |
| chr4 | 85202796 | CTAAAATTGCCTG-TATGTTTT-CCCACAGATTGCT | loxF8R | 3 | loxF8L | 3 | AAAACATA/TATGTTTT | 6 |
| chrX | 108482689 | ATAAGAGCTTGTG-TATGTTTT-TCCACAATTTTAG | loxF8R | 4 | loxF8L | 3 | AAAACATA/TATGTTTT | 7 |
| --- | | | | | | | | 0 |
| chr14 | 102662835 | ATAACTTTGTAGA-TAAGTTTT-TGCAAAGCTTTAT | loxF8L | 3 | loxF8L | 3 | AAAACTTA/TAAGTTTT | 6 |
| chr9 | 118403643 | ATAAATTTTCGAA-TAAGTTTT-CTCACCATCTTAC | loxF8L | 4 | loxF8R | 3 | AAAACTTA/TAAGTTTT | 7 |
| --- | | | | | | | | 0 |
| chr5 | 140049176 | ACAAAGCTATGGA-TATCTTTT-TGCCAAGATTTAT | loxF8L | 3 | loxF8L | 3 | AAAAGATA/TATCTTTT | 6 |
| chr1 | 100760982 | ATAATTTTGTTGA-AAAAGATA-GAAATAAACTTAG | loxF8L | 3 | loxF8R | 4 | AAAAGATA/TATCTTTT | 7 |
| --- | | | | | | | | 0 |
| chr5 | 141228582 | ATAAATCTAGGTA-AAAATATT-CATATCATCTTAG | loxF8L | 3 | loxF8R | 3 | AAAATATT/AATATTTT | 6 |
| chr1 | 238724001 | CTAATATTTTGTT-AATATTTT-TGCACATATATTT | loxF8R | 3 | loxF8L | 4 | AAAATATT/AATATTTT | 7 |
| chr3 | 83167783 | TTGACTTTGTGTG-AATATTTT-TGTACTGATTTAT | loxF8R | 4 | loxF8L | 3 | AAAATATT/AATATTTT | 7 |
| chr12 | 46633858 | ATATATCTTTGCA-AATATTTT-CTCACAGTCTGTG | loxF8L | 3 | loxF8R | 4 | AAAATATT/AATATTTT | 7 |
| chr20 | 18880908 | GCAAGTCTGTGGA-AAAATATT-CCCTCAATTTTTG | loxF8L | 3 | loxF8R | 4 | AAAATATT/AATATTTT | 7 |
| --- | | | | | | | | 0 |
| chr5 | 16485028 | AGATATCTGTGGC-AAAATCTT-GCCACAGATATCT | loxF8L | 3 | loxF8L | 3 | AAAATCTT/AAGATTTT | 6 |
| chr8 | 60519514 | CTCTTATTGTGTG-AAGATTTT-CACAGTATCCTGG | loxF8R | 3 | loxF8R | 4 | AAAATCTT/AAGATTTT | 7 |
| --- | | | | | | | | 0 |
| chr3 | 31727658 | ATAAATATGTGCC-ACCTGTTT-CAAACAAACTTAT | loxF8L | 3 | loxF8R | 3 | AAACAGGT/ACCTGTTT | 6 |
| chr7 | 158486443 | AAAATTCTTTGAA-ACCTGTTT-TCCTCATATTTAG | loxF8L | 4 | loxF8L | 3 | AAACAGGT/ACCTGTTT | 7 |
| --- | | | | | | | | 0 |
| chr18 | 22525198 | TTAAGTTTCTGTG-ACAGCTTT-TCTACATATTTCT | loxF8R | 3 | loxF8L | 3 | AAAGCTGT/ACAGCTTT | 6 |
| chr16 | 10460150 | ATAAAATTTTGGA-AAAGCTGT-AACACCATTATAG | loxF8L | 3 | loxF8R | 4 | AAAGCTGT/ACAGCTTT | 7 |
| --- | | | | | | | | 0 |
| chr19 | 9757771 | CGAGGATTGAGTG-AAAGCTTT-CCCACATTCTTTG | loxF8R | 3 | loxF8R | 3 | AAAGCTTT/AAAGCTTT | 6 |
| --- | | | | | | | | 0 |
| chr13 | 20029537 | TTAAATCTGTATA-TTTTATTT-TTCACAAACTTAG | loxF8L | 3 | loxF8R | 3 | AAATAAAA/TTTTATTT | 6 |
| chr10 | 121386481 | AAAAATTTGAGCA-AAATAAAA-AACACAGTATTAG | loxF8L | 4 | loxF8R | 3 | AAATAAAA/TTTTATTT | 7 |
| chr5 | 28325373 | ATTATTCTGTTGA-TTTTATTT-TAAACATGTTTAT | loxF8L | 3 | loxF8L | 4 | AAATAAAA/TTTTATTT | 7 |
| chr15 | 86506797 | ATAAATGTGTACA-TTTTATTT-CCTCAAGATTTAT | loxF8L | 3 | loxF8L | 4 | AAATAAAA/TTTTATTT | 7 |
| chr11 | 103469969 | CTAATTCTCTGGA-AAATAAAA-TCCATATAATGAT | loxF8L | 3 | loxF8L | 4 | AAATAAAA/TTTTATTT | 7 |
| chrX | 65796662 | CCAGGATTGGGAG-AAATAAAA-CATGCAATCTAAG | loxF8R | 4 | loxF8R | 3 | AAATAAAA/TTTTATTT | 7 |
| --- | | | | | | | | 0 |
| chr10 | 106137959 | ATAAATCTTAAGA-TATTATTT-GCCACATATTGAT | loxF8L | 3 | loxF8L | 3 | AAATAATA/TATTATTT | 6 |
| chr6 | 146291350 | AGAAATATGTATA-TATTATTT-TGAACAGATATAT | loxF8L | 4 | loxF8L | 3 | AAATAATA/TATTATTT | 7 |
| chr7 | 118205968 | TTAAATTTGTGTA-TATTATTT-TACAATTATTTAT | loxF8L | 3 | loxF8L | 4 | AAATAATA/TATTATTT | 7 |
| chrX | 26429450 | GTAAATGTCTGGA-AAATAATA-CATACACACTTAT | loxF8L | 3 | loxF8R | 4 | AAATAATA/TATTATTT | 7 |
| --- | | | | | | | | 0 |
| chr12 | 23289054 | CTCAGATAATGTG-AAATACAG-CATACTTTCTTAG | loxF8R | 3 | loxF8R | 3 | AAATACAG/CTGTATTT | 6 |
| chr6 | 6413088 | AGAAATCTGTGCT-CTGTATTT-TACATTATCTTTG | loxF8L | 3 | loxF8R | 4 | AAATACAG/CTGTATTT | 7 |
| chr11 | 23794873 | ATAAATCACTGGT-AAATACAG-TTAAAAGATGTAT | loxF8L | 3 | loxF8L | 4 | AAATACAG/CTGTATTT | 7 |
| --- | | | | | | | | 0 |
| chr8 | 131521511 | CTAAAAATGTGTA-AAATATAA-CACTTATTCTTAG | loxF8R | 3 | loxF8R | 3 | AAATATAA/TTATATTT | 6 |
| chr1 | 72765228 | AAATATATGTCGA-TTATATTT-AACACTATCTCAG | loxF8L | 4 | loxF8R | 3 | AAATATAA/TTATATTT | 7 |
| --- | | | | | | | | 0 |

| | | | | | | | | | |
|---|---|---|---|---|---|---|---|---|---|
| chr10 | 54104421 | ATAAGAGTTTGTG-TCTGATTT-TCCACAATTTTAG | loxF8R | 3 | loxF8L | 3 | AAATCAGA/TCTGATTT | | 6 |
| chr21 | 28302084 | CTAAAGTTGTGGA-AAATCAGA-CACAGGCTCTTAT | loxF8L | 3 | loxF8R | 4 | AAATCAGA/TCTGATTT | | 7 |
| chr2 | 187204455 | ATAAGAGCTTGTG-TCTGATTT-TCCACAACTTTAG | loxF8R | 4 | loxF8L | 3 | AAATCAGA/TCTGATTT | | 7 |
| chrX | 125815519 | ATAAGAGCTTGTG-TCTGATTT-TCCACAATTTCAT | loxF8R | 4 | loxF8L | 3 | AAATCAGA/TCTGATTT | | 7 |
| chr2 | 39963595 | ATAAGAGCTTGTG-TCTGATTT-TCCACAATTTTAG | loxF8R | 4 | loxF8L | 3 | AAATCAGA/TCTGATTT | | 7 |
| chrX | 94555118 | CTAAAATTGTGGA-AAATCAGA-CACAAGCTCTTAT | loxF8L | 3 | loxF8R | 4 | AAATCAGA/TCTGATTT | | 7 |
| chr9 | 85376076 | ATAAGAGCTTGTG-TCTGATTT-TCCACAGCTTCAG | loxF8R | 4 | loxF8L | 3 | AAATCAGA/TCTGATTT | | 7 |
| chr5 | 121543770 | CTAAAATTGTGGA-AAATCAGA-CACGAACTCTTAT | loxF8L | 3 | loxF8R | 4 | AAATCAGA/TCTGATTT | | 7 |
| chr12 | 120608624 | CTGAAATTGTGGA-AAATCAGA-CACACACCCTTAT | loxF8L | 4 | loxF8R | 3 | AAATCAGA/TCTGATTT | | 7 |
| --- | | | | | | | | | 0 |
| chr4 | 166592083 | AGCAATTTGTGGA-AATAATTT-ACCATAGAATTAT | loxF8L | 3 | loxF8L | 3 | AAATTATT/AATAATTT | | 6 |
| chr3 | 87457652 | CTAATATTATTTG-AAATTATT-TCCATATAATTAT | loxF8R | 3 | loxF8L | 3 | AAATTATT/AATAATTT | | 6 |
| chr1 | 102325282 | ATAAATCTCTATA-AATAATTT-TATAGAGATTTTT | loxF8L | 3 | loxF8L | 4 | AAATTATT/AATAATTT | | 7 |
| chrY | 7510031 | ATAAATCTCTTCA-AATAATTT-TCCAGAGTTTGAC | loxF8L | 3 | loxF8L | 4 | AAATTATT/AATAATTT | | 7 |
| --- | | | | | | | | | 0 |
| chr14 | 84305885 | ATAAATTCCTGGA-AACACAGC-TCCAAAGATTGAA | loxF8L | 3 | loxF8L | 3 | AACACAGC/GCTGTGTT | | 6 |
| chr12 | 28512547 | AAAGTTCTGTTGA-AACACAGC-TTCACCCATTTAT | loxF8L | 4 | loxF8L | 3 | AACACAGC/GCTGTGTT | | 7 |
| --- | | | | | | | | | 0 |
| chr3 | 111054121 | CAAAATCTGTGAA-AACAGATT-TTCACAGATTTGA | loxF8L | 3 | loxF8L | 3 | AACAGATT/AATCTGTT | | 6 |
| chr12 | 79530724 | CTTAATCTTTGGA-AATCTGTT-TGCTCCATCTTAG | loxF8L | 3 | loxF8R | 4 | AACAGATT/AATCTGTT | | 7 |
| --- | | | | | | | | | 0 |
| chr8 | 62026200 | ACAAATATGTGCA-AACATCAC-CAGTCAATTTTAG | loxF8L | 3 | loxF8R | 3 | AACATCAC/GTGATGTT | | 6 |
| chr11 | 104429305 | TTAAGTTTATGTT-GTGATGTT-TTGACACATTTAT | loxF8R | 4 | loxF8L | 3 | AACATCAC/GTGATGTT | | 7 |
| --- | | | | | | | | | 0 |
| chr7 | 122577601 | ATAAAGCTGTTTA-AACATTCA-CACACAGATTTTT | loxF8L | 3 | loxF8L | 3 | AACATTCA/TGAATGTT | | 6 |
| chr17 | 18909933 | AGAGATCTGTGTA-TGAATGTT-CACAGCAGCTTTG | loxF8L | 3 | loxF8R | 4 | AACATTCA/TGAATGTT | | 7 |
| chr10 | 6920255 | CTAGGGTAGTGGG-TGAATGTT-CACACAAGCTGTG | loxF8R | 4 | loxF8R | 3 | AACATTCA/TGAATGTT | | 7 |
| chr9 | 83661015 | GTCAATCTGTGAA-TGAATGTT-GAGACAATGTTAC | loxF8L | 3 | loxF8R | 4 | AACATTCA/TGAATGTT | | 7 |
| --- | | | | | | | | | 0 |
| chr4 | 68886998 | GTAAATTTATGGA-TGGTAGTT-TTGACAGATGTAT | loxF8L | 3 | loxF8L | 3 | AACTACCA/TGGTAGTT | | 6 |
| chr15 | 24518648 | TTAAGTCCATGGA-TGGTAGTT-TCCACAGAATACT | loxF8L | 4 | loxF8L | 3 | AACTACCA/TGGTAGTT | | 7 |
| chr15 | 24311591 | TTAAGTCCATGGA-TGGTAGTT-TCCACAGAATACT | loxF8L | 4 | loxF8L | 3 | AACTACCA/TGGTAGTT | | 7 |
| chr3 | 84470537 | ATAATTCTGCCAA-AACTACCA-TCCACAGACTCAC | loxF8L | 4 | loxF8L | 3 | AACTACCA/TGGTAGTT | | 7 |
| chrX | 62852126 | ATGCATCTGTCGA-AACTACCA-TCCATCGAATTAC | loxF8L | 3 | loxF8L | 4 | AACTACCA/TGGTAGTT | | 7 |
| chr15 | 23996379 | GTATGTCTATGGA-TGGTAGTT-TTCACAGAATAAT | loxF8L | 4 | loxF8L | 3 | AACTACCA/TGGTAGTT | | 7 |
| --- | | | | | | | | | 0 |
| chr11 | 12622404 | ATCAAACTGTCGA-AAGACAAA-CAGAGAATCTTAA | loxF8L | 3 | loxF8R | 3 | AAGACAAA/TTTGTCTT | | 6 |
| chr1 | 235441164 | CTAAGAATGTTAG-AAGACAAA-AGCACAGATAGAT | loxF8R | 3 | loxF8L | 4 | AAGACAAA/TTTGTCTT | | 7 |
| --- | | | | | | | | | 0 |
| chr13 | 65870496 | CTATAATTGTCTG-TTTCTCTT-TCTATAGATATAT | loxF8R | 3 | loxF8L | 3 | AAGAGAAA/TTTCTCTT | | 6 |
| chr12 | 91651394 | GTAAATCTTTGCA-AAGAGAAA-TACTGAGATTTGT | loxF8L | 3 | loxF8L | 4 | AAGAGAAA/TTTCTCTT | | 7 |
| --- | | | | | | | | | 0 |
| chr1 | 43155881 | CTTAGATTCTATG-TGACTCTT-TCAACAAATTCAT | loxF8R | 3 | loxF8L | 3 | AAGAGTCA/TGACTCTT | | 6 |
| chr18 | 21790865 | GTAAGATTATGGA-TGACTCTT-TCCCCAAATTTGT | loxF8R | 4 | loxF8L | 3 | AAGAGTCA/TGACTCTT | | 7 |
| --- | | | | | | | | | 0 |
| chrX | 50752343 | ATATCTCTGGGGA-AAGCAGAC-AACACAATTTGAG | loxF8L | 3 | loxF8R | 3 | AAGCAGAC/GTCTGCTT | | 6 |
| chr7 | 118905901 | CTAAAACTGTAGA-AAGCAGAC-TCCTCAGAGTTAA | loxF8L | 3 | loxF8L | 3 | AAGCAGAC/GTCTGCTT | | 6 |
| chrX | 51652939 | AGTAGTCTGTGGT-GTCTGCTT-TTCTCACATTTAT | loxF8L | 4 | loxF8L | 3 | AAGCAGAC/GTCTGCTT | | 7 |
| --- | | | | | | | | | 0 |
| chr2 | 175014157 | TTAGGAATGTGTG-GAATGCTT-TCCATACATTTTT | loxF8R | 3 | loxF8L | 3 | AAGCATTC/GAATGCTT | | 6 |
| chr2 | 222161410 | CTGAGATTGTTTC-AAGCATTC-CATACAATATTTC | loxF8R | 3 | loxF8R | 4 | AAGCATTC/GAATGCTT | | 7 |
| --- | | | | | | | | | 0 |
| chr12 | 75591636 | CCAATATTGTGTA-CATAGCTT-TCTACACATTTTT | loxF8R | 3 | loxF8L | 3 | AAGCTATG/CATAGCTT | | 6 |
| chr7 | 31383599 | ATAAATCAGGGCA-AAGCTATG-TTAACCTATTTAT | loxF8L | 3 | loxF8L | 4 | AAGCTATG/CATAGCTT | | 7 |
| chr4 | 20263758 | GTAAGATAATGTG-CATAGCTT-CACAAAATGCTGG | loxF8R | 3 | loxF8R | 4 | AAGCTATG/CATAGCTT | | 7 |
| --- | | | | | | | | | 0 |
| chr17 | 7003955 | TTAACTATGTGGA-AAGGAAGA-CAAACAGTCTAAG | loxF8L | 3 | loxF8R | 3 | AAGGAAGA/TCTTCCTT | | 6 |
| chr1 | 95520337 | ATAAAAGTGTAGA-AAGGAAGA-AACACAGGTATAT | loxF8L | 3 | loxF8L | 4 | AAGGAAGA/TCTTCCTT | | 7 |
| --- | | | | | | | | | 0 |
| chr3 | 134982774 | CTCTGATTTTGTG-TGTTACTT-TCCAAAGAGATAT | loxF8R | 3 | loxF8L | 3 | AAGTAACA/TGTTACTT | | 6 |
| chr15 | 71075569 | CCAAATATTTGGA-AAGTAACA-CACATAAAATTAG | loxF8L | 4 | loxF8R | 3 | AAGTAACA/TGTTACTT | | 7 |
| --- | | | | | | | | | 0 |
| chr8 | 126146957 | TTAATTTTGTGTG-AAGTACTT-AACAAAATTTTAG | loxF8R | 3 | loxF8R | 3 | AAGTACTT/AAGTACTT | | 6 |
| chr11 | 20525988 | TTATATATGTGTA-AAGTACTT-AACACAATCCTAA | loxF8L | 4 | loxF8R | 3 | AAGTACTT/AAGTACTT | | 7 |
| --- | | | | | | | | | 0 |
| chr6 | 20785762 | TTAAATATGTAGA-TTTCACTT-TCCATAAATTTTT | loxF8L | 3 | loxF8L | 3 | AAGTGAAA/TTTCACTT | | 6 |
| chr10 | 87813515 | ATAAAATTGTGGT-TTTCACTT-TTTACTGATTTAA | loxF8L | 3 | loxF8L | 4 | AAGTGAAA/TTTCACTT | | 7 |
| chr1 | 205907608 | CTGAAACTGTGGA-AAGTGAAA-CCCACAGTTTCAG | loxF8L | 3 | loxF8L | 4 | AAGTGAAA/TTTCACTT | | 7 |
| --- | | | | | | | | | 0 |
| chr16 | 61832869 | ATAAAAATTTGGA-AATAAATA-CACACAAAGATAG | loxF8L | 3 | loxF8R | 3 | AATAAATA/TATTTATT | | 6 |
| chr6 | 76123480 | GTATGTTTGTGTG-TATTTATT-CACTTATTCTGAG | loxF8R | 3 | loxF8R | 4 | AATAAATA/TATTTATT | | 7 |
| --- | | | | | | | | | 0 |
| chr11 | 132372152 | TTAAGAGTGTGTT-GTCTTATT-TCTACATATTTGT | loxF8R | 3 | loxF8L | 3 | AATAAGAC/GTCTTATT | | 6 |
| chr13 | 22365730 | CAAAGATTTTATG-AATAAGAC-CTCAGAAGCTTAG | loxF8R | 3 | loxF8R | 3 | AATAAGAC/GTCTTATT | | 6 |
| chr4 | 13451617 | CTATGTTTGTGAG-GTCTTATT-CATAAAATCTTTT | loxF8R | 3 | loxF8R | 4 | AATAAGAC/GTCTTATT | HG1R | 7 |
| chr5 | 88318010 | CTATGTTTTTGAG-GTCTTATT-CACAAAATCTTTT | loxF8R | 4 | loxF8R | 3 | AATAAGAC/GTCTTATT | HG1R | 7 |
| chr1 | 57504806 | CTTTGCTTTTGTG-GTCTTATT-TCCAAAAATCTAT | loxF8R | 4 | loxF8L | 3 | AATAAGAC/GTCTTATT | | 7 |
| --- | | | | | | | | | 0 |
| chr9 | 103365724 | ATATAATTGTGTG-TGTGTATT-TCCACTGATTGCT | loxF8R | 3 | loxF8L | 3 | AATACACA/TGTGTATT | | 6 |
| chr5 | 123547448 | CTGTGTGTGTGTG-TGTGTATT-CACACAAAGCTAG | loxF8R | 4 | loxF8R | 3 | AATACACA/TGTGTATT | | 7 |
| --- | | | | | | | | | 0 |
| chr7 | 15904709 | ATAAATATGTATA-TTTCTATT-AACACAATATTAA | loxF8L | 3 | loxF8R | 3 | AATAGAAA/TTTCTATT | | 6 |
| chr2 | 106833784 | ATAAATATATATA-TTTCTATT-TCTACTGATTTTT | loxF8L | 4 | loxF8L | 3 | AATAGAAA/TTTCTATT | | 7 |
| chr17 | 51419286 | ATATATATGTAGC-AATAGAAA-TACATAGTTTTAT | loxF8L | 4 | loxF8L | 3 | AATAGAAA/TTTCTATT | | 7 |
| chrX | 127511798 | ATAAAGTTTTGGT-TTTCTATT-ACCACAACTTTAT | loxF8L | 4 | loxF8L | 3 | AATAGAAA/TTTCTATT | | 7 |
| --- | | | | | | | | | 0 |
| chr21 | 23494720 | CTAGATCTATGGA-TTCCTATT-ACCACATATATAT | loxF8L | 3 | loxF8L | 3 | AATAGGAA/TTCCTATT | | 6 |

| | | | | | | | | |
|---|---|---|---|---|---|---|---|---|
| chr17 | 51419864 | ACCAAGCTGAGGA-TTCCTATT-TCCATTTATTTAT | loxF8L | 4 | loxF8L | 3 | AATAGGAA/TTCCTATT | 7 |
| --- | | | | | | | | 0 |
| chr8 | 120919650 | ATAAATGTGTATA-GTTATATT-TACATAAATTTAT | loxF8L | 3 | loxF8L | 3 | AATATAAC/GTTATATT | 6 |
| chr18 | 10667113 | AAAAATATGTGGG-AATATAAC-CACAGATACTTAT | loxF8L | 3 | loxF8R | 4 | AATATAAC/GTTATATT | 7 |
| chr3 | 110870725 | AGAAGTTTGTGAA-AATATAAC-TCTACTGATTTCT | loxF8L | 4 | loxF8L | 3 | AATATAAC/GTTATATT | 7 |
| chr13 | 62677817 | ATAAATCTGGAAA-AATATAAC-TATAGATATTTAT | loxF8L | 3 | loxF8L | 4 | AATATAAC/GTTATATT | 7 |
| chr8 | 120919684 | ATAAATATGTATA-GTTATATT-TACATAAATATAT | loxF8L | 3 | loxF8L | 4 | AATATAAC/GTTATATT | 7 |
| --- | | | | | | | | 0 |
| chrX | 115630723 | ATATATGTGTGTA-TATATATT-TACACATATATAT | loxF8L | 3 | loxF8L | 3 | AATATATA/TATATATT | 6 |
| chr13 | 45019078 | ATAAATATATGTA-TATATATT-TATATATATTTAT | loxF8L | 3 | loxF8L | 4 | AATATATA/TATATATT | 7 |
| chr16 | 63054997 | ATATATATGTATA-TATATATT-TACATATATTTAT | loxF8L | 4 | loxF8L | 3 | AATATATA/TATATATT | 7 |
| chr5 | 4190460 | ATAAATATATTTA-TATATATT-TATACATATTTAT | loxF8L | 4 | loxF8L | 3 | AATATATA/TATATATT | 7 |
| chr5 | 4190496 | ATAAATATATTTA-TATATATT-TATACATATTTAT | loxF8L | 4 | loxF8L | 3 | AATATATA/TATATATT | 7 |
| chr20 | 38780641 | ATAAATATGTGTG-TATATATT-CAAATAATGTTAA | loxF8L | 3 | loxF8R | 4 | AATATATA/TATATATT | 7 |
| chr11 | 13059586 | ATATATGTGTATA-TATATATT-TATACACATTTAT | loxF8L | 4 | loxF8L | 3 | AATATATA/TATATATT | 7 |
| chr4 | 84169954 | ATATATGTGTGTA-TATATATT-TACATATATGTAT | loxF8L | 3 | loxF8L | 4 | AATATATA/TATATATT | 7 |
| chr17 | 33546112 | ATATATATATGTA-AATATATA-TACACATATGTAT | loxF8L | 4 | loxF8L | 3 | AATATATA/TATATATT | 7 |
| chr14 | 80960632 | ATACCTATGTGTA-TATATATT-TAGACAGATATAT | loxF8L | 4 | loxF8L | 3 | AATATATA/TATATATT | 7 |
| chr14 | 32800091 | ATATATATATAGA-AATATATA-TACACATATATAT | loxF8L | 4 | loxF8L | 3 | AATATATA/TATATATT | 7 |
| chr1 | 205551964 | ATATATATGTGTA-TATATATT-TGCATATATATAT | loxF8L | 3 | loxF8L | 4 | AATATATA/TATATATT | 7 |
| chr15 | 43842567 | ATATATATGTAAA-AATATATA-CATACTAACTTAG | loxF8L | 4 | loxF8R | 3 | AATATATA/TATATATT | 7 |
| chr8 | 68158148 | ATATATATGTGTA-TATATATT-CACACACACATAT | loxF8L | 3 | loxF8R | 4 | AATATATA/TATATATT | 7 |
| chr5 | 180735741 | ATATATATGTGTA-TATATATT-CACACACATATAT | loxF8L | 3 | loxF8L | 4 | AATATATA/TATATATT | 7 |
| chr14 | 54345948 | ATCAATCTCTTTA-AATATATA-TACACATATATAT | loxF8L | 4 | loxF8L | 3 | AATATATA/TATATATT | 7 |
| chr4 | 67184633 | ATATATATGTGTA-AATATATA-TACACACACATAT | loxF8L | 3 | loxF8L | 4 | AATATATA/TATATATT | 7 |
| chr17 | 69394378 | GTATGTGTGTGTG-AATATATA-TACACACATATAT | loxF8R | 4 | loxF8L | 3 | AATATATA/TATATATT | 7 |
| --- | | | | | | | | 0 |
| chr2 | 161252753 | ACATCTCTGTGGA-CAAATATT-TGCATACATTTAT | loxF8L | 3 | loxF8L | 3 | AATATTTG/CAAATATT | 6 |
| chr4 | 176758236 | ATATATGTGAGGA-CAAATATT-TCCTCAGTTTTGA | loxF8L | 3 | loxF8L | 4 | AATATTTG/CAAATATT | 7 |
| chr2 | 2381885 | CTAAAGGTATGGA-CAAATATT-AACACAATTTTAA | loxF8L | 4 | loxF8R | 3 | AATATTTG/CAAATATT | 7 |
| --- | | | | | | | | 0 |
| chr11 | 84950529 | ATAAATCTATATA-CATTCATT-AACAAAGATTTAT | loxF8L | 3 | loxF8L | 3 | AATGAATG/CATTCATT | 6 |
| chr11 | 115207195 | GTGAGATTCTGGG-CATTCATT-ACCACAAATGTAT | loxF8R | 4 | loxF8L | 3 | AATGAATG/CATTCATT | 7 |
| --- | | | | | | | | 0 |
| chr3 | 60618197 | ATAATTCTTTGGC-TATTTTGT-TTTACAGGTTTAT | loxF8L | 3 | loxF8L | 3 | ACAAAATA/TATTTTGT | 6 |
| chr18 | 29661338 | ATACATCTTAGGA-ACAAAATA-TCAACTGAATTAC | loxF8L | 3 | loxF8L | 4 | ACAAAATA/TATTTTGT | 7 |
| chr6 | 13764416 | CTAAGTTTGGGGG-TATTTTGT-TACACAGCAATAT | loxF8R | 3 | loxF8L | 4 | ACAAAATA/TATTTTGT | 7 |
| chr3 | 2955370 | CTGATTTTGAGTG-ACAAAATA-CACACAAACTCAT | loxF8R | 4 | loxF8R | 3 | ACAAAATA/TATTTTGT | 7 |
| --- | | | | | | | | 0 |
| chr1 | 181395880 | CTATCATTGTGTA-ACAAATCA-CTCAAAAACTTAG | loxF8R | 3 | loxF8R | 3 | ACAAATCA/TGATTTGT | 6 |
| chr20 | 6733884 | ATAAAGCTGCATA-ACAAATCA-CCCCAAATCTTAG | loxF8L | 4 | loxF8R | 3 | ACAAATCA/TGATTTGT | 7 |
| chr6 | 147176755 | CTAAATCTGTACT-TGATTTGT-TCCACAGGTTTCA | loxF8L | 4 | loxF8L | 3 | ACAAATCA/TGATTTGT | 7 |
| --- | | | | | | | | 0 |
| chr3 | 29102503 | GTACAGCTGTGGA-GTGGTTGT-CCCAAAGCTTTAT | loxF8L | 3 | loxF8L | 3 | ACAACCAC/GTGGTTGT | 6 |
| chr3 | 54963118 | ATAAATCTGGAGG-GTGGTTGT-TGCATTGATTTCT | loxF8L | 3 | loxF8L | 4 | ACAACCAC/GTGGTTGT | 7 |
| --- | | | | | | | | 0 |
| chr4 | 182178872 | ATACATCTGTAAA-TTTTCTGT-TCCAGATATTTAA | loxF8L | 3 | loxF8L | 3 | ACAGAAAA/TTTTCTGT | 6 |
| chr9 | 100041713 | ATATATCAGTGCA-ACAGAAAA-TCCAGAAATAAAT | loxF8L | 3 | loxF8L | 4 | ACAGAAAA/TTTTCTGT | 7 |
| chr12 | 70828338 | CTAATATTGTCTT-ACAGAAAA-AACACATTCTTCA | loxF8R | 3 | loxF8R | 4 | ACAGAAAA/TTTTCTGT | 7 |
| chrX | 95910135 | CTAATATTGTGAA-TTTTCTGT-CACAGAATATGAA | loxF8R | 3 | loxF8R | 4 | ACAGAAAA/TTTTCTGT | 7 |
| --- | | | | | | | | 0 |
| chr11 | 19925868 | CAAAAATTGTTTG-ACCATATG-TGCAAAGGTTTAT | loxF8R | 3 | loxF8L | 3 | ACCATATG/CATATGGT | 6 |
| chr7 | 148356042 | ACAATTCTGTGCA-CATATGGT-TACACACACATAG | loxF8L | 3 | loxF8R | 4 | ACCATATG/CATATGGT | 7 |
| chr4 | 66078066 | ATAAACCTAAGCA-CATATGGT-CACCTAATCTTAA | loxF8L | 4 | loxF8R | 3 | ACCATATG/CATATGGT | 7 |
| --- | | | | | | | | 0 |
| chr2 | 202241917 | CTAAAATTGATTG-GTGATGGT-TGCACAGCTATAT | loxF8R | 3 | loxF8L | 3 | ACCATCAC/GTGATGGT | 6 |
| chr7 | 112555966 | CTAAAATGGACTG-GTGATGGT-TCCACATATGTGT | loxF8R | 4 | loxF8L | 3 | ACCATCAC/GTGATGGT | 7 |
| chr11 | 74126807 | ATAAATGTGACCA-ACCATCAC-CACTCAATTTTAA | loxF8L | 4 | loxF8R | 3 | ACCATCAC/GTGATGGT | 7 |
| chr12 | 109838677 | CTAAAATTGACTG-GTGATGGT-TGCTCATATGTAT | loxF8R | 3 | loxF8L | 4 | ACCATCAC/GTGATGGT | 7 |
| --- | | | | | | | | 0 |
| chr7 | 12991481 | GTCAGATTTTGTG-GTAATGGT-TACACAATCCTAT | loxF8R | 3 | loxF8R | 3 | ACCATTAC/GTAATGGT | 6 |
| chr1 | 34940394 | ATACATCTTTTGA-GTAATGGT-TACATAGATATAC | loxF8L | 3 | loxF8L | 4 | ACCATTAC/GTAATGGT | 7 |
| --- | | | | | | | | 0 |
| chr11 | 24130525 | CTAAGATTAGTTG-ATTGCAGT-TACACGGCTTTAT | loxF8R | 3 | loxF8L | 3 | ACTGCAAT/ATTGCAGT | 6 |
| chr7 | 107711704 | CAAAGATATTGTG-ACTGCAAT-GACACAACCTTCT | loxF8R | 3 | loxF8R | 4 | ACTGCAAT/ATTGCAGT | 7 |
| --- | | | | | | | | 0 |
| chr1 | 71764109 | ATATAACTATGGA-AGAAGAAA-CACACAATTTTTC | loxF8L | 3 | loxF8R | 3 | AGAAGAAA/TTTCTTCT | 6 |
| chr6 | 22877197 | ATAAGAAGGTGTG-AGAAGAAA-CACTCAATATGGG | loxF8R | 3 | loxF8R | 4 | AGAAGAAA/TTTCTTCT | 7 |
| --- | | | | | | | | 0 |
| chr7 | 9707071 | ATAAATATGTAGT-AGAAGGAA-TCTCCAGATTTCT | loxF8L | 3 | loxF8L | 3 | AGAAGGAA/TTCCTTCT | 6 |
| chr3 | 167497542 | ATTCATCTGTGGC-TTCCTTCT-GCCACAGATTATA | loxF8L | 3 | loxF8L | 4 | AGAAGGAA/TTCCTTCT | 7 |
| --- | | | | | | | | 0 |
| chr14 | 37449258 | AGAAATCTTTGGC-AGAGACAG-TCCACAAAATAAT | loxF8L | 3 | loxF8L | 3 | AGAGACAG/CTGTCTCT | 6 |
| chr6 | 108408515 | AAAACTCTGAGGA-CTGTCTCT-TCAACTCATTTAG | loxF8L | 3 | loxF8L | 4 | AGAGACAG/CTGTCTCT | 7 |
| chr12 | 117669154 | ATAAAGCCAGGGA-CTGTCTCT-TACACAGAATCAT | loxF8L | 4 | loxF8L | 3 | AGAGACAG/CTGTCTCT | 7 |
| --- | | | | | | | | 0 |
| chr7 | 10061664 | ATAAATGGGTGAA-AGAGTAAA-TTAACACATTTAT | loxF8L | 3 | loxF8L | 3 | AGAGTAAA/TTTACTCT | 6 |
| chr3 | 86948125 | ATAATTGTTTGGA-AGAGTAAA-AACACAGAATAAT | loxF8L | 3 | loxF8L | 4 | AGAGTAAA/TTTACTCT | 7 |
| --- | | | | | | | | 0 |
| chrY | 19317965 | ATAGAACTCTGGA-TATTATCT-CAGACAATCAGAG | loxF8L | 3 | loxF8R | 3 | AGATAATA/TATTATCT | 6 |
| chrX | 98012429 | CTAATATAGTCTA-TATTATCT-TCCATTGATTGAT | loxF8R | 4 | loxF8L | 3 | AGATAATA/TATTATCT | 7 |
| --- | | | | | | | | 0 |
| chr5 | 99095272 | ATATATATGTGGC-AGGAATAT-CACAGAAGCATAG | loxF8L | 3 | loxF8R | 3 | AGGAATAT/ATATTCCT | 6 |
| chr13 | 102963607 | CTATAATTCTGTG-AGGAATAT-TCCAGTGTTTTAG | loxF8R | 3 | loxF8L | 4 | AGGAATAT/ATATTCCT | 7 |
| --- | | | | | | | | 0 |
| chr5 | 95122087 | CAAAGATCGTTTG-AGTAATAC-CCCACAAGCTCAG | loxF8R | 3 | loxF8R | 3 | AGTAATAC/GTATTACT | 6 |
| chr12 | 10456352 | CTGTGATTGTGGG-GTATTACT-CAAAAAATCTTTG | loxF8R | 3 | loxF8R | 3 | AGTAATAC/GTATTACT | 6 |

| | | | | | | | | |
|---|---|---|---|---|---|---|---|---|
| chrX | 88193620 | CAAAGATTTTTTG-AGTAATAC-CCTACAAGCTTAG | loxF8R | 3 | loxF8R | 3 | AGTAATAC/GTATTACT | 6 |
| chr3 | 110535820 | CAAAGATTTATTG-AGTAATAC-CCCACAAGCGTAG | loxF8R | 4 | loxF8R | 3 | AGTAATAC/GTATTACT | 7 |
| chrX | 57645302 | CAAAGATTTCCTG-AGTAATAC-CACACAAGCACAG | loxF8R | 4 | loxF8R | 3 | AGTAATAC/GTATTACT | 7 |
| chr14 | 57434370 | CTTTGCTTGTGTG-GTATTACT-CAAGAAATCTTTG | loxF8R | 3 | loxF8R | 4 | AGTAATAC/GTATTACT | 7 |
| chr7 | 31140691 | CTATGCTTGTCTG-GTATTACT-CAAGAAATCTTTG | loxF8R | 3 | loxF8R | 4 | AGTAATAC/GTATTACT | 7 |
| chr10 | 61648977 | CTATGCTTGTGGG-GTATTACT-CAAGAAATCTTTG | loxF8R | 3 | loxF8R | 4 | AGTAATAC/GTATTACT | 7 |
| chr11 | 7054841 | CTATGCTTGTGGG-GTATTACT-CAAGAAATCTTTG | loxF8R | 3 | loxF8R | 4 | AGTAATAC/GTATTACT | 7 |
| chr6 | 85222791 | CAAAGATTTCTTG-AGTAATAC-CCCACAAGCATAG | loxF8R | 4 | loxF8R | 3 | AGTAATAC/GTATTACT | 7 |
| chr1 | 86671151 | CAAAGATTTCTTG-AGTAATAC-CCCACAAGCTCAG | loxF8R | 4 | loxF8R | 3 | AGTAATAC/GTATTACT | 7 |
| chr11 | 87704995 | CTGTGATTGTGGG-GTATTACT-CAAGAAATCTTTG | loxF8R | 3 | loxF8R | 4 | AGTAATAC/GTATTACT | 7 |
| chrX | 21414599 | CTGTGATTGTGGG-GTATTACT-CAAGAAATCTTTG | loxF8R | 3 | loxF8R | 4 | AGTAATAC/GTATTACT | 7 |
| chrX | 131109269 | CTATGCTTGTGAG-GTATTACT-CAAGAAATCTTTG | loxF8R | 3 | loxF8R | 4 | AGTAATAC/GTATTACT | 7 |
| chrX | 105333454 | CAAAGATTTTCTG-AGTAATAC-CCCACAAGCACAG | loxF8R | 3 | loxF8R | 4 | AGTAATAC/GTATTACT | 7 |
| chr7 | 145278833 | CTCTGCTTATGTG-GTATTACT-CAAAAAATCTTTG | loxF8R | 4 | loxF8R | 3 | AGTAATAC/GTATTACT | 7 |
| chr7 | 139207258 | CAAAGATTTTTTG-AGTAATAC-CCCACAAACACAG | loxF8R | 3 | loxF8R | 4 | AGTAATAC/GTATTACT | 7 |
| chr6 | 123167256 | CAAAGATTTTTTG-AGTAATAC-CCCACAAGCACAG | loxF8R | 3 | loxF8R | 4 | AGTAATAC/GTATTACT | 7 |
| chr6 | 118950186 | CAAAGATTTTTTG-AGTAATAC-CCCACAAGCAGAG | loxF8R | 3 | loxF8R | 4 | AGTAATAC/GTATTACT | 7 |
| chr5 | 102566183 | CTGTGCTTGTGAG-GTATTACT-CAAAAAATCTTTG | loxF8R | 4 | loxF8R | 3 | AGTAATAC/GTATTACT | 7 |
| chr1 | 49689110 | CTATGTTTGTGGA-GTATTACT-CAAAAAATCTTTG | loxF8L | 4 | loxF8R | 3 | AGTAATAC/GTATTACT | 7 |
| chrX | 76358704 | CTGTGTTTGTGGG-GTATTACT-CAGAAAATTTTAG | loxF8R | 4 | loxF8R | 3 | AGTAATAC/GTATTACT | 7 |
| chr2 | 157976962 | CTGTGCTTGTGGG-GTATTACT-CAAGAAATCTTAG | loxF8R | 4 | loxF8R | 3 | AGTAATAC/GTATTACT | 7 |
| --- | | | | | | | | 0 |
| chr11 | 17192968 | CTACGACTGTGGG-AGTACACT-CACACAATGTCAT | loxF8R | 3 | loxF8R | 3 | AGTACACT/AGTGTACT | 6 |
| chr4 | 97353465 | CTAGGTTTGTGTT-AGTACACT-CACACAAACATGA | loxF8R | 3 | loxF8R | 4 | AGTACACT/AGTGTACT | 7 |
| --- | | | | | | | | 0 |
| chr4 | 1260231 | TTAAGAGTGTGTT-GTTTAACT-TCCAGAAATTTGT | loxF8R | 3 | loxF8L | 3 | AGTTAAAC/GTTTAACT | 6 |
| chr9 | 95951182 | ACAAATATGTAGA-AGTTAAAC-AACACACACTTAA | loxF8L | 3 | loxF8R | 4 | AGTTAAAC/GTTTAACT | 7 |
| chr7 | 95624923 | ATGATTGTGTGGA-AGTTAAAC-AACACACTCCTAA | loxF8L | 3 | loxF8R | 4 | AGTTAAAC/GTTTAACT | 7 |
| --- | | | | | | | | 0 |
| chr7 | 102240725 | CTCAGATTCTGAG-CTAAGTAT-TTCACCGAGTTAT | loxF8R | 3 | loxF8L | 3 | ATACTTAG/CTAAGTAT | 6 |
| chr9 | 76873949 | TTAAAACTATGGA-ATACTTAG-AGCACAGACATAT | loxF8L | 3 | loxF8L | 4 | ATACTTAG/CTAAGTAT | 7 |
| --- | | | | | | | | 0 |
| chr4 | 135669017 | CGAAGCTTCTGTG-ATTTATAT-TCTTCAGATTTTT | loxF8R | 3 | loxF8L | 3 | ATATAAAT/ATTTATAT | 6 |
| chrX | 147514028 | AATAATCTGTTAA-ATTTATAT-TCCAATGATTTCT | loxF8L | 4 | loxF8L | 3 | ATATAAAT/ATTTATAT | 7 |
| chr14 | 45426590 | ATAAAATTGAGGA-ATATAAAT-GATACAATGTTAA | loxF8L | 3 | loxF8R | 4 | ATATAAAT/ATTTATAT | 7 |
| chr13 | 103902312 | ATAGAAATGTGCA-ATTTATAT-TCCAAATACTTAT | loxF8L | 4 | loxF8L | 3 | ATATAAAT/ATTTATAT | 7 |
| --- | | | | | | | | 0 |
| chr12 | 102500102 | ATAAAAATGTGGT-ATATATAC-CACAGAATATTAA | loxF8L | 3 | loxF8R | 3 | ATATATAC/GTATATAT | 6 |
| chr5 | 28328543 | AGTATTCTGTGGT-GTATATAT-ACCACAGTTTTAA | loxF8L | 4 | loxF8L | 3 | ATATATAC/GTATATAT | 7 |
| --- | | | | | | | | 0 |
| chr1 | 90689968 | ATAAATATATGAA-ATATATAT-TTCACATATATAT | loxF8L | 3 | loxF8L | 3 | ATATATAT/ATATATAT | 6 |
| chr4 | 115021185 | AAGAAACTGTGGT-ATATATAT-TCCATATATATAT | loxF8L | 4 | loxF8L | 3 | ATATATAT/ATATATAT | 7 |
| chr3 | 190297690 | ATAAATATATAAA-ATATATAT-TACATATATTTAT | loxF8L | 4 | loxF8L | 3 | ATATATAT/ATATATAT | 7 |
| chr3 | 19209152 | ATAAATGTGTAAA-ATATATAT-TACACCTTCTCAG | loxF8L | 3 | loxF8R | 4 | ATATATAT/ATATATAT | 7 |
| --- | | | | | | | | 0 |
| chr6 | 56053202 | ATAAAACTTTGGT-TAAGATAT-TCCAAAGATAAAT | loxF8L | 3 | loxF8L | 3 | ATATCTTA/TAAGATAT | 6 |
| chr10 | 85620457 | ATAGAACTGTGGC-ATATCTTA-CCCACTGATATTT | loxF8L | 3 | loxF8L | 4 | ATATCTTA/TAAGATAT | 7 |
| --- | | | | | | | | 0 |
| chr2 | 67955564 | ATAACTCTGTGTC-ATATGACA-CATACAATTTTAC | loxF8L | 3 | loxF8R | 3 | ATATGACA/TGTCATAT | 6 |
| chr20 | 20699289 | ATAAATATTTGCA-ATATGACA-CACACAGACAAAG | loxF8L | 3 | loxF8R | 4 | ATATGACA/TGTCATAT | 7 |
| --- | | | | | | | | 0 |
| chr4 | 105797268 | ATAAATAATTGGA-GTAAATAT-CACACAAGGATAG | loxF8L | 3 | loxF8R | 3 | ATATTTAC/GTAAATAT | 6 |
| chr4 | 173218713 | CTTAGAGTTTGTG-GTAAATAT-CAAAATATCTTAG | loxF8R | 3 | loxF8R | 3 | ATATTTAC/GTAAATAT | 6 |
| --- | | | | | | | | 0 |
| chr5 | 73086421 | TTAAGCTTCTGTG-CAGGGGAT-CACAGGATCTTAC | loxF8R | 3 | loxF8R | 3 | ATCCCCTG/CAGGGGAT | 6 |
| chr8 | 133681198 | GAACCTCTGTGGA-CAGGGGAT-TCCATAAATTGAT | loxF8L | 4 | loxF8L | 3 | ATCCCCTG/CAGGGGAT | 7 |
| --- | | | | | | | | 0 |
| chr8 | 5094848 | ATAAGATTATGTC-ATCTATAA-CACATAATTTTAC | loxF8R | 3 | loxF8R | 3 | ATCTATAA/TTATAGAT | 6 |
| chrX | 87154674 | ATAAAACTTTTGA-TTATAGAT-TAAATATATTTAT | loxF8L | 3 | loxF8L | 4 | ATCTATAA/TTATAGAT | 7 |
| --- | | | | | | | | 0 |
| chr12 | 104491991 | CAAAGGCTGTGTG-ATTTCCAT-CAGTCAGTCTTAG | loxF8R | 3 | loxF8R | 3 | ATGGAAAT/ATTTCCAT | 6 |
| chr14 | 81316366 | CTAACATTGTGGT-ATGGAAAT-CTCACATTGTTAA | loxF8R | 3 | loxF8R | 4 | ATGGAAAT/ATTTCCAT | 7 |
| --- | | | | | | | | 0 |
| chr4 | 173540165 | ATAAAATTGTGCA-ATGGCAAA-GCCACAGAATAAT | loxF8L | 3 | loxF8L | 3 | ATGGCAAA/TTTGCCAT | 6 |
| chr20 | 11065158 | ATAGTATTGTGTG-ATGGCAAA-GGCACAAACTTAG | loxF8R | 3 | loxF8R | 3 | ATGGCAAA/TTTGCCAT | 6 |
| --- | | | | | | | | 0 |
| chrX | 3505143 | GTAAGATTGGGTT-TTTCACAT-AACACTGATTTAT | loxF8R | 3 | loxF8L | 3 | ATGTGAAA/TTTCACAT | 6 |
| chr8 | 107228370 | AGAAATATGTGAA-ATGTGAAA-TCCAACGATTTCC | loxF8L | 3 | loxF8L | 4 | ATGTGAAA/TTTCACAT | 7 |
| --- | | | | | | | | 0 |
| chr12 | 60361202 | TCAAGATTCTGTG-ATGTTTGG-TCCACATATATAA | loxF8R | 3 | loxF8L | 3 | ATGTTTGG/CCAAACAT | 6 |
| chr12 | 73735901 | AGAAATATGTTGC-CCAAACAT-CATAAAATTTTAG | loxF8L | 4 | loxF8R | 3 | ATGTTTGG/CCAAACAT | 7 |
| --- | | | | | | | | 0 |
| chr18 | 66480356 | ATAAACTTGTGGC-ATTATAAG-TTCATTGATTTAT | loxF8L | 3 | loxF8L | 3 | ATTATAAG/CTTATAAT | 6 |
| chr6 | 21274984 | CTAAATCAGGGGA-ATTATAAG-CACCCAATTTAAC | loxF8L | 3 | loxF8R | 4 | ATTATAAG/CTTATAAT | 7 |
| --- | | | | | | | | 0 |
| chr14 | 35076971 | ATATCTTTGTGGA-ATTTTAAC-TCTACAGATTTTA | loxF8L | 3 | loxF8L | 3 | ATTTTAAC/GTTAAAAT | 6 |
| chr4 | 166258920 | AGATATATGTGTA-ATTTTAAC-ATCACATATTTAT | loxF8L | 4 | loxF8L | 3 | ATTTTAAC/GTTAAAAT | 7 |
| --- | | | | | | | | 0 |
| chr7 | 116424065 | CTCAGATTGTTTA-TGTATTTG-CACAGAGTGTTAG | loxF8R | 3 | loxF8R | 3 | CAAATACA/TGTATTTG | 6 |
| chr16 | 29269020 | CTAAGCCTGTCTC-CAAATACA-GTCACATTCTTAG | loxF8R | 4 | loxF8R | 3 | CAAATACA/TGTATTTG | 7 |
| --- | | | | | | | | 0 |
| chr6 | 149345864 | ATAATTATTTGGA-TATATTTG-TCCATATAATTAT | loxF8L | 3 | loxF8L | 3 | CAAATATA/TATATTTG | 6 |
| chr14 | 85348902 | ACACATTTGTGCA-TATATTTG-TGCACACATTTTT | loxF8L | 4 | loxF8L | 3 | CAAATATA/TATATTTG | 7 |
| chr1 | 198659994 | ATAAATATATAGA-TATATTTG-TCCATATATATGT | loxF8L | 3 | loxF8L | 4 | CAAATATA/TATATTTG | 7 |
| chr5 | 107800803 | TTATGAGTGTATG-TATATTTG-TGCACAGATATCT | loxF8R | 4 | loxF8L | 3 | CAAATATA/TATATTTG | 7 |
| chr14 | 25476887 | ATACACATGTGCA-TATATTTG-TACACATATGTAT | loxF8L | 4 | loxF8L | 3 | CAAATATA/TATATTTG | 7 |
| chr13 | 63485470 | CTAAAACTGTGAA-TATATTTG-CATACCATGTGAG | loxF8L | 3 | loxF8R | 4 | CAAATATA/TATATTTG | 7 |

| | | | | | | | | |
|---|---|---|---|---|---|---|---|---|
| --- | | | | | | | | 0 |
| chr21 | 38126621 | ATATATATGTGTA-CAATATCA-TACACACATATAT | loxF8L | 3 | loxF8L | 3 | CAATATCA/TGATATTG | 6 |
| chrX | 35712156 | CTAAGTTTATGAA-TGATATTG-GCCATATATTTAT | loxF8R | 4 | loxF8L | 3 | CAATATCA/TGATATTG | 7 |
| --- | | | | | | | | 0 |
| chr21 | 36341039 | ATAAAGCTGTAAA-CATTTGTG-TACACAGTTTTGT | loxF8L | 3 | loxF8L | 3 | CACAAATG/CATTTGTG | 6 |
| chr5 | 22900641 | ATACTACTGTGAA-CATTTGTG-TTCACATTTTTAT | loxF8L | 4 | loxF8L | 3 | CACAAATG/CATTTGTG | 7 |
| chr5 | 121634387 | ATAAATAAGTGTA-CACAAATG-CAAACAATACCAG | loxF8L | 3 | loxF8R | 4 | CACAAATG/CATTTGTG | 7 |
| chr8 | 134951445 | ATATATCGGTGGG-CACAAATG-AACACAATCCTTC | loxF8L | 3 | loxF8R | 4 | CACAAATG/CATTTGTG | 7 |
| --- | | | | | | | | 0 |
| chr21 | 14459264 | CTAAATCTGAGCA-CACACAAA-TCCAGAAATTTAA | loxF8L | 3 | loxF8L | 3 | CACACAAA/TTTGTGTG | 6 |
| chr5 | 120330337 | ATATATATATGCA-TTTGTGTG-TCCAAAATTTTAT | loxF8L | 4 | loxF8L | 3 | CACACAAA/TTTGTGTG | 7 |
| --- | | | | | | | | 0 |
| chr4 | 60572473 | ATATATGTGTGTA-TATGTGTG-TACACATATATAT | loxF8L | 3 | loxF8L | 3 | CACACATA/TATGTGTG | 6 |
| chr4 | 60572501 | ATATATGTGTGTA-TATGTGTG-TACACATATATAT | loxF8L | 3 | loxF8L | 3 | CACACATA/TATGTGTG | 6 |
| chr4 | 60572529 | ATATATGTGTGTA-TATGTGTG-TACACATATATAT | loxF8L | 3 | loxF8L | 3 | CACACATA/TATGTGTG | 6 |
| chr4 | 60572557 | ATATATGTGTGTA-TATGTGTG-TACACATATATAT | loxF8L | 3 | loxF8L | 3 | CACACATA/TATGTGTG | 6 |
| chr4 | 60572613 | ATATATGTGTGTA-TATGTGTG-TACACATATATAT | loxF8L | 3 | loxF8L | 3 | CACACATA/TATGTGTG | 6 |
| chr4 | 60572585 | ATATATGTGTGTA-TATGTGTG-TACACATATATAT | loxF8L | 3 | loxF8L | 3 | CACACATA/TATGTGTG | 6 |
| chr11 | 26701590 | AGAAATGTGTGTG-TATGTGTG-TACACATATTGAT | loxF8L | 4 | loxF8L | 3 | CACACATA/TATGTGTG | 7 |
| chr1 | 194502931 | ATACATTTTTGTA-CACACATA-TATACACATTTAT | loxF8L | 4 | loxF8L | 3 | CACACATA/TATGTGTG | 7 |
| chr14 | 104508910 | ATATATATATGCA-CACACATA-TACACATATATAT | loxF8L | 4 | loxF8L | 3 | CACACATA/TATGTGTG | 7 |
| chr21 | 38915366 | CTATATGTGTGTG-TATGTGTG-TACACATATATAT | loxF8R | 4 | loxF8L | 3 | CACACATA/TATGTGTG | 7 |
| chr4 | 60572641 | ATATATGTGTGTA-TATGTGTG-TGTACACATATAT | loxF8L | 3 | loxF8L | 4 | CACACATA/TATGTGTG | 7 |
| chr12 | 57347314 | ATATATGTGTGTA-TATGTGTG-CATACATACATAG | loxF8L | 3 | loxF8R | 4 | CACACATA/TATGTGTG | 7 |
| --- | | | | | | | | 0 |
| chr3 | 70244747 | CTTATATTTTGTG-CATGTGTG-TGCACACATATAT | loxF8R | 3 | loxF8L | 3 | CACACATG/CATGTGTG | 6 |
| chr9 | 2570236 | ATATATGTGTGCA-CACACATG-TGTACAGATGTGT | loxF8L | 3 | loxF8L | 4 | CACACATG/CATGTGTG | 7 |
| chr7 | 80763017 | ATAAATATGTACA-CACACATG-CACACACACATAC | loxF8L | 3 | loxF8R | 4 | CACACATG/CATGTGTG | 7 |
| chr4 | 95071926 | GTCTGTTTGTGTG-CATGTGTG-TAAAGAGATTTAT | loxF8R | 4 | loxF8L | 3 | CACACATG/CATGTGTG | 7 |
| chr19 | 42170684 | ATAAATGTATGAA-CACACATG-CACACAAACATGC | loxF8L | 3 | loxF8R | 4 | CACACATG/CATGTGTG | 7 |
| chr10 | 123043925 | ATACACCTGTGTA-CACACATG-CACACACACATAT | loxF8L | 3 | loxF8R | 4 | CACACATG/CATGTGTG | 7 |
| chrX | 111581874 | ATAGATGTATGCA-CATGTGTG-TGCACATATGTAT | loxF8L | 4 | loxF8L | 3 | CACACATG/CATGTGTG | 7 |
| chrX | 33141950 | GTATATATTTGGA-CACACATG-TCCACACATTTTA | loxF8L | 4 | loxF8L | 3 | CACACATG/CATGTGTG | 7 |
| --- | | | | | | | | 0 |
| chr16 | 11463389 | GTATGAATGTGTG-TGGATGTG-TGTACAGATGTAT | loxF8R | 3 | loxF8L | 3 | CACATCCA/TGGATGTG | 6 |
| chr7 | 74049124 | ATCCATCTGTCCA-CACATCCA-TCCATCCATTTAT | loxF8L | 4 | loxF8L | 3 | CACATCCA/TGGATGTG | 7 |
| --- | | | | | | | | 0 |
| chr1 | 159158944 | ATAAATCAGAGCA-CTCATGTG-TGAACAGATCTAT | loxF8L | 3 | loxF8L | 3 | CACATGAG/CTCATGTG | 6 |
| chr18 | 39081971 | CTACGATGGTATG-CACATGAG-TCTATGCATTTAT | loxF8R | 3 | loxF8L | 4 | CACATGAG/CTCATGTG | 7 |
| --- | | | | | | | | 0 |
| chr10 | 1405718 | CTAAGAAGCTGTG-CACCAAGA-TACACCCATTTAT | loxF8R | 3 | loxF8L | 3 | CACCAAGA/TCTTGGTG | 6 |
| chrX | 88376139 | CTAAGGTAGTGTA-CACCAAGA-CACATTATAATAG | loxF8R | 3 | loxF8R | 4 | CACCAAGA/TCTTGGTG | 7 |
| --- | | | | | | | | 0 |
| chr19 | 42865378 | CTAGGATTGTGAC-CACTGAAG-CACAGCATCATAG | loxF8R | 3 | loxF8R | 3 | CACTGAAG/CTTCAGTG | 6 |
| chr19 | 42923053 | CTAAGAGTGTGAC-CACTGAAG-CACAGCATCACAG | loxF8R | 3 | loxF8R | 4 | CACTGAAG/CTTCAGTG | 7 |
| chr19 | 42751602 | CTAGGAGTGTGAC-CACTGAAG-CACAGCATCATAG | loxF8R | 4 | loxF8R | 3 | CACTGAAG/CTTCAGTG | 7 |
| chr19 | 12712447 | CTGAGATGCTGTG-CTTCAGTG-GTCACACTCCTAG | loxF8R | 3 | loxF8R | 4 | CACTGAAG/CTTCAGTG | 7 |
| --- | | | | | | | | 0 |
| chr5 | 125980495 | GTAAGATACTGTG-CAGAAATG-GACACAGATTTAG | loxF8R | 3 | loxF8L | 3 | CAGAAATG/CATTTCTG | 6 |
| chr2 | 242060688 | ACAACTCTGAGGA-CATTTCTG-CAACCAATCTTTT | loxF8L | 3 | loxF8R | 4 | CAGAAATG/CATTTCTG | 7 |
| --- | | | | | | | | 0 |
| chr12 | 22580458 | ATAGATAGGTGGA-CAGATAAA-TACAGATATTTAT | loxF8L | 3 | loxF8L | 3 | CAGATAAA/TTTATCTG | 6 |
| chr17 | 32830648 | AAAAAACTCTGGG-CAGATAAA-CACAAAATATTAA | loxF8L | 4 | loxF8R | 3 | CAGATAAA/TTTATCTG | 7 |
| chr9 | 70108102 | ATATATCTGCAGG-TTTATCTG-CCCAGAGATTTAC | loxF8L | 4 | loxF8L | 3 | CAGATAAA/TTTATCTG | 7 |
| --- | | | | | | | | 0 |
| chrX | 51227003 | ATAAATATTTGGC-CAGCCATG-GAAACAATCATAG | loxF8L | 3 | loxF8R | 3 | CAGCCATG/CATGGCTG | 6 |
| chr16 | 56220103 | ATAGCTCTTTGGA-CAGCCATG-TGAACAGATATCT | loxF8L | 3 | loxF8L | 4 | CAGCCATG/CATGGCTG | 7 |
| --- | | | | | | | | 0 |
| chr17 | 65642617 | AAATATCTGTTGA-TTCTCCTG-TCTACTGATTTTT | loxF8L | 3 | loxF8L | 3 | CAGGAGAA/TTCTCCTG | 6 |
| chr7 | 36681996 | ATAAAACTGAGAA-CAGGAGAA-TCCCCAGATAGCT | loxF8L | 3 | loxF8L | 4 | CAGGAGAA/TTCTCCTG | 7 |
| --- | | | | | | | | 0 |
| chrY | 16293345 | ATAAATGTGAAGA-CAGTACTG-TGGATAGATTTAT | loxF8L | 3 | loxF8L | 3 | CAGTACTG/CAGTACTG | 6 |
| --- | | | | | | | | 0 |
| chr21 | 7795215 | CTAAAATTGTGGT-GCTCACTG-TACACAATCTGTG | loxF8R | 3 | loxF8R | 3 | CAGTGAGC/GCTCACTG | 6 |
| chr21 | 34425227 | CTAAAATTGTGGT-GCTCACTG-TACACAATCTGTG | loxF8R | 3 | loxF8R | 3 | CAGTGAGC/GCTCACTG | 6 |
| --- | | | | | | | | 0 |
| chr1 | 66637816 | ATATATGTGTGTA-CATATACA-TACACACATATAT | loxF8L | 3 | loxF8L | 3 | CATATACA/TGTATATG | 6 |
| chr5 | 135406991 | ATAAATATGTGTG-TGTATATG-TGCACATATATGT | loxF8L | 3 | loxF8L | 4 | CATATACA/TGTATATG | 7 |
| chr11 | 39492466 | ATAAATATATACA-CATATACA-CACACAAACATAT | loxF8L | 4 | loxF8R | 3 | CATATACA/TGTATATG | 7 |
| chr6 | 126058832 | ATAAATTCCTGGA-CATATACA-TCCCCAGTGGTAT | loxF8L | 3 | loxF8L | 4 | CATATACA/TGTATATG | 7 |
| chr6 | 34718391 | ATACATATGTGTA-TGTATATG-TATACATATGTAT | loxF8L | 3 | loxF8L | 4 | CATATACA/TGTATATG | 7 |
| chrX | 38578907 | ATACATATGTGGG-TGTATATG-TACACACATATAC | loxF8L | 3 | loxF8L | 4 | CATATACA/TGTATATG | 7 |
| chr20 | 13600766 | ATACATATGTGTA-TGTATATG-CACACATACATAC | loxF8L | 3 | loxF8R | 4 | CATATACA/TGTATATG | 7 |
| chr9 | 11673984 | ATATATATATGCA-CATATACA-TACAGAGATATAT | loxF8L | 4 | loxF8L | 3 | CATATACA/TGTATATG | 7 |
| chr17 | 15606646 | ATATATATGTGTG-TGTATATG-TACACACATATAT | loxF8L | 4 | loxF8L | 3 | CATATACA/TGTATATG | 7 |
| chr10 | 25423084 | GTATGTGTGTGTG-TGTATATG-TCTACACATATAT | loxF8R | 4 | loxF8L | 3 | CATATACA/TGTATATG | 7 |
| chr17 | 18757047 | ATATATGTGTGTA-CATATACA-CACACACATATAT | loxF8L | 3 | loxF8L | 4 | CATATACA/TGTATATG | 7 |
| chr3 | 36300379 | ATATATGTGTGTA-TGTATATG-TACACACACATAT | loxF8L | 3 | loxF8L | 4 | CATATACA/TGTATATG | 7 |
| --- | | | | | | | | 0 |
| chr5 | 3504954 | ATATATGTGTGTA-CCTATATG-TCCACATATTTTC | loxF8L | 3 | loxF8L | 3 | CATATAGG/CCTATATG | 6 |
| chr8 | 43374918 | CTACTTCTGTGTG-CCTATATG-TCTAAATATTTAT | loxF8R | 4 | loxF8L | 3 | CATATAGG/CCTATATG | 7 |
| --- | | | | | | | | 0 |
| chr5 | 22268262 | ATATATGTGTGTA-TATATATG-TACATAAATTTAT | loxF8L | 3 | loxF8L | 3 | CATATATA/TATATATG | 6 |
| chr1 | 106458142 | ATAAGTGTGTGTG-CATATATA-TACACATATATAT | loxF8R | 3 | loxF8L | 3 | CATATATA/TATATATG | 6 |
| chr21 | 30252713 | ATACATATGTGTA-TATATATG-TACACATATATAT | loxF8L | 3 | loxF8L | 3 | CATATATA/TATATATG | 6 |
| chr4 | 179679069 | ATATATATGTGTA-TATATATG-TACACACATATAT | loxF8L | 3 | loxF8L | 3 | CATATATA/TATATATG | 6 |
| chr4 | 179679041 | ATATATATGTGTA-TATATATG-TACACACATATAT | loxF8L | 3 | loxF8L | 3 | CATATATA/TATATATG | 6 |

| | | | | | | | | |
|---|---|---|---|---|---|---|---|---|
| chr2 | 194180196 | ATCAATGTGTGGG-TATATATG-TACACACATATAT | loxF8L | 3 | loxF8L | 3 | CATATATA/TATATATG | 6 |
| chr4 | 179679097 | ATATATGTGTGTA-TATATATG-TACACACATATAT | loxF8L | 3 | loxF8L | 3 | CATATATA/TATATATG | 6 |
| chr2 | 190921643 | ACAAATATGTAAA-TATATATG-TACATAGATTTTT | loxF8L | 4 | loxF8L | 3 | CATATATA/TATATATG | 7 |
| chr2 | 152229954 | ATAACTATGTAGA-TATATATG-TATAAAGATTTTT | loxF8L | 3 | loxF8L | 4 | CATATATA/TATATATG | 7 |
| chr8 | 48691409 | AAAAATGTATGCA-TATATATG-CACATAATTTTAC | loxF8L | 4 | loxF8R | 3 | CATATATA/TATATATG | 7 |
| chr6 | 66666925 | ATATATATGTGTA-TATATATG-TGTACATATTTGT | loxF8L | 3 | loxF8L | 4 | CATATATA/TATATATG | 7 |
| chr2 | 222622593 | ACAAATGTGTGTA-TATATATG-TGCAAAGATGAAT | loxF8L | 3 | loxF8L | 4 | CATATATA/TATATATG | 7 |
| chr4 | 179679149 | ACACATATGTGTA-TATATATG-TACACACATATAT | loxF8L | 4 | loxF8L | 3 | CATATATA/TATATATG | 7 |
| chr10 | 57693817 | ATATATATGTGTA-TATATATG-TCCATATATATGT | loxF8L | 3 | loxF8L | 4 | CATATATA/TATATATG | 7 |
| chr6 | 160723110 | ATATATGTGTGTA-TATATATG-TACACATATATGT | loxF8L | 3 | loxF8L | 4 | CATATATA/TATATATG | 7 |
| chr4 | 179679125 | ATATATATGTGTA-TATATATG-TACACACATATGT | loxF8L | 3 | loxF8L | 4 | CATATATA/TATATATG | 7 |
| chr9 | 5441605 | ATGAATTTGTGAA-TATATATG-TGTACATTTTTAT | loxF8L | 3 | loxF8L | 4 | CATATATA/TATATATG | 7 |
| chr21 | 37447333 | ATATATAAGTGTA-CATATATA-TACACATTTTTAT | loxF8L | 4 | loxF8L | 3 | CATATATA/TATATATG | 7 |
| chr5 | 116084804 | ATAAATATAAAGA-CATATATA-TGCACAGACTTAA | loxF8L | 4 | loxF8L | 3 | CATATATA/TATATATG | 7 |
| chr3 | 169214905 | ATAAATATATATA-CATATATA-TACACACATATAT | loxF8L | 4 | loxF8L | 3 | CATATATA/TATATATG | 7 |
| chr13 | 57932429 | ATATATATGTGTA-TATATATG-TATATATATTTAT | loxF8L | 3 | loxF8L | 4 | CATATATA/TATATATG | 7 |
| chr17 | 47361540 | ATATATATGTCCA-TATATATG-TGCATATATTTAT | loxF8L | 4 | loxF8L | 3 | CATATATA/TATATATG | 7 |
| chr2 | 170479858 | GTATATGTGTGTA-TATATATG-CACACATATTTAT | loxF8L | 4 | loxF8L | 3 | CATATATA/TATATATG | 7 |
| chr6 | 88672270 | ATAAATGTATACA-CATATATA-TACACATATATAT | loxF8L | 4 | loxF8L | 3 | CATATATA/TATATATG | 7 |
| chr3 | 21653281 | ATATATGTGTGTA-TATATATG-TCTATATATGTAT | loxF8L | 3 | loxF8L | 4 | CATATATA/TATATATG | 7 |
| chr13 | 20046620 | ATATATATGTGTA-TATATATG-TGTACATATGTAT | loxF8L | 3 | loxF8L | 4 | CATATATA/TATATATG | 7 |
| chr2 | 9998573 | ATACATATGTACA-TATATATG-TACACATATATAT | loxF8L | 4 | loxF8L | 3 | CATATATA/TATATATG | 7 |
| chr1 | 163059263 | ATACATATGTACA-CATATATA-TACACATATGTAT | loxF8L | 4 | loxF8L | 3 | CATATATA/TATATATG | 7 |
| chr11 | 30871869 | ATACATATGTATA-CATATATA-TACACATATATAT | loxF8L | 4 | loxF8L | 3 | CATATATA/TATATATG | 7 |
| chr5 | 22154502 | ATATATATGTACA-CATATATA-TACACATATGTAT | loxF8L | 4 | loxF8L | 3 | CATATATA/TATATATG | 7 |
| chr4 | 130702475 | ATATGCGTGTGTG-CATATATA-TACACAAATGTAT | loxF8R | 4 | loxF8L | 3 | CATATATA/TATATATG | 7 |
| chr4 | 34137725 | ATATATTTATGGA-TATATATG-TACACATTTATAT | loxF8L | 3 | loxF8L | 4 | CATATATA/TATATATG | 7 |
| chr10 | 57693050 | ATATATGTGTGTA-TATATATG-TCCATGTATATAT | loxF8L | 3 | loxF8L | 4 | CATATATA/TATATATG | 7 |
| chr14 | 82491298 | ATATATATATAGA-TATATATG-TACACATATATAT | loxF8L | 4 | loxF8L | 3 | CATATATA/TATATATG | 7 |
| chr5 | 17900677 | ATATATATGTGTA-CATATATA-TGCATATATATAT | loxF8L | 3 | loxF8L | 4 | CATATATA/TATATATG | 7 |
| chr5 | 17900651 | ATATATATGTGTA-CATATATA-TGCATATATATAT | loxF8L | 3 | loxF8L | 4 | CATATATA/TATATATG | 7 |
| chr5 | 17900599 | ATATATATGTGTA-CATATATA-TGCATATATATAT | loxF8L | 3 | loxF8L | 4 | CATATATA/TATATATG | 7 |
| chr5 | 17900523 | ATATATATGTGTA-CATATATA-TGCATATATATAT | loxF8L | 3 | loxF8L | 4 | CATATATA/TATATATG | 7 |
| chr5 | 17900549 | ATATATATGTGTA-CATATATA-TGCATATATATAT | loxF8L | 3 | loxF8L | 4 | CATATATA/TATATATG | 7 |
| chr5 | 17900625 | ATATATATGTGTA-CATATATA-TGCATATATATAT | loxF8L | 3 | loxF8L | 4 | CATATATA/TATATATG | 7 |
| chr1 | 198660006 | ATATATTTGTCCA-TATATATG-TCCATATATATAT | loxF8L | 4 | loxF8L | 3 | CATATATA/TATATATG | 7 |
| chr8 | 50734968 | ATATATCTATCCA-TATATATG-TCCATATATATAT | loxF8L | 4 | loxF8L | 3 | CATATATA/TATATATG | 7 |
| chr10 | 90956245 | ATATATATATGTA-CATATATA-TACACATATATAT | loxF8L | 4 | loxF8L | 3 | CATATATA/TATATATG | 7 |
| chr5 | 117993476 | ATATATATGCGTA-TATATATG-TACACATATATAT | loxF8L | 4 | loxF8L | 3 | CATATATA/TATATATG | 7 |
| chrX | 79081320 | ATATATATGTACA-CATATATA-CACACACATTTAG | loxF8L | 4 | loxF8R | 3 | CATATATA/TATATATG | 7 |
| chr3 | 115474956 | ATATATGTGTGTA-TATATATG-TGTACATATATAT | loxF8L | 3 | loxF8L | 4 | CATATATA/TATATATG | 7 |
| chr18 | 65160632 | ATATATATGTACA-TATATATG-TACACATATATAT | loxF8L | 4 | loxF8L | 3 | CATATATA/TATATATG | 7 |
| chr10 | 59407929 | ATATATGTGTGTA-TATATATG-TATACATATATAT | loxF8L | 3 | loxF8L | 4 | CATATATA/TATATATG | 7 |
| chr22 | 30623889 | ATATATATGTATA-CATATATA-TACACACATATAT | loxF8L | 4 | loxF8L | 3 | CATATATA/TATATATG | 7 |
| chr22 | 30623721 | ATATATATGTATA-CATATATA-TACACACATATAT | loxF8L | 4 | loxF8L | 3 | CATATATA/TATATATG | 7 |
| chr14 | 44441723 | CTAAAAGTGTGTA-TATATATG-TATACATATATAT | loxF8R | 3 | loxF8L | 4 | CATATATA/TATATATG | 7 |
| chr5 | 168034056 | ATATATATGTGTA-TATATATG-TATACATATATAT | loxF8L | 3 | loxF8L | 4 | CATATATA/TATATATG | 7 |
| chr2 | 49779975 | ATATATATGTATA-TATATATG-TACATAGATATAT | loxF8L | 4 | loxF8L | 3 | CATATATA/TATATATG | 7 |
| chr18 | 1910086 | ATATATATGTGCA-CATATATA-TATACATATATAT | loxF8L | 3 | loxF8L | 4 | CATATATA/TATATATG | 7 |
| chrX | 137335989 | ATATATATGTGAA-CATATATA-CACACACACATAT | loxF8L | 3 | loxF8R | 4 | CATATATA/TATATATG | 7 |
| chr3 | 167955536 | ATATATATGTGTG-CATATATA-TACACATATATAT | loxF8L | 4 | loxF8L | 3 | CATATATA/TATATATG | 7 |
| chr3 | 167955510 | ATATATATGTGTG-CATATATA-TACACATATATAT | loxF8L | 4 | loxF8L | 3 | CATATATA/TATATATG | 7 |
| chr8 | 3886002 | ATATATATGTGTA-CATATATA-TACATACATATAT | loxF8L | 3 | loxF8L | 4 | CATATATA/TATATATG | 7 |
| chr12 | 69041506 | ATATATGTGTATA-CATATATA-TACACATATATAT | loxF8L | 4 | loxF8L | 3 | CATATATA/TATATATG | 7 |
| chr10 | 15941257 | ATATATATGTGTA-TATATATG-TCTAGACATATAT | loxF8L | 3 | loxF8L | 4 | CATATATA/TATATATG | 7 |
| chrX | 35605871 | ATATATGTATGTA-TATATATG-CACACAATAACAG | loxF8L | 4 | loxF8R | 3 | CATATATA/TATATATG | 7 |
| chr4 | 179679317 | ATATATGTGCGTA-TATATATG-TACACACATATAT | loxF8L | 4 | loxF8L | 3 | CATATATA/TATATATG | 7 |
| chr11 | 29460166 | ATATATGTGTACA-CATATATA-TACACACATATAT | loxF8L | 4 | loxF8L | 3 | CATATATA/TATATATG | 7 |
| chr18 | 60697176 | ATATATGTGTGCA-CATATATA-CACACACACATAT | loxF8L | 3 | loxF8R | 4 | CATATATA/TATATATG | 7 |
| chr11 | 29989560 | ATATATGTGTGTA-CATATATA-TACACACACATAT | loxF8L | 3 | loxF8L | 4 | CATATATA/TATATATG | 7 |
| chr4 | 172177748 | ATATATGTGTGTA-TATATATG-TACACATATATAC | loxF8L | 3 | loxF8L | 4 | CATATATA/TATATATG | 7 |
| chr13 | 68710262 | ATATGTATGTGTA-CATATATA-CACACACTCATAT | loxF8L | 4 | loxF8R | 3 | CATATATA/TATATATG | 7 |
| --- | | | | | | | | 0 |
| chr9 | 70956505 | CTAGGAATCTGTG-GATATATG-TGCATATATTTAT | loxF8R | 3 | loxF8L | 3 | CATATATC/GATATATG | 6 |
| chr10 | 117981902 | ATATATATATGGA-GATATATG-TGGAGAGATATAT | loxF8L | 3 | loxF8L | 4 | CATATATC/GATATATG | 7 |
| --- | | | | | | | | 0 |
| chr7 | 83521097 | TTTAGATTCTGTG-TATAGATG-CACAAAATGTTGG | loxF8R | 3 | loxF8R | 3 | CATCTATA/TATAGATG | 6 |
| chr13 | 87405974 | ATAAGTATTTGTG-CATCTATA-TGTATAGATTTAT | loxF8R | 4 | loxF8L | 3 | CATCTATA/TATAGATG | 7 |
| --- | | | | | | | | 0 |
| chr11 | 101664396 | ACAATTCTCTGGA-TTACCATG-CACAAAATCTTGT | loxF8L | 3 | loxF8R | 3 | CATGGTAA/TTACCATG | 6 |
| chr16 | 13969063 | AAAATTCTCTGGA-TTACCATG-CATAGATTCTTAT | loxF8L | 3 | loxF8R | 4 | CATGGTAA/TTACCATG | 7 |
| chr3 | 96685082 | AGAATTCTCTGGA-TTACCATG-CAGAGACTCTTAT | loxF8L | 3 | loxF8R | 4 | CATGGTAA/TTACCATG | 7 |
| --- | | | | | | | | 0 |
| chr8 | 103789889 | CTACAAGTGTGTG-CTACCATG-TCCAGATATTTTT | loxF8R | 3 | loxF8L | 3 | CATGGTAG/CTACCATG | 6 |
| chr4 | 138972509 | CTACGGTTATGTG-CTACCATG-TCTATATATATAT | loxF8R | 3 | loxF8L | 4 | CATGGTAG/CTACCATG | 7 |
| --- | | | | | | | | 0 |
| chr10 | 2668956 | TTAAATCTATGTA-TATACATG-CACACAGATATAT | loxF8L | 3 | loxF8L | 3 | CATGTATA/TATACATG | 6 |
| chr2 | 27452437 | ATAAATCTCTTTA-TATACATG-TACACACATATAC | loxF8L | 3 | loxF8L | 4 | CATGTATA/TATACATG | 7 |
| chr1 | 15789599 | GTATATGTGTGTA-TATACATG-TCCATATATGTAT | loxF8L | 4 | loxF8L | 3 | CATGTATA/TATACATG | 7 |
| chr7 | 149781803 | TTTAGCATGTGTG-TATACATG-CACACAAACATGG | loxF8R | 4 | loxF8R | 3 | CATGTATA/TATACATG | 7 |
| --- | | | | | | | | 0 |
| chr17 | 46435776 | GTGAGTTTGTGTG-CATGTGAA-TGCACACATGTAT | loxF8R | 3 | loxF8L | 3 | CATGTGAA/TTCACATG | 6 |
| chr17 | 46653348 | GTGAGTTTGTGTG-CATGTGAA-TGCACACATGTAT | loxF8R | 3 | loxF8L | 3 | CATGTGAA/TTCACATG | 6 |
| --- | | | | | | | | 0 |
| chr4 | 150795703 | GTAAATGAGTGGA-CATTGAAA-ATGACAATCTTAG | loxF8L | 3 | loxF8R | 3 | CATTGAAA/TTTCAATG | 6 |
| chr1 | 117358792 | ATATATCTTTTGA-TTTCAATG-ACCATATTTTTAT | loxF8L | 3 | loxF8L | 4 | CATTGAAA/TTTCAATG | 7 |
| --- | | | | | | | | 0 |

| | | | | | | | | |
|---|---|---|---|---|---|---|---|---|
| chr12 | 101724307 | ATTATATTGTGTG-TAGAAATG-GCCACATATCTAT | loxF8R | 3 | loxF8L | 3 | CATTTCTA/TAGAAATG | 6 |
| chr18 | 39169882 | ATTATTTTGTGGA-TAGAAATG-TCTTCATTTTTAT | loxF8L | 3 | loxF8L | 4 | CATTTCTA/TAGAAATG | 7 |
| --- | | | | | | | | 0 |
| chr2 | 172637466 | ATAAATGTTGGGA-CTTTTTGG-TCTACAGAATTCT | loxF8L | 3 | loxF8L | 3 | CCAAAAAG/CTTTTTGG | 6 |
| chr9 | 19005240 | CAAAATCTGTGGC-CTTTTTGG-CAGAAAATCCTAT | loxF8L | 3 | loxF8R | 4 | CCAAAAAG/CTTTTTGG | 7 |
| --- | | | | | | | | 0 |
| chr1 | 21443059 | ATAAATCGATTGA-CTGTATGG-TCATCTGATTTAT | loxF8L | 3 | loxF8L | 3 | CCATACAG/CTGTATGG | 6 |
| chr1 | 146473733 | ATAAATCGGATGA-CTGTATGG-TCATCCGATTTAT | loxF8L | 3 | loxF8L | 3 | CCATACAG/CTGTATGG | 6 |
| chr1 | 16610616 | ATAAATCAGATGA-CCATACAG-TCATTCGATTTAT | loxF8L | 3 | loxF8L | 4 | CCATACAG/CTGTATGG | 7 |
| --- | | | | | | | | 0 |
| chr2 | 1733199 | AGAAATCTGTAAA-CCCACAGG-TCCAAAAATTTAA | loxF8L | 3 | loxF8L | 3 | CCCACAGG/CCTGTGGG | 6 |
| chr3 | 24960473 | GTGAGAGTGTGTG-CCTGTGGG-TACTCAGATCTTT | loxF8R | 3 | loxF8L | 4 | CCCACAGG/CCTGTGGG | 7 |
| --- | | | | | | | | 0 |
| chr20 | 9134557 | TTTAGATAGTGTG-CCCAGAAA-GACACAAAATTAG | loxF8R | 3 | loxF8R | 3 | CCCAGAAA/TTTCTGGG | 6 |
| chr2 | 187650927 | CTTAGATTGTGCC-TTTCTGGG-CAGACAATATTTA | loxF8R | 3 | loxF8R | 4 | CCCAGAAA/TTTCTGGG | 7 |
| --- | | | | | | | | 0 |
| chr7 | 136851310 | CTTAGATTGTATT-CCCAGGAA-AACACACTCTGAG | loxF8R | 3 | loxF8R | 3 | CCCAGGAA/TTCCTGGG | 6 |
| chr2 | 213241216 | CTAACTCTGTAGA-CCCAGGAA-CCCATATTTTTAG | loxF8L | 3 | loxF8R | 4 | CCCAGGAA/TTCCTGGG | 7 |
| --- | | | | | | | | 0 |
| chr13 | 81809098 | ATAAGAGTGTCTG-CCTGGCAA-TCTATAGATTTCT | loxF8R | 3 | loxF8L | 3 | CCTGGCAA/TTGCCAGG | 6 |
| chr3 | 97179653 | AGAATTCTCTGGA-TTGCCAGG-CAGAGACTCTTAT | loxF8L | 3 | loxF8R | 4 | CCTGGCAA/TTGCCAGG | 7 |
| --- | | | | | | | | 0 |
| chr5 | 54429799 | ACAAATATATGGA-CTGAAATA-CAGAGAATCTGAG | loxF8L | 3 | loxF8R | 3 | CTGAAATA/TATTTCAG | 6 |
| chr13 | 23072733 | TGAAATATGTGAA-TATTTCAG-CACACACTCCTAC | loxF8L | 4 | loxF8R | 3 | CTGAAATA/TATTTCAG | 7 |
| --- | | | | | | | | 0 |
| chr10 | 57155726 | CTAATGTTGTATG-TTTTTAAG-CACATAATCTAAA | loxF8R | 3 | loxF8R | 3 | CTTAAAAA/TTTTTAAG | 6 |
| chr13 | 80394601 | ATAAATTTGTGTT-TTTTTAAG-TCTATAAATTTTT | loxF8L | 3 | loxF8L | 4 | CTTAAAAA/TTTTTAAG | 7 |
| chr9 | 101794773 | ATAAATGTGTGTT-TTTTTAAG-TCACCAGGTTTGT | loxF8L | 3 | loxF8L | 4 | CTTAAAAA/TTTTTAAG | 7 |
| chr2 | 135102413 | ATCAATCTGTGTT-TTTTTAAG-TCACCAAATTTGT | loxF8L | 3 | loxF8L | 4 | CTTAAAAA/TTTTTAAG | 7 |
| chr1 | 232654650 | AGAAATCTGATGA-CTTAAAAA-AACACAGTCATAA | loxF8L | 3 | loxF8R | 4 | CTTAAAAA/TTTTTAAG | 7 |
| --- | | | | | | | | 0 |
| chr15 | 54645539 | ATAAGAGTCTGTG-CTTAGTAA-TCCAGAGAATTCT | loxF8R | 3 | loxF8L | 3 | CTTAGTAA/TTACTAAG | 6 |
| chr1 | 71154467 | AGAATTCTCTGGA-TTACTAAG-CAGAGACTCTTAT | loxF8L | 3 | loxF8R | 4 | CTTAGTAA/TTACTAAG | 7 |
| --- | | | | | | | | 0 |
| chr4 | 28632750 | ATAAGTGTTTGGA-GAAAAAAA-TATACAATGTTAG | loxF8L | 3 | loxF8R | 3 | GAAAAAAA/TTTTTTTC | 6 |
| chr6 | 100208206 | ATAGCTGTGTGGA-TTTTTTTC-TTCTCTGATTTTT | loxF8L | 3 | loxF8L | 4 | GAAAAAAA/TTTTTTTC | 7 |
| chr1 | 162961936 | ACAAATCTCTTCA-GAAAAAAA-GCCACAGATGCAT | loxF8L | 4 | loxF8L | 3 | GAAAAAAA/TTTTTTTC | 7 |
| chr12 | 24760998 | ATATATTTGTGGG-GAAAAAAA-TCCACCTAGTTCT | loxF8L | 3 | loxF8L | 4 | GAAAAAAA/TTTTTTTC | 7 |
| chr6 | 91265882 | CTAAGACTATTTT-GAAAAAAA-TGCACAAATGTAT | loxF8R | 4 | loxF8L | 3 | GAAAAAAA/TTTTTTTC | 7 |
| chr2 | 70441273 | ATATCTCTGACGA-TTTTTTTC-CACAGAATGTTAC | loxF8L | 4 | loxF8R | 3 | GAAAAAAA/TTTTTTTC | 7 |
| chr8 | 88535220 | GTAAATCTGTAGG-TTTTTTTC-CATACACTCATAT | loxF8L | 3 | loxF8R | 4 | GAAAAAAA/TTTTTTTC | 7 |
| chr1 | 42989524 | CTAAGTTGGTGGG-TTTTTTTC-TCCCAACATTTAA | loxF8R | 3 | loxF8L | 4 | GAAAAAAA/TTTTTTTC | 7 |
| --- | | | | | | | | 0 |
| chr9 | 104089093 | TTAACTTTGTGTG-TAATTTTC-ACCAAAGGTTTAT | loxF8R | 3 | loxF8L | 3 | GAAAATTA/TAATTTTC | 6 |
| chr8 | 39958901 | ATAAATTTGTTAA-GAAAATTA-TCCAAAGAATGAT | loxF8L | 3 | loxF8L | 3 | GAAAATTA/TAATTTTC | 6 |
| chr2 | 153932153 | TTATGATTGTGTT-TAATTTTC-TCCCCCTTTTTAT | loxF8R | 3 | loxF8L | 4 | GAAAATTA/TAATTTTC | 7 |
| --- | | | | | | | | 0 |
| chr11 | 98778300 | AGCACTCTGTGGA-TTTATTTC-TCCTCCGACTTAT | loxF8L | 3 | loxF8L | 3 | GAAATAAA/TTTATTTC | 6 |
| chr5 | 148765502 | TTGAATCTGTAAA-TTTATTTC-TTCAAAGATTTTT | loxF8L | 4 | loxF8L | 3 | GAAATAAA/TTTATTTC | 7 |
| --- | | | | | | | | 0 |
| chr3 | 108451830 | AAAAATCTGTTAA-TATATTTC-CACACATTTATAG | loxF8L | 3 | loxF8R | 3 | GAAATATA/TATATTTC | 6 |
| chr4 | 135826549 | ATAAATATGTATA-TATATTTC-TACAAATATGTAT | loxF8L | 3 | loxF8L | 4 | GAAATATA/TATATTTC | 7 |
| chr10 | 60662519 | CTAAATATGTGTA-TATATTTC-TAAACACATTCAT | loxF8L | 3 | loxF8L | 4 | GAAATATA/TATATTTC | 7 |
| --- | | | | | | | | 0 |
| chr1 | 244791681 | AAAAATCTGAAGA-GACATTTC-TCCAAAGAAATAT | loxF8L | 3 | loxF8L | 3 | GAAATGTC/GACATTTC | 6 |
| chr1 | 191153623 | ACAAATGTGTTAA-GAAATGTC-TGCACACATTTTT | loxF8L | 4 | loxF8L | 3 | GAAATGTC/GACATTTC | 7 |
| --- | | | | | | | | 0 |
| chr7 | 82818290 | AGAAATCTGCTGA-GAAGCTTA-CAAAAAATCTTTG | loxF8L | 3 | loxF8R | 3 | GAAGCTTA/TAAGCTTC | 6 |
| chr6 | 62806383 | GTAAATCTGTATA-TAAGCTTC-CAAAAAATGTCAG | loxF8L | 3 | loxF8R | 4 | GAAGCTTA/TAAGCTTC | 7 |
| --- | | | | | | | | 0 |
| chr2 | 146001046 | CCAAGATAATGTG-TTACTGTC-TCAATAGATATAT | loxF8R | 3 | loxF8L | 3 | GACAGTAA/TTACTGTC | 6 |
| chr12 | 66689212 | CAAAGCTTGTGTT-GACAGTAA-TCCAAAAATTTAG | loxF8R | 3 | loxF8L | 3 | GACAGTAA/TTACTGTC | 6 |
| --- | | | | | | | | 0 |
| chr8 | 124066153 | GTAGGATTGTGTC-TAATTCTC-CACACATCCTTAA | loxF8R | 3 | loxF8R | 3 | GAGAATTA/TAATTCTC | 6 |
| chr1 | 104605090 | ATATATCTATGTA-TAATTCTC-TCTATATATCTAT | loxF8L | 3 | loxF8L | 4 | GAGAATTA/TAATTCTC | 7 |
| --- | | | | | | | | 0 |
| chr11 | 104833869 | CTATAATTTTGTG-GTCTTATC-CATAAAATCTTTG | loxF8R | 3 | loxF8R | 3 | GATAAGAC/GTCTTATC | 6 |
| chr18 | 67006530 | CTTAGCTTTTGTG-GTCTTATC-CAAAAAATCTTTC | loxF8R | 3 | loxF8R | 4 | GATAAGAC/GTCTTATC | 7 |
| --- | | | | | | | | 0 |
| chrX | 32227151 | AAGAATATGTGGA-GATATATC-TCCACATATTCTT | loxF8L | 3 | loxF8L | 3 | GATATATC/GATATATC | 6 |
| --- | | | | | | | | 0 |
| chr2 | 155792222 | CTTAGACTGGGTG-GCTTAAGC-AACATAGATTTAT | loxF8R | 3 | loxF8L | 3 | GCTTAAGC/GCTTAAGC | 6 |
| chr10 | 82619891 | ATAAATTTGTGCT-GCTTAAGC-CAACCAATCTTGG | loxF8L | 3 | loxF8R | 3 | GCTTAAGC/GCTTAAGC | 6 |
| chr5 | 142554616 | CCCAGATTGAGTG-GCTTAAGC-AACACAAATTTAT | loxF8R | 3 | loxF8L | 3 | GCTTAAGC/GCTTAAGC | 6 |
| chr4 | 30668638 | ATAAATGTTTGTA-GCTTAAGC-CACGCAGTCTAGG | loxF8L | 3 | loxF8R | 4 | GCTTAAGC/GCTTAAGC | 7 |
| chr3 | 172277265 | CTATAATTGGGTG-GCTTAAGC-AACAAACATTTAT | loxF8R | 3 | loxF8L | 4 | GCTTAAGC/GCTTAAGC | 7 |
| chr5 | 55860824 | CTCAGACTGGGTG-GCTTAAGC-AACAGAAATTTAT | loxF8R | 3 | loxF8L | 4 | GCTTAAGC/GCTTAAGC | 7 |
| chr6 | 99423769 | CAAAGACTGGGTG-GCTTAAGC-AACAGAAATTTAT | loxF8R | 3 | loxF8L | 4 | GCTTAAGC/GCTTAAGC | 7 |
| chr5 | 65855061 | ATAAATTTTTGTT-GCTTAAGC-CACCCAATCTGTG | loxF8L | 4 | loxF8R | 3 | GCTTAAGC/GCTTAAGC | 7 |
| chr6 | 12480298 | ATAAATTTTTGTT-GCTTAAGC-CACCCAGTCTGAG | loxF8L | 4 | loxF8R | 3 | GCTTAAGC/GCTTAAGC | 7 |
| --- | | | | | | | | 0 |
| chr14 | 18550724 | TTAAAAATGTGTG-TATGTTCC-TCTACAAGTTTAT | loxF8R | 3 | loxF8L | 3 | GGAACATA/TATGTTCC | 6 |
| chr22 | 15477776 | TTAAAAATGTGTG-TATGTTCC-TCTACAAGTTTAT | loxF8R | 3 | loxF8L | 3 | GGAACATA/TATGTTCC | 6 |
| chr13 | 48256632 | CTAATATTTTTTA-TATGTTCC-CACATTTTCTTAG | loxF8R | 4 | loxF8R | 3 | GGAACATA/TATGTTCC | 7 |
| --- | | | | | | | | 0 |
| chr1 | 114645498 | ATCAATCTCTGGC-GTTTGAAC-CACTCAATCTGCG | loxF8L | 3 | loxF8R | 3 | GTTCAAAC/GTTTGAAC | 6 |
| chr6 | 40319523 | ATAAATCCCTGGT-GTTCAAAC-CACTCAGTCTATG | loxF8L | 3 | loxF8R | 4 | GTTCAAAC/GTTTGAAC | 7 |

| | | | | | | | | |
|---|---|---|---|---|---|---|---|---|
| chr7 | 88486883 | CTAAGTTTCTATT-GTTCAAAC-CACCCAGTCTTTG | loxF8R | 4 | loxF8R | 3 | GTTCAAAC/GTTTGAAC | 7 |
| --- | | | | | | | | 0 |
| chr11 | 21223437 | TTTAGATGGTGTG-TGTTTTTA-CAGGCAATATTAG | loxF8R | 3 | loxF8R | 3 | TAAAAACA/TGTTTTTA | 6 |
| chr7 | 90684959 | TTAAATATGTATA-TGTTTTTA-TGCATAGATTTTT | loxF8L | 4 | loxF8L | 3 | TAAAAACA/TGTTTTTA | 7 |
| --- | | | | | | | | 0 |
| chr15 | 95635686 | CTAAGATTATTAG-TAAAATTA-TCCAAAGTTTTTT | loxF8R | 3 | loxF8L | 3 | TAAAATTA/TAATTTTA | 6 |
| chr11 | 109370372 | AAGAATATCTGGA-TAAAATTA-TACACATATTCAT | loxF8L | 4 | loxF8L | 3 | TAAAATTA/TAATTTTA | 7 |
| chr4 | 61189028 | GTAAGGTTGTTTG-TAATTTTA-CATAGAGTGTTAG | loxF8R | 3 | loxF8R | 4 | TAAAATTA/TAATTTTA | 7 |
| --- | | | | | | | | 0 |
| chr22 | 17112696 | TCAAGAGTGTGTG-TAAATAAA-GCCACACCTTTAT | loxF8R | 3 | loxF8L | 3 | TAAATAAA/TTTATTTA | 6 |
| chr11 | 130378665 | GTTAATCTGTGTA-TTTATTTA-TTTATATATTTAT | loxF8L | 3 | loxF8L | 4 | TAAATAAA/TTTATTTA | 7 |
| --- | | | | | | | | 0 |
| chr22 | 31839985 | ATAAATATATGAA-TCCATTTA-TTCACACATTGAT | loxF8L | 3 | loxF8L | 3 | TAAATGGA/TCCATTTA | 6 |
| chr10 | 9772911 | CTAAAATGGTTTA-TAAATGGA-CACACAATGTAAA | loxF8R | 4 | loxF8R | 3 | TAAATGGA/TCCATTTA | 7 |
| chr11 | 127662808 | TTAAGAACTTGTG-TCCATTTA-TACACAGAATTAG | loxF8R | 4 | loxF8L | 3 | TAAATGGA/TCCATTTA | 7 |
| chr15 | 68098462 | GAAAATCTGTGTA-TAAATGGA-CCCACACAGTTAA | loxF8L | 3 | loxF8L | 4 | TAAATGGA/TCCATTTA | 7 |
| --- | | | | | | | | 0 |
| chr5 | 53007983 | TTATATCTGTGCA-TAAATGTA-TGGACAGAATTAT | loxF8L | 3 | loxF8L | 3 | TAAATGTA/TACATTTA | 6 |
| chr13 | 56476514 | GAAAAATTGTGTT-TACATTTA-AACACTATCTTAT | loxF8R | 4 | loxF8R | 3 | TAAATGTA/TACATTTA | 7 |
| --- | | | | | | | | 0 |
| chr11 | 119235671 | AAAAGTCTGTGTA-TAAGTGAA-TCCACACAGTTAA | loxF8L | 3 | loxF8L | 3 | TAAGTGAA/TTCACTTA | 6 |
| chr5 | 30503242 | GAAAATCTGTGTA-TAAGTGAA-TCCACACAGTTCA | loxF8L | 3 | loxF8L | 4 | TAAGTGAA/TTCACTTA | 7 |
| --- | | | | | | | | 0 |
| chr18 | 13373308 | TTAAGTTTGTGTA-TGACATTA-TACACATATGTAT | loxF8R | 3 | loxF8L | 3 | TAATGTCA/TGACATTA | 6 |
| chr6 | 114230313 | ACAAAGCTATGTA-TGACATTA-CTCAAAATCTCAG | loxF8L | 4 | loxF8R | 3 | TAATGTCA/TGACATTA | 7 |
| chr2 | 198563728 | CTAATTCTGTGCA-TAATGTCA-TTGACAGTTTGAT | loxF8L | 3 | loxF8L | 4 | TAATGTCA/TGACATTA | 7 |
| --- | | | | | | | | 0 |
| chr9 | 104494029 | ATAAATCTATCTA-TACATAGA-TCTATAGATCTAT | loxF8L | 3 | loxF8L | 3 | TACATAGA/TCTATGTA | 6 |
| chr4 | 102410704 | ATATATATGTGTA-TACATAGA-TACACACATATAT | loxF8L | 3 | loxF8L | 3 | TACATAGA/TCTATGTA | 6 |
| chr6 | 82769551 | AAAAGTCTGTTAA-TCTATGTA-TGCTCATATTTAT | loxF8L | 4 | loxF8L | 3 | TACATAGA/TCTATGTA | 7 |
| chr3 | 93888074 | CTGAGTTTGTGCA-TACATAGA-CACAAAATATTTG | loxF8R | 4 | loxF8R | 3 | TACATAGA/TCTATGTA | 7 |
| --- | | | | | | | | 0 |
| chr15 | 55602665 | ATAAGCTTGTGTA-TACATGCA-CAGACAATTTGAG | loxF8R | 3 | loxF8R | 3 | TACATGCA/TGCATGTA | 6 |
| chr1 | 115674945 | CTGTGTGTGTGTG-TGCATGTA-TGCACACATTTTT | loxF8R | 4 | loxF8L | 3 | TACATGCA/TGCATGTA | 7 |
| chr16 | 76142475 | ATATAAGTGTGTA-TACATGCA-TACACATATGTAT | loxF8L | 4 | loxF8L | 3 | TACATGCA/TGCATGTA | 7 |
| chr4 | 19303042 | ATATATATGTGTA-TACATGCA-AACACATATATAT | loxF8L | 3 | loxF8L | 4 | TACATGCA/TGCATGTA | 7 |
| --- | | | | | | | | 0 |
| chr12 | 63556521 | ATACATATGTAGA-TACATGTA-TCTACATATGTAT | loxF8L | 3 | loxF8L | 3 | TACATGTA/TACATGTA | 6 |
| chr4 | 100178707 | ACATATGTGTGTA-TACATGTA-TACACATATATAT | loxF8L | 4 | loxF8L | 3 | TACATGTA/TACATGTA | 7 |
| chr13 | 89949999 | ATAAATATATGTA-TACATGTA-TACACATATCTAC | loxF8L | 3 | loxF8L | 4 | TACATGTA/TACATGTA | 7 |
| chr9 | 76867913 | CTAGGCTCTTGTG-TACATGTA-TCTAAATATTTAT | loxF8R | 4 | loxF8L | 3 | TACATGTA/TACATGTA | 7 |
| chrX | 73878367 | ATATATATGTGTA-TACATGTA-TACATACATATAT | loxF8L | 3 | loxF8L | 4 | TACATGTA/TACATGTA | 7 |
| chr12 | 45715888 | GTATATATGTGTA-TACATGTA-TACACACATATAT | loxF8L | 4 | loxF8L | 3 | TACATGTA/TACATGTA | 7 |
| --- | | | | | | | | 0 |
| chr3 | 118372927 | ATATATCTGTATA-TACTTATA-TACACATATATAT | loxF8L | 3 | loxF8L | 3 | TACTTATA/TATAAGTA | 6 |
| chr1 | 197825414 | ATACATATGTATA-TACTTATA-TACACACATATAT | loxF8L | 4 | loxF8L | 3 | TACTTATA/TATAAGTA | 7 |
| --- | | | | | | | | 0 |
| chr8 | 3354872 | ATATATCTATAGA-TATGTCTA-TCTATAGATCTAT | loxF8L | 3 | loxF8L | 3 | TAGACATA/TATGTCTA | 6 |
| chr9 | 6904251 | AAAAATCTGTATC-TATGTCTA-TCTATATATTTAT | loxF8L | 4 | loxF8L | 3 | TAGACATA/TATGTCTA | 7 |
| chr13 | 86639120 | AAATATCTGTTTA-TATGTCTA-TGCACACATATAT | loxF8L | 4 | loxF8L | 3 | TAGACATA/TATGTCTA | 7 |
| chr8 | 69820787 | ATATATGTGTACA-TAGACATA-TACACACATATAT | loxF8L | 4 | loxF8L | 3 | TAGACATA/TATGTCTA | 7 |
| chr6 | 64633227 | TTGAAATTGTATG-TAGACATA-TACAAAGATTTAG | loxF8R | 4 | loxF8L | 3 | TAGACATA/TATGTCTA | 7 |
| --- | | | | | | | | 0 |
| chr6 | 54988532 | ATCTATCTGTCGA-TCTATCTA-TCCACCTATCTAT | loxF8L | 3 | loxF8L | 3 | TAGATAGA/TCTATCTA | 6 |
| chr12 | 53881157 | AAAAATCTGTCTA-TCTATCTA-TCTACCTATCTAT | loxF8L | 3 | loxF8L | 4 | TAGATAGA/TCTATCTA | 7 |
| chr2 | 140364184 | ACAGATATGTGGA-TAGATAGA-TATACAGATAGAT | loxF8L | 3 | loxF8L | 4 | TAGATAGA/TCTATCTA | 7 |
| chr8 | 3592206 | ATAAACAAGTGGA-TAGATAGA-TACATAGATGGAT | loxF8L | 3 | loxF8L | 4 | TAGATAGA/TCTATCTA | 7 |
| chr2 | 9020896 | ATCTATCTGTTTA-TCTATCTA-TCTACCTATTTAT | loxF8L | 4 | loxF8L | 3 | TAGATAGA/TCTATCTA | 7 |
| chr4 | 104161535 | ATCTATCTGTCTA-TCTATCTA-TCTACCTATTTAT | loxF8L | 4 | loxF8L | 3 | TAGATAGA/TCTATCTA | 7 |
| chr18 | 72820939 | ATAAATCTCTAGG-TAGATAGA-TAGATAGACTTAT | loxF8L | 3 | loxF8L | 4 | TAGATAGA/TCTATCTA | 7 |
| chr1 | 243115279 | ATATATCTGTATA-TCTATCTA-TACACACACGTAT | loxF8L | 3 | loxF8L | 4 | TAGATAGA/TCTATCTA | 7 |
| chr3 | 116019202 | ATAGATCTCTTGA-TAGATAGA-TAGACAGATAGAT | loxF8L | 3 | loxF8L | 4 | TAGATAGA/TCTATCTA | 7 |
| chr18 | 65493654 | ATAGATTTGTAGA-TAGATAGA-TACATAGATAGAT | loxF8L | 3 | loxF8L | 4 | TAGATAGA/TCTATCTA | 7 |
| chr12 | 90908837 | ATATATATGTAGA-TAGATAGA-TAGACACATATAT | loxF8L | 3 | loxF8L | 4 | TAGATAGA/TCTATCTA | 7 |
| chr5 | 102010253 | ATATATGTGTGTA-TAGATAGA-TATACATATATAT | loxF8L | 3 | loxF8L | 4 | TAGATAGA/TCTATCTA | 7 |
| chr4 | 118565245 | ATATGTGTGTGTG-TAGATAGA-TATACAGATATAT | loxF8R | 4 | loxF8L | 3 | TAGATAGA/TCTATCTA | 7 |
| --- | | | | | | | | 0 |
| chr20 | 56280704 | ATATATCTATCGA-TAGATATA-TCCACATAGATAT | loxF8L | 3 | loxF8L | 3 | TAGATATA/TATATCTA | 6 |
| chr18 | 52928578 | ACATATATGTGGA-TAGATATA-CACATAGATATAT | loxF8L | 3 | loxF8L | 4 | TAGATATA/TATATCTA | 7 |
| chr4 | 60595409 | ATAGATATGTGTA-TAGATATA-TCTATAGATATGT | loxF8L | 3 | loxF8L | 4 | TAGATATA/TATATCTA | 7 |
| chr6 | 67020925 | ATAAATATATGCA-TATATCTA-TATATAGATATAT | loxF8L | 3 | loxF8L | 4 | TAGATATA/TATATCTA | 7 |
| chr4 | 60595487 | ATAGATATGTATA-TAGATATA-TCTATAGATGTAT | loxF8L | 4 | loxF8L | 3 | TAGATATA/TATATCTA | 7 |
| chr17 | 71609858 | ATATATCTATGTA-TATATCTA-TATACATATCTAT | loxF8L | 3 | loxF8L | 4 | TAGATATA/TATATCTA | 7 |
| chr9 | 98437663 | ATCCATCTGTGCA-TATATCTA-TTCACATAACTAT | loxF8L | 3 | loxF8L | 4 | TAGATATA/TATATCTA | 7 |
| chr15 | 26606781 | ATATATCTATAGA-TAGATATA-TCTATAGATAGAT | loxF8L | 3 | loxF8L | 4 | TAGATATA/TATATCTA | 7 |
| chr20 | 56280764 | ATATCTATCTGGA-TATATCTA-TCGATAGATATAT | loxF8L | 4 | loxF8L | 3 | TAGATATA/TATATCTA | 7 |
| chr18 | 43631170 | ATATATGTATGGA-TAGATATA-TATAGAGATATAT | loxF8L | 3 | loxF8L | 4 | TAGATATA/TATATCTA | 7 |
| chr2 | 190024337 | ATATGTGTGTGTG-TATATCTA-TACACACATATAT | loxF8R | 4 | loxF8L | 3 | TAGATATA/TATATCTA | 7 |
| --- | | | | | | | | 0 |
| chr8 | 72518183 | ATGAATCTGTCTA-TAGCCTCA-GACATAATCTCAG | loxF8L | 3 | loxF8R | 3 | TAGCCTCA/TGAGGCTA | 6 |
| chr16 | 71555669 | GTTAGAGTGTGGG-TGAGGCTA-TCCACAGAGATGT | loxF8R | 4 | loxF8L | 3 | TAGCCTCA/TGAGGCTA | 7 |
| --- | | | | | | | | 0 |
| chr14 | 91019577 | ATAAATCTTTTTA-TATTTATA-CACACACACATAG | loxF8L | 3 | loxF8R | 3 | TATAAATA/TATTTATA | 6 |
| chr3 | 169499087 | ATACATATGAGTA-TATAAATA-TCCATATGTTTAT | loxF8L | 4 | loxF8L | 3 | TATAAATA/TATTTATA | 7 |
| chr18 | 1916730 | ATAAATATATATA-TATTTATA-TACACATATGTAT | loxF8L | 4 | loxF8L | 3 | TATAAATA/TATTTATA | 7 |
| chr5 | 43579416 | ATATATTTTTGCA-TATTTATA-TACATATATTTAT | loxF8L | 4 | loxF8L | 3 | TATAAATA/TATTTATA | 7 |
| chr10 | 22300402 | ATAAATATGTACA-TATTTATA-TACACACACATAT | loxF8L | 3 | loxF8L | 4 | TATAAATA/TATTTATA | 7 |

| | | | | | | | | |
|---|---|---|---|---|---|---|---|---|
| chr10 | 22300306 | ATAAATATGTACA-TATTTATA-TACACACACATAT | loxF8L | 3 | loxF8L | 4 | TATAAATA/TATTTATA | 7 |
| chr10 | 22300338 | ATAAATATGTACA-TATTTATA-TACACACACATAT | loxF8L | 3 | loxF8L | 4 | TATAAATA/TATTTATA | 7 |
| chr10 | 22300370 | ATAAATATGTACA-TATTTATA-TACACACACATAT | loxF8L | 3 | loxF8L | 4 | TATAAATA/TATTTATA | 7 |
| chr10 | 22300276 | ATAAATATGTACA-TATTTATA-TACACACATATAA | loxF8L | 3 | loxF8L | 4 | TATAAATA/TATTTATA | 7 |
| chr10 | 22300434 | ATAAATATGTACA-TATTTATA-TACACACATATAA | loxF8L | 3 | loxF8L | 4 | TATAAATA/TATTTATA | 7 |
| chr10 | 22300614 | ATAAATATGTACA-TATTTATA-TACACACATATAA | loxF8L | 3 | loxF8L | 4 | TATAAATA/TATTTATA | 7 |
| chr5 | 100493524 | ATAAATTTATATA-TATTTATA-TATACATATTTAT | loxF8L | 4 | loxF8L | 3 | TATAAATA/TATTTATA | 7 |
| chr2 | 229292149 | ATGTATATGTGTA-TATAAATA-CACACATATTTAT | loxF8L | 4 | loxF8L | 3 | TATAAATA/TATTTATA | 7 |
| chr6 | 9664117 | ATAAATGTATATA-TATTTATA-AACACATATTTAT | loxF8L | 4 | loxF8L | 3 | TATAAATA/TATTTATA | 7 |
| chr22 | 34578337 | ATAAATATGTTTA-TATAAATA-TATATAAATTTAT | loxF8L | 3 | loxF8L | 4 | TATAAATA/TATTTATA | 7 |
| chr2 | 66165733 | ATATATGTGTAGA-TATAAATA-AAAAGAGATTTAT | loxF8L | 3 | loxF8L | 4 | TATAAATA/TATTTATA | 7 |
| chr13 | 71656941 | ATATATATGCGCA-TATAAATA-TGCACATATATAT | loxF8L | 4 | loxF8L | 3 | TATAAATA/TATTTATA | 7 |
| chr3 | 104625712 | ATATATATGTGTG-TATTTATA-TACACATATATAT | loxF8L | 4 | loxF8L | 3 | TATAAATA/TATTTATA | 7 |
| chr5 | 28376893 | CTATAAATGTGTG-TATAAATA-TGCATATATTTAC | loxF8R | 3 | loxF8L | 4 | TATAAATA/TATTTATA | 7 |
| --- | | | | | | | | 0 |
| chr4 | 109232787 | CTCAGATTCTGTT-TTATTATA-CCCACATTCTTAA | loxF8R | 3 | loxF8R | 3 | TATAATAA/TTATTATA | 6 |
| chr1 | 221806280 | ATAATACTGTGTG-TATAATAA-AACAAATTCTGAG | loxF8R | 3 | loxF8R | 4 | TATAATAA/TTATTATA | 7 |
| --- | | | | | | | | 0 |
| chr4 | 45026572 | ATATATCTCTGTA-TATATGGA-TCTATAGATCTAT | loxF8L | 3 | loxF8L | 3 | TATATGGA/TCCATATA | 6 |
| chrX | 50304132 | ATAAATGTGTTTA-TATATGGA-CTCTCAATTTTAT | loxF8L | 3 | loxF8R | 4 | TATATGGA/TCCATATA | 7 |
| chr6 | 139795655 | ATATATGTATGGA-TATATGGA-TATACATATATAT | loxF8L | 3 | loxF8L | 4 | TATATGGA/TCCATATA | 7 |
| --- | | | | | | | | 0 |
| chr5 | 29267539 | ATGAATCAATGGA-TTTTGATA-CAGATATTCTTAG | loxF8L | 3 | loxF8R | 3 | TATCAAAA/TTTTGATA | 6 |
| chrX | 121068135 | GGACACCTGTGGA-TATCAAAA-TCCACAGATGCCT | loxF8L | 4 | loxF8L | 3 | TATCAAAA/TTTTGATA | 7 |
| --- | | | | | | | | 0 |
| chr3 | 161207597 | CTAAATCTATGCA-TTGAGATA-TGAAGAGATTTAT | loxF8L | 3 | loxF8L | 3 | TATCTCAA/TTGAGATA | 6 |
| chr5 | 5782890 | TTATATCTGTGTA-TTGAGATA-TACATACATTTAA | loxF8L | 3 | loxF8L | 4 | TATCTCAA/TTGAGATA | 7 |
| --- | | | | | | | | 0 |
| chr3 | 5540518 | ATACATATGTGTA-TATGCATA-TACACACATATAT | loxF8L | 3 | loxF8L | 3 | TATGCATA/TATGCATA | 6 |
| chr3 | 160647772 | AAAAATATGTATA-TATGCATA-TACACATATATAT | loxF8L | 4 | loxF8L | 3 | TATGCATA/TATGCATA | 7 |
| chrX | 134611524 | ATACATATGTATA-TATGCATA-TACACATATGTAT | loxF8L | 4 | loxF8L | 3 | TATGCATA/TATGCATA | 7 |
| chr4 | 58944537 | GAAAGTGTGTGTG-TATGCATA-TACACACATGTAT | loxF8R | 4 | loxF8L | 3 | TATGCATA/TATGCATA | 7 |
| chr7 | 124660183 | ATATATATGTATA-TATGCATA-TACACACATATAT | loxF8L | 4 | loxF8L | 3 | TATGCATA/TATGCATA | 7 |
| chr10 | 96039715 | ATATATGTGTGTA-TATGCATA-TACACAGAGATAC | loxF8L | 3 | loxF8L | 4 | TATGCATA/TATGCATA | 7 |
| --- | | | | | | | | 0 |
| chr6 | 45959339 | ATAAATGTGTCTA-TTAACATA-CACAACATCTTAA | loxF8L | 3 | loxF8R | 3 | TATGTTAA/TTAACATA | 6 |
| chr1 | 162283122 | AAAAATCCATGGA-TATGTTAA-TATACATATATAT | loxF8L | 3 | loxF8L | 4 | TATGTTAA/TTAACATA | 7 |
| --- | | | | | | | | 0 |
| chr5 | 115056499 | CTACTATTATGTG-TCCTTGGA-CACACAATTTAAA | loxF8R | 3 | loxF8R | 3 | TCCAAGGA/TCCTTGGA | 6 |
| chr9 | 10230538 | ATGTATTTGTGGT-TCCTTGGA-CCCTGAGATTTAT | loxF8L | 4 | loxF8L | 3 | TCCAAGGA/TCCTTGGA | 7 |
| --- | | | | | | | | 0 |
| chr20 | 25758335 | ATCAATCTGTCAA-TCCATGCA-TCCATCCATTTAT | loxF8L | 3 | loxF8L | 3 | TCCATGCA/TGCATGGA | 6 |
| chr19 | 42979270 | ACAAATCTGTCCA-TCCATGCA-TCTACGTATGTAT | loxF8L | 3 | loxF8L | 4 | TCCATGCA/TGCATGGA | 7 |
| chr4 | 83901149 | ATATATCTATCCA-TCCATGCA-TCCATCCATTTAT | loxF8L | 4 | loxF8L | 3 | TCCATGCA/TGCATGGA | 7 |
| chr17 | 22708980 | ATTAATCTATCAA-TCCATGCA-TCCATCCATTTAT | loxF8L | 4 | loxF8L | 3 | TCCATGCA/TGCATGGA | 7 |
| chr17 | 22056370 | ATAAATGGATGGA-TGCATGGA-TTGATAGATTGAT | loxF8L | 3 | loxF8L | 4 | TCCATGCA/TGCATGGA | 7 |
| chr17 | 21646935 | ATCAATCTATCAA-TCCATGCA-TCCATCCATTTAT | loxF8L | 4 | loxF8L | 3 | TCCATGCA/TGCATGGA | 7 |
| chr17 | 21849689 | ATCAATCTATCAA-TCCATGCA-TCCATCCATTTAT | loxF8L | 4 | loxF8L | 3 | TCCATGCA/TGCATGGA | 7 |
| chr20 | 26097861 | ATAAATGGATGGA-TGCATGGA-TTGATAGATTGAT | loxF8L | 3 | loxF8L | 4 | TCCATGCA/TGCATGGA | 7 |
| chr21 | 9564741 | ATCAATCTATCAA-TCCATGCA-TCCATCCATTTAT | loxF8L | 4 | loxF8L | 3 | TCCATGCA/TGCATGGA | 7 |
| chr14 | 101625948 | ATAGATGGATGGA-TGCATGGA-TCGACAGATGGAT | loxF8L | 4 | loxF8L | 3 | TCCATGCA/TGCATGGA | 7 |
| chr5 | 2219109 | ATCTATCTGTCCA-TCCATGCA-TCCACATATCCAT | loxF8L | 4 | loxF8L | 3 | TCCATGCA/TGCATGGA | 7 |
| --- | | | | | | | | 0 |
| chr2 | 65694231 | ATGAGATGGTGTG-TGGGGGGA-TAGACAGATTTGT | loxF8R | 3 | loxF8L | 3 | TCCCCCCA/TGGGGGGA | 6 |
| chr8 | 22081084 | CTAAGATTGCCCC-TCCCCCCA-CCCCCAGATCTAT | loxF8R | 4 | loxF8L | 3 | TCCCCCCA/TGGGGGGA | 7 |
| --- | | | | | | | | 0 |
| chrX | 27707401 | ATAAATGGGTAGA-TGGCTAGA-GACACAGATATAT | loxF8L | 3 | loxF8L | 3 | TCTAGCCA/TGGCTAGA | 6 |
| chr7 | 2395603 | ATAGATCGATAGA-TGGCTAGA-TCGACAGGTTGAT | loxF8L | 4 | loxF8L | 3 | TCTAGCCA/TGGCTAGA | 7 |
| --- | | | | | | | | 0 |
| chr2 | 81372676 | ATACATCTATAGA-TCTATAGA-TCTATAGATATAT | loxF8L | 3 | loxF8L | 3 | TCTATAGA/TCTATAGA | 6 |
| chr3 | 42598938 | AGATATCTGTAGA-TCTATAGA-TCTATAGATAGAT | loxF8L | 3 | loxF8L | 4 | TCTATAGA/TCTATAGA | 7 |
| --- | | | | | | | | 0 |
| chr11 | 113314166 | ATAAATGTGTGAG-TCTTCTAA-CTCACACATTTAT | loxF8L | 3 | loxF8L | 3 | TCTTCTAA/TTAGAAGA | 6 |
| chr7 | 93938771 | CTAAGAGTATGAC-TCTTCTAA-CACTAAACCTTAG | loxF8R | 4 | loxF8R | 3 | TCTTCTAA/TTAGAAGA | 7 |
| --- | | | | | | | | 0 |
| chr5 | 40062515 | TTCAATCTGTGGT-TGAGTGAA-TCCACAGATGCAG | loxF8L | 3 | loxF8L | 3 | TGAGTGAA/TTCACTCA | 6 |
| chr15 | 86084797 | ATCCATCTGTATA-TTCACTCA-TCCACCCATTCAT | loxF8L | 4 | loxF8L | 3 | TGAGTGAA/TTCACTCA | 7 |
| --- | | | | | | | | 0 |
| chr15 | 42846495 | TTCAATCTGTGGT-TGGTTGCA-TCCACAGATGTGG | loxF8L | 3 | loxF8L | 3 | TGCAACCA/TGGTTGCA | 6 |
| chr8 | 133439076 | CTGTATCTATGGA-TGCAACCA-ACCACAGATTGAA | loxF8L | 4 | loxF8L | 3 | TGCAACCA/TGGTTGCA | 7 |
| --- | | | | | | | | 0 |
| chr2 | 77213469 | ATAAATATGTATA-TGTATACA-TACACACATATAT | loxF8L | 3 | loxF8L | 3 | TGTATACA/TGTATACA | 6 |
| chr4 | 180358111 | ATATATATGTGTA-TGTATACA-TACACATATATAT | loxF8L | 3 | loxF8L | 3 | TGTATACA/TGTATACA | 6 |
| chr2 | 67813523 | ACATATATGTGTA-TGTATACA-TACACATATATAT | loxF8L | 4 | loxF8L | 3 | TGTATACA/TGTATACA | 7 |
| chr1 | 43952625 | ATATATATGTGTG-TGTATACA-TACATATATTTAT | loxF8L | 4 | loxF8L | 3 | TGTATACA/TGTATACA | 7 |
| chr13 | 67336931 | GTGTGTTTGTGTG-TGTATACA-TATATAGATTTAT | loxF8R | 4 | loxF8L | 3 | TGTATACA/TGTATACA | 7 |
| chr17 | 51010598 | ATACATATATGTA-TGTATACA-CACACACATTTAG | loxF8L | 4 | loxF8R | 3 | TGTATACA/TGTATACA | 7 |
| chr4 | 107262175 | ATACATATGTGCA-TGTATACA-CAAACATTCTTGC | loxF8L | 3 | loxF8R | 4 | TGTATACA/TGTATACA | 7 |
| chr16 | 64806239 | ATACATATGTGTA-TGTATACA-CACACACAATTAA | loxF8L | 3 | loxF8R | 4 | TGTATACA/TGTATACA | 7 |
| chr11 | 57226701 | ATATATACGTGGG-TGTATACA-TACACATATATAT | loxF8L | 4 | loxF8L | 3 | TGTATACA/TGTATACA | 7 |
| chr4 | 142246109 | ATATATATGTGTA-TGTATACA-CACACATATATAT | loxF8L | 3 | loxF8L | 4 | TGTATACA/TGTATACA | 7 |
| chr6 | 7957071 | ATATATCTGTGTG-TGTATACA-CACACACACACAG | loxF8L | 3 | loxF8R | 4 | TGTATACA/TGTATACA | 7 |
| chr10 | 118475743 | ATATATGTGTGTA-TGTATACA-TACACATACATAT | loxF8L | 3 | loxF8L | 4 | TGTATACA/TGTATACA | 7 |
| chr18 | 53537415 | CCAATAGTGTGTG-TGTATACA-CACACACATATAT | loxF8R | 3 | loxF8L | 4 | TGTATACA/TGTATACA | 7 |
| chr10 | 118475713 | ATGTATGTGTGGA-TGTATACA-TACACATACATAT | loxF8L | 3 | loxF8L | 4 | TGTATACA/TGTATACA | 7 |
| chr2 | 30611365 | CAGAGAGTGTGTG-TGTATACA-CTCACACTTTCAG | loxF8R | 3 | loxF8R | 4 | TGTATACA/TGTATACA | 7 |
| chrX | 26077000 | GTGAGAGTGTGTG-TGTATACA-TGCACATATGTAC | loxF8R | 3 | loxF8L | 4 | TGTATACA/TGTATACA | 7 |

| | | | | | | | | |
|---|---|---|---|---|---|---|---|---|
| --- | | | | | | | | 0 |
| chr11 | 75726640 | ATACATCTGAGGT-TGTTTGAA-TCCATAGATGTCT | loxF8L | 3 | loxF8L | 3 | TGTTTGAA/TTCAAACA | 6 |
| chr20 | 29523609 | CCACATCTGTGGA-TTCAAACA-ACCACAGATGGAA | loxF8L | 3 | loxF8L | 4 | TGTTTGAA/TTCAAACA | 7 |
| chr12 | 26693652 | TTTGATCTGTGGA-TGTTTGAA-TACACAGATGCAA | loxF8L | 3 | loxF8L | 4 | TGTTTGAA/TTCAAACA | 7 |
| --- | | | | | | | | 0 |
| chr6 | 75002457 | ATAAATATATGGG-TTAATTAA-GCCTCAGATTTAG | loxF8L | 3 | loxF8L | 3 | TTAATTAA/TTAATTAA | 6 |
| --- | | | | | | | | 0 |
| chr15 | 77305899 | TTTAATCTGTGGT-TTGTTGAA-TCCACAGATATGA | loxF8L | 3 | loxF8L | 3 | TTCAACAA/TTGTTGAA | 6 |
| chr4 | 167891669 | AATTATCTGTGGA-TTGTTGAA-CACATAATACTGG | loxF8L | 3 | loxF8R | 4 | TTCAACAA/TTGTTGAA | 7 |
| chr11 | 6114326 | CTATATATGTGGA-TTCAACAA-ACTATAGATTTAA | loxF8L | 3 | loxF8L | 4 | TTCAACAA/TTGTTGAA | 7 |
| chr14 | 58260369 | TTCTATCTGTGGT-TTGTTGAA-TCCACAGATGCAG | loxF8L | 4 | loxF8L | 3 | TTCAACAA/TTGTTGAA | 7 |
| --- | | | | | | | | 0 |
| chrX | 35122218 | TTTAATCTGTGGT-TTCTTGAA-TCCACAGATATGG | loxF8L | 3 | loxF8L | 3 | TTCAAGAA/TTCTTGAA | 6 |
| chr4 | 126287078 | TTCAATCTGTAGT-TTCTTGAA-TCCACAGATGTGG | loxF8L | 4 | loxF8L | 3 | TTCAAGAA/TTCTTGAA | 7 |
| --- | | | | | | | | 0 |
| chr4 | 80722474 | CTAAGGATGTCTG-TTTTTCAA-AAGACACTCTTAG | loxF8R | 3 | loxF8R | 3 | TTGAAAAA/TTTTTCAA | 6 |
| chr3 | 68233685 | TTGAATCTGTAGA-TTGAAAAA-GCCACACATTCCT | loxF8L | 3 | loxF8L | 4 | TTGAAAAA/TTTTTCAA | 7 |
| chr6 | 20813418 | TTAACATTGTTTT-TTGAAAAA-GCAACAGTTTTAT | loxF8R | 4 | loxF8L | 3 | TTGAAAAA/TTTTTCAA | 7 |
| --- | | | | | | | | 0 |
| chr11 | 18403814 | ATAAATCTTTCAA-TTTGGCAA-CACAGAATATTAA | loxF8L | 3 | loxF8R | 3 | TTGCCAAA/TTTGGCAA | 6 |
| chr11 | 61331634 | CTAATCTCGTGTT-TTTGGCAA-TCAACAGGTTTAA | loxF8R | 4 | loxF8L | 3 | TTGCCAAA/TTTGGCAA | 7 |
| --- | | | | | | | | 0 |
| chr5 | 42052088 | CAACTATTGTGTG-TTTAAAAA-AACACAATTTTTG | loxF8R | 3 | loxF8R | 3 | TTTAAAAA/TTTTTAAA | 6 |
| chr21 | 6517097 | ATGAATCTTTGGC-TTTTTAAA-GCCTCAGATTAAA | loxF8L | 3 | loxF8L | 4 | TTTAAAAA/TTTTTAAA | 7 |
| chr21 | 43125414 | ATGAATCTTTGGC-TTTTTAAA-GCCTCAGATTAAA | loxF8L | 3 | loxF8L | 4 | TTTAAAAA/TTTTTAAA | 7 |
| --- | | | | | | | | 0 |
| chr2 | 86736955 | AGAAATCTGTGGA-ACACCTTT-TAAACAGATTTTG | loxF8L | 1 | loxF8L | 4 | AAAGGTGT/ACACCTTT | 5 |
| chr2 | 87059744 | CAAAATCTGTTTA-AAAGGTGT-TCCACAGATTTCT | loxF8L | 4 | loxF8L | 1 | AAAGGTGT/ACACCTTT | 5 |
| chr5 | 144370662 | ATATATTTAGGGA-AAAGGTGT-TGCAAAGATTTTT | loxF8L | 4 | loxF8L | 3 | AAAGGTGT/ACACCTTT | 7 |
| --- | | | | | | | | 0 |
| chr2 | 32598312 | CAAAGATTGTGTG-AAGCTGGT-CACAATATCAAAG | loxF8R | 1 | loxF8R | 4 | AAGCTGGT/ACCAGCTT | 5 |
| chr21 | 10328007 | ATAAAGCTCAGGA-ACCAGCTT-TCACCATATTTTT | loxF8L | 3 | loxF8L | 4 | AAGCTGGT/ACCAGCTT | 7 |
| --- | | | | | | | | 0 |
| chr15 | 81413818 | CTGATATGGTCTG-AATGTGTC-TCCCCAGATTTAT | loxF8R | 4 | loxF8L | 1 | AATGTGTC/GACACATT | 5 |
| chr8 | 20163347 | ATAAATTTTGGGG-GACACATT-CACACCATATCAG | loxF8L | 4 | loxF8R | 3 | AATGTGTC/GACACATT | 7 |
| chr1 | 106001198 | CTGCTATTGTTTG-AATGTGTC-TCCTTAGAGTTAT | loxF8R | 4 | loxF8L | 3 | AATGTGTC/GACACATT | 7 |
| --- | | | | | | | | 0 |
| chr2 | 2611586 | ATAAATCTGTGCA-TTTATGGT-CAAATAATATTTG | loxF8L | 1 | loxF8R | 4 | ACCATAAA/TTTATGGT | 5 |
| chr7 | 91665600 | ATAAATCCATGCA-TTTATGGT-CAACTAATCTTTG | loxF8L | 3 | loxF8R | 4 | ACCATAAA/TTTATGGT | 7 |
| --- | | | | | | | | 0 |
| chr3 | 136847295 | AAAAATCTGTGGA-CTCTAAGT-CCCAAAATGTAAG | loxF8L | 1 | loxF8R | 4 | ACTTAGAG/CTCTAAGT | 5 |
| chr5 | 147444175 | TTAAGAATGGGGG-CTCTAAGT-CACATAAACTTAA | loxF8R | 4 | loxF8R | 3 | ACTTAGAG/CTCTAAGT | 7 |
| --- | | | | | | | | 0 |
| chrX | 31606879 | ATAAATCTCTGGA-AGAGAGCA-TTCACAAATAAAT | loxF8L | 1 | loxF8L | 4 | AGAGAGCA/TGCTCTCT | 5 |
| chr17 | 12903841 | CAAAGTTTGAGGG-AGAGAGCA-AACAGTATCTTAG | loxF8R | 4 | loxF8R | 3 | AGAGAGCA/TGCTCTCT | 7 |
| --- | | | | | | | | 0 |
| chrX | 126162779 | CTAAGATTTTGTG-ATTAAAAA-GGCACAGATACAT | loxF8R | 1 | loxF8L | 4 | ATTAAAAA/TTTTTAAT | 5 |
| chr3 | 57894258 | ATAAATATATGCA-ATTAAAAA-TTCATAGATAAAT | loxF8L | 3 | loxF8L | 4 | ATTAAAAA/TTTTTAAT | 7 |
| chr9 | 14604621 | ATTAATCTTGGTA-TTTTTAAT-TACACAAATTTAG | loxF8L | 4 | loxF8L | 3 | ATTAAAAA/TTTTTAAT | 7 |
| chr11 | 115391947 | TTAAATCTGTGTT-ATTAAAAA-AACAAAATATCAG | loxF8L | 3 | loxF8R | 4 | ATTAAAAA/TTTTTAAT | 7 |
| --- | | | | | | | | 0 |
| chr16 | 47721528 | AAAGCTCTGTGCA-CAACTCAC-TCCACAGAGTTAT | loxF8L | 4 | loxF8L | 1 | CAACTCAC/GTGAGTTG | 5 |
| chr7 | 95558707 | CTTAGATGGTGGG-CAACTCAC-TACCCACAATTAT | loxF8R | 3 | loxF8L | 4 | CAACTCAC/GTGAGTTG | 7 |
| --- | | | | | | | | 0 |
| chr4 | 142007048 | CTAAGATTGGGTG-GCTGTAGG-TGTGCAGCTTTAT | loxF8R | 1 | loxF8L | 4 | CCTACAGC/GCTGTAGG | 5 |
| chr1 | 226551015 | ATAAAACAGGTGA-GCTGTAGG-TGCACAGATTTCT | loxF8L | 4 | loxF8L | 2 | CCTACAGC/GCTGTAGG | 6 |
| chr3 | 78771187 | CAAAGATTAGATG-GCTGTAGG-TGCACGGCTTTAT | loxF8R | 4 | loxF8L | 3 | CCTACAGC/GCTGTAGG | 7 |
| chr9 | 87689226 | CGCAGATTGGGTG-GCTGTAGG-TGTGCAGCTTTAT | loxF8R | 3 | loxF8L | 4 | CCTACAGC/GCTGTAGG | 7 |
| --- | | | | | | | | 0 |
| chr5 | 97821426 | AGAAATCTGTGGA-TTATGAGG-CAGAGATTCTTTG | loxF8L | 1 | loxF8R | 4 | CCTCATAA/TTATGAGG | 5 |
| chr10 | 36295602 | AGAATTCTCTGGA-TTATGAGG-CAGACACTCTTGT | loxF8L | 3 | loxF8R | 4 | CCTCATAA/TTATGAGG | 7 |
| chr10 | 81386860 | AGAAATATCTGGA-TTATGAGG-CAGAGACTCTTAC | loxF8L | 3 | loxF8R | 4 | CCTCATAA/TTATGAGG | 7 |
| --- | | | | | | | | 0 |
| chr3 | 97041503 | CTATGACTGCCTG-CTCCAAAA-TCCACATATTTAT | loxF8R | 4 | loxF8L | 1 | CTCCAAAA/TTTTGGAG | 5 |
| chr1 | 199418023 | CTAATACTATTTG-TTTTGGAG-CACACACATTTAT | loxF8R | 4 | loxF8L | 3 | CTCCAAAA/TTTTGGAG | 7 |
| --- | | | | | | | | 0 |
| chr3 | 102972306 | ATAAACCTGTGGA-GTATTTTC-TCCAAATCTTAAT | loxF8L | 1 | loxF8L | 4 | GAAAATAC/GTATTTTC | 5 |
| chr18 | 58208797 | ATCAAGCTGTGCT-GAAAATAC-TCCTCAGATATAT | loxF8L | 4 | loxF8L | 2 | GAAAATAC/GTATTTTC | 6 |
| --- | | | | | | | | 0 |
| chr16 | 60210754 | ACAGTTCTGTGGC-TGGAAGTC-TCCACAGATTTGT | loxF8L | 4 | loxF8L | 1 | GACTTCCA/TGGAAGTC | 5 |
| chr18 | 66971051 | CTATGGTTGAGTT-TGGAAGTC-CACACATTCTAGG | loxF8R | 4 | loxF8R | 3 | GACTTCCA/TGGAAGTC | 7 |
| --- | | | | | | | | 0 |
| chrX | 95540146 | ATAAATATGTGGA-GGTTAAAC-AACATACTCTTAA | loxF8L | 1 | loxF8R | 4 | GGTTAAAC/GTTTAACC | 5 |
| chr13 | 85347506 | ATAAATGTCTGTG-GTTTAACC-CACCCAATCTTTG | loxF8L | 4 | loxF8R | 2 | GGTTAAAC/GTTTAACC | 6 |
| chr2 | 169462352 | CATAGACTGGGTG-GGTTAAAC-AGCAAAGATTTAT | loxF8R | 4 | loxF8L | 3 | GGTTAAAC/GTTTAACC | 7 |
| chr7 | 136654558 | ATAAATGTCTGTT-GTTTAACC-CACCCAATCTATG | loxF8L | 4 | loxF8R | 3 | GGTTAAAC/GTTTAACC | 7 |
| --- | | | | | | | | 0 |
| chr1 | 202519927 | CTTAGATTGTGTG-TACACCCA-CACAGAAACTTCA | loxF8R | 1 | loxF8R | 4 | TACACCCA/TGGGTGTA | 5 |
| chrY | 26254857 | TGAAGTTTCTGTG-TGGGTGTA-CACACAATCTAAA | loxF8R | 4 | loxF8R | 2 | TACACCCA/TGGGTGTA | 6 |
| chrY | 23415728 | TTTAGATTGTGTG-TACACCCA-CACAGAAACTTCA | loxF8R | 2 | loxF8R | 4 | TACACCCA/TGGGTGTA | 6 |
| --- | | | | | | | | 0 |
| chr3 | 107741884 | ATTCATGTGTAGA-TCAGTGTA-TCCACAGATATAT | loxF8L | 4 | loxF8L | 1 | TACACTGA/TCAGTGTA | 5 |
| chr1 | 102586546 | GTTAATCTGTGAT-TCAGTGTA-CACCCAATCATTG | loxF8L | 4 | loxF8R | 3 | TACACTGA/TCAGTGTA | 7 |
| --- | | | | | | | | 0 |
| chr12 | 105615891 | ATATATCTGTGGA-TACATTCA-GTTACATTCTTAG | loxF8L | 1 | loxF8R | 4 | TACATTCA/TGAATGTA | 5 |
| chr13 | 26626820 | TTAAATATGTGGC-TACATTCA-ACAACAGATCTTT | loxF8L | 3 | loxF8L | 4 | TACATTCA/TGAATGTA | 7 |
| --- | | | | | | | | 0 |

| | | | | | | | | |
|---|---|---|---|---|---|---|---|---|
| chr18 | 43960729 | AAATATTTTTGGA-GATTTTTT-TCCACTCATTTAT | loxF8L | 4 | loxF8L | 2 | AAAAAATC/GATTTTTT | 6 |
| chrX | 89835494 | ATGAATCTGTAGA-AAAAAATC-TCCATTGATGTAG | loxF8L | 2 | loxF8L | 4 | AAAAAATC/GATTTTTT | 6 |
| chrY | 3490237 | ATGAATCTGTAGA-AAAAAATC-TCCATTGATGTAG | loxF8L | 2 | loxF8L | 4 | AAAAAATC/GATTTTTT | 6 |
| chrX | 32153427 | ATAAATCTTTTTA-AAAAAATC-ACCAAAATCTCAG | loxF8L | 3 | loxF8R | 4 | AAAAAATC/GATTTTTT | 7 |
| chr6 | 102367637 | GTAAATAAGTGGA-GATTTTTT-TCCAGGGAAATAT | loxF8L | 3 | loxF8L | 4 | AAAAAATC/GATTTTTT | 7 |
| --- | | | | | | | | 0 |
| chr7 | 11988712 | ATTAGATTGTATT-AAAAACAA-AACAAAATCTTAG | loxF8R | 4 | loxF8R | 2 | AAAAACAA/TTGTTTTT | 6 |
| chr9 | 104298947 | ATAAATTTGATGT-TTGTTTTT-CCCTCAATTTTAG | loxF8L | 4 | loxF8R | 3 | AAAAACAA/TTGTTTTT | 7 |
| --- | | | | | | | | 0 |
| chr8 | 24460959 | ATAGATCTGTGGC-AAAAATTC-ACTTCAGTTTTAT | loxF8L | 2 | loxF8L | 4 | AAAAATTC/GAATTTTT | 6 |
| chr5 | 65327487 | CAAGGATTTTGTG-GAATTTTT-TCCCCATTTTTAA | loxF8R | 3 | loxF8L | 4 | AAAAATTC/GAATTTTT | 7 |
| --- | | | | | | | | 0 |
| chr1 | 103394379 | ATTAATCTGTTGA-ATGGTTTT-TCCACCTCTCTAT | loxF8L | 2 | loxF8L | 4 | AAAACCAT/ATGGTTTT | 6 |
| chr9 | 10411875 | ATAAACTTGGGGT-AAAACCAT-TTCAAATATTTAT | loxF8L | 4 | loxF8L | 3 | AAAACCAT/ATGGTTTT | 7 |
| --- | | | | | | | | 0 |
| chr13 | 97041389 | AAAAGATTCTTTG-GTTCTTTT-TCCAAAGATTAAT | loxF8R | 4 | loxF8L | 2 | AAAAGAAC/GTTCTTTT | 6 |
| chr8 | 65630335 | ATAAATCAGGGAA-AAAAGAAC-CAAACACTCTGGG | loxF8L | 3 | loxF8R | 4 | AAAAGAAC/GTTCTTTT | 7 |
| --- | | | | | | | | 0 |
| chr5 | 44808441 | ATAAATATATGGA-ATTATTTT-CCCAAAGATGAAT | loxF8L | 2 | loxF8L | 4 | AAAATAAT/ATTATTTT | 6 |
| chr18 | 52413498 | ATAAATATTTGAA-ATTATTTT-CACAAAATGATAA | loxF8L | 3 | loxF8R | 4 | AAAATAAT/ATTATTTT | 7 |
| chr1 | 188380446 | CCAATAGTGTGTT-AAAATAAT-TACAAAATCTTAT | loxF8R | 4 | loxF8R | 3 | AAAATAAT/ATTATTTT | 7 |
| chr11 | 129000227 | ATTAATATTTGAA-AAAATAAT-TCCACCGACTGAT | loxF8L | 4 | loxF8L | 3 | AAAATAAT/ATTATTTT | 7 |
| chr16 | 75904573 | CTAATATTTTATC-AAAATAAT-TCCTAAGATTTAG | loxF8R | 4 | loxF8L | 3 | AAAATAAT/ATTATTTT | 7 |
| chr6 | 145191447 | CTAAGATTATTTA-ATTATTTT-TCCACAAACCTAC | loxF8R | 3 | loxF8L | 4 | AAAATAAT/ATTATTTT | 7 |
| --- | | | | | | | | 0 |
| chr15 | 57058807 | ATGAATGTCTGGG-AAAATACA-TTCATAGATTTAT | loxF8L | 4 | loxF8L | 2 | AAAATACA/TGTATTTT | 6 |
| chr6 | 141532913 | CTAACATTTAGTG-TGTATTTT-TAAACAGAATTCT | loxF8R | 3 | loxF8L | 4 | AAAATACA/TGTATTTT | 7 |
| chr17 | 30509530 | ATAAGTAGTTGGA-AAAATACA-TGCAGATATTTAT | loxF8L | 4 | loxF8L | 3 | AAAATACA/TGTATTTT | 7 |
| --- | | | | | | | | 0 |
| chr13 | 67576288 | ATAAATCTGACGA-CATATTTT-TCCACATAATTTG | loxF8L | 2 | loxF8L | 4 | AAAATATG/CATATTTT | 6 |
| chr8 | 33576554 | CTACGGTTGTTTT-AAAATATG-TCCACACATTCTT | loxF8R | 4 | loxF8L | 3 | AAAATATG/CATATTTT | 7 |
| chr7 | 84376813 | TTCAGATTATTTG-CATATTTT-TAATCAGATTTAT | loxF8R | 4 | loxF8L | 3 | AAAATATG/CATATTTT | 7 |
| --- | | | | | | | | 0 |
| chr21 | 16811924 | ATTTATCTGTTTA-CACATTTT-CACACAATTTTGG | loxF8L | 4 | loxF8R | 2 | AAAATGTG/CACATTTT | 6 |
| chr13 | 82181678 | ATTAATATCTTGA-AAAATGTG-TCCAAATATTTTT | loxF8L | 4 | loxF8L | 3 | AAAATGTG/CACATTTT | 7 |
| chr3 | 26456314 | ATAAAATTGTGAT-CACATTTT-TGCACACATTCAT | loxF8L | 4 | loxF8L | 3 | AAAATGTG/CACATTTT | 7 |
| --- | | | | | | | | 0 |
| chr8 | 75587323 | ATAAATTACTGTA-AAAATGTT-ACCACATATTTAT | loxF8L | 4 | loxF8L | 2 | AAAATGTT/AACATTTT | 6 |
| chr4 | 155148142 | CAAAGCTGCTGTG-AACATTTT-TCCATATATTTTT | loxF8R | 4 | loxF8L | 3 | AAAATGTT/AACATTTT | 7 |
| --- | | | | | | | | 0 |
| chr10 | 33602274 | ACAAATATGTGGA-AAAGAAAC-AACACACTCTTGA | loxF8L | 2 | loxF8R | 4 | AAAGAAAC/GTTTCTTT | 6 |
| chr19 | 10599721 | AAAAAAGTGTGTG-GTTTCTTT-CACACAACATTAT | loxF8R | 4 | loxF8R | 3 | AAAGAAAC/GTTTCTTT | 7 |
| chr11 | 30406180 | CTCAGATTCTGAG-GTTTCTTT-CAAATAATCTTGT | loxF8R | 3 | loxF8R | 4 | AAAGAAAC/GTTTCTTT | 7 |
| --- | | | | | | | | 0 |
| chr12 | 9929529 | ATAATTGTGTGGA-CTTTCTTT-CAAAAAATTTTAA | loxF8L | 2 | loxF8R | 4 | AAAGAAAG/CTTTCTTT | 6 |
| chr6 | 45085149 | ATAAAAATGTAGA-AAAGAAAG-TGCAAACATCTAT | loxF8L | 3 | loxF8L | 4 | AAAGAAAG/CTTTCTTT | 7 |
| chr5 | 26957778 | ATAAAAATGTATG-CTTTCTTT-ACCACAAATTAAT | loxF8R | 4 | loxF8L | 3 | AAAGAAAG/CTTTCTTT | 7 |
| chr6 | 121973785 | TTAAAATTCTGTC-CTTTCTTT-TTCACATATTAAT | loxF8R | 4 | loxF8L | 3 | AAAGAAAG/CTTTCTTT | 7 |
| --- | | | | | | | | 0 |
| chr14 | 39290671 | GTAAATCTTTGGA-TAATCTTT-AGTACAATCTTAC | loxF8L | 2 | loxF8R | 4 | AAAGATTA/TAATCTTT | 6 |
| chr5 | 29514342 | AGTAATGTGTGCA-TAATCTTT-TCCACCTATTTTT | loxF8L | 4 | loxF8L | 3 | AAAGATTA/TAATCTTT | 7 |
| --- | | | | | | | | 0 |
| chrX | 95964809 | ATAAGAGCTTGTG-TCTGCTTT-TACACAATTTTAG | loxF8R | 4 | loxF8R | 2 | AAAGCAGA/TCTGCTTT | 6 |
| chrY | 21684112 | CTAAAATTGTGGA-AAAGCAGA-CACAGGCTCTTAT | loxF8L | 3 | loxF8R | 4 | AAAGCAGA/TCTGCTTT | 7 |
| chr3 | 190249999 | ATAAGAGCCTGTG-TCTGCTTT-TCCACAGTTTCAG | loxF8R | 4 | loxF8L | 3 | AAAGCAGA/TCTGCTTT | 7 |
| chr3 | 193923144 | CTGAAACTGTGGA-AAAGCAGA-CACAAGCTCTTAC | loxF8L | 3 | loxF8R | 4 | AAAGCAGA/TCTGCTTT | 7 |
| --- | | | | | | | | 0 |
| chr16 | 72700406 | CAAAGCCTATGTG-GTTTATTT-GACACAGATTTAT | loxF8R | 4 | loxF8L | 2 | AAATAAAC/GTTTATTT | 6 |
| chr6 | 115135614 | TTATTTCTTTGGA-AAATAAAC-TGCACACATATAT | loxF8L | 4 | loxF8L | 3 | AAATAAAC/GTTTATTT | 7 |
| --- | | | | | | | | 0 |
| chr6 | 51744614 | CTAATGTTGTTTA-AAATAAAT-TCAACAGATTTTT | loxF8R | 4 | loxF8L | 2 | AAATAAAT/ATTTATTT | 6 |
| chr5 | 167965607 | AGAAATTTGTTAA-ATTTATTT-CACACAATCATAT | loxF8L | 4 | loxF8R | 2 | AAATAAAT/ATTTATTT | 6 |
| chr4 | 110256504 | CTGAGATTGAGTA-ATTTATTT-TAAAAAGGTTTAT | loxF8R | 3 | loxF8L | 4 | AAATAAAT/ATTTATTT | 7 |
| chr11 | 104984386 | CTAGAATTGTGAG-AAATAAAT-TTCAATCATTTAT | loxF8R | 3 | loxF8L | 4 | AAATAAAT/ATTTATTT | 7 |
| chr1 | 89279530 | CTAAAACTGTGAA-AAATAAAT-TTCTCTGATGTAT | loxF8L | 3 | loxF8L | 4 | AAATAAAT/ATTTATTT | 7 |
| chr12 | 110657563 | CTGTGAATGTGTT-AAATAAAT-GCCACAGAATCAT | loxF8R | 4 | loxF8L | 3 | AAATAAAT/ATTTATTT | 7 |
| --- | | | | | | | | 0 |
| chr2 | 30285654 | ATAAATTTGTTTT-TAATATTT-ACCACATATTTAT | loxF8L | 4 | loxF8L | 2 | AAATATTA/TAATATTT | 6 |
| chr14 | 81975512 | ATTTATCTGTGGC-TAATATTT-TACCCAAGATTAG | loxF8L | 3 | loxF8R | 4 | AAATATTA/TAATATTT | 7 |
| --- | | | | | | | | 0 |
| chr14 | 75495371 | ATAAAACTGTTGA-TTTCATTT-AACAGAATTTTAA | loxF8L | 2 | loxF8R | 4 | AAATGAAA/TTTCATTT | 6 |
| chrX | 50654376 | ATCAAGCTGTTCA-TTTCATTT-TGCATAGCTTTAT | loxF8L | 4 | loxF8L | 3 | AAATGAAA/TTTCATTT | 7 |
| chr6 | 90597303 | ATAAATAAGGGTA-TTTCATTT-CACACAGCCTCAG | loxF8L | 4 | loxF8R | 3 | AAATGAAA/TTTCATTT | 7 |
| --- | | | | | | | | 0 |
| chr12 | 81522637 | CTAATATTTTGTG-CTTCATTT-CACCAAATTTTAA | loxF8R | 2 | loxF8R | 4 | AAATGAAG/CTTCATTT | 6 |
| chr18 | 30207371 | ATAAAGTTTTGGA-CTTCATTT-TCCATTAATTTTT | loxF8L | 3 | loxF8L | 4 | AAATGAAG/CTTCATTT | 7 |
| --- | | | | | | | | 0 |
| chr9 | 95455533 | ATAATTCTTTGAT-AAATGTAT-ACAACAGATTTAT | loxF8L | 4 | loxF8L | 2 | AAATGTAT/ATACATTT | 6 |
| chr3 | 26180229 | GAAAGTCTGTGTA-AAATGTAT-TCCACATTTTTCT | loxF8L | 4 | loxF8L | 3 | AAATGTAT/ATACATTT | 7 |
| --- | | | | | | | | 0 |
| chr22 | 42841123 | CTAAGTTTGTGTC-AACAAGGA-CCCACACTTTTGG | loxF8R | 2 | loxF8R | 4 | AACAAGGA/TCCTTGTT | 6 |
| chr2 | 200466227 | ATAAAGCAGTTGA-TCCTTGTT-CAGCCAAGCATAG | loxF8L | 3 | loxF8R | 4 | AACAAGGA/TCCTTGTT | 7 |
| --- | | | | | | | | 0 |
| chr11 | 55694939 | ATTAATCTGTGGT-AACAGAAG-TCAACATAGTAAT | loxF8L | 2 | loxF8L | 4 | AACAGAAG/CTTCTGTT | 6 |
| chr11 | 8569298 | ACAAATCAATGAA-AACAGAAG-TCAACTGATTTTT | loxF8L | 4 | loxF8L | 3 | AACAGAAG/CTTCTGTT | 7 |
| --- | | | | | | | | 0 |
| chr3 | 139899257 | AGAAATCTTTGGA-AACAGTTG-TTCCCAATCTTAA | loxF8L | 2 | loxF8R | 4 | AACAGTTG/CAACTGTT | 6 |

| | | | | | | | | |
|---|---|---|---|---|---|---|---|---|
| chr3 | 18614441 | ATAATGCTTTGCA-CAACTGTT-CAAAAAATCTTAG | loxF8L | 4 | loxF8R | 2 | AACAGTTG/CAACTGTT | 6 |
| --- | | | | | | | | 0 |
| chr3 | 107461243 | ATTAACCTCTGTA-TCTATGTT-TCCTCAAATTTAT | loxF8L | 4 | loxF8L | 2 | AACATAGA/TCTATGTT | 6 |
| chr10 | 54018541 | CAAAGATTAAGTG-AACATAGA-TGCCTAGAGTTAT | loxF8R | 3 | loxF8L | 4 | AACATAGA/TCTATGTT | 7 |
| --- | | | | | | | | 0 |
| chr16 | 60119586 | ATTATTCTGATGA-AACATTAT-CACACATTCTTAT | loxF8L | 4 | loxF8R | 2 | AACATTAT/ATAATGTT | 6 |
| chr9 | 16956046 | CTGAAATTTTGAG-ATAATGTT-TCCGGAGATGTAT | loxF8R | 4 | loxF8L | 3 | AACATTAT/ATAATGTT | 7 |
| chr8 | 23637456 | CTACCATTATGAG-ATAATGTT-CACACTATATCAG | loxF8R | 4 | loxF8R | 3 | AACATTAT/ATAATGTT | 7 |
| --- | | | | | | | | 0 |
| chr9 | 91505867 | AAAAATCTTAGCA-AACCAAAA-TGCACACATTTAT | loxF8L | 4 | loxF8L | 2 | AACCAAAA/TTTTGGTT | 6 |
| chr2 | 33687676 | TCAAGAGTGTGTT-TTTTGGTT-TCCAAATATTTTT | loxF8R | 4 | loxF8L | 3 | AACCAAAA/TTTTGGTT | 7 |
| chr4 | 9940583 | ATAAATCTCTATA-AACCAAAA-CCCACAGCTTTGG | loxF8L | 3 | loxF8L | 4 | AACCAAAA/TTTTGGTT | 7 |
| chr5 | 150447211 | TTACATATGAGGA-AACCAAAA-CTCAGAATCTGAG | loxF8L | 4 | loxF8R | 3 | AACCAAAA/TTTTGGTT | 7 |
| --- | | | | | | | | 0 |
| chr1 | 76849237 | TCAAGAGTGTGTT-ATTTAGTT-GCCACATATTTAT | loxF8R | 4 | loxF8L | 2 | AACTAAAT/ATTTAGTT | 6 |
| chr7 | 84792006 | AAAAATTTATAGA-AACTAAAT-CACATCATCTTGG | loxF8L | 4 | loxF8R | 3 | AACTAAAT/ATTTAGTT | 7 |
| chrX | 69335777 | TTAGGAATGTGTT-ATTTAGTT-TCCATAGATTAGT | loxF8R | 4 | loxF8L | 3 | AACTAAAT/ATTTAGTT | 7 |
| chr4 | 2347629 | ATAAATATTTGAA-AACTAAAT-AACACACTTTTAA | loxF8L | 3 | loxF8R | 4 | AACTAAAT/ATTTAGTT | 7 |
| --- | | | | | | | | 0 |
| chr3 | 78688054 | ATAAATCTCTGTA-TTTCAGTT-TCCATATATATAA | loxF8L | 2 | loxF8L | 4 | AACTGAAA/TTTCAGTT | 6 |
| chr4 | 94826771 | ATTAAATTGTTGA-AACTGAAA-TCAATAGTTTTAT | loxF8L | 4 | loxF8L | 3 | AACTGAAA/TTTCAGTT | 7 |
| --- | | | | | | | | 0 |
| chr12 | 111177839 | ATACCTCTGTGGA-CCTCAGTT-TCCTCATCTCTAT | loxF8L | 2 | loxF8L | 4 | AACTGAGG/CCTCAGTT | 6 |
| chr7 | 105100134 | TTACCTCTTTGGA-CCTCAGTT-TCCTCAGAGTCAT | loxF8L | 4 | loxF8L | 3 | AACTGAGG/CCTCAGTT | 7 |
| chr1 | 32891633 | TTTAATCTGAGGA-AACTGAGG-CTCACAGATACAT | loxF8L | 3 | loxF8L | 4 | AACTGAGG/CCTCAGTT | 7 |
| chr14 | 94037764 | TTACAGCTGAGGA-AACTGAGG-CACACAAATTAAG | loxF8L | 4 | loxF8R | 3 | AACTGAGG/CCTCAGTT | 7 |
| --- | | | | | | | | 0 |
| chr16 | 49595861 | AAACATCTGTGGA-AACTGCTT-TCCATGTAATTAT | loxF8L | 2 | loxF8L | 4 | AACTGCTT/AAGCAGTT | 6 |
| chr1 | 173753779 | CTAAAACTAGGTG-AAGCAGTT-TCCAGAGGTTTTT | loxF8R | 4 | loxF8L | 3 | AACTGCTT/AAGCAGTT | 7 |
| --- | | | | | | | | 0 |
| chr3 | 10422817 | ACACACGTGTGGA-CACAAGTT-TCCACTGATTTAA | loxF8L | 4 | loxF8L | 2 | AACTTGTG/CACAAGTT | 6 |
| chr13 | 90352267 | ATAAATGTCTGGT-CACAAGTT-CTCAACCTCTTAG | loxF8L | 3 | loxF8R | 4 | AACTTGTG/CACAAGTT | 7 |
| --- | | | | | | | | 0 |
| chr1 | 95573890 | CAAATAATGTCTG-AACTTTCA-CACAGAAACTTAG | loxF8R | 4 | loxF8R | 2 | AACTTTCA/TGAAAGTT | 6 |
| chr9 | 39165617 | CTATGATATTGTT-TGAAAGTT-TCCACTTATTTAC | loxF8R | 4 | loxF8L | 3 | AACTTTCA/TGAAAGTT | 7 |
| chr9 | 41985852 | CTATGATATTGTT-TGAAAGTT-TCCACTTATTTAC | loxF8R | 4 | loxF8L | 3 | AACTTTCA/TGAAAGTT | 7 |
| chr9 | 61453311 | GTAAATAAGTGGA-AACTTTCA-AACAATATCATAG | loxF8L | 3 | loxF8R | 4 | AACTTTCA/TGAAAGTT | 7 |
| chr9 | 67201500 | GTAAATAAGTGGA-AACTTTCA-AACAATATCATAG | loxF8L | 3 | loxF8R | 4 | AACTTTCA/TGAAAGTT | 7 |
| --- | | | | | | | | 0 |
| chr4 | 43544445 | ATAAAGCTGTATT-TTTTTCTT-TTCACATATTTAT | loxF8L | 4 | loxF8L | 2 | AAGAAAAA/TTTTTCTT | 6 |
| chrX | 109425820 | TTAAGTCTATGAA-AAGAAAAA-AAGACAATCTTAG | loxF8L | 4 | loxF8R | 2 | AAGAAAAA/TTTTTCTT | 6 |
| chr2 | 224965855 | ATAAAACTCTGTA-TTTTTCTT-CTCACAGTATTAT | loxF8L | 3 | loxF8L | 4 | AAGAAAAA/TTTTTCTT | 7 |
| --- | | | | | | | | 0 |
| chr4 | 187352831 | AAAAATCTGTGTA-CTGTTCTT-ATCATACATTTAT | loxF8L | 2 | loxF8L | 4 | AAGAACAG/CTGTTCTT | 6 |
| chr19 | 41796098 | ATAATGCTGCAGA-AAGAACAG-CTCAGAATCTTAG | loxF8L | 4 | loxF8R | 2 | AAGAACAG/CTGTTCTT | 6 |
| --- | | | | | | | | 0 |
| chrX | 9057747 | GTAAAGCTGGGGG-AAGAAGAA-TCCCCAGATTTAA | loxF8L | 4 | loxF8L | 2 | AAGAAGAA/TTCTTCTT | 6 |
| chr12 | 59829746 | AAAAAGCTGTGGC-TTCTTCTT-CACACAACAGAAG | loxF8L | 3 | loxF8R | 4 | AAGAAGAA/TTCTTCTT | 7 |
| --- | | | | | | | | 0 |
| chrX | 111581744 | TTCAATCTTTTGA-AAGAGTTA-TCCACATATTTAA | loxF8L | 4 | loxF8L | 2 | AAGAGTTA/TAACTCTT | 6 |
| chr20 | 20341876 | ATTGATCTCAGGA-AAGAGTTA-CGCCAAATCTTAG | loxF8L | 4 | loxF8R | 3 | AAGAGTTA/TAACTCTT | 7 |
| --- | | | | | | | | 0 |
| chr18 | 44504503 | ATAAATTTCTGGA-AAGATTTA-TACAAAATATAAG | loxF8L | 2 | loxF8R | 4 | AAGATTTA/TAAATCTT | 6 |
| chr5 | 71775364 | ATAAAATTGTGGT-AAGATTTA-TCCAGACATATTT | loxF8L | 3 | loxF8L | 4 | AAGATTTA/TAAATCTT | 7 |
| --- | | | | | | | | 0 |
| chr3 | 16268190 | ATAAAGCTTTGGA-AAGCACTT-TAAACAGAATTTT | loxF8L | 2 | loxF8L | 4 | AAGCACTT/AAGTGCTT | 6 |
| chr3 | 157806325 | CTAGCATTGTGCT-AAGCACTT-TCCATACATTAAT | loxF8R | 4 | loxF8L | 3 | AAGCACTT/AAGTGCTT | 7 |
| --- | | | | | | | | 0 |
| chrX | 121575550 | CTAAAATTGAGTG-AATAAAAG-GTCAAAATCTAAG | loxF8R | 2 | loxF8R | 4 | AATAAAAG/CTTTTATT | 6 |
| chr2 | 64586534 | TAAACTTTGTGTG-CTTTTATT-CCCCCAGATTTTT | loxF8R | 4 | loxF8L | 3 | AATAAAAG/CTTTTATT | 7 |
| chr2 | 223513424 | TTAAAGATGAGGA-AATAAAAG-TCCACAGAAGTAG | loxF8L | 4 | loxF8L | 3 | AATAAAAG/CTTTTATT | 7 |
| chr11 | 85744306 | TTAAAACTGAGGA-AATAAAAG-TCCAAGGAGTTAA | loxF8L | 3 | loxF8L | 4 | AATAAAAG/CTTTTATT | 7 |
| --- | | | | | | | | 0 |
| chr13 | 27893204 | TTAAGAGTATGTT-GCTTTATT-TCCACAGGTTTGT | loxF8R | 4 | loxF8L | 2 | AATAAAGC/GCTTTATT | 6 |
| chr1 | 228230898 | TTAAAAGTGTGTT-GCTTTATT-TCCACATATTTGT | loxF8R | 4 | loxF8L | 2 | AATAAAGC/GCTTTATT | 6 |
| chr7 | 40936865 | TTAAGGATGTGTT-GCTTTATT-TCCACATATATAA | loxF8R | 4 | loxF8L | 3 | AATAAAGC/GCTTTATT | 7 |
| --- | | | | | | | | 0 |
| chr11 | 113055495 | TTAAGATTGTATG-GTAATATT-TGCACAGAAACAT | loxF8R | 2 | loxF8L | 4 | AATATTAC/GTAATATT | 6 |
| chr8 | 120919616 | ATAAATGTGTATA-GTAATATT-TACATAAATATAT | loxF8L | 3 | loxF8L | 4 | AATATTAC/GTAATATT | 7 |
| --- | | | | | | | | 0 |
| chr6 | 92796109 | TTAAGAATGTATT-AATTGATT-TCCACAAATTTGT | loxF8R | 4 | loxF8L | 2 | AATCAATT/AATTGATT | 6 |
| chr9 | 9169059 | TTTTATCAGTGGA-AATCAATT-TGCATAAATTTAT | loxF8L | 4 | loxF8L | 3 | AATCAATT/AATTGATT | 7 |
| --- | | | | | | | | 0 |
| chr4 | 21020762 | CAAAGATTGAGTG-AATGAATA-GAGACAATTTTAA | loxF8R | 2 | loxF8R | 4 | AATGAATA/TATTCATT | 6 |
| chr13 | 50601386 | AGAACATTGTGTT-AATGAATA-TCCATACATCTAT | loxF8R | 4 | loxF8L | 3 | AATGAATA/TATTCATT | 7 |
| chr5 | 103374498 | GTAAACCTATGGA-AATGAATA-CTCACAATCTGCA | loxF8L | 3 | loxF8R | 4 | AATGAATA/TATTCATT | 7 |
| --- | | | | | | | | 0 |
| chr3 | 48814828 | ATAAATATTTAGT-TATTAATT-TCAACAGATTTTT | loxF8L | 4 | loxF8L | 2 | AATTAATA/TATTAATT | 6 |
| chr4 | 71587673 | TTAAGAGTGTATT-TATTAATT-TCCATATATTTGT | loxF8R | 4 | loxF8L | 3 | AATTAATA/TATTAATT | 7 |
| chr4 | 86213058 | AGAAATTTGTAGA-AATTAATA-AAAACACTTTTAG | loxF8L | 3 | loxF8R | 4 | AATTAATA/TATTAATT | 7 |
| --- | | | | | | | | 0 |
| chr2 | 107233402 | ACAAATATGTGGA-AATTAGAA-TACTCATTCTTAA | loxF8L | 2 | loxF8R | 4 | AATTAGAA/TTCTAATT | 6 |
| chr14 | 41626763 | ATACAACTGTGTA-TTCTAATT-TACTCTGATGTAT | loxF8L | 3 | loxF8L | 4 | AATTAGAA/TTCTAATT | 7 |
| --- | | | | | | | | 0 |
| chr11 | 77534448 | ATAAATATGTGAA-AATTAGAC-AACACACTCTTGA | loxF8L | 2 | loxF8R | 4 | AATTAGAC/GTCTAATT | 6 |
| chr16 | 21288856 | TTAAGTTTGTATT-GTCTAATT-TACACATATTTGT | loxF8R | 4 | loxF8L | 3 | AATTAGAC/GTCTAATT | 7 |
| chr3 | 112591645 | TTAAGAGTATGTT-GTCTAATT-TCCTCAAATTTGT | loxF8R | 4 | loxF8L | 3 | AATTAGAC/GTCTAATT | 7 |

| | | | | | | | | |
|---|---|---|---|---|---|---|---|---|
| --- | | | | | | | | 0 |
| chr4 | 188744406 | ATAAGATTCTGTG-CTCTAATT-TTCACAAATTTCA | loxF8R | 2 | loxF8L | 4 | AATTAGAG/CTCTAATT | 6 |
| chr22 | 22085749 | CCAAGTCTGTGGA-AATTAGAG-AACAAAATTTTAC | loxF8L | 3 | loxF8R | 4 | AATTAGAG/CTCTAATT | 7 |
| --- | | | | | | | | 0 |
| chr1 | 92219854 | ATAAATTTTTTGG-AATTATTC-TGCACAGATTTGT | loxF8L | 4 | loxF8L | 2 | AATTATTC/GAATAATT | 6 |
| chr4 | 162080959 | ATTATTCTATGCA-AATTATTC-TCTAAGGATTTAT | loxF8L | 4 | loxF8L | 3 | AATTATTC/GAATAATT | 7 |
| --- | | | | | | | | 0 |
| chr17 | 72595367 | AGAAATCTGTGCA-GAAGAATT-TCAAAAAATGTAT | loxF8L | 2 | loxF8L | 4 | AATTCTTC/GAAGAATT | 6 |
| chr6 | 58449624 | ATAAACCTGCAGA-GAAGAATT-CACAGTAACTTAC | loxF8L | 3 | loxF8R | 4 | AATTCTTC/GAAGAATT | 7 |
| --- | | | | | | | | 0 |
| chr3 | 89266495 | ATAAATCAGTGAA-CTTCAATT-ACTACAGATTTTG | loxF8L | 2 | loxF8L | 4 | AATTGAAG/CTTCAATT | 6 |
| chr1 | 236514501 | TTAAGACTGTAGG-AATTGAAG-ACCACAGAATTAT | loxF8R | 4 | loxF8L | 2 | AATTGAAG/CTTCAATT | 6 |
| chr9 | 2974658 | ACAAATTTGAGCA-AATTGAAG-AACACCATCTTAA | loxF8L | 4 | loxF8R | 3 | AATTGAAG/CTTCAATT | 7 |
| --- | | | | | | | | 0 |
| chrX | 5453963 | TTTCATCTCTGGA-AATTGTGT-TCAACAGAATTAT | loxF8L | 4 | loxF8L | 2 | AATTGTGT/ACACAATT | 6 |
| chr13 | 72570705 | CTGACTTTGTGGG-ACACAATT-TCAACAGTTTTCT | loxF8R | 4 | loxF8L | 3 | AATTGTGT/ACACAATT | 7 |
| --- | | | | | | | | 0 |
| chr18 | 51688773 | ATAAATCACTTGG-AATTGTTC-TTCACAGATTTGT | loxF8L | 4 | loxF8L | 2 | AATTGTTC/GAACAATT | 6 |
| chr11 | 40161817 | ATAAATCTGTGCT-GAACAATT-TACAGAATCTAGG | loxF8L | 2 | loxF8R | 4 | AATTGTTC/GAACAATT | 6 |
| --- | | | | | | | | 0 |
| chr12 | 2707744 | ACAAATTTGTGGA-AATTTAAC-AGCACACTCTTAA | loxF8L | 2 | loxF8R | 4 | AATTTAAC/GTTAAATT | 6 |
| chrY | 12705223 | CTCAGAATGTTTT-GTTAAATT-TTCACACATTTGT | loxF8R | 4 | loxF8L | 3 | AATTTAAC/GTTAAATT | 7 |
| chr2 | 108095631 | ACACATATGTGGA-AATTTAAC-AACACTCTCTTAA | loxF8L | 3 | loxF8R | 4 | AATTTAAC/GTTAAATT | 7 |
| --- | | | | | | | | 0 |
| chr6 | 147130605 | AAAAATCTGTTGA-ATTAAATT-CACACAAACACAC | loxF8L | 2 | loxF8R | 4 | AATTTAAT/ATTAAATT | 6 |
| chr13 | 90956987 | ATAAATTTGTATA-AATTTAAT-CACACCAGCTAAA | loxF8L | 3 | loxF8R | 4 | AATTTAAT/ATTAAATT | 7 |
| --- | | | | | | | | 0 |
| chrY | 13410403 | AAAAATTTGAGAA-ATCAAATT-TCCACAGAATTAA | loxF8L | 4 | loxF8L | 2 | AATTTGAT/ATCAAATT | 6 |
| chr6 | 128752872 | ATAAGACTGCGTA-AATTTGAT-TCCATACCTTTAT | loxF8L | 4 | loxF8L | 3 | AATTTGAT/ATCAAATT | 7 |
| chr8 | 115559699 | ATTAAAATTTGGA-ATCAAATT-TCCAAAGATTTTA | loxF8L | 4 | loxF8L | 3 | AATTTGAT/ATCAAATT | 7 |
| --- | | | | | | | | 0 |
| chr2 | 31544414 | ATAAATATGTGAA-AATTTTAA-AACACATTCTTTA | loxF8L | 2 | loxF8R | 4 | AATTTTAA/TTAAAATT | 6 |
| chr8 | 511791 | TCAAATATGTGGA-AATTTTAA-AACTCACTCTTAA | loxF8L | 3 | loxF8R | 4 | AATTTTAA/TTAAAATT | 7 |
| --- | | | | | | | | 0 |
| chr1 | 89888921 | ACAAATCTGTGCA-GAAAAATT-CCAAATGATTTAT | loxF8L | 2 | loxF8L | 4 | AATTTTTC/GAAAAATT | 6 |
| chr11 | 99779187 | CTAAATTTGTAGA-AATTTTTC-TCTACAAGTTTGT | loxF8L | 3 | loxF8L | 4 | AATTTTTC/GAAAAATT | 7 |
| --- | | | | | | | | 0 |
| chr17 | 55856092 | CTGAGATTAAGGG-TAGTTTGT-CACACAATATTAT | loxF8R | 4 | loxF8R | 2 | ACAAACTA/TAGTTTGT | 6 |
| chr4 | 33025191 | ATAATTCTGTGAA-ACAAACTA-TCTAGAAATTTAG | loxF8L | 2 | loxF8L | 4 | ACAAACTA/TAGTTTGT | 6 |
| chr3 | 127993862 | CTATTGTTGTGTA-ACAAACTA-CACCCAAACTTAG | loxF8R | 4 | loxF8R | 2 | ACAAACTA/TAGTTTGT | 6 |
| chr2 | 198469952 | ATAATGCTTAGGA-TAGTTTGT-TACACAGCATTAT | loxF8L | 4 | loxF8L | 3 | ACAAACTA/TAGTTTGT | 7 |
| chr8 | 20412313 | CTATGGCTGTGTA-ACAAACTA-CTCCCAAACTTAG | loxF8R | 4 | loxF8R | 3 | ACAAACTA/TAGTTTGT | 7 |
| --- | | | | | | | | 0 |
| chr10 | 83944193 | TAAAATCAGTAGA-ACAATAAT-TCCAAAGAATTAT | loxF8L | 4 | loxF8L | 2 | ACAATAAT/ATTATTGT | 6 |
| chr6 | 155933344 | ATAATTGTGTTCA-ACAATAAT-TGAACACATTTAT | loxF8L | 4 | loxF8L | 3 | ACAATAAT/ATTATTGT | 7 |
| chr6 | 71694574 | TTATGATTGTTTG-ATTATTGT-TTTACTGATCTAT | loxF8R | 3 | loxF8L | 4 | ACAATAAT/ATTATTGT | 7 |
| chr8 | 29867696 | ATTAGTCTGAAGA-ACAATAAT-CACATAATCATAT | loxF8L | 4 | loxF8R | 3 | ACAATAAT/ATTATTGT | 7 |
| --- | | | | | | | | 0 |
| chr11 | 21681532 | CTAAGAATGTATG-ACATATTA-CACACACTCAGAA | loxF8R | 2 | loxF8R | 4 | ACATATTA/TAATATGT | 6 |
| chr14 | 90694918 | ATAAATATATGTA-TAATATGT-CACATATATTTAT | loxF8L | 3 | loxF8L | 4 | ACATATTA/TAATATGT | 7 |
| chr11 | 86043094 | CCAAGTTTGGATG-ACATATTA-CACATCACCTTAG | loxF8R | 4 | loxF8R | 3 | ACATATTA/TAATATGT | 7 |
| --- | | | | | | | | 0 |
| chrX | 93045379 | GTAAGTTTGTGTG-CTTAATGT-TCCACATTTCTGT | loxF8R | 2 | loxF8L | 4 | ACATTAAG/CTTAATGT | 6 |
| chrY | 6161711 | GTAAGTTTGTGTG-CTTAATGT-TCCACATTTCTGT | loxF8R | 2 | loxF8L | 4 | ACATTAAG/CTTAATGT | 6 |
| --- | | | | | | | | 0 |
| chrX | 894662 | CCAACATTGTGTG-ACCAGTCA-TGTCCACATTTAT | loxF8R | 2 | loxF8L | 4 | ACCAGTCA/TGACTGGT | 6 |
| chrY | 894662 | CCAACATTGTGTG-ACCAGTCA-TGTCCACATTTAT | loxF8R | 2 | loxF8L | 4 | ACCAGTCA/TGACTGGT | 6 |
| --- | | | | | | | | 0 |
| chr16 | 66958067 | AAAAATCTGTAGA-GATAGGGT-CTCACTATGTTAC | loxF8L | 2 | loxF8R | 4 | ACCCTATC/GATAGGGT | 6 |
| chr16 | 25680357 | ATTTTTCTGTGGA-GATAGGGT-CTCACTATGTTAC | loxF8L | 3 | loxF8R | 4 | ACCCTATC/GATAGGGT | 7 |
| --- | | | | | | | | 0 |
| chr11 | 27050145 | CAAAGATTAATTG-ACCTTATA-TGCACACATTTAT | loxF8R | 4 | loxF8L | 2 | ACCTTATA/TATAAGGT | 6 |
| chrX | 129488694 | ATAAAGCATTAGA-TATAAGGT-CACTCAATTTTAA | loxF8L | 4 | loxF8R | 3 | ACCTTATA/TATAAGGT | 7 |
| --- | | | | | | | | 0 |
| chr14 | 27283238 | ATACAGATGTAGA-ACCTTTTC-TCCATAGGTTTAT | loxF8L | 4 | loxF8L | 2 | ACCTTTTC/GAAAAGGT | 6 |
| chrX | 128665108 | ATAAATTTGGTGA-ACCTTTTC-CATACACTATTAT | loxF8L | 3 | loxF8R | 4 | ACCTTTTC/GAAAAGGT | 7 |
| --- | | | | | | | | 0 |
| chr1 | 77674400 | ATAATTCTGTGGT-ACTAGACA-TCAAAAGAGTGAT | loxF8L | 2 | loxF8L | 4 | ACTAGACA/TGTCTAGT | 6 |
| chr5 | 84577633 | AAGTATCTGTGAA-TGTCTAGT-TCTAGAAATTTAT | loxF8L | 4 | loxF8L | 3 | ACTAGACA/TGTCTAGT | 7 |
| --- | | | | | | | | 0 |
| chr7 | 64589786 | TCAAGATTGTCTT-TATCTAGT-ACCTCAGATTTAT | loxF8R | 4 | loxF8L | 2 | ACTAGATA/TATCTAGT | 6 |
| chr19 | 19907131 | CCCAGATTGTCTT-TATCTAGT-ACTTCAGATTTAT | loxF8R | 4 | loxF8L | 3 | ACTAGATA/TATCTAGT | 7 |
| --- | | | | | | | | 0 |
| chr13 | 87068021 | ATAAAATAGTCTG-TGTATAGT-TCTAGAGATTTAT | loxF8R | 4 | loxF8L | 2 | ACTATACA/TGTATAGT | 6 |
| chr2 | 112945052 | CAAAGATTAGTTG-ACTATACA-TGCACGGGTTTAT | loxF8R | 4 | loxF8L | 3 | ACTATACA/TGTATAGT | 7 |
| chr12 | 108193494 | ATAGATGAATGGA-TGTATAGT-TGGACAGATTAAT | loxF8L | 4 | loxF8L | 3 | ACTATACA/TGTATAGT | 7 |
| --- | | | | | | | | 0 |
| chr13 | 71094684 | AGAAATCTGTTTT-ACTCAAAG-TCCACTGATTTAA | loxF8L | 4 | loxF8L | 2 | ACTCAAAG/CTTTGAGT | 6 |
| chr18 | 32919542 | ATAAATCTGCTTT-ACTCAAAG-GCCACTGATTTAA | loxF8L | 4 | loxF8L | 3 | ACTCAAAG/CTTTGAGT | 7 |
| --- | | | | | | | | 0 |
| chr6 | 156766018 | CTAAGAGGGTCTT-ACTCATCT-TCTACAGTTTTAT | loxF8R | 4 | loxF8L | 2 | ACTCATCT/AGATGAGT | 6 |
| chr4 | 150531243 | CTTATATTTGTA-ACTCATCT-TTCCCAGATTTTT | loxF8R | 4 | loxF8L | 3 | ACTCATCT/AGATGAGT | 7 |
| --- | | | | | | | | 0 |
| chr11 | 134306502 | ACAGATCTTTGGG-CTTAGAGT-GAGACAATCTTAG | loxF8L | 4 | loxF8R | 2 | ACTCTAAG/CTTAGAGT | 6 |
| chr20 | 7539620 | ATGATTCTGTGTG-ACTCTAAG-TCCAAAAATTTGT | loxF8L | 4 | loxF8L | 3 | ACTCTAAG/CTTAGAGT | 7 |
| chr9 | 97994691 | CTGAATTTGTGAA-CTTAGAGT-TCCTCAGTTTAAT | loxF8L | 4 | loxF8L | 3 | ACTCTAAG/CTTAGAGT | 7 |
| --- | | | | | | | | 0 |

| | | | | | | | | |
|---|---|---|---|---|---|---|---|---|
| chr17 | 66767078 | ATAAATGTGTCAG-TATTCAGT-TCCACATATTAAT | loxF8L | 4 | loxF8L | 2 | ACTGAATA/TATTCAGT | 6 |
| chr6 | 9794855 | CTAATTCTGAGGC-ACTGAATA-TGCACAGATTAAA | loxF8L | 4 | loxF8L | 3 | ACTGAATA/TATTCAGT | 7 |
| --- | | | | | | | | 0 |
| chr7 | 18871822 | AGAGATCTGTGGA-GTCTCAGT-TTCATTCATTTAT | loxF8L | 2 | loxF8L | 4 | ACTGAGAC/GTCTCAGT | 6 |
| chr4 | 108551482 | CGAAGAATGTATG-ACTGAGAC-CTCAAAAGCATAG | loxF8R | 3 | loxF8R | 4 | ACTGAGAC/GTCTCAGT | 7 |
| --- | | | | | | | | 0 |
| chr11 | 130712456 | ATAATTGTGAGTA-TGGGCAGT-TTCACAGATTAAT | loxF8L | 4 | loxF8L | 2 | ACTGCCCA/TGGGCAGT | 6 |
| chr10 | 103690069 | CAAGGACTGTGTG-TGGGCAGT-CACCCCATCGGAG | loxF8R | 3 | loxF8R | 4 | ACTGCCCA/TGGGCAGT | 7 |
| --- | | | | | | | | 0 |
| chr8 | 64651823 | ATAAATTTGTGGC-TTATAAGT-CACCCAGTCTATG | loxF8L | 2 | loxF8R | 4 | ACTTATAA/TTATAAGT | 6 |
| chrX | 155035396 | ATAAATAGGTATA-TTATAAGT-TGCACATATTTAA | loxF8L | 4 | loxF8L | 3 | ACTTATAA/TTATAAGT | 7 |
| chr4 | 96134024 | TTAAAAATGTGGA-ACTTATAA-TTCATGAATTTAT | loxF8L | 3 | loxF8L | 4 | ACTTATAA/TTATAAGT | 7 |
| chr9 | 25270328 | ATTAATCTGTCAA-TTATAAGT-TCAACATAAATAT | loxF8L | 3 | loxF8L | 4 | ACTTATAA/TTATAAGT | 7 |
| --- | | | | | | | | 0 |
| chr8 | 6659856 | CCAAGATAGAGAG-TTCAAAGT-CACAAAATCTAAG | loxF8R | 4 | loxF8R | 2 | ACTTTGAA/TTCAAAGT | 6 |
| chr6 | 108966843 | CTGAGAATCTGTG-TTCAAAGT-TCCACAGGTGAGT | loxF8R | 3 | loxF8L | 4 | ACTTTGAA/TTCAAAGT | 7 |
| chr18 | 32205261 | TTGTGATTTTGTG-TTCAAAGT-CACAAAGTCTTAA | loxF8R | 4 | loxF8R | 3 | ACTTTGAA/TTCAAAGT | 7 |
| --- | | | | | | | | 0 |
| chr3 | 162766717 | CAAAGATTGTGTC-ACTTTTTA-AACAAAGTCTTAT | loxF8R | 2 | loxF8R | 4 | ACTTTTTA/TAAAAAGT | 6 |
| chr13 | 61594010 | CCAAGACTATGTT-ACTTTTTA-GAGACAATCTTAT | loxF8R | 4 | loxF8R | 3 | ACTTTTTA/TAAAAAGT | 7 |
| --- | | | | | | | | 0 |
| chr15 | 53275392 | ATAAACGAGTGAA-AGACTTCA-TCCAATGATTTAT | loxF8L | 4 | loxF8L | 2 | AGACTTCA/TGAAGTCT | 6 |
| chr9 | 81459361 | ACAAATTTGTGGC-AGACTTCA-TCTAAAAATTAAT | loxF8L | 3 | loxF8L | 4 | AGACTTCA/TGAAGTCT | 7 |
| --- | | | | | | | | 0 |
| chr17 | 5565522 | CTATGCTTGTGTG-TTATTGCT-CAAGAAATCTTTG | loxF8R | 2 | loxF8R | 4 | AGCAATAA/TTATTGCT | 6 |
| chr19 | 33371583 | TTATGATTTTGTG-AGCAATAA-ATCACAATCCTAT | loxF8R | 3 | loxF8R | 4 | AGCAATAA/TTATTGCT | 7 |
| --- | | | | | | | | 0 |
| chr4 | 106802325 | TTAAGATGGTGTG-AGCCAAAG-ACCACAGTGTTTT | loxF8R | 2 | loxF8L | 4 | AGCCAAAG/CTTTGGCT | 6 |
| chr14 | 90907989 | AGAAATATATGCA-CTTTGGCT-GACATCATCTTAG | loxF8L | 4 | loxF8R | 3 | AGCCAAAG/CTTTGGCT | 7 |
| --- | | | | | | | | 0 |
| chr12 | 83945670 | TTAAGATTATGTG-TGCTTCCT-CACATTTACTTAG | loxF8R | 2 | loxF8R | 4 | AGGAAGCA/TGCTTCCT | 6 |
| chr12 | 46532909 | AAAAAACTGTGAA-AGGAAGCA-TACCAAGGTTTAT | loxF8L | 3 | loxF8L | 4 | AGGAAGCA/TGCTTCCT | 7 |
| --- | | | | | | | | 0 |
| chr18 | 42433538 | CTAATATTGTGTA-GTTTACCT-CCCACATACTTGG | loxF8R | 2 | loxF8R | 4 | AGGTAAAC/GTTTACCT | 6 |
| chr8 | 13020047 | ATGCCTCTGTGGA-AGGTAAAC-TACAAAGATTAGT | loxF8L | 3 | loxF8L | 4 | AGGTAAAC/GTTTACCT | 7 |
| --- | | | | | | | | 0 |
| chr2 | 233011807 | AGAAAGCTGTGGA-ATGTTACT-CACACTGTCATTG | loxF8L | 2 | loxF8R | 4 | AGTAACAT/ATGTTACT | 6 |
| chr12 | 24655438 | CTATGCTTGTGGG-ATGTTACT-CAAGAAATCTTTG | loxF8R | 3 | loxF8R | 4 | AGTAACAT/ATGTTACT | 7 |
| --- | | | | | | | | 0 |
| chr2 | 226299567 | ATAAATTATTTGA-AGTTCTTC-TGCACAGATTTGT | loxF8L | 4 | loxF8L | 2 | AGTTCTTC/GAAGAACT | 6 |
| chr7 | 81386146 | ATAAATTAGAAGA-GAAGAACT-TTCACAGATATGT | loxF8L | 4 | loxF8L | 3 | AGTTCTTC/GAAGAACT | 7 |
| --- | | | | | | | | 0 |
| chr3 | 119482765 | CTGATATTTTGGG-AGTTTGTG-CCCACTGATTTAT | loxF8R | 4 | loxF8L | 2 | AGTTTGTG/CACAAACT | 6 |
| chr1 | 192819805 | CAAAGGTTGCGTG-AGTTTGTG-TTCACTGACTGAT | loxF8R | 3 | loxF8L | 4 | AGTTTGTG/CACAAACT | 7 |
| --- | | | | | | | | 0 |
| chr12 | 41593227 | ATTAATCTGTAGA-AGTTTTTA-TATACAATGATAG | loxF8L | 2 | loxF8R | 4 | AGTTTTTA/TAAAAACT | 6 |
| chr13 | 45073487 | ATCAATCTGAAGA-TAAAAACT-TTCAGAGTTTTTT | loxF8L | 3 | loxF8L | 4 | AGTTTTTA/TAAAAACT | 7 |
| chr2 | 136406055 | AAAAATGTTTGGA-AGTTTTTA-AAAACAGCTTTAT | loxF8L | 3 | loxF8L | 4 | AGTTTTTA/TAAAAACT | 7 |
| chrX | 76227245 | AGAAATATCTGAA-AGTTTTTA-TTCTCAGATTTAG | loxF8L | 4 | loxF8L | 3 | AGTTTTTA/TAAAAACT | 7 |
| chrY | 20443527 | ATAAAAATGTGCA-TAAAAACT-GACACAAAATTAA | loxF8L | 3 | loxF8R | 4 | AGTTTTTA/TAAAAACT | 7 |
| chr17 | 10422854 | GTATGATTGTATG-AGTTTTTA-TACACGTATTGAT | loxF8R | 3 | loxF8L | 4 | AGTTTTTA/TAAAAACT | 7 |
| chr1 | 56162027 | CTAATAAAATGTG-AGTTTTTA-CACTTGATCTTAG | loxF8R | 4 | loxF8R | 3 | AGTTTTTA/TAAAAACT | 7 |
| --- | | | | | | | | 0 |
| chr20 | 23676943 | TTGCATCTCTGGA-ATAAAACC-CACTCAATCTTTG | loxF8L | 4 | loxF8R | 2 | ATAAAACC/GGTTTTAT | 6 |
| chr20 | 23808060 | TTGCATCTCTGGA-ATAAAACC-CACTCAATCTTGG | loxF8L | 4 | loxF8R | 2 | ATAAAACC/GGTTTTAT | 6 |
| chr20 | 23732203 | TTGCATCTCTGGA-ATAAAACC-CACTCAATCTTGT | loxF8L | 4 | loxF8R | 3 | ATAAAACC/GGTTTTAT | 7 |
| chr1 | 187017932 | CCACGATTGAGTG-GGTTTTAT-TCCACGTATGTAA | loxF8R | 3 | loxF8L | 4 | ATAAAACC/GGTTTTAT | 7 |
| --- | | | | | | | | 0 |
| chr5 | 137325755 | AGAAAACAGTGGC-ATAAGCAA-TCCACAGAGTGAT | loxF8L | 4 | loxF8L | 2 | ATAAGCAA/TTGCTTAT | 6 |
| chr14 | 33354032 | AAAAAATAGTGTG-TTGCTTAT-TTCTCAGCTTTAT | loxF8R | 4 | loxF8L | 3 | ATAAGCAA/TTGCTTAT | 7 |
| --- | | | | | | | | 0 |
| chr5 | 71061458 | ATTAACTTATGGA-ATACAATA-CACACAATTATAG | loxF8L | 4 | loxF8R | 2 | ATACAATA/TATTGTAT | 6 |
| chr5 | 70421552 | CTATAATTGTGTG-TATTGTAT-TCCATAAGTTAAT | loxF8R | 2 | loxF8L | 4 | ATACAATA/TATTGTAT | 6 |
| chr5 | 69566403 | CTATAATTATGTG-TATTGTAT-TCCATAAGTTAAT | loxF8R | 3 | loxF8L | 4 | ATACAATA/TATTGTAT | 7 |
| --- | | | | | | | | 0 |
| chr3 | 97961034 | ATAAAACTATGGA-ATACATAA-GCCAAAGAGTTAA | loxF8L | 2 | loxF8L | 4 | ATACATAA/TTATGTAT | 6 |
| chr12 | 76384063 | ATATATGTGATGA-ATACATAA-GCCACACCTTTAT | loxF8L | 4 | loxF8L | 3 | ATACATAA/TTATGTAT | 7 |
| chr4 | 171418842 | TTGAGATTGGGTC-TTATGTAT-TCCTCAAATTCAT | loxF8R | 4 | loxF8L | 3 | ATACATAA/TTATGTAT | 7 |
| --- | | | | | | | | 0 |
| chr4 | 5269790 | CTAACATTATGTG-ATATGAAA-ACCAAAGATATAA | loxF8R | 2 | loxF8L | 4 | ATATGAAA/TTTCATAT | 6 |
| chr3 | 36047964 | TTTTATGTGTGGA-TTTCATAT-CCCACAAATTTAC | loxF8L | 4 | loxF8L | 3 | ATATGAAA/TTTCATAT | 7 |
| --- | | | | | | | | 0 |
| chr8 | 95132622 | GAAAGATTGTGTG-TGTGTGAT-CATACAATTTTCC | loxF8R | 2 | loxF8R | 4 | ATCACACA/TGTGTGAT | 6 |
| chr8 | 113024313 | GTGTGTTTGTGTG-TGTGTGAT-CACACTTTCTTAA | loxF8R | 4 | loxF8R | 3 | ATCACACA/TGTGTGAT | 7 |
| --- | | | | | | | | 0 |
| chr4 | 37316935 | CTAAATCTGTGAA-ATCTATTC-CCCACAATATGTG | loxF8L | 2 | loxF8R | 4 | ATCTATTC/GAATAGAT | 6 |
| chrY | 14174502 | ATACATGTATGCA-GAATAGAT-CATAAAATCTTAA | loxF8L | 4 | loxF8R | 3 | ATCTATTC/GAATAGAT | 7 |
| --- | | | | | | | | 0 |
| chrX | 106545582 | ATAAATGTGACAA-ATTAAATG-CACACATTCTTTG | loxF8L | 4 | loxF8R | 2 | ATTAAATG/CATTTAAT | 6 |
| chr14 | 84769962 | ATAGGATTGTCTT-CATTTAAT-TTCCCATATTTAT | loxF8R | 4 | loxF8L | 3 | ATTAAATG/CATTTAAT | 7 |
| chr9 | 128935934 | ATAGCTCAGTGGT-ATTAAATG-CATTCAATCTTAT | loxF8L | 4 | loxF8R | 3 | ATTAAATG/CATTTAAT | 7 |
| --- | | | | | | | | 0 |
| chr3 | 35524764 | AAAATTCTGGGTA-TATGTAAT-TCCACAGATTACT | loxF8L | 4 | loxF8L | 2 | ATTACATA/TATGTAAT | 6 |
| chr9 | 70051828 | CTTGGAGTGTTTG-ATTACATA-TCCATCTATTTAT | loxF8R | 4 | loxF8L | 3 | ATTACATA/TATGTAAT | 7 |
| --- | | | | | | | | 0 |
| chr9 | 126207157 | ATAAACCTGTAGA-TATGCAAT-ACTACAGTTTTGT | loxF8L | 2 | loxF8L | 4 | ATTGCATA/TATGCAAT | 6 |
| chr4 | 172268053 | CCAAATATCTGGA-TATGCAAT-TCTAAAGATTTAT | loxF8L | 4 | loxF8L | 2 | ATTGCATA/TATGCAAT | 6 |

| | | | | | | | | |
|---|---|---|---|---|---|---|---|---|
| chr13 | 63695846 | ATAAAATTGTATT-TATGCAAT-TTATCAGATTTAT | loxF8R | 4 | loxF8L | 3 | ATTGCATA/TATGCAAT | 7 |
| --- | | | | | | | | 0 |
| chr4 | 86587782 | AAAAATCACTGGC-ATTTAAAA-CACACACTCTTAA | loxF8L | 4 | loxF8R | 2 | ATTTAAAA/TTTTAAAT | 6 |
| chr5 | 108916130 | AAAAATATTTGGA-ATTTAAAA-TAAAAATATTTAT | loxF8L | 3 | loxF8L | 4 | ATTTAAAA/TTTTAAAT | 7 |
| chr19 | 39606518 | AAAAATGTGTGCA-TTTTAAAT-TCGAAAAATTAAT | loxF8L | 3 | loxF8L | 4 | ATTTAAAA/TTTTAAAT | 7 |
| chr8 | 56576383 | CTAAGCTTGACTA-TTTTAAAT-TCCACATATAAAT | loxF8R | 4 | loxF8L | 3 | ATTTAAAA/TTTTAAAT | 7 |
| chr3 | 127824803 | CTAACATTTTATG-ATTTAAAA-TACACAGCTTGAA | loxF8R | 3 | loxF8L | 4 | ATTTAAAA/TTTTAAAT | 7 |
| chr2 | 237710159 | TTAAATATATGTA-TTTTAAAT-GACAAAATCTCAG | loxF8L | 4 | loxF8R | 3 | ATTTAAAA/TTTTAAAT | 7 |
| chr2 | 161273631 | TTAAAATTGTCTA-TTTTAAAT-TGCACAGATTGAA | loxF8R | 4 | loxF8L | 3 | ATTTAAAA/TTTTAAAT | 7 |
| --- | | | | | | | | 0 |
| chr6 | 79025736 | CCAAGGTTATGTT-ATTTAAAG-TCCATATATTTAT | loxF8R | 4 | loxF8L | 2 | ATTTAAAG/CTTTAAAT | 6 |
| chr11 | 115852683 | CTATGATTGTTTC-CTTTAAAT-TCCTAAGGGTTAT | loxF8R | 3 | loxF8L | 4 | ATTTAAAG/CTTTAAAT | 7 |
| --- | | | | | | | | 0 |
| chr7 | 88953944 | TCACCTCTGTGGA-TATTTTTG-TCCACATATTTGT | loxF8L | 4 | loxF8L | 2 | CAAAAATA/TATTTTTG | 6 |
| chr2 | 70911973 | TTAAATTTGTGTT-TATTTTTG-CACACAGTGTGAG | loxF8L | 4 | loxF8R | 3 | CAAAAATA/TATTTTTG | 7 |
| --- | | | | | | | | 0 |
| chr9 | 26965217 | ATAAATGAATAGA-CAAAAATG-TCCATAAATTTAT | loxF8L | 4 | loxF8L | 2 | CAAAAATG/CATTTTTG | 6 |
| chr13 | 26055301 | CTAAACTTATGTT-CAAAAATG-TTCATAAATTTAT | loxF8R | 4 | loxF8L | 3 | CAAAAATG/CATTTTTG | 7 |
| --- | | | | | | | | 0 |
| chr18 | 7246062 | ATAAATGTGTGTA-CAAACACA-CACACAGATAGAT | loxF8L | 2 | loxF8L | 4 | CAAACACA/TGTGTTTG | 6 |
| chr21 | 17472524 | GCAAAACTATGGA-CAAACACA-GCCACAAATTTAT | loxF8L | 4 | loxF8L | 2 | CAAACACA/TGTGTTTG | 6 |
| chr6 | 68259142 | ATATGTGTGTGTG-TGTGTTTG-TGCACAGTTTTTT | loxF8R | 4 | loxF8L | 3 | CAAACACA/TGTGTTTG | 7 |
| chr4 | 6985489 | TTAGGGGTGTGTG-TGTGTTTG-TACACACATATAT | loxF8R | 4 | loxF8L | 3 | CAAACACA/TGTGTTTG | 7 |
| chr4 | 39620829 | CAAAAATGGTGTG-CAAACACA-CACACACACATAC | loxF8R | 3 | loxF8R | 4 | CAAACACA/TGTGTTTG | 7 |
| --- | | | | | | | | 0 |
| chr13 | 87603556 | ATAAATATATACA-CACACAGA-CACACACACTTAG | loxF8L | 4 | loxF8R | 2 | CACACAGA/TCTGTGTG | 6 |
| chr10 | 70855336 | CTAATAGTGTTTG-TCTGTGTG-TATACACATGTAT | loxF8R | 3 | loxF8L | 4 | CACACAGA/TCTGTGTG | 7 |
| --- | | | | | | | | 0 |
| chr6 | 58077225 | CTAGTGTTGTGTT-CCATCGTG-TCAACAGATATAT | loxF8R | 4 | loxF8L | 2 | CACGATGG/CCATCGTG | 6 |
| chr6 | 60829002 | CTAGTGTTGTGTT-CCATCGTG-TCAACGGATATAT | loxF8R | 4 | loxF8L | 3 | CACGATGG/CCATCGTG | 7 |
| --- | | | | | | | | 0 |
| chr2 | 52218258 | ATAAATCTTTAGA-GAACTCTG-AACACATATTTTT | loxF8L | 2 | loxF8L | 4 | CAGAGTTC/GAACTCTG | 6 |
| chr4 | 55491630 | AGAAAAATGTGAA-CAGAGTTC-ACAGCAGATTTAT | loxF8L | 4 | loxF8L | 3 | CAGAGTTC/GAACTCTG | 7 |
| --- | | | | | | | | 0 |
| chrX | 32671342 | TTAAATTTATGTA-CATACACA-CACACACTCTGAG | loxF8L | 4 | loxF8R | 2 | CATACACA/TGTGTATG | 6 |
| chr1 | 21027165 | ATATATATGTGTA-CATACACA-CATACAATTTTTT | loxF8L | 3 | loxF8R | 4 | CATACACA/TGTGTATG | 7 |
| chr3 | 117566900 | ACAAATGTGTGCA-TGTGTATG-CATACACTCTTCA | loxF8L | 3 | loxF8R | 4 | CATACACA/TGTGTATG | 7 |
| chr2 | 15556082 | ATACATGTATGCA-CATACACA-CACACATACATAG | loxF8L | 4 | loxF8R | 3 | CATACACA/TGTGTATG | 7 |
| chr2 | 140303042 | ATATATATATGGA-CATACACA-CACACATATATAT | loxF8L | 3 | loxF8L | 4 | CATACACA/TGTGTATG | 7 |
| chr4 | 187427473 | ATATATATGTATA-CATACACA-CACACATTCAGAG | loxF8L | 4 | loxF8R | 3 | CATACACA/TGTGTATG | 7 |
| chr3 | 65826068 | ATATATGTGTAGA-TGTGTATG-TGCACATATATAC | loxF8L | 3 | loxF8L | 4 | CATACACA/TGTGTATG | 7 |
| chr13 | 76835165 | ATGTATGTGTGTA-CATACACA-CACACAAAGCTAG | loxF8L | 4 | loxF8R | 3 | CATACACA/TGTGTATG | 7 |
| --- | | | | | | | | 0 |
| chr7 | 120052645 | ATAAATGTGTGGC-TATGTATG-TATGCATATTTAT | loxF8L | 2 | loxF8L | 4 | CATACATA/TATGTATG | 6 |
| chr20 | 35628989 | ATACATGTGTGTA-CATACATA-AACACATATTTTT | loxF8L | 3 | loxF8L | 4 | CATACATA/TATGTATG | 7 |
| chr1 | 213109852 | TTTAGAGGGTGTG-TATGTATG-TACATATATTTAT | loxF8R | 4 | loxF8L | 3 | CATACATA/TATGTATG | 7 |
| chrX | 12446190 | ATACATTTCAGGA-CATACATA-TGTACATATTTAT | loxF8L | 4 | loxF8L | 3 | CATACATA/TATGTATG | 7 |
| chr9 | 75842873 | ATATATGTGTGTG-CATACATA-TACACACACTTAT | loxF8L | 4 | loxF8L | 3 | CATACATA/TATGTATG | 7 |
| chrY | 2994058 | ATATGTTTCTGTG-TATGTATG-TCCATATATATAT | loxF8R | 4 | loxF8L | 3 | CATACATA/TATGTATG | 7 |
| chr3 | 50174933 | CCAAGCCTGTGTG-TATGTATG-TACACACGCATAG | loxF8R | 3 | loxF8R | 4 | CATACATA/TATGTATG | 7 |
| --- | | | | | | | | 0 |
| chr6 | 21987998 | ATAGATCCGTACA-CATAGACA-CACACAATCTCAT | loxF8L | 4 | loxF8R | 2 | CATAGACA/TGTCTATG | 6 |
| chrX | 11416034 | GTACGATTTTGTG-TGTCTATG-TGCATATATGTAT | loxF8R | 3 | loxF8L | 4 | CATAGACA/TGTCTATG | 7 |
| chr3 | 153017058 | ATGTATCTGTGTG-CATAGACA-TCCACAAATCTAA | loxF8L | 4 | loxF8L | 3 | CATAGACA/TGTCTATG | 7 |
| --- | | | | | | | | 0 |
| chrX | 21390228 | ATAAATGTGTGTA-TAAATATG-CACACACATTTAC | loxF8L | 2 | loxF8L | 4 | CATATTTA/TAAATATG | 6 |
| chr7 | 21519694 | CTAAAAGTCTGTG-TAAATATG-TCCACCAATTTGA | loxF8R | 3 | loxF8L | 4 | CATATTTA/TAAATATG | 7 |
| --- | | | | | | | | 0 |
| chr16 | 62944959 | ATAAATTTGTATG-TGTGCATG-TCTACATATTTAT | loxF8L | 4 | loxF8L | 2 | CATGCACA/TGTGCATG | 6 |
| chr9 | 39434522 | GTATGATTGTGTG-TGTGCATG-TGCACATGTATAT | loxF8R | 2 | loxF8L | 4 | CATGCACA/TGTGCATG | 6 |
| chr9 | 60993265 | GTATGATTGTGTG-TGTGCATG-TGCACATGTATAT | loxF8R | 2 | loxF8L | 4 | CATGCACA/TGTGCATG | 6 |
| chr6 | 120990392 | TCAAGTTTGTGTG-TGTGCATG-TGCACATATCTTT | loxF8R | 3 | loxF8L | 4 | CATGCACA/TGTGCATG | 7 |
| chr10 | 9518524 | ATACATGTGTGCA-TGTGCATG-CACACACACATAC | loxF8L | 3 | loxF8R | 4 | CATGCACA/TGTGCATG | 7 |
| chr9 | 7687338 | ATATATATGTGCA-TGTGCATG-TATACATATATAT | loxF8L | 3 | loxF8L | 4 | CATGCACA/TGTGCATG | 7 |
| chr2 | 167160315 | TTTATTCTATGGA-TGTGCATG-CACACAAGCATAT | loxF8L | 4 | loxF8R | 3 | CATGCACA/TGTGCATG | 7 |
| --- | | | | | | | | 0 |
| chr17 | 42600631 | ATAATATTATGTA-CATGGTGG-CACACAATATTAT | loxF8R | 4 | loxF8R | 2 | CATGGTGG/CCACCATG | 6 |
| chr11 | 112002634 | AAAATTAGGTGGA-CATGGTGG-CACACACTTGTAG | loxF8L | 4 | loxF8R | 3 | CATGGTGG/CCACCATG | 7 |
| chr10 | 38651775 | AAAATTTTGTGGG-CATGGTGG-CACACAACTGTAG | loxF8L | 4 | loxF8R | 3 | CATGGTGG/CCACCATG | 7 |
| --- | | | | | | | | 0 |
| chr1 | 197021573 | CTGAGATTGTCTG-TTAACATG-CACCCACTCTTCT | loxF8R | 2 | loxF8R | 4 | CATGTTAA/TTAACATG | 6 |
| chr18 | 71383625 | CTAAAATTTTATT-CATGTTAA-ACCACAGTTCTAT | loxF8R | 4 | loxF8L | 3 | CATGTTAA/TTAACATG | 7 |
| --- | | | | | | | | 0 |
| chr5 | 150853198 | TTAAAAGTGTGTT-GTTTAATG-TCCACATATTTGT | loxF8R | 4 | loxF8L | 2 | CATTAAAC/GTTTAATG | 6 |
| chr10 | 54940298 | TAAAGAGTGTGTT-GTTTAATG-TCCAAATATTTGT | loxF8R | 4 | loxF8L | 3 | CATTAAAC/GTTTAATG | 7 |
| chr14 | 65009199 | CACAGATGGGGTG-CATTAAAC-CACAGAGATTTAT | loxF8R | 4 | loxF8L | 3 | CATTAAAC/GTTTAATG | 7 |
| chr2 | 84095088 | CAAAATATGTAGA-CATTAAAC-AACACATTCTTAA | loxF8L | 4 | loxF8R | 3 | CATTAAAC/GTTTAATG | 7 |
| --- | | | | | | | | 0 |
| chr5 | 128681786 | ATTGATCTGTGGA-CATTTACA-TGCATACATTTTT | loxF8L | 2 | loxF8L | 4 | CATTTACA/TGTAAATG | 6 |
| chr3 | 148458548 | ATAAATATGTGTA-TGTAAATG-TCTATATATGTAT | loxF8L | 2 | loxF8L | 4 | CATTTACA/TGTAAATG | 6 |
| --- | | | | | | | | 0 |
| chr22 | 44323199 | ATCCATCTGTCCA-CCCATTCA-TCCACCCATTTAT | loxF8L | 4 | loxF8L | 2 | CCCATTCA/TGAATGGG | 6 |
| chr5 | 127694168 | CTAAGTTTCTGAG-CCCATTCA-TCCTCATATTCAA | loxF8R | 3 | loxF8L | 4 | CCCATTCA/TGAATGGG | 7 |
| --- | | | | | | | | 0 |
| chr3 | 108516022 | ATAAATATGGGGA-TGAGGGGG-TGCAGAAATTCAT | loxF8L | 2 | loxF8L | 4 | CCCCCTCA/TGAGGGGG | 6 |
| chrX | 94312418 | GCAAATATGTGAA-TGAGGGGG-TACAAAATCTGAG | loxF8L | 4 | loxF8R | 3 | CCCCCTCA/TGAGGGGG | 7 |
| --- | | | | | | | | 0 |

| | | | | | | | |
|---|---|---|---|---|---|---|---|
| chr11 | 26020488 | ATAAGACTGTGTG-TTTCTAGG-TGGACAGATTCCT | loxF8R | 2 | loxF8L | 4 | CCTAGAAA/TTTCTAGG | 6 |
| chr8 | 108560878 | ATAAACGTGTGTA-CCTAGAAA-TTCAAAAATTCAT | loxF8L | 3 | loxF8L | 4 | CCTAGAAA/TTTCTAGG | 7 |
| --- | | | | | | | | 0 |
| chrX | 142137238 | ACAAGAGTCTGTG-CCTAGTAA-TCCAGAGAATTAT | loxF8R | 4 | loxF8L | 2 | CCTAGTAA/TTACTAGG | 6 |
| chr2 | 163827115 | AGAATTCTCTGGA-TTACTAGG-CAGACACTCTTGT | loxF8L | 3 | loxF8R | 4 | CCTAGTAA/TTACTAGG | 7 |
| chr2 | 129152605 | AGAATTATCTGGA-TTACTAGG-CAGATAATCTTAC | loxF8L | 4 | loxF8R | 3 | CCTAGTAA/TTACTAGG | 7 |
| chr10 | 108274018 | AGAATTCTCTGGA-TTACTAGG-CAAAGACTCTTAT | loxF8L | 3 | loxF8R | 4 | CCTAGTAA/TTACTAGG | 7 |
| chr3 | 17862624 | AGAATTCTCTGGA-TTACTAGG-CAGAGACTCTTAT | loxF8L | 3 | loxF8R | 4 | CCTAGTAA/TTACTAGG | 7 |
| chr5 | 84894598 | ATAACTTTCTGGA-TTACTAGG-CAGACACTCTTTA | loxF8L | 3 | loxF8R | 4 | CCTAGTAA/TTACTAGG | 7 |
| --- | | | | | | | | 0 |
| chr2 | 31586587 | ATAAGAGTATCTG-CCTGGTAA-TCCAGAGAATTAT | loxF8R | 4 | loxF8L | 2 | CCTGGTAA/TTACCAGG | 6 |
| chr3 | 180093199 | AAAATTCTCTGGA-TTACCAGG-CAGAGACTCTTAT | loxF8L | 3 | loxF8R | 4 | CCTGGTAA/TTACCAGG | 7 |
| chr8 | 130620536 | AGAAATCCCTGGA-TTACCAGG-CACAGACTCTTGT | loxF8L | 3 | loxF8R | 4 | CCTGGTAA/TTACCAGG | 7 |
| chr1 | 192715392 | ACAAGAGTGTCTG-CCTGGTAA-TCCAGAGAATTCT | loxF8R | 4 | loxF8L | 3 | CCTGGTAA/TTACCAGG | 7 |
| chr4 | 76388847 | ACAAGAGTGTCTG-CCTGGTAA-TCCAGAGAATTCT | loxF8R | 4 | loxF8L | 3 | CCTGGTAA/TTACCAGG | 7 |
| chr14 | 30087166 | ACAAGATTCTCTG-CCTGGTAA-TCCAGAGAATTCT | loxF8R | 4 | loxF8L | 3 | CCTGGTAA/TTACCAGG | 7 |
| chrX | 14173487 | AGAATTCTCTGGA-TTACCAGG-CAGAAACTCTTAT | loxF8L | 3 | loxF8R | 4 | CCTGGTAA/TTACCAGG | 7 |
| chrX | 122798441 | GCAAGATTCTCTG-CCTGGTAA-TCCAGAGAATTCT | loxF8R | 4 | loxF8L | 3 | CCTGGTAA/TTACCAGG | 7 |
| chr5 | 100290005 | AGAATTCTCTGGA-TTACCAGG-CAGAGAATTTTAT | loxF8L | 3 | loxF8R | 4 | CCTGGTAA/TTACCAGG | 7 |
| chr9 | 2560855 | GTAAGAGTCTCTG-CCTGGTAA-TCCAGAGAATTCT | loxF8R | 4 | loxF8L | 3 | CCTGGTAA/TTACCAGG | 7 |
| chr13 | 38077069 | ATAAGAGTCTCTG-CCTGGTAA-TCCAGAGAATTCT | loxF8R | 4 | loxF8L | 3 | CCTGGTAA/TTACCAGG | 7 |
| chr15 | 44706548 | AGAATTCTCTGGA-TTACCAGG-CAGAGACTCTTAT | loxF8L | 3 | loxF8R | 4 | CCTGGTAA/TTACCAGG | 7 |
| chr4 | 101876393 | ATAAGAGTCTCTG-CCTGGTAA-TCCAGAGAATTCT | loxF8R | 4 | loxF8L | 3 | CCTGGTAA/TTACCAGG | 7 |
| chr7 | 28339419 | AGAATTCTCTGGA-TTACCAGG-CAGAGACTCTTAT | loxF8L | 3 | loxF8R | 4 | CCTGGTAA/TTACCAGG | 7 |
| chr9 | 122644674 | ATAAGAGTCTCTG-CCTGGTAA-TCCAGAGAATTCT | loxF8R | 4 | loxF8L | 3 | CCTGGTAA/TTACCAGG | 7 |
| chr19 | 12219496 | AGAATTCTCTGGA-TTACCAGG-CAGAGACTCTTGG | loxF8L | 3 | loxF8R | 4 | CCTGGTAA/TTACCAGG | 7 |
| chr14 | 53469310 | AGAATTCTCTGGA-TTACCAGG-CATAGACTCTTAT | loxF8L | 3 | loxF8R | 4 | CCTGGTAA/TTACCAGG | 7 |
| chr7 | 119900115 | ATAATTCTCTGGT-TTACCAGG-CAGATACTCTTAT | loxF8L | 3 | loxF8R | 4 | CCTGGTAA/TTACCAGG | 7 |
| chrX | 20938331 | ATATTTCTCTGGA-TTACCAGG-CAGAGACTCTTAT | loxF8L | 3 | loxF8R | 4 | CCTGGTAA/TTACCAGG | 7 |
| chrX | 93090517 | ATAATGCTCTGGA-TTACCAGG-CAGAGACTCTTAT | loxF8L | 3 | loxF8R | 4 | CCTGGTAA/TTACCAGG | 7 |
| --- | | | | | | | | 0 |
| chr20 | 42225139 | AAAAATCTGTGAA-GTTTAAGG-AACACCATCTAAA | loxF8L | 2 | loxF8R | 4 | CCTTAAAC/GTTTAAGG | 6 |
| chr15 | 38064334 | CTAAGTTTTTGTT-GTTTAAGG-CACCCAGTCTGTG | loxF8R | 3 | loxF8R | 4 | CCTTAAAC/GTTTAAGG | 7 |
| --- | | | | | | | | 0 |
| chr11 | 127980466 | CTAAGATTGTTAG-GCTTTTAG-TGTAAAGTTTTAT | loxF8R | 2 | loxF8L | 4 | CTAAAAGC/GCTTTTAG | 6 |
| chr4 | 80007886 | ATAAATTTGTGTT-GCTTTTAG-TCAACATATGTGT | loxF8L | 3 | loxF8L | 4 | CTAAAAGC/GCTTTTAG | 7 |
| --- | | | | | | | | 0 |
| chrX | 24356621 | CATAGACTGGGTG-GCTTATAG-ACAACAGATTTAT | loxF8R | 4 | loxF8L | 2 | CTATAAGC/GCTTATAG | 6 |
| chr2 | 144270003 | ATTAATCTATATA-CTATAAGC-TTAACAGATTTTT | loxF8L | 4 | loxF8L | 3 | CTATAAGC/GCTTATAG | 7 |
| --- | | | | | | | | 0 |
| chr9 | 19009580 | TTAAATCTGTGCA-GAGAGGAG-TACACAGAATGAA | loxF8L | 2 | loxF8L | 4 | CTCCTCTC/GAGAGGAG | 6 |
| chr1 | 38108883 | ATAAATGTTGGGA-GAGAGGAG-TAAATAATCATAG | loxF8L | 3 | loxF8R | 4 | CTCCTCTC/GAGAGGAG | 7 |
| --- | | | | | | | | 0 |
| chr7 | 99928012 | ATAAATCTATAGA-TATTAGAG-GACACTGACTTAT | loxF8L | 2 | loxF8L | 4 | CTCTAATA/TATTAGAG | 6 |
| chr10 | 93459729 | CCAAGATTATTTG-CTCTAATA-CACAAAATATTGC | loxF8R | 3 | loxF8R | 4 | CTCTAATA/TATTAGAG | 7 |
| --- | | | | | | | | 0 |
| chr5 | 175881287 | TTGAGATTGTGTG-TGTTGCAG-TCATCATATCTAT | loxF8R | 2 | loxF8L | 4 | CTGCAACA/TGTTGCAG | 6 |
| chr3 | 192928263 | ATAAATGGCTGCA-TGTTGCAG-TCCATAGCTATAT | loxF8L | 4 | loxF8L | 3 | CTGCAACA/TGTTGCAG | 7 |
| --- | | | | | | | | 0 |
| chr6 | 64431361 | ATGACTCTGTAGC-TAAATAAG-CACACATTCTGAG | loxF8L | 4 | loxF8R | 2 | CTTATTTA/TAAATAAG | 6 |
| chr1 | 219112913 | CAAAGATTGAAGG-TAAATAAG-CCCACAAGCTGAG | loxF8R | 4 | loxF8R | 3 | CTTATTTA/TAAATAAG | 7 |
| --- | | | | | | | | 0 |
| chrX | 113700770 | CTTAGAATATATG-TGTTCAAG-TTAACAGATTTAT | loxF8R | 4 | loxF8L | 2 | CTTGAACA/TGTTCAAG | 6 |
| chr7 | 41827282 | CAAAATCTATGGA-TGTTCAAG-TCCATATATACAT | loxF8L | 3 | loxF8L | 4 | CTTGAACA/TGTTCAAG | 7 |
| --- | | | | | | | | 0 |
| chr19 | 52382622 | CAAAGTTTCTGTT-GAAAAATA-CACATAATCTTAT | loxF8R | 4 | loxF8R | 2 | GAAAAATA/TATTTTTC | 6 |
| chr2 | 129259792 | TTATTTTTGTGTG-TATTTTTC-TGCACCGAATTAT | loxF8R | 4 | loxF8L | 3 | GAAAAATA/TATTTTTC | 7 |
| --- | | | | | | | | 0 |
| chr19 | 40668831 | CTAAGATTGCATG-GATCTTTC-TCCACATGTATAA | loxF8R | 2 | loxF8L | 4 | GAAAGATC/GATCTTTC | 6 |
| chr18 | 1883387 | GTAACTTTCTGTG-GATCTTTC-CAGACAATTTTAC | loxF8R | 4 | loxF8R | 3 | GAAAGATC/GATCTTTC | 7 |
| --- | | | | | | | | 0 |
| chr4 | 27441833 | ATAAATATTTATA-GAACATTA-CACTCAATGTTAG | loxF8L | 4 | loxF8R | 2 | GAACATTA/TAATGTTC | 6 |
| chr12 | 100205824 | ATAAACTTATGAA-TAATGTTC-TCCGCACATTTAA | loxF8L | 4 | loxF8L | 3 | GAACATTA/TAATGTTC | 7 |
| chr10 | 30717997 | CCATGATGGTTTG-GAACATTA-TCTACAGTTTTAC | loxF8R | 4 | loxF8L | 3 | GAACATTA/TAATGTTC | 7 |
| --- | | | | | | | | 0 |
| chr2 | 199589112 | CTAAGAGAGGGGG-GAATATCA-TCCATAGATTTTT | loxF8R | 4 | loxF8L | 2 | GAATATCA/TGATATTC | 6 |
| chr5 | 83308956 | ACAAATCCTTAGA-GAATATCA-TCCATAGAAATAT | loxF8L | 4 | loxF8L | 3 | GAATATCA/TGATATTC | 7 |
| --- | | | | | | | | 0 |
| chr11 | 18881741 | ATATTTCTGTCCA-GTCCATTC-CACACAATCATAA | loxF8L | 4 | loxF8R | 2 | GAATGGAC/GTCCATTC | 6 |
| chr8 | 131720506 | ATAAATGCGTGGT-GAATGGAC-CACAGTCTCTCAG | loxF8L | 3 | loxF8R | 4 | GAATGGAC/GTCCATTC | 7 |
| --- | | | | | | | | 0 |
| chrX | 142836862 | ATAAATATATGGA-GACATACA-CACACCCACTTAC | loxF8L | 2 | loxF8R | 4 | GACATACA/TGTATGTC | 6 |
| chr7 | 83258527 | ATGATTCTGTGTG-TGTATGTC-TCTACAAATATAT | loxF8L | 4 | loxF8L | 3 | GACATACA/TGTATGTC | 7 |
| --- | | | | | | | | 0 |
| chr11 | 113980873 | ATAAATATGTTGA-GACCAAGA-TCCAAGAAGTTAT | loxF8L | 2 | loxF8L | 4 | GACCAAGA/TCTTGGTC | 6 |
| chr7 | 121747275 | CTACGAATATGTG-GACCAAGA-TCCACAAATACCT | loxF8R | 3 | loxF8L | 4 | GACCAAGA/TCTTGGTC | 7 |
| --- | | | | | | | | 0 |
| chr8 | 5584423 | ACAAATATGTGGA-GACTGAAA-ACCACATGTATAT | loxF8L | 2 | loxF8L | 4 | GACTGAAA/TTTCAGTC | 6 |
| chr13 | 25683445 | ATACTTCTGTGGG-TTTCAGTC-TACACATTTGTAT | loxF8L | 3 | loxF8L | 4 | GACTGAAA/TTTCAGTC | 7 |
| --- | | | | | | | | 0 |
| chr2 | 154055743 | CTTAAATTATATG-GTTTTCTC-GCCACAGATTTAG | loxF8R | 4 | loxF8L | 2 | GAGAAAAC/GTTTTCTC | 6 |
| chr6 | 10890725 | CTGTGATTTTGAG-GTTTTCTC-CATAAAATCTTTG | loxF8R | 4 | loxF8R | 3 | GAGAAAAC/GTTTTCTC | 7 |
| --- | | | | | | | | 0 |
| chr14 | 19266618 | ATAATACTGTGGA-GAGTAATA-TCAGCACTTTTAT | loxF8L | 2 | loxF8L | 4 | GAGTAATA/TATTACTC | 6 |
| chr14 | 19133352 | ATAAAAGTGCTGA-TATTACTC-TCCACAGTATTAT | loxF8L | 4 | loxF8L | 2 | GAGTAATA/TATTACTC | 6 |
| chr22 | 15855705 | ATAAAAGTGCTGA-TATTACTC-TCCACAGTATTAT | loxF8L | 4 | loxF8L | 2 | GAGTAATA/TATTACTC | 6 |
| --- | | | | | | | | 0 |

| | | | | | | | | |
|---|---|---|---|---|---|---|---|---|
| chr4 | 101644183 | CTAAGAATGTGTA-TTATTATC-TCAATAGAGTTTT | loxF8R | 2 | loxF8L | 4 | GATAATAA/TTATTATC | 6 |
| chr18 | 66530320 | ATCAATATATGGA-TTATTATC-CCCCCAATTTTAC | loxF8L | 3 | loxF8R | 4 | GATAATAA/TTATTATC | 7 |
| --- | | | | | | | | 0 |
| chr21 | 30534452 | GTAAATCTGTGGG-GATAATGA-CTCTCACTGTTAG | loxF8L | 2 | loxF8R | 4 | GATAATGA/TCATTATC | 6 |
| chr8 | 116316776 | TAAAAATTGTGTG-GATAATGA-ATGAAAATCTTAG | loxF8R | 3 | loxF8R | 4 | GATAATGA/TCATTATC | 7 |
| --- | | | | | | | | 0 |
| chr2 | 185129609 | ACACATATGTGTA-TATATATC-TCCACACATATAT | loxF8L | 4 | loxF8L | 2 | GATATATA/TATATATC | 6 |
| chr1 | 84987343 | ATAGATATATGTA-GATATATA-TCCATATATTTAT | loxF8L | 4 | loxF8L | 2 | GATATATA/TATATATC | 6 |
| chr10 | 27994571 | ATATATATGTGTA-TATATATC-TCCACATATGGGT | loxF8L | 3 | loxF8L | 4 | GATATATA/TATATATC | 7 |
| chr3 | 46741274 | AGATATATGTAGA-TATATATC-TCTAGAGATATAT | loxF8L | 4 | loxF8L | 3 | GATATATA/TATATATC | 7 |
| chr20 | 56281139 | ATAAATCTATATA-GATATATA-TACATAAATCTAT | loxF8L | 3 | loxF8L | 4 | GATATATA/TATATATC | 7 |
| chr20 | 56281187 | ATAAATCTATATA-GATATATA-TACATAAATCTAT | loxF8L | 3 | loxF8L | 4 | GATATATA/TATATATC | 7 |
| chr20 | 56281211 | ATAAATCTATATA-GATATATA-TACATAAATCTAT | loxF8L | 3 | loxF8L | 4 | GATATATA/TATATATC | 7 |
| chr20 | 56281163 | ATAAATCTATATA-GATATATA-TACATAAATCTAT | loxF8L | 3 | loxF8L | 4 | GATATATA/TATATATC | 7 |
| chr20 | 56281057 | ATAAATCTATATA-GATATATA-TACATAAATCTAT | loxF8L | 3 | loxF8L | 4 | GATATATA/TATATATC | 7 |
| chr20 | 56281081 | ATAAATCTATATA-GATATATA-TACATAAATCTAT | loxF8L | 3 | loxF8L | 4 | GATATATA/TATATATC | 7 |
| chr20 | 56281033 | ATAAATCTATATA-GATATATA-TACATAAATCTAT | loxF8L | 3 | loxF8L | 4 | GATATATA/TATATATC | 7 |
| chr20 | 56281257 | ATAAATCTATATA-GATATATA-TACATAAATCTAT | loxF8L | 3 | loxF8L | 4 | GATATATA/TATATATC | 7 |
| chr20 | 56281281 | ATAAATCTATATA-GATATATA-TACATAAATCTAT | loxF8L | 3 | loxF8L | 4 | GATATATA/TATATATC | 7 |
| chr20 | 56281305 | ATAAATCTATATA-GATATATA-TACATAAATCTAT | loxF8L | 3 | loxF8L | 4 | GATATATA/TATATATC | 7 |
| chr3 | 154631544 | ATGAATGTGTGTA-GATATATA-GCCATATAATTAT | loxF8L | 3 | loxF8L | 4 | GATATATA/TATATATC | 7 |
| chr2 | 210512835 | ATACATATATGGA-GATATATA-TATATAGATATAT | loxF8L | 3 | loxF8L | 4 | GATATATA/TATATATC | 7 |
| chr11 | 32755844 | ATATATGTGTGTG-TATATATC-TCCATATATATAT | loxF8L | 4 | loxF8L | 3 | GATATATA/TATATATC | 7 |
| chrX | 55611196 | ATATATATATGGA-GATATATA-TACGGAGATATAT | loxF8L | 3 | loxF8L | 4 | GATATATA/TATATATC | 7 |
| chr6 | 67118006 | ATATATCTGCATA-TATATATC-TCCATATATATAT | loxF8L | 4 | loxF8L | 3 | GATATATA/TATATATC | 7 |
| chr6 | 67118034 | ATATATCTGCATA-TATATATC-TCCATATATATAT | loxF8L | 4 | loxF8L | 3 | GATATATA/TATATATC | 7 |
| chr10 | 83834664 | ATATGTGTGTGTA-TATATATC-TGCACACATATAT | loxF8L | 4 | loxF8L | 3 | GATATATA/TATATATC | 7 |
| chr6 | 9604207 | CTACATATGTGTA-GATATATA-TACACACATATAT | loxF8L | 4 | loxF8L | 3 | GATATATA/TATATATC | 7 |
| --- | | | | | | | | 0 |
| chr1 | 230466909 | ACATATCTATGTA-GTCATATC-CACACAATGGTAG | loxF8L | 4 | loxF8R | 2 | GATATGAC/GTCATATC | 6 |
| chr3 | 109890872 | CTGTGATTTTGAG-GTCATATC-CAAAAAATCTTTG | loxF8R | 4 | loxF8R | 3 | GATATGAC/GTCATATC | 7 |
| chr5 | 67905992 | CAATGATTTTGTG-GATATGAC-CCCAAAAGCTCAG | loxF8R | 3 | loxF8R | 4 | GATATGAC/GTCATATC | 7 |
| --- | | | | | | | | 0 |
| chr12 | 20988046 | TTAAAAATGTGTT-TTTTAATC-TCCACATATTTGT | loxF8R | 4 | loxF8L | 2 | GATTAAAA/TTTTAATC | 6 |
| chrX | 113020309 | ATAAATATGTGGG-GATTAAAA-AACACAGTCCTAC | loxF8L | 2 | loxF8R | 4 | GATTAAAA/TTTTAATC | 6 |
| chr17 | 5830994 | ATAAAACTGCTGA-GATTAAAA-AATACAGATTTTT | loxF8L | 3 | loxF8L | 4 | GATTAAAA/TTTTAATC | 7 |
| --- | | | | | | | | 0 |
| chr1 | 22442035 | TTAAGAATGCGTT-GTTTAATC-TCCACAAATTTGT | loxF8R | 4 | loxF8L | 2 | GATTAAAC/GTTTAATC | 6 |
| chr18 | 30655343 | AAAAATGTTTGGA-GATTAAAC-AACACATTTTTAA | loxF8L | 3 | loxF8R | 4 | GATTAAAC/GTTTAATC | 7 |
| chrX | 102610040 | TTAAGAGTGTGTT-GTTTAATC-TCCATATATATGT | loxF8R | 3 | loxF8L | 4 | GATTAAAC/GTTTAATC | 7 |
| chr13 | 88510476 | ATAAATATCTGTT-GTTTAATC-CACACAGTCTCTG | loxF8L | 4 | loxF8R | 3 | GATTAAAC/GTTTAATC | 7 |
| chrY | 7323306 | CATAGATTGGGTG-GATTAAAC-AACATTGATTTAT | loxF8R | 3 | loxF8L | 4 | GATTAAAC/GTTTAATC | 7 |
| chrX | 3657505 | ATAAATCAGTGTT-GTTTAATC-CACACCGTCTATG | loxF8L | 3 | loxF8R | 4 | GATTAAAC/GTTTAATC | 7 |
| chr3 | 157061024 | ATAAATGTCTGTT-GTTTAATC-CACCCAGTCTTTG | loxF8L | 4 | loxF8R | 3 | GATTAAAC/GTTTAATC | 7 |
| chr6 | 50428615 | CATAGACTGTGTA-GATTAAAC-TACAGAAATTTAT | loxF8R | 4 | loxF8L | 3 | GATTAAAC/GTTTAATC | 7 |
| chr2 | 8986950 | ATAAATTTCTGTT-GTTTAATC-CACACAGTCTATG | loxF8L | 4 | loxF8R | 3 | GATTAAAC/GTTTAATC | 7 |
| chr8 | 57149056 | CATAGACTGTGTG-GATTAAAC-AACAGAAATTTAT | loxF8R | 3 | loxF8L | 4 | GATTAAAC/GTTTAATC | 7 |
| chr11 | 83378545 | CATAGATTGGGTG-GTTTAATC-AACAGAAATTTAT | loxF8R | 3 | loxF8L | 4 | GATTAAAC/GTTTAATC | 7 |
| chr7 | 116003846 | CAAAATATTTGGA-GATTAAAC-CACACACTTTTAC | loxF8L | 4 | loxF8R | 3 | GATTAAAC/GTTTAATC | 7 |
| chr5 | 13437019 | GTATGACTGGGTG-GATTAAAC-AACAGAAACTTAG | loxF8R | 4 | loxF8R | 3 | GATTAAAC/GTTTAATC | 7 |
| --- | | | | | | | | 0 |
| chr2 | 44466387 | ATAAATATGTGGT-GATTAATC-AACACACTCTTGA | loxF8L | 2 | loxF8R | 4 | GATTAATC/GATTAATC | 6 |
| --- | | | | | | | | 0 |
| chr2 | 171104880 | ATAAATCTATGCA-TTTATGGC-CAACTAATCTTCG | loxF8L | 2 | loxF8R | 4 | GCCATAAA/TTTATGGC | 6 |
| chr14 | 59686007 | ATAAATCCATGCA-TTTATGGC-CAACCAATTTTTG | loxF8L | 3 | loxF8R | 4 | GCCATAAA/TTTATGGC | 7 |
| --- | | | | | | | | 0 |
| chrX | 131031041 | ATAAAACTGTGGT-TATGTAGC-TACTCATATTCAT | loxF8L | 2 | loxF8L | 4 | GCTACATA/TATGTAGC | 6 |
| chr2 | 209643037 | GTTCATCTGTGGA-TATGTAGC-AGCACAAAATTAT | loxF8L | 3 | loxF8L | 4 | GCTACATA/TATGTAGC | 7 |
| --- | | | | | | | | 0 |
| chr4 | 57030765 | AGCAGAATGTGTG-TTCTCAGC-TCCACTCATTTAT | loxF8R | 4 | loxF8L | 2 | GCTGAGAA/TTCTCAGC | 6 |
| chr8 | 129862841 | CTAAGAGTGGGTC-TTCTCAGC-TCCATAGATACTT | loxF8R | 3 | loxF8L | 4 | GCTGAGAA/TTCTCAGC | 7 |
| --- | | | | | | | | 0 |
| chr4 | 115925417 | ATAAATCTATGCA-TTTACAGC-CCCCCAATTTTTG | loxF8L | 2 | loxF8R | 4 | GCTGTAAA/TTTACAGC | 6 |
| chr13 | 72339813 | ATAAATCCATGCA-TTTACAGC-CAAATCATCTTTG | loxF8L | 3 | loxF8R | 4 | GCTGTAAA/TTTACAGC | 7 |
| chr18 | 9229102 | CAAAGATCATTTG-GCTGTAAA-TATGCAGATTTAT | loxF8R | 4 | loxF8L | 3 | GCTGTAAA/TTTACAGC | 7 |
| --- | | | | | | | | 0 |
| chrX | 34121227 | CATAGACTGGGTG-GCTTATAA-ACAACAGATTTAT | loxF8R | 4 | loxF8L | 2 | GCTTATAA/TTATAAGC | 6 |
| chr6 | 68394378 | CATAGATTGGGTA-GCTTATAA-ACAACAGATTTTT | loxF8R | 4 | loxF8L | 3 | GCTTATAA/TTATAAGC | 7 |
| chr8 | 5325556 | ATAAATCTTTGTT-TTATAAGC-CCCTCAGATTTCA | loxF8L | 3 | loxF8L | 4 | GCTTATAA/TTATAAGC | 7 |
| chr2 | 78678897 | CCTAGATTGGGTG-GCTTATAA-ACAAGACATTTAT | loxF8R | 3 | loxF8L | 4 | GCTTATAA/TTATAAGC | 7 |
| chr4 | 80790697 | ATAAATGTGTTGT-TTATAAGC-CATCCAATCTATG | loxF8L | 3 | loxF8R | 4 | GCTTATAA/TTATAAGC | 7 |
| chr2 | 21757042 | ATATGCTTGGGTG-GCTTATAA-ACAACAAATTTAT | loxF8R | 4 | loxF8L | 3 | GCTTATAA/TTATAAGC | 7 |
| chr9 | 95768765 | ATAAATTTGTTGT-TTATAAGC-CACCCAGTCTATG | loxF8L | 3 | loxF8R | 4 | GCTTATAA/TTATAAGC | 7 |
| chr4 | 28575182 | CATAGATTGAGTG-GCTTATAA-ACAACAGAAATAT | loxF8R | 3 | loxF8L | 4 | GCTTATAA/TTATAAGC | 7 |
| chr9 | 118112068 | CTTAAATTCTGTG-GCTTATAA-ACCACAGAATTTA | loxF8R | 3 | loxF8L | 4 | GCTTATAA/TTATAAGC | 7 |
| --- | | | | | | | | 0 |
| chr6 | 146852039 | CAAAGACTGTGTG-GCTTGAAC-CACAGAAATTTAT | loxF8R | 2 | loxF8L | 4 | GCTTGAAC/GTTCAAGC | 6 |
| chr5 | 93731478 | ATAAATTTCTGTT-GTTCAAGC-CACCCAATCTGTG | loxF8L | 4 | loxF8R | 3 | GCTTGAAC/GTTCAAGC | 7 |
| --- | | | | | | | | 0 |
| chr2 | 160337652 | AAAAAACTGGGGG-GGAGGGGA-ACCACATATTTAT | loxF8L | 4 | loxF8L | 2 | GGAGGGGA/TCCCCTCC | 6 |
| chr1 | 76290874 | ATAAAACAGGGGA-GGAGGGGA-TCCACACATACCT | loxF8L | 3 | loxF8L | 4 | GGAGGGGA/TCCCCTCC | 7 |
| --- | | | | | | | | 0 |
| chr9 | 63885238 | ACACATCTGTGGA-TTCAAACC-ACCACAGATGGAA | loxF8L | 2 | loxF8L | 4 | GGTTTGAA/TTCAAACC | 6 |
| chr9 | 40966745 | TTCCATCTGTGGT-GGTTTGAA-TCCACAGATGTGT | loxF8L | 4 | loxF8L | 2 | GGTTTGAA/TTCAAACC | 6 |
| chr21 | 5216674 | CAACATCTGTGGA-TTCAAACC-ACCACAGATGGAA | loxF8L | 3 | loxF8L | 4 | GGTTTGAA/TTCAAACC | 7 |
| chr4 | 189908610 | TTCCATCTGTGGT-GGTTTGAA-TCCACAGATGTGG | loxF8L | 4 | loxF8L | 3 | GGTTTGAA/TTCAAACC | 7 |
| chr20 | 28630343 | TCACATCTGTGGA-TTCAAACC-ACCACAGATGGAA | loxF8L | 3 | loxF8L | 4 | GGTTTGAA/TTCAAACC | 7 |

| | | | | | | | | |
|---|---|---|---|---|---|---|---|---|
| --- | | | | | | | | 0 |
| chr1 | 35171158 | ACAGATATGTGCA-TCCATCAC-CACTCAATTTTAG | loxF8L | 4 | loxF8R | 2 | GTGATGGA/TCCATCAC | 6 |
| chr6 | 55056784 | TCAAATCTGAGTA-TCCATCAC-TTCAAATATTTAT | loxF8L | 4 | loxF8L | 3 | GTGATGGA/TCCATCAC | 7 |
| --- | | | | | | | | 0 |
| chr1 | 77255551 | CTAACACTGTGTG-GTGTATAA-ACCACTCATTTAC | loxF8R | 2 | loxF8L | 4 | GTGTATAA/TTATACAC | 6 |
| chr8 | 26750015 | ATAAATAGGTGGC-TTATACAC-ATCAGAAATTTAT | loxF8L | 3 | loxF8L | 4 | GTGTATAA/TTATACAC | 7 |
| --- | | | | | | | | 0 |
| chr8 | 71732757 | CTAAGATTGAGTT-GTTTCAAC-TAAACACATTTAG | loxF8R | 2 | loxF8L | 4 | GTTGAAAC/GTTTCAAC | 6 |
| chr6 | 55237877 | CCAGGAGTATGTG-GTTGAAAC-TGCACAGTTTTCT | loxF8R | 4 | loxF8L | 3 | GTTGAAAC/GTTTCAAC | 7 |
| chr3 | 21656284 | CATAGACTGGGTG-GTTTCAAC-AACACACATTTAT | loxF8R | 4 | loxF8L | 3 | GTTGAAAC/GTTTCAAC | 7 |
| --- | | | | | | | | 0 |
| chr6 | 26113677 | CATAGATTGTGTG-GTTTAAAC-AACAAAATTTTAT | loxF8R | 2 | loxF8R | 4 | GTTTAAAC/GTTTAAAC | 6 |
| chr5 | 87191176 | CATAGAATGAGTG-GTTTAAAC-TACACACATTTAT | loxF8R | 4 | loxF8L | 2 | GTTTAAAC/GTTTAAAC | 6 |
| chr13 | 60651017 | ATAAATGTTTGTT-GTTTAAAC-CACACAATCTATG | loxF8L | 4 | loxF8R | 2 | GTTTAAAC/GTTTAAAC | 6 |
| chr8 | 38614727 | ATAAAAGTGTGTT-GTTTAAAC-CACCCAATCTATG | loxF8L | 4 | loxF8R | 3 | GTTTAAAC/GTTTAAAC | 7 |
| chr16 | 58686115 | CATAGACTGGGTG-GTTTAAAC-GACAGAGATTTAT | loxF8R | 4 | loxF8L | 3 | GTTTAAAC/GTTTAAAC | 7 |
| chr4 | 116154909 | CTCTAATTGGGTG-GTTTAAAC-AACAGAGATTTAT | loxF8R | 4 | loxF8L | 3 | GTTTAAAC/GTTTAAAC | 7 |
| chr14 | 93853735 | ATAAATCTTTGTT-GTTTAAAC-CACCCAGTCTGGG | loxF8L | 3 | loxF8R | 4 | GTTTAAAC/GTTTAAAC | 7 |
| chr1 | 178936727 | CATAGACTGGGTG-GTTTAAAC-AACACACATTTAT | loxF8R | 4 | loxF8L | 3 | GTTTAAAC/GTTTAAAC | 7 |
| chr4 | 87787176 | ATAAATGTGTGTT-GTTTAAAC-CACCCAGTCTATG | loxF8L | 3 | loxF8R | 4 | GTTTAAAC/GTTTAAAC | 7 |
| chr14 | 31446514 | ATAGGACTGGGTG-GTTTAAAC-ACCAGAAATTTAT | loxF8R | 4 | loxF8L | 3 | GTTTAAAC/GTTTAAAC | 7 |
| chr16 | 5777090 | CTCACACTTTGTG-GTTTAAAC-AACACAAATTTAT | loxF8R | 4 | loxF8L | 3 | GTTTAAAC/GTTTAAAC | 7 |
| chr2 | 31921304 | ATAAATTTGTGTT-GTTTAAAC-CACCTAATCTGTG | loxF8L | 3 | loxF8R | 4 | GTTTAAAC/GTTTAAAC | 7 |
| chr2 | 14793218 | ATAAATTTTTGTA-GTTTAAAC-CACCCAGTCTATG | loxF8L | 3 | loxF8R | 4 | GTTTAAAC/GTTTAAAC | 7 |
| chr1 | 29916180 | CACAGCCTGTGTG-GTTTAAAC-CACAGAAACTTAT | loxF8R | 4 | loxF8R | 3 | GTTTAAAC/GTTTAAAC | 7 |
| --- | | | | | | | | 0 |
| chr2 | 209454860 | CATAGATTGTGTG-TCTTAAAC-AGCAGACATTTAT | loxF8R | 2 | loxF8L | 4 | GTTTAAGA/TCTTAAAC | 6 |
| chr6 | 33478811 | CCCAGACTGTGGG-TCTTAAAC-AGCAGAGATTTAT | loxF8R | 4 | loxF8L | 3 | GTTTAAGA/TCTTAAAC | 7 |
| chr6 | 135846543 | CACAGACTGAGTG-TCTTAAAC-AACAGAGATTTAT | loxF8R | 4 | loxF8L | 3 | GTTTAAGA/TCTTAAAC | 7 |
| chr16 | 61207735 | ATAAATCTCTGTT-GTTTAAGA-CACTGAATCTATG | loxF8L | 3 | loxF8R | 4 | GTTTAAGA/TCTTAAAC | 7 |
| chr5 | 13504326 | CACAGATGGGGTG-TCTTAAAC-ACCAGAAATTTAT | loxF8R | 4 | loxF8L | 3 | GTTTAAGA/TCTTAAAC | 7 |
| chr10 | 121442703 | ATAAATTTCTGTC-GTTTAAGA-CACTCAATCTATG | loxF8L | 4 | loxF8R | 3 | GTTTAAGA/TCTTAAAC | 7 |
| chr2 | 138953810 | CACAGATTGAGTG-TCTTAAAC-AACAGAAATTTAT | loxF8R | 3 | loxF8L | 4 | GTTTAAGA/TCTTAAAC | 7 |
| chr3 | 86882987 | ATAAATTTTTGTT-GTTTAAGA-CACTCAGTCTTGG | loxF8L | 4 | loxF8R | 3 | GTTTAAGA/TCTTAAAC | 7 |
| chr8 | 41216078 | CACAGATTGAGTG-TCTTAAAC-AACAGAAACTTGG | loxF8R | 3 | loxF8R | 4 | GTTTAAGA/TCTTAAAC | 7 |
| --- | | | | | | | | 0 |
| chr10 | 127395187 | CACAGACTGGGTG-GTTTATAA-ACCACAGATTCAT | loxF8R | 4 | loxF8L | 2 | GTTTATAA/TTATAAAC | 6 |
| chr16 | 74234503 | AAACATCTGTTGT-TTATAAAC-CACCCAATCTGTG | loxF8L | 4 | loxF8R | 3 | GTTTATAA/TTATAAAC | 7 |
| chr8 | 124807787 | ATAAAATTCTGTG-GTTTATAA-GCCACAAGTCTAT | loxF8R | 3 | loxF8L | 4 | GTTTATAA/TTATAAAC | 7 |
| chr5 | 106209157 | ATAAATAGGTAGC-TTATAAAC-AACACAAATTTAT | loxF8L | 4 | loxF8L | 3 | GTTTATAA/TTATAAAC | 7 |
| chr16 | 5995853 | ATCAATGTGTAGT-TTATAAAC-AACAGAGATTTAT | loxF8L | 4 | loxF8L | 3 | GTTTATAA/TTATAAAC | 7 |
| chr13 | 86483245 | ATAAATTTCTGTT-GTTTATAA-GCCACCCATTTAT | loxF8L | 4 | loxF8L | 3 | GTTTATAA/TTATAAAC | 7 |
| chr3 | 6521139 | ATAAATGGGTGGC-TTATAAAC-AGCAGAAATTTAT | loxF8L | 3 | loxF8L | 4 | GTTTATAA/TTATAAAC | 7 |
| chr3 | 150143244 | CAAAGATATTGTG-TTATAAAC-TGCCCAATCCTAG | loxF8R | 3 | loxF8R | 4 | GTTTATAA/TTATAAAC | 7 |
| --- | | | | | | | | 0 |
| chr3 | 97188078 | GTAAAGCTGTGGA-GTTTATGA-TCCAGAAGTTTCT | loxF8L | 2 | loxF8L | 4 | GTTTATGA/TCATAAAC | 6 |
| chr1 | 241948406 | CTCAGTTTTAGTG-GTTTATGA-TCCAAGCATTTAT | loxF8R | 4 | loxF8L | 3 | GTTTATGA/TCATAAAC | 7 |
| --- | | | | | | | | 0 |
| chr17 | 41051966 | AACAGATTTTGTG-TAACAAAC-TCCAGAGCTTTAT | loxF8R | 4 | loxF8L | 2 | GTTTGTTA/TAACAAAC | 6 |
| chr20 | 40734448 | ATCAATGTGTGGA-TAACAAAC-CACACACCCCTAT | loxF8L | 2 | loxF8R | 4 | GTTTGTTA/TAACAAAC | 6 |
| chr1 | 173933064 | ATAAACATTAGGA-TAACAAAC-TCCACAGGATTTT | loxF8L | 4 | loxF8L | 3 | GTTTGTTA/TAACAAAC | 7 |
| --- | | | | | | | | 0 |
| chr4 | 14299861 | CTATGTTTGTGTG-TGTGTTTA-TACACAGACATAC | loxF8R | 2 | loxF8L | 4 | TAAACACA/TGTGTTTA | 6 |
| chr4 | 14305367 | CATACATTGTGTG-TGTGTTTA-TACACAGACATAC | loxF8R | 3 | loxF8L | 4 | TAAACACA/TGTGTTTA | 7 |
| --- | | | | | | | | 0 |
| chr4 | 21714029 | GATATATTGTGTG-TAAATACA-TCATCAGATTTAT | loxF8R | 4 | loxF8L | 2 | TAAATACA/TGTATTTA | 6 |
| chrX | 89211835 | AAAATTCTCTGGA-TAAATACA-CTGACAGTTTTAT | loxF8L | 3 | loxF8L | 4 | TAAATACA/TGTATTTA | 7 |
| chrY | 6541139 | AAAATTCTCTGGA-TAAATACA-CTGACAGTTTTAT | loxF8L | 3 | loxF8L | 4 | TAAATACA/TGTATTTA | 7 |
| chr4 | 166089418 | CTAGGTTTTTGGG-TAAATACA-TTGACAGGTTTAT | loxF8R | 4 | loxF8L | 3 | TAAATACA/TGTATTTA | 7 |
| chrX | 85088073 | ATACATATATGTA-TAAATACA-TATACATATTTAT | loxF8L | 4 | loxF8L | 3 | TAAATACA/TGTATTTA | 7 |
| chr8 | 2898826 | ATACATATATGCA-TAAATACA-TACACACATATAT | loxF8L | 4 | loxF8L | 3 | TAAATACA/TGTATTTA | 7 |
| --- | | | | | | | | 0 |
| chr2 | 157051883 | ATAAATCTATGTA-TAAATGCA-GGTTCAGATTTAT | loxF8L | 2 | loxF8L | 4 | TAAATGCA/TGCATTTA | 6 |
| chr8 | 52636279 | AAAACTCTGTGTA-TGCATTTA-TCTAGAGATGTCT | loxF8L | 3 | loxF8L | 4 | TAAATGCA/TGCATTTA | 7 |
| chr9 | 33948062 | AAAATTCTGTCCA-TGCATTTA-CACAAAGTGTTAG | loxF8L | 4 | loxF8R | 3 | TAAATGCA/TGCATTTA | 7 |
| --- | | | | | | | | 0 |
| chr18 | 34965962 | ATTAATTTGTGGA-TGTGGTTA-AATGCTATCTTAG | loxF8L | 2 | loxF8R | 4 | TAACCACA/TGTGGTTA | 6 |
| chr2 | 113422724 | ACACATCTGTGGT-TGTGGTTA-TACAGATATATAT | loxF8L | 3 | loxF8L | 4 | TAACCACA/TGTGGTTA | 7 |
| chr9 | 68226856 | ACACATCTGTGGT-TGTGGTTA-TACAGATATATAT | loxF8L | 3 | loxF8L | 4 | TAACCACA/TGTGGTTA | 7 |
| chr9 | 194101 | ATATATATCTGTA-TAACCACA-ACCACAGATGTGT | loxF8L | 4 | loxF8L | 3 | TAACCACA/TGTGGTTA | 7 |
| chr9 | 41204413 | ATATATATCTGTA-TAACCACA-ACCACAGATGTGT | loxF8L | 4 | loxF8L | 3 | TAACCACA/TGTGGTTA | 7 |
| chr9 | 65660802 | ACACATCTGTGGT-TGTGGTTA-TACAGATATATAT | loxF8L | 3 | loxF8L | 4 | TAACCACA/TGTGGTTA | 7 |
| --- | | | | | | | | 0 |
| chr5 | 114368022 | TGAACTGTGTGGA-TCCACTTA-TACACAGATTTTT | loxF8L | 4 | loxF8L | 2 | TAAGTGGA/TCCACTTA | 6 |
| chr1 | 71434062 | AGAATTATGTGGG-TCCACTTA-TACATAGATTTTT | loxF8L | 4 | loxF8L | 3 | TAAGTGGA/TCCACTTA | 7 |
| chr9 | 77063059 | TGAAGAATGTGGG-TCCACTTA-TACAAAGATTTTT | loxF8R | 4 | loxF8L | 3 | TAAGTGGA/TCCACTTA | 7 |
| chr2 | 62807116 | AAAAATTTGTGTA-TAAGTGGA-TCCACACAGTTCA | loxF8L | 3 | loxF8L | 4 | TAAGTGGA/TCCACTTA | 7 |
| chr7 | 135091464 | AAATATCTGTGTA-TAAGTGGA-TCCACACAGTTCA | loxF8L | 3 | loxF8L | 4 | TAAGTGGA/TCCACTTA | 7 |
| chr14 | 84339381 | TTAACTATGTGGA-TCCACTTA-TAGACAGATTCTT | loxF8L | 3 | loxF8L | 4 | TAAGTGGA/TCCACTTA | 7 |
| chr2 | 17703548 | TGAAATATGTGGG-TCCACTTA-TACACAGATTTTC | loxF8L | 4 | loxF8L | 3 | TAAGTGGA/TCCACTTA | 7 |
| --- | | | | | | | | 0 |
| chr15 | 70898885 | ATATATATGTGGA-TATTTGTA-TACCCACATATAT | loxF8L | 2 | loxF8L | 4 | TACAAATA/TATTTGTA | 6 |
| chr11 | 64276243 | ATATATATGTAAA-TACAAATA-TACATATATTTAT | loxF8L | 4 | loxF8L | 3 | TACAAATA/TATTTGTA | 7 |
| chr10 | 85397374 | GTAAGACTGTCTT-TATTTGTA-TGCACAATCTTAT | loxF8R | 4 | loxF8R | 3 | TACAAATA/TATTTGTA | 7 |
| --- | | | | | | | | 0 |
| chr2 | 178603687 | ATATATCTGTGCA-TGTATGTA-TACACACATAAAT | loxF8L | 2 | loxF8L | 4 | TACATACA/TGTATGTA | 6 |
| chr1 | 24523039 | CTGATATTCTGTG-TGTATGTA-CACACACTTTTTT | loxF8R | 3 | loxF8R | 4 | TACATACA/TGTATGTA | 7 |

| | | | | | | | | |
|---|---|---|---|---|---|---|---|---|
| chr1 | 101943710 | AAAAATGTATAGA-TGTATGTA-TCTATATATTTAT | loxF8L | 4 | loxF8L | 3 | TACATACA/TGTATGTA | 7 |
| chr4 | 10930920 | ATAGATATGTGTA-TGTATGTA-TACATAGATATGT | loxF8L | 3 | loxF8L | 4 | TACATACA/TGTATGTA | 7 |
| chr12 | 76116236 | ACATATTTGTGGA-TGTATGTA-CACACACGCAGAG | loxF8L | 3 | loxF8R | 4 | TACATACA/TGTATGTA | 7 |
| chr13 | 22245750 | AGGTATCTGTGAA-TGTATGTA-TCTACATATGTAT | loxF8L | 4 | loxF8L | 3 | TACATACA/TGTATGTA | 7 |
| chr14 | 36823420 | ATACATATATGTA-TGTATGTA-TACACATATTTAA | loxF8L | 4 | loxF8L | 3 | TACATACA/TGTATGTA | 7 |
| chrX | 39104187 | ATACATATGTACA-TACATACA-TACATAGATTGAT | loxF8L | 4 | loxF8L | 3 | TACATACA/TGTATGTA | 7 |
| chr9 | 112418724 | ATACATATGTATA-TACATACA-TGCACACATATAT | loxF8L | 4 | loxF8L | 3 | TACATACA/TGTATGTA | 7 |
| chr10 | 50957204 | ATACATGTGTGTA-TACATACA-CACACACACAGAG | loxF8L | 3 | loxF8R | 4 | TACATACA/TGTATGTA | 7 |
| chr18 | 31115814 | ATAGATATGTAGA-TACATACA-TACACAAATAGAT | loxF8L | 3 | loxF8L | 4 | TACATACA/TGTATGTA | 7 |
| chrX | 14021939 | ATATATATGTATA-TGTATGTA-TACACATATATAT | loxF8L | 4 | loxF8L | 3 | TACATACA/TGTATGTA | 7 |
| chr13 | 38609226 | ATATATATGTCTA-TACATACA-TACACACATTTAC | loxF8L | 4 | loxF8L | 3 | TACATACA/TGTATGTA | 7 |
| chr7 | 48825981 | ATATATATGTGTA-TGTATGTA-TACATACATATAT | loxF8L | 3 | loxF8L | 4 | TACATACA/TGTATGTA | 7 |
| chr15 | 51704216 | ATATATGTGTGTA-TACATACA-TACATACATATAT | loxF8L | 3 | loxF8L | 4 | TACATACA/TGTATGTA | 7 |
| chr4 | 167780621 | ATATATGTGTGTA-TACATACA-TACATACATATAT | loxF8L | 3 | loxF8L | 4 | TACATACA/TGTATGTA | 7 |
| chr5 | 130135171 | ATATATGTGTGTA-TACATACA-TACATACATATAT | loxF8L | 3 | loxF8L | 4 | TACATACA/TGTATGTA | 7 |
| chr5 | 96665857 | CTAGATTTATGTG-TGTATGTA-TCTACAGATATAA | loxF8R | 4 | loxF8L | 3 | TACATACA/TGTATGTA | 7 |
| chr16 | 6904163 | CTGTGTCTGTGTG-TGTATGTA-AACACCATCTTAA | loxF8R | 4 | loxF8R | 3 | TACATACA/TGTATGTA | 7 |
| --- | | | | | | | | 0 |
| chr4 | 131056530 | AGAGATATGTAGA-TATAGGTA-TACACAGATATAT | loxF8L | 4 | loxF8L | 2 | TACCTATA/TATAGGTA | 6 |
| chr13 | 63025366 | ATATATATGTGCA-TATAGGTA-TACATATATTTAC | loxF8L | 3 | loxF8L | 4 | TACCTATA/TATAGGTA | 7 |
| chr4 | 109202772 | ATATATGTGTGTA-TACCTATA-TCTATAGAGATAT | loxF8L | 3 | loxF8L | 4 | TACCTATA/TATAGGTA | 7 |
| --- | | | | | | | | 0 |
| chr10 | 87539175 | TTATGATTCTGTA-TATTTCTA-TCCAGACATTTAT | loxF8R | 4 | loxF8L | 2 | TAGAAATA/TATTTCTA | 6 |
| chr14 | 27949482 | ATAAATCTATATA-TAGAAATA-TATATAAATTTAT | loxF8L | 3 | loxF8L | 4 | TAGAAATA/TATTTCTA | 7 |
| --- | | | | | | | | 0 |
| chr13 | 28633587 | AAATATATGTGTA-TATATAAA-TACACATATTTAT | loxF8L | 4 | loxF8L | 2 | TATATAAA/TTTATATA | 6 |
| chr6 | 74808473 | ATATATATGTGGA-TATATAAA-CACACACACATAA | loxF8L | 2 | loxF8R | 4 | TATATAAA/TTTATATA | 6 |
| chrX | 79884516 | ATATATGTGTGGA-TTTATATA-TACACATATATAC | loxF8L | 2 | loxF8L | 4 | TATATAAA/TTTATATA | 6 |
| chr4 | 136451064 | AAATATATGTGTA-TATATAAA-TACACACATATAT | loxF8L | 4 | loxF8L | 3 | TATATAAA/TTTATATA | 7 |
| chr6 | 160723032 | ACATATGTGTGCA-TATATAAA-TACACACATATAT | loxF8L | 4 | loxF8L | 3 | TATATAAA/TTTATATA | 7 |
| chr14 | 59749702 | ATATATCTATATA-TTTATATA-TCTATATATTTAT | loxF8L | 4 | loxF8L | 3 | TATATAAA/TTTATATA | 7 |
| chr6 | 166288844 | ATAAATATATAGA-TATATAAA-TATATAGATATAT | loxF8L | 3 | loxF8L | 4 | TATATAAA/TTTATATA | 7 |
| chr7 | 136463805 | ATAAATATATATA-TTTATATA-TACATAAATTTAT | loxF8L | 4 | loxF8L | 3 | TATATAAA/TTTATATA | 7 |
| chr5 | 164329166 | ATAAATATATGTA-TATATAAA-TATATATATTTAT | loxF8L | 3 | loxF8L | 4 | TATATAAA/TTTATATA | 7 |
| chrX | 141891710 | ATAAATATATATA-TTTATATA-TGCATATATTTAT | loxF8L | 4 | loxF8L | 3 | TATATAAA/TTTATATA | 7 |
| chr2 | 208590732 | ATATAACTATGTA-TTTATATA-TACAGATATTTAT | loxF8L | 4 | loxF8L | 3 | TATATAAA/TTTATATA | 7 |
| chr3 | 36879777 | ATAAATATGTAAA-TATATAAA-TATACAAATATAT | loxF8L | 3 | loxF8L | 4 | TATATAAA/TTTATATA | 7 |
| chr21 | 21595693 | ATAAATGTGTACA-TTTATATA-TCCACAGAGGGAA | loxF8L | 3 | loxF8L | 4 | TATATAAA/TTTATATA | 7 |
| chr15 | 62976266 | ATACATGTGTGTA-TATATAAA-TCTACACGTGTAT | loxF8L | 3 | loxF8L | 4 | TATATAAA/TTTATATA | 7 |
| chr20 | 12105660 | ATACATATATGTA-TATATAAA-TCCATACATATAT | loxF8L | 4 | loxF8L | 3 | TATATAAA/TTTATATA | 7 |
| chr5 | 130417451 | ATATATATGTATA-TATATAAA-TACACATATATAT | loxF8L | 4 | loxF8L | 3 | TATATAAA/TTTATATA | 7 |
| chr4 | 16766383 | ATATATGTGTGCA-TTTATATA-TATACACATATAT | loxF8L | 3 | loxF8L | 4 | TATATAAA/TTTATATA | 7 |
| chr13 | 55037079 | ATATATGTGTGTG-TTTATATA-TACACACATATAT | loxF8L | 4 | loxF8L | 3 | TATATAAA/TTTATATA | 7 |
| chr3 | 84316708 | ATATATGTGTGTA-TATATAAA-CACACAGATACAT | loxF8L | 3 | loxF8L | 4 | TATATAAA/TTTATATA | 7 |
| chr7 | 12709508 | CTAATATTTTGTC-TTTATATA-TTCACACATTAAC | loxF8R | 3 | loxF8L | 4 | TATATAAA/TTTATATA | 7 |
| --- | | | | | | | | 0 |
| chr18 | 55078062 | ATAAATGTGTGTA-TCTATATA-TACACACACATAT | loxF8L | 2 | loxF8L | 4 | TATATAGA/TCTATATA | 6 |
| chr3 | 83845678 | CTATATCTATAGA-TCTATATA-TCTATATATTTAT | loxF8L | 4 | loxF8L | 3 | TATATAGA/TCTATATA | 7 |
| chr4 | 185224592 | ATAAATATGTAAA-TATATAGA-TATATAAATTTAT | loxF8L | 3 | loxF8L | 4 | TATATAGA/TCTATATA | 7 |
| chr9 | 22061839 | ATAAATCTATATA-TCTATATA-TACACTTATATAT | loxF8L | 3 | loxF8L | 4 | TATATAGA/TCTATATA | 7 |
| chr7 | 152466027 | ATAAATGTATAGA-TATATAGA-TCTATAGAGATAT | loxF8L | 3 | loxF8L | 4 | TATATAGA/TCTATATA | 7 |
| chr12 | 110432140 | ATACATATCTAGA-TATATAGA-TCGATAGATCTAT | loxF8L | 4 | loxF8L | 3 | TATATAGA/TCTATATA | 7 |
| chr13 | 79801258 | ATATATGTGTGTG-TCTATATA-TCTACACATGTAT | loxF8L | 4 | loxF8L | 3 | TATATAGA/TCTATATA | 7 |
| chr3 | 63550330 | ATACGTGTGTGTA-TATATAGA-TACACAAATATAT | loxF8L | 4 | loxF8L | 3 | TATATAGA/TCTATATA | 7 |
| chr5 | 100593917 | ATATATCTGTATA-TCTATATA-TCTATATATCTAT | loxF8L | 3 | loxF8L | 4 | TATATAGA/TCTATATA | 7 |
| chr7 | 47111707 | ATATATATGTATA-TCTATATA-TCTATAGATATAT | loxF8L | 4 | loxF8L | 3 | TATATAGA/TCTATATA | 7 |
| chr7 | 47111651 | ATATATATGTATA-TCTATATA-TCTATAGATATAT | loxF8L | 4 | loxF8L | 3 | TATATAGA/TCTATATA | 7 |
| chr7 | 47111623 | ATATATATGTATA-TCTATATA-TCTATAGATATAT | loxF8L | 4 | loxF8L | 3 | TATATAGA/TCTATATA | 7 |
| chr7 | 47111679 | ATATATATGTATA-TCTATATA-TCTATAGATATAT | loxF8L | 4 | loxF8L | 3 | TATATAGA/TCTATATA | 7 |
| chr7 | 47111567 | ATATATATGTATA-TCTATATA-TCTATAGATATAT | loxF8L | 4 | loxF8L | 3 | TATATAGA/TCTATATA | 7 |
| chr7 | 47111595 | ATATATATGTATA-TCTATATA-TCTATAGATATAT | loxF8L | 4 | loxF8L | 3 | TATATAGA/TCTATATA | 7 |
| chr13 | 32842009 | ATATATATGTGTA-TCTATATA-CACACACACATAC | loxF8L | 3 | loxF8R | 4 | TATATAGA/TCTATATA | 7 |
| chr4 | 60845385 | ATATATGTGTGTA-TATATAGA-TGAACATATATAT | loxF8L | 3 | loxF8L | 4 | TATATAGA/TCTATATA | 7 |
| chr7 | 47111539 | GTATATGTATGGA-TCTATATA-TCTATAGATATAT | loxF8L | 4 | loxF8L | 3 | TATATAGA/TCTATATA | 7 |
| chr6 | 49041762 | ATATATGTGTGTA-TATATAGA-CACACACATATAT | loxF8L | 3 | loxF8L | 4 | TATATAGA/TCTATATA | 7 |
| chr9 | 102011565 | ATATGTGTGTGTA-TCTATATA-TACACACATATAT | loxF8L | 4 | loxF8L | 3 | TATATAGA/TCTATATA | 7 |
| chr2 | 83452205 | CTCAGATTATATA-TCTATATA-TACACACATATAT | loxF8R | 4 | loxF8L | 3 | TATATAGA/TCTATATA | 7 |
| chr2 | 50354504 | ATATGTGTGTGTA-TCTATATA-TACACTACCTTAG | loxF8L | 4 | loxF8R | 3 | TATATAGA/TCTATATA | 7 |
| --- | | | | | | | | 0 |
| chr2 | 187142382 | ATATATATGTATA-TATCCATA-AACACAGATTTAT | loxF8L | 4 | loxF8L | 2 | TATCCATA/TATGGATA | 6 |
| chr5 | 127482187 | ATATATATGTGTA-TATCCATA-TATACATATATAT | loxF8L | 3 | loxF8L | 4 | TATCCATA/TATGGATA | 7 |
| --- | | | | | | | | 0 |
| chr7 | 15145453 | ATAAATATGTAGA-TATGTAGA-TACATAAGCTTAT | loxF8L | 2 | loxF8R | 4 | TATGTAGA/TCTACATA | 6 |
| chr14 | 85923701 | ATACAGCTGTGTA-TATGTAGA-AACACAAAGATAG | loxF8L | 3 | loxF8R | 4 | TATGTAGA/TCTACATA | 7 |
| chr9 | 120100028 | GTAGATATGTGTA-TATGTAGA-TACACATATGTAT | loxF8L | 4 | loxF8L | 3 | TATGTAGA/TCTACATA | 7 |
| --- | | | | | | | | 0 |
| chr7 | 119191493 | ATAAATGTGTGTA-TATGTGCA-CACACACATATAT | loxF8L | 2 | loxF8L | 4 | TATGTGCA/TGCACATA | 6 |
| chr3 | 57212767 | TTAAATAGGAGGA-TGCACATA-TGCACATATTTTT | loxF8L | 4 | loxF8L | 3 | TATGTGCA/TGCACATA | 7 |
| chr6 | 18031906 | ATAAATAAGTGTA-TATGTGCA-TAAAAAGATTTGT | loxF8L | 3 | loxF8L | 4 | TATGTGCA/TGCACATA | 7 |
| chr3 | 80645930 | ATAAACATGTGTA-TATGTGCA-TATACACATATAT | loxF8L | 3 | loxF8L | 4 | TATGTGCA/TGCACATA | 7 |
| chr4 | 46763800 | ATATACATGTGCA-TGCACATA-CACACATACTTTG | loxF8L | 4 | loxF8R | 3 | TATGTGCA/TGCACATA | 7 |
| --- | | | | | | | | 0 |
| chr20 | 22057454 | ATAAATATATGGA-TGGATGGA-TCTATAGATAGAT | loxF8L | 2 | loxF8L | 4 | TCCATCCA/TGGATGGA | 6 |
| chr10 | 123272868 | ATAAATCTGTCCA-TCCATCCA-TCCAAACAGCTAT | loxF8L | 2 | loxF8L | 4 | TCCATCCA/TGGATGGA | 6 |
| chrX | 1525763 | ATCCATCTGTCCA-TCCATCCA-TCCATCCATTTAT | loxF8L | 4 | loxF8L | 3 | TCCATCCA/TGGATGGA | 7 |
| chrY | 1525763 | ATCCATCTGTCCA-TCCATCCA-TCCATCCATTTAT | loxF8L | 4 | loxF8L | 3 | TCCATCCA/TGGATGGA | 7 |
| chr1 | 163339136 | ATAGGTATGTGTG-TGGATGGA-TCCATAGATTTGA | loxF8R | 4 | loxF8L | 3 | TCCATCCA/TGGATGGA | 7 |

| | | | | | | | | |
|---|---|---|---|---|---|---|---|---|
| chr12 | 113439223 | ATCCATCAGTGTA-TCCATCCA-TCCACATATCCAT | loxF8L | 4 | loxF8L | 3 | TCCATCCA/TGGATGGA | 7 |
| chr22 | 35438504 | ATCCATCTATGCA-TCCATCCA-TCTACATATTCAT | loxF8L | 4 | loxF8L | 3 | TCCATCCA/TGGATGGA | 7 |
| chrX | 39533617 | ATCCATCTCTGCA-TCCATCCA-TCCACCTATTCAT | loxF8L | 4 | loxF8L | 3 | TCCATCCA/TGGATGGA | 7 |
| chr1 | 12583667 | ATTCATCTGTCCA-TCCATCCA-TCCACCCATTCAT | loxF8L | 4 | loxF8L | 3 | TCCATCCA/TGGATGGA | 7 |
| --- | | | | | | | | 0 |
| chr9 | 9363552 | CTGAGATTGGCTC-TCTGTTCA-AACAAAATCTTAG | loxF8R | 4 | loxF8R | 2 | TCTGTTCA/TGAACAGA | 6 |
| chr1 | 222095389 | CTACACTTGTGTG-TGAACAGA-CACACACACATAT | loxF8R | 3 | loxF8R | 4 | TCTGTTCA/TGAACAGA | 7 |
| --- | | | | | | | | 0 |
| chr18 | 41169127 | CAAACATAATGTG-TGAATACA-TGCACAAATTTAT | loxF8R | 4 | loxF8L | 2 | TGAATACA/TGTATTCA | 6 |
| chr14 | 32571679 | ATTACTCTGAGGA-TGAATACA-CACACATTCACAT | loxF8L | 3 | loxF8R | 4 | TGAATACA/TGTATTCA | 7 |
| --- | | | | | | | | 0 |
| chr2 | 59450258 | ATAAATCTGTGCT-TTCAGTCA-TACACAAGTATAG | loxF8L | 2 | loxF8R | 4 | TGACTGAA/TTCAGTCA | 6 |
| chr7 | 126558732 | TTCAACCTGTGGT-TGACTGAA-TCCACAGATGTGG | loxF8L | 4 | loxF8L | 3 | TGACTGAA/TTCAGTCA | 7 |
| --- | | | | | | | | 0 |
| chr11 | 112171060 | CTGAATCTGTGGA-TTCAATCA-ACCACACATTGAA | loxF8L | 2 | loxF8L | 4 | TGATTGAA/TTCAATCA | 6 |
| chr11 | 110326355 | ATAAAAGTGTCTA-TTCAATCA-CACACAAACAAAG | loxF8L | 4 | loxF8R | 3 | TGATTGAA/TTCAATCA | 7 |
| chr3 | 89326716 | ATTGATCTGTGGT-TGATTGAA-TCCATAGATGCAG | loxF8L | 3 | loxF8L | 4 | TGATTGAA/TTCAATCA | 7 |
| chr7 | 93144292 | CCACATCTGTGGG-TTCAATCA-ACCACAGATTGAA | loxF8L | 4 | loxF8L | 3 | TGATTGAA/TTCAATCA | 7 |
| chr15 | 70955136 | TTAAATCTGAGGT-TGATTGAA-TTCATAGATGTAG | loxF8L | 3 | loxF8L | 4 | TGATTGAA/TTCAATCA | 7 |
| chr14 | 38474471 | TTCAATCTGTGGT-TGATTGAA-TCCTCAGATGCAG | loxF8L | 3 | loxF8L | 4 | TGATTGAA/TTCAATCA | 7 |
| chr12 | 49591971 | TTTGATCTGTGGT-TGATTGAA-TCCACAGATGCAG | loxF8L | 4 | loxF8L | 3 | TGATTGAA/TTCAATCA | 7 |
| --- | | | | | | | | 0 |
| chr22 | 36027699 | CTCACACTCTGTG-TGATTTAA-CCCACAATCTCAG | loxF8R | 4 | loxF8R | 2 | TGATTTAA/TTAAATCA | 6 |
| chr11 | 102800713 | AGAAAGCTGGGGA-TTAAATCA-TACCAAATCCTAG | loxF8L | 3 | loxF8R | 4 | TGATTTAA/TTAAATCA | 7 |
| chr8 | 121616331 | CTCAGATTTTTTT-TTAAATCA-ACCACATATTGAT | loxF8R | 4 | loxF8L | 3 | TGATTTAA/TTAAATCA | 7 |
| --- | | | | | | | | 0 |
| chr9 | 25807277 | CAAAGATTGAGTG-TTCATGCA-TCAAACGATTTTT | loxF8R | 2 | loxF8L | 4 | TGCATGAA/TTCATGCA | 6 |
| chr16 | 65865811 | ATACATATGTGTA-TTCATGCA-TCTATATATTTAC | loxF8L | 3 | loxF8L | 4 | TGCATGAA/TTCATGCA | 7 |
| --- | | | | | | | | 0 |
| chr9 | 88441046 | TTACATCTGTGGA-TTCATCCA-AACACAGATTGAA | loxF8L | 2 | loxF8L | 4 | TGGATGAA/TTCATCCA | 6 |
| chr5 | 60365554 | ATAAATAGATGGA-TGGATGAA-TCAATAGATAGAT | loxF8L | 3 | loxF8L | 4 | TGGATGAA/TTCATCCA | 7 |
| --- | | | | | | | | 0 |
| chr14 | 59784513 | TTTAGTTTGTATG-TGGTCAAA-TCTACAGATTTTT | loxF8R | 4 | loxF8L | 2 | TGGTCAAA/TTTGACCA | 6 |
| chr16 | 25338039 | CCACATCTGTGGA-TTTGACCA-ACCACAGATGGAA | loxF8L | 3 | loxF8L | 4 | TGGTCAAA/TTTGACCA | 7 |
| --- | | | | | | | | 0 |
| chr5 | 29265550 | CCAAATCTGTGGA-TTCCACCA-ACCACAGATGGAA | loxF8L | 2 | loxF8L | 4 | TGGTGGAA/TTCCACCA | 6 |
| chr15 | 101242080 | TTCAATCCATGGA-TGGTGGAA-TCCACAGATGTAG | loxF8L | 4 | loxF8L | 2 | TGGTGGAA/TTCCACCA | 6 |
| chr7 | 91812641 | CTAACAATTTCTG-TTCCACCA-TACACACTTTTAT | loxF8R | 4 | loxF8L | 3 | TGGTGGAA/TTCCACCA | 7 |
| --- | | | | | | | | 0 |
| chr2 | 205289707 | ATAAATTAGTGGA-TTAAACCA-TTCATGGATTGAT | loxF8L | 2 | loxF8L | 4 | TGGTTTAA/TTAAACCA | 6 |
| chr3 | 145994473 | CCACATCTGTAGA-TTAAACCA-ACCACAGATTAAA | loxF8L | 4 | loxF8L | 3 | TGGTTTAA/TTAAACCA | 7 |
| chrX | 129261651 | TTTAATCTGTGGT-TGGTTTAA-TCCACAAATGCAG | loxF8L | 3 | loxF8L | 4 | TGGTTTAA/TTAAACCA | 7 |
| --- | | | | | | | | 0 |
| chr14 | 21599907 | ATAAATCACTGGA-TTAAAGAA-GCTAAAGTTTTAT | loxF8L | 2 | loxF8L | 4 | TTAAAGAA/TTCTTTAA | 6 |
| chr5 | 7701257 | ATAACACATTGGA-TTAAAGAA-TCCACCCATTTAC | loxF8L | 4 | loxF8L | 3 | TTAAAGAA/TTCTTTAA | 7 |
| --- | | | | | | | | 0 |
| chr1 | 116970633 | CTAAGGTTTTATT-TTATTTAA-TCCACAGAGTTGT | loxF8R | 4 | loxF8L | 2 | TTAAATAA/TTATTTAA | 6 |
| chr1 | 170018589 | ATTAATCCTTTGA-TTATTTAA-TCAAAGGATTTAT | loxF8L | 4 | loxF8L | 3 | TTAAATAA/TTATTTAA | 7 |
| --- | | | | | | | | 0 |
| chr4 | 28278636 | CTTAAATACTGTG-TTAGAAAA-TCCACTGATTTTT | loxF8R | 4 | loxF8L | 2 | TTAGAAAA/TTTTCTAA | 6 |
| chr13 | 105268338 | ATAAAACTGAGGT-TTAGAAAA-CAAATAATTTTAC | loxF8L | 3 | loxF8R | 4 | TTAGAAAA/TTTTCTAA | 7 |
| chr5 | 42429421 | CTCCTAATGTGTG-TTTTCTAA-ACAAAAGATTTAT | loxF8R | 4 | loxF8L | 3 | TTAGAAAA/TTTTCTAA | 7 |
| --- | | | | | | | | 0 |
| chr12 | 73493873 | GTAAAATTGTCTA-TTTGTGAA-CACATAATCTTAC | loxF8R | 4 | loxF8R | 2 | TTCACAAA/TTTGTGAA | 6 |
| chr14 | 35644323 | ATGCATCTGTTCA-TTTGTGAA-AAAACAATCATAG | loxF8L | 4 | loxF8R | 3 | TTCACAAA/TTTGTGAA | 7 |
| --- | | | | | | | | 0 |
| chr1 | 190455414 | GTAAAACTGTGGA-TTTAAGAA-TACATAAATATAT | loxF8L | 2 | loxF8L | 4 | TTCTTAAA/TTTAAGAA | 6 |
| chr12 | 69568099 | CTCAGTTGGAGTG-TTCTTAAA-CACTCAATGTCAG | loxF8R | 4 | loxF8R | 3 | TTCTTAAA/TTTAAGAA | 7 |
| --- | | | | | | | | 0 |
| chr14 | 48544432 | ATAATCCTTTGGG-TTTATAAA-TCCAAAGGTTTAT | loxF8L | 4 | loxF8L | 2 | TTTATAAA/TTTATAAA | 6 |
| chr4 | 12080313 | TTAAATCTGTTCA-TTTATAAA-ACCCCAAATTTTT | loxF8L | 3 | loxF8L | 4 | TTTATAAA/TTTATAAA | 7 |
| --- | | | | | | | | 0 |
| chr2 | 21158851 | ATATATATGTGGT-CTTTTTTT-CACAAATTTTTAT | loxF8L | 3 | loxF8R | 4 | AAAAAAAG/CTTTTTTT | 7 |
| chr12 | 39303548 | ATAAACCTGTGCC-CTTTTTTT-CACACTACTTTTG | loxF8L | 3 | loxF8R | 4 | AAAAAAAG/CTTTTTTT | 7 |
| --- | | | | | | | | 0 |
| chr2 | 80936010 | AAAAATATTTGGA-AAAAAAAT-TCTACAAAATTCT | loxF8L | 3 | loxF8L | 4 | AAAAAAAT/ATTTTTTT | 7 |
| chr12 | 63864877 | AAACATCTGTGAA-AAAAAAAT-TCCACTCATATAA | loxF8L | 3 | loxF8L | 4 | AAAAAAAT/ATTTTTTT | 7 |
| chr2 | 135636446 | TATAGTTTGTGTG-ATTTTTTT-TCCCCAGACTTGT | loxF8R | 4 | loxF8L | 3 | AAAAAAAT/ATTTTTTT | 7 |
| chr1 | 31023487 | AGAAAGATGTGGG-AAAAAAAT-TTAACAGATTTAG | loxF8L | 4 | loxF8L | 3 | AAAAAAAT/ATTTTTTT | 7 |
| chr15 | 96611629 | ATAAACCTTTAGT-ATTTTTTT-TCCAAAGGTTTAA | loxF8L | 4 | loxF8L | 3 | AAAAAAAT/ATTTTTTT | 7 |
| chr10 | 63960376 | ATATAACAGTGGG-AAAAAAAT-AATACATTCTTAG | loxF8L | 4 | loxF8R | 3 | AAAAAAAT/ATTTTTTT | 7 |
| --- | | | | | | | | 0 |
| chr5 | 130364546 | ATATATTTTTGTA-TCTTTTTT-TCAACATGTTTAT | loxF8L | 4 | loxF8L | 3 | AAAAAAGA/TCTTTTTT | 7 |
| chr10 | 94651378 | CTGAAATTGTGGG-AAAAAAGA-CACAAGCTCTTAT | loxF8R | 3 | loxF8R | 4 | AAAAAAGA/TCTTTTTT | 7 |
| --- | | | | | | | | 0 |
| chr10 | 110752601 | ATAAATATGTACT-TATTTTTT-TCCAATGATTTTT | loxF8L | 4 | loxF8L | 3 | AAAAAATA/TATTTTTT | 7 |
| chr1 | 101053013 | ATAAATCTCTTTA-AAAAAATA-CACACTCTCATAT | loxF8L | 3 | loxF8R | 4 | AAAAAATA/TATTTTTT | 7 |
| chr3 | 101096334 | ATCATTCGGGGGA-AAAAAATA-CACGCAATCACAG | loxF8L | 4 | loxF8R | 3 | AAAAAATA/TATTTTTT | 7 |
| chr1 | 96754893 | TAAAATCTCTAGA-AAAAAATA-CAGAAAATCTTCG | loxF8L | 4 | loxF8R | 3 | AAAAAATA/TATTTTTT | 7 |
| --- | | | | | | | | 0 |
| chr5 | 115091939 | ACAAATATGGGGA-AAAAAATG-CACCTAAACTTGG | loxF8L | 3 | loxF8R | 4 | AAAAAATG/CATTTTTT | 7 |
| chr2 | 153592622 | ATAAATGGGTGCT-AAAAAATG-TACACACATTGAT | loxF8L | 4 | loxF8L | 3 | AAAAAATG/CATTTTTT | 7 |
| --- | | | | | | | | 0 |
| chr2 | 141089649 | AGGAATCTGGGTA-AAAAACAC-ACCTCAGAATTAT | loxF8L | 4 | loxF8L | 3 | AAAAACAC/GTGTTTTT | 7 |
| chr3 | 1040392 | ATAAGTTTGTTGA-GTGTTTTT-ACCATGGAATTAT | loxF8L | 3 | loxF8L | 4 | AAAAACAC/GTGTTTTT | 7 |
| chr6 | 146235528 | GTTAGGTTGTTTG-GTGTTTTT-TTCACAGATTTTG | loxF8R | 4 | loxF8L | 3 | AAAAACAC/GTGTTTTT | 7 |
| --- | | | | | | | | 0 |

| | | | | | | | | |
|---|---|---|---|---|---|---|---|---|
| chr7 | 114901404 | AGAAATTTATAGA-AAAAACTA-CACAAAATATCAG | loxF8L | 4 | loxF8R | 3 | AAAAACTA/TAGTTTTT | 7 |
| chr12 | 67062845 | CTATGTTTCAGTG-TAGTTTTT-CACAAAACATTAG | loxF8R | 4 | loxF8R | 3 | AAAAACTA/TAGTTTTT | 7 |
| --- | | | | | | | | 0 |
| chr2 | 183182319 | ATAAATCTTTCAC-CTCTTTTT-TCCCTAGGTTTAT | loxF8L | 4 | loxF8L | 3 | AAAAAGAG/CTCTTTTT | 7 |
| chr14 | 30806261 | ATAATTTTGTATG-CTCTTTTT-TCCACATATACAT | loxF8R | 4 | loxF8L | 3 | AAAAAGAG/CTCTTTTT | 7 |
| --- | | | | | | | | 0 |
| chr11 | 115292955 | ATAGATCTTTTGA-AAAAAGAT-GCCACATGTTTAC | loxF8L | 3 | loxF8L | 4 | AAAAAGAT/ATCTTTTT | 7 |
| chr9 | 27803699 | TTAAATCAGTGGG-ATCTTTTT-TCCTCAGATCAAA | loxF8L | 3 | loxF8L | 4 | AAAAAGAT/ATCTTTTT | 7 |
| --- | | | | | | | | 0 |
| chr6 | 13623503 | ATAAACGTGTGCA-AAAAAGGG-CACATAGATTGAT | loxF8L | 3 | loxF8L | 4 | AAAAAGGG/CCCTTTTT | 7 |
| chr18 | 13549546 | TTAACATTGTCTG-AAAAAGGG-AACTGAATCTGAG | loxF8R | 3 | loxF8R | 4 | AAAAAGGG/CCCTTTTT | 7 |
| --- | | | | | | | | 0 |
| chr4 | 88563001 | ATAAATGTCAGGA-TTATTTTT-TCTACATATGTAA | loxF8L | 3 | loxF8L | 4 | AAAAATAA/TTATTTTT | 7 |
| chr2 | 111059456 | CTCAGATTGGATG-AAAAATAA-CACCCAACCATAC | loxF8R | 3 | loxF8R | 4 | AAAAATAA/TTATTTTT | 7 |
| --- | | | | | | | | 0 |
| chr1 | 78786223 | AACAAACTATGGA-AAAAATAG-CACACAAAATAAG | loxF8L | 4 | loxF8R | 3 | AAAAATAG/CTATTTTT | 7 |
| chr18 | 41914273 | ATAAATTTGGGGT-CTATTTTT-TACAACAGCTTAG | loxF8L | 3 | loxF8R | 4 | AAAAATAG/CTATTTTT | 7 |
| --- | | | | | | | | 0 |
| chr3 | 127348752 | AAAAATCGGTAGA-AAAAATAT-TCCACCTAATTAA | loxF8L | 3 | loxF8L | 4 | AAAAATAT/ATATTTTT | 7 |
| chr6 | 49346548 | AAAAATCTGAGAA-AAAAATAT-TCCAGAGAGATAA | loxF8L | 3 | loxF8L | 4 | AAAAATAT/ATATTTTT | 7 |
| chr2 | 69005756 | ATCAAAATGTGGA-AAAAATAT-TCCAAATATGTCT | loxF8L | 3 | loxF8L | 4 | AAAAATAT/ATATTTTT | 7 |
| chr5 | 100419048 | ATAAATTTTAGGT-ATATTTTT-CACTCAATTTTAT | loxF8L | 4 | loxF8R | 3 | AAAAATAT/ATATTTTT | 7 |
| chr8 | 50811104 | ATATTTCTGTGGC-ATATTTTT-CCCATAGTTTTAC | loxF8L | 3 | loxF8L | 4 | AAAAATAT/ATATTTTT | 7 |
| chr6 | 8877626 | CTATGATTTTGTT-AAAAATAT-ACCACAGATATCC | loxF8R | 3 | loxF8L | 4 | AAAAATAT/ATATTTTT | 7 |
| --- | | | | | | | | 0 |
| chr1 | 81485372 | TTAAGATTTTGAG-TGATTTTT-TGAACATGTTTAT | loxF8R | 3 | loxF8L | 4 | AAAAATCA/TGATTTTT | 7 |
| chr12 | 63280952 | CCATAATTGTGGG-TGATTTTT-AACACATCCTTAG | loxF8R | 4 | loxF8R | 3 | AAAAATCA/TGATTTTT | 7 |
| --- | | | | | | | | 0 |
| chr3 | 73996843 | AGAAATATGTGCA-AAAAATTA-TGAACAGACTCAT | loxF8L | 3 | loxF8L | 4 | AAAAATTA/TAATTTTT | 7 |
| chr7 | 140371777 | CAAGGAATCTGTG-AAAAATTA-TCCTCAACTTTAT | loxF8R | 4 | loxF8L | 3 | AAAAATTA/TAATTTTT | 7 |
| --- | | | | | | | | 0 |
| chr20 | 48748432 | AAAAATCTATGCA-AAAACAAA-CACACACACATAC | loxF8L | 3 | loxF8R | 4 | AAAACAAA/TTTGTTTT | 7 |
| chr17 | 36218130 | CTGAAATTGTGAA-AAAACAAA-CACAAATTCTTAT | loxF8R | 4 | loxF8R | 3 | AAAACAAA/TTTGTTTT | 7 |
| chr17 | 31184796 | CTGAAATTGTGAA-AAAACAAA-CACAAACTCTTAT | loxF8R | 4 | loxF8R | 3 | AAAACAAA/TTTGTTTT | 7 |
| --- | | | | | | | | 0 |
| chr3 | 169435812 | ATAAATCAGTATA-AAAACAAG-CAGACAATAGCAG | loxF8L | 3 | loxF8R | 4 | AAAACAAG/CTTGTTTT | 7 |
| chrX | 35660470 | ATTAAGATGTTGA-AAAACAAG-TTCACAGATATAA | loxF8L | 4 | loxF8L | 3 | AAAACAAG/CTTGTTTT | 7 |
| --- | | | | | | | | 0 |
| chr14 | 82148544 | AGATATCAGTGGC-AAAACTAC-TACTGAGATTTAT | loxF8L | 4 | loxF8L | 3 | AAAACTAC/GTAGTTTT | 7 |
| chr21 | 21358675 | CTAGTATTACGTG-GTAGTTTT-AACACTGATTTAT | loxF8R | 4 | loxF8L | 3 | AAAACTAC/GTAGTTTT | 7 |
| --- | | | | | | | | 0 |
| chr2 | 216039118 | ATATATTTCAGGA-ATAGTTTT-TCCATAGATATAA | loxF8L | 4 | loxF8L | 3 | AAAACTAT/ATAGTTTT | 7 |
| chr7 | 47877824 | CTGAGACTCAGTG-ATAGTTTT-CACATACCCTTAG | loxF8R | 4 | loxF8R | 3 | AAAACTAT/ATAGTTTT | 7 |
| --- | | | | | | | | 0 |
| chr8 | 17220504 | ATAAATCCAAGGA-CCAGTTTT-TACCCAAATATAT | loxF8L | 3 | loxF8L | 4 | AAAACTGG/CCAGTTTT | 7 |
| chr1 | 105810182 | ATAAGAGCTTGTG-CCAGTTTT-TCCACAATTTTAG | loxF8R | 4 | loxF8L | 3 | AAAACTGG/CCAGTTTT | 7 |
| chr6 | 136323976 | TGAACAGTGTGTG-CCAGTTTT-TCCACAGAATTGC | loxF8R | 4 | loxF8L | 3 | AAAACTGG/CCAGTTTT | 7 |
| --- | | | | | | | | 0 |
| chr14 | 35779048 | ATACATCTGTAAA-AAAAGAAT-ACTAAATATTTAT | loxF8L | 3 | loxF8L | 4 | AAAAGAAT/ATTCTTTT | 7 |
| chr2 | 162304681 | ATGAATTTGTGAA-ATTCTTTT-TCAGCAGAAATAT | loxF8L | 3 | loxF8L | 4 | AAAAGAAT/ATTCTTTT | 7 |
| --- | | | | | | | | 0 |
| chrX | 96114035 | ATAAAACTAAGGA-AAAAGCTA-TCCACTAATCTTT | loxF8L | 3 | loxF8L | 4 | AAAAGCTA/TAGCTTTT | 7 |
| chr8 | 28545732 | TTAAGTTTTGGTG-TAGCTTTT-TACACAGTCATAG | loxF8R | 4 | loxF8R | 3 | AAAAGCTA/TAGCTTTT | 7 |
| --- | | | | | | | | 0 |
| chr12 | 13215472 | ACAATTCGGTGGT-AAAAGTAC-CACACAAACTATG | loxF8L | 4 | loxF8R | 3 | AAAAGTAC/GTACTTTT | 7 |
| chr15 | 50993981 | ATATATCTTTGTA-AAAAGTAC-TCCTCAAAATTGT | loxF8L | 3 | loxF8L | 4 | AAAAGTAC/GTACTTTT | 7 |
| --- | | | | | | | | 0 |
| chr1 | 198715388 | CAAAGTGTGTTTG-TTTATTTT-TCCTCACATTTTT | loxF8R | 4 | loxF8L | 3 | AAAATAAA/TTTATTTT | 7 |
| chr13 | 48500045 | AGAAATCAGTGGG-TTTATTTT-TATAAAATCTAAG | loxF8L | 3 | loxF8R | 4 | AAAATAAA/TTTATTTT | 7 |
| chr10 | 123301027 | ATAAAACTGTTTA-AAAATAAA-TCCACGGATACAG | loxF8L | 3 | loxF8L | 4 | AAAATAAA/TTTATTTT | 7 |
| chr8 | 15268726 | TTACAAATGTGTG-AAAATAAA-ACAACATATTTAT | loxF8R | 4 | loxF8L | 3 | AAAATAAA/TTTATTTT | 7 |
| chr3 | 196920201 | ATATATCTTTGCA-TTTATTTT-TTCTCAGTATTAT | loxF8L | 3 | loxF8L | 4 | AAAATAAA/TTTATTTT | 7 |
| chr2 | 170135223 | ATGAACCTGAGGA-AAAATAAA-TCCAGAAAGTAAT | loxF8L | 3 | loxF8L | 4 | AAAATAAA/TTTATTTT | 7 |
| chr13 | 103469012 | CTAAGAGTTTTTG-AAAATAAA-TATAAAATCATAG | loxF8R | 3 | loxF8R | 4 | AAAATAAA/TTTATTTT | 7 |
| --- | | | | | | | | 0 |
| chrX | 126098657 | ATAAGATCTGGTG-TCTATTTT-TCCACAATTTTAG | loxF8R | 4 | loxF8L | 3 | AAAATAGA/TCTATTTT | 7 |
| chr2 | 154981732 | ATGACATTGTGTA-TCTATTTT-CATACAGTCTTTG | loxF8R | 4 | loxF8R | 3 | AAAATAGA/TCTATTTT | 7 |
| --- | | | | | | | | 0 |
| chr9 | 103270388 | AAATATATGTGAA-TATATTTT-TGCATATATTTAT | loxF8L | 4 | loxF8L | 3 | AAAATATA/TATATTTT | 7 |
| chr1 | 86371753 | ATAAACTTTTAGA-TATATTTT-CACACATTCTTGT | loxF8L | 4 | loxF8R | 3 | AAAATATA/TATATTTT | 7 |
| chr1 | 56160578 | ATAAATCTAAAGA-AAAATATA-TGCATATACTTAT | loxF8L | 3 | loxF8L | 4 | AAAATATA/TATATTTT | 7 |
| chr5 | 121093773 | ATAAATCTGAAGT-TATATTTT-TCTCCAGATAGAT | loxF8L | 3 | loxF8L | 4 | AAAATATA/TATATTTT | 7 |
| chrX | 146048812 | ATATATCTGGGTA-TATATTTT-GAAAAAATCATAG | loxF8L | 3 | loxF8R | 4 | AAAATATA/TATATTTT | 7 |
| chr17 | 71607455 | TTAAATGTGTGTA-AAAATATA-AAGACATTTTTAG | loxF8L | 3 | loxF8R | 4 | AAAATATA/TATATTTT | 7 |
| --- | | | | | | | | 0 |
| chr1 | 105495964 | TCATATGTGTGGA-ATGATTTT-TCTCCAGGTTTAT | loxF8L | 4 | loxF8L | 3 | AAAATCAT/ATGATTTT | 7 |
| chr2 | 8142543 | CTTATAGTGGGTG-ATGATTTT-CATAGAATCATAG | loxF8R | 4 | loxF8R | 3 | AAAATCAT/ATGATTTT | 7 |
| --- | | | | | | | | 0 |
| chr14 | 92866700 | ATGAATCATTGGG-GCCATTTT-CCCCCACATTTAT | loxF8L | 4 | loxF8L | 3 | AAAATGGC/GCCATTTT | 7 |
| chr13 | 101282833 | ATAACTCTATGGT-AAAATGGC-CATATAAACATAG | loxF8L | 3 | loxF8R | 4 | AAAATGGC/GCCATTTT | 7 |
| chr12 | 99707065 | CTAAATTTGTGGT-AAAATGGC-CACACAGCAATAG | loxF8L | 3 | loxF8R | 4 | AAAATGGC/GCCATTTT | 7 |
| --- | | | | | | | | 0 |
| chr15 | 24801549 | ATAAATGTCAGGA-TTAATTTT-TCAACATTTCTAT | loxF8L | 3 | loxF8L | 4 | AAAATTAA/TTAATTTT | 7 |
| chr10 | 86201270 | CTAATGCTGAGTG-AAAATTAA-AAAACAATCTCAG | loxF8R | 4 | loxF8R | 3 | AAAATTAA/TTAATTTT | 7 |
| --- | | | | | | | | 0 |
| chr2 | 116466048 | ATAAATTTATGTA-AAAATTAT-TCCAAATATTGAC | loxF8L | 3 | loxF8L | 4 | AAAATTAT/ATAATTTT | 7 |
| chr2 | 223916219 | CTAACTTTTTGTG-AAAATTAT-TACAGAATCTAAA | loxF8R | 3 | loxF8R | 4 | AAAATTAT/ATAATTTT | 7 |

| | | | | | | | | |
|---|---|---|---|---|---|---|---|---|
| --- | | | | | | | | 0 |
| chr14 | 85724499 | ATAAATCTAGTGA-TGAATTTT-TCCATTTAATTAT | loxF8L | 3 | loxF8L | 4 | AAAATTCA/TGAATTTT | 7 |
| chr2 | 95842042 | ATAAATTTGGGAA-AAAATTCA-TTAACAGAATGAT | loxF8L | 3 | loxF8L | 4 | AAAATTCA/TGAATTTT | 7 |
| chr11 | 49333853 | CAATGCTTATGTG-TGAATTTT-TACACAGAAATAT | loxF8R | 4 | loxF8L | 3 | AAAATTCA/TGAATTTT | 7 |
| chr11 | 89374272 | ATATTTCTGTGTA-AAAATTCA-CACATAAGCATTG | loxF8L | 3 | loxF8R | 4 | AAAATTCA/TGAATTTT | 7 |
| chr4 | 146176767 | ATTAGATTGTTTT-TGAATTTT-TACATCATCTTAG | loxF8R | 4 | loxF8R | 3 | AAAATTCA/TGAATTTT | 7 |
| --- | | | | | | | | 0 |
| chr8 | 123334661 | CAAAGAATGTGAA-AGAATTTT-GCAAGAGATTTAT | loxF8R | 4 | loxF8L | 3 | AAAATTCT/AGAATTTT | 7 |
| chr3 | 65641665 | ATAAGGAAGTGGA-AAAATTCT-AACACCCTCTTAG | loxF8L | 4 | loxF8R | 3 | AAAATTCT/AGAATTTT | 7 |
| --- | | | | | | | | 0 |
| chr7 | 121663240 | AAAAATTTGTGGT-AAAATTGT-TTAAGAGATTTAA | loxF8L | 3 | loxF8L | 4 | AAAATTGT/ACAATTTT | 7 |
| chr6 | 79072339 | AGAAATCAGTAGC-ACAATTTT-TCCAGAGGTCTAT | loxF8L | 4 | loxF8L | 3 | AAAATTGT/ACAATTTT | 7 |
| --- | | | | | | | | 0 |
| chr14 | 84678202 | CAAAGACTGGGAG-GAAATTTT-TGCACACATTTGT | loxF8R | 4 | loxF8L | 3 | AAAATTTC/GAAATTTT | 7 |
| chr6 | 63059346 | ATATATTTGTAGA-GAAATTTT-CCCACATTTTTAA | loxF8L | 3 | loxF8L | 4 | AAAATTTC/GAAATTTT | 7 |
| --- | | | | | | | | 0 |
| chr3 | 29866292 | ATAAAGCTGTTAA-AAACAAAC-AACAAAAACTTAC | loxF8L | 3 | loxF8R | 4 | AAACAAAC/GTTTGTTT | 7 |
| chr7 | 40235177 | TTTTATCTGTGGA-GTTTGTTT-TACATACTCTTAT | loxF8L | 3 | loxF8R | 4 | AAACAAAC/GTTTGTTT | 7 |
| chr20 | 51593995 | TTAAATCTCTTGA-AAACAAAC-ACCACAGGCTTAG | loxF8L | 3 | loxF8L | 4 | AAACAAAC/GTTTGTTT | 7 |
| --- | | | | | | | | 0 |
| chr10 | 29575466 | ATATTTCTGTGAA-ACTTGTTT-TGCACACTTCTAT | loxF8L | 3 | loxF8L | 4 | AAACAAGT/ACTTGTTT | 7 |
| chr5 | 138189209 | TTAAGATGCTGTT-AAACAAGT-TCCACAGATCAAG | loxF8R | 4 | loxF8L | 3 | AAACAAGT/ACTTGTTT | 7 |
| --- | | | | | | | | 0 |
| chr8 | 102439269 | ATTGATTGGTGGA-AAACAATA-TACACAGTTTTTT | loxF8L | 4 | loxF8L | 3 | AAACAATA/TATTGTTT | 7 |
| chr13 | 33140510 | ATATTTCTCTGGA-TATTGTTT-TTCACTGTTTTAC | loxF8L | 3 | loxF8L | 4 | AAACAATA/TATTGTTT | 7 |
| --- | | | | | | | | 0 |
| chr6 | 19363142 | ATAAATATGTATA-AAACAATG-TCTGCACATTTTT | loxF8L | 3 | loxF8L | 4 | AAACAATG/CATTGTTT | 7 |
| chr13 | 60421641 | CAAAATCTGTGTA-AAACAATG-CTCTCAACCTTAT | loxF8L | 3 | loxF8R | 4 | AAACAATG/CATTGTTT | 7 |
| --- | | | | | | | | 0 |
| chr3 | 124699420 | AGAATTATGTTGA-AAACACCT-TCCACAGTATTTT | loxF8L | 4 | loxF8L | 3 | AAACACCT/AGGTGTTT | 7 |
| chr5 | 156004419 | CTAGCATTATGTG-AGGTGTTT-TCCATATATTGTT | loxF8R | 3 | loxF8L | 4 | AAACACCT/AGGTGTTT | 7 |
| chr1 | 58067461 | ATTAATGTGTGTA-AAACACCT-ACCACAGATTCCA | loxF8L | 3 | loxF8L | 4 | AAACACCT/AGGTGTTT | 7 |
| chr6 | 154914855 | CTGAGATTATAGG-AAACACCT-TCGACATATTTAA | loxF8R | 4 | loxF8L | 3 | AAACACCT/AGGTGTTT | 7 |
| --- | | | | | | | | 0 |
| chr2 | 241288151 | ACAAATCAGTGAA-AAACAGAG-TCACAAAATTTAT | loxF8L | 3 | loxF8L | 4 | AAACAGAG/CTCTGTTT | 7 |
| chr4 | 96420085 | ATGAATCAATGGA-AAACAGAG-TCCAAAGATAGAC | loxF8L | 3 | loxF8L | 4 | AAACAGAG/CTCTGTTT | 7 |
| --- | | | | | | | | 0 |
| chr20 | 39213161 | CTCAGAGTGTCTG-AAACCAAA-CACACCAACTTCT | loxF8R | 3 | loxF8R | 4 | AAACCAAA/TTTGGTTT | 7 |
| chr18 | 74812506 | TTATGATTTTGTC-TTTGGTTT-TCAACACATTGAT | loxF8R | 4 | loxF8L | 3 | AAACCAAA/TTTGGTTT | 7 |
| --- | | | | | | | | 0 |
| chr7 | 143822101 | CTACATCTGGGTA-CATGGTTT-TCGACAGGTATAT | loxF8L | 4 | loxF8L | 3 | AAACCATG/CATGGTTT | 7 |
| chr7 | 143635506 | ATATACCTGTCGA-AAACCATG-TACCCAGATGTAG | loxF8L | 3 | loxF8L | 4 | AAACCATG/CATGGTTT | 7 |
| chr7 | 143714724 | ATATACCTGTCGA-AAACCATG-TACCCAGATGTAG | loxF8L | 3 | loxF8L | 4 | AAACCATG/CATGGTTT | 7 |
| --- | | | | | | | | 0 |
| chr10 | 80782578 | CAAAGGTTTGGTG-TTAGGTTT-TTCAAAGTTTTAT | loxF8R | 4 | loxF8L | 3 | AAACCTAA/TTAGGTTT | 7 |
| chrX | 81138196 | ATAAAACTTTGGC-AAACCTAA-TCCAACAATGTAT | loxF8L | 3 | loxF8L | 4 | AAACCTAA/TTAGGTTT | 7 |
| --- | | | | | | | | 0 |
| chr20 | 52989814 | AAAAGATTGTGTA-AATAGTTT-CACACTATATTTT | loxF8R | 3 | loxF8R | 4 | AAACTATT/AATAGTTT | 7 |
| chr7 | 112768004 | ATAAAATTAAGGA-AATAGTTT-TACACAGTTGTAT | loxF8L | 4 | loxF8L | 3 | AAACTATT/AATAGTTT | 7 |
| --- | | | | | | | | 0 |
| chrX | 47797616 | AAAAATCTTTTGA-AAACTCCA-TCCAGAGAGTCTT | loxF8L | 3 | loxF8L | 4 | AAACTCCA/TGGAGTTT | 7 |
| chr3 | 117374574 | CTATGATTTTGGG-TGGAGTTT-TCCACCTCTTTGT | loxF8R | 3 | loxF8L | 4 | AAACTCCA/TGGAGTTT | 7 |
| --- | | | | | | | | 0 |
| chr7 | 131932150 | ATACATCTTTGGC-AAGAGTTT-CACAGAAACTTCT | loxF8L | 3 | loxF8R | 4 | AAACTCTT/AAGAGTTT | 7 |
| chr5 | 156158406 | CTGAGAATGTGGG-AAGAGTTT-AAAGCAATCTTGG | loxF8R | 3 | loxF8R | 4 | AAACTCTT/AAGAGTTT | 7 |
| --- | | | | | | | | 0 |
| chr1 | 74163297 | AAAAATCTGTGTG-AAACTTCC-CACACTAGCATGG | loxF8L | 3 | loxF8R | 4 | AAACTTCC/GGAAGTTT | 7 |
| chr8 | 34737104 | AGAAATATTTGGA-GGAAGTTT-TAAATATATTTAT | loxF8L | 3 | loxF8L | 4 | AAACTTCC/GGAAGTTT | 7 |
| --- | | | | | | | | 0 |
| chr10 | 94589597 | ATAAACCTACGTA-TTTTCTTT-TCAAAATATTTAT | loxF8L | 4 | loxF8L | 3 | AAAGAAAA/TTTTCTTT | 7 |
| chr2 | 213873618 | TTACATCTGTGAA-TTTTCTTT-CAAGCAATGCTAG | loxF8L | 3 | loxF8R | 4 | AAAGAAAA/TTTTCTTT | 7 |
| --- | | | | | | | | 0 |
| chr21 | 18220454 | CTAACAATGTTTG-ATTTCTTT-TTCATTTATTTAT | loxF8R | 3 | loxF8L | 4 | AAAGAAAT/ATTTCTTT | 7 |
| chr6 | 50755259 | TTAAATGTGTGTA-ATTTCTTT-ACTGCACATTTAT | loxF8L | 3 | loxF8L | 4 | AAAGAAAT/ATTTCTTT | 7 |
| chr2 | 98079632 | ATGGCTCTGTGGA-ATTTCTTT-TTCACATTATTAG | loxF8L | 3 | loxF8R | 4 | AAAGAAAT/ATTTCTTT | 7 |
| --- | | | | | | | | 0 |
| chr10 | 49947933 | AAAAAAATGGGGA-AAAGACAA-TCCACAGACTGAG | loxF8L | 4 | loxF8L | 3 | AAAGACAA/TTGTCTTT | 7 |
| chr2 | 203723169 | TTAAGATAATATG-AAAGACAA-TCCACAGAATGAG | loxF8R | 4 | loxF8L | 3 | AAAGACAA/TTGTCTTT | 7 |
| --- | | | | | | | | 0 |
| chr4 | 111123392 | ACTAATCTGTGAA-AAAGAGAA-TGCTCAACTTTAT | loxF8L | 3 | loxF8L | 4 | AAAGAGAA/TTCTCTTT | 7 |
| chr9 | 15279304 | ATAAATGTCTCAA-AAAGAGAA-CACAGAACCTTAT | loxF8L | 4 | loxF8R | 3 | AAAGAGAA/TTCTCTTT | 7 |
| --- | | | | | | | | 0 |
| chr8 | 23921584 | ATAAAACAGTCAA-CACTCTTT-TCCACTGAATTAA | loxF8L | 4 | loxF8L | 3 | AAAGAGTG/CACTCTTT | 7 |
| chr18 | 53951565 | GTAAATCTATTCA-CACTCTTT-CACACATTCCTCG | loxF8L | 4 | loxF8R | 3 | AAAGAGTG/CACTCTTT | 7 |
| --- | | | | | | | | 0 |
| chr17 | 73479467 | CTATATTTGTGGA-TTTGCTTT-TGCAAAGATTCTT | loxF8L | 3 | loxF8L | 4 | AAAGCAAA/TTTGCTTT | 7 |
| chr15 | 35932145 | CTAAGTTAGTTTA-TTTGCTTT-CTCACTAGCTTAG | loxF8R | 4 | loxF8R | 3 | AAAGCAAA/TTTGCTTT | 7 |
| --- | | | | | | | | 0 |
| chr1 | 111713847 | ATTAAGTTCTGGA-AAAGGAAA-GAAACAAGCTTAG | loxF8L | 4 | loxF8R | 3 | AAAGGAAA/TTTCCTTT | 7 |
| chr9 | 93331661 | CTAGATTTGGGTG-TTTCCTTT-CCCTCAGATTAAT | loxF8R | 4 | loxF8L | 3 | AAAGGAAA/TTTCCTTT | 7 |
| chr18 | 27876324 | TTAAGATTTTTTT-TTTCCTTT-CATACATTGTTAG | loxF8R | 4 | loxF8R | 3 | AAAGGAAA/TTTCCTTT | 7 |
| chr11 | 111339380 | CTAAGGTAGTGTT-AAAGGAAA-TTTATAGATTTAA | loxF8R | 3 | loxF8L | 4 | AAAGGAAA/TTTCCTTT | 7 |
| --- | | | | | | | | 0 |
| chr22 | 28369419 | AGAAATATTTGAA-AAAGGTTG-TCCACTGAATGAT | loxF8L | 4 | loxF8L | 3 | AAAGGTTG/CAACCTTT | 7 |
| chr1 | 105587224 | ATTTATCTTTGAA-AAAGGTTG-AAAAAAATCTTAG | loxF8L | 4 | loxF8R | 3 | AAAGGTTG/CAACCTTT | 7 |
| --- | | | | | | | | 0 |
| chr8 | 110040944 | ATATATTTGTGGT-TTTACTTT-TCTTCAATTTTAT | loxF8L | 3 | loxF8L | 4 | AAAGTAAA/TTTACTTT | 7 |

| | | | | | | | | |
|---|---|---|---|---|---|---|---|---|
| chr2 | 165755750 | CTAAATGTATGCA-AAAGTAAA-TACTCAATCTTAT | loxF8L | 4 | loxF8R | 3 | AAAGTAAA/TTTACTTT | 7 |
| chrX | 116129839 | GTAAATCTGTTTA-AAAGTAAA-TCTACAAAAATAT | loxF8L | 3 | loxF8L | 4 | AAAGTAAA/TTTACTTT | 7 |
| --- | | | | | | | | 0 |
| chr20 | 18899129 | ATAAATATATGAA-AAAGTACA-TATATAGATATAT | loxF8L | 3 | loxF8L | 4 | AAAGTACA/TGTACTTT | 7 |
| chrX | 21789692 | TGAAGATTGTGCT-TGTACTTT-TCCATGCATTTAT | loxF8R | 4 | loxF8L | 3 | AAAGTACA/TGTACTTT | 7 |
| --- | | | | | | | | 0 |
| chr2 | 59576122 | ATAATGCTATGTA-AAAGTCAC-CATAAAATCTCAG | loxF8L | 4 | loxF8R | 3 | AAAGTCAC/GTGACTTT | 7 |
| chrX | 140194006 | CTAATATTGATTG-GTGACTTT-AAGAAAATCTTAA | loxF8R | 3 | loxF8R | 4 | AAAGTCAC/GTGACTTT | 7 |
| --- | | | | | | | | 0 |
| chr12 | 44996204 | ATAAAAATGTGTG-CTTTATTT-CAAAGAATCTAAA | loxF8R | 3 | loxF8R | 4 | AAATAAAG/CTTTATTT | 7 |
| chr22 | 47905042 | CTGAGTTTTTGTG-AAATAAAG-GCAACAAATTTAA | loxF8R | 3 | loxF8L | 4 | AAATAAAG/CTTTATTT | 7 |
| --- | | | | | | | | 0 |
| chr4 | 60513944 | CTAAAAATGAGTA-AAATAAGA-TGCACATATTTCT | loxF8R | 4 | loxF8L | 3 | AAATAAGA/TCTTATTT | 7 |
| chr12 | 67459630 | ATAAAAATGTAGA-TCTTATTT-TGCAGGGGTTTAT | loxF8L | 3 | loxF8L | 4 | AAATAAGA/TCTTATTT | 7 |
| chr5 | 52299562 | TTTAATCAGTGGA-TCTTATTT-AACACTATTTTTG | loxF8L | 3 | loxF8R | 4 | AAATAAGA/TCTTATTT | 7 |
| --- | | | | | | | | 0 |
| chr2 | 228520614 | ATAAATCTATTTA-AAATAAGT-TATAGAGATTTGT | loxF8L | 3 | loxF8L | 4 | AAATAAGT/ACTTATTT | 7 |
| chr5 | 145236860 | TTATGTCTGTGGC-ACTTATTT-TCCTCAGATGCAT | loxF8L | 4 | loxF8L | 3 | AAATAAGT/ACTTATTT | 7 |
| --- | | | | | | | | 0 |
| chr14 | 38602416 | ATAAATGTGTGAT-AAATACAT-AACACAACTTTAT | loxF8L | 3 | loxF8L | 4 | AAATACAT/ATGTATTT | 7 |
| chr9 | 26014216 | ATACATCTGAAGA-AAATACAT-CACAAAAAATCAG | loxF8L | 3 | loxF8R | 4 | AAATACAT/ATGTATTT | 7 |
| --- | | | | | | | | 0 |
| chr4 | 162888209 | ATAAGTCACTGTA-GGGTATTT-CACACAATTTTCA | loxF8L | 4 | loxF8R | 3 | AAATACCC/GGGTATTT | 7 |
| chr14 | 62587672 | CTGAAGTTGTGTG-GGGTATTT-TCTAAATATCTAT | loxF8R | 3 | loxF8L | 4 | AAATACCC/GGGTATTT | 7 |
| --- | | | | | | | | 0 |
| chr6 | 60780786 | TTCACATTCTGTG-AAATAGTT-TCCACAGCATTGT | loxF8R | 4 | loxF8L | 3 | AAATAGTT/AACTATTT | 7 |
| chr6 | 48049053 | AGAAATCTGTGAG-AACTATTT-CACTCAATGTTCA | loxF8L | 3 | loxF8R | 4 | AAATAGTT/AACTATTT | 7 |
| chr16 | 76790503 | CTAACATTTTATG-AAATAGTT-TCAAAAAATATAT | loxF8R | 3 | loxF8L | 4 | AAATAGTT/AACTATTT | 7 |
| chr4 | 90976135 | ATTAACCTAAGGA-AACTATTT-GCCACAGATTTTG | loxF8L | 4 | loxF8L | 3 | AAATAGTT/AACTATTT | 7 |
| chr7 | 45513480 | TCAAAACTGTGTG-AAATAGTT-TGCAAAGATTTAA | loxF8R | 4 | loxF8L | 3 | AAATAGTT/AACTATTT | 7 |
| --- | | | | | | | | 0 |
| chr4 | 90022831 | ATATATCTCTGTA-AAATATAC-TCCACTATTTTCT | loxF8L | 3 | loxF8L | 4 | AAATATAC/GTATATTT | 7 |
| chr2 | 155190372 | TTAAAACTTTGTG-GTATATTT-ACCACATATGTAT | loxF8R | 4 | loxF8L | 3 | AAATATAC/GTATATTT | 7 |
| --- | | | | | | | | 0 |
| chr5 | 76127391 | ACATATTTGTGCA-AAATATCA-CACACAATGATTG | loxF8L | 4 | loxF8R | 3 | AAATATCA/TGATATTT | 7 |
| chr9 | 30554691 | ATAAGTGTGTCAA-TGATATTT-TCCAGTGCTTTAT | loxF8L | 4 | loxF8L | 3 | AAATATCA/TGATATTT | 7 |
| --- | | | | | | | | 0 |
| chr10 | 73563797 | ATATATATGTTGA-AGATATTT-TCTCCAGGTTTGT | loxF8L | 3 | loxF8L | 4 | AAATATCT/AGATATTT | 7 |
| chr6 | 149298737 | ATTAGAATGTGGA-AGATATTT-TACAAAATCTAAG | loxF8L | 4 | loxF8R | 3 | AAATATCT/AGATATTT | 7 |
| --- | | | | | | | | 0 |
| chr13 | 103699844 | AAAAATCTGAGTA-AAATATGA-GCAAAAGATATAT | loxF8L | 3 | loxF8L | 4 | AAATATGA/TCATATTT | 7 |
| chr5 | 109722840 | CTAACAGTGTCTG-TCATATTT-TAGACAGTTTTCT | loxF8R | 3 | loxF8L | 4 | AAATATGA/TCATATTT | 7 |
| chr1 | 92994418 | ATACATCTGAGAG-AAATATGA-TCTATAGATTTAC | loxF8L | 4 | loxF8L | 3 | AAATATGA/TCATATTT | 7 |
| chr4 | 168125579 | CTATGAATAAGTG-AAATATGA-TCCATATATTGAT | loxF8R | 4 | loxF8L | 3 | AAATATGA/TCATATTT | 7 |
| --- | | | | | | | | 0 |
| chr9 | 81018489 | ATAAGTCTTTTAA-AAATATTT-CTCACAGATTTCT | loxF8L | 4 | loxF8L | 3 | AAATATTT/AAATATTT | 7 |
| chr8 | 64904908 | ATCAAACTCTGTA-AAATATTT-TTAAAAGATTTAT | loxF8L | 4 | loxF8L | 3 | AAATATTT/AAATATTT | 7 |
| --- | | | | | | | | 0 |
| chr17 | 18593030 | ATAAAGCTTTGGC-AAATCAAA-TCCAATGATACAT | loxF8L | 3 | loxF8L | 4 | AAATCAAA/TTTGATTT | 7 |
| chr17 | 16722306 | ATGTATCATTGGA-TTTGATTT-GCCAAAGCTTTAT | loxF8L | 4 | loxF8L | 3 | AAATCAAA/TTTGATTT | 7 |
| chr17 | 20353053 | ATGTATCATTGGA-TTTGATTT-GCCAAAGCTTTAT | loxF8L | 4 | loxF8L | 3 | AAATCAAA/TTTGATTT | 7 |
| chr10 | 65509150 | ATAAGAACTTGTG-TTTGATTT-TCCACAATTTTAC | loxF8R | 4 | loxF8L | 3 | AAATCAAA/TTTGATTT | 7 |
| --- | | | | | | | | 0 |
| chr7 | 86088635 | CTAAAATTGTGCA-AAATCAGT-CACAAGTTCTTAT | loxF8R | 3 | loxF8R | 4 | AAATCAGT/ACTGATTT | 7 |
| chr2 | 38048117 | GTATGTTTGTGTG-AAATCAGT-TGCTGAGATATAT | loxF8R | 3 | loxF8L | 4 | AAATCAGT/ACTGATTT | 7 |
| --- | | | | | | | | 0 |
| chr18 | 65355853 | TTAAAATTGTATT-AAATCCAA-TCCACAAATATTT | loxF8R | 4 | loxF8L | 3 | AAATCCAA/TTGGATTT | 7 |
| chr7 | 46438652 | CGAAATCTGTTAA-TTGGATTT-TCCACAGTTTGAA | loxF8L | 4 | loxF8L | 3 | AAATCCAA/TTGGATTT | 7 |
| --- | | | | | | | | 0 |
| chr6 | 23490577 | AGAAATATGGGAA-ATAGATTT-CACAAAATTTTAA | loxF8L | 4 | loxF8R | 3 | AAATCTAT/ATAGATTT | 7 |
| chr11 | 127669154 | CAAAGATTCACTG-AAATCTAT-GTCAAAATCTTAG | loxF8R | 4 | loxF8R | 3 | AAATCTAT/ATAGATTT | 7 |
| --- | | | | | | | | 0 |
| chr3 | 27878018 | ATGTTTCTGTTGA-AAATCTTA-TTCACACATTTGT | loxF8L | 4 | loxF8L | 3 | AAATCTTA/TAAGATTT | 7 |
| chr8 | 23185384 | ACACATCTTTGGC-AAATCTTA-TGTACAGAATTAT | loxF8L | 4 | loxF8L | 3 | AAATCTTA/TAAGATTT | 7 |
| chr3 | 75181992 | ATAAATGTGTATT-AAATCTTA-TCAATTGATTTAT | loxF8L | 4 | loxF8L | 3 | AAATCTTA/TAAGATTT | 7 |
| --- | | | | | | | | 0 |
| chr20 | 42886026 | TTTAAACTCTGGA-GAAGATTT-TCCATATATATAT | loxF8L | 4 | loxF8L | 3 | AAATCTTC/GAAGATTT | 7 |
| chr18 | 69610531 | TTAAATCCTTTGA-GAAGATTT-CAATCAAACTTAG | loxF8L | 4 | loxF8R | 3 | AAATCTTC/GAAGATTT | 7 |
| --- | | | | | | | | 0 |
| chr8 | 51634501 | ACAATTCTGTGAA-AAATGACA-TCCGTAGTTTGAT | loxF8L | 3 | loxF8L | 4 | AAATGACA/TGTCATTT | 7 |
| chr4 | 33112245 | ATAAAACTGCTGT-TGTCATTT-TTCACAGAATTAG | loxF8L | 4 | loxF8L | 3 | AAATGACA/TGTCATTT | 7 |
| chr3 | 84544055 | CTAATTCTGTGAA-AAATGACA-TTGGCATATTTAT | loxF8L | 3 | loxF8L | 4 | AAATGACA/TGTCATTT | 7 |
| chr13 | 71413505 | TTATTTCTGTGGA-AAATGACA-TTCATAGTTTGAT | loxF8L | 3 | loxF8L | 4 | AAATGACA/TGTCATTT | 7 |
| chr10 | 123534128 | ATCAAACTATGAA-TGTCATTT-TTCACAGAATTAG | loxF8L | 4 | loxF8L | 3 | AAATGACA/TGTCATTT | 7 |
| chrX | 73734453 | CTAATTCTGTGAA-AAATGACA-TTGACAGTTTGAT | loxF8L | 3 | loxF8L | 4 | AAATGACA/TGTCATTT | 7 |
| chr14 | 28533143 | CTAATTCTGTGAA-AAATGACA-TTGACAGTTTAAT | loxF8L | 3 | loxF8L | 4 | AAATGACA/TGTCATTT | 7 |
| --- | | | | | | | | 0 |
| chr13 | 86736387 | ATAAATATATGTA-CATCATTT-TTCACAGAAATAG | loxF8L | 3 | loxF8L | 4 | AAATGATG/CATCATTT | 7 |
| chr3 | 88848614 | CTAATTCTGTGAA-AAATGATG-TTGACAGTTTGAT | loxF8L | 3 | loxF8L | 4 | AAATGATG/CATCATTT | 7 |
| chrX | 76040797 | CTAATTATGTGGA-AAATGATG-TTAATAGATTGAT | loxF8L | 3 | loxF8L | 4 | AAATGATG/CATCATTT | 7 |
| --- | | | | | | | | 0 |
| chr3 | 37608521 | CTGAAACTGTGTG-AATCATTT-TAAACAGGTTTGT | loxF8R | 3 | loxF8L | 4 | AAATGATT/AATCATTT | 7 |
| chr6 | 81864514 | ATATGTTTGTCTG-AATCATTT-CTAACAATGTTAG | loxF8R | 4 | loxF8R | 3 | AAATGATT/AATCATTT | 7 |
| --- | | | | | | | | 0 |
| chr11 | 110988899 | AAAAATGGGTGGA-GTGCATTT-TCCCAACATTTTT | loxF8L | 3 | loxF8L | 4 | AAATGCAC/GTGCATTT | 7 |
| chr8 | 58478405 | ACATCTCTGTGGA-AAATGCAC-TTAACAAATATAT | loxF8L | 3 | loxF8L | 4 | AAATGCAC/GTGCATTT | 7 |
| --- | | | | | | | | 0 |

| | | | | | | | | |
|---|---|---|---|---|---|---|---|---|
| chr5 | 176755087 | ATAAAACTGTTTA-AAATGGGA-AACATAATGTTAA | loxF8L | 3 | loxF8R | 4 | AAATGGGA/TCCCATTT | 7 |
| chrY | 13209129 | CTCATGTTATGTG-TCCCATTT-TCAACAGAAGTAT | loxF8R | 4 | loxF8L | 3 | AAATGGGA/TCCCATTT | 7 |
| --- | | | | | | | | 0 |
| chrX | 101031804 | AAAAACCTTTTGA-AAATTAAA-GACACAATGATAG | loxF8L | 4 | loxF8R | 3 | AAATTAAA/TTTAATTT | 7 |
| chr10 | 63373658 | TTAAATTTGTAGA-AAATTAAA-GACAAAGTATTAG | loxF8L | 3 | loxF8R | 4 | AAATTAAA/TTTAATTT | 7 |
| --- | | | | | | | | 0 |
| chr1 | 88010587 | AAATGATTCTGTG-AAATTACA-GCCTCGGATTTAT | loxF8R | 4 | loxF8L | 3 | AAATTACA/TGTAATTT | 7 |
| chr13 | 108306411 | ATATATATGTGTG-TGTAATTT-TTCACATTTTTAT | loxF8L | 4 | loxF8L | 3 | AAATTACA/TGTAATTT | 7 |
| --- | | | | | | | | 0 |
| chr12 | 122244733 | AAATTTCTCTGGA-GGTAATTT-CACATAACCTGAG | loxF8L | 4 | loxF8R | 3 | AAATTACC/GGTAATTT | 7 |
| chr12 | 101585335 | CTAAAATTTTGGG-GGTAATTT-TTCACACATTTCC | loxF8R | 3 | loxF8L | 4 | AAATTACC/GGTAATTT | 7 |
| --- | | | | | | | | 0 |
| chr11 | 56189480 | ATAAATATGGAGA-AAATTATA-TGTACATAATTAT | loxF8L | 3 | loxF8L | 4 | AAATTATA/TATAATTT | 7 |
| chr3 | 73150352 | ATAAATATGGGGC-AAATTATA-TAATCAATCTTTG | loxF8L | 3 | loxF8R | 4 | AAATTATA/TATAATTT | 7 |
| chr6 | 123716943 | ATAAATGTGTAAA-AAATTATA-CACACACTTTAAT | loxF8L | 3 | loxF8R | 4 | AAATTATA/TATAATTT | 7 |
| --- | | | | | | | | 0 |
| chr5 | 21021544 | AAAAATTTGTTGA-GATAATTT-CATAAAATATTAA | loxF8L | 3 | loxF8R | 4 | AAATTATC/GATAATTT | 7 |
| chr6 | 78238825 | ATAAAACTATAGA-AAATTATC-CACATAATTTCAA | loxF8L | 3 | loxF8R | 4 | AAATTATC/GATAATTT | 7 |
| --- | | | | | | | | 0 |
| chr12 | 30097413 | ATATATGTGAGGA-AAATTATG-CACAGAAAGATAG | loxF8L | 3 | loxF8R | 4 | AAATTATG/CATAATTT | 7 |
| chr18 | 39660087 | ATCACATTGTGTG-CATAATTT-GCAACAGATAGAT | loxF8R | 3 | loxF8L | 4 | AAATTATG/CATAATTT | 7 |
| --- | | | | | | | | 0 |
| chr11 | 75007302 | TTAAAACACTGGA-CAGAATTT-CCCACAGTCTTTG | loxF8L | 4 | loxF8R | 3 | AAATTCTG/CAGAATTT | 7 |
| chr3 | 135132250 | GTGAGATTTTGTG-CAGAATTT-CAGAATATGTTAG | loxF8R | 3 | loxF8R | 4 | AAATTCTG/CAGAATTT | 7 |
| --- | | | | | | | | 0 |
| chr15 | 29348227 | ATAAAACTGTCAA-GTAAATTT-CACACAGTGAGAG | loxF8L | 3 | loxF8R | 4 | AAATTTAC/GTAAATTT | 7 |
| chr2 | 231720180 | TTAAGGTTGTTTT-GTAAATTT-TCCACACATAGAT | loxF8R | 4 | loxF8L | 3 | AAATTTAC/GTAAATTT | 7 |
| --- | | | | | | | | 0 |
| chr2 | 135866525 | CTGAGAATTTGTT-AAATTTGA-GCCACAGATTACT | loxF8R | 4 | loxF8L | 3 | AAATTTGA/TCAAATTT | 7 |
| chr14 | 40633894 | GTAAAATTGTATT-AAATTTGA-TACACATACTTAG | loxF8R | 4 | loxF8R | 3 | AAATTTGA/TCAAATTT | 7 |
| --- | | | | | | | | 0 |
| chr1 | 214327973 | ATAAATTTGAGTC-AAATTTGT-CCCAAAGCTTTAT | loxF8L | 4 | loxF8L | 3 | AAATTTGT/ACAAATTT | 7 |
| chr10 | 16840682 | TAAAGATTGTATC-AAATTTGT-ACCACAGATTAAA | loxF8R | 4 | loxF8L | 3 | AAATTTGT/ACAAATTT | 7 |
| --- | | | | | | | | 0 |
| chr18 | 60669255 | ATAAATCTTTTAA-AAATTTTC-TTGCCAGTTTTAT | loxF8L | 3 | loxF8L | 4 | AAATTTTC/GAAAATTT | 7 |
| chr14 | 46983363 | ATAAAATTTTCTG-AAATTTTC-TTCACAGTTATAT | loxF8R | 4 | loxF8L | 3 | AAATTTTC/GAAAATTT | 7 |
| --- | | | | | | | | 0 |
| chr12 | 58638682 | AAAAATCAGTTAA-AAATTTTG-CCTACAAATTTAT | loxF8L | 4 | loxF8L | 3 | AAATTTTG/CAAAATTT | 7 |
| chr1 | 198504460 | ATAAAAGTGTGTG-AAATTTTG-TCCAATGTTTTCT | loxF8R | 3 | loxF8L | 4 | AAATTTTG/CAAAATTT | 7 |
| chr6 | 121633403 | ATAAATATGTTCC-AAATTTTG-TTCACAGAAGTAT | loxF8L | 4 | loxF8L | 3 | AAATTTTG/CAAAATTT | 7 |
| chr14 | 47157116 | ATAAATCGATGGG-AAATTTTG-TTCAAATATTGAT | loxF8L | 3 | loxF8L | 4 | AAATTTTG/CAAAATTT | 7 |
| --- | | | | | | | | 0 |
| chr2 | 4947568 | CAAAGAATGAGTG-AACAAAAT-ACCACAAATATGT | loxF8R | 3 | loxF8L | 4 | AACAAAAT/ATTTTGTT | 7 |
| chr13 | 114326681 | ATAAATTTGAGAA-ATTTTGTT-TCCTTACATGTAT | loxF8L | 3 | loxF8L | 4 | AACAAAAT/ATTTTGTT | 7 |
| --- | | | | | | | | 0 |
| chr12 | 78407682 | GTAATTCTGTGGT-AACAAACA-ACCACAAAATTTT | loxF8L | 3 | loxF8L | 4 | AACAAACA/TGTTTGTT | 7 |
| chr3 | 174513480 | CTTAGACTGTATC-TGTTTGTT-TCCAAAGATTGGT | loxF8R | 4 | loxF8L | 3 | AACAAACA/TGTTTGTT | 7 |
| --- | | | | | | | | 0 |
| chr11 | 26020937 | ATAAAAATTTGGA-AACAAACT-ACCAGAAATTCAT | loxF8L | 3 | loxF8L | 4 | AACAAACT/AGTTTGTT | 7 |
| chrX | 114410857 | CTAAACTTCTCTG-AGTTTGTT-TCCATATATTTAA | loxF8R | 4 | loxF8L | 3 | AACAAACT/AGTTTGTT | 7 |
| --- | | | | | | | | 0 |
| chr3 | 106544798 | TTAAGATTGGATA-TATTTGTT-TACAAAGTTTTAT | loxF8R | 4 | loxF8L | 3 | AACAAATA/TATTTGTT | 7 |
| chr21 | 28712904 | CTATTTTTGTGTA-TATTTGTT-TCCATAGATACAT | loxF8R | 4 | loxF8L | 3 | AACAAATA/TATTTGTT | 7 |
| --- | | | | | | | | 0 |
| chr8 | 64781318 | CTAAGAATGTTAT-AACAAATT-ACCACAAATTTAG | loxF8R | 4 | loxF8L | 3 | AACAAATT/AATTTGTT | 7 |
| chr6 | 160377864 | TTTAACCTGTGGA-AATTTGTT-CACTCATTCTAAC | loxF8L | 3 | loxF8R | 4 | AACAAATT/AATTTGTT | 7 |
| --- | | | | | | | | 0 |
| chr8 | 84720579 | CAAAGATTTTTTG-AACAAGAA-CTCAAAAGCATAG | loxF8R | 3 | loxF8R | 4 | AACAAGAA/TTCTTGTT | 7 |
| chr1 | 74422156 | CTAAGGTTATGTA-TTCTTGTT-CTCACTATTTAAG | loxF8R | 3 | loxF8R | 4 | AACAAGAA/TTCTTGTT | 7 |
| chr2 | 53156028 | TTAAGCTTGTGTT-TTCTTGTT-CAAACATTCTCAA | loxF8R | 3 | loxF8R | 4 | AACAAGAA/TTCTTGTT | 7 |
| --- | | | | | | | | 0 |
| chr6 | 164576412 | AAAAATCTGCTGA-AACAAGTA-ACCTCATATTTAG | loxF8L | 3 | loxF8L | 4 | AACAAGTA/TACTTGTT | 7 |
| chr1 | 42649649 | CTGAAATTGTGTC-TACTTGTT-GACATGATCTTAT | loxF8R | 3 | loxF8R | 4 | AACAAGTA/TACTTGTT | 7 |
| --- | | | | | | | | 0 |
| chr20 | 12912175 | TTTAATTTGGGGA-TTATTGTT-TCCATATATTTTT | loxF8L | 4 | loxF8L | 3 | AACAATAA/TTATTGTT | 7 |
| chr6 | 151074436 | CTAAATATATGGC-AACAATAA-CACAAAGTCTAAG | loxF8L | 4 | loxF8R | 3 | AACAATAA/TTATTGTT | 7 |
| --- | | | | | | | | 0 |
| chr2 | 230165225 | ATTAATTTATGAA-AACAATAG-AACACATTCTAAG | loxF8L | 4 | loxF8R | 3 | AACAATAG/CTATTGTT | 7 |
| chr10 | 52987590 | TTAAATCTGTACA-CTATTGTT-TACAAAAATTTAA | loxF8L | 3 | loxF8L | 4 | AACAATAG/CTATTGTT | 7 |
| --- | | | | | | | | 0 |
| chr9 | 32391408 | ATAGACCTATTGA-AACAATAT-CACACTCTCTGAG | loxF8L | 4 | loxF8R | 3 | AACAATAT/ATATTGTT | 7 |
| chr10 | 37615443 | CTCAGATTTTCTG-ATATTGTT-TCCACAGTATTCC | loxF8R | 3 | loxF8L | 4 | AACAATAT/ATATTGTT | 7 |
| --- | | | | | | | | 0 |
| chr6 | 55978631 | AAAAATCTATAGC-TCATTGTT-TGCAGAGATTTTT | loxF8L | 4 | loxF8L | 3 | AACAATGA/TCATTGTT | 7 |
| chr16 | 51472514 | TTAAGTTTGTGAT-TCATTGTT-TTCATACATTTAT | loxF8R | 4 | loxF8L | 3 | AACAATGA/TCATTGTT | 7 |
| --- | | | | | | | | 0 |
| chr5 | 93740118 | CCTGGATTTTGTG-TTTGTGTT-TACACAGAGTTCT | loxF8R | 4 | loxF8L | 3 | AACACAAA/TTTGTGTT | 7 |
| chr6 | 114995410 | CCAAATCTATTGA-AACACAAA-GCCAGAGAGTTAT | loxF8L | 4 | loxF8L | 3 | AACACAAA/TTTGTGTT | 7 |
| --- | | | | | | | | 0 |
| chr3 | 29810676 | ACATATATGTGTA-TGTGTGTT-TCCATACATTTTT | loxF8L | 4 | loxF8L | 3 | AACACACA/TGTGTGTT | 7 |
| chr6 | 115823778 | ACATATATGTGCA-AACACACA-CACACATTCATAA | loxF8L | 4 | loxF8R | 3 | AACACACA/TGTGTGTT | 7 |
| chr3 | 65534133 | GTATGTTTGTGTG-TGTGTGTT-TCTACAGCATGAT | loxF8R | 3 | loxF8L | 4 | AACACACA/TGTGTGTT | 7 |
| chrX | 18025697 | GAGAGAGTGTGTG-TGTGTGTT-CACATAAGTTTAG | loxF8R | 4 | loxF8R | 3 | AACACACA/TGTGTGTT | 7 |
| --- | | | | | | | | 0 |
| chr8 | 50461474 | ATAATTATTTAGA-ATTCTGTT-AACATAGATTTAT | loxF8L | 4 | loxF8L | 3 | AACAGAAT/ATTCTGTT | 7 |
| chr6 | 145127915 | CTAAATTTGTAGA-ATTCTGTT-TCCATCTATTCAT | loxF8L | 3 | loxF8L | 4 | AACAGAAT/ATTCTGTT | 7 |
| --- | | | | | | | | 0 |

| | | | | | | | | |
|---|---|---|---|---|---|---|---|---|
| chr8 | 41203023 | ACACATATGTGTA-AACAGACA-TCCACACATACAT | loxF8L | 4 | loxF8L | 3 | AACAGACA/TGTCTGTT | 7 |
| chr12 | 109341272 | GTAATATTGTCTA-TGTCTGTT-TCTTCAGATTAAT | loxF8R | 4 | loxF8L | 3 | AACAGACA/TGTCTGTT | 7 |
| chr5 | 148967500 | CTCACATTAGGTG-TGTCTGTT-CCCACAATTTAAG | loxF8R | 4 | loxF8R | 3 | AACAGACA/TGTCTGTT | 7 |
| --- | | | | | | | | 0 |
| chr2 | 154445680 | ATAAATCCTTGAA-AACATAAA-TCCAATAATTTGT | loxF8L | 3 | loxF8L | 4 | AACATAAA/TTTATGTT | 7 |
| chr2 | 184441024 | CTATATCTGTGGT-AACATAAA-TCTAATGATTAAT | loxF8L | 3 | loxF8L | 4 | AACATAAA/TTTATGTT | 7 |
| --- | | | | | | | | 0 |
| chr7 | 57544907 | CCAAGAATGTGAG-AACATATA-TTTACTGATTTTT | loxF8R | 3 | loxF8L | 4 | AACATATA/TATATGTT | 7 |
| chr10 | 81705438 | ATACATGTGTATA-AACATATA-TACACATATATAT | loxF8L | 4 | loxF8L | 3 | AACATATA/TATATGTT | 7 |
| chr2 | 226485526 | ATATATATGTGTA-AACATATA-CACACACATATAT | loxF8L | 3 | loxF8L | 4 | AACATATA/TATATGTT | 7 |
| chr10 | 17552962 | ATATATGTGTGTA-TATATGTT-TACACATATATAC | loxF8L | 3 | loxF8L | 4 | AACATATA/TATATGTT | 7 |
| --- | | | | | | | | 0 |
| chr7 | 145516509 | CCTAGTTTGTGTG-CAGATGTT-TTCATAGGTTTTT | loxF8R | 3 | loxF8L | 4 | AACATCTG/CAGATGTT | 7 |
| chr1 | 194236157 | ATAAATGTATTAA-AACATCTG-TACAAATATTTAT | loxF8L | 4 | loxF8L | 3 | AACATCTG/CAGATGTT | 7 |
| --- | | | | | | | | 0 |
| chr11 | 74437919 | ATAAATATATGTG-CACATGTT-TACACATATGTAT | loxF8L | 4 | loxF8L | 3 | AACATGTG/CACATGTT | 7 |
| chr13 | 93892870 | ATAATCATGTGCA-AACATGTG-TGAACATATTTAT | loxF8L | 4 | loxF8L | 3 | AACATGTG/CACATGTT | 7 |
| --- | | | | | | | | 0 |
| chr6 | 66314540 | ATAAATATTTTAA-AACATGTT-TCCTCAGATCTCT | loxF8L | 4 | loxF8L | 3 | AACATGTT/AACATGTT | 7 |
| --- | | | | | | | | 0 |
| chr2 | 88817487 | CTGAAATAGGGTG-AACCACTG-TATACTATCTTAG | loxF8R | 4 | loxF8R | 3 | AACCACTG/CAGTGGTT | 7 |
| chr8 | 104745913 | CTAGGCTGGAGTG-CAGTGGTT-CGTACAATCATAG | loxF8R | 4 | loxF8R | 3 | AACCACTG/CAGTGGTT | 7 |
| --- | | | | | | | | 0 |
| chr21 | 40674085 | CTAAAATTTTTTG-AACCAGAA-TCCACATAATCTT | loxF8R | 3 | loxF8L | 4 | AACCAGAA/TTCTGGTT | 7 |
| chr4 | 153004823 | CTGGGAGTGTGTG-TTCTGGTT-TCCCCTGAGCTAT | loxF8R | 3 | loxF8L | 4 | AACCAGAA/TTCTGGTT | 7 |
| --- | | | | | | | | 0 |
| chr20 | 9503070 | ATTAATATGTTGA-AACCCTAA-TCCCCAAAGTGAT | loxF8L | 3 | loxF8L | 4 | AACCCTAA/TTAGGGTT | 7 |
| chr10 | 48223706 | ATCCATATGTTGA-AACCCTAA-CACCCAATCTGAT | loxF8L | 4 | loxF8R | 3 | AACCCTAA/TTAGGGTT | 7 |
| chr4 | 26472032 | ATTCATATGTTGA-AACCCTAA-TCCACAATTTCAT | loxF8L | 4 | loxF8L | 3 | AACCCTAA/TTAGGGTT | 7 |
| --- | | | | | | | | 0 |
| chr9 | 39071087 | ATAAAACTGAGGG-AGCAGGTT-AAAAAAATCTTAC | loxF8L | 3 | loxF8R | 4 | AACCTGCT/AGCAGGTT | 7 |
| chr9 | 67296227 | ATAAGATTTTTTT-AACCTGCT-CCCTCAGTTTTAT | loxF8R | 4 | loxF8L | 3 | AACCTGCT/AGCAGGTT | 7 |
| chr9 | 41891200 | ATAAAACTGAGGG-AGCAGGTT-AAAAAAATCTTAT | loxF8L | 3 | loxF8R | 4 | AACCTGCT/AGCAGGTT | 7 |
| --- | | | | | | | | 0 |
| chr7 | 84394629 | ATAAATGTTTTAA-AACTAAAA-CCTACAGATTTAA | loxF8L | 4 | loxF8L | 3 | AACTAAAA/TTTTAGTT | 7 |
| chr6 | 66682409 | CAAAGATTATTTA-TTTTAGTT-AACAGAACCTTAG | loxF8R | 4 | loxF8R | 3 | AACTAAAA/TTTTAGTT | 7 |
| --- | | | | | | | | 0 |
| chr4 | 35529194 | TCAAGAGTGTGTT-AGTTAGTT-TCCACATGTTTGT | loxF8R | 4 | loxF8L | 3 | AACTAACT/AGTTAGTT | 7 |
| chrX | 89476683 | ATAGAGCAGTTGA-AGTTAGTT-TCCAGGTATTTAT | loxF8L | 4 | loxF8L | 3 | AACTAACT/AGTTAGTT | 7 |
| chrY | 3132873 | ATAGAGCAGTTGA-AGTTAGTT-TCCAGGTATTTAT | loxF8L | 4 | loxF8L | 3 | AACTAACT/AGTTAGTT | 7 |
| --- | | | | | | | | 0 |
| chr1 | 67030416 | ATAATTCTCTGAA-CCTTAGTT-TCTACATATGTAA | loxF8L | 3 | loxF8L | 4 | AACTAAGG/CCTTAGTT | 7 |
| chr2 | 143360641 | TTATATCTCTGGA-CCTTAGTT-TCCTCACCTGTAT | loxF8L | 3 | loxF8L | 4 | AACTAAGG/CCTTAGTT | 7 |
| --- | | | | | | | | 0 |
| chr17 | 62129011 | AAAAATGAATGGA-AATTAGTT-TTAACCGATTTAT | loxF8L | 4 | loxF8L | 3 | AACTAATT/AATTAGTT | 7 |
| chr7 | 48666134 | ATATAACTGTATA-AACTAATT-TCCAAATATCTAT | loxF8L | 4 | loxF8L | 3 | AACTAATT/AATTAGTT | 7 |
| chr12 | 127073485 | ATTAATATATGGA-AACTAATT-CACACAATTAAAA | loxF8L | 3 | loxF8R | 4 | AACTAATT/AATTAGTT | 7 |
| --- | | | | | | | | 0 |
| chr19 | 31863225 | ATAAATATGAAGA-AACTACAA-CTCAGAATATGAG | loxF8L | 3 | loxF8R | 4 | AACTACAA/TTGTAGTT | 7 |
| chr7 | 26737815 | TTAAAATTATGTG-AACTACAA-CTGAGAGATTTAT | loxF8R | 3 | loxF8L | 4 | AACTACAA/TTGTAGTT | 7 |
| --- | | | | | | | | 0 |
| chr6 | 80814400 | GTAAATCCATGGA-TAGTAGTT-TTAACAGAAGTAT | loxF8L | 3 | loxF8L | 4 | AACTACTA/TAGTAGTT | 7 |
| chr14 | 40441632 | GTAAGTCTGCAGA-TAGTAGTT-TTCACAGAATCAT | loxF8L | 4 | loxF8L | 3 | AACTACTA/TAGTAGTT | 7 |
| --- | | | | | | | | 0 |
| chr2 | 181806338 | TTCACTGTGTGGA-AACTATCA-CACACACATTTAT | loxF8L | 4 | loxF8L | 3 | AACTATCA/TGATAGTT | 7 |
| chr3 | 145668780 | ATACATCTGCAAA-AACTATCA-TCTACAAACTTAT | loxF8L | 4 | loxF8L | 3 | AACTATCA/TGATAGTT | 7 |
| --- | | | | | | | | 0 |
| chrX | 101888880 | CTCAGATTATATT-TCAGAGTT-TTTACATATTTAT | loxF8R | 4 | loxF8L | 3 | AACTCTGA/TCAGAGTT | 7 |
| chr2 | 149429891 | CTAACATGGTGTC-TCAGAGTT-TCAGCACAATTAT | loxF8R | 3 | loxF8L | 4 | AACTCTGA/TCAGAGTT | 7 |
| --- | | | | | | | | 0 |
| chr13 | 68167076 | ATATATATATGGA-ATAAAGTT-CACATAGGTTTAT | loxF8L | 3 | loxF8L | 4 | AACTTTAT/ATAAAGTT | 7 |
| chr9 | 132250669 | ATCAAGCTGTTGA-ATAAAGTT-CAGCCAGTCCTAG | loxF8L | 3 | loxF8R | 4 | AACTTTAT/ATAAAGTT | 7 |
| --- | | | | | | | | 0 |
| chr2 | 221163967 | TTAAGATTATCTA-TCAAAGTT-TGCAAAGATTTTT | loxF8R | 4 | loxF8L | 3 | AACTTTGA/TCAAAGTT | 7 |
| chr14 | 57609997 | AAAACTCTAGGGA-AACTTTGA-CCCACACATCTAT | loxF8L | 4 | loxF8L | 3 | AACTTTGA/TCAAAGTT | 7 |
| --- | | | | | | | | 0 |
| chr3 | 74430176 | AAAAATCTGTACA-TAAAAGTT-CGCAGAGTTTTAT | loxF8L | 3 | loxF8L | 4 | AACTTTTA/TAAAAGTT | 7 |
| chr3 | 1094455 | ATATAAATGTGGA-AACTTTTA-TATAAAGCTTTAT | loxF8L | 3 | loxF8L | 4 | AACTTTTA/TAAAAGTT | 7 |
| chr3 | 116828236 | ATAAAGTTGTGTA-TAAAAGTT-TACATATAGTTAT | loxF8L | 3 | loxF8L | 4 | AACTTTTA/TAAAAGTT | 7 |
| --- | | | | | | | | 0 |
| chr2 | 147882819 | AAAAATTTCTGGG-AAGAAAGC-CAGACAATGTTTG | loxF8L | 4 | loxF8R | 3 | AAGAAAGC/GCTTTCTT | 7 |
| chr10 | 53077919 | CTGAGATCTTTTG-GCTTTCTT-TCCAGAGGTTGAT | loxF8R | 4 | loxF8L | 3 | AAGAAAGC/GCTTTCTT | 7 |
| --- | | | | | | | | 0 |
| chr5 | 90440224 | ATGAATCTGTCTT-CATTTCTT-TCCACTAATTTTT | loxF8L | 4 | loxF8L | 3 | AAGAAATG/CATTTCTT | 7 |
| chr13 | 83262348 | CTAATATCTTTTG-CATTTCTT-TCCACAAAGTTTT | loxF8R | 4 | loxF8L | 3 | AAGAAATG/CATTTCTT | 7 |
| chr10 | 37433800 | TCAAATTTATGGA-AAGAAATG-AAGACAATTTTAG | loxF8L | 4 | loxF8R | 3 | AAGAAATG/CATTTCTT | 7 |
| --- | | | | | | | | 0 |
| chr6 | 92708776 | CTAAAATTGAAGG-AAGAACAA-ACCTCAAATTTAT | loxF8R | 4 | loxF8L | 3 | AAGAACAA/TTGTTCTT | 7 |
| chr14 | 27923314 | CTAAAATTGAGTC-AAGAACAA-ACATCAAATTTAT | loxF8R | 3 | loxF8L | 4 | AAGAACAA/TTGTTCTT | 7 |
| chr3 | 82299052 | CTAAAATTGTGAC-AAGAACAA-ACACCAAATTTAT | loxF8R | 3 | loxF8L | 4 | AAGAACAA/TTGTTCTT | 7 |
| chr19 | 41755049 | ATAACACTGCAGA-AAGAACAA-TTCAGAATCTTAG | loxF8L | 4 | loxF8R | 3 | AAGAACAA/TTGTTCTT | 7 |
| --- | | | | | | | | 0 |
| chr8 | 137546325 | ACAATTCTGTGTA-TACTTCTT-TTCACATGATTAT | loxF8L | 3 | loxF8L | 4 | AAGAAGTA/TACTTCTT | 7 |
| chr3 | 148710374 | TTAAAATTGTGAA-AAGAAGTA-CCCACAGTTATAT | loxF8L | 4 | loxF8L | 3 | AAGAAGTA/TACTTCTT | 7 |
| --- | | | | | | | | 0 |
| chr6 | 91554096 | AGAAAGATCTGGA-GACTTCTT-TTCACATATTTGT | loxF8L | 4 | loxF8L | 3 | AAGAAGTC/GACTTCTT | 7 |
| chrX | 105519275 | TGAAGATTCTGTC-AAGAAGTC-TCCCCAGATATCT | loxF8R | 4 | loxF8L | 3 | AAGAAGTC/GACTTCTT | 7 |

| | | | | | | | | |
|---|---|---|---|---|---|---|---|---|
| --- | | | | | | | | 0 |
| chr2 | 113533106 | CTAGGATTTAATG-AAGAATAC-TGCAGAATCTTAG | loxF8R | 4 | loxF8R | 3 | AAGAATAC/GTATTCTT | 7 |
| chr9 | 82228 | CTAAGATTCTGCA-GTATTCTT-CATTAAATCCTAG | loxF8R | 3 | loxF8R | 4 | AAGAATAC/GTATTCTT | 7 |
| chr9 | 68348234 | CTAGGATTTAATG-AAGAATAC-TGCAGAATCTTAG | loxF8R | 4 | loxF8R | 3 | AAGAATAC/GTATTCTT | 7 |
| chr9 | 41083305 | CTAAGATTCTGCA-GTATTCTT-CATTAAATCCTAG | loxF8R | 3 | loxF8R | 4 | AAGAATAC/GTATTCTT | 7 |
| chr2 | 96980870 | TTAGATCTGTGAA-GTATTCTT-TACACAGTCTTGA | loxF8L | 3 | loxF8R | 4 | AAGAATAC/GTATTCTT | 7 |
| --- | | | | | | | | 0 |
| chr3 | 44341952 | CTAGCATTTTGTC-TGTGTCTT-TCCACCATTTTAT | loxF8R | 4 | loxF8L | 3 | AAGACACA/TGTGTCTT | 7 |
| chr6 | 27714197 | GTGGGAGTGTGTG-TGTGTCTT-TCCTCACAATTAT | loxF8R | 4 | loxF8L | 3 | AAGACACA/TGTGTCTT | 7 |
| --- | | | | | | | | 0 |
| chr2 | 158307306 | ATAAACATATGAA-AAGACACT-TGGACAGCTTTAT | loxF8L | 4 | loxF8L | 3 | AAGACACT/AGTGTCTT | 7 |
| chr16 | 78052139 | GTGAGATTAGGTG-AGTGTCTT-CTCAGAAACTTAG | loxF8R | 4 | loxF8R | 3 | AAGACACT/AGTGTCTT | 7 |
| --- | | | | | | | | 0 |
| chr7 | 41952158 | AAAATATTGAGTG-TCTGTCTT-TGCACAGAACTAT | loxF8R | 4 | loxF8L | 3 | AAGACAGA/TCTGTCTT | 7 |
| chrX | 102692410 | TTAGAACTGTTGA-AAGACAGA-ACCACATATTTAA | loxF8L | 4 | loxF8L | 3 | AAGACAGA/TCTGTCTT | 7 |
| --- | | | | | | | | 0 |
| chr8 | 81127638 | ATAATGGTGAGGA-AAGACATA-CACACCATCTTGT | loxF8L | 4 | loxF8R | 3 | AAGACATA/TATGTCTT | 7 |
| chr5 | 11443476 | ATATATATGTGCA-TATGTCTT-CACACAAGCATGT | loxF8L | 3 | loxF8R | 4 | AAGACATA/TATGTCTT | 7 |
| chr1 | 99549833 | TTATGTATGTGGA-AAGACATA-TACATATATTTAT | loxF8L | 4 | loxF8L | 3 | AAGACATA/TATGTCTT | 7 |
| chr1 | 97818777 | ATATATGTGTATA-AAGACATA-TACACACATTCAT | loxF8L | 4 | loxF8L | 3 | AAGACATA/TATGTCTT | 7 |
| --- | | | | | | | | 0 |
| chr11 | 5222535 | CTAAGATTATATA-CATGTCTT-CCCATATTTTTAT | loxF8R | 3 | loxF8L | 4 | AAGACATG/CATGTCTT | 7 |
| chr11 | 74420250 | CTAACTTTCTGTT-CATGTCTT-TCCCCATTTTTAT | loxF8R | 4 | loxF8L | 3 | AAGACATG/CATGTCTT | 7 |
| --- | | | | | | | | 0 |
| chr18 | 22347526 | AGAAGATTGTATG-TTCCTCTT-TCCAGAGATACTT | loxF8R | 3 | loxF8L | 4 | AAGAGGAA/TTCCTCTT | 7 |
| chr13 | 22550099 | TTGGGATTGTGTG-AAGAGGAA-CACATTAGCTTTG | loxF8R | 3 | loxF8R | 4 | AAGAGGAA/TTCCTCTT | 7 |
| --- | | | | | | | | 0 |
| chr15 | 22634119 | GTAAGATTTTATG-ATTATCTT-TCACCAGATTAAA | loxF8R | 3 | loxF8L | 4 | AAGATAAT/ATTATCTT | 7 |
| chr15 | 32567107 | TTTAATCTGGTGA-AAGATAAT-CATAAAATCTTAC | loxF8L | 4 | loxF8R | 3 | AAGATAAT/ATTATCTT | 7 |
| chr15 | 30440593 | GTAAGATTTTATG-ATTATCTT-TCACCAGATTAAA | loxF8R | 3 | loxF8L | 4 | AAGATAAT/ATTATCTT | 7 |
| chr13 | 92068602 | GTAAGATTTTATG-ATTATCTT-TCATCAGATTAAA | loxF8R | 3 | loxF8L | 4 | AAGATAAT/ATTATCTT | 7 |
| --- | | | | | | | | 0 |
| chr16 | 35104486 | TTAACATTTTGTG-TATATCTT-TCCAGACTTTTGT | loxF8R | 3 | loxF8L | 4 | AAGATATA/TATATCTT | 7 |
| chr4 | 25927073 | AGAAATCTTTTGA-TATATCTT-GAGAGAATCTTGG | loxF8L | 3 | loxF8R | 4 | AAGATATA/TATATCTT | 7 |
| chr6 | 97193600 | CCAAAATTATCTG-TATATCTT-TCTACAAATTTAC | loxF8R | 4 | loxF8L | 3 | AAGATATA/TATATCTT | 7 |
| --- | | | | | | | | 0 |
| chr10 | 45938720 | ATAAATCATGGGC-AAGATTAA-AACACAGAATTAT | loxF8L | 4 | loxF8L | 3 | AAGATTAA/TTAATCTT | 7 |
| chr10 | 49907814 | ATAAATCATGGGC-AAGATTAA-AACACAGAATTAT | loxF8L | 4 | loxF8L | 3 | AAGATTAA/TTAATCTT | 7 |
| --- | | | | | | | | 0 |
| chr4 | 172204061 | CTAAGTTAATGTG-AAGATTCT-ACAATATATTTAT | loxF8R | 3 | loxF8L | 4 | AAGATTCT/AGAATCTT | 7 |
| chr9 | 73278768 | CTGAGATTATATG-AGAATCTT-TGCACAAAGTAAT | loxF8R | 3 | loxF8L | 4 | AAGATTCT/AGAATCTT | 7 |
| --- | | | | | | | | 0 |
| chr6 | 134882757 | ACCAAACTGCGGA-AAGATTTC-TCCGCAGGTTAAT | loxF8L | 4 | loxF8L | 3 | AAGATTTC/GAAATCTT | 7 |
| chr7 | 49570809 | CTAAGATTATCAG-AAGATTTC-TCAGCAGAAATAT | loxF8R | 3 | loxF8L | 4 | AAGATTTC/GAAATCTT | 7 |
| chr3 | 106504917 | TTTATTCTGTGGA-AAGATTTC-TTCTAAGATTAAT | loxF8L | 3 | loxF8L | 4 | AAGATTTC/GAAATCTT | 7 |
| --- | | | | | | | | 0 |
| chr15 | 21021254 | CTCACAGTGTGTG-TTGAGCTT-TGCACCCTTTTAT | loxF8R | 3 | loxF8L | 4 | AAGCTCAA/TTGAGCTT | 7 |
| chr15 | 20014606 | CTCACAGTGTGTG-TTGAGCTT-TGCACCCTTTTAT | loxF8R | 3 | loxF8L | 4 | AAGCTCAA/TTGAGCTT | 7 |
| --- | | | | | | | | 0 |
| chrX | 78959795 | CCAAGATTATATG-AAGGAAAA-TACTCTATTTTAG | loxF8R | 3 | loxF8R | 4 | AAGGAAAA/TTTTCCTT | 7 |
| chr13 | 22659722 | TGAAGTTTGTGGG-TTTTCCTT-TACACAGTTTTAA | loxF8R | 4 | loxF8L | 3 | AAGGAAAA/TTTTCCTT | 7 |
| --- | | | | | | | | 0 |
| chr11 | 35331874 | CTTAGACTCTGTG-AAGGAAGG-GCCATACATCTAT | loxF8R | 3 | loxF8L | 4 | AAGGAAGG/CCTTCCTT | 7 |
| chr10 | 92106354 | CAAACATGGTCTG-AAGGAAGG-CACACCATTTTAA | loxF8R | 4 | loxF8R | 3 | AAGGAAGG/CCTTCCTT | 7 |
| --- | | | | | | | | 0 |
| chr3 | 102089690 | ATGAAGCTGTGAA-AATGCCTT-CAGACAATGTTTT | loxF8L | 3 | loxF8R | 4 | AAGGCATT/AATGCCTT | 7 |
| chr1 | 153168646 | ATGATCCTATGGA-AAGGCATT-GATACATTCTTAG | loxF8L | 4 | loxF8R | 3 | AAGGCATT/AATGCCTT | 7 |
| chr1 | 240125880 | TTAATGTTGTGTG-AAGGCATT-TCCTCAGAATTGG | loxF8R | 3 | loxF8L | 4 | AAGGCATT/AATGCCTT | 7 |
| --- | | | | | | | | 0 |
| chr11 | 30539222 | TTCATATTGTGTG-AAGGCCCT-CACACTATATTTT | loxF8R | 3 | loxF8R | 4 | AAGGCCCT/AGGGCCTT | 7 |
| chrX | 39967611 | CTCAGACTGAGTC-AGGGCCTT-TACACCGTCTTAG | loxF8R | 4 | loxF8R | 3 | AAGGCCCT/AGGGCCTT | 7 |
| --- | | | | | | | | 0 |
| chr11 | 84516 | ATACATCTGATGA-AAGGCTAA-TCTCCAAATATAT | loxF8L | 3 | loxF8L | 4 | AAGGCTAA/TTAGCCTT | 7 |
| chr15 | 101878996 | ATATATTTGGAGA-TTAGCCTT-TCATCAGATGTAT | loxF8L | 4 | loxF8L | 3 | AAGGCTAA/TTAGCCTT | 7 |
| chr19 | 154146 | ATACATCTGATGA-AAGGCTAA-TCTCCAAATATAT | loxF8L | 3 | loxF8L | 4 | AAGGCTAA/TTAGCCTT | 7 |
| chr1 | 448603 | ATACATCTGATGA-AAGGCTAA-TCTCCAAATATAT | loxF8L | 3 | loxF8L | 4 | AAGGCTAA/TTAGCCTT | 7 |
| chr1 | 683579 | ATACATCTGATGA-AAGGCTAA-TCTCCAAATATAT | loxF8L | 3 | loxF8L | 4 | AAGGCTAA/TTAGCCTT | 7 |
| chr5 | 181370327 | ATATATTTGGAGA-TTAGCCTT-TCATCAGATGTAT | loxF8L | 4 | loxF8L | 3 | AAGGCTAA/TTAGCCTT | 7 |
| chr6 | 170642641 | ATATATTTGGAGA-TTAGCCTT-TCATCAGATGTAT | loxF8L | 4 | loxF8L | 3 | AAGGCTAA/TTAGCCTT | 7 |
| chr8 | 163953 | ATACATCTGATGA-AAGGCTAA-TCTCCAAATATAT | loxF8L | 3 | loxF8L | 4 | AAGGCTAA/TTAGCCTT | 7 |
| chr9 | 138322933 | ATATATTTGGAGA-TTAGCCTT-TCATCAGATGTAT | loxF8L | 4 | loxF8L | 3 | AAGGCTAA/TTAGCCTT | 7 |
| --- | | | | | | | | 0 |
| chr4 | 161448028 | AAGAAACTGTGGA-AAGGGAAG-GTCACAATTTCAG | loxF8L | 3 | loxF8R | 4 | AAGGGAAG/CTTCCCTT | 7 |
| chr8 | 136013304 | TGAAGATTTTGTG-CTTCCCTT-CACACCTTATTGG | loxF8R | 3 | loxF8R | 4 | AAGGGAAG/CTTCCCTT | 7 |
| --- | | | | | | | | 0 |
| chr8 | 55045839 | GTCAAGCTGTGGA-AAGTCAAA-TACAAAGAATTCT | loxF8L | 3 | loxF8L | 4 | AAGTCAAA/TTTGACTT | 7 |
| chr12 | 44719936 | CTAAGATTCTGCA-AAGTCAAA-CAAAAAATTTTAA | loxF8R | 3 | loxF8R | 4 | AAGTCAAA/TTTGACTT | 7 |
| --- | | | | | | | | 0 |
| chr2 | 37926936 | AAAAAGCTATGTA-AATGACTT-CATACAACCTTTG | loxF8L | 4 | loxF8R | 3 | AAGTCATT/AATGACTT | 7 |
| chr6 | 18725343 | CTATGACTGTTTA-AAGTCATT-TCCACAGTTAGAT | loxF8R | 4 | loxF8L | 3 | AAGTCATT/AATGACTT | 7 |
| --- | | | | | | | | 0 |
| chr5 | 119477566 | GAAAGATTATGTG-AAGTGTTG-TGAAATGATTTAT | loxF8R | 3 | loxF8L | 4 | AAGTGTTG/CAACACTT | 7 |
| chr3 | 106420852 | TTACATATGTTGA-CAACACTT-AACACAATATTAT | loxF8L | 4 | loxF8R | 3 | AAGTGTTG/CAACACTT | 7 |
| --- | | | | | | | | 0 |
| chr21 | 42303989 | ATAATGCTGTGTG-GATAACTT-TTCACTGATTTAG | loxF8L | 4 | loxF8L | 3 | AAGTTATC/GATAACTT | 7 |
| chr14 | 28793282 | CAAAATATGTGCA-AAGTTATC-TTAACAGATTAAT | loxF8L | 4 | loxF8L | 3 | AAGTTATC/GATAACTT | 7 |
| --- | | | | | | | | 0 |

| | | | | | | | | |
|---|---|---|---|---|---|---|---|---|
| chr7 | 153626935 | CAGAGATTTTGTT-AAGTTGTG-TCCCTATATTTAT | loxF8R | 4 | loxF8L | 3 | AAGTTGTG/CACAACTT | 7 |
| chr6 | 1067771 | CAAAGATTTTGGA-AAGTTGTG-TCCCTATATTTAT | loxF8R | 4 | loxF8L | 3 | AAGTTGTG/CACAACTT | 7 |
| --- | | | | | | | | 0 |
| chr7 | 78400519 | ATCAGATCTTGTG-TTTTTATT-CACACAATTTTTT | loxF8R | 4 | loxF8R | 3 | AATAAAAA/TTTTTATT | 7 |
| chr13 | 39598216 | ATAAGTTTTTGTG-AATAAAAA-CCCAAACAATTAT | loxF8R | 3 | loxF8L | 4 | AATAAAAA/TTTTTATT | 7 |
| chr9 | 68543752 | CTAAATCTGTACA-AATAAAAA-CAAACAAATTTAA | loxF8L | 3 | loxF8R | 4 | AATAAAAA/TTTTTATT | 7 |
| chr6 | 130963770 | CTAATATTGGGTA-TTTTTATT-CACATAACCATAA | loxF8R | 3 | loxF8R | 4 | AATAAAAA/TTTTTATT | 7 |
| --- | | | | | | | | 0 |
| chr6 | 94653126 | CTTAAATTTTGTG-CAGTTATT-TAAACTGTTTTAT | loxF8R | 3 | loxF8L | 4 | AATAACTG/CAGTTATT | 7 |
| chr4 | 86061403 | CTAATAATTTGAG-CAGTTATT-TCAAAATATTTAT | loxF8R | 4 | loxF8L | 3 | AATAACTG/CAGTTATT | 7 |
| --- | | | | | | | | 0 |
| chr3 | 119284105 | CTAGGATTTTGAG-ATCTTATT-CACAAAATATTTT | loxF8R | 3 | loxF8R | 4 | AATAAGAT/ATCTTATT | 7 |
| chr17 | 12170007 | TTAAGAGTATGTT-ATCTTATT-TTCACATATTTGT | loxF8R | 4 | loxF8L | 3 | AATAAGAT/ATCTTATT | 7 |
| --- | | | | | | | | 0 |
| chr4 | 96064336 | ATAAAAATGTGAT-AATAATAA-TTAACATATTTAT | loxF8L | 4 | loxF8L | 3 | AATAATAA/TTATTATT | 7 |
| chr8 | 32039161 | GTAGAATTGTGTT-AATAATAA-TCCACAGGCTTAC | loxF8R | 4 | loxF8L | 3 | AATAATAA/TTATTATT | 7 |
| --- | | | | | | | | 0 |
| chr1 | 98894394 | TAAGAACTGTGGA-AATAATAC-ATCACAGTTTTAT | loxF8L | 4 | loxF8L | 3 | AATAATAC/GTATTATT | 7 |
| chr13 | 68113257 | CAAAGATTTCTTG-AATAATAC-CACACAAGCACAG | loxF8R | 4 | loxF8R | 3 | AATAATAC/GTATTATT | 7 |
| chr2 | 46603797 | CTAAGATTTCTTG-AATAATAC-CCCACAAGCACAG | loxF8R | 3 | loxF8R | 4 | AATAATAC/GTATTATT | 7 |
| chr3 | 122871413 | CTGTGAGTGTGTG-AATAATAC-ATAACAGATTTAA | loxF8R | 3 | loxF8L | 4 | AATAATAC/GTATTATT | 7 |
| --- | | | | | | | | 0 |
| chr9 | 20150675 | AAAAATCCCTGGA-AATAATTG-TATTCAGCTTTAT | loxF8L | 3 | loxF8L | 4 | AATAATTG/CAATTATT | 7 |
| chr10 | 27611753 | TTTGAATTGTGTG-AATAATTG-TGTACATATTTAT | loxF8R | 4 | loxF8L | 3 | AATAATTG/CAATTATT | 7 |
| --- | | | | | | | | 0 |
| chr13 | 22616399 | ATACATGTTTGTA-GTTGTATT-TGCAGACATTTAT | loxF8L | 4 | loxF8L | 3 | AATACAAC/GTTGTATT | 7 |
| chr3 | 177107807 | CTGAGATTATCTG-AATACAAC-TACTCCATCTTAC | loxF8R | 3 | loxF8R | 4 | AATACAAC/GTTGTATT | 7 |
| --- | | | | | | | | 0 |
| chr3 | 121018407 | ATAGATCTGTATA-AATACAAT-CAACTAATCTTTG | loxF8L | 3 | loxF8R | 4 | AATACAAT/ATTGTATT | 7 |
| chr3 | 171574049 | TTTAATCTGTAGA-AATACAAT-TACACATATTTTC | loxF8L | 3 | loxF8L | 4 | AATACAAT/ATTGTATT | 7 |
| --- | | | | | | | | 0 |
| chr4 | 110982289 | ATAAATGAGTAGA-AATACAGA-CACACAAACCTGT | loxF8L | 3 | loxF8R | 4 | AATACAGA/TCTGTATT | 7 |
| chrX | 91059538 | CTGAGATTGTGGA-AATACAGA-CACAAGCTCTTAT | loxF8R | 3 | loxF8R | 4 | AATACAGA/TCTGTATT | 7 |
| chr5 | 128404180 | TTCAATCTGAGAA-AATACAGA-CTCAGAATTTTAG | loxF8L | 4 | loxF8R | 3 | AATACAGA/TCTGTATT | 7 |
| --- | | | | | | | | 0 |
| chr13 | 56383021 | AGATTTCTGTGGG-AATACAGC-CAAACCATCTCAG | loxF8L | 4 | loxF8R | 3 | AATACAGC/GCTGTATT | 7 |
| chr9 | 42387447 | ATACCTCTGTGAA-GCTGTATT-TCTATCCATTTAT | loxF8L | 3 | loxF8L | 4 | AATACAGC/GCTGTATT | 7 |
| --- | | | | | | | | 0 |
| chr1 | 163605457 | AGAAATATGTGTA-TATGTATT-TGTATACATTTAT | loxF8L | 3 | loxF8L | 4 | AATACATA/TATGTATT | 7 |
| chr4 | 58801647 | ATATATATATGAA-TATGTATT-TACATATATTTAT | loxF8L | 4 | loxF8L | 3 | AATACATA/TATGTATT | 7 |
| chr21 | 39052134 | ATATGTGTGTGTG-TATGTATT-CACACATTCTGAT | loxF8R | 4 | loxF8R | 3 | AATACATA/TATGTATT | 7 |
| --- | | | | | | | | 0 |
| chr5 | 67928375 | ATAAATATTTGCA-AATACATT-TCCCCAGACTTTG | loxF8L | 3 | loxF8L | 4 | AATACATT/AATGTATT | 7 |
| chr4 | 41517013 | GTAACACTGTGGA-AATGTATT-TCCACTCTTGTAT | loxF8L | 3 | loxF8L | 4 | AATACATT/AATGTATT | 7 |
| chr12 | 102981708 | CTATGGTAGTGGG-AATGTATT-TACCCAGATCTAT | loxF8R | 4 | loxF8L | 3 | AATACATT/AATGTATT | 7 |
| --- | | | | | | | | 0 |
| chr12 | 72626896 | CCAAGTTGGTGTG-GTCCTATT-TCCTCAGTTCTTT | loxF8R | 3 | loxF8L | 4 | AATAGGAC/GTCCTATT | 7 |
| chr9 | 29826983 | ATGTATCTGTGGT-GTCCTATT-TTCTCAGGTTCAT | loxF8L | 3 | loxF8L | 4 | AATAGGAC/GTCCTATT | 7 |
| --- | | | | | | | | 0 |
| chr2 | 224676138 | AAAAATCTTTGAA-AATATAAA-ACAAAAGATTTTT | loxF8L | 3 | loxF8L | 4 | AATATAAA/TTTATATT | 7 |
| chr18 | 68396832 | AAAAATTGGTGGC-AATATAAA-TTCACATATTTAG | loxF8L | 4 | loxF8L | 3 | AATATAAA/TTTATATT | 7 |
| chr2 | 213490563 | ATGAATCTGCGTA-TTTATATT-CATACACATTTAT | loxF8L | 3 | loxF8L | 4 | AATATAAA/TTTATATT | 7 |
| --- | | | | | | | | 0 |
| chrX | 6685182 | AGTAATGTGAGGA-AATATACA-AACACCATCTTAT | loxF8L | 4 | loxF8R | 3 | AATATACA/TGTATATT | 7 |
| chr15 | 35857763 | TAAAGCATGTGTG-AATATACA-TTCTCATATTTAT | loxF8R | 4 | loxF8L | 3 | AATATACA/TGTATATT | 7 |
| --- | | | | | | | | 0 |
| chr4 | 55787473 | ATACATCTGTACA-AATATAGA-AACATAATTTTAT | loxF8L | 3 | loxF8R | 4 | AATATAGA/TCTATATT | 7 |
| chr3 | 133050077 | GAAAAGATGTGGA-AATATAGA-CACACAATTTTTC | loxF8L | 4 | loxF8R | 3 | AATATAGA/TCTATATT | 7 |
| --- | | | | | | | | 0 |
| chr4 | 44764263 | ATAAATATGTCTC-AATATATT-TCAAAAGATTGAT | loxF8L | 4 | loxF8L | 3 | AATATATT/AATATATT | 7 |
| --- | | | | | | | | 0 |
| chr6 | 161692964 | AGAAAAAAGTGGA-TGGATATT-ACCACATATTTTT | loxF8L | 4 | loxF8L | 3 | AATATCCA/TGGATATT | 7 |
| chr11 | 30662375 | TTAAGCTTTTGTG-AATATCCA-CACATACTGTTTG | loxF8R | 3 | loxF8R | 4 | AATATCCA/TGGATATT | 7 |
| --- | | | | | | | | 0 |
| chr2 | 52425650 | AAATATCTGGGAA-CAGATATT-TACAAATATTTAT | loxF8L | 4 | loxF8L | 3 | AATATCTG/CAGATATT | 7 |
| chr3 | 42202180 | CTTAGTCTGTGTG-CAGATATT-CATCCCACCTTAG | loxF8R | 3 | loxF8R | 4 | AATATCTG/CAGATATT | 7 |
| --- | | | | | | | | 0 |
| chr9 | 14143284 | CCTAGATTTTGTG-TTAATATT-TTGCCTGATTTAT | loxF8R | 3 | loxF8L | 4 | AATATTAA/TTAATATT | 7 |
| chr9 | 39502276 | ATATATATATGGG-TTAATATT-CAGACAATCTAAA | loxF8L | 4 | loxF8R | 3 | AATATTAA/TTAATATT | 7 |
| chr9 | 66863002 | TTTAGATTGTCTG-AATATTAA-CCCATATATATAT | loxF8R | 3 | loxF8L | 4 | AATATTAA/TTAATATT | 7 |
| --- | | | | | | | | 0 |
| chr15 | 98040269 | ATGAATCTAAGAA-AATATTAT-TATATAGATTTAT | loxF8L | 4 | loxF8L | 3 | AATATTAT/ATAATATT | 7 |
| chr3 | 175492750 | ATACAGCTGTGTA-AATATTAT-TCCACTAATTTTA | loxF8L | 3 | loxF8L | 4 | AATATTAT/ATAATATT | 7 |
| --- | | | | | | | | 0 |
| chr3 | 114740815 | ATTAATATCTGAA-TGAATATT-TCCATTGATTTGT | loxF8L | 4 | loxF8L | 3 | AATATTCA/TGAATATT | 7 |
| chr11 | 97699305 | CAAATATTGTCTA-AATATTCA-CACACATTCATAT | loxF8R | 4 | loxF8R | 3 | AATATTCA/TGAATATT | 7 |
| --- | | | | | | | | 0 |
| chr5 | 147983260 | ATACATTTGTTCA-TAAATATT-TCCATACAGTTAT | loxF8L | 4 | loxF8L | 3 | AATATTTA/TAAATATT | 7 |
| chr7 | 19830232 | ATGAATCTGTTAA-AATATTTA-TACTCAAATATAT | loxF8L | 3 | loxF8L | 4 | AATATTTA/TAAATATT | 7 |
| chr7 | 138464627 | TTAAAAATGTGTG-AATATTTA-TACACATAAATAT | loxF8R | 3 | loxF8L | 4 | AATATTTA/TAAATATT | 7 |
| --- | | | | | | | | 0 |
| chr2 | 87846528 | ATAAATATGTATA-AATATTTC-TCAACAATTTTTT | loxF8L | 3 | loxF8L | 4 | AATATTTC/GAAATATT | 7 |
| chrX | 144191940 | AAAAAATTGTTGA-GAAATATT-TATACATATTTAT | loxF8L | 4 | loxF8L | 3 | AATATTTC/GAAATATT | 7 |
| chr10 | 53580832 | ATAAATATCTGTA-AATATTTC-TACATATATATAT | loxF8L | 3 | loxF8L | 4 | AATATTTC/GAAATATT | 7 |
| --- | | | | | | | | 0 |
| chr14 | 36125196 | AAAAGTCTGGAGA-AATCATCT-TTCCCAGATTTAA | loxF8L | 4 | loxF8L | 3 | AATCATCT/AGATGATT | 7 |
| chr6 | 119922506 | ATTAATCTTTGTA-AATCATCT-CACTCCCTCTTGG | loxF8L | 3 | loxF8R | 4 | AATCATCT/AGATGATT | 7 |

| | | | | | | | | |
|---|---|---|---|---|---|---|---|---|
| --- | | | | | | | | 0 |
| chr13 | 103585013 | ATCACCCTGGGGA-TTAGGATT-TCCACCTATTTTT | loxF8L | 4 | loxF8L | 3 | AATCCTAA/TTAGGATT | 7 |
| chr4 | 16593099 | ATATTTCTGTTGA-TTAGGATT-GCGACTGATTTTT | loxF8L | 3 | loxF8L | 4 | AATCCTAA/TTAGGATT | 7 |
| chr2 | 189293511 | AGAATTGTGTTGA-AATCCTAA-CCCACAAGGTTAG | loxF8L | 4 | loxF8R | 3 | AATCCTAA/TTAGGATT | 7 |
| chr3 | 168719150 | ATTCATATGTTGA-AATCCTAA-CCCCCAGGTTTAT | loxF8L | 4 | loxF8L | 3 | AATCCTAA/TTAGGATT | 7 |
| chr10 | 9578937 | ATTCATATGTGGA-AATCCTAA-CCCCCAAAATTAT | loxF8L | 3 | loxF8L | 4 | AATCCTAA/TTAGGATT | 7 |
| chr3 | 56523765 | ATTCATATGTTGA-AATCCTAA-CACACAATGTGAT | loxF8L | 4 | loxF8R | 3 | AATCCTAA/TTAGGATT | 7 |
| chr4 | 149504506 | ATTCATATGTTGA-AATCCTAA-CCCACAATGTTAC | loxF8L | 4 | loxF8R | 3 | AATCCTAA/TTAGGATT | 7 |
| chr4 | 153680389 | CTCACATTGGGGG-TTAGGATT-TCAACAGATGAAT | loxF8R | 4 | loxF8L | 3 | AATCCTAA/TTAGGATT | 7 |
| --- | | | | | | | | 0 |
| chr3 | 18411810 | ATAAATTAGTGTC-AATCGATT-TGGGCAGATTTAT | loxF8L | 4 | loxF8L | 3 | AATCGATT/AATCGATT | 7 |
| --- | | | | | | | | 0 |
| chr7 | 144965649 | TGAAGATTTTGTG-AATCTGGT-AACACAATCCTTT | loxF8R | 3 | loxF8R | 4 | AATCTGGT/ACCAGATT | 7 |
| chr11 | 64041157 | ATTAATCGTTGGC-ACCAGATT-ACAACAGATTTAG | loxF8L | 4 | loxF8L | 3 | AATCTGGT/ACCAGATT | 7 |
| --- | | | | | | | | 0 |
| chr2 | 186186076 | ACAAAATTGTGTA-AATGAAAT-GACACACTCTTAA | loxF8L | 4 | loxF8R | 3 | AATGAAAT/ATTTCATT | 7 |
| chr15 | 70325587 | ATAATTCTGGGCA-AATGAAAT-CACAGATTATTAA | loxF8L | 3 | loxF8R | 4 | AATGAAAT/ATTTCATT | 7 |
| chr9 | 22708594 | ATTAATCTGTACA-AATGAAAT-CATACAATATGTG | loxF8L | 3 | loxF8R | 4 | AATGAAAT/ATTTCATT | 7 |
| --- | | | | | | | | 0 |
| chr11 | 81103111 | ATAAATTTGAAGA-AATGCAAA-CACACAAACATGT | loxF8L | 3 | loxF8R | 4 | AATGCAAA/TTTGCATT | 7 |
| chr20 | 14742429 | ATAAAGGTCTGGA-AATGCAAA-CAGTCAAATTTAG | loxF8L | 3 | loxF8R | 4 | AATGCAAA/TTTGCATT | 7 |
| --- | | | | | | | | 0 |
| chr17 | 33003390 | ATATTTTTGTGGA-AATGCATT-TCCTCAGACCTTT | loxF8L | 3 | loxF8L | 4 | AATGCATT/AATGCATT | 7 |
| --- | | | | | | | | 0 |
| chr11 | 46029191 | TTAATGTTGTGGA-AATGCTCT-TTCACTGATTAAT | loxF8L | 4 | loxF8L | 3 | AATGCTCT/AGAGCATT | 7 |
| chr6 | 83455275 | CTAACTTTGTGAG-AGAGCATT-TCCAAAATTTTAA | loxF8R | 3 | loxF8L | 4 | AATGCTCT/AGAGCATT | 7 |
| --- | | | | | | | | 0 |
| chr12 | 118268401 | AAAAAACTATGGA-CTTCCATT-TTCACATTTTTCT | loxF8L | 3 | loxF8L | 4 | AATGGAAG/CTTCCATT | 7 |
| chr4 | 162649911 | ACAAATCTGTTCA-AATGGAAG-TATACATCTTTAT | loxF8L | 3 | loxF8L | 4 | AATGGAAG/CTTCCATT | 7 |
| --- | | | | | | | | 0 |
| chr2 | 102657230 | ATAAAATTGTGTT-ATTCCATT-TCCACAGTTCCAG | loxF8R | 3 | loxF8L | 4 | AATGGAAT/ATTCCATT | 7 |
| chr13 | 43032459 | ATTATTTTGTGGA-ATTCCATT-TACACTGTCTTAT | loxF8L | 3 | loxF8L | 4 | AATGGAAT/ATTCCATT | 7 |
| --- | | | | | | | | 0 |
| chr5 | 170904693 | CTAAAATTGAGAG-AATGGAGT-TCTTCAGTATTAT | loxF8R | 3 | loxF8L | 4 | AATGGAGT/ACTCCATT | 7 |
| chr11 | 88074388 | CTAAATATGGGGA-AATGGAGT-TACAGAGTTTTAA | loxF8L | 3 | loxF8L | 4 | AATGGAGT/ACTCCATT | 7 |
| --- | | | | | | | | 0 |
| chr3 | 39688365 | ATAAAAATGGAGA-AATGGGAA-CACACAAGATTAA | loxF8L | 4 | loxF8R | 3 | AATGGGAA/TTCCCATT | 7 |
| chr11 | 1066124 | ATAAATAATTGGA-AATGGGAA-CAAACAATATCAT | loxF8L | 3 | loxF8R | 4 | AATGGGAA/TTCCCATT | 7 |
| --- | | | | | | | | 0 |
| chr10 | 91152396 | CAAAAATTCTGTG-ATCCCATT-TCCCCGTATTTGT | loxF8R | 3 | loxF8L | 4 | AATGGGAT/ATCCCATT | 7 |
| chr5 | 21464281 | CTAACACAGTGAG-ATCCCATT-CAAAAAATCTTAT | loxF8R | 4 | loxF8R | 3 | AATGGGAT/ATCCCATT | 7 |
| chr5 | 37385331 | CTAACACAGTGAG-ATCCCATT-CAAAAAATCTTAC | loxF8R | 4 | loxF8R | 3 | AATGGGAT/ATCCCATT | 7 |
| --- | | | | | | | | 0 |
| chr3 | 166700630 | AAAAATGTGTGAA-TTTACATT-CCCACGAACTGAG | loxF8L | 3 | loxF8R | 4 | AATGTAAA/TTTACATT | 7 |
| chr10 | 126450529 | ATACATCTATGTA-TTTACATT-CAACTAATCTTTG | loxF8L | 3 | loxF8R | 4 | AATGTAAA/TTTACATT | 7 |
| --- | | | | | | | | 0 |
| chr6 | 158126079 | ATATATGTGTACA-TATACATT-TACACATATTTGT | loxF8L | 4 | loxF8L | 3 | AATGTATA/TATACATT | 7 |
| chr6 | 93554347 | ATAATTATTTGTA-AATGTATA-CACACACATTTAG | loxF8L | 4 | loxF8R | 3 | AATGTATA/TATACATT | 7 |
| chr3 | 85092334 | ATAATTCACTGGA-AATGTATA-CATAAAATTTAAG | loxF8L | 3 | loxF8R | 4 | AATGTATA/TATACATT | 7 |
| chr2 | 192427272 | ATTAATACATGGA-AATGTATA-ACCACAGAATAAT | loxF8L | 4 | loxF8L | 3 | AATGTATA/TATACATT | 7 |
| --- | | | | | | | | 0 |
| chr2 | 122424752 | TTAAATCTATTCA-TTCACATT-TGCAAACATTTAT | loxF8L | 4 | loxF8L | 3 | AATGTGAA/TTCACATT | 7 |
| chr1 | 81959704 | ATAACAAAGTGTG-TTCACATT-TTCATAAATTTAT | loxF8R | 4 | loxF8L | 3 | AATGTGAA/TTCACATT | 7 |
| chr14 | 101446371 | CAAAATATGTTGA-TTCACATT-ACCACTGACTTAT | loxF8L | 4 | loxF8L | 3 | AATGTGAA/TTCACATT | 7 |
| --- | | | | | | | | 0 |
| chr1 | 31807073 | AAATATGTTTGGA-AATTAAGA-AATACACTCTTAG | loxF8L | 4 | loxF8R | 3 | AATTAAGA/TCTTAATT | 7 |
| chr5 | 24539021 | TTAAAACTGTAGA-TCTTAATT-TCTACAAAAATAT | loxF8L | 3 | loxF8L | 4 | AATTAAGA/TCTTAATT | 7 |
| --- | | | | | | | | 0 |
| chr3 | 198083525 | ACAAATATGTGGG-AATTACAG-AATACACTCTTAA | loxF8L | 3 | loxF8R | 4 | AATTACAG/CTGTAATT | 7 |
| chrX | 133572202 | ACTAATATGTGGA-AATTACAG-AATACAATCCTAC | loxF8L | 3 | loxF8R | 4 | AATTACAG/CTGTAATT | 7 |
| --- | | | | | | | | 0 |
| chr11 | 55953873 | CTAAATATGAGTA-TTATAATT-TCTTCAGATTTTT | loxF8L | 4 | loxF8L | 3 | AATTATAA/TTATAATT | 7 |
| chr2 | 139807037 | ATATTTCTGTTCA-TTATAATT-CACCCAGTCTTTG | loxF8L | 4 | loxF8R | 3 | AATTATAA/TTATAATT | 7 |
| --- | | | | | | | | 0 |
| chr3 | 162329385 | TTAAGAATGTGTT-TCATAATT-TCCATATATCTGT | loxF8R | 3 | loxF8L | 4 | AATTATGA/TCATAATT | 7 |
| chr2 | 108160388 | ATAAATATGTCAT-TCATAATT-AACACAGATTTAA | loxF8L | 4 | loxF8L | 3 | AATTATGA/TCATAATT | 7 |
| chr14 | 48178737 | ATATATCTGTCTA-TCATAATT-TGCAAATATATAT | loxF8L | 3 | loxF8L | 4 | AATTATGA/TCATAATT | 7 |
| --- | | | | | | | | 0 |
| chr5 | 110316721 | ATAAGAGCTTGTG-TCTGAATT-TCCACAATTTTAG | loxF8R | 4 | loxF8L | 3 | AATTCAGA/TCTGAATT | 7 |
| chr14 | 29363104 | GGCAGATTGTTTG-AATTCAGA-CACACAATTTGCG | loxF8R | 4 | loxF8R | 3 | AATTCAGA/TCTGAATT | 7 |
| --- | | | | | | | | 0 |
| chr10 | 43935710 | ATAAATTTGGGGG-ATAGAATT-CAATCCATCGTAG | loxF8L | 3 | loxF8R | 4 | AATTCTAT/ATAGAATT | 7 |
| chr18 | 43533560 | ATAAGTTGGTGCA-AATTCTAT-GAAACAATGTTAG | loxF8L | 4 | loxF8R | 3 | AATTCTAT/ATAGAATT | 7 |
| --- | | | | | | | | 0 |
| chr5 | 24665545 | CTGAGATTGCATC-CCAGAATT-TCCTAAGATTTTT | loxF8R | 4 | loxF8L | 3 | AATTCTGG/CCAGAATT | 7 |
| chr11 | 107022747 | ATAAAGTTATGGA-AATTCTGG-CACAGAGATTTAA | loxF8L | 3 | loxF8L | 4 | AATTCTGG/CCAGAATT | 7 |
| --- | | | | | | | | 0 |
| chr10 | 130910105 | TTAGAAGTGTGTG-GCTCAATT-TCCACACGTTTGT | loxF8R | 4 | loxF8L | 3 | AATTGAGC/GCTCAATT | 7 |
| chr11 | 34585719 | CTATAATTACGTG-GCTCAATT-TACACAATAATAG | loxF8R | 4 | loxF8R | 3 | AATTGAGC/GCTCAATT | 7 |
| --- | | | | | | | | 0 |
| chr1 | 83256117 | ATAAATATTAGGA-AATTGATG-TCTACAGACATGT | loxF8L | 3 | loxF8L | 4 | AATTGATG/CATCAATT | 7 |
| chr7 | 76720981 | ATAAATATTAGGA-AATTGATG-TCTACAGACATGT | loxF8L | 3 | loxF8L | 4 | AATTGATG/CATCAATT | 7 |
| chrX | 147372091 | CTAATTCTGTGAA-AATTGATG-TTCAGAGTTTGAT | loxF8L | 3 | loxF8L | 4 | AATTGATG/CATCAATT | 7 |
| --- | | | | | | | | 0 |
| chr19 | 6087026 | ACAAATTTGAGGA-CTCCAATT-CAGACGAGCTTCG | loxF8L | 3 | loxF8R | 4 | AATTGGAG/CTCCAATT | 7 |
| chr9 | 36991365 | CAAAAATCGTCTG-CTCCAATT-TCCACATTTTAAT | loxF8R | 4 | loxF8L | 3 | AATTGGAG/CTCCAATT | 7 |
| --- | | | | | | | | 0 |

| | | | | | | | | |
|---|---|---|---|---|---|---|---|---|
| chr4 | 81790375 | TTAAATTTGTCAA-ATACAATT-CACCTAATCTTAT | loxF8L | 4 | loxF8R | 3 | AATTGTAT/ATACAATT | 7 |
| chrX | 33034556 | TTAATTATGTGGA-ATACAATT-TCCTCAACTTTAA | loxF8L | 3 | loxF8L | 4 | AATTGTAT/ATACAATT | 7 |
| --- | | | | | | | | 0 |
| chr10 | 100909891 | TTAAATGTGTGGC-AATTTCAG-AGCAAAAATTTAT | loxF8L | 3 | loxF8L | 4 | AATTTCAG/CTGAAATT | 7 |
| chr8 | 31809263 | ATACCTCAGGGGA-AATTTCAG-TCTAAAGATGTAT | loxF8L | 4 | loxF8L | 3 | AATTTCAG/CTGAAATT | 7 |
| --- | | | | | | | | 0 |
| chr13 | 67654145 | ATGAAGCTGTTAA-AATTTCTA-TACACAGAAATAT | loxF8L | 4 | loxF8L | 3 | AATTTCTA/TAGAAATT | 7 |
| chr6 | 70841557 | TTTAATCTGGGGA-AATTTCTA-TCTAAACATTCAT | loxF8L | 3 | loxF8L | 4 | AATTTCTA/TAGAAATT | 7 |
| --- | | | | | | | | 0 |
| chr16 | 33374771 | GTATAGTTGTGTG-AATTTGAG-CCCAGTGATTTAT | loxF8R | 4 | loxF8L | 3 | AATTTGAG/CTCAAATT | 7 |
| chr16 | 33473542 | GTATAGTTGTGTG-AATTTGAG-CCCAGTGATTTAT | loxF8R | 4 | loxF8L | 3 | AATTTGAG/CTCAAATT | 7 |
| chr16 | 32662200 | ATAAATCACTGGG-CTCAAATT-CACACAACTATAC | loxF8L | 3 | loxF8R | 4 | AATTTGAG/CTCAAATT | 7 |
| chr16 | 33318232 | GTATAGTTGTGTG-AATTTGAG-CCCAGTGATTTAT | loxF8R | 4 | loxF8L | 3 | AATTTGAG/CTCAAATT | 7 |
| chr16 | 32267203 | GTATAGTTGTGTG-AATTTGAG-CCCAGTGATTTAT | loxF8R | 4 | loxF8L | 3 | AATTTGAG/CTCAAATT | 7 |
| chr16 | 33208152 | GTATAGTTGTGTG-AATTTGAG-CCCAGTGATTTAT | loxF8R | 4 | loxF8L | 3 | AATTTGAG/CTCAAATT | 7 |
| --- | | | | | | | | 0 |
| chr6 | 72141636 | ATAAATTTATTTA-AATTTTAC-CACACAATTTAAA | loxF8L | 4 | loxF8R | 3 | AATTTTAC/GTAAAATT | 7 |
| chr12 | 93583445 | CTTAGATTCTTTG-GTAAAATT-AAAAGAATCTTGG | loxF8R | 3 | loxF8R | 4 | AATTTTAC/GTAAAATT | 7 |
| --- | | | | | | | | 0 |
| chr11 | 16518149 | ATAAATGTGGCCA-AATTTTCT-TTCACAGCTTCAT | loxF8L | 4 | loxF8L | 3 | AATTTTCT/AGAAAATT | 7 |
| chr1 | 98469093 | ATAATATCGTGTG-AATTTTCT-CACATCAACTAAG | loxF8R | 3 | loxF8R | 4 | AATTTTCT/AGAAAATT | 7 |
| chr5 | 29878765 | ATTAATTTTTTGA-AATTTTCT-CACATTTTCTTAG | loxF8L | 4 | loxF8R | 3 | AATTTTCT/AGAAAATT | 7 |
| --- | | | | | | | | 0 |
| chr1 | 190004162 | AAAATTCTGTGAA-TAAAAATT-TCAACACATTTTA | loxF8L | 3 | loxF8L | 4 | AATTTTTA/TAAAAATT | 7 |
| chr7 | 31451916 | ATAATTTTATGGA-AATTTTTA-AATAAAATCTTAA | loxF8L | 3 | loxF8R | 4 | AATTTTTA/TAAAAATT | 7 |
| --- | | | | | | | | 0 |
| chr10 | 63543137 | CTAATACTGTGTA-ACAAACCA-CCCAATAACTTAG | loxF8R | 3 | loxF8R | 4 | ACAAACCA/TGGTTTGT | 7 |
| chr3 | 26312049 | GTGATATTGAGTG-ACAAACCA-CTCCCAAACTTAG | loxF8R | 4 | loxF8R | 3 | ACAAACCA/TGGTTTGT | 7 |
| --- | | | | | | | | 0 |
| chr8 | 91098648 | ATGAACCGGTGGC-TACTTTGT-TCCTCTGATTTTT | loxF8L | 4 | loxF8L | 3 | ACAAAGTA/TACTTTGT | 7 |
| chr10 | 86410939 | CTATAGCTGTGTA-ACAAAGTA-CACCCAAACTTGG | loxF8L | 4 | loxF8R | 3 | ACAAAGTA/TACTTTGT | 7 |
| --- | | | | | | | | 0 |
| chr9 | 1937159 | ATAAATATATATA-ACAAATAT-TACATATATTTAT | loxF8L | 4 | loxF8L | 3 | ACAAATAT/ATATTTGT | 7 |
| chr9 | 132379270 | ATACATGTGTGAT-ACAAATAT-GACACAATTTTTG | loxF8L | 4 | loxF8R | 3 | ACAAATAT/ATATTTGT | 7 |
| --- | | | | | | | | 0 |
| chr6 | 88804274 | ATGACTCTCTGAA-ACAAATGC-CCCAAAGATTTCT | loxF8L | 4 | loxF8L | 3 | ACAAATGC/GCATTTGT | 7 |
| chr11 | 66489595 | CTATTGCTGTGGA-ACAAATGC-TCCGCAAATTTAG | loxF8L | 4 | loxF8L | 3 | ACAAATGC/GCATTTGT | 7 |
| --- | | | | | | | | 0 |
| chr8 | 108767545 | ACAGATCTGTGTG-ACAAATTC-TCCATAGAATTTT | loxF8L | 4 | loxF8L | 3 | ACAAATTC/GAATTTGT | 7 |
| chr15 | 66098062 | CTAAGTTTGAGAG-GAATTTGT-TCCACAAATGCAG | loxF8R | 3 | loxF8L | 4 | ACAAATTC/GAATTTGT | 7 |
| --- | | | | | | | | 0 |
| chr2 | 61730245 | ATAAATCCCTGAA-ACAACTGG-CAAATAATTTGAG | loxF8L | 3 | loxF8R | 4 | ACAACTGG/CCAGTTGT | 7 |
| chr7 | 152933674 | ATAACTCTTTCAA-CCAGTTGT-CAGAAAATCTTTG | loxF8L | 4 | loxF8R | 3 | ACAACTGG/CCAGTTGT | 7 |
| --- | | | | | | | | 0 |
| chr4 | 20872858 | CTAAGCTTGTTTA-ATTCTTGT-TTCAAAAATTTGT | loxF8R | 3 | loxF8L | 4 | ACAAGAAT/ATTCTTGT | 7 |
| chr13 | 54749148 | ATAAATGCATGGT-ACAAGAAT-TAAACAAATTTAT | loxF8L | 4 | loxF8L | 3 | ACAAGAAT/ATTCTTGT | 7 |
| --- | | | | | | | | 0 |
| chr7 | 120502915 | AATAATCTCTGCA-TATCTTGT-ACCTCAGATTTAC | loxF8L | 4 | loxF8L | 3 | ACAAGATA/TATCTTGT | 7 |
| chr3 | 197361580 | CTAAGTTTGGGGG-TATCTTGT-TACACAGCATTAG | loxF8R | 3 | loxF8L | 4 | ACAAGATA/TATCTTGT | 7 |
| --- | | | | | | | | 0 |
| chr8 | 17620890 | CTAGGGGTGTTTG-ACAAGTCA-AACACATTCTTAT | loxF8R | 4 | loxF8R | 3 | ACAAGTCA/TGACTTGT | 7 |
| chrX | 39527641 | CTAAGTTTGGGAA-TGACTTGT-TACATAATCATAG | loxF8R | 4 | loxF8R | 3 | ACAAGTCA/TGACTTGT | 7 |
| --- | | | | | | | | 0 |
| chr9 | 17278810 | ATAAAATTCTGTG-AGCATTGT-TCTACATTTTTCT | loxF8R | 3 | loxF8L | 4 | ACAATGCT/AGCATTGT | 7 |
| chr11 | 19689159 | CAAAGGCTGTGTG-ACAATGCT-GGGACAGAGTTAT | loxF8R | 3 | loxF8L | 4 | ACAATGCT/AGCATTGT | 7 |
| --- | | | | | | | | 0 |
| chr2 | 97259239 | ATCATTCTGTTAA-TGAATTGT-TTCCCAAATTTAT | loxF8L | 4 | loxF8L | 3 | ACAATTCA/TGAATTGT | 7 |
| chr2 | 97497934 | ATAAATTTGGGAA-ACAATTCA-TTAACAGAATGAT | loxF8L | 3 | loxF8L | 4 | ACAATTCA/TGAATTGT | 7 |
| chr2 | 113400783 | ATCATTCTGTTAA-TGAATTGT-TTCCCAAATTTAT | loxF8L | 4 | loxF8L | 3 | ACAATTCA/TGAATTGT | 7 |
| --- | | | | | | | | 0 |
| chr4 | 129610035 | AAAAACCTGTACA-TAAATTGT-TTATCAGATTTAT | loxF8L | 4 | loxF8L | 3 | ACAATTTA/TAAATTGT | 7 |
| chr17 | 11380631 | CTAGCACTGTGAG-ACAATTTA-CTCTCAAACTTAG | loxF8R | 4 | loxF8R | 3 | ACAATTTA/TAAATTGT | 7 |
| chr12 | 85846214 | CTAAGTTTTGGAG-TAAATTGT-TACACAGAGTTAA | loxF8R | 4 | loxF8L | 3 | ACAATTTA/TAAATTGT | 7 |
| --- | | | | | | | | 0 |
| chr6 | 127993517 | AAAAGTCTGTGGT-ACAATTTG-TCCACATACTGGT | loxF8L | 3 | loxF8L | 4 | ACAATTTG/CAAATTGT | 7 |
| chr15 | 95677544 | CTAAAATCATGTG-CAAATTGT-ACCCTAGCTTTAT | loxF8R | 3 | loxF8L | 4 | ACAATTTG/CAAATTGT | 7 |
| --- | | | | | | | | 0 |
| chr3 | 28671221 | ACAAACCTCTGGC-ACACAAAT-CTCAGAGTCTTAG | loxF8L | 4 | loxF8R | 3 | ACACAAAT/ATTTGTGT | 7 |
| chr6 | 160378084 | TCAAGATTGTTTG-ATTTGTGT-TCCCTAATTTTAT | loxF8R | 3 | loxF8L | 4 | ACACAAAT/ATTTGTGT | 7 |
| --- | | | | | | | | 0 |
| chr5 | 4145361 | CTTACTGTGTGTG-TTCTGTGT-GCCACAGTTTTGT | loxF8R | 4 | loxF8L | 3 | ACACAGAA/TTCTGTGT | 7 |
| chr10 | 117187161 | CTAACACTGTCTC-TTCTGTGT-TCCAGAGCTTTCT | loxF8R | 4 | loxF8L | 3 | ACACAGAA/TTCTGTGT | 7 |
| --- | | | | | | | | 0 |
| chr6 | 153643928 | ATAAAGCAGTTGA-ACAGAACA-GCCACAGATAAAA | loxF8L | 3 | loxF8L | 4 | ACAGAACA/TGTTCTGT | 7 |
| chr5 | 123688501 | ATAGATCACTGGG-ACAGAACA-CAGAGAATCTAAG | loxF8L | 4 | loxF8R | 3 | ACAGAACA/TGTTCTGT | 7 |
| --- | | | | | | | | 0 |
| chr7 | 94319170 | TTAAAATTGTGTC-ACAGAATG-TCCACTGAAAGAT | loxF8R | 3 | loxF8L | 4 | ACAGAATG/CATTCTGT | 7 |
| chr11 | 42255878 | CTGAGTTTGGGGG-CATTCTGT-CACACAGTAGTAG | loxF8R | 4 | loxF8R | 3 | ACAGAATG/CATTCTGT | 7 |
| --- | | | | | | | | 0 |
| chr5 | 29461676 | AAAAAATTGTGAA-ACAGATAT-ACCACAGCTATAT | loxF8L | 4 | loxF8L | 3 | ACAGATAT/ATATCTGT | 7 |
| chr12 | 121120847 | AACAAAATGTGGA-ATATCTGT-ACAACAGACTTAT | loxF8L | 4 | loxF8L | 3 | ACAGATAT/ATATCTGT | 7 |
| --- | | | | | | | | 0 |
| chr17 | 6326552 | AGAAGAATGCGTG-ACAGCACA-CACACACACGTAG | loxF8R | 4 | loxF8R | 3 | ACAGCACA/TGTGCTGT | 7 |
| chr13 | 71114018 | GTAAAATTGTATG-ACAGCACA-CTCACAACCATGG | loxF8R | 3 | loxF8R | 4 | ACAGCACA/TGTGCTGT | 7 |
| --- | | | | | | | | 0 |
| chr2 | 141863575 | CTTATTTTTTGTG-ACAGGAGA-TCAACAAATTTTT | loxF8R | 4 | loxF8L | 3 | ACAGGAGA/TCTCCTGT | 7 |
| chr2 | 225801801 | ATAAGATTTTGTA-TCTCCTGT-TGCACAGAGCTAG | loxF8R | 3 | loxF8L | 4 | ACAGGAGA/TCTCCTGT | 7 |

| | | | | | | | | |
|---|---|---|---|---|---|---|---|---|
| --- | | | | | | | | 0 |
| chr1 | 236222660 | ATAAATATGTCTA-GTTTATGT-TTCACTTATTTTT | loxF8L | 3 | loxF8L | 4 | ACATAAAC/GTTTATGT | 7 |
| chrX | 83829105 | ATAAATTTCTGTT-GTTTATGT-CGCACAATCTGTG | loxF8L | 4 | loxF8R | 3 | ACATAAAC/GTTTATGT | 7 |
| --- | | | | | | | | 0 |
| chr13 | 94553092 | ATAAAACTGAAGA-GAGTATGT-TGCATATAGTTAT | loxF8L | 3 | loxF8L | 4 | ACATACTC/GAGTATGT | 7 |
| chrX | 129464093 | ATAAATTTGATGG-ACATACTC-CACACACTCTAAA | loxF8L | 4 | loxF8R | 3 | ACATACTC/GAGTATGT | 7 |
| --- | | | | | | | | 0 |
| chr9 | 115897804 | ATATATATGAGGA-AGATATGT-ACCACAGGTGTCT | loxF8L | 3 | loxF8L | 4 | ACATATCT/AGATATGT | 7 |
| chr8 | 52832456 | ATAGATATTTTGA-ACATATCT-TGCACAGTTTAAT | loxF8L | 4 | loxF8L | 3 | ACATATCT/AGATATGT | 7 |
| --- | | | | | | | | 0 |
| chr14 | 59574479 | CTACTTCAGTGGA-ACATATGT-CAAATAATCTTTG | loxF8L | 4 | loxF8R | 3 | ACATATGT/ACATATGT | 7 |
| --- | | | | | | | | 0 |
| chr2 | 54672379 | AATATTCTGTGGT-ATGGATGT-ACCACAGTTTTTT | loxF8L | 4 | loxF8L | 3 | ACATCCAT/ATGGATGT | 7 |
| chr4 | 74990245 | GGAAATCTGTAGT-ACATCCAT-TGCACAGGGTTAT | loxF8L | 4 | loxF8L | 3 | ACATCCAT/ATGGATGT | 7 |
| --- | | | | | | | | 0 |
| chr6 | 40167057 | TTTAGATTTGGTG-ACATTACT-TCCACAGACTGGT | loxF8R | 4 | loxF8L | 3 | ACATTACT/AGTAATGT | 7 |
| chr20 | 23689773 | ATAAATGGGTGAA-AGTAATGT-TTAAAATATTTAT | loxF8L | 3 | loxF8L | 4 | ACATTACT/AGTAATGT | 7 |
| chr20 | 23751671 | ATAAATGGGTGAA-AGTAATGT-TTAAAATATTTAT | loxF8L | 3 | loxF8L | 4 | ACATTACT/AGTAATGT | 7 |
| --- | | | | | | | | 0 |
| chr6 | 85104822 | TTCATTCTGTTGA-CATAATGT-ATCACTGATTTAT | loxF8L | 4 | loxF8L | 3 | ACATTATG/CATAATGT | 7 |
| chr1 | 80079869 | ATACGTGTGTGGC-ACATTATG-TAAAAAGATTTAT | loxF8L | 4 | loxF8L | 3 | ACATTATG/CATAATGT | 7 |
| --- | | | | | | | | 0 |
| chr4 | 10745064 | ATAAGGATGTGGA-GAAAATGT-TAAGCAGAGTTAT | loxF8L | 3 | loxF8L | 4 | ACATTTTC/GAAAATGT | 7 |
| chr3 | 20706243 | ATAAGATTTTGTA-ACATTTTC-TTTACAGTTTTAC | loxF8R | 3 | loxF8L | 4 | ACATTTTC/GAAAATGT | 7 |
| chr2 | 119304785 | GTAAGACTCTGTG-GAAAATGT-TTCAGAGAATGAT | loxF8R | 3 | loxF8L | 4 | ACATTTTC/GAAAATGT | 7 |
| --- | | | | | | | | 0 |
| chr3 | 137553475 | AAAACTCTCTGTA-ACCACAGT-TCCCAAGATTTCT | loxF8L | 4 | loxF8L | 3 | ACCACAGT/ACTGTGGT | 7 |
| chr18 | 1109860 | TTAAATCAGTGAA-ACTGTGGT-CACTCAACATGAG | loxF8L | 3 | loxF8R | 4 | ACCACAGT/ACTGTGGT | 7 |
| --- | | | | | | | | 0 |
| chr1 | 31546346 | ATCCATCTGTGCA-ACCATCCA-TCCATCCATTTGT | loxF8L | 3 | loxF8L | 4 | ACCATCCA/TGGATGGT | 7 |
| chr1 | 31546306 | ATCCATCTGTGCA-ACCATCCA-TCCATCCATGTAT | loxF8L | 3 | loxF8L | 4 | ACCATCCA/TGGATGGT | 7 |
| chr11 | 112185838 | ATACGATTGTGCT-ACCATCCA-TCCACAGAATTCA | loxF8R | 4 | loxF8L | 3 | ACCATCCA/TGGATGGT | 7 |
| --- | | | | | | | | 0 |
| chr1 | 151439262 | AGAAATCTTTACA-TAAATGGT-TCCCCATCTTTAT | loxF8L | 4 | loxF8L | 3 | ACCATTTA/TAAATGGT | 7 |
| chr10 | 66499760 | AGAGATCTTGGGA-TAAATGGT-TCCACACATTAAC | loxF8L | 4 | loxF8L | 3 | ACCATTTA/TAAATGGT | 7 |
| chr9 | 79244125 | ATGATTCTGTAGA-TAAATGGT-TGCTCAGTTATAT | loxF8L | 3 | loxF8L | 4 | ACCATTTA/TAAATGGT | 7 |
| --- | | | | | | | | 0 |
| chr19 | 45887386 | CTCTGATTCTGTT-ACCCAGAC-CTCACAATTTTAA | loxF8R | 4 | loxF8R | 3 | ACCCAGAC/GTCTGGGT | 7 |
| chr7 | 70204118 | GTAAGATAGTGGT-ACCCAGAC-CCCATAATCTAAG | loxF8R | 4 | loxF8R | 3 | ACCCAGAC/GTCTGGGT | 7 |
| --- | | | | | | | | 0 |
| chr12 | 94960955 | GTAAGATTAAGTG-ACCCCATC-TCCACTCTTTGAT | loxF8R | 3 | loxF8L | 4 | ACCCCATC/GATGGGGT | 7 |
| chr17 | 76231602 | GTAATATAGTGAG-ACCCCATC-TCCACTTATTTAA | loxF8R | 4 | loxF8L | 3 | ACCCCATC/GATGGGGT | 7 |
| --- | | | | | | | | 0 |
| chr13 | 26056482 | GTAACATAGTGAG-ACCCTGTC-TCTACAAATTTTT | loxF8R | 4 | loxF8L | 3 | ACCCTGTC/GACAGGGT | 7 |
| chr18 | 14082860 | AAAATTCTGTAGA-GACAGGGT-CACACTATGTTGC | loxF8L | 3 | loxF8R | 4 | ACCCTGTC/GACAGGGT | 7 |
| chr3 | 29132838 | GCAAGATGGTGAG-ACCCTGTC-TCTACAAATATAT | loxF8R | 4 | loxF8L | 3 | ACCCTGTC/GACAGGGT | 7 |
| chr5 | 14853386 | GTAACATAGTGAG-ACCCTGTC-TCCACAGAAAAAT | loxF8R | 4 | loxF8L | 3 | ACCCTGTC/GACAGGGT | 7 |
| --- | | | | | | | | 0 |
| chr14 | 103442611 | ACAAACATGTGGA-ACCTAAAA-CACACAGAATTCT | loxF8L | 3 | loxF8L | 4 | ACCTAAAA/TTTTAGGT | 7 |
| chr21 | 26143752 | ATGAGTTTGAGTG-TTTTAGGT-TCCACATATAAAT | loxF8R | 4 | loxF8L | 3 | ACCTAAAA/TTTTAGGT | 7 |
| chr1 | 43765521 | CTAAAACTGTATG-TTTTAGGT-TCCCCACACTTAA | loxF8R | 3 | loxF8L | 4 | ACCTAAAA/TTTTAGGT | 7 |
| --- | | | | | | | | 0 |
| chr8 | 11215874 | CACAGATTGGGTG-ACTTAGGT-AACAGAAACTTAT | loxF8R | 3 | loxF8R | 4 | ACCTAAGT/ACTTAGGT | 7 |
| chr20 | 40817354 | CCAAGATTGAGAA-ACTTAGGT-TACAGAATCTCAG | loxF8R | 4 | loxF8R | 3 | ACCTAAGT/ACTTAGGT | 7 |
| --- | | | | | | | | 0 |
| chrY | 15801478 | ACAACACGGTGGA-ACCTCATC-TGCACAAATATAT | loxF8L | 4 | loxF8L | 3 | ACCTCATC/GATGAGGT | 7 |
| chr7 | 87547538 | GCAAGATAGTGAG-ACCTCATC-TCTACAAATATAT | loxF8R | 4 | loxF8L | 3 | ACCTCATC/GATGAGGT | 7 |
| --- | | | | | | | | 0 |
| chr5 | 33521444 | ATATATTTGTAGA-GACAAGGT-CTCACTATGTTAC | loxF8L | 3 | loxF8R | 4 | ACCTTGTC/GACAAGGT | 7 |
| chr3 | 149516770 | ATATTTTTGTAGA-GACAAGGT-CTCACAATGTTAC | loxF8L | 4 | loxF8R | 3 | ACCTTGTC/GACAAGGT | 7 |
| --- | | | | | | | | 0 |
| chrX | 106168014 | ATATATGTAAGGA-ACGTTTTA-ACCACATCTTTAT | loxF8L | 4 | loxF8L | 3 | ACGTTTTA/TAAAACGT | 7 |
| chr9 | 117545128 | TTAACATTTTGTC-ACGTTTTA-TCTGCATATTTAT | loxF8R | 4 | loxF8L | 3 | ACGTTTTA/TAAAACGT | 7 |
| --- | | | | | | | | 0 |
| chr6 | 6075037 | ACAAAAATGTGGA-ACTAAAAG-ATAAAAGATTTAT | loxF8L | 3 | loxF8L | 4 | ACTAAAAG/CTTTTAGT | 7 |
| chr3 | 93920006 | AGACATGTGTGGA-CTTTTAGT-TCCTCAGTTAAAT | loxF8L | 3 | loxF8L | 4 | ACTAAAAG/CTTTTAGT | 7 |
| --- | | | | | | | | 0 |
| chr6 | 92071292 | CAAAGACTGAGTG-ACTAAGAC-AACAAACATTTAT | loxF8R | 3 | loxF8L | 4 | ACTAAGAC/GTCTTAGT | 7 |
| chr8 | 11610281 | CAAAGATTTTCTG-ACTAAGAC-CTCAAAAGCATAG | loxF8R | 3 | loxF8R | 4 | ACTAAGAC/GTCTTAGT | 7 |
| chr13 | 71898475 | CAAAGATTTTTTG-ACTAAGAC-CCCAAAAGCATAG | loxF8R | 3 | loxF8R | 4 | ACTAAGAC/GTCTTAGT | 7 |
| --- | | | | | | | | 0 |
| chr15 | 33387566 | TTAAATATGTGCA-ACTAAGGA-TCACCAGATATTT | loxF8L | 3 | loxF8L | 4 | ACTAAGGA/TCCTTAGT | 7 |
| chr13 | 44479718 | CTAAATCTGAGTC-ACTAAGGA-CACACAGTCTTCT | loxF8L | 4 | loxF8R | 3 | ACTAAGGA/TCCTTAGT | 7 |
| --- | | | | | | | | 0 |
| chr4 | 122139748 | CTAGGCTTGTGGG-GTATTAGT-CAAGAAATCTTTG | loxF8R | 3 | loxF8R | 4 | ACTAATAC/GTATTAGT | 7 |
| chr4 | 106770616 | CAAAGATTTCTTG-ACTAATAC-CCCACAATCACAG | loxF8R | 4 | loxF8R | 3 | ACTAATAC/GTATTAGT | 7 |
| --- | | | | | | | | 0 |
| chr11 | 49120195 | CTAAGAATCAGTG-GGTGTAGT-AAGATAATTTTAG | loxF8R | 3 | loxF8R | 4 | ACTACACC/GGTGTAGT | 7 |
| chr11 | 89714665 | CTAAAATTATCTT-ACTACACC-CACTGATTCTTAG | loxF8R | 4 | loxF8R | 3 | ACTACACC/GGTGTAGT | 7 |
| --- | | | | | | | | 0 |
| chr8 | 59849337 | ATAAATTGGTGTA-ACTACCAC-CACACAATCAAGA | loxF8L | 3 | loxF8R | 4 | ACTACCAC/GTGGTAGT | 7 |
| chr16 | 59925195 | ATAAATTTGGAGA-ACTACCAC-CACCCAATTTAAA | loxF8L | 3 | loxF8R | 4 | ACTACCAC/GTGGTAGT | 7 |
| --- | | | | | | | | 0 |
| chr1 | 102716057 | TTATGATTGTGGT-TATATAGT-TCTATAGATTTGT | loxF8R | 4 | loxF8L | 3 | ACTATATA/TATATAGT | 7 |
| chrX | 145419759 | CAAAGATTGGTTG-ACTATATA-TGTATGGATTTAT | loxF8R | 3 | loxF8L | 4 | ACTATATA/TATATAGT | 7 |
| chr11 | 6035427 | ATAAGTATGTGTG-TATATAGT-CTCATATTCTTAT | loxF8R | 3 | loxF8R | 4 | ACTATATA/TATATAGT | 7 |
| --- | | | | | | | | 0 |

| | | | | | | | | |
|---|---|---|---|---|---|---|---|---|
| chr7 | 78747951 | AGAATTCTGTGGT-ACTATATG-CACTCAATATCAA | loxF8L | 3 | loxF8R | 4 | ACTATATG/CATATAGT | 7 |
| chrX | 125652189 | CAAAGATTTGTTG-ACTATATG-TGCATAGGTTTAT | loxF8R | 4 | loxF8L | 3 | ACTATATG/CATATAGT | 7 |
| --- | | | | | | | | 0 |
| chr18 | 64438496 | ATAAATTTCTAGA-GTGATAGT-CACACTTTCTAAA | loxF8L | 3 | loxF8R | 4 | ACTATCAC/GTGATAGT | 7 |
| chr6 | 140127031 | GTAAGATTATAGG-GTGATAGT-AACAGAATCTAAG | loxF8R | 4 | loxF8R | 3 | ACTATCAC/GTGATAGT | 7 |
| --- | | | | | | | | 0 |
| chr1 | 56951766 | CCAAGTCAGTGTG-ACTCCAAA-TCCTAAGCTTTAT | loxF8R | 4 | loxF8L | 3 | ACTCCAAA/TTTGGAGT | 7 |
| chr1 | 65045852 | ATCAAGCTTGGGA-TTTGGAGT-CAGACAAACATAG | loxF8L | 4 | loxF8R | 3 | ACTCCAAA/TTTGGAGT | 7 |
| --- | | | | | | | | 0 |
| chr6 | 95323675 | AAAGACCTGTGGA-ACTCTAGC-TTCTCAGACTTAA | loxF8L | 3 | loxF8L | 4 | ACTCTAGC/GCTAGAGT | 7 |
| chr13 | 70732280 | TTAGCAATGTGTG-ACTCTAGC-CACAAAATGTTAC | loxF8R | 4 | loxF8R | 3 | ACTCTAGC/GCTAGAGT | 7 |
| --- | | | | | | | | 0 |
| chr3 | 167835714 | ATATGATTGAGTG-ACTCTCTC-TCCTGATATTTTT | loxF8R | 3 | loxF8L | 4 | ACTCTCTC/GAGAGAGT | 7 |
| chr7 | 69809636 | ATAAAACCCTAGA-GAGAGAGT-AACACAATCCTGG | loxF8L | 4 | loxF8R | 3 | ACTCTCTC/GAGAGAGT | 7 |
| chr19 | 30584484 | ATAAAAGTGTGTG-ACTCTCTC-TCCACATGGATAT | loxF8R | 3 | loxF8L | 4 | ACTCTCTC/GAGAGAGT | 7 |
| --- | | | | | | | | 0 |
| chr2 | 233494848 | AAAAAACTGTGCA-ATTTCAGT-TCTAGGAATTTAT | loxF8L | 3 | loxF8L | 4 | ACTGAAAT/ATTTCAGT | 7 |
| chr3 | 87136164 | ATAAATGTTTGTT-ATTTCAGT-CACACCATCTATG | loxF8L | 4 | loxF8R | 3 | ACTGAAAT/ATTTCAGT | 7 |
| --- | | | | | | | | 0 |
| chr10 | 119939732 | CCCAGCTTGTGTG-AGTGCAGT-GGCGCAATCTTGG | loxF8R | 3 | loxF8R | 4 | ACTGCACT/AGTGCAGT | 7 |
| chr3 | 3174587 | CTGAGATTGTGCC-ACTGCACT-CACTCGAGCTTGG | loxF8R | 3 | loxF8R | 4 | ACTGCACT/AGTGCAGT | 7 |
| --- | | | | | | | | 0 |
| chr2 | 37489352 | GTAAGTCAGTGTA-ACTGCCAA-GTCACAGATTTTT | loxF8L | 4 | loxF8L | 3 | ACTGCCAA/TTGGCAGT | 7 |
| chr4 | 27560893 | CTAAAACTGAGGA-ACTGCCAA-AACAGAATTTTAC | loxF8L | 3 | loxF8R | 4 | ACTGCCAA/TTGGCAGT | 7 |
| --- | | | | | | | | 0 |
| chr7 | 10615800 | ATAAACCTATGCA-TATACAGT-CAACCAATCTTCA | loxF8L | 3 | loxF8R | 4 | ACTGTATA/TATACAGT | 7 |
| chr19 | 12855615 | TTCAAATTGTGTC-TATACAGT-CACACAGATATAT | loxF8R | 4 | loxF8L | 3 | ACTGTATA/TATACAGT | 7 |
| --- | | | | | | | | 0 |
| chr2 | 216597529 | AGTAATCTGTAGC-ATCACAGT-GCCAATGATTTAT | loxF8L | 4 | loxF8L | 3 | ACTGTGAT/ATCACAGT | 7 |
| chr2 | 153633734 | CCCAATCTTTGGA-ACTGTGAT-GCTAGAGATTTAT | loxF8L | 4 | loxF8L | 3 | ACTGTGAT/ATCACAGT | 7 |
| --- | | | | | | | | 0 |
| chr2 | 149264858 | CATAGACTGGGTG-ACTTAAAC-TGCAGATATTTAT | loxF8R | 4 | loxF8L | 3 | ACTTAAAC/GTTTAAGT | 7 |
| chr1 | 190621529 | CATAGACTGGGTG-ACTTAAAC-AGCAGAGATTTAT | loxF8R | 4 | loxF8L | 3 | ACTTAAAC/GTTTAAGT | 7 |
| chr17 | 30004983 | ATAAATCTCTGTT-GTTTAAGT-CACCCAGTCTATG | loxF8L | 3 | loxF8R | 4 | ACTTAAAC/GTTTAAGT | 7 |
| chrX | 43997237 | ATAAATCTTTGTT-GTTTAAGT-CACCCAGTCTGTG | loxF8L | 3 | loxF8R | 4 | ACTTAAAC/GTTTAAGT | 7 |
| chr8 | 61026372 | ATAAATGTCTGCT-GTTTAAGT-CACACAGTCTGTG | loxF8L | 4 | loxF8R | 3 | ACTTAAAC/GTTTAAGT | 7 |
| chr2 | 208378739 | ATAAATGTTTGCT-GTTTAAGT-CACCCAGTCTAAG | loxF8L | 4 | loxF8R | 3 | ACTTAAAC/GTTTAAGT | 7 |
| chr7 | 52733658 | CATAGATTGGGTG-ACTTAAAC-AACAAACATTTAT | loxF8R | 3 | loxF8L | 4 | ACTTAAAC/GTTTAAGT | 7 |
| chr2 | 216528502 | ATAAATTTATGTT-GTTTAAGT-CACCCAGTCTTTG | loxF8L | 4 | loxF8R | 3 | ACTTAAAC/GTTTAAGT | 7 |
| chr11 | 106466127 | ATAAATTTCTGGT-GTTTAAGT-CACCCAGTCTGTG | loxF8L | 3 | loxF8R | 4 | ACTTAAAC/GTTTAAGT | 7 |
| chr3 | 137810777 | ATAAATTTCTGTA-GTTTAAGT-CACCCAGTCTGTG | loxF8L | 3 | loxF8R | 4 | ACTTAAAC/GTTTAAGT | 7 |
| chr3 | 176025841 | CAAAGACTGGGTG-ACTTAAAC-AACAGAAATTTAT | loxF8R | 3 | loxF8L | 4 | ACTTAAAC/GTTTAAGT | 7 |
| chr15 | 49654801 | CATAGATTGAGTG-ACTTAAAC-AACAGAAATTTAT | loxF8R | 3 | loxF8L | 4 | ACTTAAAC/GTTTAAGT | 7 |
| chr15 | 97104453 | ATAAATTTGTGTT-GTTTAAGT-CACCCAGTCTGTG | loxF8L | 3 | loxF8R | 4 | ACTTAAAC/GTTTAAGT | 7 |
| chr14 | 33544687 | ATAAGTTTCTGTT-GTTTAAGT-CACCCAATCTGTG | loxF8R | 4 | loxF8R | 3 | ACTTAAAC/GTTTAAGT | 7 |
| chr12 | 118947624 | CACAGATTGGGTG-ACTTAAAC-AACACAAATCTAT | loxF8R | 3 | loxF8L | 4 | ACTTAAAC/GTTTAAGT | 7 |
| --- | | | | | | | | 0 |
| chr13 | 80385208 | CACAGATTGGGTG-ACTTAAAT-AACAGAAATTTAT | loxF8R | 3 | loxF8L | 4 | ACTTAAAT/ATTTAAGT | 7 |
| chr7 | 20088946 | TGAAGATGGTGTT-ATTTAAGT-TCCATAAAGTTAT | loxF8R | 4 | loxF8L | 3 | ACTTAAAT/ATTTAAGT | 7 |
| chr2 | 176832871 | GTTAATCTTTGGA-ATTTAAGT-TCTACATCTTAAT | loxF8L | 3 | loxF8L | 4 | ACTTAAAT/ATTTAAGT | 7 |
| --- | | | | | | | | 0 |
| chr8 | 55389248 | ATAAATTTCTGTT-GTTGAAGT-CACACAGTCTGTG | loxF8L | 4 | loxF8R | 3 | ACTTCAAC/GTTGAAGT | 7 |
| chrX | 33759056 | CCAAGATTGTGGT-GTTGAAGT-CCCCCACAATTAT | loxF8R | 3 | loxF8L | 4 | ACTTCAAC/GTTGAAGT | 7 |
| --- | | | | | | | | 0 |
| chr20 | 8218923 | ATTGAGCTGTGAA-TTAGAAGT-TTCAGAGTTTTAT | loxF8L | 4 | loxF8L | 3 | ACTTCTAA/TTAGAAGT | 7 |
| chr13 | 50781864 | TTAAGATTTTATG-TTAGAAGT-CAGAAAATCTTGA | loxF8R | 3 | loxF8R | 4 | ACTTCTAA/TTAGAAGT | 7 |
| --- | | | | | | | | 0 |
| chr18 | 60412420 | ATAAATGTATGTA-GTTCAAGT-CACCCAGTCTGTG | loxF8L | 3 | loxF8R | 4 | ACTTGAAC/GTTCAAGT | 7 |
| chr6 | 96959353 | CACAGACTGGGTG-ACTTGAAC-AACACAAATTTAT | loxF8R | 4 | loxF8L | 3 | ACTTGAAC/GTTCAAGT | 7 |
| --- | | | | | | | | 0 |
| chr19 | 52904068 | CTCAGGTGGAGTG-GGGCAAGT-TCCAAAGATTAGT | loxF8R | 4 | loxF8L | 3 | ACTTGCCC/GGGCAAGT | 7 |
| chr10 | 84171682 | CTCAGATGGAGTG-GGGCAAGT-TCCAAAGACTAGT | loxF8R | 3 | loxF8L | 4 | ACTTGCCC/GGGCAAGT | 7 |
| chr10 | 97561021 | CTCAGATGGAGTG-GGGCAAGT-TCCAAAGACTAGT | loxF8R | 3 | loxF8L | 4 | ACTTGCCC/GGGCAAGT | 7 |
| chr13 | 22884300 | ACTAGTCTTTGGA-ACTTGCCC-CACTCCATCTGAG | loxF8L | 4 | loxF8R | 3 | ACTTGCCC/GGGCAAGT | 7 |
| chr19 | 20431816 | CTCAGATGGAGTG-GGGCAAGT-TCCAAAGACTAGT | loxF8R | 3 | loxF8L | 4 | ACTTGCCC/GGGCAAGT | 7 |
| chr19 | 52903955 | CTCAGATGGAGTG-GGGCAAGT-TCCAAAGACTAGT | loxF8R | 3 | loxF8L | 4 | ACTTGCCC/GGGCAAGT | 7 |
| chr1 | 46331785 | CTCAGATGGAGTG-GGGCAAGT-TCCAAAGACTAGT | loxF8R | 3 | loxF8L | 4 | ACTTGCCC/GGGCAAGT | 7 |
| chr20 | 15985463 | CTCAGATGGAGTG-GGGCAAGT-TCCAAAGACTAGT | loxF8R | 3 | loxF8L | 4 | ACTTGCCC/GGGCAAGT | 7 |
| chr2 | 34678045 | ACTAGTCTTTGGA-ACTTGCCC-CACTCCATCTGAG | loxF8L | 4 | loxF8R | 3 | ACTTGCCC/GGGCAAGT | 7 |
| chr4 | 115951130 | ACTAGTCTTTGGA-ACTTGCCC-CACTCCATCTGAG | loxF8L | 4 | loxF8R | 3 | ACTTGCCC/GGGCAAGT | 7 |
| chr5 | 42814150 | ACTAGTCTTTGGA-ACTTGCCC-CACTCCATCTGAG | loxF8L | 4 | loxF8R | 3 | ACTTGCCC/GGGCAAGT | 7 |
| chr5 | 42813924 | ACTAGTCTTTGGA-ACTTGCCC-CACTCCATCTGAG | loxF8L | 4 | loxF8R | 3 | ACTTGCCC/GGGCAAGT | 7 |
| chr5 | 82272859 | CTCAGATGGAGTG-GGGCAAGT-TCCAAAGACTAGT | loxF8R | 3 | loxF8L | 4 | ACTTGCCC/GGGCAAGT | 7 |
| chr5 | 42814037 | ACTAGTCTTTGGA-ACTTGCCC-CACTCCATCTGAG | loxF8L | 4 | loxF8R | 3 | ACTTGCCC/GGGCAAGT | 7 |
| chr5 | 139526617 | ACTAGTCTTTGGA-ACTTGCCC-CACTCCATCTGAG | loxF8L | 4 | loxF8R | 3 | ACTTGCCC/GGGCAAGT | 7 |
| chr5 | 139526730 | ACTAGTCTTTGGA-ACTTGCCC-CACTCCATCTGAG | loxF8L | 4 | loxF8R | 3 | ACTTGCCC/GGGCAAGT | 7 |
| chr6 | 31185986 | CTCAGATGGAGTG-GGGCAAGT-TCCAAAGACTAGT | loxF8R | 3 | loxF8L | 4 | ACTTGCCC/GGGCAAGT | 7 |
| chr6 | 111289840 | CTCAGATGGAGTG-GGGCAAGT-TCCAAAGACTAGT | loxF8R | 3 | loxF8L | 4 | ACTTGCCC/GGGCAAGT | 7 |
| chr7 | 135172201 | ACTAGTCTTTGGA-ACTTGCCC-CACTCCATCTGAG | loxF8L | 4 | loxF8R | 3 | ACTTGCCC/GGGCAAGT | 7 |
| chrX | 3966586 | ACTAGTCTTTGGA-ACTTGCCC-CACTCCATCTGAG | loxF8L | 4 | loxF8R | 3 | ACTTGCCC/GGGCAAGT | 7 |
| chr19 | 20431703 | TTCAGATGGGGTG-GGGCAAGT-TCCAAAGACTAAT | loxF8R | 4 | loxF8L | 3 | ACTTGCCC/GGGCAAGT | 7 |
| --- | | | | | | | | 0 |
| chrX | 77632614 | ATAAACATATGAA-GAAAAAGT-TGCACAAATTAAT | loxF8L | 4 | loxF8L | 3 | ACTTTTTC/GAAAAAGT | 7 |
| chr8 | 76699377 | ATACTGCAGTGGA-GAAAAAGT-TACACAATCTTTC | loxF8L | 4 | loxF8R | 3 | ACTTTTTC/GAAAAAGT | 7 |
| --- | | | | | | | | 0 |
| chr1 | 66258840 | CTTTGAATGTTTG-CTTTTTCT-TTCACTGTTTTAT | loxF8R | 4 | loxF8L | 3 | AGAAAAAG/CTTTTTCT | 7 |
| chr4 | 25265390 | TTTAGTTTGTTTG-CTTTTTCT-TTTATAGATTTAT | loxF8R | 4 | loxF8L | 3 | AGAAAAAG/CTTTTTCT | 7 |

| | | | | | | | | |
|---|---|---|---|---|---|---|---|---|
| --- | | | | | | | | 0 |
| chr1 | 174484240 | ATAAGATTGTTAG-ATTTTTCT-TCTATAGAGTTGT | loxF8R | 3 | loxF8L | 4 | AGAAAAAT/ATTTTTCT | 7 |
| chr7 | 42272609 | ATAGAACTGTGGG-AGAAAAAT-GTCAAAAATTTAT | loxF8L | 3 | loxF8L | 4 | AGAAAAAT/ATTTTTCT | 7 |
| chr4 | 80438488 | TTAAGGTTGTCTG-ATTTTTCT-GAGACAATTTGAG | loxF8R | 3 | loxF8R | 4 | AGAAAAAT/ATTTTTCT | 7 |
| --- | | | | | | | | 0 |
| chr8 | 20946733 | AAAAAGCTGTGGG-AGAAAACC-TTCACAATCTTGA | loxF8L | 3 | loxF8R | 4 | AGAAAACC/GGTTTTCT | 7 |
| chr3 | 134998493 | CTAAGTCTGGGGA-GGTTTTCT-CAAACCATCTCTG | loxF8L | 3 | loxF8R | 4 | AGAAAACC/GGTTTTCT | 7 |
| chr5 | 132744963 | CTAGGATGGTGGG-GGTTTTCT-CTCAAAATTTTAA | loxF8R | 3 | loxF8R | 4 | AGAAAACC/GGTTTTCT | 7 |
| --- | | | | | | | | 0 |
| chr13 | 109239886 | AGAAATCTGAGTA-TCTTTTCT-TACATAATTTTTG | loxF8L | 3 | loxF8R | 4 | AGAAAAGA/TCTTTTCT | 7 |
| chr6 | 161454882 | CTCAGTTTCTCTG-TCTTTTCT-TCCCCAGCTATAT | loxF8R | 4 | loxF8L | 3 | AGAAAAGA/TCTTTTCT | 7 |
| --- | | | | | | | | 0 |
| chr6 | 18246907 | AAAAATCTGTGTC-AGAAAATA-AACAGAATATTAA | loxF8L | 3 | loxF8R | 4 | AGAAAATA/TATTTTCT | 7 |
| chr13 | 45726118 | ATGACTATGTGGG-AGAAAATA-CCCACAGTCTTAT | loxF8L | 4 | loxF8L | 3 | AGAAAATA/TATTTTCT | 7 |
| chr9 | 27446946 | ATGATTATGTGAA-AGAAAATA-TGGACAGAATTAT | loxF8L | 4 | loxF8L | 3 | AGAAAATA/TATTTTCT | 7 |
| --- | | | | | | | | 0 |
| chr5 | 117818426 | ATGATATTGTGGA-GATTTTCT-TCCAGGGTTTTAT | loxF8L | 4 | loxF8L | 3 | AGAAAATC/GATTTTCT | 7 |
| chr2 | 215796805 | ATAAATCTATTCA-AGAAAATC-TACTAAATCTTGG | loxF8L | 3 | loxF8R | 4 | AGAAAATC/GATTTTCT | 7 |
| chr6 | 103439048 | ATAAATTTGGGAA-AGAAAATC-TAGAATATCTTAG | loxF8L | 3 | loxF8R | 4 | AGAAAATC/GATTTTCT | 7 |
| --- | | | | | | | | 0 |
| chr12 | 43477913 | AAAAATCTCTAGA-AGAAAGCA-CACACTTTTTGAG | loxF8L | 3 | loxF8R | 4 | AGAAAGCA/TGCTTTCT | 7 |
| chr3 | 103436430 | ATATTTGTGTGGA-AGAAAGCA-CAGAAAATCTGAT | loxF8L | 3 | loxF8R | 4 | AGAAAGCA/TGCTTTCT | 7 |
| chr3 | 84584853 | CTAAGAGTATGTT-AGAAAGCA-CACAAAATTTGTG | loxF8R | 3 | loxF8R | 4 | AGAAAGCA/TGCTTTCT | 7 |
| --- | | | | | | | | 0 |
| chr13 | 55229015 | AGTATTTTGTGGA-AGAAAGTA-TCCAGAGACTTAG | loxF8L | 4 | loxF8L | 3 | AGAAAGTA/TACTTTCT | 7 |
| chr10 | 56337593 | CAGAGATTGAGTT-AGAAAGTA-TTCCCAGATCTAT | loxF8R | 4 | loxF8L | 3 | AGAAAGTA/TACTTTCT | 7 |
| --- | | | | | | | | 0 |
| chr8 | 66182757 | AATAATTTGTTGA-AGAAATCT-TCCCCAGATTCCT | loxF8L | 4 | loxF8L | 3 | AGAAATCT/AGATTTCT | 7 |
| chr6 | 32368612 | AGAAATCTCTGGT-AGATTTCT-CCCACTCACTTAG | loxF8L | 3 | loxF8R | 4 | AGAAATCT/AGATTTCT | 7 |
| chr2 | 104130235 | ATATGATTGAGAG-AGATTTCT-TCCACAGATACAA | loxF8R | 4 | loxF8L | 3 | AGAAATCT/AGATTTCT | 7 |
| chr3 | 107279371 | ATTGATCTGTGGT-AGAAATCT-AACAAAATATTTG | loxF8L | 3 | loxF8R | 4 | AGAAATCT/AGATTTCT | 7 |
| --- | | | | | | | | 0 |
| chr2 | 8617647 | CTCAGGTTCTGTG-AGAACACA-CACACATATTTTT | loxF8R | 3 | loxF8L | 4 | AGAACACA/TGTGTTCT | 7 |
| chr14 | 83071199 | ATATATATGTGTA-AGAACACA-CACACATATATAT | loxF8L | 3 | loxF8L | 4 | AGAACACA/TGTGTTCT | 7 |
| --- | | | | | | | | 0 |
| chr13 | 93003510 | ATAAGATGTTGTG-AGAACTCA-CTCACTATCTTGA | loxF8R | 3 | loxF8R | 4 | AGAACTCA/TGAGTTCT | 7 |
| chr2 | 99283115 | CTATGATAGTGAG-TGAGTTCT-CACAAGATCTGAT | loxF8R | 3 | loxF8R | 4 | AGAACTCA/TGAGTTCT | 7 |
| chr15 | 20586070 | ATCAGATTTTATG-AGAACTCA-CTCACTATCTGAG | loxF8R | 4 | loxF8R | 3 | AGAACTCA/TGAGTTCT | 7 |
| chr15 | 22414281 | CTCAGATAGTGAG-TGAGTTCT-CATAAAATCTGAT | loxF8R | 3 | loxF8R | 4 | AGAACTCA/TGAGTTCT | 7 |
| chr17 | 49517270 | CTGTGATGGTGAG-TGAGTTCT-CACAAAATCTGAT | loxF8R | 4 | loxF8R | 3 | AGAACTCA/TGAGTTCT | 7 |
| chr7 | 104516243 | ATCAGATTTTGTG-AGAACTCA-CTCACTATCACAG | loxF8R | 3 | loxF8R | 4 | AGAACTCA/TGAGTTCT | 7 |
| chr14 | 50406325 | CTCTGATAGTGAG-TGAGTTCT-CACAAAATCTGAT | loxF8R | 4 | loxF8R | 3 | AGAACTCA/TGAGTTCT | 7 |
| chr9 | 12232960 | ATCAGATTTTGTG-AGAACTCA-CTCACTATCATAA | loxF8R | 3 | loxF8R | 4 | AGAACTCA/TGAGTTCT | 7 |
| --- | | | | | | | | 0 |
| chr5 | 67810662 | AAAACTATCTGGA-AGAATATG-CACAGAATATTAA | loxF8L | 4 | loxF8R | 3 | AGAATATG/CATATTCT | 7 |
| chr6 | 110299472 | CTTAGATTTTCTT-AGAATATG-TCCACAAATATAA | loxF8R | 4 | loxF8L | 3 | AGAATATG/CATATTCT | 7 |
| chr11 | 88577179 | GCAAAATTGTATG-CATATTCT-GCCACAGATTTTA | loxF8R | 4 | loxF8L | 3 | AGAATATG/CATATTCT | 7 |
| --- | | | | | | | | 0 |
| chr10 | 49658889 | AAAATTCAGTGGA-AGAATTAG-AAGATAATCTGAG | loxF8L | 3 | loxF8R | 4 | AGAATTAG/CTAATTCT | 7 |
| chr15 | 94892635 | ATAGAGCTGAGGA-AGAATTAG-TATTCATATTTAT | loxF8L | 3 | loxF8L | 4 | AGAATTAG/CTAATTCT | 7 |
| --- | | | | | | | | 0 |
| chr10 | 117021439 | GTAAGATTAAATG-AGAATTCT-GATACATTCTTAG | loxF8R | 4 | loxF8R | 3 | AGAATTCT/AGAATTCT | 7 |
| --- | | | | | | | | 0 |
| chr16 | 12083581 | CTCCGATTGGGAG-TTAGGTCT-TCCACTGGTTCAT | loxF8R | 4 | loxF8L | 3 | AGACCTAA/TTAGGTCT | 7 |
| chr9 | 43275470 | GTAATTCTCTGGG-TTAGGTCT-TATACAATCTTGG | loxF8L | 4 | loxF8R | 3 | AGACCTAA/TTAGGTCT | 7 |
| chr10 | 42182322 | CTAAGATTGTACA-AGACCTAA-CCCAGAGAATTAC | loxF8R | 3 | loxF8L | 4 | AGACCTAA/TTAGGTCT | 7 |
| --- | | | | | | | | 0 |
| chr5 | 23546981 | CTGTGGTTGTGGG-AGGAGTCT-TCTATAGATATAT | loxF8R | 4 | loxF8L | 3 | AGACTCCT/AGGAGTCT | 7 |
| chr7 | 33438902 | CTAAAGCTGATGA-AGGAGTCT-CTTACAACCTTAG | loxF8L | 4 | loxF8R | 3 | AGACTCCT/AGGAGTCT | 7 |
| --- | | | | | | | | 0 |
| chr16 | 66063071 | AAAAATATGTGAA-AGACTTAG-CACACATTTTCTG | loxF8L | 3 | loxF8R | 4 | AGACTTAG/CTAAGTCT | 7 |
| chr6 | 140343278 | CCAAGTTTTTGTG-AGACTTAG-CAAACAACCTTTA | loxF8R | 3 | loxF8R | 4 | AGACTTAG/CTAAGTCT | 7 |
| --- | | | | | | | | 0 |
| chr8 | 79059887 | CTTATATTGTGCT-TAAAGTCT-AACACAATCCTTG | loxF8R | 4 | loxF8R | 3 | AGACTTTA/TAAAGTCT | 7 |
| chr20 | 52126706 | TTAGGATTTTATG-TAAAGTCT-CATAAAAGCTTAG | loxF8R | 4 | loxF8R | 3 | AGACTTTA/TAAAGTCT | 7 |
| --- | | | | | | | | 0 |
| chr7 | 10568922 | ATAAACCTTTAGA-ATTTCTCT-TTCACAAATATGT | loxF8L | 3 | loxF8L | 4 | AGAGAAAT/ATTTCTCT | 7 |
| chr4 | 71288824 | AGAAGTCTGTGGT-ATTTCTCT-CATTCCATCTTAT | loxF8L | 3 | loxF8R | 4 | AGAGAAAT/ATTTCTCT | 7 |
| chr10 | 37692513 | CAAAGATGGTGCC-AGAGAAAT-CAGACATTCATAG | loxF8R | 4 | loxF8R | 3 | AGAGAAAT/ATTTCTCT | 7 |
| chr1 | 232740112 | TTTAAACTATGGA-AGAGAAAT-TATACAATCTAAG | loxF8L | 4 | loxF8R | 3 | AGAGAAAT/ATTTCTCT | 7 |
| --- | | | | | | | | 0 |
| chr8 | 83345460 | CTAATATTATGAA-AGAGAATA-CACAGGTTCTTAG | loxF8R | 4 | loxF8R | 3 | AGAGAATA/TATTCTCT | 7 |
| chr15 | 83052399 | CTAAGATTAAGGA-TATTCTCT-TACATAATCATAG | loxF8R | 4 | loxF8R | 3 | AGAGAATA/TATTCTCT | 7 |
| --- | | | | | | | | 0 |
| chr7 | 10433364 | ATAAGTTTTTGGA-AGAGACCT-TTTACAAATTTAA | loxF8L | 3 | loxF8L | 4 | AGAGACCT/AGGTCTCT | 7 |
| chr13 | 35970074 | ATGACATTGTTTG-AGGTCTCT-CTCACAATCTAAA | loxF8R | 4 | loxF8R | 3 | AGAGACCT/AGGTCTCT | 7 |
| --- | | | | | | | | 0 |
| chr8 | 65954218 | AAAAATCTATGAC-TTTCCTCT-CCCAGACATTTAT | loxF8L | 4 | loxF8L | 3 | AGAGGAAA/TTTCCTCT | 7 |
| chr14 | 40530044 | AAAAATCTGTCTA-TTTCCTCT-GCCATTCATTTAT | loxF8L | 3 | loxF8L | 4 | AGAGGAAA/TTTCCTCT | 7 |
| --- | | | | | | | | 0 |
| chr17 | 42968629 | ACAAATCTGGGCA-AGAGGAAG-CACCCCAGCTTGG | loxF8L | 3 | loxF8R | 4 | AGAGGAAG/CTTCCTCT | 7 |
| chr7 | 133282081 | ATAGATCTCTGGT-CTTCCTCT-CACAAATGCTAAG | loxF8L | 3 | loxF8R | 4 | AGAGGAAG/CTTCCTCT | 7 |
| chr2 | 75005754 | CTATGTTTCTCTG-AGAGGAAG-CCTACAATCTTAA | loxF8R | 4 | loxF8R | 3 | AGAGGAAG/CTTCCTCT | 7 |
| --- | | | | | | | | 0 |
| chr12 | 79847134 | AAAAATCTTTCCA-GTACCTCT-GCCATAAATTTAT | loxF8L | 4 | loxF8L | 3 | AGAGGTAC/GTACCTCT | 7 |
| chr1 | 114211066 | TTAAATCTGAGAA-GTACCTCT-TCCACAGCCTCGT | loxF8L | 3 | loxF8L | 4 | AGAGGTAC/GTACCTCT | 7 |
| --- | | | | | | | | 0 |

| | | | | | | | | |
|---|---|---|---|---|---|---|---|---|
| chr2 | 200177416 | CTAAGTTTCTATG-GTTACTCT-TTGACAGTTTTGT | loxF8R | 3 | loxF8L | 4 | AGAGTAAC/GTTACTCT | 7 |
| chr2 | 159363289 | CACAGATAGTGTA-GTTACTCT-AACACACTCTGAG | loxF8R | 4 | loxF8R | 3 | AGAGTAAC/GTTACTCT | 7 |
| --- | | | | | | | | 0 |
| chr16 | 22887970 | ACAAAGATGTGGA-AGATAAAT-ACAACAGCTTTCT | loxF8L | 3 | loxF8L | 4 | AGATAAAT/ATTTATCT | 7 |
| chr13 | 55484121 | CAAACATTGTTTC-ATTTATCT-TCTACTTATTTAT | loxF8R | 4 | loxF8L | 3 | AGATAAAT/ATTTATCT | 7 |
| --- | | | | | | | | 0 |
| chrX | 34491975 | AAAAATATGTAGC-AGATGCTA-GGCATAGATTTAT | loxF8L | 4 | loxF8L | 3 | AGATGCTA/TAGCATCT | 7 |
| chr8 | 85458533 | CCAGGATTGTATC-TAGCATCT-TCCAGACATTCAT | loxF8R | 4 | loxF8L | 3 | AGATGCTA/TAGCATCT | 7 |
| chr3 | 148829385 | CTAAGATTCTTTT-TAGCATCT-TAAACAGTCATAG | loxF8R | 3 | loxF8R | 4 | AGATGCTA/TAGCATCT | 7 |
| chr3 | 129730229 | TTAATTCTCTCGA-TAGCATCT-TTCACAGATTAAA | loxF8L | 4 | loxF8L | 3 | AGATGCTA/TAGCATCT | 7 |
| --- | | | | | | | | 0 |
| chr17 | 19182566 | ATAAATTTATGAA-TATTTGCT-CACAAACTCAAAG | loxF8L | 3 | loxF8R | 4 | AGCAAATA/TATTTGCT | 7 |
| chr17 | 19069595 | CTTTGAGTTTGTG-AGCAAATA-TTCATAAATTTAT | loxF8R | 4 | loxF8L | 3 | AGCAAATA/TATTTGCT | 7 |
| chr17 | 19195660 | CTTTGAGTTTGTG-AGCAAATA-TTCATAAATTTAT | loxF8R | 4 | loxF8L | 3 | AGCAAATA/TATTTGCT | 7 |
| chr4 | 140278010 | ATAATTCTATGTA-TATTTGCT-TCTTCTGACTTAT | loxF8L | 3 | loxF8L | 4 | AGCAAATA/TATTTGCT | 7 |
| --- | | | | | | | | 0 |
| chr4 | 101372488 | CTGAGCTTGTGGG-GTATTGCT-CAATAAATCTTTG | loxF8R | 3 | loxF8R | 4 | AGCAATAC/GTATTGCT | 7 |
| chr9 | 12692113 | CTGTGCTTGTGTG-GTATTGCT-CAAGAAATCTTTG | loxF8R | 3 | loxF8R | 4 | AGCAATAC/GTATTGCT | 7 |
| chrX | 74646644 | CTATGCTTGTGGG-GTATTGCT-CAAGAAATCTTTG | loxF8R | 3 | loxF8R | 4 | AGCAATAC/GTATTGCT | 7 |
| chrX | 140396533 | CAAAGATTTCTTG-AGCAATAC-CTCACAAGCATAG | loxF8R | 4 | loxF8R | 3 | AGCAATAC/GTATTGCT | 7 |
| chrX | 56274551 | CTATGCTTGTGGG-GTATTGCT-CAACAAATCTTTG | loxF8R | 3 | loxF8R | 4 | AGCAATAC/GTATTGCT | 7 |
| chrX | 94653363 | CTGTGCTTGTGGG-GTATTGCT-CAAAAAATCTTTG | loxF8R | 4 | loxF8R | 3 | AGCAATAC/GTATTGCT | 7 |
| chrX | 65059260 | CTGCGCTTGTGCG-GTATTGCT-CAAAAAATCTTTG | loxF8R | 4 | loxF8R | 3 | AGCAATAC/GTATTGCT | 7 |
| --- | | | | | | | | 0 |
| chrX | 55062637 | CTGTGCTTGTGGG-ATATTGCT-CACGAAATCTTTG | loxF8R | 4 | loxF8R | 3 | AGCAATAT/ATATTGCT | 7 |
| chr3 | 163309496 | CAAAGATTTTTTG-AGCAATAT-CCCACAAACACAG | loxF8R | 3 | loxF8R | 4 | AGCAATAT/ATATTGCT | 7 |
| --- | | | | | | | | 0 |
| chr11 | 98548575 | AAAAATCTGAAAA-AGCACAAA-CAGGCAATCTAAG | loxF8L | 4 | loxF8R | 3 | AGCACAAA/TTTGTGCT | 7 |
| chr18 | 39428310 | AAAAATCTGAAGG-AGCACAAA-TAGACAATCTAAG | loxF8L | 4 | loxF8R | 3 | AGCACAAA/TTTGTGCT | 7 |
| chr11 | 59740442 | CTTAGATTGTCTA-TTTGTGCT-TCTTCAGACTTTT | loxF8R | 3 | loxF8L | 4 | AGCACAAA/TTTGTGCT | 7 |
| chrX | 95255089 | CTTAGATTGTCTA-TTTGTGCT-TTTTCAGACTTAT | loxF8R | 3 | loxF8L | 4 | AGCACAAA/TTTGTGCT | 7 |
| --- | | | | | | | | 0 |
| chr7 | 118271064 | GTTAGATTGTCTA-TATGTGCT-TCCTCAGACTTTT | loxF8R | 4 | loxF8L | 3 | AGCACATA/TATGTGCT | 7 |
| chr3 | 117351438 | CTGGGTTGGTGTG-TATGTGCT-TCTACAAATATAT | loxF8R | 4 | loxF8L | 3 | AGCACATA/TATGTGCT | 7 |
| --- | | | | | | | | 0 |
| chr2 | 147741729 | AAAAGTATTTGGA-AGCAGACA-TACACGGATTTAA | loxF8L | 4 | loxF8L | 3 | AGCAGACA/TGTCTGCT | 7 |
| chr6 | 146657898 | ATAAATATGGCAA-AGCAGACA-CACACTAAATTAG | loxF8L | 4 | loxF8R | 3 | AGCAGACA/TGTCTGCT | 7 |
| --- | | | | | | | | 0 |
| chr13 | 107978591 | ATTAATATGCAGA-TTTGGGCT-TCTGCAGATTTAA | loxF8L | 4 | loxF8L | 3 | AGCCCAAA/TTTGGGCT | 7 |
| chr12 | 21635406 | CTTTGATTCTGTG-AGCCCAAA-TTCACATATTGAA | loxF8R | 3 | loxF8L | 4 | AGCCCAAA/TTTGGGCT | 7 |
| --- | | | | | | | | 0 |
| chr1 | 228443317 | ACTCATCTGTGGA-AGCCCCAA-CCCTCAATGTGAG | loxF8L | 3 | loxF8R | 4 | AGCCCCAA/TTGGGGCT | 7 |
| chr2 | 146807230 | GTCACATTGTGGG-TTGGGGCT-TCAACATATTCAT | loxF8R | 4 | loxF8L | 3 | AGCCCCAA/TTGGGGCT | 7 |
| --- | | | | | | | | 0 |
| chr14 | 61611711 | CAAAATCTCTAGA-CTGGGGCT-TCCCCATATTTCT | loxF8L | 4 | loxF8L | 3 | AGCCCCAG/CTGGGGCT | 7 |
| chr10 | 77215708 | ATTCATATGTTGA-AGCCCCAG-TCCCCAGTGTTAT | loxF8L | 4 | loxF8L | 3 | AGCCCCAG/CTGGGGCT | 7 |
| --- | | | | | | | | 0 |
| chr13 | 90828160 | ATAAAATTGAGGG-TTAGGGCT-TCAACATATTAAT | loxF8L | 4 | loxF8L | 3 | AGCCCTAA/TTAGGGCT | 7 |
| chr5 | 84736635 | GTCACATTGGGTG-TTAGGGCT-TCAACACATTGAT | loxF8R | 4 | loxF8L | 3 | AGCCCTAA/TTAGGGCT | 7 |
| chr6 | 110512060 | ATTAATATGTTGA-AGCCCTAA-TTCACAGTGTGAT | loxF8L | 3 | loxF8L | 4 | AGCCCTAA/TTAGGGCT | 7 |
| chr6 | 115524212 | ATTCATATGTGGA-AGCCCTAA-CACCCAATATGAT | loxF8L | 3 | loxF8R | 4 | AGCCCTAA/TTAGGGCT | 7 |
| chr2 | 106715211 | GTAAGATTAGGGG-TTAGGGCT-TCCACATATGGAT | loxF8R | 4 | loxF8L | 3 | AGCCCTAA/TTAGGGCT | 7 |
| chr8 | 21184993 | ATTAATATGTTGA-AGCCCTAA-CACCCAATGTAAT | loxF8L | 3 | loxF8R | 4 | AGCCCTAA/TTAGGGCT | 7 |
| chr1 | 82939703 | GTCAAACTGTGGG-TTAGGGCT-TCAACATATTAAT | loxF8L | 4 | loxF8L | 3 | AGCCCTAA/TTAGGGCT | 7 |
| chr14 | 92234426 | ATTCATCTGTTGA-AGCCCTAA-CCCTCAATGTTAC | loxF8L | 3 | loxF8R | 4 | AGCCCTAA/TTAGGGCT | 7 |
| --- | | | | | | | | 0 |
| chr17 | 7139321 | ATCACACTGTGGA-GTAGGGCT-TCCACCTATGAAT | loxF8L | 3 | loxF8L | 4 | AGCCCTAC/GTAGGGCT | 7 |
| chr13 | 96127111 | ATTCATCTGTTGA-AGCCCTAC-CAAACAATGTGAT | loxF8L | 3 | loxF8R | 4 | AGCCCTAC/GTAGGGCT | 7 |
| --- | | | | | | | | 0 |
| chr8 | 98464364 | AAAACACTGTTGA-TGGAGGCT-GCCACACATTTAA | loxF8L | 4 | loxF8L | 3 | AGCCTCCA/TGGAGGCT | 7 |
| chr12 | 57642690 | ATAAAGATTAGGA-TGGAGGCT-ACCAGAGATTCAT | loxF8L | 4 | loxF8L | 3 | AGCCTCCA/TGGAGGCT | 7 |
| --- | | | | | | | | 0 |
| chr17 | 83222735 | ATATGTGTGTGTG-AGCGTGTG-TGCACAGATGCAT | loxF8R | 4 | loxF8L | 3 | AGCGTGTG/CACACGCT | 7 |
| chr1 | 363812 | ATGCATCTGTGCA-CACACGCT-CACACACACATAT | loxF8L | 3 | loxF8R | 4 | AGCGTGTG/CACACGCT | 7 |
| chr1 | 592876 | ATGCATCTGTGCA-CACACGCT-CACACACACATAT | loxF8L | 3 | loxF8R | 4 | AGCGTGTG/CACACGCT | 7 |
| chr5 | 181456547 | ATATGTGTGTGTG-AGCGTGTG-TGCACAGATGCAT | loxF8R | 4 | loxF8L | 3 | AGCGTGTG/CACACGCT | 7 |
| chr6 | 170729776 | ATATGTGTGTGTG-AGCGTGTG-TGCACAGATGCAT | loxF8R | 4 | loxF8L | 3 | AGCGTGTG/CACACGCT | 7 |
| chr8 | 77546 | ATGCATCTGTGCA-CACACGCT-CACACACACATAT | loxF8L | 3 | loxF8R | 4 | AGCGTGTG/CACACGCT | 7 |
| --- | | | | | | | | 0 |
| chr19 | 42574450 | GTATAATTTTGTG-AGCTAAAT-TTGAAAGATTTAT | loxF8R | 4 | loxF8L | 3 | AGCTAAAT/ATTTAGCT | 7 |
| chr14 | 33604815 | TCAAATATGTGGA-AGCTAAAT-AACACAATCCAAA | loxF8L | 3 | loxF8R | 4 | AGCTAAAT/ATTTAGCT | 7 |
| --- | | | | | | | | 0 |
| chr6 | 134133245 | CTTATATTCTGTC-TTAGAGCT-GCCAGAGATTTTT | loxF8R | 4 | loxF8L | 3 | AGCTCTAA/TTAGAGCT | 7 |
| chr4 | 57902699 | ATACATATGTTGA-AGCTCTAA-CCCACAATGTGAT | loxF8L | 3 | loxF8R | 4 | AGCTCTAA/TTAGAGCT | 7 |
| --- | | | | | | | | 0 |
| chr9 | 28281663 | ATAGAGATGTGGA-TAGCAGCT-CATACAAGGATAG | loxF8L | 3 | loxF8R | 4 | AGCTGCTA/TAGCAGCT | 7 |
| chr18 | 30541680 | CTTAGTTAGTGTG-AGCTGCTA-CACAAAGTAATAG | loxF8R | 3 | loxF8R | 4 | AGCTGCTA/TAGCAGCT | 7 |
| --- | | | | | | | | 0 |
| chr12 | 31423766 | AGAAGTCTGGTGA-CCCAAGCT-CACTCATTCCTAG | loxF8L | 4 | loxF8R | 3 | AGCTTGGG/CCCAAGCT | 7 |
| chr6 | 15780769 | ATACAGCTGTGAA-AGCTTGGG-CAAATTGTCTTAG | loxF8L | 3 | loxF8R | 4 | AGCTTGGG/CCCAAGCT | 7 |
| --- | | | | | | | | 0 |
| chr3 | 112896286 | AAGAAACTGTTGA-TTGTTCCT-TCTGCAGAGTTAT | loxF8L | 4 | loxF8L | 3 | AGGAACAA/TTGTTCCT | 7 |
| chr10 | 112620118 | TTAACAGTGTGTG-TTGTTCCT-TTCACTTAATTAT | loxF8R | 3 | loxF8L | 4 | AGGAACAA/TTGTTCCT | 7 |
| --- | | | | | | | | 0 |
| chr11 | 3255109 | CTGTTATTGTGTG-AGGAACCA-GGCAGAGATTTGT | loxF8R | 3 | loxF8L | 4 | AGGAACCA/TGGTTCCT | 7 |
| chr11 | 3300912 | CTGTTATTGTGTG-AGGAACCA-GGCAGAGATTTGT | loxF8R | 3 | loxF8L | 4 | AGGAACCA/TGGTTCCT | 7 |
| --- | | | | | | | | 0 |

| | | | | | | | | |
|---|---|---|---|---|---|---|---|---|
| chr9 | 78943769 | AAAAATCCCTGGC-CCCTTCCT-TCTAAGGATTTAT | loxF8L | 4 | loxF8L | 3 | AGGAAGGG/CCCTTCCT | 7 |
| chr13 | 97828865 | AGATTTCTGTGGA-AGGAAGGG-TCCAGATATTGAA | loxF8L | 3 | loxF8L | 4 | AGGAAGGG/CCCTTCCT | 7 |
| --- | | | | | | | | 0 |
| chr6 | 108489241 | AGAAATCTGGGAA-AGGAGAGA-ACCACAGAGGAAT | loxF8L | 3 | loxF8L | 4 | AGGAGAGA/TCTCTCCT | 7 |
| chr16 | 70963124 | CTCTGATGGTGTG-TCTCTCCT-CACACTCTCTTCA | loxF8R | 3 | loxF8R | 4 | AGGAGAGA/TCTCTCCT | 7 |
| --- | | | | | | | | 0 |
| chr4 | 95813075 | CTACTATTATGTC-AGGATAAA-ACAACAAATTTAT | loxF8R | 4 | loxF8L | 3 | AGGATAAA/TTTATCCT | 7 |
| chr6 | 141444491 | CTATGAGTGTATG-AGGATAAA-AATATAATCTTAT | loxF8R | 3 | loxF8R | 4 | AGGATAAA/TTTATCCT | 7 |
| --- | | | | | | | | 0 |
| chrX | 120589543 | AAAAATTTCTGGA-AGGATATA-CACACAAATGTAC | loxF8L | 3 | loxF8R | 4 | AGGATATA/TATATCCT | 7 |
| chr2 | 170893276 | TTAATGTTTTGTG-TATATCCT-TCCAGACTTTTAT | loxF8R | 4 | loxF8L | 3 | AGGATATA/TATATCCT | 7 |
| --- | | | | | | | | 0 |
| chr1 | 50523395 | CTAAATTTGTGTC-TACATCCT-CATAAATTCTTAT | loxF8R | 3 | loxF8R | 4 | AGGATGTA/TACATCCT | 7 |
| chrX | 31704845 | CTGAGATTGAGTT-AGGATGTA-TACAAAATCAAAG | loxF8R | 3 | loxF8R | 4 | AGGATGTA/TACATCCT | 7 |
| --- | | | | | | | | 0 |
| chr19 | 32190269 | AAAAATGTGGGGA-AGGTACTC-CCCACAGAGTGAA | loxF8L | 3 | loxF8L | 4 | AGGTACTC/GAGTACCT | 7 |
| chr8 | 29233502 | CTAAAATTGCGGT-GAGTACCT-CAAACAATTTTAA | loxF8R | 4 | loxF8R | 3 | AGGTACTC/GAGTACCT | 7 |
| --- | | | | | | | | 0 |
| chr4 | 175211386 | CTACCATTGTGTC-CAGAACCT-ACCTCAGATTACT | loxF8R | 3 | loxF8L | 4 | AGGTTCTG/CAGAACCT | 7 |
| chr11 | 87444627 | TTAATTTTGGGGA-AGGTTCTG-GCCACAGAGTCAT | loxF8L | 4 | loxF8L | 3 | AGGTTCTG/CAGAACCT | 7 |
| --- | | | | | | | | 0 |
| chr14 | 40647083 | ACAAATATGTGTA-AGTAAAAC-CAGACACTCCTAA | loxF8L | 3 | loxF8R | 4 | AGTAAAAC/GTTTTACT | 7 |
| chr21 | 22701604 | TTAAATTTTTTGA-GTTTTACT-TACTCAGTTTTAT | loxF8L | 4 | loxF8L | 3 | AGTAAAAC/GTTTTACT | 7 |
| --- | | | | | | | | 0 |
| chr14 | 79047899 | CTAAGACTTCGTA-AGTTTACT-TTTACATATTTAT | loxF8R | 4 | loxF8L | 3 | AGTAAACT/AGTTTACT | 7 |
| chr2 | 209007026 | ATAAATTTATGTA-AGTAAACT-CAGTCAATCTGTG | loxF8L | 3 | loxF8R | 4 | AGTAAACT/AGTTTACT | 7 |
| --- | | | | | | | | 0 |
| chr15 | 55361025 | ATAAGCTTTTGTG-TCTTTACT-TCTACAGATACAA | loxF8R | 3 | loxF8L | 4 | AGTAAAGA/TCTTTACT | 7 |
| chr12 | 79026330 | ATAGAAATGAGGA-AGTAAAGA-TCCAAAGAGTTAG | loxF8L | 4 | loxF8L | 3 | AGTAAAGA/TCTTTACT | 7 |
| --- | | | | | | | | 0 |
| chr1 | 220737625 | CTATGCTTGTGGG-GTGTTACT-CAAGAAATCTTTG | loxF8R | 3 | loxF8R | 4 | AGTAACAC/GTGTTACT | 7 |
| chr3 | 163136752 | CTGAGCTTGTGGG-GTGTTACT-CAAGAAATCTTTG | loxF8R | 3 | loxF8R | 4 | AGTAACAC/GTGTTACT | 7 |
| --- | | | | | | | | 0 |
| chr10 | 120353963 | CAAAGATTTCTTG-AGTAATAA-CCCACAAGCCTAG | loxF8R | 4 | loxF8R | 3 | AGTAATAA/TTATTACT | 7 |
| chr15 | 94216423 | CAAAGATTTTTTG-AGTAATAA-CCCACAGTTATAG | loxF8R | 3 | loxF8L | 4 | AGTAATAA/TTATTACT | 7 |
| --- | | | | | | | | 0 |
| chr5 | 18647187 | CTATGCTTGTGGG-ATATTACT-CAAGAAATCTTTG | loxF8R | 3 | loxF8R | 4 | AGTAATAT/ATATTACT | 7 |
| chr4 | 69904998 | CAAAGATTTTTTG-AGTAATAT-CCCACAAGCACAG | loxF8R | 3 | loxF8R | 4 | AGTAATAT/ATATTACT | 7 |
| chr9 | 108736999 | GAAAGATTATTTG-AGTAATAT-CCCACAAGCATAG | loxF8R | 4 | loxF8R | 3 | AGTAATAT/ATATTACT | 7 |
| --- | | | | | | | | 0 |
| chr4 | 63629748 | CAAAGATTTTTTG-AGTAATGC-CCCAGACTCATAG | loxF8R | 3 | loxF8R | 4 | AGTAATGC/GCATTACT | 7 |
| chr3 | 136823448 | CTATGCTGGTGTG-GCATTACT-CAAGAAATTTTAG | loxF8R | 3 | loxF8R | 4 | AGTAATGC/GCATTACT | 7 |
| --- | | | | | | | | 0 |
| chr1 | 160603295 | CTTAGATTGTCTA-TTTGTACT-CCTTCAGATTTTT | loxF8R | 3 | loxF8L | 4 | AGTACAAA/TTTGTACT | 7 |
| chr5 | 159624791 | AAAAATCTGAAGG-AGTACAAA-TAGACAATCTAAG | loxF8L | 4 | loxF8R | 3 | AGTACAAA/TTTGTACT | 7 |
| --- | | | | | | | | 0 |
| chr11 | 102667841 | ATATATATATGCA-TATGTACT-TGCACATATATAT | loxF8L | 4 | loxF8L | 3 | AGTACATA/TATGTACT | 7 |
| chr3 | 153334034 | ATATATATATGTA-TATGTACT-TCCATATATATAT | loxF8L | 4 | loxF8L | 3 | AGTACATA/TATGTACT | 7 |
| --- | | | | | | | | 0 |
| chr6 | 77459487 | GTAATATTGAGTG-AGTATACA-TTCATATATTTCT | loxF8R | 3 | loxF8L | 4 | AGTATACA/TGTATACT | 7 |
| chr20 | 8161284 | CATACATAGTGTG-TGTATACT-TACACATATATAT | loxF8R | 4 | loxF8L | 3 | AGTATACA/TGTATACT | 7 |
| --- | | | | | | | | 0 |
| chr5 | 118761635 | ATAAATATATGTA-AGTATATA-TATATACATTTAT | loxF8L | 3 | loxF8L | 4 | AGTATATA/TATATACT | 7 |
| chr1 | 239899465 | ATATATGTGTGTA-TATATACT-CACACACATGTAT | loxF8L | 3 | loxF8L | 4 | AGTATATA/TATATACT | 7 |
| --- | | | | | | | | 0 |
| chrY | 16036002 | AAAATTCTGGGGA-TTAGGACT-TCCACACATGAGT | loxF8L | 3 | loxF8L | 4 | AGTCCTAA/TTAGGACT | 7 |
| chr2 | 156624838 | ATCAAGCTGGGTA-TTAGGACT-TCAACATATTAAT | loxF8L | 4 | loxF8L | 3 | AGTCCTAA/TTAGGACT | 7 |
| chr5 | 61230452 | ATCAATATGTTGA-AGTCCTAA-CCCCCAGAGTGAT | loxF8L | 3 | loxF8L | 4 | AGTCCTAA/TTAGGACT | 7 |
| chr13 | 91788374 | ATCAATTTGGGGA-TTAGGACT-TCAACATATGAAT | loxF8L | 3 | loxF8L | 4 | AGTCCTAA/TTAGGACT | 7 |
| chr13 | 48812120 | ATCACACTGTGGA-TTAGGACT-TCAACATATGAAT | loxF8L | 3 | loxF8L | 4 | AGTCCTAA/TTAGGACT | 7 |
| --- | | | | | | | | 0 |
| chr18 | 10332328 | CTAAGCTTATTTG-AGTCCTGA-CCCCCAACCTGAG | loxF8R | 3 | loxF8R | 4 | AGTCCTGA/TCAGGACT | 7 |
| chr14 | 72045924 | GTTTATATGTGGA-AGTCCTGA-CACCCAATATGAG | loxF8L | 4 | loxF8R | 3 | AGTCCTGA/TCAGGACT | 7 |
| --- | | | | | | | | 0 |
| chr3 | 5270867 | ACACAACTATGGA-AGTGACAT-CACGTAATCTTTG | loxF8L | 4 | loxF8R | 3 | AGTGACAT/ATGTCACT | 7 |
| chrX | 78268068 | CTAAGATAGTGCT-AGTGACAT-GCAAAAGATTTCT | loxF8R | 3 | loxF8L | 4 | AGTGACAT/ATGTCACT | 7 |
| --- | | | | | | | | 0 |
| chr18 | 53297924 | ATAAAACTGATAA-TAATCACT-TCCAAAGATGAAT | loxF8L | 4 | loxF8L | 3 | AGTGATTA/TAATCACT | 7 |
| chr19 | 35446206 | TTAAGAATGTGGA-AGTGATTA-CACAGAACTTTAG | loxF8L | 4 | loxF8R | 3 | AGTGATTA/TAATCACT | 7 |
| --- | | | | | | | | 0 |
| chr2 | 172899327 | AGAAATCTGTTTT-AGTGCATG-ACCAAAGATTAAT | loxF8L | 4 | loxF8L | 3 | AGTGCATG/CATGCACT | 7 |
| chr18 | 25026035 | ATAATTCTATGAT-CATGCACT-TACAAAATCTTAA | loxF8L | 4 | loxF8R | 3 | AGTGCATG/CATGCACT | 7 |
| --- | | | | | | | | 0 |
| chr6 | 7689689 | ATGAACTTGTGTG-AGTGTGTG-CGCACAATCTTGT | loxF8R | 4 | loxF8R | 3 | AGTGTGTG/CACACACT | 7 |
| chr4 | 120325485 | ACACATATGTGCA-CACACACT-CACACAAACACAG | loxF8L | 4 | loxF8R | 3 | AGTGTGTG/CACACACT | 7 |
| --- | | | | | | | | 0 |
| chr5 | 16164679 | ATAAATAACTTGA-AGTTAAAG-TCCACATGTTTTT | loxF8L | 4 | loxF8L | 3 | AGTTAAAG/CTTTAACT | 7 |
| chr12 | 45266182 | ACAAATCTTTGCA-AGTTAAAG-ACTATAGGTTTAT | loxF8L | 3 | loxF8L | 4 | AGTTAAAG/CTTTAACT | 7 |
| --- | | | | | | | | 0 |
| chr2 | 124406100 | ATAAAACTGTGTT-GATTAACT-TCCAAATATTTGA | loxF8L | 3 | loxF8L | 4 | AGTTAATC/GATTAACT | 7 |
| chr4 | 172715848 | ATATGACTTTGTG-AGTTAATC-CAAACAAACTAAG | loxF8R | 4 | loxF8R | 3 | AGTTAATC/GATTAACT | 7 |
| chr12 | 21146701 | TTAAAAGTGTGGT-AGTTAATC-TCCACATATTTTG | loxF8L | 4 | loxF8L | 3 | AGTTAATC/GATTAACT | 7 |
| --- | | | | | | | | 0 |
| chr8 | 53630097 | CGAATATGGTGTG-AGTTAGAG-TCCACTGGGCTAT | loxF8R | 3 | loxF8L | 4 | AGTTAGAG/CTCTAACT | 7 |
| chr4 | 70251492 | CAAATTTAGTGTG-AGTTAGAG-TACACAAATTTAC | loxF8R | 4 | loxF8L | 3 | AGTTAGAG/CTCTAACT | 7 |
| --- | | | | | | | | 0 |
| chr13 | 97653496 | ATAATTCTGAGGC-AGTTGGGG-TCCACAGAAGCCT | loxF8L | 3 | loxF8L | 4 | AGTTGGGG/CCCCAACT | 7 |

| | | | | | | | | |
|---|---|---|---|---|---|---|---|---|
| chr13 | 98712755 | CTAAGAGCGTTTG-AGTTGGGG-TGAACAGAATTAA | loxF8R | 3 | loxF8L | 4 | AGTTGGGG/CCCCAACT | 7 |
| --- | | | | | | | | 0 |
| chr7 | 93685426 | AAAAATATTAGGA-AGTTTAAA-TCAACATAGTTAT | loxF8L | 4 | loxF8L | 3 | AGTTTAAA/TTTAAACT | 7 |
| chr4 | 181400544 | CTAAAAGAGTGTT-AGTTTAAA-CACACATGCTTGG | loxF8R | 4 | loxF8R | 3 | AGTTTAAA/TTTAAACT | 7 |
| --- | | | | | | | | 0 |
| chr12 | 10205100 | CTAATAATGGGAG-TTAAAACT-TCCACATCTTTTT | loxF8R | 4 | loxF8L | 3 | AGTTTTAA/TTAAAACT | 7 |
| chr3 | 98384033 | TTTTATCTGTGGA-AGTTTTAA-CAAGCAATCATAT | loxF8L | 3 | loxF8R | 4 | AGTTTTAA/TTAAAACT | 7 |
| --- | | | | | | | | 0 |
| chr2 | 26955454 | TTCAGATTATGTG-AGTTTTAT-TGAACAAATTGAT | loxF8R | 3 | loxF8L | 4 | AGTTTTAT/ATAAAACT | 7 |
| chr5 | 91654757 | CCATGATCGTGTG-AGTTTTAT-TCCAGAGATGCAA | loxF8R | 3 | loxF8L | 4 | AGTTTTAT/ATAAAACT | 7 |
| --- | | | | | | | | 0 |
| chr11 | 40468975 | AAAAATCTGCAAA-ATAAAAAT-AATACAATGTTAG | loxF8L | 4 | loxF8R | 3 | ATAAAAAT/ATTTTTAT | 7 |
| chr13 | 42256587 | ATAAATCTGACTA-ATAAAAAT-GCAAAAGCTTTAT | loxF8L | 3 | loxF8L | 4 | ATAAAAAT/ATTTTTAT | 7 |
| chr16 | 34929687 | ATAAATATTTGAA-ATAAAAAT-TCCAGAGATAGAA | loxF8L | 3 | loxF8L | 4 | ATAAAAAT/ATTTTTAT | 7 |
| chr14 | 43198274 | ATAAATCTTGTCA-ATAAAAAT-TACATAAATTTAT | loxF8L | 4 | loxF8L | 3 | ATAAAAAT/ATTTTTAT | 7 |
| chr5 | 111354540 | CTAAATGTGTGTA-ATTTTTAT-TTCTCAGTTTTAC | loxF8L | 3 | loxF8L | 4 | ATAAAAAT/ATTTTTAT | 7 |
| --- | | | | | | | | 0 |
| chrX | 47688701 | ATAAACGTGTGGC-TATTTTAT-TGTACAGATTATT | loxF8L | 3 | loxF8L | 4 | ATAAAATA/TATTTTAT | 7 |
| chr4 | 165921261 | ATAAAAATCAGGA-TATTTTAT-CCAACAACCTTAG | loxF8L | 4 | loxF8R | 3 | ATAAAATA/TATTTTAT | 7 |
| chr15 | 94221916 | GTAAATTTCTTGA-ATAAAATA-CAAAGAATCTTTG | loxF8L | 4 | loxF8R | 3 | ATAAAATA/TATTTTAT | 7 |
| chr13 | 46757196 | CTAGAAATGTGTA-ATAAAATA-AACACATACTTAG | loxF8R | 4 | loxF8R | 3 | ATAAAATA/TATTTTAT | 7 |
| --- | | | | | | | | 0 |
| chr18 | 41807133 | CTAAGATAAAGTC-TGGTTTAT-CACAGAATCACAG | loxF8R | 4 | loxF8R | 3 | ATAAACCA/TGGTTTAT | 7 |
| chr18 | 63381625 | TTAAGATTTTGAG-TGGTTTAT-TATACACTCATAG | loxF8R | 3 | loxF8R | 4 | ATAAACCA/TGGTTTAT | 7 |
| --- | | | | | | | | 0 |
| chrX | 85270322 | CTATGCTTGTTTG-CTCTTTAT-TCCAAAGATACTT | loxF8R | 3 | loxF8L | 4 | ATAAAGAG/CTCTTTAT | 7 |
| chr10 | 26005954 | ATGAATAAGTGGA-CTCTTTAT-TTTAGAGCTTTAT | loxF8L | 3 | loxF8L | 4 | ATAAAGAG/CTCTTTAT | 7 |
| --- | | | | | | | | 0 |
| chr6 | 94983221 | AAAATCCTGTGGA-ATAAATAG-CTCACCTTCTTAA | loxF8L | 3 | loxF8R | 4 | ATAAATAG/CTATTTAT | 7 |
| chr5 | 143332104 | ATTAATTTGTAGC-ATAAATAG-CTGACAACCTTAG | loxF8L | 4 | loxF8R | 3 | ATAAATAG/CTATTTAT | 7 |
| --- | | | | | | | | 0 |
| chrX | 91370346 | ATAAATATCTAGA-ATAAATCC-CACATAGACATAG | loxF8L | 3 | loxF8R | 4 | ATAAATCC/GGATTTAT | 7 |
| chr7 | 14249173 | TTACATCCTTGGA-ATAAATCC-CACTTAATCATAG | loxF8L | 4 | loxF8R | 3 | ATAAATCC/GGATTTAT | 7 |
| --- | | | | | | | | 0 |
| chr11 | 98923337 | AAAGAGCTGTGGC-TAATTTAT-TCCATAGAGTTAG | loxF8L | 4 | loxF8L | 3 | ATAAATTA/TAATTTAT | 7 |
| chr20 | 52149643 | ACAATTCTGTGTA-TAATTTAT-TCCAAAGAGTAAG | loxF8L | 3 | loxF8L | 4 | ATAAATTA/TAATTTAT | 7 |
| chr13 | 81126897 | AGATATATGTTGA-TAATTTAT-TCTACAGCTTTAC | loxF8L | 4 | loxF8L | 3 | ATAAATTA/TAATTTAT | 7 |
| chr6 | 138306163 | ATATATATGAGAA-ATAAATTA-TCCAAAGATAAAT | loxF8L | 4 | loxF8L | 3 | ATAAATTA/TAATTTAT | 7 |
| --- | | | | | | | | 0 |
| chr3 | 3892189 | ATAAATCACTAGA-ATAACAAG-ACAATAGATTTAA | loxF8L | 3 | loxF8L | 4 | ATAACAAG/CTTGTTAT | 7 |
| chr13 | 68030649 | CTAAGATTCTGCT-CTTGTTAT-GACACCATCTTTA | loxF8R | 3 | loxF8R | 4 | ATAACAAG/CTTGTTAT | 7 |
| --- | | | | | | | | 0 |
| chr6 | 159247402 | TTAGATCTGTTGA-TTTCTTAT-CATAGAATCTTTT | loxF8L | 3 | loxF8R | 4 | ATAAGAAA/TTTCTTAT | 7 |
| chr7 | 8609152 | ATATATCTCTGGC-TTTCTTAT-TCTACAGATGGTT | loxF8L | 3 | loxF8L | 4 | ATAAGAAA/TTTCTTAT | 7 |
| --- | | | | | | | | 0 |
| chr10 | 22362023 | ATAAATATATAGA-TATATTAT-TTTATATATTTAT | loxF8L | 3 | loxF8L | 4 | ATAATATA/TATATTAT | 7 |
| chrX | 26378599 | CTTGTATTGAGTG-ATAATATA-CACACACTCACAG | loxF8R | 4 | loxF8R | 3 | ATAATATA/TATATTAT | 7 |
| --- | | | | | | | | 0 |
| chr14 | 41816258 | ATATATATATGCA-TAGATTAT-TGAACAGATTTTT | loxF8L | 4 | loxF8L | 3 | ATAATCTA/TAGATTAT | 7 |
| chr20 | 1239693 | CTAGTATGGTTTG-ATAATCTA-CATACAATTTGAG | loxF8R | 4 | loxF8R | 3 | ATAATCTA/TAGATTAT | 7 |
| chr4 | 171774018 | TGAAGATTTTGTG-TAGATTAT-CCAAGAGATTTAC | loxF8R | 3 | loxF8L | 4 | ATAATCTA/TAGATTAT | 7 |
| --- | | | | | | | | 0 |
| chr1 | 83370530 | ATACATATGTTAA-TTCATTAT-TCCACAAATATTT | loxF8L | 4 | loxF8L | 3 | ATAATGAA/TTCATTAT | 7 |
| chr7 | 76826672 | ATACATATGTTAA-TTCATTAT-TCCACAAATATTT | loxF8L | 4 | loxF8L | 3 | ATAATGAA/TTCATTAT | 7 |
| --- | | | | | | | | 0 |
| chr16 | 65583031 | ATATACTTCTGGA-TTAATTAT-TCAACATATTTTT | loxF8L | 4 | loxF8L | 3 | ATAATTAA/TTAATTAT | 7 |
| chr18 | 58062465 | CTGAGTTTATGTT-TTAATTAT-GGCACAGATTAAT | loxF8R | 4 | loxF8L | 3 | ATAATTAA/TTAATTAT | 7 |
| chr17 | 56856435 | CTAAGATTGACAT-TTAATTAT-GACTCAAGCTTAG | loxF8R | 4 | loxF8R | 3 | ATAATTAA/TTAATTAT | 7 |
| --- | | | | | | | | 0 |
| chr3 | 150356787 | ATCACTCTGTGGG-ATAATTAT-TTCACAGTCTGAG | loxF8L | 3 | loxF8R | 4 | ATAATTAT/ATAATTAT | 7 |
| --- | | | | | | | | 0 |
| chr4 | 28968063 | AAATATATGTGCA-TCAATTAT-TAAACAATCTTAT | loxF8L | 4 | loxF8R | 3 | ATAATTGA/TCAATTAT | 7 |
| chr16 | 80655141 | CTAAGGCTGGGTG-TCAATTAT-TGAACAGTCTTAT | loxF8R | 3 | loxF8L | 4 | ATAATTGA/TCAATTAT | 7 |
| chr15 | 58892042 | GTAAAGTTGTGTG-TCAATTAT-TGTACAGAATAAT | loxF8R | 3 | loxF8L | 4 | ATAATTGA/TCAATTAT | 7 |
| --- | | | | | | | | 0 |
| chr13 | 90499655 | AGAATTCTCTGCA-ATACAAAT-TTCACATATTTTT | loxF8L | 4 | loxF8L | 3 | ATACAAAT/ATTTGTAT | 7 |
| chr10 | 92330558 | CTGAAATTGTCTT-ATTTGTAT-TCCAGAGCTATAT | loxF8R | 4 | loxF8L | 3 | ATACAAAT/ATTTGTAT | 7 |
| --- | | | | | | | | 0 |
| chr18 | 43950181 | ACAAATCTGTTTG-TAATGTAT-TCTGCAGCTTTAT | loxF8L | 4 | loxF8L | 3 | ATACATTA/TAATGTAT | 7 |
| chrX | 143773002 | ATAATACTGTCGA-TAATGTAT-CAAAATATCTTAA | loxF8L | 3 | loxF8R | 4 | ATACATTA/TAATGTAT | 7 |
| --- | | | | | | | | 0 |
| chr16 | 80403509 | ATAACTCTGAGTA-TTGAGTAT-TTCAAATATTTTT | loxF8L | 3 | loxF8L | 4 | ATACTCAA/TTGAGTAT | 7 |
| chr11 | 80780150 | ATAACCCTGTGCA-TTGAGTAT-TACATTATCTTTG | loxF8L | 3 | loxF8R | 4 | ATACTCAA/TTGAGTAT | 7 |
| --- | | | | | | | | 0 |
| chr5 | 51095107 | ACAAATCTATGAA-TTTTCTAT-CACACATTTTTTT | loxF8L | 3 | loxF8R | 4 | ATAGAAAA/TTTTCTAT | 7 |
| chr9 | 131565948 | CTAAGATTTGATG-TTTTCTAT-TCCTCATATGTTT | loxF8R | 3 | loxF8L | 4 | ATAGAAAA/TTTTCTAT | 7 |
| chr10 | 73280874 | CTATTTTTGTCTG-TTTTCTAT-TTCACTGATTTCT | loxF8R | 4 | loxF8L | 3 | ATAGAAAA/TTTTCTAT | 7 |
| chr11 | 107451451 | TTAAATCAATGAA-ATAGAAAA-CACACAACCAGAG | loxF8L | 4 | loxF8R | 3 | ATAGAAAA/TTTTCTAT | 7 |
| --- | | | | | | | | 0 |
| chr17 | 67537288 | ATATGATTGTCTG-ATAGAAAT-TTCAGACATTTCT | loxF8R | 3 | loxF8L | 4 | ATAGAAAT/ATTTCTAT | 7 |
| chr7 | 108617160 | CTAAGTATGTGTA-ATTTCTAT-AAAACAATCAGAG | loxF8R | 3 | loxF8R | 4 | ATAGAAAT/ATTTCTAT | 7 |
| --- | | | | | | | | 0 |
| chr2 | 204942741 | ATAAATATTTGAA-ATAGGAAA-TCCCTAAACTTAT | loxF8L | 3 | loxF8L | 4 | ATAGGAAA/TTTCCTAT | 7 |
| chr17 | 54993685 | TAGAATCTGTGGA-ATAGGAAA-CACAAACTCATAA | loxF8L | 3 | loxF8R | 4 | ATAGGAAA/TTTCCTAT | 7 |
| --- | | | | | | | | 0 |
| chr4 | 164324469 | ATAAAACTTTGAT-GAACCTAT-TATACAATTTTAG | loxF8L | 4 | loxF8R | 3 | ATAGGTTC/GAACCTAT | 7 |
| chr5 | 33269303 | CTAAACATGTGGC-ATAGGTTC-TACACAAATGTAT | loxF8L | 4 | loxF8L | 3 | ATAGGTTC/GAACCTAT | 7 |

| | | | | | | | | |
|---|---|---|---|---|---|---|---|---|
| --- | | | | | | | | 0 |
| chr21 | 9558001 | CTCAGATTCAGTG-CATACTAT-TACAGAATATGAG | loxF8R | 3 | loxF8R | 4 | ATAGTATG/CATACTAT | 7 |
| chr17 | 21640196 | CTTAGATTCAGTG-CATACTAT-TACAGAATATGAG | loxF8R | 3 | loxF8R | 4 | ATAGTATG/CATACTAT | 7 |
| chr17 | 22063093 | CTCATATTCTGTA-ATAGTATG-CACTGAATCTAAG | loxF8R | 4 | loxF8R | 3 | ATAGTATG/CATACTAT | 7 |
| --- | | | | | | | | 0 |
| chr18 | 33193828 | CTAAGACTGGGTA-ATATATAA-TCAACAGAAATTT | loxF8R | 3 | loxF8L | 4 | ATATATAA/TTATATAT | 7 |
| chr6 | 165374395 | AGAAATATGTGTA-ATATATAA-CAGAAAATTTTAA | loxF8L | 3 | loxF8R | 4 | ATATATAA/TTATATAT | 7 |
| chr16 | 63792875 | TTATATCTGTGGT-TTATATAT-CACAGGATTTCAG | loxF8L | 3 | loxF8R | 4 | ATATATAA/TTATATAT | 7 |
| --- | | | | | | | | 0 |
| chr12 | 90485174 | ATATATGTGTGTA-ATATATTA-TCCATATATGTGT | loxF8L | 3 | loxF8L | 4 | ATATATTA/TAATATAT | 7 |
| chr8 | 31672861 | ATCAATCTTTGAA-TAATATAT-TTCCAAGATATAT | loxF8L | 3 | loxF8L | 4 | ATATATTA/TAATATAT | 7 |
| --- | | | | | | | | 0 |
| chr5 | 86629168 | AAAAAATTGTGGA-ATATGGAA-TAAAAAGATGTAT | loxF8L | 3 | loxF8L | 4 | ATATGGAA/TTCCATAT | 7 |
| chr5 | 66735967 | ACAAGATAGGGTG-ATATGGAA-CATTTAATCTTAG | loxF8R | 4 | loxF8R | 3 | ATATGGAA/TTCCATAT | 7 |
| --- | | | | | | | | 0 |
| chr3 | 89475667 | TTAAGTGTGTGGA-ATGAATAT-TAAGCAGATTCAT | loxF8L | 3 | loxF8L | 4 | ATATTCAT/ATGAATAT | 7 |
| chr2 | 145332218 | GTAATTCTTTGGA-ATGAATAT-TTCCCAGTTTTAC | loxF8L | 3 | loxF8L | 4 | ATATTCAT/ATGAATAT | 7 |
| --- | | | | | | | | 0 |
| chr4 | 68023204 | CCAAGATGGTTTT-CTCAATAT-TCCACTGATCTGT | loxF8R | 4 | loxF8L | 3 | ATATTGAG/CTCAATAT | 7 |
| chr14 | 84769268 | TTCAGATTGCTTG-CTCAATAT-ACAACAGAATTAT | loxF8R | 4 | loxF8L | 3 | ATATTGAG/CTCAATAT | 7 |
| --- | | | | | | | | 0 |
| chr2 | 221967553 | ATAAATAAATTGA-ATATTTGA-TCCATATATTTCT | loxF8L | 4 | loxF8L | 3 | ATATTTGA/TCAAATAT | 7 |
| chr6 | 76827412 | GTAAATCTTTTGT-TCAAATAT-TCTAAAGATTTAC | loxF8L | 4 | loxF8L | 3 | ATATTTGA/TCAAATAT | 7 |
| --- | | | | | | | | 0 |
| chr10 | 65496793 | ACAATTCTGTGGT-ATATTTGG-CAGACACTTTTTG | loxF8L | 3 | loxF8R | 4 | ATATTTGG/CCAAATAT | 7 |
| chr1 | 101799157 | AGAAATGTTGGGA-ATATTTGG-TCCATGGATTTCT | loxF8L | 4 | loxF8L | 3 | ATATTTGG/CCAAATAT | 7 |
| --- | | | | | | | | 0 |
| chr7 | 57545822 | ATCAAGCTGTGTA-TTCTTGAT-ACCACAGAATATT | loxF8L | 3 | loxF8L | 4 | ATCAAGAA/TTCTTGAT | 7 |
| chr4 | 180840953 | ATGAATCTGTATC-TTCTTGAT-GCCACAGCTTTAA | loxF8L | 4 | loxF8L | 3 | ATCAAGAA/TTCTTGAT | 7 |
| --- | | | | | | | | 0 |
| chr17 | 32604123 | TTAGATATGTGGA-CATCTGAT-TCCAAATATGTGT | loxF8L | 3 | loxF8L | 4 | ATCAGATG/CATCTGAT | 7 |
| chr8 | 128738611 | CCAAATTTATGTG-CATCTGAT-TCCAAAAATCTAT | loxF8R | 4 | loxF8L | 3 | ATCAGATG/CATCTGAT | 7 |
| --- | | | | | | | | 0 |
| chr4 | 72207413 | ATAAATGTGGAGA-ATCATGAT-CTCATCATCTTGG | loxF8L | 3 | loxF8R | 4 | ATCATGAT/ATCATGAT | 7 |
| --- | | | | | | | | 0 |
| chr7 | 140916957 | ATAAAACAGTAGA-ATCCAAAA-CTCACAAACATAA | loxF8L | 3 | loxF8R | 4 | ATCCAAAA/TTTTGGAT | 7 |
| chr3 | 75248298 | CTCAGTTTGTCTG-ATCCAAAA-TGGAAAAATTTAT | loxF8R | 3 | loxF8L | 4 | ATCCAAAA/TTTTGGAT | 7 |
| --- | | | | | | | | 0 |
| chr3 | 33948319 | CTAAGCATGTATA-GTTAGGAT-AACACAGATTTTT | loxF8R | 4 | loxF8L | 3 | ATCCTAAC/GTTAGGAT | 7 |
| chr5 | 145533772 | CTGAAATTCTGTA-GTTAGGAT-TCCAAATATTTGT | loxF8R | 4 | loxF8L | 3 | ATCCTAAC/GTTAGGAT | 7 |
| --- | | | | | | | | 0 |
| chr4 | 94679952 | AGAAATCATTGTA-ATCCTCAC-CCCACACATTTTT | loxF8L | 4 | loxF8L | 3 | ATCCTCAC/GTGAGGAT | 7 |
| chr4 | 119265146 | GTAAGATTTTGGG-GTGAGGAT-TACACAATTGTAT | loxF8R | 3 | loxF8L | 4 | ATCCTCAC/GTGAGGAT | 7 |
| --- | | | | | | | | 0 |
| chr9 | 11466534 | CAAAGTCTGTGGA-ATCTAAAT-TCCAGATTTTTTT | loxF8L | 3 | loxF8L | 4 | ATCTAAAT/ATTTAGAT | 7 |
| chr5 | 122797879 | ATATATATGTTGA-ATCTAAAT-CCTACAATTTTTG | loxF8L | 3 | loxF8R | 4 | ATCTAAAT/ATTTAGAT | 7 |
| --- | | | | | | | | 0 |
| chr5 | 20890620 | CAAAGATTATCTG-ATCTGTTG-TTCAGAATCTTAA | loxF8R | 3 | loxF8R | 4 | ATCTGTTG/CAACAGAT | 7 |
| chr5 | 34314667 | TTAAGATTCTGAA-CAACAGAT-CAGATAATCTTTG | loxF8R | 4 | loxF8R | 3 | ATCTGTTG/CAACAGAT | 7 |
| --- | | | | | | | | 0 |
| chr5 | 132168345 | CTCAGACTGGGTG-ATCTTAAG-TTCACTGATTCTT | loxF8R | 3 | loxF8L | 4 | ATCTTAAG/CTTAAGAT | 7 |
| chr14 | 28014709 | ATTAGTCTATGGA-ATCTTAAG-TAGACAAACTTAC | loxF8L | 3 | loxF8R | 4 | ATCTTAAG/CTTAAGAT | 7 |
| --- | | | | | | | | 0 |
| chr2 | 143248619 | ATACAGTTGTGAA-TGTAAGAT-TTCACAGAATCAT | loxF8L | 4 | loxF8L | 3 | ATCTTACA/TGTAAGAT | 7 |
| chr4 | 132370054 | CTATGATTTTCAG-ATCTTACA-CAAAAAATCTTTG | loxF8R | 4 | loxF8R | 3 | ATCTTACA/TGTAAGAT | 7 |
| chr8 | 75159357 | CTATGCTTTTGAG-ATCTTACA-CACAAAATCTTTA | loxF8R | 4 | loxF8R | 3 | ATCTTACA/TGTAAGAT | 7 |
| --- | | | | | | | | 0 |
| chr14 | 84321491 | CAAACATTTTGTG-GCTAAGAT-CTCAAAAGCTGAG | loxF8R | 3 | loxF8R | 4 | ATCTTAGC/GCTAAGAT | 7 |
| chr16 | 74000208 | CTATGCTTTTGGG-ATCTTAGC-CATAAAATCTTTG | loxF8R | 4 | loxF8R | 3 | ATCTTAGC/GCTAAGAT | 7 |
| chr11 | 107291132 | CTGTGATTTTGAG-ATCTTAGC-CATAAAATCTTTG | loxF8R | 4 | loxF8R | 3 | ATCTTAGC/GCTAAGAT | 7 |
| chr5 | 20638708 | CAAAGATTTTATG-GCTAAGAT-TTCAAAATCATAG | loxF8R | 3 | loxF8R | 4 | ATCTTAGC/GCTAAGAT | 7 |
| --- | | | | | | | | 0 |
| chr2 | 16975150 | CTAAATCTTATGA-ATGAAGGA-TACACACTTTTAT | loxF8L | 4 | loxF8L | 3 | ATGAAGGA/TCCTTCAT | 7 |
| chr9 | 30396720 | ATATATCTGAGAA-ATGAAGGA-AAAACAAACTAAG | loxF8L | 3 | loxF8R | 4 | ATGAAGGA/TCCTTCAT | 7 |
| --- | | | | | | | | 0 |
| chr2 | 162874341 | ACAAAACTATGAA-ATGACTAC-AAGAAAATCTTAG | loxF8L | 4 | loxF8R | 3 | ATGACTAC/GTAGTCAT | 7 |
| chr14 | 20692223 | CTCAGTATGTGTG-ATGACTAC-CAAGAAATCATAG | loxF8R | 3 | loxF8R | 4 | ATGACTAC/GTAGTCAT | 7 |
| --- | | | | | | | | 0 |
| chr10 | 67421631 | ATAAATCTGAATA-ATGATTTC-TTCTTAGATTTCT | loxF8L | 3 | loxF8L | 4 | ATGATTTC/GAAATCAT | 7 |
| chrX | 74386584 | ATAAATCAGGAGA-ATGATTTC-TCAACAGATAGAG | loxF8L | 3 | loxF8L | 4 | ATGATTTC/GAAATCAT | 7 |
| --- | | | | | | | | 0 |
| chr18 | 66000249 | AGATATCTGTGTT-TTTTCCAT-TTCACAGTTATAT | loxF8L | 4 | loxF8L | 3 | ATGGAAAA/TTTTCCAT | 7 |
| chr3 | 166929677 | TTATATCTGTATA-ATGGAAAA-TTTACAGAGTTAT | loxF8L | 4 | loxF8L | 3 | ATGGAAAA/TTTTCCAT | 7 |
| chr9 | 122105573 | CTAAGTCTGTGTT-TTTTCCAT-TCCACCACTGTAT | loxF8R | 3 | loxF8L | 4 | ATGGAAAA/TTTTCCAT | 7 |
| --- | | | | | | | | 0 |
| chr4 | 26118836 | ATAACTATTTGCA-CTCTCCAT-CACACACTCTTGA | loxF8L | 4 | loxF8R | 3 | ATGGAGAG/CTCTCCAT | 7 |
| chr8 | 68033854 | CTGAGATGCTGTA-CTCTCCAT-TGAACAGATTTAA | loxF8R | 4 | loxF8L | 3 | ATGGAGAG/CTCTCCAT | 7 |
| --- | | | | | | | | 0 |
| chr3 | 163032575 | CTAATATTTTGCT-ATGGCACA-TGCAGAAATTTAT | loxF8R | 4 | loxF8L | 3 | ATGGCACA/TGTGCCAT | 7 |
| chr2 | 4189405 | ATAATTGTATGGC-ATGGCACA-CACATAATCTTTA | loxF8L | 4 | loxF8R | 3 | ATGGCACA/TGTGCCAT | 7 |
| --- | | | | | | | | 0 |
| chr7 | 93616283 | ATAAATCAGAGAA-ATGGCCAT-TTCAGAGATTCAC | loxF8L | 3 | loxF8L | 4 | ATGGCCAT/ATGGCCAT | 7 |
| --- | | | | | | | | 0 |
| chr11 | 25483909 | CAAGGATCGTGTG-ATGGGTGC-ACCAAAATCTCAG | loxF8R | 3 | loxF8R | 4 | ATGGGTGC/GCACCCAT | 7 |
| chr2 | 66360259 | CTGAGATTTTGTT-GCACCCAT-CACTCAAGCATTG | loxF8R | 3 | loxF8R | 4 | ATGGGTGC/GCACCCAT | 7 |
| chr13 | 47537578 | CTCTGCTTGGGTG-ATGGGTGC-ACCAAAATCTTAG | loxF8R | 4 | loxF8R | 3 | ATGGGTGC/GCACCCAT | 7 |
| chr7 | 72258017 | CTCTGCTTGGGTG-ATGGGTGC-AACAAAATCTCAG | loxF8R | 4 | loxF8R | 3 | ATGGGTGC/GCACCCAT | 7 |

| | | | | | | | | |
|---|---|---|---|---|---|---|---|---|
| --- | | | | | | | | 0 |
| chr5 | 156101164 | ATAAAATTCTGTT-GTTTACAT-CACCCAATCTATG | loxF8R | 4 | loxF8R | 3 | ATGTAAAC/GTTTACAT | 7 |
| chr8 | 49614400 | TTAAATCTGTAAA-ATGTAAAC-TCTAAACATCTAT | loxF8L | 3 | loxF8L | 4 | ATGTAAAC/GTTTACAT | 7 |
| --- | | | | | | | | 0 |
| chr8 | 100308694 | AAAAAAATGTGGA-ATGTAAAG-ATCACAATTTAAG | loxF8L | 3 | loxF8R | 4 | ATGTAAAG/CTTTACAT | 7 |
| chr15 | 64527060 | AGAAACCTCTGGA-ATGTAAAG-CACACAAGATTGA | loxF8L | 3 | loxF8R | 4 | ATGTAAAG/CTTTACAT | 7 |
| --- | | | | | | | | 0 |
| chr8 | 92779978 | AAAAATCTATGAA-ATTTACAT-TACATGGCTTTAT | loxF8L | 3 | loxF8L | 4 | ATGTAAAT/ATTTACAT | 7 |
| chr10 | 130878416 | CTAAGACTGAGTA-ATTTACAT-TCAACAGAAATGT | loxF8R | 3 | loxF8L | 4 | ATGTAAAT/ATTTACAT | 7 |
| --- | | | | | | | | 0 |
| chr13 | 35653078 | ATAACTCTTTGCA-GCAGACAT-TCAAAATATTTTT | loxF8L | 3 | loxF8L | 4 | ATGTCTGC/GCAGACAT | 7 |
| chr11 | 106935981 | ATAAAACTGAGGT-GCAGACAT-TAAAAAGATTTAC | loxF8L | 3 | loxF8L | 4 | ATGTCTGC/GCAGACAT | 7 |
| --- | | | | | | | | 0 |
| chr6 | 108429691 | TTGAATCTGAGTA-ATGTGATA-CCCCCAGATTTGT | loxF8L | 4 | loxF8L | 3 | ATGTGATA/TATCACAT | 7 |
| chr20 | 10467219 | CGTTGATTTTGTG-TATCACAT-TGAAAAGATTTAT | loxF8R | 4 | loxF8L | 3 | ATGTGATA/TATCACAT | 7 |
| --- | | | | | | | | 0 |
| chr3 | 133186604 | ATATCTCTGGGAA-ATGTGTCC-TCCATTGATTTGT | loxF8L | 4 | loxF8L | 3 | ATGTGTCC/GGACACAT | 7 |
| chr16 | 31814563 | TTAATATTGTGTT-GGACACAT-TCAAAAGAGTGAT | loxF8R | 3 | loxF8L | 4 | ATGTGTCC/GGACACAT | 7 |
| --- | | | | | | | | 0 |
| chr1 | 101406794 | ATAGATATTTGGA-ATTAAAAC-AAAACAATCTTTT | loxF8L | 3 | loxF8R | 4 | ATTAAAAC/GTTTTAAT | 7 |
| chr2 | 61045110 | ATAAATATGCTCA-ATTAAAAC-CACATACTCATAG | loxF8L | 4 | loxF8R | 3 | ATTAAAAC/GTTTTAAT | 7 |
| chr11 | 41194801 | ATGAATGTGTGTT-GTTTTAAT-TCACCACATTTAT | loxF8L | 4 | loxF8L | 3 | ATTAAAAC/GTTTTAAT | 7 |
| chr5 | 152725036 | ATTAATGTGTGTG-ATTAAAAC-TACACACATTAAT | loxF8L | 4 | loxF8L | 3 | ATTAAAAC/GTTTTAAT | 7 |
| --- | | | | | | | | 0 |
| chr11 | 65675054 | CTTAGATTAGGTA-ATTAACAA-ACAACAAATTTAT | loxF8R | 4 | loxF8L | 3 | ATTAACAA/TTGTTAAT | 7 |
| chr14 | 86895254 | TTAAATATGTGTA-ATTAACAA-CACAAATAATTAG | loxF8L | 3 | loxF8R | 4 | ATTAACAA/TTGTTAAT | 7 |
| --- | | | | | | | | 0 |
| chr4 | 69913968 | ATAAAACTACTGA-TTATTAAT-TACACAGTTTTTT | loxF8L | 4 | loxF8L | 3 | ATTAATAA/TTATTAAT | 7 |
| chr9 | 30688397 | CTTAGACTGGGTG-ATTAATAA-ACTACTGATTTTT | loxF8R | 3 | loxF8L | 4 | ATTAATAA/TTATTAAT | 7 |
| --- | | | | | | | | 0 |
| chr2 | 110232529 | ATATATCTGTATC-ATTAGAAA-TGCACAGACTTTT | loxF8L | 4 | loxF8L | 3 | ATTAGAAA/TTTCTAAT | 7 |
| chr2 | 109934584 | ATATATCTGTATC-ATTAGAAA-TGCACAGACTTTT | loxF8L | 4 | loxF8L | 3 | ATTAGAAA/TTTCTAAT | 7 |
| chr2 | 110436889 | AAAAGTCTGTGCA-TTTCTAAT-GATACAGATATAT | loxF8L | 3 | loxF8L | 4 | ATTAGAAA/TTTCTAAT | 7 |
| --- | | | | | | | | 0 |
| chr18 | 6565701 | TTAAATTTTTGGA-ATTAGAGA-TATACAGATTCTT | loxF8L | 3 | loxF8L | 4 | ATTAGAGA/TCTCTAAT | 7 |
| chr2 | 4857068 | ATCAATGTGTCTA-TCTCTAAT-TCCATTGGTTTAT | loxF8L | 4 | loxF8L | 3 | ATTAGAGA/TCTCTAAT | 7 |
| --- | | | | | | | | 0 |
| chr7 | 94351183 | ATAAAACACTGGA-ATTATAAT-TCCACATGTAGAT | loxF8L | 3 | loxF8L | 4 | ATTATAAT/ATTATAAT | 7 |
| chr7 | 156294499 | ATAATTCTCTTGA-ATTATAAT-TCAAAACTTTTAT | loxF8L | 3 | loxF8L | 4 | ATTATAAT/ATTATAAT | 7 |
| --- | | | | | | | | 0 |
| chr16 | 55570062 | AAAAATATGTGAA-ATTATACA-TCTACATTTTTAA | loxF8L | 3 | loxF8L | 4 | ATTATACA/TGTATAAT | 7 |
| chr2 | 62060367 | TTAAATTTCAGGA-ATTATACA-ACCACAGAATTCT | loxF8L | 4 | loxF8L | 3 | ATTATACA/TGTATAAT | 7 |
| --- | | | | | | | | 0 |
| chr3 | 70367026 | AAAATTGTATGGA-TATATAAT-TGTACATATTTAT | loxF8L | 4 | loxF8L | 3 | ATTATATA/TATATAAT | 7 |
| chr7 | 114790600 | TTATTACTGTGTG-ATTATATA-TCCAAAAATTTCT | loxF8R | 4 | loxF8L | 3 | ATTATATA/TATATAAT | 7 |
| --- | | | | | | | | 0 |
| chr9 | 16361500 | ATATAATTTTGTG-TTATGAAT-TCCACAAATATTT | loxF8R | 4 | loxF8L | 3 | ATTCATAA/TTATGAAT | 7 |
| chr10 | 82085387 | CTGAGACTGAGTA-ATTCATAA-AGAACAGATTTAT | loxF8R | 4 | loxF8L | 3 | ATTCATAA/TTATGAAT | 7 |
| chrX | 146589305 | ATAAATTTGTTGC-TTATGAAT-TACCCAGTCTTGG | loxF8L | 3 | loxF8R | 4 | ATTCATAA/TTATGAAT | 7 |
| --- | | | | | | | | 0 |
| chr2 | 81878448 | ATAACTCTGTCCC-ATTCTAAG-CACAGTCTCTTAG | loxF8L | 4 | loxF8R | 3 | ATTCTAAG/CTTAGAAT | 7 |
| chr2 | 149627439 | TTAAGATTAAGTT-ATTCTAAG-CACACTATCTCTG | loxF8R | 4 | loxF8R | 3 | ATTCTAAG/CTTAGAAT | 7 |
| --- | | | | | | | | 0 |
| chr2 | 126224006 | AAAAAACTTTAGA-ATTCTATG-TCTACAGAATTAG | loxF8L | 4 | loxF8L | 3 | ATTCTATG/CATAGAAT | 7 |
| chr5 | 25638496 | ATTAACCTATGGA-ATTCTATG-TTGACAGTTATAT | loxF8L | 3 | loxF8L | 4 | ATTCTATG/CATAGAAT | 7 |
| chr4 | 173456194 | CTAGGACTCTGTG-ATTCTATG-TACACAGAATTCG | loxF8R | 3 | loxF8L | 4 | ATTCTATG/CATAGAAT | 7 |
| --- | | | | | | | | 0 |
| chr8 | 5616097 | GTAATTCTGTGAA-TTTTCAAT-TTAATACATTTAT | loxF8L | 3 | loxF8L | 4 | ATTGAAAA/TTTTCAAT | 7 |
| chr18 | 65550702 | TTAAATCTTCAGA-TTTTCAAT-TCCACACATGTAA | loxF8L | 4 | loxF8L | 3 | ATTGAAAA/TTTTCAAT | 7 |
| --- | | | | | | | | 0 |
| chrX | 141040816 | CTTAAAATGTTTG-ATTGAAGA-AATACAATCATAG | loxF8R | 4 | loxF8R | 3 | ATTGAAGA/TCTTCAAT | 7 |
| chrX | 141546541 | CTATGATTGTATT-TCTTCAAT-CAAACATTTTAAG | loxF8R | 3 | loxF8R | 4 | ATTGAAGA/TCTTCAAT | 7 |
| --- | | | | | | | | 0 |
| chr4 | 109416528 | ATAAATCTGGGTT-ATTGAAGC-CACCCAGTCTGTG | loxF8L | 3 | loxF8R | 4 | ATTGAAGC/GCTTCAAT | 7 |
| chr7 | 147641836 | ATAAATTTCTGTC-ATTGAAGC-CACTCAATCTATG | loxF8L | 4 | loxF8R | 3 | ATTGAAGC/GCTTCAAT | 7 |
| chr7 | 138061893 | CTCAGATTATTTG-ATTGAAGC-CATTCAATGTAAG | loxF8R | 3 | loxF8R | 4 | ATTGAAGC/GCTTCAAT | 7 |
| --- | | | | | | | | 0 |
| chr1 | 78869805 | ATAGAACTCTGGG-CTCTCAAT-CACACAATCATTT | loxF8L | 4 | loxF8R | 3 | ATTGAGAG/CTCTCAAT | 7 |
| chr15 | 45554606 | ATTTATTTCTGGA-CTCTCAAT-TCTACTGATGTAT | loxF8L | 4 | loxF8L | 3 | ATTGAGAG/CTCTCAAT | 7 |
| --- | | | | | | | | 0 |
| chr17 | 8675226 | AAAAGTCTGAGGA-CACTCAAT-TCCCCAAATTCCT | loxF8L | 3 | loxF8L | 4 | ATTGAGTG/CACTCAAT | 7 |
| chr11 | 120534336 | ATATCTATGTGGA-ATTGAGTG-CACACACCCAGAG | loxF8L | 3 | loxF8R | 4 | ATTGAGTG/CACTCAAT | 7 |
| --- | | | | | | | | 0 |
| chr12 | 7282165 | TTAAATCTTTCAA-CTGCCAAT-CAGAAAATCTTTG | loxF8L | 4 | loxF8R | 3 | ATTGGCAG/CTGCCAAT | 7 |
| chr7 | 81975844 | CTCAGTTTCTGTC-ATTGGCAG-CAAATACTCTTAG | loxF8R | 4 | loxF8R | 3 | ATTGGCAG/CTGCCAAT | 7 |
| --- | | | | | | | | 0 |
| chr8 | 128284823 | ATATTTCTCTGAA-CATACAAT-TCAACAGAATGAT | loxF8L | 4 | loxF8L | 3 | ATTGTATG/CATACAAT | 7 |
| chr19 | 47838037 | GTATGTTTGTGTG-ATTGTATG-CACATATGATTAG | loxF8R | 3 | loxF8R | 4 | ATTGTATG/CATACAAT | 7 |
| --- | | | | | | | | 0 |
| chr1 | 36866782 | CTATGATGATGTG-TTGACAAT-TTCACGGATTACT | loxF8R | 3 | loxF8L | 4 | ATTGTCAA/TTGACAAT | 7 |
| chr6 | 119975677 | ATAAATATGTACT-ATTGTCAA-CACACAGCCTTAA | loxF8L | 4 | loxF8R | 3 | ATTGTCAA/TTGACAAT | 7 |
| --- | | | | | | | | 0 |
| chr6 | 15751888 | ACAAATCAGTGCA-GTAACAAT-CCCCAAATCTCAG | loxF8L | 3 | loxF8R | 4 | ATTGTTAC/GTAACAAT | 7 |
| chr10 | 67936495 | ATAAAACTTTAGA-ATTGTTAC-TATAAAATATTAG | loxF8L | 3 | loxF8R | 4 | ATTGTTAC/GTAACAAT | 7 |
| --- | | | | | | | | 0 |
| chr5 | 22517356 | AAAAATCTCTAGA-ATTTAAAT-TCATCACTTTTAT | loxF8L | 3 | loxF8L | 4 | ATTTAAAT/ATTTAAAT | 7 |
| chr3 | 182559241 | ATGAATCTCTGTT-ATTTAAAT-CACGCAATCTATG | loxF8L | 4 | loxF8R | 3 | ATTTAAAT/ATTTAAAT | 7 |

| | | | | | | | | |
|---|---|---|---|---|---|---|---|---|
| chr15 | 49285805 | TTAAATCTGAGCA-ATTTAAAT-TCCCCTGCTTAAT | loxF8L | 3 | loxF8L | 4 | ATTTAAAT/ATTTAAAT | 7 |
| chr8 | 92500375 | CTGTGACTGTGTG-ATTTAAAT-CCCAAAGATTTTA | loxF8R | 3 | loxF8L | 4 | ATTTAAAT/ATTTAAAT | 7 |
| --- | | | | | | | | 0 |
| chr2 | 180325145 | AAAAATCTGTAAA-TCTTAAAT-TCAAATGATTTCT | loxF8L | 3 | loxF8L | 4 | ATTTAAGA/TCTTAAAT | 7 |
| chr2 | 27631935 | AAAAATCTGTGAC-TCTTAAAT-TCCTTAGCTTGAT | loxF8L | 3 | loxF8L | 4 | ATTTAAGA/TCTTAAAT | 7 |
| --- | | | | | | | | 0 |
| chr1 | 110073419 | CTAAGATTGGTTT-ATTTAATC-TTCACAGCTCTGT | loxF8R | 3 | loxF8L | 4 | ATTTAATC/GATTAAAT | 7 |
| chr3 | 131587093 | CTAGGATTGTCTC-ATTTAATC-TTCACAAAATTCT | loxF8R | 3 | loxF8L | 4 | ATTTAATC/GATTAAAT | 7 |
| chr9 | 67586856 | TTAAGAGTGTGTT-GATTAAAT-TCTGCATATTCAT | loxF8R | 3 | loxF8L | 4 | ATTTAATC/GATTAAAT | 7 |
| chr9 | 62001822 | ATGAATATGCAGA-ATTTAATC-AACACACTCTTAA | loxF8L | 4 | loxF8R | 3 | ATTTAATC/GATTAAAT | 7 |
| --- | | | | | | | | 0 |
| chr4 | 99275028 | ATAAATTTCTTCA-TTGTAAAT-CACCCAGTCTCAG | loxF8L | 4 | loxF8R | 3 | ATTTACAA/TTGTAAAT | 7 |
| chr7 | 78145711 | ATATATCTGTTCA-TTGTAAAT-CACCCAGTCTGTG | loxF8L | 3 | loxF8R | 4 | ATTTACAA/TTGTAAAT | 7 |
| --- | | | | | | | | 0 |
| chr2 | 221253375 | CTGAGACTGGGTG-ATTTAGAA-TAAACAGATATTT | loxF8R | 3 | loxF8L | 4 | ATTTAGAA/TTCTAAAT | 7 |
| chr6 | 98739220 | ACAAATATTTGGT-ATTTAGAA-CAGACAATGCTAG | loxF8L | 4 | loxF8R | 3 | ATTTAGAA/TTCTAAAT | 7 |
| --- | | | | | | | | 0 |
| chr4 | 148624933 | CTGAGACTGTGTA-ATTTATTA-TAAAAAAATTTAT | loxF8R | 3 | loxF8L | 4 | ATTTATTA/TAATAAAT | 7 |
| chr18 | 66563925 | ATTAGACTGTGTT-TAATAAAT-CAAACAATATTAT | loxF8R | 4 | loxF8R | 3 | ATTTATTA/TAATAAAT | 7 |
| chr15 | 97658943 | CTAATTCTGGTGA-TAATAAAT-CAAACATTTTTAG | loxF8L | 4 | loxF8R | 3 | ATTTATTA/TAATAAAT | 7 |
| --- | | | | | | | | 0 |
| chr15 | 55269140 | ATAAATTTCTGTT-TTTGAAAT-CACACAGTTTTAA | loxF8L | 4 | loxF8R | 3 | ATTTCAAA/TTTGAAAT | 7 |
| chr13 | 69879386 | CTAACATATTGTG-TTTGAAAT-TCCACATATTCTA | loxF8R | 3 | loxF8L | 4 | ATTTCAAA/TTTGAAAT | 7 |
| --- | | | | | | | | 0 |
| chr22 | 33823053 | ATAAATCTTTGTT-ATTTCAAC-CACACAGAGATAT | loxF8L | 3 | loxF8L | 4 | ATTTCAAC/GTTGAAAT | 7 |
| chrX | 127496190 | ATAATTCTTGGGA-ATTTCAAC-ACCCCAATATTAG | loxF8L | 3 | loxF8R | 4 | ATTTCAAC/GTTGAAAT | 7 |
| --- | | | | | | | | 0 |
| chr8 | 27790018 | CTGACATTGTCTG-CTTCAAAT-TTGACAGCTTTCT | loxF8R | 3 | loxF8L | 4 | ATTTGAAG/CTTCAAAT | 7 |
| chr2 | 220392940 | ATAAGTTTTTGTT-ATTTGAAG-TCACCAAATTTAT | loxF8R | 4 | loxF8L | 3 | ATTTGAAG/CTTCAAAT | 7 |
| --- | | | | | | | | 0 |
| chr4 | 61253281 | ATAAAGCTGACTA-TTCCAAAT-TCCAAAGATTCAG | loxF8L | 4 | loxF8L | 3 | ATTTGGAA/TTCCAAAT | 7 |
| chrX | 120160706 | ATAAATATATGTA-TTCCAAAT-TACAAATATATAT | loxF8L | 3 | loxF8L | 4 | ATTTGGAA/TTCCAAAT | 7 |
| chrX | 120075562 | ATATATATTGTA-ATTTGGAA-TACATATATTTAT | loxF8L | 4 | loxF8L | 3 | ATTTGGAA/TTCCAAAT | 7 |
| --- | | | | | | | | 0 |
| chrX | 113879444 | CTTAGACTGTGTA-ATTTGTAA-ACAACAGAATTTT | loxF8R | 3 | loxF8L | 4 | ATTTGTAA/TTACAAAT | 7 |
| chr8 | 19772243 | ATAAGTTTGTGTT-ATTTGTAA-GCCACATTCTTAT | loxF8R | 3 | loxF8L | 4 | ATTTGTAA/TTACAAAT | 7 |
| chr12 | 47690105 | TTCAGATTTTGTG-ATTTGTAA-TACACAAACCTAT | loxF8R | 3 | loxF8L | 4 | ATTTGTAA/TTACAAAT | 7 |
| --- | | | | | | | | 0 |
| chr4 | 38507487 | CTTAGACTGGGTG-ATTTGTGA-ACAACAGAATTTT | loxF8R | 3 | loxF8L | 4 | ATTTGTGA/TCACAAAT | 7 |
| chr6 | 21650977 | CTAAAGATATGGA-ATTTGTGA-TTGGCAGATTTAT | loxF8L | 4 | loxF8L | 3 | ATTTGTGA/TCACAAAT | 7 |
| chr3 | 164712102 | ATAAATCTTTCCT-TCACAAAT-TACCCAATCTCAG | loxF8L | 4 | loxF8R | 3 | ATTTGTGA/TCACAAAT | 7 |
| chr2 | 227111148 | CTCAAATTGTGGC-ATTTGTGA-TTCACACATTTAC | loxF8R | 4 | loxF8L | 3 | ATTTGTGA/TCACAAAT | 7 |
| --- | | | | | | | | 0 |
| chrY | 15642459 | ACAAATTTAGGGA-CTTAAAAT-GACACATATTTAT | loxF8L | 4 | loxF8L | 3 | ATTTTAAG/CTTAAAAT | 7 |
| chr6 | 154719526 | ATAAATGTGTGTT-ATTTTAAG-TCACCAAGTTTAT | loxF8L | 3 | loxF8L | 4 | ATTTTAAG/CTTAAAAT | 7 |
| --- | | | | | | | | 0 |
| chr1 | 97264492 | ATACATTTCTGAA-TTGAAAAT-TCCAGTGATTTGT | loxF8L | 4 | loxF8L | 3 | ATTTTCAA/TTGAAAAT | 7 |
| chr7 | 20722131 | CTATGATTCTGTC-ATTTTCAA-ACATCAGCTTTAT | loxF8R | 3 | loxF8L | 4 | ATTTTCAA/TTGAAAAT | 7 |
| chr3 | 24189193 | ATATGATTGTGAG-ATTTTCAA-AACACAATTTAAA | loxF8R | 3 | loxF8R | 4 | ATTTTCAA/TTGAAAAT | 7 |
| --- | | | | | | | | 0 |
| chr3 | 46766193 | TGAAGCTTGTATG-TAGAAAAT-TCTACAGAATTTT | loxF8R | 4 | loxF8L | 3 | ATTTTCTA/TAGAAAAT | 7 |
| chr4 | 161095285 | ATAACTATGTGAA-TAGAAAAT-TCAAAATATATAT | loxF8L | 3 | loxF8L | 4 | ATTTTCTA/TAGAAAAT | 7 |
| chr13 | 57495304 | CCAAGACTGTGGG-ATTTTCTA-GATACAGAATTAT | loxF8R | 3 | loxF8L | 4 | ATTTTCTA/TAGAAAAT | 7 |
| --- | | | | | | | | 0 |
| chr2 | 84048334 | ACAAATTTGAGGA-GTAAAAAT-CATACAATGTATG | loxF8L | 3 | loxF8R | 4 | ATTTTTAC/GTAAAAAT | 7 |
| chr12 | 123980359 | CTAAAAGGATGTG-GTAAAAAT-GACACACTCTTAT | loxF8R | 4 | loxF8R | 3 | ATTTTTAC/GTAAAAAT | 7 |
| chr2 | 36535686 | TTGTATCTGTGGA-GTAAAAAT-TTTACAATATTAG | loxF8L | 3 | loxF8R | 4 | ATTTTTAC/GTAAAAAT | 7 |
| --- | | | | | | | | 0 |
| chr4 | 120493427 | ATAACTTTGTGAA-ATTTTTTC-CCCAAATATTTTT | loxF8L | 3 | loxF8L | 4 | ATTTTTTC/GAAAAAAT | 7 |
| chr5 | 144693013 | GTTAGAGTGTGTG-GAAAAAAT-CTTATAATCTTAT | loxF8R | 3 | loxF8R | 4 | ATTTTTTC/GAAAAAAT | 7 |
| --- | | | | | | | | 0 |
| chr15 | 39597342 | CTAGGAATGTGCT-TTTTTTTG-TACACATTTTTAT | loxF8R | 4 | loxF8L | 3 | CAAAAAAA/TTTTTTTG | 7 |
| chr17 | 56387906 | CTAAGAGTCTGTC-TTTTTTTG-TATACACATTTAA | loxF8R | 3 | loxF8L | 4 | CAAAAAAA/TTTTTTTG | 7 |
| chr4 | 58175040 | CTCACATTGTTTG-CAAAAAAA-CCCTCAGCTTTAC | loxF8R | 3 | loxF8L | 4 | CAAAAAAA/TTTTTTTG | 7 |
| --- | | | | | | | | 0 |
| chr11 | 29618242 | TTAATATTTTGTA-GTTTTTTG-TTTACACATTTAT | loxF8R | 4 | loxF8L | 3 | CAAAAAAC/GTTTTTTG | 7 |
| chr2 | 99326573 | TCAAATCTGTGCA-CAAAAAAC-CACAAAGGCTTGG | loxF8L | 3 | loxF8R | 4 | CAAAAAAC/GTTTTTTG | 7 |
| --- | | | | | | | | 0 |
| chr3 | 144532740 | TGAGAGCTGTGGA-CAAAACCA-TCTACTAATTTAT | loxF8L | 4 | loxF8L | 3 | CAAAACCA/TGGTTTTG | 7 |
| chr1 | 43727336 | ATAAGATGATGAG-CAAAACCA-GACACAATTTCAG | loxF8R | 4 | loxF8R | 3 | CAAAACCA/TGGTTTTG | 7 |
| --- | | | | | | | | 0 |
| chr12 | 93399687 | ATCAGTCTGTGGT-TTATTTTG-TCATCAGATTACT | loxF8L | 3 | loxF8L | 4 | CAAAATAA/TTATTTTG | 7 |
| chr1 | 146299023 | ATCAGTCTGTGGT-TTATTTTG-TCATCAGATTACT | loxF8L | 3 | loxF8L | 4 | CAAAATAA/TTATTTTG | 7 |
| chr1 | 148749806 | ATCAGTCTGTGGT-TTATTTTG-TCATCAGATTACT | loxF8L | 3 | loxF8L | 4 | CAAAATAA/TTATTTTG | 7 |
| --- | | | | | | | | 0 |
| chr8 | 108740867 | CTAAAACTATGTG-CAAAATAG-TTCACAGTGTCAT | loxF8R | 3 | loxF8L | 4 | CAAAATAG/CTATTTTG | 7 |
| chr2 | 76382233 | ATTAATCTGTTGC-CTATTTTG-TCTACAGATGAAG | loxF8L | 3 | loxF8L | 4 | CAAAATAG/CTATTTTG | 7 |
| --- | | | | | | | | 0 |
| chr2 | 82008040 | ATAAAAATGTGGT-TATGTTTG-CACATATTTATAG | loxF8L | 3 | loxF8R | 4 | CAAACATA/TATGTTTG | 7 |
| chr1 | 35255829 | CCAAGAGAATGTG-CAAACATA-TAGAGAGATTTAT | loxF8R | 4 | loxF8L | 3 | CAAACATA/TATGTTTG | 7 |
| --- | | | | | | | | 0 |
| chr15 | 101274318 | CTTAATTTATGGA-CAAAGAAA-CAAACAATCTTGT | loxF8L | 4 | loxF8R | 3 | CAAAGAAA/TTTCTTTG | 7 |
| chr7 | 9482867 | CTAAGCCTGTATG-TTTCTTTG-TGGAAAGTTTTAT | loxF8R | 3 | loxF8L | 4 | CAAAGAAA/TTTCTTTG | 7 |
| --- | | | | | | | | 0 |
| chr7 | 124241782 | AAAACTCTGGGTA-CAAAGTTC-TGGACATATTTAT | loxF8L | 4 | loxF8L | 3 | CAAAGTTC/GAACTTTG | 7 |
| chr1 | 61144363 | ATAGCTCTCTGGA-GAACTTTG-TCTCCTGATTCAT | loxF8L | 3 | loxF8L | 4 | CAAAGTTC/GAACTTTG | 7 |
| chr8 | 39411400 | CAAAGTTGGTGTG-GAACTTTG-CACTGAAACTCAG | loxF8R | 3 | loxF8R | 4 | CAAAGTTC/GAACTTTG | 7 |

| | | | | | | | | |
|---|---|---|---|---|---|---|---|---|
| --- | | | | | | | | 0 |
| chr2 | 89299266 | ATAAAACTGGTAA-TGGATTTG-CACCCAATCATAT | loxF8L | 4 | loxF8R | 3 | CAAATCCA/TGGATTTG | 7 |
| chr2 | 89883177 | ATATGATTGGGTG-CAAATCCA-TTACCAGTTTTAT | loxF8R | 3 | loxF8L | 4 | CAAATCCA/TGGATTTG | 7 |
| --- | | | | | | | | 0 |
| chr12 | 8095173 | ATAAAGCTGCCGT-GGCATTTG-TATACAGATTTTT | loxF8L | 4 | loxF8L | 3 | CAAATGCC/GGCATTTG | 7 |
| chr2 | 188615574 | CTAAATGTCTGGT-GGCATTTG-TTCAGAAATTTAT | loxF8L | 4 | loxF8L | 3 | CAAATGCC/GGCATTTG | 7 |
| --- | | | | | | | | 0 |
| chr8 | 78980019 | AGATATGTGTGGT-CAAATGTG-CACACAGTGTTAT | loxF8L | 4 | loxF8R | 3 | CAAATGTG/CACATTTG | 7 |
| chr3 | 68369780 | CTAGAATGGTGTG-CACATTTG-CACATTCTATTAG | loxF8R | 3 | loxF8R | 4 | CAAATGTG/CACATTTG | 7 |
| --- | | | | | | | | 0 |
| chr5 | 159958055 | AAAAATGTCTAGA-CAAATTCA-TCAATACATTTAT | loxF8L | 4 | loxF8L | 3 | CAAATTCA/TGAATTTG | 7 |
| chr1 | 33661154 | GCAAATCTGTGGG-TGAATTTG-TCAGCATAATTAT | loxF8L | 3 | loxF8L | 4 | CAAATTCA/TGAATTTG | 7 |
| --- | | | | | | | | 0 |
| chr4 | 36107783 | AAACATCTGTAGG-CAACATTA-TACACAATCATAT | loxF8L | 4 | loxF8R | 3 | CAACATTA/TAATGTTG | 7 |
| chr12 | 44787396 | ATTTAGCTGTAGA-CAACATTA-TCCTAAAATTTAT | loxF8L | 4 | loxF8L | 3 | CAACATTA/TAATGTTG | 7 |
| --- | | | | | | | | 0 |
| chr6 | 54517191 | TCAAGATTGTTTA-CAACTCAA-ACCACAGAGTTTT | loxF8R | 4 | loxF8L | 3 | CAACTCAA/TTGAGTTG | 7 |
| chr14 | 96722928 | CTAAGATTGCAAG-TTGAGTTG-TCCATTTATTCAT | loxF8R | 3 | loxF8L | 4 | CAACTCAA/TTGAGTTG | 7 |
| --- | | | | | | | | 0 |
| chr4 | 86345297 | TTCAATCTGTGTG-CAACTTAA-TCCACATAATTTT | loxF8L | 4 | loxF8L | 3 | CAACTTAA/TTAAGTTG | 7 |
| chr14 | 32704245 | ATAAAATTTTGGA-TTAAGTTG-CAAAAAATTTCAG | loxF8L | 3 | loxF8R | 4 | CAACTTAA/TTAAGTTG | 7 |
| --- | | | | | | | | 0 |
| chr1 | 108635491 | AAAAATCTGTGAT-CAAGAGGC-TCAATAGATTTGG | loxF8L | 3 | loxF8L | 4 | CAAGAGGC/GCCTCTTG | 7 |
| chr15 | 93120631 | ATAAAAATGTGGG-GCCTCTTG-TACAAATATGTAT | loxF8L | 3 | loxF8L | 4 | CAAGAGGC/GCCTCTTG | 7 |
| --- | | | | | | | | 0 |
| chr3 | 164616641 | CTAAGATATTGAG-TTTTATTG-TTAATAGTTTTAT | loxF8R | 3 | loxF8L | 4 | CAATAAAA/TTTTATTG | 7 |
| chr11 | 79957446 | ATAGTTCTATGGT-TTTTATTG-TGCAAAAATTTAT | loxF8L | 4 | loxF8L | 3 | CAATAAAA/TTTTATTG | 7 |
| --- | | | | | | | | 0 |
| chr4 | 132963789 | ATAAGAATCTGTG-CAATAAAG-AAGACAATTTTAT | loxF8R | 3 | loxF8R | 4 | CAATAAAG/CTTTATTG | 7 |
| chr15 | 52026020 | ATACATCTGTGTT-CAATAAAG-TAGATAATCTAAG | loxF8L | 3 | loxF8R | 4 | CAATAAAG/CTTTATTG | 7 |
| --- | | | | | | | | 0 |
| chr14 | 52056031 | CTGATTTTGTGTG-CAATATGC-TATACATATTTGT | loxF8R | 3 | loxF8L | 4 | CAATATGC/GCATATTG | 7 |
| chr4 | 92685891 | GTATGATTTTGTA-CAATATGC-CACAAAATTTTAA | loxF8R | 4 | loxF8R | 3 | CAATATGC/GCATATTG | 7 |
| --- | | | | | | | | 0 |
| chr5 | 12563523 | ACAAGACTGTATG-GAATATTG-CAGACACTCTTAC | loxF8R | 4 | loxF8R | 3 | CAATATTC/GAATATTG | 7 |
| chr13 | 37748701 | ATATATCTTTGTA-GAATATTG-CAGAAAATTTTAT | loxF8L | 3 | loxF8R | 4 | CAATATTC/GAATATTG | 7 |
| --- | | | | | | | | 0 |
| chr7 | 72741434 | AAAAAAGTGTGAA-GAAAATTG-AACAGAGTCTTAG | loxF8L | 4 | loxF8R | 3 | CAATTTTC/GAAAATTG | 7 |
| chr1 | 163830437 | ATCAATCTATGGT-GAAAATTG-TGCACAAACTAAG | loxF8L | 3 | loxF8R | 4 | CAATTTTC/GAAAATTG | 7 |
| --- | | | | | | | | 0 |
| chr2 | 55398126 | ACAAAACTGTGCT-GCTTTGTG-TCTACAAATCTAT | loxF8L | 4 | loxF8L | 3 | CACAAAGC/GCTTTGTG | 7 |
| chr2 | 7332244 | ATTAAGGTGGGGA-CACAAAGC-CAAACCATATTAG | loxF8L | 4 | loxF8R | 3 | CACAAAGC/GCTTTGTG | 7 |
| --- | | | | | | | | 0 |
| chr18 | 69923965 | AAAAATCAATGGA-GTATTGTG-TCCTCAGACTTCA | loxF8L | 3 | loxF8L | 4 | CACAATAC/GTATTGTG | 7 |
| chr3 | 108521896 | GTAAAAGGGTGTG-GTATTGTG-TCCAGACATTTGT | loxF8R | 4 | loxF8L | 3 | CACAATAC/GTATTGTG | 7 |
| --- | | | | | | | | 0 |
| chr1 | 102530207 | ATATATATGTATA-GATGTGTG-TGCACACATATAT | loxF8L | 4 | loxF8L | 3 | CACACATC/GATGTGTG | 7 |
| chr3 | 28146546 | CCAAATGTGTGCA-CACACATC-CACACTCACTTAG | loxF8L | 4 | loxF8R | 3 | CACACATC/GATGTGTG | 7 |
| --- | | | | | | | | 0 |
| chr15 | 84648378 | CTAAATCTGTGTG-TATCTGTG-TGCACACACTTGT | loxF8L | 3 | loxF8L | 4 | CACAGATA/TATCTGTG | 7 |
| chr2 | 208637419 | ATATATATGTGTA-TATCTGTG-TGCATACATATAT | loxF8L | 3 | loxF8L | 4 | CACAGATA/TATCTGTG | 7 |
| chr2 | 208637391 | ATATATATGTGTA-TATCTGTG-TGCATACATATAT | loxF8L | 3 | loxF8L | 4 | CACAGATA/TATCTGTG | 7 |
| chr2 | 208637335 | ATATATATGTGTA-TATCTGTG-TGCATACATATAT | loxF8L | 3 | loxF8L | 4 | CACAGATA/TATCTGTG | 7 |
| chr2 | 208637363 | ATATATATGTGTA-TATCTGTG-TGCATACATATAT | loxF8L | 3 | loxF8L | 4 | CACAGATA/TATCTGTG | 7 |
| chr2 | 208637447 | ATATATATGTGTA-TATCTGTG-TGCATACATATAT | loxF8L | 3 | loxF8L | 4 | CACAGATA/TATCTGTG | 7 |
| --- | | | | | | | | 0 |
| chr21 | 32407641 | AGACATCAGTTGA-CTGCTGTG-CACATATTCTTAA | loxF8L | 4 | loxF8R | 3 | CACAGCAG/CTGCTGTG | 7 |
| chr2 | 240360975 | ATAGATGTTTGAA-CACAGCAG-ATCACAGAATTAT | loxF8L | 4 | loxF8L | 3 | CACAGCAG/CTGCTGTG | 7 |
| --- | | | | | | | | 0 |
| chr12 | 196957 | CTGTGTGTGTGTG-TACCTGTG-CACACACTTTAAG | loxF8R | 4 | loxF8R | 3 | CACAGGTA/TACCTGTG | 7 |
| chr11 | 96689283 | TAATGATTGTGTG-TACCTGTG-TTCACATGTTTAA | loxF8R | 3 | loxF8L | 4 | CACAGGTA/TACCTGTG | 7 |
| --- | | | | | | | | 0 |
| chrX | 118945357 | ATATGTCTGTGTA-CACAGTCA-GGCACATATTTTT | loxF8L | 3 | loxF8L | 4 | CACAGTCA/TGACTGTG | 7 |
| chr13 | 55743685 | ATATATATATGGA-CACAGTCA-CACACATATTTGT | loxF8L | 3 | loxF8L | 4 | CACAGTCA/TGACTGTG | 7 |
| --- | | | | | | | | 0 |
| chr6 | 166558031 | ATATAGGTGTGTA-TGTATGTG-TCTATAGATTTGT | loxF8L | 4 | loxF8L | 3 | CACATACA/TGTATGTG | 7 |
| chr6 | 65605797 | ATAAATATGTGTG-TGTATGTG-TGTATATATTTAT | loxF8L | 3 | loxF8L | 4 | CACATACA/TGTATGTG | 7 |
| chr3 | 70306952 | ATAAATATGTATA-TGTATGTG-TGTATATATTTAT | loxF8L | 3 | loxF8L | 4 | CACATACA/TGTATGTG | 7 |
| chr10 | 65013200 | ATTTATGTGTGTA-CACATACA-TACATATATTTAT | loxF8L | 4 | loxF8L | 3 | CACATACA/TGTATGTG | 7 |
| chr2 | 152934697 | ATAAATATATGTG-TGTATGTG-TGCACACATATAT | loxF8L | 4 | loxF8L | 3 | CACATACA/TGTATGTG | 7 |
| chr3 | 63725657 | ATAAGCCTGTGCA-CACATACA-TACACTCATATAT | loxF8L | 3 | loxF8L | 4 | CACATACA/TGTATGTG | 7 |
| chr11 | 16948564 | ATACATGTGTTCA-CACATACA-TGCACACATATAT | loxF8L | 4 | loxF8L | 3 | CACATACA/TGTATGTG | 7 |
| chr21 | 40962393 | CTATGTGTGTGTG-TGTATGTG-TGCATATATATAT | loxF8R | 3 | loxF8L | 4 | CACATACA/TGTATGTG | 7 |
| chr4 | 123846948 | ATATATATATGTA-CACATACA-TACACATATATAT | loxF8L | 4 | loxF8L | 3 | CACATACA/TGTATGTG | 7 |
| chr18 | 56401757 | ATATATATGTACA-CACATACA-CACACAATGTTTA | loxF8L | 4 | loxF8R | 3 | CACATACA/TGTATGTG | 7 |
| chrY | 20004118 | ATATATATGTATA-CACATACA-TACACATATATAT | loxF8L | 4 | loxF8L | 3 | CACATACA/TGTATGTG | 7 |
| chr13 | 76929104 | CTGTGTGTGTGTG-TGTATGTG-TACACACATATAT | loxF8R | 4 | loxF8L | 3 | CACATACA/TGTATGTG | 7 |
| chr1 | 87134524 | ATATATGTGTGTA-TGTATGTG-TCTATAAATATAT | loxF8L | 3 | loxF8L | 4 | CACATACA/TGTATGTG | 7 |
| chr7 | 14204926 | CTTTCATTGTGTG-TGTATGTG-TCCACAACTGTAG | loxF8R | 3 | loxF8L | 4 | CACATACA/TGTATGTG | 7 |
| chr18 | 56853069 | TGAAGATTTTGTG-TGTATGTG-TGTACAGCTTTAA | loxF8R | 3 | loxF8L | 4 | CACATACA/TGTATGTG | 7 |
| --- | | | | | | | | 0 |
| chr11 | 11423165 | ATACATGTGTGCA-CACATGCA-CACACACACTTGT | loxF8L | 3 | loxF8R | 4 | CACATGCA/TGCATGTG | 7 |
| chr14 | 69215440 | AGGAGTTTGTGTG-CACATGCA-CGCACAATTTTAA | loxF8R | 4 | loxF8R | 3 | CACATGCA/TGCATGTG | 7 |
| chr7 | 109928520 | TTAAATTTCTGGA-CACATGCA-CACACAAAATAAT | loxF8L | 3 | loxF8R | 4 | CACATGCA/TGCATGTG | 7 |
| --- | | | | | | | | 0 |
| chr3 | 53379651 | ATACATGTATGCA-CACATGTA-TACACACATGTAT | loxF8L | 4 | loxF8L | 3 | CACATGTA/TACATGTG | 7 |
| chr3 | 144512485 | ATATATGTGTGTA-CACATGTA-TACACATATTTGC | loxF8L | 3 | loxF8L | 4 | CACATGTA/TACATGTG | 7 |

| | | | | | | | | |
|---|---|---|---|---|---|---|---|---|
| chrX | 126586488 | CACAGTTTGTGTG-TACATGTG-TATACAATTTTAA | loxF8R | 3 | loxF8R | 4 | CACATGTA/TACATGTG | 7 |
| --- | | | | | | | | 0 |
| chr12 | 77974035 | GAAAGTTTGTGTG-TGCACGTG-CACACACTTTTGT | loxF8R | 3 | loxF8R | 4 | CACGTGCA/TGCACGTG | 7 |
| chr6 | 26830647 | ATGCATGTGTGTA-CACGTGCA-TGCACACATGTAT | loxF8L | 4 | loxF8L | 3 | CACGTGCA/TGCACGTG | 7 |
| chr6 | 57872804 | ATGCATGTGTGTA-CACGTGCA-TGCACACATGTAT | loxF8L | 4 | loxF8L | 3 | CACGTGCA/TGCACGTG | 7 |
| --- | | | | | | | | 0 |
| chr1 | 43814222 | CTATGTTTATGTC-CACTTTGG-CCCCCAGATTTCT | loxF8R | 4 | loxF8L | 3 | CACTTTGG/CCAAAGTG | 7 |
| chr8 | 75526355 | ATCAATACGTGGA-CACTTTGG-CATTCATTCTTGG | loxF8L | 3 | loxF8R | 4 | CACTTTGG/CCAAAGTG | 7 |
| --- | | | | | | | | 0 |
| chr3 | 171917111 | CAAAAATTGTTGG-TAAAAGTG-TAAACAGATTTTT | loxF8R | 4 | loxF8L | 3 | CACTTTTA/TAAAAGTG | 7 |
| chr14 | 100488830 | CTTGATCTGTGGG-CACTTTTA-TCCACAGGTGTTT | loxF8L | 4 | loxF8L | 3 | CACTTTTA/TAAAAGTG | 7 |
| chrX | 131119677 | ATAAATTTGGGAA-TAAAAGTG-TCAACTGCTATAT | loxF8L | 3 | loxF8L | 4 | CACTTTTA/TAAAAGTG | 7 |
| --- | | | | | | | | 0 |
| chrX | 56261467 | AAAAATCTGATGA-CAGAACAA-TCCTCAGATGAGT | loxF8L | 3 | loxF8L | 4 | CAGAACAA/TTGTTCTG | 7 |
| chr20 | 24256143 | CTAAGAGCCTGTG-CAGAACAA-GCCTCCGATTCAT | loxF8R | 3 | loxF8L | 4 | CAGAACAA/TTGTTCTG | 7 |
| --- | | | | | | | | 0 |
| chr20 | 50545031 | ATTAGATTGTTAG-CAGAAGTA-GCCACAGGTTTCT | loxF8R | 4 | loxF8L | 3 | CAGAAGTA/TACTTCTG | 7 |
| chr2 | 190815128 | GTATATCTGTGCA-TACTTCTG-GACACAAGCCAAG | loxF8L | 3 | loxF8R | 4 | CAGAAGTA/TACTTCTG | 7 |
| --- | | | | | | | | 0 |
| chr11 | 82767692 | ATAAATCTGCCTA-GTTCTCTG-CTCTTAGATTTAT | loxF8L | 3 | loxF8L | 4 | CAGAGAAC/GTTCTCTG | 7 |
| chr8 | 101943077 | CAAAGAGTGTGTA-CAGAGAAC-CAAACATTGTTGG | loxF8R | 3 | loxF8R | 4 | CAGAGAAC/GTTCTCTG | 7 |
| chr15 | 61762468 | CTAAAACTGGGGA-CAGAGAAC-CAAATAATATAAG | loxF8L | 3 | loxF8R | 4 | CAGAGAAC/GTTCTCTG | 7 |
| --- | | | | | | | | 0 |
| chr14 | 44360681 | TTAAGACTATGTG-TATATCTG-TACTCTGATGTAT | loxF8R | 3 | loxF8L | 4 | CAGATATA/TATATCTG | 7 |
| chr14 | 58962277 | ATATATCTATAGA-TATATCTG-TATATAGATATAT | loxF8L | 3 | loxF8L | 4 | CAGATATA/TATATCTG | 7 |
| chr2 | 35147240 | ATATATGTGTGTA-TATATCTG-TCAATATATTAAT | loxF8L | 3 | loxF8L | 4 | CAGATATA/TATATCTG | 7 |
| --- | | | | | | | | 0 |
| chr10 | 84076288 | ATAACACTGGGTA-GAAATCTG-AAGACACTCTTAG | loxF8L | 4 | loxF8R | 3 | CAGATTTC/GAAATCTG | 7 |
| chr1 | 98770535 | ATAATTCAGTGGC-CAGATTTC-ACCACAAATTCAA | loxF8L | 3 | loxF8L | 4 | CAGATTTC/GAAATCTG | 7 |
| --- | | | | | | | | 0 |
| chr13 | 105797853 | CTAAGATCCTTTG-CAGGAAGA-AATAAAGATTTAT | loxF8R | 3 | loxF8L | 4 | CAGGAAGA/TCTTCCTG | 7 |
| chr2 | 227187119 | CTATCTTTGTGTG-CAGGAAGA-CACACTGTCTTTA | loxF8R | 3 | loxF8R | 4 | CAGGAAGA/TCTTCCTG | 7 |
| --- | | | | | | | | 0 |
| chr14 | 67498293 | ATCACTCTGTGGT-TCTCCCTG-TGCACACATTAGT | loxF8L | 3 | loxF8L | 4 | CAGGGAGA/TCTCCCTG | 7 |
| chr15 | 86590050 | ATTACTGTGTGCA-CAGGGAGA-ACCACAGAGTGAT | loxF8L | 4 | loxF8L | 3 | CAGGGAGA/TCTCCCTG | 7 |
| --- | | | | | | | | 0 |
| chr3 | 141896467 | CTATGATTGATTG-TGCCACTG-CACTCCAGCTTGG | loxF8R | 3 | loxF8R | 4 | CAGTGGCA/TGCCACTG | 7 |
| chr10 | 21569295 | CTAGGCTGGAGTG-CAGTGGCA-TGCACAATCATAG | loxF8R | 4 | loxF8R | 3 | CAGTGGCA/TGCCACTG | 7 |
| --- | | | | | | | | 0 |
| chr3 | 132313513 | CATAGATTCTGTG-TTACACTG-CCCAAGGATTCAT | loxF8R | 3 | loxF8L | 4 | CAGTGTAA/TTACACTG | 7 |
| chr5 | 38497301 | ATGAATCTCTCAA-CAGTGTAA-AACACAGATTTAA | loxF8L | 4 | loxF8L | 3 | CAGTGTAA/TTACACTG | 7 |
| --- | | | | | | | | 0 |
| chr17 | 55154390 | ATAAATTTTTGGT-GGACACTG-CCAACAGGTATAT | loxF8L | 3 | loxF8L | 4 | CAGTGTCC/GGACACTG | 7 |
| chr1 | 247744609 | ATCTATTGGTGGA-GGACACTG-TCCATAGATCTAG | loxF8L | 4 | loxF8L | 3 | CAGTGTCC/GGACACTG | 7 |
| --- | | | | | | | | 0 |
| chr18 | 46611728 | CTAACATTTTGTC-CAGTTTTC-TCCAGACCTTTCT | loxF8R | 3 | loxF8L | 4 | CAGTTTTC/GAAAACTG | 7 |
| chr6 | 98283895 | GTAAATCTGATGA-CAGTTTTC-TCTTCAGTATTAT | loxF8L | 3 | loxF8L | 4 | CAGTTTTC/GAAAACTG | 7 |
| chr12 | 102994264 | TTTAATTTGGGGA-CAGTTTTC-TACACAGCATTAT | loxF8L | 4 | loxF8L | 3 | CAGTTTTC/GAAAACTG | 7 |
| --- | | | | | | | | 0 |
| chr2 | 2141289 | ATAAACATATGGA-TTTTTATG-TCAATAAATTTAA | loxF8L | 3 | loxF8L | 4 | CATAAAAA/TTTTTATG | 7 |
| chr18 | 77766182 | CAAAAATTATGTT-CATAAAAA-TTCAAAGATGTAT | loxF8R | 4 | loxF8L | 3 | CATAAAAA/TTTTTATG | 7 |
| --- | | | | | | | | 0 |
| chr2 | 62062683 | AAAAATCTCTGTT-CATTTATG-TCTTAAGATTTAT | loxF8L | 4 | loxF8L | 3 | CATAAATG/CATTTATG | 7 |
| chr10 | 64432748 | ATATGTGTGTGTA-CATTTATG-TACACATATGTAT | loxF8L | 4 | loxF8L | 3 | CATAAATG/CATTTATG | 7 |
| --- | | | | | | | | 0 |
| chr8 | 91144961 | ACAAATATGTGCA-CATACAGA-CAAGCATTCCTAG | loxF8L | 3 | loxF8R | 4 | CATACAGA/TCTGTATG | 7 |
| chr4 | 36291468 | ATAACTCTGAGGC-TCTGTATG-ATCACAAATTCAT | loxF8L | 3 | loxF8L | 4 | CATACAGA/TCTGTATG | 7 |
| --- | | | | | | | | 0 |
| chr9 | 2260309 | ATATGTTTGTGTG-CATGTATG-TACACAGGTATGT | loxF8R | 3 | loxF8L | 4 | CATACATG/CATGTATG | 7 |
| chr5 | 100499233 | AGAAATCAATGAA-CATACATG-CTCACACTTTTAG | loxF8L | 4 | loxF8R | 3 | CATACATG/CATGTATG | 7 |
| chr5 | 7833600 | ATACATATCTGTA-CATACATG-TACACACATATAT | loxF8L | 4 | loxF8L | 3 | CATACATG/CATGTATG | 7 |
| --- | | | | | | | | 0 |
| chr2 | 129174513 | ATAATACTATGGA-CATAGATG-CAAAAATTCTTAA | loxF8L | 3 | loxF8R | 4 | CATAGATG/CATCTATG | 7 |
| chr10 | 52743926 | CTCAGAGTGTGTT-CATCTATG-AATACAGTATTAG | loxF8R | 3 | loxF8R | 4 | CATAGATG/CATCTATG | 7 |
| --- | | | | | | | | 0 |
| chr5 | 134762115 | ATAAATAGGTGGG-CATAGTGG-CACACGCTTTTAT | loxF8L | 3 | loxF8R | 4 | CATAGTGG/CCACTATG | 7 |
| chr8 | 27847417 | ATTAGTGTGTGTG-CATAGTGG-CACACAACTGTAG | loxF8R | 4 | loxF8R | 3 | CATAGTGG/CCACTATG | 7 |
| --- | | | | | | | | 0 |
| chr4 | 60595367 | ATAGATATGTGTA-TAGATATG-TATATAGATATAT | loxF8L | 3 | loxF8L | 4 | CATATCTA/TAGATATG | 7 |
| chr14 | 58962224 | ATGTATCTGTATA-TAGATATG-TCTATAGATATAT | loxF8L | 4 | loxF8L | 3 | CATATCTA/TAGATATG | 7 |
| --- | | | | | | | | 0 |
| chr12 | 17692877 | AAATTTCTGTGGA-TTAATATG-CTTCCAATGTTAG | loxF8L | 3 | loxF8R | 4 | CATATTAA/TTAATATG | 7 |
| chr2 | 135138573 | ATATATGTGTGTA-CATATTAA-TACATGGTTTTAT | loxF8L | 3 | loxF8L | 4 | CATATTAA/TTAATATG | 7 |
| chr9 | 5485981 | ATAATTATGTGAA-TTAATATG-TGCAAAAATTTAA | loxF8L | 3 | loxF8L | 4 | CATATTAA/TTAATATG | 7 |
| --- | | | | | | | | 0 |
| chr3 | 37166365 | ACAAAACTGTGTA-CATCAAAG-CACACTATCAAAA | loxF8L | 3 | loxF8R | 4 | CATCAAAG/CTTTGATG | 7 |
| chr5 | 96181342 | TCCAGATTCTGTG-CATCAAAG-CACAGCATCTTTG | loxF8R | 4 | loxF8R | 3 | CATCAAAG/CTTTGATG | 7 |
| --- | | | | | | | | 0 |
| chrX | 155832838 | ATCAAACTATTGA-CATCATTA-TTCACAGAATTAG | loxF8L | 4 | loxF8L | 3 | CATCATTA/TAATGATG | 7 |
| chrY | 57019358 | ATCAAACTATTGA-CATCATTA-TTCACAGAATTAG | loxF8L | 4 | loxF8L | 3 | CATCATTA/TAATGATG | 7 |
| --- | | | | | | | | 0 |
| chr15 | 51453239 | ATAAAAATGTTGA-GAATGATG-GAGACAATATCAG | loxF8L | 3 | loxF8R | 4 | CATCATTC/GAATGATG | 7 |
| chrX | 21365759 | CTAATTCTGTGAA-GAATGATG-TTCATAGTTTGAT | loxF8L | 3 | loxF8L | 4 | CATCATTC/GAATGATG | 7 |
| chrX | 25901334 | CTAGTTCTGTGAA-GAATGATG-TTCATAGATTGAT | loxF8L | 4 | loxF8L | 3 | CATCATTC/GAATGATG | 7 |
| --- | | | | | | | | 0 |
| chr1 | 25704769 | AAAAATCTGTTGT-CATCCAGA-TACTCTGATTTAG | loxF8L | 3 | loxF8L | 4 | CATCCAGA/TCTGGATG | 7 |
| chr4 | 164453587 | AAAGAGATGTGGA-CATCCAGA-TCCCCAAATTGAT | loxF8L | 4 | loxF8L | 3 | CATCCAGA/TCTGGATG | 7 |

| | | | | | | | | |
|---|---|---|---|---|---|---|---|---|
| --- | | | | | | | | 0 |
| chr4 | 169479715 | AAAAATCTCTGGT-CTCAGATG-TACACATAGTCAT | loxF8L | 3 | loxF8L | 4 | CATCTGAG/CTCAGATG | 7 |
| chr6 | 34048142 | TCAAGAATGTGTC-CTCAGATG-TCCACAGACCTAG | loxF8R | 4 | loxF8L | 3 | CATCTGAG/CTCAGATG | 7 |
| --- | | | | | | | | 0 |
| chr15 | 47203807 | ATTGTTTTGTGGA-CATGAAAA-TCCACACATTCCT | loxF8L | 4 | loxF8L | 3 | CATGAAAA/TTTTCATG | 7 |
| chr14 | 75200838 | ATTAAATTATGTG-TTTTCATG-TACACAGACATAT | loxF8R | 4 | loxF8L | 3 | CATGAAAA/TTTTCATG | 7 |
| --- | | | | | | | | 0 |
| chr15 | 64545946 | ATAAAGGTGTGAA-CATTCATG-TACACAAAGTTTT | loxF8L | 3 | loxF8L | 4 | CATGAATG/CATTCATG | 7 |
| chr4 | 78264437 | ATAAATATGTGTC-CATTCATG-TAAATAAATTTAT | loxF8L | 3 | loxF8L | 4 | CATGAATG/CATTCATG | 7 |
| chr5 | 16177660 | CTAAGCTTAAGTG-CATTCATG-TTAACAAAATTAT | loxF8R | 3 | loxF8L | 4 | CATGAATG/CATTCATG | 7 |
| --- | | | | | | | | 0 |
| chr7 | 49082842 | CCAACATTCTGTG-CATGATGA-GCCACATATATCT | loxF8R | 3 | loxF8L | 4 | CATGATGA/TCATCATG | 7 |
| chr4 | 120010304 | ATATATCTGTATG-TCATCATG-TGCAAATATTTAT | loxF8L | 4 | loxF8L | 3 | CATGATGA/TCATCATG | 7 |
| --- | | | | | | | | 0 |
| chr4 | 120835377 | CTATGCCTGTGTG-CATGCATG-TACACACATATTT | loxF8R | 3 | loxF8L | 4 | CATGCATG/CATGCATG | 7 |
| chr12 | 103411834 | ATACATGTGTGCA-CATGCATG-CACACAAACACAC | loxF8L | 3 | loxF8R | 4 | CATGCATG/CATGCATG | 7 |
| --- | | | | | | | | 0 |
| chr5 | 168850446 | GTAATATTGTGTC-CATGTAAA-TACAAATATATAT | loxF8R | 3 | loxF8L | 4 | CATGTAAA/TTTACATG | 7 |
| chr8 | 50295629 | CAAAGGTTGTTTG-CATGTAAA-CAAACAACCTTCC | loxF8R | 3 | loxF8R | 4 | CATGTAAA/TTTACATG | 7 |
| --- | | | | | | | | 0 |
| chr13 | 78331377 | AGAAATATGTGAA-CATGTACA-CACATAATCACAC | loxF8L | 3 | loxF8R | 4 | CATGTACA/TGTACATG | 7 |
| chr19 | 51420412 | TAAAGACTGTCTG-CATGTACA-GCCACATTCTTAG | loxF8R | 4 | loxF8R | 3 | CATGTACA/TGTACATG | 7 |
| --- | | | | | | | | 0 |
| chr12 | 47633971 | ATAAATATATGCA-CATGTGTA-TATACACATATAT | loxF8L | 3 | loxF8L | 4 | CATGTGTA/TACACATG | 7 |
| chr10 | 129328041 | ATAAATATGAATA-TACACATG-TCCACCTACTTAT | loxF8L | 4 | loxF8L | 3 | CATGTGTA/TACACATG | 7 |
| chr6 | 103746485 | ATACATGTGTGTA-TACACATG-CACATGATTTTGG | loxF8L | 3 | loxF8R | 4 | CATGTGTA/TACACATG | 7 |
| chr7 | 129297058 | TGAAGACTGTGTG-TACACATG-CACACTGATCTAT | loxF8R | 3 | loxF8L | 4 | CATGTGTA/TACACATG | 7 |
| --- | | | | | | | | 0 |
| chr10 | 91870399 | AAAAAGGAGTGGA-CATTAATG-CACAAAATGTTAA | loxF8L | 4 | loxF8R | 3 | CATTAATG/CATTAATG | 7 |
| --- | | | | | | | | 0 |
| chr3 | 90708483 | AAAAGTTTCTGTG-CATTCTTC-TGTATAGATTTAT | loxF8R | 4 | loxF8L | 3 | CATTCTTC/GAAGAATG | 7 |
| chr11 | 94154342 | ATAAGTTTGTGCA-GAAGAATG-TTCTCTGAGTTAT | loxF8L | 3 | loxF8L | 4 | CATTCTTC/GAAGAATG | 7 |
| chr6 | 15155995 | CTAAGACAGTGAT-GAAGAATG-TATACAATCTTAA | loxF8R | 4 | loxF8R | 3 | CATTCTTC/GAAGAATG | 7 |
| --- | | | | | | | | 0 |
| chr2 | 123454845 | CCAAGTTTCTGTG-GGACAATG-TCCTTAGATTCCT | loxF8R | 3 | loxF8L | 4 | CATTGTCC/GGACAATG | 7 |
| chr11 | 31772479 | CTAAGTATCTGTG-CATTGTCC-CTCCTAATCTTTG | loxF8R | 3 | loxF8R | 4 | CATTGTCC/GGACAATG | 7 |
| --- | | | | | | | | 0 |
| chr4 | 135677970 | ACAAATATGTGAA-CATTTAAC-AATACAGTCTTAA | loxF8L | 3 | loxF8R | 4 | CATTTAAC/GTTAAATG | 7 |
| chr1 | 78427235 | CTACTATGGTGTG-CATTTAAC-TTAAAATATTTAT | loxF8R | 3 | loxF8L | 4 | CATTTAAC/GTTAAATG | 7 |
| --- | | | | | | | | 0 |
| chr8 | 81235089 | ATAGATGTGTGGC-CATTTGAA-ACCAGAGATATCT | loxF8L | 3 | loxF8L | 4 | CATTTGAA/TTCAAATG | 7 |
| chr3 | 48328521 | TAAAATCTTTGGA-CATTTGAA-AAAAAAATCTTAA | loxF8L | 3 | loxF8R | 4 | CATTTGAA/TTCAAATG | 7 |
| --- | | | | | | | | 0 |
| chr15 | 43459000 | AGAAAGATGTGCA-TCCAAATG-ATCACAGAATTAT | loxF8L | 4 | loxF8L | 3 | CATTTGGA/TCCAAATG | 7 |
| chr5 | 44871071 | CTAAAAGTGTGTT-TCCAAATG-TCCACAGCTTACC | loxF8R | 3 | loxF8L | 4 | CATTTGGA/TCCAAATG | 7 |
| --- | | | | | | | | 0 |
| chr3 | 106647673 | TCAAGATAGTGGG-TGAAAATG-TTCACTCATTTAT | loxF8R | 4 | loxF8L | 3 | CATTTTCA/TGAAAATG | 7 |
| chr2 | 73419034 | ATACATGAGTGGA-CATTTTCA-TCTAAATATTAAT | loxF8L | 3 | loxF8L | 4 | CATTTTCA/TGAAAATG | 7 |
| --- | | | | | | | | 0 |
| chrX | 48215074 | ATAATTCATTGTA-TAAAAATG-CTAACAATCTTCG | loxF8L | 4 | loxF8R | 3 | CATTTTTA/TAAAAATG | 7 |
| chr8 | 75196118 | CTAAATTTTTTGA-TAAAAATG-AACACAATGGTAG | loxF8L | 4 | loxF8R | 3 | CATTTTTA/TAAAAATG | 7 |
| --- | | | | | | | | 0 |
| chr6 | 21993956 | ATAAAGATGGGGA-CCAACACA-CACACACTCACAA | loxF8L | 3 | loxF8R | 4 | CCAACACA/TGTGTTGG | 7 |
| chr7 | 124029780 | CAGAGATTTTGTC-TGTGTTGG-TTCACAGCTATAT | loxF8R | 4 | loxF8L | 3 | CCAACACA/TGTGTTGG | 7 |
| --- | | | | | | | | 0 |
| chr18 | 30350694 | AAAAATAGTTGGA-TGTGGTGG-CACACATTTGTAG | loxF8L | 4 | loxF8R | 3 | CCACCACA/TGTGGTGG | 7 |
| chr12 | 65530838 | ATATATGGCTGGA-TGTGGTGG-CTCACAATTGTAG | loxF8L | 4 | loxF8R | 3 | CCACCACA/TGTGGTGG | 7 |
| --- | | | | | | | | 0 |
| chrX | 3979573 | CTAAAGGTGTGTG-CCACCACG-CCCAGCGATTTTT | loxF8R | 3 | loxF8L | 4 | CCACCACG/CGTGGTGG | 7 |
| chr7 | 57098954 | ATAAATCTCTTTT-CCACCACG-ACCAAAGATTTTT | loxF8L | 4 | loxF8L | 3 | CCACCACG/CGTGGTGG | 7 |
| --- | | | | | | | | 0 |
| chr19 | 57587859 | CTGAGAGTCTGTG-CCACTGAG-GCCACTGATTTTG | loxF8R | 3 | loxF8L | 4 | CCACTGAG/CTCAGTGG | 7 |
| chr14 | 72862099 | TTAAGAATGGGAG-CTCAGTGG-CACACGACCTGAG | loxF8R | 4 | loxF8R | 3 | CCACTGAG/CTCAGTGG | 7 |
| --- | | | | | | | | 0 |
| chr5 | 154625236 | AAAAATCAGTGGG-GATGATGG-AACAAAATGCTAG | loxF8L | 3 | loxF8R | 4 | CCATCATC/GATGATGG | 7 |
| chr7 | 63476089 | AAAAATCTTTGGT-GATGATGG-AAAAGAGATTTAT | loxF8L | 3 | loxF8L | 4 | CCATCATC/GATGATGG | 7 |
| --- | | | | | | | | 0 |
| chr13 | 107102404 | ACAAATTTGTGGT-TGAGATGG-AAAACAATTTTAA | loxF8L | 3 | loxF8R | 4 | CCATCTCA/TGAGATGG | 7 |
| chr6 | 24539301 | TTAAAATTGTTTG-CCATCTCA-ACCACAGCCTTGT | loxF8R | 3 | loxF8L | 4 | CCATCTCA/TGAGATGG | 7 |
| chr4 | 105085356 | TTAAAGTTGTTTG-CCATCTCA-ACCACAGACTTGT | loxF8R | 4 | loxF8L | 3 | CCATCTCA/TGAGATGG | 7 |
| --- | | | | | | | | 0 |
| chr6 | 25014763 | CGAAAGTTGTGTG-TAACATGG-CACAAATTATTAT | loxF8R | 3 | loxF8R | 4 | CCATGTTA/TAACATGG | 7 |
| chr8 | 31004125 | ATGAATCTTTGCA-TAACATGG-CAAACAATTTAAT | loxF8L | 3 | loxF8R | 4 | CCATGTTA/TAACATGG | 7 |
| --- | | | | | | | | 0 |
| chr4 | 109981849 | AAAAAGCTGTGTA-GTCAATGG-TCAAGAGTTTTAA | loxF8L | 3 | loxF8L | 4 | CCATTGAC/GTCAATGG | 7 |
| chr13 | 89116668 | TTAAGAATGTGTT-CCATTGAC-TCCTAACGTTTAT | loxF8R | 3 | loxF8L | 4 | CCATTGAC/GTCAATGG | 7 |
| --- | | | | | | | | 0 |
| chr12 | 34061528 | CTAAAATTATTTC-CCCAACAC-TTCTCAGAATTAT | loxF8R | 4 | loxF8L | 3 | CCCAACAC/GTGTTGGG | 7 |
| chr12 | 38360037 | CTAAAATTATTTC-CCCAACAC-TTCTCAGAATTAT | loxF8R | 4 | loxF8L | 3 | CCCAACAC/GTGTTGGG | 7 |
| --- | | | | | | | | 0 |
| chr9 | 13677990 | ATACATATATGCA-CCCACACA-TGCACAGACATAT | loxF8L | 4 | loxF8L | 3 | CCCACACA/TGTGTGGG | 7 |
| chr18 | 65406994 | ATATATATGTGTA-TGTGTGGG-TACACACACATAT | loxF8L | 3 | loxF8L | 4 | CCCACACA/TGTGTGGG | 7 |
| --- | | | | | | | | 0 |
| chr5 | 90156760 | ATAAATATGTGAC-CCCATTAA-GACAAAAACTTGG | loxF8L | 3 | loxF8R | 4 | CCCATTAA/TTAATGGG | 7 |
| chr10 | 22797715 | ATATATATGTGAC-TTAATGGG-TCTACAGTTTTAA | loxF8L | 4 | loxF8L | 3 | CCCATTAA/TTAATGGG | 7 |
| chr7 | 128340292 | ATCAATTTATGGA-TTAATGGG-TTAATAGATTAAT | loxF8L | 3 | loxF8L | 4 | CCCATTAA/TTAATGGG | 7 |
| chr3 | 62606335 | ATCTATTTATGGA-TTAATGGG-TTAACAGATTAAT | loxF8L | 4 | loxF8L | 3 | CCCATTAA/TTAATGGG | 7 |

| | | | | | | | | |
|---|---|---|---|---|---|---|---|---|
| --- | | | | | | | | 0 |
| chr12 | 5815331 | AGAAATCAGAGCA-GTGTGGGG-CAAACAATCCTAA | loxF8L | 4 | loxF8R | 3 | CCCCACAC/GTGTGGGG | 7 |
| chr19 | 30132319 | CTGACTTTGTGTG-CCCCACAC-CACCCAAACCTGG | loxF8R | 3 | loxF8R | 4 | CCCCACAC/GTGTGGGG | 7 |
| --- | | | | | | | | 0 |
| chr5 | 35324716 | AGAGAACAGTGGA-CCCTCCCC-CACCCACTCTGAG | loxF8L | 4 | loxF8R | 3 | CCCTCCCC/GGGGAGGG | 7 |
| chr7 | 104999147 | GTGAGATTCCGTG-CCCTCCCC-CCCAAAATCTTAT | loxF8R | 4 | loxF8R | 3 | CCCTCCCC/GGGGAGGG | 7 |
| --- | | | | | | | | 0 |
| chrX | 20316614 | CTCAGAGTCTGTT-TTCCAGGG-CACACAACCTGTG | loxF8R | 4 | loxF8R | 3 | CCCTGGAA/TTCCAGGG | 7 |
| chr15 | 57579957 | CTCAGAGTCTGTG-TTCCAGGG-AACACAAGCAAAG | loxF8R | 3 | loxF8R | 4 | CCCTGGAA/TTCCAGGG | 7 |
| --- | | | | | | | | 0 |
| chr3 | 158395017 | ATAAAGCTGCACA-CCTACAAC-CACTCAATCTTTA | loxF8L | 4 | loxF8R | 3 | CCTACAAC/GTTGTAGG | 7 |
| chr5 | 147146282 | ATAAAGCTGCACA-CCTACAAC-CATCCAATCTTTG | loxF8L | 4 | loxF8R | 3 | CCTACAAC/GTTGTAGG | 7 |
| chr8 | 70817799 | ATAAAGCTGTACA-CCTACAAC-CACCTGATCTTTG | loxF8L | 3 | loxF8R | 4 | CCTACAAC/GTTGTAGG | 7 |
| chrX | 57080589 | ATAAAGCTGTACA-CCTACAAC-CACTTAATCTTCA | loxF8L | 3 | loxF8R | 4 | CCTACAAC/GTTGTAGG | 7 |
| chr17 | 6744420 | ATAAATCTGCACA-CCTACAAC-CACCTGATCTTTG | loxF8L | 3 | loxF8R | 4 | CCTACAAC/GTTGTAGG | 7 |
| chr1 | 189049189 | CAAAGATTCTATG-GTTGTAGG-TGTACAGCGTTAT | loxF8R | 3 | loxF8L | 4 | CCTACAAC/GTTGTAGG | 7 |
| chr17 | 34230587 | ATAATATTGTGTA-CCTACAAC-CACCTGATCTTTG | loxF8R | 3 | loxF8R | 4 | CCTACAAC/GTTGTAGG | 7 |
| chrX | 64243991 | CGAAGATTATATG-GTTGTAGG-TCTGCAGCATTAT | loxF8R | 3 | loxF8L | 4 | CCTACAAC/GTTGTAGG | 7 |
| --- | | | | | | | | 0 |
| chr9 | 7347999 | ATAAAACTTTGAA-GATGTAGG-CAAACAGTATTTG | loxF8L | 3 | loxF8R | 4 | CCTACATC/GATGTAGG | 7 |
| chr9 | 105791573 | CTAAGATTAGATG-GATGTAGG-TGTGCAGCTTTAT | loxF8R | 3 | loxF8L | 4 | CCTACATC/GATGTAGG | 7 |
| --- | | | | | | | | 0 |
| chr11 | 8502322 | ATAATGCTGTGCA-CCTACCAG-CATCAAATCTTTG | loxF8L | 3 | loxF8R | 4 | CCTACCAG/CTGGTAGG | 7 |
| chr3 | 112428744 | GTAGATCTGTAGA-CCTACCAG-TATGCAATGTTAG | loxF8L | 3 | loxF8R | 4 | CCTACCAG/CTGGTAGG | 7 |
| --- | | | | | | | | 0 |
| chrX | 85613813 | CTACGATCATGTG-GCTATAGG-TGTTCAGCTTTAT | loxF8R | 3 | loxF8L | 4 | CCTATAGC/GCTATAGG | 7 |
| chr2 | 84380433 | ATAAAGCTGCGCA-CCTATAGC-CATCTGATCTTAG | loxF8L | 3 | loxF8R | 4 | CCTATAGC/GCTATAGG | 7 |
| chr2 | 68680199 | CAAAGATTTGATG-GCTATAGG-TGTACAGCTTTAT | loxF8R | 4 | loxF8L | 3 | CCTATAGC/GCTATAGG | 7 |
| --- | | | | | | | | 0 |
| chr8 | 7364181 | ATAGATATGTAGA-TATATAGG-TGCATATGTTTAT | loxF8L | 3 | loxF8L | 4 | CCTATATA/TATATAGG | 7 |
| chr18 | 22768100 | ATAAATGGGAGGA-TATATAGG-TATACAATGTTAA | loxF8L | 3 | loxF8R | 4 | CCTATATA/TATATAGG | 7 |
| chr12 | 118823682 | ATATATATGTGTA-TATATAGG-CTCAGAATCTTGA | loxF8L | 3 | loxF8R | 4 | CCTATATA/TATATAGG | 7 |
| --- | | | | | | | | 0 |
| chr2 | 39081919 | AAAATTTTGTGGG-TACATAGG-TGCATATATTTAT | loxF8L | 4 | loxF8L | 3 | CCTATGTA/TACATAGG | 7 |
| chr7 | 8083132 | AAATTTCTGTGGA-TACATAGG-TGTATATATTTAT | loxF8L | 3 | loxF8L | 4 | CCTATGTA/TACATAGG | 7 |
| chr13 | 108491511 | ATAAATATATACA-CCTATGTA-TCCACAAAATTAA | loxF8L | 4 | loxF8L | 3 | CCTATGTA/TACATAGG | 7 |
| --- | | | | | | | | 0 |
| chr2 | 124069682 | ACAAGATCCTGTG-CCTGATAA-TCCAGAGAATTCT | loxF8R | 4 | loxF8L | 3 | CCTGATAA/TTATCAGG | 7 |
| chr5 | 85897968 | AGAATTCTCTGGA-TTATCAGG-CAAAGACTCTTAT | loxF8L | 3 | loxF8R | 4 | CCTGATAA/TTATCAGG | 7 |
| chr5 | 108240912 | TTAAGAGTCTCTG-CCTGATAA-TTCACAGAATTCT | loxF8R | 4 | loxF8L | 3 | CCTGATAA/TTATCAGG | 7 |
| chrX | 109102221 | ATACTTCTCTGGA-TTATCAGG-CAAAGACTCTTAT | loxF8L | 3 | loxF8R | 4 | CCTGATAA/TTATCAGG | 7 |
| --- | | | | | | | | 0 |
| chr5 | 84025417 | ATAAAACTGTCTA-GTGGCAGG-AAAACAAGCTCAG | loxF8L | 3 | loxF8R | 4 | CCTGCCAC/GTGGCAGG | 7 |
| chr20 | 52421100 | ATAGATTTGTAGA-CCTGCCAC-CACCAATTCATAG | loxF8L | 3 | loxF8R | 4 | CCTGCCAC/GTGGCAGG | 7 |
| --- | | | | | | | | 0 |
| chr5 | 137165262 | CCAAGAGTCTCTG-CCTGCTAA-TCCAGAGAATTCT | loxF8R | 4 | loxF8L | 3 | CCTGCTAA/TTAGCAGG | 7 |
| chr7 | 42274729 | CTGAGAAGGTCTG-TTAGCAGG-TCCCCAGAACTAT | loxF8R | 4 | loxF8L | 3 | CCTGCTAA/TTAGCAGG | 7 |
| --- | | | | | | | | 0 |
| chr3 | 153866165 | ACAAGAGTGTCTG-CCTGGGAA-TCCAGAGAATTCT | loxF8R | 4 | loxF8L | 3 | CCTGGGAA/TTCCCAGG | 7 |
| chr12 | 77345138 | ATATAGCTGGGAA-CCTGGGAA-CATACAATATTTG | loxF8L | 4 | loxF8R | 3 | CCTGGGAA/TTCCCAGG | 7 |
| --- | | | | | | | | 0 |
| chrX | 131676941 | ATCTGATTCTGTG-CTCCCAGG-ATCAAAGATTTAT | loxF8R | 4 | loxF8L | 3 | CCTGGGAG/CTCCCAGG | 7 |
| chr21 | 32662485 | ATCCCTCTGTGGC-CTCCCAGG-TCCACAGTTTTTG | loxF8L | 4 | loxF8L | 3 | CCTGGGAG/CTCCCAGG | 7 |
| --- | | | | | | | | 0 |
| chr15 | 78702240 | AGAATTCTCTGGA-CTACCAGG-CAGAGACTCTTAT | loxF8L | 3 | loxF8R | 4 | CCTGGTAG/CTACCAGG | 7 |
| chr15 | 84596321 | ATAAGAATCTCTG-CCTGGTAG-TCCAGAGAATTCT | loxF8R | 4 | loxF8L | 3 | CCTGGTAG/CTACCAGG | 7 |
| chr6 | 102154832 | CAAAGAATCTCTG-CCTGGTAG-TCCAGAGAATTCT | loxF8R | 4 | loxF8L | 3 | CCTGGTAG/CTACCAGG | 7 |
| --- | | | | | | | | 0 |
| chr10 | 120329955 | ACAAATCTGTCCC-CCTGTATA-TCAACAGATCTAA | loxF8L | 4 | loxF8L | 3 | CCTGTATA/TATACAGG | 7 |
| chr17 | 50066844 | ACAAATGTATGTA-CCTGTATA-ACCAAAGATATAT | loxF8L | 4 | loxF8L | 3 | CCTGTATA/TATACAGG | 7 |
| chr13 | 104159760 | ATAAATCTATATG-CCTGTATA-TACATAGATTTAC | loxF8L | 4 | loxF8L | 3 | CCTGTATA/TATACAGG | 7 |
| --- | | | | | | | | 0 |
| chr17 | 9650349 | ACAAATGTGTGAA-CCTGTGTA-GTCACCATCTTAC | loxF8L | 3 | loxF8R | 4 | CCTGTGTA/TACACAGG | 7 |
| chr3 | 15543946 | ATAAAATTGTGGG-TACACAGG-TGTATAAATTTAT | loxF8L | 3 | loxF8L | 4 | CCTGTGTA/TACACAGG | 7 |
| --- | | | | | | | | 0 |
| chr6 | 157641335 | ACAAGTCTGTGCT-CCTGTGTC-CCCACACCTTTAT | loxF8L | 4 | loxF8L | 3 | CCTGTGTC/GACACAGG | 7 |
| chr3 | 62540616 | TTCAGATTGTCTG-CCTGTGTC-TCCATACATGTCT | loxF8R | 3 | loxF8L | 4 | CCTGTGTC/GACACAGG | 7 |
| --- | | | | | | | | 0 |
| chr5 | 161180792 | ATAAGAGTCTCTG-CCTTCCAC-TCCAGAGAATTCT | loxF8R | 4 | loxF8L | 3 | CCTTCCAC/GTGGAAGG | 7 |
| chr3 | 174348855 | CTATTATTGTGTT-CCTTCCAC-TCCAGATACTTAA | loxF8R | 3 | loxF8L | 4 | CCTTCCAC/GTGGAAGG | 7 |
| --- | | | | | | | | 0 |
| chr15 | 82679056 | AGAAAAGTGTGAA-CCTTGGAG-TCAGCAGAATTAT | loxF8L | 4 | loxF8L | 3 | CCTTGGAG/CTCCAAGG | 7 |
| chr20 | 16306976 | CTAGGATGATCTG-CTCCAAGG-TCCACACTCTTAG | loxF8R | 4 | loxF8R | 3 | CCTTGGAG/CTCCAAGG | 7 |
| --- | | | | | | | | 0 |
| chr15 | 60924581 | ATAAGTCACTGGC-CCTTTCTA-TCCTCAGATTCCT | loxF8L | 4 | loxF8L | 3 | CCTTTCTA/TAGAAAGG | 7 |
| chr4 | 139485382 | GAAAACCTGTGGA-CCTTTCTA-CAAACTATCAGAG | loxF8L | 3 | loxF8R | 4 | CCTTTCTA/TAGAAAGG | 7 |
| --- | | | | | | | | 0 |
| chr6 | 62169138 | ATATATGTGTGTA-TATATACG-TACACATATATGT | loxF8L | 3 | loxF8L | 4 | CGTATATA/TATATACG | 7 |
| chr3 | 18477998 | GTATGTGTGTGTG-TATATACG-CACACTATATTTG | loxF8R | 4 | loxF8R | 3 | CGTATATA/TATATACG | 7 |
| --- | | | | | | | | 0 |
| chr6 | 16589376 | AGAAATGTGTGTA-TATACACG-CACACATATATAT | loxF8L | 3 | loxF8L | 4 | CGTGTATA/TATACACG | 7 |
| chr9 | 119228285 | ATACATGTATGTA-CGTGTATA-CACACACACATAG | loxF8L | 4 | loxF8R | 3 | CGTGTATA/TATACACG | 7 |
| chrX | 67970312 | ATATATATGTGTA-TATACACG-CACACAATGGAAT | loxF8L | 3 | loxF8R | 4 | CGTGTATA/TATACACG | 7 |
| --- | | | | | | | | 0 |
| chr18 | 15132929 | AAATATCTATTGA-CTAAAGAA-AAGGCAATCTTAG | loxF8L | 4 | loxF8R | 3 | CTAAAGAA/TTCTTTAG | 7 |
| chr21 | 27874362 | CTAAGATTGCCTT-TTCTTTAG-TCAATAGATATTT | loxF8R | 3 | loxF8L | 4 | CTAAAGAA/TTCTTTAG | 7 |
| chr2 | 131977544 | AAATATCTATTGA-CTAAAGAA-AAGGCAATCTTAG | loxF8L | 4 | loxF8R | 3 | CTAAAGAA/TTCTTTAG | 7 |

| | | | | | | | | |
|---|---|---|---|---|---|---|---|---|
| --- | | | | | | | | 0 |
| chr10 | 35014503 | CTAAGACATTCTG-CTAAGGAA-TGGACAGATTCAT | loxF8R | 4 | loxF8L | 3 | CTAAGGAA/TTCCTTAG | 7 |
| chr1 | 71776835 | CAAAATTTGTGTT-CTAAGGAA-CACAAATTCTAAG | loxF8R | 4 | loxF8R | 3 | CTAAGGAA/TTCCTTAG | 7 |
| --- | | | | | | | | 0 |
| chrY | 17811537 | GTAAAATTGTGAT-CTACAATC-TCTACAGCTTTAA | loxF8R | 4 | loxF8L | 3 | CTACAATC/GATTGTAG | 7 |
| chrY | 18094438 | TTAAAGCTGTAGA-GATTGTAG-ATCACAATTTTAC | loxF8L | 3 | loxF8R | 4 | CTACAATC/GATTGTAG | 7 |
| --- | | | | | | | | 0 |
| chr9 | 32848002 | AAAAGTCTGTTGA-CTACTCAA-GCTACAGTTTCAT | loxF8L | 3 | loxF8L | 4 | CTACTCAA/TTGAGTAG | 7 |
| chr7 | 38274582 | ATTAAATCGTGTG-CTACTCAA-CAGAGAATCTGAG | loxF8R | 4 | loxF8R | 3 | CTACTCAA/TTGAGTAG | 7 |
| --- | | | | | | | | 0 |
| chr2 | 235914169 | ATGTTTCTGTTGA-TTCTCTAG-CACACAGTCATAA | loxF8L | 4 | loxF8R | 3 | CTAGAGAA/TTCTCTAG | 7 |
| chr8 | 30450342 | CTTAGATTTGGTG-TTCTCTAG-CAGGCAGACTTAG | loxF8R | 3 | loxF8R | 4 | CTAGAGAA/TTCTCTAG | 7 |
| --- | | | | | | | | 0 |
| chr3 | 81190131 | GTAAGAAAATGTG-CTATGAAA-ACCACTGATTTTT | loxF8R | 4 | loxF8L | 3 | CTATGAAA/TTTCATAG | 7 |
| chr2 | 183293168 | ATAACTCAGTGGG-TTTCATAG-CACATATATTTAT | loxF8L | 3 | loxF8L | 4 | CTATGAAA/TTTCATAG | 7 |
| --- | | | | | | | | 0 |
| chr11 | 26998759 | CCAAGAAGCTGTG-TCCAATAG-TTCACAGACTTTT | loxF8R | 4 | loxF8L | 3 | CTATTGGA/TCCAATAG | 7 |
| chr5 | 41111956 | ATATTTTTGAGGA-CTATTGGA-CACAGAGTCTGAG | loxF8L | 4 | loxF8R | 3 | CTATTGGA/TCCAATAG | 7 |
| --- | | | | | | | | 0 |
| chr10 | 18709664 | GTATATATATGGA-CTCACATA-CACACAAATATAG | loxF8L | 4 | loxF8R | 3 | CTCACATA/TATGTGAG | 7 |
| chr20 | 6689584 | GTGAGCATGTGTG-CTCACATA-CACACAAACTCAC | loxF8R | 4 | loxF8R | 3 | CTCACATA/TATGTGAG | 7 |
| --- | | | | | | | | 0 |
| chr17 | 64663722 | ATACATCAGTGAA-CTCACTTC-TCCAGATAGTTAC | loxF8L | 3 | loxF8L | 4 | CTCACTTC/GAAGTGAG | 7 |
| chr5 | 59650779 | ATATATCTATGCA-CTCACTTC-TCCATAATTTGAT | loxF8L | 3 | loxF8L | 4 | CTCACTTC/GAAGTGAG | 7 |
| --- | | | | | | | | 0 |
| chrY | 25782351 | ATATATATTTGGA-GAATGGAG-ACAACAAACTTAG | loxF8L | 3 | loxF8R | 4 | CTCCATTC/GAATGGAG | 7 |
| chrY | 23887721 | CTAAGTTTGTTGT-CTCCATTC-TCCAAATATATAT | loxF8R | 4 | loxF8L | 3 | CTCCATTC/GAATGGAG | 7 |
| --- | | | | | | | | 0 |
| chrX | 1055392 | CTAAGATGTTTTG-GTGGGGAG-CCCTGATATTTAT | loxF8R | 3 | loxF8L | 4 | CTCCCCAC/GTGGGGAG | 7 |
| chrY | 1055392 | CTAAGATGTTTTG-GTGGGGAG-CCCTGATATTTAT | loxF8R | 3 | loxF8L | 4 | CTCCCCAC/GTGGGGAG | 7 |
| --- | | | | | | | | 0 |
| chr5 | 59882366 | AAAAATGTGTGTA-TAGAAGAG-TCCAAACTTTTAC | loxF8L | 3 | loxF8L | 4 | CTCTTCTA/TAGAAGAG | 7 |
| chr13 | 89295906 | TTAACATTGTCTT-TAGAAGAG-CAGACATTCTTAA | loxF8R | 4 | loxF8R | 3 | CTCTTCTA/TAGAAGAG | 7 |
| --- | | | | | | | | 0 |
| chr18 | 34294175 | ATGAATTTGTGGG-CTCTTTGA-TTTACCCATTTAT | loxF8L | 3 | loxF8L | 4 | CTCTTTGA/TCAAAGAG | 7 |
| chr5 | 55703412 | ATAAGTAAGTTGA-CTCTTTGA-GACACAGATTCAT | loxF8L | 4 | loxF8L | 3 | CTCTTTGA/TCAAAGAG | 7 |
| --- | | | | | | | | 0 |
| chr14 | 62585075 | CCAACACTGTGTG-CTGAAAAA-AACACATTCTTTC | loxF8R | 3 | loxF8R | 4 | CTGAAAAA/TTTTTCAG | 7 |
| chr7 | 121376218 | TTTAGGATGTGTG-CTGAAAAA-CAAACAAGTTTAG | loxF8R | 4 | loxF8R | 3 | CTGAAAAA/TTTTTCAG | 7 |
| --- | | | | | | | | 0 |
| chr18 | 27591702 | CTCAGTTTGGGTT-CTGATAAA-TGCACTGATTTTT | loxF8R | 4 | loxF8L | 3 | CTGATAAA/TTTATCAG | 7 |
| chr11 | 129701184 | GTAAATATGAGGA-CTGATAAA-TCCTCAGATCAAA | loxF8L | 3 | loxF8L | 4 | CTGATAAA/TTTATCAG | 7 |
| --- | | | | | | | | 0 |
| chr6 | 18885688 | CAAAGAATTTGTG-CTGCTGCC-CACACAGAATAAG | loxF8R | 3 | loxF8R | 4 | CTGCTGCC/GGCAGCAG | 7 |
| chr5 | 32632176 | CTCCATCTGTGGA-GGCAGCAG-CACAGCATCTTCC | loxF8L | 3 | loxF8L | 4 | CTGCTGCC/GGCAGCAG | 7 |
| --- | | | | | | | | 0 |
| chr12 | 24794276 | ATAAATATCTGAT-CTGTGAAG-TACACATATTTAA | loxF8L | 4 | loxF8L | 3 | CTGTGAAG/CTTCACAG | 7 |
| chr2 | 59170128 | GTAAGACTCTCTG-CTGTGAAG-AACACAACCTTGG | loxF8R | 4 | loxF8R | 3 | CTGTGAAG/CTTCACAG | 7 |
| --- | | | | | | | | 0 |
| chr10 | 37150892 | CTAAGAATTTGAA-CTGTGCCC-CACAGCTTCTTAG | loxF8R | 4 | loxF8R | 3 | CTGTGCCC/GGGCACAG | 7 |
| chr10 | 37163928 | CTAAGAATTTGAA-CTGTGCCC-CACAGCTTCTTAG | loxF8R | 4 | loxF8R | 3 | CTGTGCCC/GGGCACAG | 7 |
| chr10 | 37186791 | CTAAGAATTTGAA-CTGTGCCC-CACAGCTTCTTAG | loxF8R | 4 | loxF8R | 3 | CTGTGCCC/GGGCACAG | 7 |
| chr10 | 37174990 | CTAAGAATTTGAA-CTGTGCCC-CACAGCTTCTTAG | loxF8R | 4 | loxF8R | 3 | CTGTGCCC/GGGCACAG | 7 |
| chr18 | 14785583 | CTAAGAATTTGAA-CTGTGCCC-CACAGCTTCTTAG | loxF8R | 4 | loxF8R | 3 | CTGTGCCC/GGGCACAG | 7 |
| --- | | | | | | | | 0 |
| chr8 | 66003469 | GTAAGTCTATGGA-CTGTGTCA-TCTATAGAGATAT | loxF8L | 3 | loxF8L | 4 | CTGTGTCA/TGACACAG | 7 |
| chr5 | 1698206 | ATTTCTCTGTGGA-TGACACAG-TGCATAGACTTAC | loxF8L | 3 | loxF8L | 4 | CTGTGTCA/TGACACAG | 7 |
| chr1 | 222107495 | CTTAGTCTGTGAA-TGACACAG-GACACATTCTTGG | loxF8L | 4 | loxF8R | 3 | CTGTGTCA/TGACACAG | 7 |
| --- | | | | | | | | 0 |
| chr19 | 30786178 | ATACATCTGTGTT-GCTTTAAG-TCATGAGATTTGT | loxF8L | 3 | loxF8L | 4 | CTTAAAGC/GCTTTAAG | 7 |
| chr16 | 13548052 | ATAAATCCATGCA-CTTAAAGC-CAGCCAATTTTTG | loxF8L | 3 | loxF8R | 4 | CTTAAAGC/GCTTTAAG | 7 |
| --- | | | | | | | | 0 |
| chr15 | 34781418 | CAAAGATCAGGTG-GCTGTAAG-TGTACAGCTTTAT | loxF8R | 4 | loxF8L | 3 | CTTACAGC/GCTGTAAG | 7 |
| chr20 | 22671535 | ATAAATCTATACA-CTTACAGC-CAACTAATTTTAG | loxF8L | 3 | loxF8R | 4 | CTTACAGC/GCTGTAAG | 7 |
| chr10 | 45320385 | CAAAGATTAGATG-GCTGTAAG-TGTGCAGATTTAT | loxF8R | 4 | loxF8L | 3 | CTTACAGC/GCTGTAAG | 7 |
| --- | | | | | | | | 0 |
| chr6 | 142473515 | ACAAAACTGACGA-CTTAGAAC-AGCACAAATTTAT | loxF8L | 4 | loxF8L | 3 | CTTAGAAC/GTTCTAAG | 7 |
| chrX | 107103552 | ATAAATTTCTGTT-GTTCTAAG-CCACCAGATTTAT | loxF8L | 4 | loxF8L | 3 | CTTAGAAC/GTTCTAAG | 7 |
| --- | | | | | | | | 0 |
| chr6 | 125047153 | ATATATGTTTGGA-CTTAGATA-CATAGAAACTGAG | loxF8L | 3 | loxF8R | 4 | CTTAGATA/TATCTAAG | 7 |
| chr3 | 99125239 | GAAAAATTGTGTG-CTTAGATA-TACAAAGGTTTAC | loxF8R | 3 | loxF8L | 4 | CTTAGATA/TATCTAAG | 7 |
| --- | | | | | | | | 0 |
| chr5 | 4520312 | ATAAATAATTTGA-TCACTAAG-ACCATATATTTAT | loxF8L | 4 | loxF8L | 3 | CTTAGTGA/TCACTAAG | 7 |
| chr11 | 27496246 | CTAAGATTGAGAA-TCACTAAG-TCAACACATTTTG | loxF8R | 3 | loxF8L | 4 | CTTAGTGA/TCACTAAG | 7 |
| --- | | | | | | | | 0 |
| chr13 | 56433428 | ATATAACAGTGCA-TAATGAAG-TATACATATTTAT | loxF8L | 4 | loxF8L | 3 | CTTCATTA/TAATGAAG | 7 |
| chr7 | 64178577 | CTATAATTGTAAG-TAATGAAG-CACACAAAATAAG | loxF8R | 4 | loxF8R | 3 | CTTCATTA/TAATGAAG | 7 |
| --- | | | | | | | | 0 |
| chr12 | 16351381 | CTAACATACTGTG-CTTTCAAG-ACTAGAGACTTAT | loxF8R | 3 | loxF8L | 4 | CTTGAAAG/CTTTCAAG | 7 |
| chr11 | 36895489 | ATAAGTTTGTTTT-CTTTCAAG-CCCACTGAATTAT | loxF8R | 4 | loxF8L | 3 | CTTGAAAG/CTTTCAAG | 7 |
| --- | | | | | | | | 0 |
| chr4 | 69389233 | ATAAAACAGGGTA-CTTGCAAG-GTCATAATCTTAG | loxF8L | 4 | loxF8R | 3 | CTTGCAAG/CTTGCAAG | 7 |
| --- | | | | | | | | 0 |
| chrX | 70795679 | CAAAGAGTCTCTG-CTTGGTAA-TCCAGAGAATTTT | loxF8R | 4 | loxF8L | 3 | CTTGGTAA/TTACCAAG | 7 |
| chr14 | 23217614 | ATAAGAGTCTCTG-CTTGGTAA-TCCATAGAATTCT | loxF8R | 4 | loxF8L | 3 | CTTGGTAA/TTACCAAG | 7 |
| chr15 | 85722286 | GTAACTCAGCGGA-TTACCAAG-TACCCAGTTTTAT | loxF8L | 4 | loxF8L | 3 | CTTGGTAA/TTACCAAG | 7 |
| chr9 | 120680415 | GGAATTCTGTGGA-TTACCAAG-CAGAAACTCTTAT | loxF8L | 3 | loxF8R | 4 | CTTGGTAA/TTACCAAG | 7 |

| | | | | | | | | |
|---|---|---|---|---|---|---|---|---|
| --- | | | | | | | | 0 |
| chr12 | 92720091 | ATAAATCAGGTGA-CTTTAAAA-AACAGAAATTTAT | loxF8L | 3 | loxF8L | 4 | CTTTAAAA/TTTTAAAG | 7 |
| chr5 | 161343451 | GTCAGATTGTGGG-TTTTAAAG-ACCATATATCTAT | loxF8R | 3 | loxF8L | 4 | CTTTAAAA/TTTTAAAG | 7 |
| --- | | | | | | | | 0 |
| chr2 | 124721806 | CACAGATTGAGTG-GCTTAAAG-AACAAAATTTTAT | loxF8R | 3 | loxF8R | 4 | CTTTAAGC/GCTTAAAG | 7 |
| chr1 | 186085252 | CAGAGACTGTGTG-GCTTAAAG-AACAGAAATTTAT | loxF8R | 3 | loxF8L | 4 | CTTTAAGC/GCTTAAAG | 7 |
| chr20 | 18440185 | ATAAATTTCTGTT-CTTTAAGC-CACACAGTCTGTG | loxF8L | 4 | loxF8R | 3 | CTTTAAGC/GCTTAAAG | 7 |
| --- | | | | | | | | 0 |
| chr17 | 14469593 | CTTAGATTGGGTA-CTTTACAA-ACAAGAGATTTGT | loxF8R | 3 | loxF8L | 4 | CTTTACAA/TTGTAAAG | 7 |
| chr6 | 77190321 | TTAAATCAGTGCA-CTTTACAA-TACAGATATTGAT | loxF8L | 3 | loxF8L | 4 | CTTTACAA/TTGTAAAG | 7 |
| --- | | | | | | | | 0 |
| chrX | 910064 | CTGAGACTGTGGT-TTATAAAG-AGCAGAGATTTAT | loxF8R | 4 | loxF8L | 3 | CTTTATAA/TTATAAAG | 7 |
| chrY | 910064 | CTGAGACTGTGGT-TTATAAAG-AGCAGAGATTTAT | loxF8R | 4 | loxF8L | 3 | CTTTATAA/TTATAAAG | 7 |
| --- | | | | | | | | 0 |
| chr8 | 75073118 | AAAAATCTCTGTA-TATGAAAG-TCCAGAGGTATAC | loxF8L | 3 | loxF8L | 4 | CTTTCATA/TATGAAAG | 7 |
| chr20 | 55576387 | ATATATATGAGGA-CTTTCATA-TATATAGATTTAA | loxF8L | 3 | loxF8L | 4 | CTTTCATA/TATGAAAG | 7 |
| --- | | | | | | | | 0 |
| chrX | 18929047 | AGAATTCTGTTGC-CTTTGGAA-GCCACTGAGTTAT | loxF8L | 4 | loxF8L | 3 | CTTTGGAA/TTCCAAAG | 7 |
| chr12 | 66201913 | ATTAATCTAAGTA-CTTTGGAA-CACAGCATCTTAA | loxF8L | 4 | loxF8R | 3 | CTTTGGAA/TTCCAAAG | 7 |
| --- | | | | | | | | 0 |
| chr17 | 70055015 | AAATATTTCTGGA-GTTAAAAG-TCCTGAGATGTAT | loxF8L | 4 | loxF8L | 3 | CTTTTAAC/GTTAAAAG | 7 |
| chr21 | 29989906 | GGAAATTTGTGGA-GTTAAAAG-AAGCCAATCTCAG | loxF8L | 3 | loxF8R | 4 | CTTTTAAC/GTTAAAAG | 7 |
| --- | | | | | | | | 0 |
| chr11 | 124437094 | ATACATTTGTAGG-CTTTTCTC-TCCATTTATTTAT | loxF8L | 4 | loxF8L | 3 | CTTTTCTC/GAGAAAAG | 7 |
| chr5 | 106203261 | GAAAGATTGTTAG-GAGAAAAG-CCTACAAATTTAT | loxF8R | 4 | loxF8L | 3 | CTTTTCTC/GAGAAAAG | 7 |
| --- | | | | | | | | 0 |
| chr3 | 154744867 | AAAACTCTGAGGA-GAAAACAA-GAGTAAATCTTAG | loxF8L | 3 | loxF8R | 4 | GAAAACAA/TTGTTTTC | 7 |
| chr5 | 99826994 | ACAAATATGTGAA-TTGTTTTC-TGCATATATTAAT | loxF8L | 3 | loxF8L | 4 | GAAAACAA/TTGTTTTC | 7 |
| --- | | | | | | | | 0 |
| chr4 | 148730451 | AACAAGCTGTGAA-TTCTTTTC-TACAGAATCTTAT | loxF8L | 4 | loxF8R | 3 | GAAAAGAA/TTCTTTTC | 7 |
| chr18 | 33936986 | ATAAATATATAGA-GAAAAGAA-CCCACAAATTTTG | loxF8L | 3 | loxF8L | 4 | GAAAAGAA/TTCTTTTC | 7 |
| chr2 | 116638655 | CAAAGATTTTATG-GAAAAGAA-CTCAAAAGCATAG | loxF8R | 3 | loxF8R | 4 | GAAAAGAA/TTCTTTTC | 7 |
| --- | | | | | | | | 0 |
| chr20 | 64168319 | ATACAAGTGTGGA-GTCTTTTC-TCCCCAGGGTTAA | loxF8L | 3 | loxF8L | 4 | GAAAAGAC/GTCTTTTC | 7 |
| chr7 | 91766025 | CAAAGATTTTATG-GAAAAGAC-CTCAAAATCACAG | loxF8R | 3 | loxF8R | 4 | GAAAAGAC/GTCTTTTC | 7 |
| --- | | | | | | | | 0 |
| chr22 | 25691331 | ATGCATCTGTGCA-GAAAATAA-ACCACAGATCCTT | loxF8L | 3 | loxF8L | 4 | GAAAATAA/TTATTTTC | 7 |
| chr6 | 80954225 | ATAAAATTTTGAG-GAAAATAA-CCCACAGAATTCT | loxF8R | 4 | loxF8L | 3 | GAAAATAA/TTATTTTC | 7 |
| --- | | | | | | | | 0 |
| chr12 | 46733301 | ATAAGAGCTTGTG-TCTGTTTC-CCCACAATTTCAG | loxF8R | 4 | loxF8R | 3 | GAAACAGA/TCTGTTTC | 7 |
| chr1 | 29795168 | CTGAAATTGTGGA-GAAACAGA-CACAAACTCTTAT | loxF8L | 4 | loxF8R | 3 | GAAACAGA/TCTGTTTC | 7 |
| --- | | | | | | | | 0 |
| chr11 | 44751969 | CTAATATTTTGCG-TATGTTTC-TTCCCAGTTTTTT | loxF8R | 3 | loxF8L | 4 | GAAACATA/TATGTTTC | 7 |
| chr18 | 32863341 | GTCCATCTATGGA-TATGTTTC-TACACACATATAT | loxF8L | 4 | loxF8L | 3 | GAAACATA/TATGTTTC | 7 |
| --- | | | | | | | | 0 |
| chr11 | 130983543 | ACAAATCTGAAGA-GAAAGAAA-TCAGCAAATTTAG | loxF8L | 3 | loxF8L | 4 | GAAAGAAA/TTTCTTTC | 7 |
| chr7 | 19876678 | TCAAGCTTCTGTG-TTTCTTTC-TTTAAAGATTTAT | loxF8R | 4 | loxF8L | 3 | GAAAGAAA/TTTCTTTC | 7 |
| --- | | | | | | | | 0 |
| chr5 | 72958938 | ATACTTCTCTGAA-TATCTTTC-TACACATTTTTAT | loxF8L | 4 | loxF8L | 3 | GAAAGATA/TATCTTTC | 7 |
| chr12 | 108064368 | CTAAATATGTGGG-GAAAGATA-ACAACAGAATCAT | loxF8L | 3 | loxF8L | 4 | GAAAGATA/TATCTTTC | 7 |
| --- | | | | | | | | 0 |
| chrY | 19835621 | AGAACTCTGTGAA-GAAAGTCA-TTGGTAGATTTAT | loxF8L | 3 | loxF8L | 4 | GAAAGTCA/TGACTTTC | 7 |
| chr5 | 129148703 | CTAATTCTGTGAA-GAAAGTCA-ACCATAGCTTGAT | loxF8L | 3 | loxF8L | 4 | GAAAGTCA/TGACTTTC | 7 |
| --- | | | | | | | | 0 |
| chr8 | 50235546 | CTGTGAGTGAGTG-GAAATCAA-TCAACACATTTTT | loxF8R | 4 | loxF8L | 3 | GAAATCAA/TTGATTTC | 7 |
| chrX | 45702834 | TTAAATTTGTGCA-TTGATTTC-TGCATTGATTCAT | loxF8L | 3 | loxF8L | 4 | GAAATCAA/TTGATTTC | 7 |
| --- | | | | | | | | 0 |
| chr9 | 118120787 | ACAAAGCTCTGAA-GAAATGGA-CATACAACTTTAG | loxF8L | 4 | loxF8R | 3 | GAAATGGA/TCCATTTC | 7 |
| chr2 | 62874165 | ATAAATCTGGATA-GAAATGGA-CCAAAAGATTAAT | loxF8L | 3 | loxF8L | 4 | GAAATGGA/TCCATTTC | 7 |
| --- | | | | | | | | 0 |
| chr10 | 74617360 | CTAAGAGTTTTTG-TTTTGTTC-TTAAGAGATTTTT | loxF8R | 3 | loxF8L | 4 | GAACAAAA/TTTTGTTC | 7 |
| chr13 | 40288745 | AAAAATCTGTGAT-GAACAAAA-TACCCAGTTTTAA | loxF8L | 3 | loxF8L | 4 | GAACAAAA/TTTTGTTC | 7 |
| --- | | | | | | | | 0 |
| chr1 | 9134264 | AAACATGTGTGGA-GTTTGTTC-TGCAGAGAGTTAA | loxF8L | 3 | loxF8L | 4 | GAACAAAC/GTTTGTTC | 7 |
| chr7 | 114672074 | CTAACATTGTTTA-GTTTGTTC-AACATTATCTTCG | loxF8R | 3 | loxF8R | 4 | GAACAAAC/GTTTGTTC | 7 |
| --- | | | | | | | | 0 |
| chr7 | 89244838 | ATCAATCTGAGAA-TTAAGTTC-TCAGCAGAGTTTT | loxF8L | 3 | loxF8L | 4 | GAACTTAA/TTAAGTTC | 7 |
| chr1 | 156978484 | ATTATTCTGTAGA-TTAAGTTC-TCCTCACTTTTAA | loxF8L | 3 | loxF8L | 4 | GAACTTAA/TTAAGTTC | 7 |
| --- | | | | | | | | 0 |
| chr7 | 92803980 | ATTAATCTTAAGA-GTCACTTC-TCCACTGGTTTGT | loxF8L | 4 | loxF8L | 3 | GAAGTGAC/GTCACTTC | 7 |
| chr17 | 68098402 | GTAACTATTTGGA-GAAGTGAC-TTCACTGATTGAT | loxF8L | 4 | loxF8L | 3 | GAAGTGAC/GTCACTTC | 7 |
| --- | | | | | | | | 0 |
| chr18 | 39133724 | ACAAATCCTTGGA-TGTTATTC-TTCAAACATTTCT | loxF8L | 3 | loxF8L | 4 | GAATAACA/TGTTATTC | 7 |
| chr1 | 241023050 | GTAAAATTGTGTA-GAATAACA-ACTACAGATATAG | loxF8R | 3 | loxF8L | 4 | GAATAACA/TGTTATTC | 7 |
| --- | | | | | | | | 0 |
| chr3 | 54763913 | ATAAAGTTAGGGA-GGATATTC-TCCAAAGAATTAA | loxF8L | 4 | loxF8L | 3 | GAATATCC/GGATATTC | 7 |
| chr18 | 57530423 | CTATGATTGTTGT-GGATATTC-CACACACACATAG | loxF8R | 4 | loxF8R | 3 | GAATATCC/GGATATTC | 7 |
| --- | | | | | | | | 0 |
| chr2 | 212960250 | ATATGATTGTAGG-GAATCAAA-ACCACAGATCTGT | loxF8R | 4 | loxF8L | 3 | GAATCAAA/TTTGATTC | 7 |
| chr11 | 22283300 | ATAAATCTCAAGA-GAATCAAA-TTCACAGAGTGAA | loxF8L | 3 | loxF8L | 4 | GAATCAAA/TTTGATTC | 7 |
| --- | | | | | | | | 0 |
| chr21 | 9649765 | CTAAGGTTATGTT-GAATCTAA-CCCAAACTCTTAT | loxF8R | 3 | loxF8R | 4 | GAATCTAA/TTAGATTC | 7 |
| chr4 | 189706747 | ATAAGAGTTTGGG-TTAGATTC-AACATAACCTTAG | loxF8R | 4 | loxF8R | 3 | GAATCTAA/TTAGATTC | 7 |
| --- | | | | | | | | 0 |
| chr1 | 95820514 | CTAGATCTGTGAA-GAATGTCA-TCAGTAGTTTTAT | loxF8L | 3 | loxF8L | 4 | GAATGTCA/TGACATTC | 7 |
| chr2 | 17023735 | CTAGTTCTGTGGA-GAATGTCA-TTGACAGTTTGAT | loxF8L | 3 | loxF8L | 4 | GAATGTCA/TGACATTC | 7 |
| chr8 | 14064226 | CTAATTCTCTGAA-GAATGTCA-TTGACAGATTGAT | loxF8L | 4 | loxF8L | 3 | GAATGTCA/TGACATTC | 7 |

| | | | | | | | | |
|---|---|---|---|---|---|---|---|---|
| chr1 | 168478951 | ATTAAACTATGGT-TGACATTC-TTCACAGAATTAC | loxF8L | 4 | loxF8L | 3 | GAATGTCA/TGACATTC | 7 |
| --- | | | | | | | | 0 |
| chrX | 124572262 | AAAAATCTATCAA-TGTAATTC-ACCACTGAATTAT | loxF8L | 4 | loxF8L | 3 | GAATTACA/TGTAATTC | 7 |
| chr9 | 102851355 | AAATATTTGTGCA-TGTAATTC-TCCATATTTTTAT | loxF8L | 4 | loxF8L | 3 | GAATTACA/TGTAATTC | 7 |
| --- | | | | | | | | 0 |
| chr9 | 76546125 | ATAAATTTATGGG-TTTGTGTC-AACATAAATTTAT | loxF8L | 3 | loxF8L | 4 | GACACAAA/TTTGTGTC | 7 |
| chr9 | 7927729 | ATGAATTTGGGGA-GACACAAA-CATTCATTCTTAT | loxF8L | 3 | loxF8R | 4 | GACACAAA/TTTGTGTC | 7 |
| --- | | | | | | | | 0 |
| chr17 | 30432232 | ATAAAATTGTATT-GACACACA-GCCACAGCTATAT | loxF8R | 4 | loxF8L | 3 | GACACACA/TGTGTGTC | 7 |
| chr13 | 85377625 | CATATAATGTGTG-TGTGTGTC-TCACCAGAATTAT | loxF8R | 4 | loxF8L | 3 | GACACACA/TGTGTGTC | 7 |
| chr5 | 94463558 | ATATATATGGAGA-GACACACA-TACACAATCTGAT | loxF8L | 4 | loxF8R | 3 | GACACACA/TGTGTGTC | 7 |
| --- | | | | | | | | 0 |
| chr13 | 78316056 | GTGTGTTTGTGTG-TATGTGTC-TACACATATTTGT | loxF8R | 4 | loxF8L | 3 | GACACATA/TATGTGTC | 7 |
| chr12 | 72111852 | CTACATGTGTGAA-TATGTGTC-TACACAGAGTCAT | loxF8L | 4 | loxF8L | 3 | GACACATA/TATGTGTC | 7 |
| --- | | | | | | | | 0 |
| chr4 | 45025288 | ATATATCTATATA-TATCTGTC-TCTATAGATATAT | loxF8L | 4 | loxF8L | 3 | GACAGATA/TATCTGTC | 7 |
| chr13 | 92207719 | ATTACTCTGCGGT-TATCTGTC-TACTCATTCTTAG | loxF8L | 4 | loxF8R | 3 | GACAGATA/TATCTGTC | 7 |
| --- | | | | | | | | 0 |
| chr1 | 200383937 | AAAAGATTGTGTT-GACATTTA-TCCACACAATTTG | loxF8R | 3 | loxF8L | 4 | GACATTTA/TAAATGTC | 7 |
| chr10 | 112009667 | ATAAATATGTAAA-TAAATGTC-TACAAAATCTAAC | loxF8L | 3 | loxF8R | 4 | GACATTTA/TAAATGTC | 7 |
| chrX | 143679107 | GTAAATTTTTGGA-GACATTTA-CACACACACATAT | loxF8L | 3 | loxF8R | 4 | GACATTTA/TAAATGTC | 7 |
| --- | | | | | | | | 0 |
| chr1 | 102834719 | AGAAATGTGTGCA-GACCAGGC-TAGACAGAATTGT | loxF8L | 3 | loxF8L | 4 | GACCAGGC/GCCTGGTC | 7 |
| chr11 | 112928625 | ATGATTCTGTTTA-GCCTGGTC-TGCACACATTTCT | loxF8L | 4 | loxF8L | 3 | GACCAGGC/GCCTGGTC | 7 |
| chr12 | 111824137 | ATGATTCTGTCTA-GCCTGGTC-TGCACACATTTCT | loxF8L | 4 | loxF8L | 3 | GACCAGGC/GCCTGGTC | 7 |
| chr19 | 11859402 | ATGATTCTGTCTA-GCCTGGTC-TGCACACATTTCT | loxF8L | 4 | loxF8L | 3 | GACCAGGC/GCCTGGTC | 7 |
| chr20 | 25401458 | ATGATTCTGTCTA-GCCTGGTC-TGCACACATTTCT | loxF8L | 4 | loxF8L | 3 | GACCAGGC/GCCTGGTC | 7 |
| chr6 | 32565891 | ATGATTCTGTCTA-GCCTGGTC-TGCACACATTTCT | loxF8L | 4 | loxF8L | 3 | GACCAGGC/GCCTGGTC | 7 |
| --- | | | | | | | | 0 |
| chr21 | 38288479 | ATCAATTTGTAGA-GACTGACA-ACTCCACATTTAT | loxF8L | 3 | loxF8L | 4 | GACTGACA/TGTCAGTC | 7 |
| chr22 | 20620827 | CTCAGCTTGTTTG-GACTGACA-CACAGGCTCTTTG | loxF8R | 3 | loxF8R | 4 | GACTGACA/TGTCAGTC | 7 |
| --- | | | | | | | | 0 |
| chr6 | 134407341 | ACAAAATAGTGTG-TCTCAGTC-TTCATAGTTTTAT | loxF8R | 4 | loxF8L | 3 | GACTGAGA/TCTCAGTC | 7 |
| chr2 | 29233102 | ATATCTCTCTGGA-TCTCAGTC-TGCTCATCTTTAT | loxF8L | 3 | loxF8L | 4 | GACTGAGA/TCTCAGTC | 7 |
| --- | | | | | | | | 0 |
| chr9 | 14414866 | AAAAATCCTTGTA-GACTTCCC-CACACACTGTCAG | loxF8L | 4 | loxF8R | 3 | GACTTCCC/GGGAAGTC | 7 |
| chrX | 136391241 | CTAGGCTTTTGTG-GGGAAGTC-CTCAGAGATTTTT | loxF8R | 3 | loxF8L | 4 | GACTTCCC/GGGAAGTC | 7 |
| --- | | | | | | | | 0 |
| chr12 | 11648259 | AAAATATTGTGTG-TACAAGTC-TCAACACATTTGA | loxF8R | 3 | loxF8L | 4 | GACTTGTA/TACAAGTC | 7 |
| chr8 | 119443722 | CTAAGTTTCTATA-TACAAGTC-TACTCAGGTTTAT | loxF8R | 4 | loxF8L | 3 | GACTTGTA/TACAAGTC | 7 |
| --- | | | | | | | | 0 |
| chr3 | 105932015 | ATAAAACTGTAAA-GACTTTGC-TCAACACATCTAG | loxF8L | 3 | loxF8L | 4 | GACTTTGC/GCAAAGTC | 7 |
| chr8 | 24574717 | ATAAATATTCTGA-GACTTTGC-TCCTCAAATTAAT | loxF8L | 4 | loxF8L | 3 | GACTTTGC/GCAAAGTC | 7 |
| --- | | | | | | | | 0 |
| chrX | 36305657 | AAAAATATATAGA-GAGAAATA-TCCATCTATTTAT | loxF8L | 4 | loxF8L | 3 | GAGAAATA/TATTTCTC | 7 |
| chrX | 68134402 | ACAAATGTGTGGT-GAGAAATA-TCTACAAATGTGT | loxF8L | 3 | loxF8L | 4 | GAGAAATA/TATTTCTC | 7 |
| --- | | | | | | | | 0 |
| chr2 | 197944424 | CTCACATTCTCTG-GAGAATAA-ACCCCAGATTTCT | loxF8R | 4 | loxF8L | 3 | GAGAATAA/TTATTCTC | 7 |
| chr2 | 225020262 | TTGACATTTTGTG-GAGAATAA-TACACATATTAAT | loxF8R | 4 | loxF8L | 3 | GAGAATAA/TTATTCTC | 7 |
| --- | | | | | | | | 0 |
| chr2 | 50549923 | ATATACATGTAGA-GAGAGACA-TGCACACATATAT | loxF8L | 4 | loxF8L | 3 | GAGAGACA/TGTCTCTC | 7 |
| chr14 | 94285460 | ATATATATATGTA-TGTCTCTC-TCCATATATATAT | loxF8L | 4 | loxF8L | 3 | GAGAGACA/TGTCTCTC | 7 |
| --- | | | | | | | | 0 |
| chr2 | 146966149 | ATATGTATGTGGA-GAGAGAGA-GAGAAAATCTTAT | loxF8L | 3 | loxF8R | 4 | GAGAGAGA/TCTCTCTC | 7 |
| chr8 | 24659892 | ATATATATATGGA-GAGAGAGA-CACACACACAGAG | loxF8L | 3 | loxF8R | 4 | GAGAGAGA/TCTCTCTC | 7 |
| --- | | | | | | | | 0 |
| chr9 | 226965 | ATAAATATATGCA-TAACTCTC-TGCAGAGATAAAT | loxF8L | 3 | loxF8L | 4 | GAGAGTTA/TAACTCTC | 7 |
| chr15 | 79380353 | ATATATATATAGA-GAGAGTTA-TACACATATATAT | loxF8L | 4 | loxF8L | 3 | GAGAGTTA/TAACTCTC | 7 |
| --- | | | | | | | | 0 |
| chr4 | 127019003 | CCTAGATTATGTT-TCTATCTC-CAAACAGATTTAT | loxF8R | 4 | loxF8L | 3 | GAGATAGA/TCTATCTC | 7 |
| chr7 | 15896637 | ATATATATATAGA-GAGATAGA-TCGACAGATAGAT | loxF8L | 4 | loxF8L | 3 | GAGATAGA/TCTATCTC | 7 |
| chr4 | 166400141 | ATATATATATGGA-GAGATAGA-TACACACACATAT | loxF8L | 3 | loxF8L | 4 | GAGATAGA/TCTATCTC | 7 |
| chr4 | 163886182 | ATGAATCTATAGA-TCTATCTC-TCAATATATATAT | loxF8L | 3 | loxF8L | 4 | GAGATAGA/TCTATCTC | 7 |
| --- | | | | | | | | 0 |
| chr1 | 2673769 | ATCTGATTGTCTG-GAGCATCA-CACACAACCACAG | loxF8R | 4 | loxF8R | 3 | GAGCATCA/TGATGCTC | 7 |
| chr1 | 2659359 | ATCTGATTGTCTG-GAGCATCA-CACACAACCACAG | loxF8R | 4 | loxF8R | 3 | GAGCATCA/TGATGCTC | 7 |
| chr1 | 2668591 | ATCTGATTGTCTG-GAGCATCA-CACACAACCACAG | loxF8R | 4 | loxF8R | 3 | GAGCATCA/TGATGCTC | 7 |
| chr1 | 2663975 | ATCTGATTGTCTG-GAGCATCA-CACACAACCACAG | loxF8R | 4 | loxF8R | 3 | GAGCATCA/TGATGCTC | 7 |
| --- | | | | | | | | 0 |
| chr1 | 248488005 | ATGAGAATGTGAG-GCTCGCTC-CACGCAATCACAG | loxF8R | 4 | loxF8R | 3 | GAGCGAGC/GCTCGCTC | 7 |
| chr1 | 248560040 | CTGTGATTGCGTG-GAGCGAGC-CTCACATTCTCAT | loxF8R | 3 | loxF8R | 4 | GAGCGAGC/GCTCGCTC | 7 |
| --- | | | | | | | | 0 |
| chr2 | 185877077 | CTGAGATTGTTGG-TCTAGCTC-TGCCCAGCTTTGT | loxF8R | 3 | loxF8L | 4 | GAGCTAGA/TCTAGCTC | 7 |
| chr3 | 78866554 | CTAATATTATATA-GAGCTAGA-ACCAGAGATATAT | loxF8R | 4 | loxF8L | 3 | GAGCTAGA/TCTAGCTC | 7 |
| --- | | | | | | | | 0 |
| chr13 | 59150670 | ACAAGTTTGTGGA-GAGGAAAA-TCCAATGAATTCT | loxF8L | 3 | loxF8L | 4 | GAGGAAAA/TTTTCCTC | 7 |
| chr18 | 66368495 | CTGTGGTTGTGTG-TTTTCCTC-TAGACATGCTTAG | loxF8R | 3 | loxF8R | 4 | GAGGAAAA/TTTTCCTC | 7 |
| --- | | | | | | | | 0 |
| chr6 | 143281733 | ATAAATTTCTAGA-GAGGGAGA-TACACAGTCTTTA | loxF8L | 3 | loxF8R | 4 | GAGGGAGA/TCTCCCTC | 7 |
| chr12 | 12007134 | CACAGATTGTGGG-GAGGGAGA-CACATAATGTGTG | loxF8R | 3 | loxF8R | 4 | GAGGGAGA/TCTCCCTC | 7 |
| --- | | | | | | | | 0 |
| chr14 | 18314892 | GTAAGATTGTAGT-GAGGGTGA-ATAACAGATTTAT | loxF8R | 4 | loxF8L | 3 | GAGGGTGA/TCACCCTC | 7 |
| chr15 | 19861563 | GTAAGATTGTAGT-GAGGGTGA-ATAACAGATTTAT | loxF8R | 4 | loxF8L | 3 | GAGGGTGA/TCACCCTC | 7 |
| chr18 | 15349276 | ATAAATCTGTTAT-TCACCCTC-ACTACAATCTTAC | loxF8L | 3 | loxF8R | 4 | GAGGGTGA/TCACCCTC | 7 |
| chr21 | 13018862 | GTAAGATTGTAGT-GAGGGTGA-ATAACAGATTTAT | loxF8R | 4 | loxF8L | 3 | GAGGGTGA/TCACCCTC | 7 |
| chr22 | 15245699 | GTAAGATTGTAGT-GAGGGTGA-ATAACAGATTTAT | loxF8R | 4 | loxF8L | 3 | GAGGGTGA/TCACCCTC | 7 |
| chr2 | 132180556 | ATAAATCTGTTAT-TCACCCTC-ACTACAATCTTAC | loxF8L | 3 | loxF8R | 4 | GAGGGTGA/TCACCCTC | 7 |

| | | | | | | | | |
|---|---|---|---|---|---|---|---|---|
| chr9 | 40787044 | ATAAATCTGTTAT-TCACCCTC-ACTACAATCTTAC | loxF8L | 3 | loxF8R | 4 | GAGGGTGA/TCACCCTC | 7 |
| chr9 | 64907845 | ATAAATCTGTTAT-TCACCCTC-ACTACAATCTTAC | loxF8L | 3 | loxF8R | 4 | GAGGGTGA/TCACCCTC | 7 |
| --- | | | | | | | | 0 |
| chr9 | 37748305 | AGAAAGCTGTGAA-GAGGTAAA-AACACAACCTATG | loxF8L | 3 | loxF8R | 4 | GAGGTAAA/TTTACCTC | 7 |
| chr5 | 25433057 | ATAGATAGGTAGA-GAGGTAAA-TACACAATTTTAA | loxF8L | 4 | loxF8R | 3 | GAGGTAAA/TTTACCTC | 7 |
| --- | | | | | | | | 0 |
| chr8 | 120858355 | ATAAATATATGTA-GATACACA-CACACATATATAT | loxF8L | 3 | loxF8L | 4 | GATACACA/TGTGTATC | 7 |
| chr8 | 96503510 | ATAGAACTGTGTA-GATACACA-TACACAAACATAC | loxF8L | 3 | loxF8R | 4 | GATACACA/TGTGTATC | 7 |
| chrY | 9174616 | CTATATATGTAGA-GATACACA-CACACAAACAAAG | loxF8L | 4 | loxF8R | 3 | GATACACA/TGTGTATC | 7 |
| --- | | | | | | | | 0 |
| chr20 | 3638029 | AGAAATACGAGGA-GATAGAGC-CACACCATTTTAT | loxF8L | 4 | loxF8R | 3 | GATAGAGC/GCTCTATC | 7 |
| chr2 | 155736607 | TTAAAATTGTTGA-GATAGAGC-TCCAAGAATTTAT | loxF8L | 4 | loxF8L | 3 | GATAGAGC/GCTCTATC | 7 |
| --- | | | | | | | | 0 |
| chr8 | 97685152 | CTATTATGGTATG-TGGATATC-TCCACATATCTTT | loxF8R | 4 | loxF8L | 3 | GATATCCA/TGGATATC | 7 |
| chr1 | 84072857 | ATTAATATGTAGA-GATATCCA-TCCTTGGATCTAT | loxF8L | 3 | loxF8L | 4 | GATATCCA/TGGATATC | 7 |
| --- | | | | | | | | 0 |
| chr11 | 88343670 | AGATCTCTGTGGA-GATCTGAC-TACATACATGTAT | loxF8L | 3 | loxF8L | 4 | GATCTGAC/GTCAGATC | 7 |
| chr11 | 112803488 | CAATGATTGTTTA-GATCTGAC-CACAAAAGCATAG | loxF8R | 4 | loxF8R | 3 | GATCTGAC/GTCAGATC | 7 |
| --- | | | | | | | | 0 |
| chr3 | 175168118 | ATAAGTCAGTTGT-GATGAGAA-ACCACAGATATGT | loxF8L | 4 | loxF8L | 3 | GATGAGAA/TTCTCATC | 7 |
| chr2 | 66990538 | GTAAGAATGAGTG-GATGAGAA-ACAACAGCTTTAC | loxF8R | 3 | loxF8L | 4 | GATGAGAA/TTCTCATC | 7 |
| --- | | | | | | | | 0 |
| chr6 | 48304378 | ATAAATCAGGAGA-GATGAGGA-TACTAAGACTTAT | loxF8L | 3 | loxF8L | 4 | GATGAGGA/TCCTCATC | 7 |
| chr13 | 27518442 | GAAAATCTGAGGA-TCCTCATC-CACACAAAACTAT | loxF8L | 3 | loxF8R | 4 | GATGAGGA/TCCTCATC | 7 |
| --- | | | | | | | | 0 |
| chr10 | 117637274 | AAAAATATGTGTA-TGTACATC-TACATACATTTGT | loxF8L | 3 | loxF8L | 4 | GATGTACA/TGTACATC | 7 |
| chr4 | 4160591 | ATAAAGCAGTGGG-TGTACATC-ACCCCAGTGTTAT | loxF8L | 3 | loxF8L | 4 | GATGTACA/TGTACATC | 7 |
| --- | | | | | | | | 0 |
| chr17 | 18721959 | CAAGGATGGTGTG-GGGTAATC-CACAAAATGCTTG | loxF8R | 3 | loxF8R | 4 | GATTACCC/GGGTAATC | 7 |
| chr17 | 15643059 | CAAGCATTTTGTG-GATTACCC-CACACCATCCTTG | loxF8R | 4 | loxF8R | 3 | GATTACCC/GGGTAATC | 7 |
| --- | | | | | | | | 0 |
| chr7 | 69161294 | AAAATTCTGTGGT-TTATAATC-CACCCAGTCTATG | loxF8L | 3 | loxF8R | 4 | GATTATAA/TTATAATC | 7 |
| chr4 | 96423726 | ATATTTCTGTTGC-TTATAATC-CACCCAATCTGTG | loxF8L | 4 | loxF8R | 3 | GATTATAA/TTATAATC | 7 |
| --- | | | | | | | | 0 |
| chr6 | 120932614 | ATAAACCTCTGAA-TTAAAATC-TCCATAAATACAT | loxF8L | 3 | loxF8L | 4 | GATTTTAA/TTAAAATC | 7 |
| chr5 | 108377001 | CTACATCTGTTCA-GATTTTAA-TTCACATATATAT | loxF8L | 4 | loxF8L | 3 | GATTTTAA/TTAAAATC | 7 |
| --- | | | | | | | | 0 |
| chr9 | 83144931 | ACAAATTTGTATA-GATTTTTA-CACAAAATATTAA | loxF8L | 4 | loxF8R | 3 | GATTTTTA/TAAAAATC | 7 |
| chrX | 138433031 | ATAAATATGTGAT-TAAAAATC-CATGCAAATTTAG | loxF8L | 3 | loxF8R | 4 | GATTTTTA/TAAAAATC | 7 |
| chr3 | 17148107 | CAAAGAGTGCATG-TAAAAATC-AGCACAATCTCAG | loxF8R | 4 | loxF8R | 3 | GATTTTTA/TAAAAATC | 7 |
| chr19 | 56356964 | CTAAGATTACCTT-GATTTTTA-TCCTCATATTTAA | loxF8R | 4 | loxF8L | 3 | GATTTTTA/TAAAAATC | 7 |
| --- | | | | | | | | 0 |
| chr4 | 136921956 | ATGAGTCTATGGA-TAATTTGC-CATAAAATGTAAG | loxF8L | 3 | loxF8R | 4 | GCAAATTA/TAATTTGC | 7 |
| chr12 | 41396474 | CTATTATTGTGTA-GCAAATTA-CCCAAAATTGTAG | loxF8R | 3 | loxF8R | 4 | GCAAATTA/TAATTTGC | 7 |
| --- | | | | | | | | 0 |
| chr1 | 186021597 | ATAAATGTATGAA-TGTGTTGC-TTAACATGTTTAT | loxF8L | 3 | loxF8L | 4 | GCAACACA/TGTGTTGC | 7 |
| chr8 | 16523679 | CTAGCTTTGTGTA-GCAACACA-CAAATAATTTTAG | loxF8R | 4 | loxF8R | 3 | GCAACACA/TGTGTTGC | 7 |
| --- | | | | | | | | 0 |
| chr7 | 14984432 | GGATATCTGTGAA-GCAAGGGA-TTCACAGAGTTCT | loxF8L | 4 | loxF8L | 3 | GCAAGGGA/TCCCTTGC | 7 |
| chr18 | 5932522 | ATAAACCTGGGTT-TCCCTTGC-TACACACAGTTAT | loxF8L | 4 | loxF8L | 3 | GCAAGGGA/TCCCTTGC | 7 |
| --- | | | | | | | | 0 |
| chr15 | 95943928 | ATAATTGTGGTGA-GTGATTGC-TCCAGAGACTTAA | loxF8L | 4 | loxF8L | 3 | GCAATCAC/GTGATTGC | 7 |
| chr6 | 55396186 | TTAAGATTGTGGA-GTGATTGC-TAGACAAATTTAA | loxF8R | 3 | loxF8L | 4 | GCAATCAC/GTGATTGC | 7 |
| --- | | | | | | | | 0 |
| chr1 | 147871187 | CTAAGAGTTCCTG-GCACCTCA-TCAACAGAATTCT | loxF8R | 4 | loxF8L | 3 | GCACCTCA/TGAGGTGC | 7 |
| chr6 | 47765402 | ATAAATCTCTCCA-TGAGGTGC-TACTTAATCTTTG | loxF8L | 3 | loxF8R | 4 | GCACCTCA/TGAGGTGC | 7 |
| --- | | | | | | | | 0 |
| chr7 | 30888500 | CTTTGATTGTGTC-TCTTCTGC-TCCTCAGAGGTGT | loxF8R | 3 | loxF8L | 4 | GCAGAAGA/TCTTCTGC | 7 |
| chr6 | 49033475 | CTAAATGTTTGTG-TCTTCTGC-TCCCCCAATTTAT | loxF8R | 4 | loxF8L | 3 | GCAGAAGA/TCTTCTGC | 7 |
| --- | | | | | | | | 0 |
| chr10 | 118247851 | ATAAATCTCTGTT-GTTCATGC-CACCCAGTCTGTG | loxF8L | 3 | loxF8R | 4 | GCATGAAC/GTTCATGC | 7 |
| chr12 | 66887074 | TTAAAACTGTGGT-GCATGAAC-CACTTGATCTTAT | loxF8L | 3 | loxF8R | 4 | GCATGAAC/GTTCATGC | 7 |
| --- | | | | | | | | 0 |
| chr13 | 66926853 | AAAAATCAGTGAA-GCATGACA-TAAAGAGTTTTAT | loxF8L | 3 | loxF8L | 4 | GCATGACA/TGTCATGC | 7 |
| chr9 | 106223995 | ATATATCTTTCAA-GCATGACA-TACAAAGAATTAT | loxF8L | 4 | loxF8L | 3 | GCATGACA/TGTCATGC | 7 |
| --- | | | | | | | | 0 |
| chr18 | 78199751 | AAAACATTCTGTG-TTAAATGC-CACACAGATTTTT | loxF8R | 4 | loxF8L | 3 | GCATTTAA/TTAAATGC | 7 |
| chr5 | 150971349 | TTGAGATTCTGTG-GCATTTAA-GCCACAATCTTGT | loxF8R | 3 | loxF8R | 4 | GCATTTAA/TTAAATGC | 7 |
| --- | | | | | | | | 0 |
| chr11 | 88058878 | CCAAAAATATGTG-GCCACTCA-CCTGCAGATTTAT | loxF8R | 4 | loxF8L | 3 | GCCACTCA/TGAGTGGC | 7 |
| chr17 | 28757250 | ATGTACCTGTGGG-GCCACTCA-CACACAATGCTAC | loxF8L | 4 | loxF8R | 3 | GCCACTCA/TGAGTGGC | 7 |
| --- | | | | | | | | 0 |
| chr7 | 14952286 | ATAAATCTCTGTT-GCCCAAGC-CACCCAATTTATG | loxF8L | 3 | loxF8R | 4 | GCCCAAGC/GCTTGGGC | 7 |
| chr4 | 63703725 | CAAAGAGTGAGTG-GCTTGGGC-AACAAAAATTTAT | loxF8R | 3 | loxF8L | 4 | GCCCAAGC/GCTTGGGC | 7 |
| --- | | | | | | | | 0 |
| chr13 | 105560181 | CTTAGACTGGGTG-GCCTAAAA-TGCAGGAATTTAT | loxF8R | 3 | loxF8L | 4 | GCCTAAAA/TTTTAGGC | 7 |
| chr21 | 32938081 | TACAAATTGTGTG-GCCTAAAA-CACAAAAACTTAT | loxF8R | 4 | loxF8R | 3 | GCCTAAAA/TTTTAGGC | 7 |
| --- | | | | | | | | 0 |
| chr10 | 104759317 | ATAAATATCTGTT-GTTTAGGC-CACCCAAACTGAG | loxF8L | 4 | loxF8R | 3 | GCCTAAAC/GTTTAGGC | 7 |
| chr4 | 17500682 | ATAAATCTCTGTT-GTTTAGGC-CACCCAGTCTGTG | loxF8L | 3 | loxF8R | 4 | GCCTAAAC/GTTTAGGC | 7 |
| chr16 | 82086766 | CACAAATCGTGTG-GCCTAAAC-AACACAAATTTAT | loxF8R | 4 | loxF8L | 3 | GCCTAAAC/GTTTAGGC | 7 |
| chr13 | 68787920 | ATAAATTTTTGTT-GTTTAGGC-CACACAGTCTATG | loxF8L | 4 | loxF8R | 3 | GCCTAAAC/GTTTAGGC | 7 |
| --- | | | | | | | | 0 |
| chr3 | 164763163 | CAAAGATTAATTG-GCCTGATA-TGCATATATTTAT | loxF8R | 4 | loxF8L | 3 | GCCTGATA/TATCAGGC | 7 |
| chr10 | 120518518 | TTAAATCTGTTCA-TATCAGGC-CAGAAAATGATAG | loxF8L | 3 | loxF8R | 4 | GCCTGATA/TATCAGGC | 7 |
| --- | | | | | | | | 0 |
| chr2 | 167991926 | AAAAAGCTGTAGA-GCCTTACA-TCTACAGCCTGAT | loxF8L | 3 | loxF8L | 4 | GCCTTACA/TGTAAGGC | 7 |

| | | | | | | | | |
|---|---|---|---|---|---|---|---|---|
| chr6 | 51459922 | TTACATCTGTGTA-GCCTTACA-TCAACTAGTTTAT | loxF8L | 3 | loxF8L | 4 | GCCTTACA/TGTAAGGC | 7 |
| --- | | | | | | | | 0 |
| chr16 | 19469778 | AAAAAGCTGTGAA-GCTAAAAC-AGAAGAATCTTAG | loxF8L | 3 | loxF8R | 4 | GCTAAAAC/GTTTTAGC | 7 |
| chr2 | 114426896 | CACAGACTGTGTG-GCTAAAAC-AACAGAGATTTGT | loxF8R | 3 | loxF8L | 4 | GCTAAAAC/GTTTTAGC | 7 |
| chr11 | 10333749 | ATAAATTTCTGGT-GTTTTAGC-CACCCAGTCTGTG | loxF8L | 3 | loxF8R | 4 | GCTAAAAC/GTTTTAGC | 7 |
| chr4 | 174745577 | CAAAGATTTTATG-GCTAAAAC-CTCAAAAGCATAG | loxF8R | 3 | loxF8R | 4 | GCTAAAAC/GTTTTAGC | 7 |
| --- | | | | | | | | 0 |
| chr2 | 220863139 | ATAATTCTGGGGT-TATTTAGC-TTCACTGATTCTT | loxF8L | 3 | loxF8L | 4 | GCTAAATA/TATTTAGC | 7 |
| chr14 | 80405990 | CTCAATTTGTGTG-TATTTAGC-TGCAAAATTTTAT | loxF8R | 3 | loxF8L | 4 | GCTAAATA/TATTTAGC | 7 |
| --- | | | | | | | | 0 |
| chr10 | 84070473 | CAAAGATTTTATG-GCTAAGAC-CACAAAAGCATGG | loxF8R | 3 | loxF8R | 4 | GCTAAGAC/GTCTTAGC | 7 |
| chr2 | 175336080 | CTGAGCTTTTGGG-GTCTTAGC-CATAAAATCTTTG | loxF8R | 4 | loxF8R | 3 | GCTAAGAC/GTCTTAGC | 7 |
| chr2 | 44467945 | CAAAGATTTTATG-GCTAAGAC-CTCAAAAACATAG | loxF8R | 3 | loxF8R | 4 | GCTAAGAC/GTCTTAGC | 7 |
| chrX | 87830765 | CTATGTTTTTGAG-GTCTTAGC-CATAAAATCTTTG | loxF8R | 4 | loxF8R | 3 | GCTAAGAC/GTCTTAGC | 7 |
| chr11 | 9123294 | CAAAGATTTTATG-GCTAAGAC-CTCAAAAGCATAG | loxF8R | 3 | loxF8R | 4 | GCTAAGAC/GTCTTAGC | 7 |
| chr14 | 60898794 | CTATGCTTTTGAG-GTCTTAGC-CATAAAATCTTTG | loxF8R | 4 | loxF8R | 3 | GCTAAGAC/GTCTTAGC | 7 |
| chr14 | 81349179 | CAAAGATTTTATG-GCTAAGAC-CTCAAAAGCATAG | loxF8R | 3 | loxF8R | 4 | GCTAAGAC/GTCTTAGC | 7 |
| chr4 | 15859298 | CAAAGATTTTATG-GCTAAGAC-CTCAAAAGCATAG | loxF8R | 3 | loxF8R | 4 | GCTAAGAC/GTCTTAGC | 7 |
| chr5 | 8969665 | CTATGCTTTTGAG-GTCTTAGC-CATAAAATCTTTG | loxF8R | 4 | loxF8R | 3 | GCTAAGAC/GTCTTAGC | 7 |
| chr13 | 84061259 | CTGAGCTTCTGAG-GTCTTAGC-CATAAAATCTTTG | loxF8R | 4 | loxF8R | 3 | GCTAAGAC/GTCTTAGC | 7 |
| chr11 | 128257098 | CAAATATTTTGTG-GCTAAGAC-CTCAAAAGCATAG | loxF8R | 3 | loxF8R | 4 | GCTAAGAC/GTCTTAGC | 7 |
| chr3 | 148019648 | CGAAGATTTTATG-GCTAAGAC-CTCAAAAGCATAG | loxF8R | 3 | loxF8R | 4 | GCTAAGAC/GTCTTAGC | 7 |
| chrX | 22657100 | TTAACATTTTGTG-GCTAAGAC-CACAAAAGCACAG | loxF8R | 3 | loxF8R | 4 | GCTAAGAC/GTCTTAGC | 7 |
| --- | | | | | | | | 0 |
| chr16 | 75949770 | ATAAATATCTGTT-GTTTGAGC-CACGCAGTCTTAA | loxF8L | 4 | loxF8R | 3 | GCTCAAAC/GTTTGAGC | 7 |
| chr15 | 94706681 | ATAAATGTCTGTT-GTTTGAGC-CACCCAATCTATG | loxF8L | 4 | loxF8R | 3 | GCTCAAAC/GTTTGAGC | 7 |
| chr18 | 69391715 | CAAAGACTGGGTG-GCTCAAAC-AACAGAAATTTAT | loxF8R | 3 | loxF8L | 4 | GCTCAAAC/GTTTGAGC | 7 |
| chr22 | 19452112 | CAAAGATAGGGTG-GCTCAAAC-AACAGAAATTTAT | loxF8R | 3 | loxF8L | 4 | GCTCAAAC/GTTTGAGC | 7 |
| --- | | | | | | | | 0 |
| chr5 | 161653254 | CAAAGATTCTGAG-GGAAGAGC-CAGAAAATCTCAA | loxF8R | 3 | loxF8R | 4 | GCTCTTCC/GGAAGAGC | 7 |
| chr5 | 166301584 | TTAGAGATGTGGA-GGAAGAGC-CACACATGTTTAG | loxF8L | 4 | loxF8R | 3 | GCTCTTCC/GGAAGAGC | 7 |
| --- | | | | | | | | 0 |
| chr17 | 9938665 | ATAAATGTCTGTA-GTTTCAGC-CACCCAGTCTGGG | loxF8L | 3 | loxF8R | 4 | GCTGAAAC/GTTTCAGC | 7 |
| chr3 | 14010415 | CATAGATTGGGTG-GCTGAAAC-AACAAACATTTAT | loxF8R | 3 | loxF8L | 4 | GCTGAAAC/GTTTCAGC | 7 |
| chr6 | 152652302 | CTGACACTGAGTG-GCTGAAAC-TACAGAAATTTAT | loxF8R | 4 | loxF8L | 3 | GCTGAAAC/GTTTCAGC | 7 |
| --- | | | | | | | | 0 |
| chr2 | 62335876 | ATAAATCTCTGTT-GCTGAAGC-CACCCAGTCTGTG | loxF8L | 3 | loxF8R | 4 | GCTGAAGC/GCTTCAGC | 7 |
| chr12 | 7448026 | ATAAATTTCTGTA-GCTGAAGC-CACCCAGTCTGTG | loxF8L | 3 | loxF8R | 4 | GCTGAAGC/GCTTCAGC | 7 |
| chr7 | 18698437 | ATAGCCCTGGGGA-GCTTCAGC-GACACAATCATAA | loxF8L | 4 | loxF8R | 3 | GCTGAAGC/GCTTCAGC | 7 |
| --- | | | | | | | | 0 |
| chr3 | 178407468 | ATAAATAAATGGA-GCTGGTGA-TACAAAATCAAAG | loxF8L | 3 | loxF8R | 4 | GCTGGTGA/TCACCAGC | 7 |
| chr2 | 4241320 | TTAAGATAGTTTT-GCTGGTGA-CAAAGTATCTTAG | loxF8R | 4 | loxF8R | 3 | GCTGGTGA/TCACCAGC | 7 |
| --- | | | | | | | | 0 |
| chr5 | 37508395 | ATAAATGTATGGT-GGTTAAGC-CACCCAGTCTATG | loxF8L | 3 | loxF8R | 4 | GCTTAACC/GGTTAAGC | 7 |
| chr22 | 22540151 | CATAGATTGGGTG-GCTTAACC-AACAGAAATTTAT | loxF8R | 3 | loxF8L | 4 | GCTTAACC/GGTTAAGC | 7 |
| --- | | | | | | | | 0 |
| chr13 | 44240979 | ATAAATCTGTCTT-GTGTAAGC-CACTAAATCTATG | loxF8L | 3 | loxF8R | 4 | GCTTACAC/GTGTAAGC | 7 |
| chr13 | 76874525 | CATAGACTGTGTG-GCTTACAC-AACAGAAATTTAT | loxF8R | 3 | loxF8L | 4 | GCTTACAC/GTGTAAGC | 7 |
| --- | | | | | | | | 0 |
| chr3 | 107072780 | CAGAGATTGGGTG-GCTTAGAC-AACAGAAATTTAT | loxF8R | 3 | loxF8L | 4 | GCTTAGAC/GTCTAAGC | 7 |
| chr18 | 45246310 | GTAAATTTCTGTA-GTCTAAGC-CACCCAATCTGTG | loxF8L | 4 | loxF8R | 3 | GCTTAGAC/GTCTAAGC | 7 |
| --- | | | | | | | | 0 |
| chr12 | 125019923 | CATAGACTGTGTG-GCTTATCA-ACAACAGACTTTT | loxF8R | 3 | loxF8L | 4 | GCTTATCA/TGATAAGC | 7 |
| chr8 | 71572495 | AAATTTCTGTGGT-TGATAAGC-CACCCATTCATAG | loxF8L | 4 | loxF8R | 3 | GCTTATCA/TGATAAGC | 7 |
| --- | | | | | | | | 0 |
| chr13 | 100859351 | CTGAAACTGGGTG-GCTTCAAC-AACAGAGATTTAT | loxF8R | 4 | loxF8L | 3 | GCTTCAAC/GTTGAAGC | 7 |
| chr5 | 109505935 | ATAAATGTCTGTA-GTTGAAGC-CACCCAGTCTATG | loxF8L | 3 | loxF8R | 4 | GCTTCAAC/GTTGAAGC | 7 |
| chr1 | 106144428 | CATAGACTGTGTG-GCTTCAAC-AACATAAATTTAT | loxF8R | 3 | loxF8L | 4 | GCTTCAAC/GTTGAAGC | 7 |
| chr7 | 24153102 | CTTAGACTGGGTG-GCTTCAAC-AGCAGAAATTTAT | loxF8R | 3 | loxF8L | 4 | GCTTCAAC/GTTGAAGC | 7 |
| chr21 | 28676149 | CACAGACTGGGTG-GCTTCAAC-TACAGAAATTTAT | loxF8R | 4 | loxF8L | 3 | GCTTCAAC/GTTGAAGC | 7 |
| chr16 | 30842323 | ATAAATTTCTGTT-GTTGAAGC-CACCCAATCTGTG | loxF8L | 4 | loxF8R | 3 | GCTTCAAC/GTTGAAGC | 7 |
| chr1 | 80727749 | CCAAGACTGGGTG-GCTTCAAC-AACAGAAATTTAT | loxF8R | 3 | loxF8L | 4 | GCTTCAAC/GTTGAAGC | 7 |
| chr11 | 36917838 | ATAAATTTCTGTT-GTTGAAGC-CACTCAATCTATG | loxF8L | 4 | loxF8R | 3 | GCTTCAAC/GTTGAAGC | 7 |
| chr4 | 25953171 | GTAAAAGTGTGTG-GTTGAAGC-CACCCAGTCTAGG | loxF8R | 3 | loxF8R | 4 | GCTTCAAC/GTTGAAGC | 7 |
| --- | | | | | | | | 0 |
| chr16 | 64529031 | ATAAAATTATGTT-GTTAAAGC-CACCCAATCTGTG | loxF8R | 4 | loxF8R | 3 | GCTTTAAC/GTTAAAGC | 7 |
| chr8 | 56943697 | ATAAATGTTTGTT-GTTAAAGC-CACACAGTCTATG | loxF8L | 4 | loxF8R | 3 | GCTTTAAC/GTTAAAGC | 7 |
| chr7 | 90456256 | ATAAATTTCTGTT-GTTAAAGC-CACCCAATCTGTG | loxF8L | 4 | loxF8R | 3 | GCTTTAAC/GTTAAAGC | 7 |
| --- | | | | | | | | 0 |
| chr11 | 78539449 | AAAAGGCTTTGGA-GGAAAAAA-TCAAGAGGTTTAT | loxF8L | 4 | loxF8L | 3 | GGAAAAAA/TTTTTTCC | 7 |
| chr13 | 100131026 | TGAGATCTGTGGA-GGAAAAAA-TGGAGAGCTTTAT | loxF8L | 3 | loxF8L | 4 | GGAAAAAA/TTTTTTCC | 7 |
| chr8 | 33125496 | ATAAGAGTTTGGG-GGAAAAAA-CCCACAAAATTAT | loxF8R | 4 | loxF8L | 3 | GGAAAAAA/TTTTTTCC | 7 |
| chr1 | 28333093 | ATAAGATTTTGGA-GGAAAAAA-TCAACAGTTTGAT | loxF8L | 4 | loxF8L | 3 | GGAAAAAA/TTTTTTCC | 7 |
| chr5 | 42721773 | TTATGTCTGTGGA-TTTTTTCC-TTCAAAGTTTAAT | loxF8L | 3 | loxF8L | 4 | GGAAAAAA/TTTTTTCC | 7 |
| --- | | | | | | | | 0 |
| chr2 | 197735324 | ATTAATTTATTGA-TAATTTCC-TCCAATGATTTTT | loxF8L | 4 | loxF8L | 3 | GGAAATTA/TAATTTCC | 7 |
| chr2 | 165989742 | ATCAACCTGAAGA-TAATTTCC-TCTTCATATTTAT | loxF8L | 4 | loxF8L | 3 | GGAAATTA/TAATTTCC | 7 |
| --- | | | | | | | | 0 |
| chr10 | 37051259 | AAAAATATGTTGA-GGACAAAA-TCCAGAGTTCTCT | loxF8L | 3 | loxF8L | 4 | GGACAAAA/TTTTGTCC | 7 |
| chr11 | 77914362 | ATAAAACTGTCAA-GGACAAAA-TCTAAATGTTTAT | loxF8L | 3 | loxF8L | 4 | GGACAAAA/TTTTGTCC | 7 |
| chr7 | 65699566 | ATAAACATTTAGA-TTTTGTCC-TTGACAGTTTTAT | loxF8L | 4 | loxF8L | 3 | GGACAAAA/TTTTGTCC | 7 |
| chr7 | 65193853 | ATAAACATTTAGA-TTTTGTCC-TTGACAGTTTTAT | loxF8L | 4 | loxF8L | 3 | GGACAAAA/TTTTGTCC | 7 |
| --- | | | | | | | | 0 |
| chr1 | 214945122 | ATGACTCTCTGTA-GGATTCAA-TCCAGAGAGTAAT | loxF8L | 4 | loxF8L | 3 | GGATTCAA/TTGAATCC | 7 |
| chr8 | 137660747 | CTAGAAATGTTTG-TTGAATCC-CAGGCAATCTCAG | loxF8R | 4 | loxF8R | 3 | GGATTCAA/TTGAATCC | 7 |
| --- | | | | | | | | 0 |
| chr17 | 6919878 | GTAGATCTGAGGA-GGCCAAAC-TCCACCTAATTTT | loxF8L | 3 | loxF8L | 4 | GGCCAAAC/GTTTGGCC | 7 |

| | | | | | | | | |
|---|---|---|---|---|---|---|---|---|
| chrX | 130869038 | CAAATACTGTGTG-GGCCAAAC-AAAACACATTTAT | loxF8R | 3 | loxF8L | 4 | GGCCAAAC/GTTTGGCC | 7 |
| chr3 | 156559251 | CAAATATTCTGTA-GGCCAAAC-TCAACAGATTTGG | loxF8R | 4 | loxF8L | 3 | GGCCAAAC/GTTTGGCC | 7 |
| --- | | | | | | | | 0 |
| chr10 | 11026084 | AGAAATCTGGACA-GTTGGGCC-TCCACGGATGTTT | loxF8L | 4 | loxF8L | 3 | GGCCCAAC/GTTGGGCC | 7 |
| chr6 | 75609372 | ACAAATCTCTTGA-GGCCCAAC-TTCAGAAATTTAC | loxF8L | 3 | loxF8L | 4 | GGCCCAAC/GTTGGGCC | 7 |
| --- | | | | | | | | 0 |
| chr2 | 148085866 | AAAAAGATCTGGA-GGGGAAAA-TCTACTGTTTTAT | loxF8L | 4 | loxF8L | 3 | GGGGAAAA/TTTTCCCC | 7 |
| chr20 | 15779641 | CTTTGTTTGTGTG-TTTTCCCC-TCCACAGCCTAAG | loxF8R | 3 | loxF8L | 4 | GGGGAAAA/TTTTCCCC | 7 |
| --- | | | | | | | | 0 |
| chr3 | 105021837 | ATAAATCTTTGTG-TCCCACCC-CCACCAAATTTAT | loxF8L | 3 | loxF8L | 4 | GGGTGGGA/TCCCACCC | 7 |
| chr7 | 37514605 | ATGAAGCTTTGGA-GGGTGGGA-CCCAATCTCTTAG | loxF8L | 3 | loxF8R | 4 | GGGTGGGA/TCCCACCC | 7 |
| --- | | | | | | | | 0 |
| chr3 | 150530351 | TTCAATCTTTGGA-GGGTTGAA-TTCACAGATGCAC | loxF8L | 3 | loxF8L | 4 | GGGTTGAA/TTCAACCC | 7 |
| chr4 | 13211748 | TTCAATCTGTGGT-GGGTTGAA-TCCACGGACATAA | loxF8L | 3 | loxF8L | 4 | GGGTTGAA/TTCAACCC | 7 |
| --- | | | | | | | | 0 |
| chr19 | 37492073 | CTGAGCTTTTGAG-GTCTTACC-CAAACAATTTTTG | loxF8R | 4 | loxF8R | 3 | GGTAAGAC/GTCTTACC | 7 |
| chr5 | 117887552 | CTGAGATTTTGAG-GTCTTACC-CATGAAATCTTTG | loxF8R | 3 | loxF8R | 4 | GGTAAGAC/GTCTTACC | 7 |
| chr9 | 134107264 | CAAAGATTTTTTG-GGTAAGAC-CTCAAAAGCATAG | loxF8R | 3 | loxF8R | 4 | GGTAAGAC/GTCTTACC | 7 |
| chr1 | 104898266 | CTATAATTGTGAG-GTCTTACC-CCCAAAATATTTG | loxF8R | 3 | loxF8R | 4 | GGTAAGAC/GTCTTACC | 7 |
| chr4 | 8339226 | CTGAGTTTTTGAG-GTCTTACC-CAAAAAATCTTCG | loxF8R | 4 | loxF8R | 3 | GGTAAGAC/GTCTTACC | 7 |
| --- | | | | | | | | 0 |
| chr8 | 59431727 | ATAAGATGGTTTT-GGTAGAAA-AAGACAATTTTAG | loxF8R | 4 | loxF8R | 3 | GGTAGAAA/TTTCTACC | 7 |
| chr3 | 1987831 | CTGAGATTAAGTT-TTTCTACC-CACACACTTTTAA | loxF8R | 4 | loxF8R | 3 | GGTAGAAA/TTTCTACC | 7 |
| --- | | | | | | | | 0 |
| chr11 | 8570537 | ATAAATCGATGGT-GGTATACC-CACACAATGCAAC | loxF8L | 3 | loxF8R | 4 | GGTATACC/GGTATACC | 7 |
| --- | | | | | | | | 0 |
| chr4 | 79376037 | ATAAATGTATGTA-GGTATATA-TATACATATATAT | loxF8L | 3 | loxF8L | 4 | GGTATATA/TATATACC | 7 |
| chr4 | 146162958 | ATATATATGTATA-TATATACC-TACACATATTTAA | loxF8L | 4 | loxF8L | 3 | GGTATATA/TATATACC | 7 |
| --- | | | | | | | | 0 |
| chr6 | 70571222 | TTTAATCTGTTGA-GGTGGTAA-ATTACAGATTTTT | loxF8L | 3 | loxF8L | 4 | GGTGGTAA/TTACCACC | 7 |
| chrX | 26656252 | ATAAAGCAGTTGT-GGTGGTAA-CACAATTTCTTAG | loxF8L | 4 | loxF8R | 3 | GGTGGTAA/TTACCACC | 7 |
| --- | | | | | | | | 0 |
| chr9 | 19385448 | CACAGATTGGGTG-GGTTAACC-AACAGAAATTTAT | loxF8R | 3 | loxF8L | 4 | GGTTAACC/GGTTAACC | 7 |
| --- | | | | | | | | 0 |
| chr16 | 62523464 | AAAAATCTATGAA-GGTTTGTA-TTCAAATATTCAT | loxF8L | 3 | loxF8L | 4 | GGTTTGTA/TACAAACC | 7 |
| chr2 | 161466783 | CCATGATTTTGTG-GGTTTGTA-TACAGAATGTGAG | loxF8R | 3 | loxF8R | 4 | GGTTTGTA/TACAAACC | 7 |
| --- | | | | | | | | 0 |
| chr17 | 45077742 | GAAAATATCTGGA-GTAGGAAA-ACCACAGAGTTGT | loxF8L | 4 | loxF8L | 3 | GTAGGAAA/TTTCCTAC | 7 |
| chr3 | 188751204 | ATAAATTTGGAGA-TTTCCTAC-CACAAAAGCTAAA | loxF8L | 3 | loxF8R | 4 | GTAGGAAA/TTTCCTAC | 7 |
| --- | | | | | | | | 0 |
| chr20 | 34394397 | TTATGTTTGTTTG-GTATTGTA-TTAACAGCTTTAT | loxF8R | 4 | loxF8L | 3 | GTATTGTA/TACAATAC | 7 |
| chr10 | 116979780 | CTATTAGTCTGTG-GTATTGTA-ATCAGAGATTTAT | loxF8R | 4 | loxF8L | 3 | GTATTGTA/TACAATAC | 7 |
| --- | | | | | | | | 0 |
| chr7 | 22541722 | AGAAATCTTTGCA-TCCTTGAC-CTCCCAGGCTTAG | loxF8L | 3 | loxF8R | 4 | GTCAAGGA/TCCTTGAC | 7 |
| chr9 | 7293874 | TTAAGAATGTGTT-TCCTTGAC-AACACCATTTAAG | loxF8R | 3 | loxF8R | 4 | GTCAAGGA/TCCTTGAC | 7 |
| --- | | | | | | | | 0 |
| chr19 | 6680684 | ACAAGTCTGTTGT-GTTAAGAC-CCCATATTCTTAG | loxF8L | 4 | loxF8R | 3 | GTCTTAAC/GTTAAGAC | 7 |
| chr2 | 7501315 | CAAAGATTTTATG-GTTAAGAC-CACAAAAACACAG | loxF8R | 3 | loxF8R | 4 | GTCTTAAC/GTTAAGAC | 7 |
| chr14 | 24583391 | CTGTGCTTTTGTG-GTCTTAAC-CATAAAATCTTTG | loxF8R | 4 | loxF8R | 3 | GTCTTAAC/GTTAAGAC | 7 |
| chr8 | 109323991 | CAAAGATTTTATG-GTTAAGAC-CTCAAAAGCATAG | loxF8R | 3 | loxF8R | 4 | GTCTTAAC/GTTAAGAC | 7 |
| --- | | | | | | | | 0 |
| chr3 | 82525717 | CTATGCTTTTGTG-GTCTTACA-CAAAAGATCTTTG | loxF8R | 3 | loxF8R | 4 | GTCTTACA/TGTAAGAC | 7 |
| chr3 | 138312695 | CAAAGATTTTTTG-TGTAAGAC-CTCAAAAGCATAG | loxF8R | 3 | loxF8R | 4 | GTCTTACA/TGTAAGAC | 7 |
| chr7 | 16832401 | CAAAGATTTTTTG-TGTAAGAC-CTCAAAAGCATAG | loxF8R | 3 | loxF8R | 4 | GTCTTACA/TGTAAGAC | 7 |
| chrX | 131968434 | CAAAGATTTTTTG-TGTAAGAC-CTCAAAAGCATAG | loxF8R | 3 | loxF8R | 4 | GTCTTACA/TGTAAGAC | 7 |
| --- | | | | | | | | 0 |
| chr7 | 7821455 | TTAAAATTATATG-TGTTTCAC-CTCACTGATTTAT | loxF8R | 4 | loxF8L | 3 | GTGAAACA/TGTTTCAC | 7 |
| chr2 | 5676305 | CTATGTGTGTATG-TGTTTCAC-CCAACAGAATTAT | loxF8R | 4 | loxF8L | 3 | GTGAAACA/TGTTTCAC | 7 |
| --- | | | | | | | | 0 |
| chr21 | 9906237 | ACAAATCTCTCTA-GTGACACA-TCTAAAGATTAAT | loxF8L | 4 | loxF8L | 3 | GTGACACA/TGTGTCAC | 7 |
| chr9 | 41575257 | ATTAATCTTTAGA-TGTGTCAC-TAGAGAGATTTGT | loxF8L | 3 | loxF8L | 4 | GTGACACA/TGTGTCAC | 7 |
| chr9 | 61785501 | ATTAATCTTTAGA-TGTGTCAC-TAGAGAGATTTGT | loxF8L | 3 | loxF8L | 4 | GTGACACA/TGTGTCAC | 7 |
| chr9 | 62359937 | ATTAATCTTTAGA-TGTGTCAC-TAGAGAGATTTGT | loxF8L | 3 | loxF8L | 4 | GTGACACA/TGTGTCAC | 7 |
| chr9 | 39890753 | ACAAATCTCTCTA-GTGACACA-TCTAAAGATTAAT | loxF8L | 4 | loxF8L | 3 | GTGACACA/TGTGTCAC | 7 |
| chr4 | 131766477 | ACAAATCTCTCTA-GTGACACA-TCTAGAGATTAAT | loxF8L | 4 | loxF8L | 3 | GTGACACA/TGTGTCAC | 7 |
| --- | | | | | | | | 0 |
| chr18 | 69817423 | CACAGACTGGGTG-GTGTAAAC-AACAGAGATTTAT | loxF8R | 4 | loxF8L | 3 | GTGTAAAC/GTTTACAC | 7 |
| chr17 | 67585954 | ATTAGATTTTGTT-GTTTACAC-CACCCAATCTGTG | loxF8R | 4 | loxF8R | 3 | GTGTAAAC/GTTTACAC | 7 |
| --- | | | | | | | | 0 |
| chr17 | 45928653 | AGACATCAGTGGA-GTTATATA-TACATATATATAT | loxF8L | 3 | loxF8L | 4 | GTTATATA/TATATAAC | 7 |
| chr4 | 58073578 | ATAAAGCTATACA-TATATAAC-CACCCATTCTTTG | loxF8L | 4 | loxF8R | 3 | GTTATATA/TATATAAC | 7 |
| --- | | | | | | | | 0 |
| chr6 | 9606378 | ATAGATATGGGAA-GTTCCAAA-TACACTGAATTAT | loxF8L | 4 | loxF8L | 3 | GTTCCAAA/TTTGGAAC | 7 |
| chr3 | 167337000 | CTATGAGTATGGG-TTTGGAAC-TCAACAGACTTAA | loxF8R | 4 | loxF8L | 3 | GTTCCAAA/TTTGGAAC | 7 |
| --- | | | | | | | | 0 |
| chr20 | 56403917 | ACAGATCTGTGCA-GTTCTACA-TTCACTGATTATT | loxF8L | 3 | loxF8L | 4 | GTTCTACA/TGTAGAAC | 7 |
| chr2 | 72671103 | CAAAGATTTTTTG-TGTAGAAC-CTCAAAAACATAG | loxF8R | 3 | loxF8R | 4 | GTTCTACA/TGTAGAAC | 7 |
| --- | | | | | | | | 0 |
| chrX | 45785872 | AAAAATCTGTATA-TCTACAAC-TCCACATTTTTTA | loxF8L | 3 | loxF8L | 4 | GTTGTAGA/TCTACAAC | 7 |
| chr2 | 139507017 | ATAAAGCTGTACA-TCTACAAC-CACCTGATCTTTG | loxF8L | 3 | loxF8R | 4 | GTTGTAGA/TCTACAAC | 7 |
| chrX | 106454535 | ATAAATTTGCACA-TCTACAAC-CATATAATCTTTG | loxF8L | 4 | loxF8R | 3 | GTTGTAGA/TCTACAAC | 7 |
| --- | | | | | | | | 0 |
| chr5 | 19961906 | CTAAGTTTGTATT-TTTTAAAC-CAAACAATTTTTT | loxF8R | 3 | loxF8R | 4 | GTTTAAAA/TTTTAAAC | 7 |
| chr3 | 181165642 | CATAGATTGTGTT-TTTTAAAC-AACAGACATTTAT | loxF8R | 3 | loxF8L | 4 | GTTTAAAA/TTTTAAAC | 7 |
| chr8 | 73063039 | ATAAATGTTTGTT-TTTTAAAC-CACCCAATCTATG | loxF8L | 4 | loxF8R | 3 | GTTTAAAA/TTTTAAAC | 7 |
| chr6 | 126288439 | CATAGATTGGGTG-GTTTAAAA-AACAGAAATTTAT | loxF8R | 3 | loxF8L | 4 | GTTTAAAA/TTTTAAAC | 7 |
| --- | | | | | | | | 0 |

| | | | | | | | | |
|---|---|---|---|---|---|---|---|---|
| chr1 | 212562828 | CGAAGACTGGGTG-TTGTAAAC-AACAGATATTTAT | loxF8R | 3 | loxF8L | 4 | GTTTACAA/TTGTAAAC | 7 |
| chr9 | 107902019 | ATAACTATGTGGC-TTGTAAAC-AACAGAAATTTAT | loxF8L | 3 | loxF8L | 4 | GTTTACAA/TTGTAAAC | 7 |
| --- | | | | | | | | 0 |
| chrX | 86698234 | AAAAATATGTATA-TTTAAAAC-CCCACAAACCTAG | loxF8L | 4 | loxF8R | 3 | GTTTTAAA/TTTAAAAC | 7 |
| chr7 | 145732403 | ATAAATGTATGGT-GTTTTAAA-CCACCAGATTTGT | loxF8L | 3 | loxF8L | 4 | GTTTTAAA/TTTAAAAC | 7 |
| --- | | | | | | | | 0 |
| chr12 | 101838812 | CTAATGTTCTGTG-GTTTTCTA-TTCATTGATTTGT | loxF8R | 3 | loxF8L | 4 | GTTTTCTA/TAGAAAAC | 7 |
| chr8 | 91687199 | ATAAATCTATATA-TAGAAAAC-ACCAAAGATTTCA | loxF8L | 3 | loxF8L | 4 | GTTTTCTA/TAGAAAAC | 7 |
| chr9 | 103967244 | CTGAGACTGTGGG-GTTTTCTA-GCTATAGAATTAT | loxF8R | 3 | loxF8L | 4 | GTTTTCTA/TAGAAAAC | 7 |
| chr1 | 193877677 | CTGAGACAGTGTG-GTTTTCTA-GACATAGAATTAT | loxF8R | 3 | loxF8L | 4 | GTTTTCTA/TAGAAAAC | 7 |
| chr5 | 3406939 | CTGAGACTGTGGG-GTTTTCTA-GACATAGAATTAT | loxF8R | 3 | loxF8L | 4 | GTTTTCTA/TAGAAAAC | 7 |
| chr14 | 79957779 | ATAATTCTGTACC-TAGAAAAC-CACATCATCTCAG | loxF8L | 4 | loxF8R | 3 | GTTTTCTA/TAGAAAAC | 7 |
| chr5 | 136582575 | CAGAGATTGTGGG-GTTTTCTA-TCTATAGAATCAT | loxF8R | 3 | loxF8L | 4 | GTTTTCTA/TAGAAAAC | 7 |
| --- | | | | | | | | 0 |
| chr21 | 20329651 | ATCAGAATGTCTG-TTCAAAAC-TCTACAGATGTAA | loxF8R | 4 | loxF8L | 3 | GTTTTGAA/TTCAAAAC | 7 |
| chr20 | 30353214 | TTCCATCTGTGGT-GTTTTGAA-TCCACAGATGTGG | loxF8L | 4 | loxF8L | 3 | GTTTTGAA/TTCAAAAC | 7 |
| --- | | | | | | | | 0 |
| chr5 | 24174173 | ATATAGCTATGAA-TTTTTTTA-TCCACAAATATTT | loxF8L | 4 | loxF8L | 3 | TAAAAAAA/TTTTTTTA | 7 |
| chr10 | 56633552 | ATAAAATAGTGTG-TTTTTTTA-CTTACAATATTAT | loxF8R | 3 | loxF8R | 4 | TAAAAAAA/TTTTTTTA | 7 |
| chr1 | 85063833 | GAGAATCTGTGGA-TTTTTTTA-ATTACATATTTAT | loxF8L | 3 | loxF8L | 4 | TAAAAAAA/TTTTTTTA | 7 |
| chr6 | 121986013 | ATAAATGTATAGA-TAAAAAAA-TTAACAAACTTAT | loxF8L | 3 | loxF8L | 4 | TAAAAAAA/TTTTTTTA | 7 |
| chrX | 145183605 | ATAAATTTTTGAA-TTTTTTTA-CACTCAATTTAAC | loxF8L | 3 | loxF8R | 4 | TAAAAAAA/TTTTTTTA | 7 |
| --- | | | | | | | | 0 |
| chr13 | 101302772 | TTACATATGTGAA-TAAAAATA-TTCACAGAATTTT | loxF8L | 4 | loxF8L | 3 | TAAAAATA/TATTTTTA | 7 |
| chr4 | 156761779 | ATACTTCTGTACA-TAAAAATA-TCCATGTATTTAT | loxF8L | 4 | loxF8L | 3 | TAAAAATA/TATTTTTA | 7 |
| chr5 | 100493699 | ATAAATATGTATA-TAAAAATA-TAAATATATTTAT | loxF8L | 3 | loxF8L | 4 | TAAAAATA/TATTTTTA | 7 |
| --- | | | | | | | | 0 |
| chrX | 78516093 | CTTAGATTCTGTC-TTCTTTTA-CAAAGAATTTGAG | loxF8R | 3 | loxF8R | 4 | TAAAAGAA/TTCTTTTA | 7 |
| chr3 | 31964818 | GTAAGTTAGTCTG-TAAAAGAA-CACACAAATTCAG | loxF8R | 4 | loxF8R | 3 | TAAAAGAA/TTCTTTTA | 7 |
| --- | | | | | | | | 0 |
| chr10 | 3899231 | ATAAATTTTAGGA-TGCTTTTA-TCCAGAGCTGTTT | loxF8L | 3 | loxF8L | 4 | TAAAAGCA/TGCTTTTA | 7 |
| chr13 | 67678411 | ATAAATATTTTTA-TAAAAGCA-AACACTATTTTAG | loxF8L | 4 | loxF8R | 3 | TAAAAGCA/TGCTTTTA | 7 |
| chr4 | 48212124 | ATATATATGTATA-TAAAAGCA-TTCACAGAAATAT | loxF8L | 4 | loxF8L | 3 | TAAAAGCA/TGCTTTTA | 7 |
| --- | | | | | | | | 0 |
| chr16 | 33612550 | AATATCCTGTGGA-TAAAATAA-TCAACTGATCTAT | loxF8L | 4 | loxF8L | 3 | TAAAATAA/TTATTTTA | 7 |
| chr14 | 70946820 | GTAAATGTGTTTA-TTATTTTA-GATACAGATTTAT | loxF8L | 4 | loxF8L | 3 | TAAAATAA/TTATTTTA | 7 |
| --- | | | | | | | | 0 |
| chr6 | 50459238 | ACAAGTTTGTGCA-TCATTTTA-CACTAAATCATAG | loxF8L | 4 | loxF8R | 3 | TAAAATGA/TCATTTTA | 7 |
| chr8 | 4322110 | ATAAAATATTGGA-TCATTTTA-TCCTCCGATTTCT | loxF8L | 4 | loxF8L | 3 | TAAAATGA/TCATTTTA | 7 |
| --- | | | | | | | | 0 |
| chr2 | 200764572 | ACAGATATGTAGA-TAAACATA-TCTACAGATATGT | loxF8L | 4 | loxF8L | 3 | TAAACATA/TATGTTTA | 7 |
| chr2 | 200764596 | ACAGATATGTAGA-TAAACATA-TCTACAGATATGT | loxF8L | 4 | loxF8L | 3 | TAAACATA/TATGTTTA | 7 |
| chr2 | 213659865 | ATACATCTGTATA-TATGTTTA-TGTGCAGGTTTAT | loxF8L | 3 | loxF8L | 4 | TAAACATA/TATGTTTA | 7 |
| chr1 | 70782014 | ATAAATATGTACA-TATGTTTA-TTCACTGGTTCAT | loxF8L | 3 | loxF8L | 4 | TAAACATA/TATGTTTA | 7 |
| chr15 | 94244097 | ATACATATGTGTA-TATGTTTA-TATACACATGTAT | loxF8L | 3 | loxF8L | 4 | TAAACATA/TATGTTTA | 7 |
| --- | | | | | | | | 0 |
| chr14 | 26926657 | ATAAATATGTAAA-TAGGTTTA-TAGACTATCTCAG | loxF8L | 3 | loxF8R | 4 | TAAACCTA/TAGGTTTA | 7 |
| chr15 | 79553657 | ATGGATTTGTTGA-TAAACCTA-TCCAAATATTTAG | loxF8L | 4 | loxF8L | 3 | TAAACCTA/TAGGTTTA | 7 |
| --- | | | | | | | | 0 |
| chr4 | 22416217 | CAAAGTTTGTAAG-TTCATTTA-ACCATACATTTAT | loxF8R | 4 | loxF8L | 3 | TAAATGAA/TTCATTTA | 7 |
| chr2 | 100435008 | ATAAGATTCTATG-TTCATTTA-CAAACAGTTTTTG | loxF8R | 3 | loxF8R | 4 | TAAATGAA/TTCATTTA | 7 |
| chr5 | 122132206 | CTATAATTTTGTA-TTCATTTA-TCCATTGATGTAT | loxF8R | 4 | loxF8L | 3 | TAAATGAA/TTCATTTA | 7 |
| --- | | | | | | | | 0 |
| chr4 | 84954487 | AGAAATCTGACTA-TAAATTTA-TCTATAAATTTAT | loxF8L | 4 | loxF8L | 3 | TAAATTTA/TAAATTTA | 7 |
| chrX | 11132517 | ATTTATCTATTGA-TAAATTTA-TGCAACGATTTAT | loxF8L | 4 | loxF8L | 3 | TAAATTTA/TAAATTTA | 7 |
| --- | | | | | | | | 0 |
| chr4 | 24822025 | ACAAATCTTTGTA-TGTTCTTA-GATTCAATTTTAG | loxF8L | 3 | loxF8R | 4 | TAAGAACA/TGTTCTTA | 7 |
| chr1 | 185896384 | ATAAACCTGTGTG-TAAGAACA-CACAAAAACTCAC | loxF8L | 3 | loxF8R | 4 | TAAGAACA/TGTTCTTA | 7 |
| --- | | | | | | | | 0 |
| chr9 | 15057753 | ATACATATATGTA-TAAGCACA-CACACAATATTTT | loxF8L | 4 | loxF8R | 3 | TAAGCACA/TGTGCTTA | 7 |
| chr15 | 101926474 | ATATATATGTGTA-TGTGCTTA-TATACAGATATAA | loxF8L | 3 | loxF8L | 4 | TAAGCACA/TGTGCTTA | 7 |
| chr19 | 105732 | TTATATCTGTATA-TAAGCACA-TACACATATATAT | loxF8L | 4 | loxF8L | 3 | TAAGCACA/TGTGCTTA | 7 |
| chr1 | 64147 | TTATATCTGTATA-TAAGCACA-TACACATATATAT | loxF8L | 4 | loxF8L | 3 | TAAGCACA/TGTGCTTA | 7 |
| --- | | | | | | | | 0 |
| chr5 | 44863030 | AAAAATCTCTAGC-TAAGCTGA-CAGAGATTCTTAG | loxF8L | 4 | loxF8R | 3 | TAAGCTGA/TCAGCTTA | 7 |
| chr13 | 18612276 | AAAACTCTGAGGC-TCAGCTTA-AACTCAATCCTAG | loxF8L | 4 | loxF8R | 3 | TAAGCTGA/TCAGCTTA | 7 |
| chr15 | 21341948 | CTAGGATTGAGTT-TAAGCTGA-GCCTCAGAGTTTT | loxF8R | 3 | loxF8L | 4 | TAAGCTGA/TCAGCTTA | 7 |
| chr15 | 20772861 | CTAGGATTGAGTT-TAAGCTGA-GCCTCAGAGTTTT | loxF8R | 3 | loxF8L | 4 | TAAGCTGA/TCAGCTTA | 7 |
| chr18 | 14447617 | CTAGGATTGAGTT-TAAGCTGA-GCCTCAGAGTTTT | loxF8R | 3 | loxF8L | 4 | TAAGCTGA/TCAGCTTA | 7 |
| --- | | | | | | | | 0 |
| chr16 | 89724881 | GTATATCTATGTA-TCTACTTA-TCTACCTATTTAT | loxF8L | 4 | loxF8L | 3 | TAAGTAGA/TCTACTTA | 7 |
| chr5 | 8755947 | ATTAATCTATGTA-TCTACTTA-TCTACCTACTTAT | loxF8L | 3 | loxF8L | 4 | TAAGTAGA/TCTACTTA | 7 |
| --- | | | | | | | | 0 |
| chr6 | 126879053 | ATAAATCTAAGAA-TAATAATA-TCCGCATATATAG | loxF8L | 3 | loxF8L | 4 | TAATAATA/TATTATTA | 7 |
| chr12 | 89705942 | GTAAGACTATATG-TAATAATA-TCCACATCTTAAT | loxF8R | 4 | loxF8L | 3 | TAATAATA/TATTATTA | 7 |
| --- | | | | | | | | 0 |
| chr2 | 22323210 | ATAACTATTTGTA-TAATATTA-TACAAATATTTAT | loxF8L | 4 | loxF8L | 3 | TAATATTA/TAATATTA | 7 |
| --- | | | | | | | | 0 |
| chr13 | 63222626 | AAAAATCTGGTTA-TAATCCTA-TGCACCTATTTAT | loxF8L | 4 | loxF8L | 3 | TAATCCTA/TAGGATTA | 7 |
| chr3 | 6006547 | ATATATTTGTGGT-TAGGATTA-AACGCATCCTTAG | loxF8L | 3 | loxF8R | 4 | TAATCCTA/TAGGATTA | 7 |
| --- | | | | | | | | 0 |
| chr6 | 65891821 | CAATGATTGTGTA-TTACATTA-TACAAAGTTTTCT | loxF8R | 3 | loxF8L | 4 | TAATGTAA/TTACATTA | 7 |
| chr5 | 165678653 | GTAAGTATCTGGA-TTACATTA-ACCACATACTTAT | loxF8L | 4 | loxF8L | 3 | TAATGTAA/TTACATTA | 7 |
| --- | | | | | | | | 0 |
| chr16 | 4031420 | ACAAATGTGTGTA-TAATTATA-TTCACACATACAT | loxF8L | 3 | loxF8L | 4 | TAATTATA/TATAATTA | 7 |
| chr3 | 167118694 | ATGAATTTGTGTC-TAATTATA-TCTACACATATAT | loxF8L | 4 | loxF8L | 3 | TAATTATA/TATAATTA | 7 |
| --- | | | | | | | | 0 |

| | | | | | | | | |
|---|---|---|---|---|---|---|---|---|
| chr1 | 74534647 | ATATCTCTTTGGA-TTAAATTA-CATAGAGTTTTAG | loxF8L | 3 | loxF8R | 4 | TAATTTAA/TTAAATTA | 7 |
| chr2 | 14466784 | CTAAGTATTTGTT-TAATTTAA-CACAATATCTTTG | loxF8R | 4 | loxF8R | 3 | TAATTTAA/TTAAATTA | 7 |
| --- | | | | | | | | 0 |
| chr4 | 146595893 | ATAATTCTGAGTA-TACAAAAA-CCCACTGAATTGT | loxF8L | 3 | loxF8L | 4 | TACAAAAA/TTTTTGTA | 7 |
| chr3 | 103331907 | ATAACTTTGTGGT-TTTTTGTA-TATACAGTTTTAC | loxF8L | 3 | loxF8L | 4 | TACAAAAA/TTTTTGTA | 7 |
| chr6 | 93856070 | ATAGATCTGTGTT-TTTTTGTA-TTGCCAGAATTAT | loxF8L | 3 | loxF8L | 4 | TACAAAAA/TTTTTGTA | 7 |
| chr2 | 171289447 | ATATAACTCTGGA-TACAAAAA-TCCAAAGAAGAAT | loxF8L | 3 | loxF8L | 4 | TACAAAAA/TTTTTGTA | 7 |
| --- | | | | | | | | 0 |
| chr9 | 31362614 | AAAAATATCTGGA-TACACACA-ACCGCATATTTAC | loxF8L | 3 | loxF8L | 4 | TACACACA/TGTGTGTA | 7 |
| chr4 | 66234388 | CTGTGTGTGTGTG-TGTGTGTA-TACACATATTTTT | loxF8R | 4 | loxF8L | 3 | TACACACA/TGTGTGTA | 7 |
| chr14 | 20101226 | ATATATATGTGTG-TGTGTGTA-TACACATATTTTT | loxF8L | 4 | loxF8L | 3 | TACACACA/TGTGTGTA | 7 |
| chr14 | 52775137 | AAAACACTGTGGT-TACACACA-CACACAAAATTAT | loxF8L | 4 | loxF8R | 3 | TACACACA/TGTGTGTA | 7 |
| chr2 | 140935869 | AGAAATCTGCTGA-TACACACA-CACACATATATAT | loxF8L | 3 | loxF8L | 4 | TACACACA/TGTGTGTA | 7 |
| chr7 | 152554542 | TAAACATTGTGTG-TGTGTGTA-TGTATATATTTAT | loxF8R | 3 | loxF8L | 4 | TACACACA/TGTGTGTA | 7 |
| chrX | 4442737 | ATAAATATATATA-TACACACA-CACACATTTTTAA | loxF8L | 4 | loxF8R | 3 | TACACACA/TGTGTGTA | 7 |
| chr9 | 30668367 | ATACAGATGTGTA-TACACACA-CACACATATTTAT | loxF8L | 4 | loxF8L | 3 | TACACACA/TGTGTGTA | 7 |
| chr19 | 45087906 | ATAAATATGTGTG-TGTGTGTA-TATATAGATATAT | loxF8L | 3 | loxF8L | 4 | TACACACA/TGTGTGTA | 7 |
| chr11 | 55943145 | ATATATGTGTGTG-TGTGTGTA-TAGATAGATTTAT | loxF8L | 4 | loxF8L | 3 | TACACACA/TGTGTGTA | 7 |
| chr2 | 221125688 | ATATATGTTTGTA-TGTGTGTA-TACATACATTTAT | loxF8L | 4 | loxF8L | 3 | TACACACA/TGTGTGTA | 7 |
| chr9 | 1283871 | ATAACTCTGTGTG-TGTGTGTA-CACACACACATAC | loxF8L | 3 | loxF8R | 4 | TACACACA/TGTGTGTA | 7 |
| chr9 | 4764442 | ATGATTTTGTGTA-TGTGTGTA-CACACATCCTTAT | loxF8L | 4 | loxF8R | 3 | TACACACA/TGTGTGTA | 7 |
| chr1 | 194717634 | ATATATATATGTA-TACACACA-CACACACACTTAT | loxF8L | 4 | loxF8R | 3 | TACACACA/TGTGTGTA | 7 |
| chr21 | 9547398 | GTAAGAGTGTATG-TACACACA-TGCACATGTGTAT | loxF8R | 3 | loxF8L | 4 | TACACACA/TGTGTGTA | 7 |
| chr7 | 57598333 | GTAAGAGTGTATG-TACACACA-TGCACATGTGTAT | loxF8R | 3 | loxF8L | 4 | TACACACA/TGTGTGTA | 7 |
| chrX | 147052311 | ATACATATGTGTA-TACACACA-CACACATATATAT | loxF8L | 3 | loxF8L | 4 | TACACACA/TGTGTGTA | 7 |
| chr11 | 37899127 | ATACATGTGTGTA-TACACACA-CACACACACATAC | loxF8L | 3 | loxF8R | 4 | TACACACA/TGTGTGTA | 7 |
| chr9 | 26104572 | ATACGTGTGTGTG-TACACACA-TACACACATATAT | loxF8R | 4 | loxF8L | 3 | TACACACA/TGTGTGTA | 7 |
| chr9 | 26104604 | ATACGTGTGTGTG-TACACACA-TACACACATATAT | loxF8R | 4 | loxF8L | 3 | TACACACA/TGTGTGTA | 7 |
| chr9 | 26104540 | ATACGTGTGTGTG-TACACACA-TACACACATATAT | loxF8R | 4 | loxF8L | 3 | TACACACA/TGTGTGTA | 7 |
| chr5 | 25001972 | GTGTGTTTGTGTG-TGTGTGTA-TACACACATCTAT | loxF8R | 4 | loxF8L | 3 | TACACACA/TGTGTGTA | 7 |
| chr14 | 94188570 | ATATATGTGTGTA-TACACACA-CACACACTTATAT | loxF8L | 3 | loxF8R | 4 | TACACACA/TGTGTGTA | 7 |
| chr3 | 60792333 | ATATAGTTGTGTA-TACACACA-CACACACACTTAC | loxF8L | 4 | loxF8R | 3 | TACACACA/TGTGTGTA | 7 |
| chr6 | 142381245 | ATATATATGTGTA-TACACACA-TATACATATATAT | loxF8L | 3 | loxF8L | 4 | TACACACA/TGTGTGTA | 7 |
| chr8 | 110104881 | ATATATATGTGCA-TGTGTGTA-TATACATATATAT | loxF8L | 3 | loxF8L | 4 | TACACACA/TGTGTGTA | 7 |
| chr11 | 79083962 | ATATATGTGTGTG-TGTGTGTA-TACACATATATAT | loxF8L | 4 | loxF8L | 3 | TACACACA/TGTGTGTA | 7 |
| chr15 | 61700272 | ATATATATGTGTA-TACACACA-CACACACATATAT | loxF8L | 3 | loxF8L | 4 | TACACACA/TGTGTGTA | 7 |
| chr1 | 66152583 | ATATATGTGTGTG-TGTGTGTA-TACACATATATAT | loxF8L | 4 | loxF8L | 3 | TACACACA/TGTGTGTA | 7 |
| chr6 | 135756158 | ATATATGTGTGTG-TGTGTGTA-TACACATATATAT | loxF8L | 4 | loxF8L | 3 | TACACACA/TGTGTGTA | 7 |
| chrX | 69518426 | ATATATATGTGTA-TACACACA-CACACACCCCAAG | loxF8L | 3 | loxF8R | 4 | TACACACA/TGTGTGTA | 7 |
| chr2 | 230436985 | ATATATATGTGTA-TACACACA-CACACATATATAT | loxF8L | 3 | loxF8L | 4 | TACACACA/TGTGTGTA | 7 |
| chr6 | 6329355 | ATATATATGTGTA-TACACACA-CACACATATATAT | loxF8L | 3 | loxF8L | 4 | TACACACA/TGTGTGTA | 7 |
| chr8 | 83983155 | ATATATATGTGTA-TACACACA-CACACATATATAT | loxF8L | 3 | loxF8L | 4 | TACACACA/TGTGTGTA | 7 |
| chr3 | 126242926 | ATATATATGTGTA-TGTGTGTA-TACACAGACGTAG | loxF8L | 3 | loxF8L | 4 | TACACACA/TGTGTGTA | 7 |
| chr3 | 25630261 | ATGTATGTGTGCA-TGTGTGTA-TACACATATATAT | loxF8L | 4 | loxF8L | 3 | TACACACA/TGTGTGTA | 7 |
| chr1 | 193588128 | ATATATGTGTGTA-TACACACA-CACACATATATAT | loxF8L | 3 | loxF8L | 4 | TACACACA/TGTGTGTA | 7 |
| chr1 | 193588158 | ATATATGTGTGTA-TACACACA-CACACATATATAT | loxF8L | 3 | loxF8L | 4 | TACACACA/TGTGTGTA | 7 |
| chr8 | 50148091 | GTAACATAGCGTG-TGTGTGTA-TACATAGATATAT | loxF8R | 4 | loxF8L | 3 | TACACACA/TGTGTGTA | 7 |
| chr6 | 154525290 | ATATATGTGTATA-TACACACA-TACATAGATATAT | loxF8L | 4 | loxF8L | 3 | TACACACA/TGTGTGTA | 7 |
| chr5 | 101474973 | ATATATGTGTGAA-TACACACA-CACACACACATAT | loxF8L | 3 | loxF8R | 4 | TACACACA/TGTGTGTA | 7 |
| chr7 | 104522971 | CTGTGTGTGTGTG-TGTGTGTA-TGCACACATATAT | loxF8R | 4 | loxF8L | 3 | TACACACA/TGTGTGTA | 7 |
| chr7 | 37826262 | ATCTATCTATAGA-TACACACA-TGCACACATATAT | loxF8L | 4 | loxF8L | 3 | TACACACA/TGTGTGTA | 7 |
| chr11 | 94898233 | ATATATGTGTGTA-TACACACA-CACACAAACACAC | loxF8L | 3 | loxF8R | 4 | TACACACA/TGTGTGTA | 7 |
| chr10 | 53415380 | ATATATGTGTGTA-TACACACA-CACACAAACACAT | loxF8L | 3 | loxF8R | 4 | TACACACA/TGTGTGTA | 7 |
| chr13 | 39906352 | CTGTGTGTGTGTG-TGTGTGTA-TACACACATATAT | loxF8R | 4 | loxF8L | 3 | TACACACA/TGTGTGTA | 7 |
| chr12 | 96883941 | ATATGTGTGTGTG-TGTGTGTA-TACACACATATAT | loxF8R | 4 | loxF8L | 3 | TACACACA/TGTGTGTA | 7 |
| chr12 | 114928811 | ATATGTATGTGTG-TGTGTGTA-TACACACATATAT | loxF8R | 4 | loxF8L | 3 | TACACACA/TGTGTGTA | 7 |
| chrX | 146080625 | ATATATGTGTGTA-TGTGTGTA-TACACAAAAATAT | loxF8L | 3 | loxF8L | 4 | TACACACA/TGTGTGTA | 7 |
| chr7 | 22416606 | GAAAGTGTGTGTG-TGTGTGTA-TCAACACATATAT | loxF8R | 4 | loxF8L | 3 | TACACACA/TGTGTGTA | 7 |
| chr14 | 66507131 | ATATGAGTGTGTG-TACACACA-CACACACGCATAT | loxF8R | 3 | loxF8R | 4 | TACACACA/TGTGTGTA | 7 |
| chr13 | 34673601 | TTAAAAAAGTGTG-TACACACA-CACACACTCATAT | loxF8R | 4 | loxF8R | 3 | TACACACA/TGTGTGTA | 7 |
| chr3 | 2015754 | CCAAGAATGAATG-TGTGTGTA-CATAGAATCTTTG | loxF8R | 4 | loxF8R | 3 | TACACACA/TGTGTGTA | 7 |
| chr2 | 57468830 | GTATATATGTGTA-TACACACA-CACAATGTCTTAG | loxF8L | 4 | loxF8R | 3 | TACACACA/TGTGTGTA | 7 |
| chr13 | 23601655 | GTATGCTTGTGTG-TACACACA-CACACACACATAA | loxF8R | 3 | loxF8R | 4 | TACACACA/TGTGTGTA | 7 |
| --- | | | | | | | | 0 |
| chr6 | 117923871 | ATACATATGTAGA-TACACATA-TACACATATATGT | loxF8L | 3 | loxF8L | 4 | TACACATA/TATGTGTA | 7 |
| chrX | 44601610 | ACATATATGTGTA-TATGTGTA-TGCACACATATAT | loxF8L | 4 | loxF8L | 3 | TACACATA/TATGTGTA | 7 |
| chrX | 44601584 | ACATATATGTGTA-TATGTGTA-TGCACACATATAT | loxF8L | 4 | loxF8L | 3 | TACACATA/TATGTGTA | 7 |
| chrX | 44601636 | ACATATATGTGTA-TATGTGTA-TGCACACATATAT | loxF8L | 4 | loxF8L | 3 | TACACATA/TATGTGTA | 7 |
| chrX | 44601688 | ACATATATGTGTA-TATGTGTA-TGCACACATATAT | loxF8L | 4 | loxF8L | 3 | TACACATA/TATGTGTA | 7 |
| chrX | 44601714 | ACATATATGTGTA-TATGTGTA-TGCACACATATAT | loxF8L | 4 | loxF8L | 3 | TACACATA/TATGTGTA | 7 |
| chrX | 44601662 | ACATATATGTGTA-TATGTGTA-TGCACACATATAT | loxF8L | 4 | loxF8L | 3 | TACACATA/TATGTGTA | 7 |
| chr22 | 45767893 | CTTACACTGTGTG-TATGTGTA-CACACTGTTTTAT | loxF8R | 3 | loxF8L | 4 | TACACATA/TATGTGTA | 7 |
| chr5 | 22659751 | ATACATATGTCTA-TACACATA-CACACAGTTTTAT | loxF8L | 4 | loxF8L | 3 | TACACATA/TATGTGTA | 7 |
| chrX | 86156340 | ATACAACTATGCA-TACACATA-TACACACGTTTAT | loxF8L | 4 | loxF8L | 3 | TACACATA/TATGTGTA | 7 |
| chr7 | 120750745 | ATAAATTTGTTCA-TACACATA-TACATAGATTTTC | loxF8L | 3 | loxF8L | 4 | TACACATA/TATGTGTA | 7 |
| chr6 | 67015997 | ATAAGTGTGTGTA-TATGTGTA-TACACACATACAT | loxF8L | 3 | loxF8L | 4 | TACACATA/TATGTGTA | 7 |
| chr12 | 130745906 | GTATATATGTGGA-TATGTGTA-TATACACATGTAT | loxF8L | 3 | loxF8L | 4 | TACACATA/TATGTGTA | 7 |
| chr6 | 60562278 | ATACATGTGTGTA-TATGTGTA-TACACATACATAT | loxF8L | 3 | loxF8L | 4 | TACACATA/TATGTGTA | 7 |
| chr6 | 153551008 | ATATATATATGTA-TACACATA-TGCACACATATAT | loxF8L | 4 | loxF8L | 3 | TACACATA/TATGTGTA | 7 |
| chr13 | 81600309 | ATATATATGTGTA-TATGTGTA-TATACATATATAT | loxF8L | 3 | loxF8L | 4 | TACACATA/TATGTGTA | 7 |
| chr3 | 36734544 | ATATATGTATGTA-TACACATA-TACACATATATAT | loxF8L | 4 | loxF8L | 3 | TACACATA/TATGTGTA | 7 |
| chr13 | 81600335 | ATATATATGTGTA-TATGTGTA-TATACACATATAT | loxF8L | 3 | loxF8L | 4 | TACACATA/TATGTGTA | 7 |
| chr3 | 163414720 | ATATATATGTGTA-TACACATA-TCTATACATATAT | loxF8L | 3 | loxF8L | 4 | TACACATA/TATGTGTA | 7 |
| chr20 | 14057631 | CTAAAGCTGTGGG-TATGTGTA-TACATACATATAT | loxF8L | 3 | loxF8L | 4 | TACACATA/TATGTGTA | 7 |
| chr2 | 227203426 | ATATATGTGTGTA-TATGTGTA-TACATACATATAT | loxF8L | 3 | loxF8L | 4 | TACACATA/TATGTGTA | 7 |
| chr3 | 78682545 | ATATATGTGTGTA-TATGTGTA-TATACACATATAT | loxF8L | 3 | loxF8L | 4 | TACACATA/TATGTGTA | 7 |
| chr2 | 142173195 | ATATTTATGTGTA-TATGTGTA-TGCACACATATAT | loxF8L | 4 | loxF8L | 3 | TACACATA/TATGTGTA | 7 |

| | | | | | | | | |
|---|---|---|---|---|---|---|---|---|
| --- | | | | | | | | 0 |
| chr2 | 64573088 | ATAAATATGTTTA-TGCGTGTA-TATACACATATAT | loxF8L | 3 | loxF8L | 4 | TACACGCA/TGCGTGTA | 7 |
| chr10 | 96801237 | ATAGATGTGTGTA-TACACGCA-TGCACACATATAC | loxF8L | 3 | loxF8L | 4 | TACACGCA/TGCGTGTA | 7 |
| --- | | | | | | | | 0 |
| chr12 | 87485920 | ATACATATGTGTA-TACGTGTA-TACACATATGCAT | loxF8L | 3 | loxF8L | 4 | TACACGTA/TACGTGTA | 7 |
| chr3 | 91357182 | ATATGTGTGTGTG-TACACGTA-TACACACATGTAT | loxF8R | 4 | loxF8L | 3 | TACACGTA/TACGTGTA | 7 |
| --- | | | | | | | | 0 |
| chr8 | 998028 | ATAAACATGTGCA-TACAGTCA-TACACAGAAACAT | loxF8L | 3 | loxF8L | 4 | TACAGTCA/TGACTGTA | 7 |
| chr5 | 131463777 | ATAAATTAGTAGA-TACAGTCA-TTCAGATATATAT | loxF8L | 3 | loxF8L | 4 | TACAGTCA/TGACTGTA | 7 |
| --- | | | | | | | | 0 |
| chr4 | 47855953 | ATAAATCTGCCTA-TACATAAA-TACATAATAGTAG | loxF8L | 3 | loxF8R | 4 | TACATAAA/TTTATGTA | 7 |
| chrX | 5069279 | ATAGATCTATAGA-TACATAAA-TACATAGATACAT | loxF8L | 3 | loxF8L | 4 | TACATAAA/TTTATGTA | 7 |
| --- | | | | | | | | 0 |
| chr16 | 25873545 | ATCTATCTGTCTA-TACATCCA-TCCACCCATTTGT | loxF8L | 4 | loxF8L | 3 | TACATCCA/TGGATGTA | 7 |
| chr2 | 66714640 | ATATATATATGGA-TGGATGTA-TACACATACATAT | loxF8L | 3 | loxF8L | 4 | TACATCCA/TGGATGTA | 7 |
| --- | | | | | | | | 0 |
| chr17 | 75127757 | CTAATTGTGTGTG-TTTTGGTA-GCGACAGAGTTTT | loxF8R | 3 | loxF8L | 4 | TACCAAAA/TTTTGGTA | 7 |
| chr16 | 3529075 | ACCCACCTGTGGA-TACCAAAA-TCCACAGATGTTC | loxF8L | 4 | loxF8L | 3 | TACCAAAA/TTTTGGTA | 7 |
| chr4 | 39208874 | GAGCATCTGTGGA-TTTTGGTA-TCCACAGAGGGAT | loxF8L | 4 | loxF8L | 3 | TACCAAAA/TTTTGGTA | 7 |
| chrX | 6980879 | GGACATCTGTGGA-TTTTGGTA-TCCACAGGGTTCC | loxF8L | 3 | loxF8L | 4 | TACCAAAA/TTTTGGTA | 7 |
| --- | | | | | | | | 0 |
| chr3 | 79096669 | ATATATATGTGTG-TATACGTA-CACACATATTTAT | loxF8L | 4 | loxF8L | 3 | TACGTATA/TATACGTA | 7 |
| chr18 | 56262466 | ATACATATGTGTA-TATACGTA-TATACATATGTAT | loxF8L | 3 | loxF8L | 4 | TACGTATA/TATACGTA | 7 |
| chr12 | 39895585 | ATATATATGTGTA-TATACGTA-TATACACATATAT | loxF8L | 3 | loxF8L | 4 | TACGTATA/TATACGTA | 7 |
| chr7 | 75664455 | ATATGTGTGTGTG-TATACGTA-TACACACATATAT | loxF8R | 4 | loxF8L | 3 | TACGTATA/TATACGTA | 7 |
| chr3 | 78478142 | TTAAATATGTGCA-TACGTATA-TACACATATATAG | loxF8L | 3 | loxF8L | 4 | TACGTATA/TATACGTA | 7 |
| --- | | | | | | | | 0 |
| chr1 | 32048748 | ATAATTCAGTGGT-TTTTAGTA-TATACAGAGTTGT | loxF8L | 3 | loxF8L | 4 | TACTAAAA/TTTTAGTA | 7 |
| chr1 | 164402388 | ACAACTCTGTGAA-TACTAAAA-ACCACTCAATTAT | loxF8L | 3 | loxF8L | 4 | TACTAAAA/TTTTAGTA | 7 |
| chr6 | 158797868 | ATAATTCAGTGGT-TTTTAGTA-TATTCAGAGTTAT | loxF8L | 3 | loxF8L | 4 | TACTAAAA/TTTTAGTA | 7 |
| chr6 | 26996239 | ATTTATATTTGGA-TTTTAGTA-TAGACAGATTAAT | loxF8L | 4 | loxF8L | 3 | TACTAAAA/TTTTAGTA | 7 |
| --- | | | | | | | | 0 |
| chr3 | 32526070 | ATATATATATGTA-TAGACAAA-TCCAAAGATTGTT | loxF8L | 4 | loxF8L | 3 | TAGACAAA/TTTGTCTA | 7 |
| chr19 | 22904710 | ATAAAATTATGAG-TTTGTCTA-TGCAAATATTTAT | loxF8R | 4 | loxF8L | 3 | TAGACAAA/TTTGTCTA | 7 |
| --- | | | | | | | | 0 |
| chr12 | 113895951 | ACATATCTATGTA-TCTCTCTA-TACACATATCTAT | loxF8L | 4 | loxF8L | 3 | TAGAGAGA/TCTCTCTA | 7 |
| chr14 | 47684198 | ATATATCTGTATA-TCTCTCTA-TATAGAGATATAT | loxF8L | 3 | loxF8L | 4 | TAGAGAGA/TCTCTCTA | 7 |
| --- | | | | | | | | 0 |
| chr1 | 32844778 | ATATATCTCTGTA-TAGATCTA-TAGAGAGATATAT | loxF8L | 3 | loxF8L | 4 | TAGATCTA/TAGATCTA | 7 |
| --- | | | | | | | | 0 |
| chr10 | 37055786 | ATATATGTATGTA-TGAATCTA-TCCACAGAATGTT | loxF8L | 4 | loxF8L | 3 | TAGATTCA/TGAATCTA | 7 |
| chr1 | 169242832 | ATAAAATTGGGTG-TGAATCTA-ACCAAAAATGTAT | loxF8R | 3 | loxF8L | 4 | TAGATTCA/TGAATCTA | 7 |
| chr8 | 37777071 | TTAATCTTATGTG-TAGATTCA-CCAACAATCTTAT | loxF8R | 4 | loxF8R | 3 | TAGATTCA/TGAATCTA | 7 |
| --- | | | | | | | | 0 |
| chr15 | 81483724 | CCAATATGGTGTC-TTTGCCTA-CTCACCCTCTTAG | loxF8R | 4 | loxF8R | 3 | TAGGCAAA/TTTGCCTA | 7 |
| chr9 | 90819268 | CTGAATTTGTGAA-TTTGCCTA-CTCACAATCACAG | loxF8L | 4 | loxF8R | 3 | TAGGCAAA/TTTGCCTA | 7 |
| --- | | | | | | | | 0 |
| chr5 | 34121118 | CCCATAGTGTGTG-TAGGTATA-TTCACATATTTTT | loxF8R | 4 | loxF8L | 3 | TAGGTATA/TATACCTA | 7 |
| chr5 | 20740335 | AAAAATATGTGAA-TATACCTA-CACACACTATGGG | loxF8L | 3 | loxF8R | 4 | TAGGTATA/TATACCTA | 7 |
| chr8 | 116774532 | ATATATATGTATA-TAGGTATA-TACACATATGTAT | loxF8L | 4 | loxF8L | 3 | TAGGTATA/TATACCTA | 7 |
| chr3 | 79256968 | TTAAATCAGTTAA-TATACCTA-AACACAGTCTTAC | loxF8L | 4 | loxF8R | 3 | TAGGTATA/TATACCTA | 7 |
| --- | | | | | | | | 0 |
| chr4 | 4278709 | ATAAATGAATGAA-TATTACTA-TCCACATTTTTTT | loxF8L | 4 | loxF8L | 3 | TAGTAATA/TATTACTA | 7 |
| chr3 | 160903034 | TTAAAAAGGTGGA-TAGTAATA-CACACCTTGTTAG | loxF8L | 4 | loxF8R | 3 | TAGTAATA/TATTACTA | 7 |
| --- | | | | | | | | 0 |
| chr14 | 91538123 | TTCAATCTGTGGT-TAGTTGAA-TCCATGGATGCAT | loxF8L | 3 | loxF8L | 4 | TAGTTGAA/TTCAACTA | 7 |
| chr9 | 88401674 | GAAAATCTGAGGT-TAGTTGAA-TCCACAGATGTGG | loxF8L | 4 | loxF8L | 3 | TAGTTGAA/TTCAACTA | 7 |
| --- | | | | | | | | 0 |
| chr2 | 225118588 | ATAAATATATGTA-TATAAAAA-TATATATATTTAT | loxF8L | 3 | loxF8L | 4 | TATAAAAA/TTTTTATA | 7 |
| chr3 | 83308110 | ATTAATTTGTGCA-TTTTTATA-TTAATACATTTAT | loxF8L | 3 | loxF8L | 4 | TATAAAAA/TTTTTATA | 7 |
| --- | | | | | | | | 0 |
| chr7 | 46877934 | ACAGGATTGTGTG-TGTTTATA-TACATATATGTAT | loxF8R | 3 | loxF8L | 4 | TATAAACA/TGTTTATA | 7 |
| chr7 | 151903173 | ATAGGTCTGTGTA-TGTTTATA-TACATACACTTAT | loxF8L | 3 | loxF8L | 4 | TATAAACA/TGTTTATA | 7 |
| chr1 | 223457457 | ATAATAGTGTGTG-TGTTTATA-TACACACATACAT | loxF8R | 3 | loxF8L | 4 | TATAAACA/TGTTTATA | 7 |
| chr11 | 83668820 | ATATATATGTGTA-TATAAACA-CACACACACATAT | loxF8L | 3 | loxF8R | 4 | TATAAACA/TGTTTATA | 7 |
| chr17 | 58890618 | ATGTATGTGTGTA-TATAAACA-TACACACATATAT | loxF8L | 4 | loxF8L | 3 | TATAAACA/TGTTTATA | 7 |
| --- | | | | | | | | 0 |
| chr4 | 159921782 | ATATAACTGTGAA-TGGTTATA-TGTACATACTTAT | loxF8L | 3 | loxF8L | 4 | TATAACCA/TGGTTATA | 7 |
| chr3 | 181513691 | TTAAAATTTTGTG-TGGTTATA-TCCGAAGATTTTG | loxF8R | 3 | loxF8L | 4 | TATAACCA/TGGTTATA | 7 |
| --- | | | | | | | | 0 |
| chr7 | 79446388 | ACATATCTGTGTT-TGGGTATA-TCCAATGATATAT | loxF8L | 4 | loxF8L | 3 | TATACCCA/TGGGTATA | 7 |
| chr4 | 65427085 | ATATATATGCGAA-TATACCCA-CACACAATTTCAT | loxF8L | 4 | loxF8R | 3 | TATACCCA/TGGGTATA | 7 |
| --- | | | | | | | | 0 |
| chr11 | 118355876 | ATATATGTGTGTG-TATACGCA-TACACACATATAT | loxF8L | 4 | loxF8L | 3 | TATACGCA/TGCGTATA | 7 |
| chr15 | 81258780 | ATATATTTGTGCA-TATACGCA-CACACAACATGTG | loxF8L | 3 | loxF8R | 4 | TATACGCA/TGCGTATA | 7 |
| --- | | | | | | | | 0 |
| chr5 | 102377640 | ATATATCTATAGA-TATAGACA-TAGACATATATAT | loxF8L | 3 | loxF8L | 4 | TATAGACA/TGTCTATA | 7 |
| chr4 | 111462548 | ATATGTGTGTAGA-TATAGACA-TACACACATATAT | loxF8L | 4 | loxF8L | 3 | TATAGACA/TGTCTATA | 7 |
| --- | | | | | | | | 0 |
| chr10 | 64478160 | GTATAAATGTGTG-TATATCCA-GACACATATTTAT | loxF8R | 4 | loxF8L | 3 | TATATCCA/TGGATATA | 7 |
| chrX | 68501612 | ATATATAAGTGGA-TATATCCA-TTCATATATATAT | loxF8L | 3 | loxF8L | 4 | TATATCCA/TGGATATA | 7 |
| --- | | | | | | | | 0 |
| chr6 | 64268308 | ATAAATAAGTGTA-TATATGAA-TAGACAACATTAG | loxF8L | 3 | loxF8R | 4 | TATATGAA/TTCATATA | 7 |
| chr5 | 59737793 | GTATATATGTGCA-TATATGAA-TCCATATAGTTAT | loxF8L | 4 | loxF8L | 3 | TATATGAA/TTCATATA | 7 |
| chr5 | 27701917 | CTAAAATTCTGTC-TATATGAA-AACAAAAAATTAG | loxF8R | 3 | loxF8R | 4 | TATATGAA/TTCATATA | 7 |
| --- | | | | | | | | 0 |
| chr4 | 113585414 | ATACATGTGTGTA-TATATGCA-CACACATATTTGT | loxF8L | 3 | loxF8L | 4 | TATATGCA/TGCATATA | 7 |

| | | | | | | | | |
|---|---|---|---|---|---|---|---|---|
| chr8 | 71067676 | ACACATCTGTGTA-TGCATATA-TACACACACATAT | loxF8L | 3 | loxF8L | 4 | TATATGCA/TGCATATA | 7 |
| chr13 | 81926530 | ACATATATGTGTA-TATATGCA-TACACACATATAT | loxF8L | 4 | loxF8L | 3 | TATATGCA/TGCATATA | 7 |
| chr16 | 79945618 | ATAAATATATATA-TGCATATA-TCCAAATATATAT | loxF8L | 4 | loxF8L | 3 | TATATGCA/TGCATATA | 7 |
| chr2 | 34239556 | ATAAATGTATGTA-TATATGCA-CACACATACATAC | loxF8L | 3 | loxF8R | 4 | TATATGCA/TGCATATA | 7 |
| chr7 | 27585168 | ATATATGTGTGTA-TATATGCA-CACACATATGTAT | loxF8L | 3 | loxF8L | 4 | TATATGCA/TGCATATA | 7 |
| chr18 | 22395241 | ATATGTCTGTGTA-TGCATATA-TAAACATATATAT | loxF8L | 3 | loxF8L | 4 | TATATGCA/TGCATATA | 7 |
| chrX | 135621334 | ATATATGTGTGTA-TATATGCA-TACATAATGTTAA | loxF8L | 3 | loxF8R | 4 | TATATGCA/TGCATATA | 7 |
| chrX | 135667648 | TTAACATTATGTA-TGCATATA-TACACACATATAT | loxF8R | 4 | loxF8L | 3 | TATATGCA/TGCATATA | 7 |
| chr2 | 203968517 | GTAAATGTGTGTA-TATATGCA-TCCACATACACAT | loxF8L | 3 | loxF8L | 4 | TATATGCA/TGCATATA | 7 |
| --- | | | | | | | | 0 |
| chr7 | 97579504 | ATACATATGTGTA-TATATTCA-CACATATATTTAT | loxF8L | 3 | loxF8L | 4 | TATATTCA/TGAATATA | 7 |
| chr7 | 97579197 | ATACATATGTGTA-TATATTCA-CACATATATTTAT | loxF8L | 3 | loxF8L | 4 | TATATTCA/TGAATATA | 7 |
| chr7 | 97579233 | ATACATATGTGTA-TATATTCA-CACATATATTTAT | loxF8L | 3 | loxF8L | 4 | TATATTCA/TGAATATA | 7 |
| chr7 | 97579161 | ATACATATGTGTA-TATATTCA-CACATATATTTAT | loxF8L | 3 | loxF8L | 4 | TATATTCA/TGAATATA | 7 |
| chr7 | 97579364 | ATACATATGTGTA-TATATTCA-CACATATATTTAT | loxF8L | 3 | loxF8L | 4 | TATATTCA/TGAATATA | 7 |
| chr22 | 27041011 | ATATATATATGTA-TATATTCA-CACACAGATATAT | loxF8L | 4 | loxF8L | 3 | TATATTCA/TGAATATA | 7 |
| chr6 | 104727690 | ATATATATGTATA-TGAATATA-TACACACATATAT | loxF8L | 4 | loxF8L | 3 | TATATTCA/TGAATATA | 7 |
| chr6 | 104727718 | ATATATATGTATA-TGAATATA-TACACACATATAT | loxF8L | 4 | loxF8L | 3 | TATATTCA/TGAATATA | 7 |
| --- | | | | | | | | 0 |
| chr2 | 60835449 | ATAAATCCAGGGA-TCAATATA-AAAACAGATTTTT | loxF8L | 3 | loxF8L | 4 | TATATTGA/TCAATATA | 7 |
| chr11 | 121313818 | ATAAGATTTTGTA-TATATTGA-CAAAAATTCTTTG | loxF8R | 3 | loxF8R | 4 | TATATTGA/TCAATATA | 7 |
| --- | | | | | | | | 0 |
| chr3 | 35655232 | ATATATGTGTGTA-TATTGATA-TCTACATTTATAT | loxF8L | 3 | loxF8L | 4 | TATCAATA/TATTGATA | 7 |
| chr5 | 41411809 | ATATATATGTCCA-TATTGATA-TCCATATATATAT | loxF8L | 4 | loxF8L | 3 | TATCAATA/TATTGATA | 7 |
| --- | | | | | | | | 0 |
| chr7 | 74137692 | ATATATCTATAGA-TATCGATA-TCTATAGATATAG | loxF8L | 3 | loxF8L | 4 | TATCGATA/TATCGATA | 7 |
| --- | | | | | | | | 0 |
| chr15 | 98633015 | GCAAAATTGTGTG-TATGAATA-GAGACAGATTTTT | loxF8R | 3 | loxF8L | 4 | TATGAATA/TATTCATA | 7 |
| chr5 | 38820695 | ATACATATGTATA-TATGAATA-TACACATATGTAT | loxF8L | 4 | loxF8L | 3 | TATGAATA/TATTCATA | 7 |
| chr1 | 195473276 | ATATATATGTGTA-TATTCATA-TATACATATATAT | loxF8L | 3 | loxF8L | 4 | TATGAATA/TATTCATA | 7 |
| chr12 | 69849214 | ATATATATGTGCA-TATGAATA-TATATAATCTTAA | loxF8L | 3 | loxF8R | 4 | TATGAATA/TATTCATA | 7 |
| --- | | | | | | | | 0 |
| chr14 | 33237703 | ATATATCTATAGA-TTTGCATA-TACATATATTTTT | loxF8L | 3 | loxF8L | 4 | TATGCAAA/TTTGCATA | 7 |
| chr18 | 1871595 | ATAAATTTGTTAA-TTTGCATA-GCAGCAGAATTAT | loxF8L | 3 | loxF8L | 4 | TATGCAAA/TTTGCATA | 7 |
| chr3 | 56839211 | ATACATTTGTGCA-TTTGCATA-CACACATTTTAAA | loxF8L | 3 | loxF8R | 4 | TATGCAAA/TTTGCATA | 7 |
| --- | | | | | | | | 0 |
| chr4 | 122716602 | ACACATGTGTGTA-TATGCACA-TACACATATGTAT | loxF8L | 4 | loxF8L | 3 | TATGCACA/TGTGCATA | 7 |
| chr11 | 102807518 | ATAAAATTGTTTG-TATGCACA-TACTCAATTTTAT | loxF8R | 3 | loxF8L | 4 | TATGCACA/TGTGCATA | 7 |
| chr3 | 743016 | ATAAATATATGTA-TATGCACA-CACACACATATAT | loxF8L | 3 | loxF8L | 4 | TATGCACA/TGTGCATA | 7 |
| chr2 | 156891635 | ATAGCAGTGTGTG-TGTGCATA-CACACATTTATAG | loxF8R | 4 | loxF8R | 3 | TATGCACA/TGTGCATA | 7 |
| chr5 | 30406500 | ATAGGGCTGTGTG-TGTGCATA-CACACACTCATAC | loxF8R | 4 | loxF8R | 3 | TATGCACA/TGTGCATA | 7 |
| chr4 | 19214347 | ATATGTTTGTGTG-TGTGCATA-TATACATATATAT | loxF8R | 3 | loxF8L | 4 | TATGCACA/TGTGCATA | 7 |
| chr18 | 66237558 | CTAAGAATGTATT-TGTGCATA-TATACACATATAT | loxF8R | 3 | loxF8L | 4 | TATGCACA/TGTGCATA | 7 |
| --- | | | | | | | | 0 |
| chr17 | 12703200 | AGAATTCTGTGGC-TATGTAAA-TCAACATTTTTCT | loxF8L | 3 | loxF8L | 4 | TATGTAAA/TTTACATA | 7 |
| chr11 | 91374544 | GTAAATTTATGTA-TTTACATA-TTCACTTATTTAT | loxF8L | 4 | loxF8L | 3 | TATGTAAA/TTTACATA | 7 |
| chr2 | 64576160 | ATATATATATGGA-TTTACATA-TACATATATATAT | loxF8L | 3 | loxF8L | 4 | TATGTAAA/TTTACATA | 7 |
| chr13 | 32778026 | GTATTATTTTGTG-TATGTAAA-TCAACATATATAT | loxF8R | 4 | loxF8L | 3 | TATGTAAA/TTTACATA | 7 |
| chr11 | 4780890 | ATGTATCTGTAGA-TATGTAAA-TTTACATATTTAA | loxF8L | 3 | loxF8L | 4 | TATGTAAA/TTTACATA | 7 |
| --- | | | | | | | | 0 |
| chr8 | 82686045 | ATAAAGCAGTAAA-TATTAAAA-TTCACATATTTTT | loxF8L | 4 | loxF8L | 3 | TATTAAAA/TTTTAATA | 7 |
| chr17 | 37136124 | ACAAGTCTGTGAA-TTTTAATA-CACATAGTCATAT | loxF8L | 3 | loxF8R | 4 | TATTAAAA/TTTTAATA | 7 |
| chr2 | 69909255 | ACAATTCAGTGGT-TTTTAATA-TACTCAGAGTTAT | loxF8L | 4 | loxF8L | 3 | TATTAAAA/TTTTAATA | 7 |
| --- | | | | | | | | 0 |
| chr7 | 26756058 | ATAGATCTGTGTG-TATTAATA-TGGAAAGATATAT | loxF8L | 3 | loxF8L | 4 | TATTAATA/TATTAATA | 7 |
| --- | | | | | | | | 0 |
| chr12 | 81935422 | ATGAGTCTACGGA-TGGTAATA-TCGACAGAAGTAT | loxF8L | 4 | loxF8L | 3 | TATTACCA/TGGTAATA | 7 |
| chr12 | 102314709 | CTAAGATTCTATT-TGGTAATA-TCCAATGTTTTAA | loxF8R | 3 | loxF8L | 4 | TATTACCA/TGGTAATA | 7 |
| --- | | | | | | | | 0 |
| chr2 | 152732154 | ATAAAATTGTATA-TTATAATA-TACAGAGATATAT | loxF8L | 4 | loxF8L | 3 | TATTATAA/TTATAATA | 7 |
| chr3 | 20385461 | CTAAGATTCTATT-TTATAATA-TCCAATGTTTTAA | loxF8R | 3 | loxF8L | 4 | TATTATAA/TTATAATA | 7 |
| --- | | | | | | | | 0 |
| chr1 | 232642430 | ATATATGTATGTA-TATTTCCA-TACACAGGTTTTT | loxF8L | 4 | loxF8L | 3 | TATTTCCA/TGGAAATA | 7 |
| chr1 | 70498640 | CTAATTTTGTATT-TGGAAATA-ATCATAGATTTAT | loxF8R | 4 | loxF8L | 3 | TATTTCCA/TGGAAATA | 7 |
| --- | | | | | | | | 0 |
| chr1 | 218015624 | CTAAGACTTTGTA-TTTTTTGA-TCAAAAGTTTTTT | loxF8R | 3 | loxF8L | 4 | TCAAAAAA/TTTTTTGA | 7 |
| chr4 | 90320384 | ATAAAGCTCTGGC-TTTTTTGA-CACTAAATCCTTG | loxF8L | 3 | loxF8R | 4 | TCAAAAAA/TTTTTTGA | 7 |
| chr8 | 140649673 | ATAAATATATGTA-TTTTTTGA-CACAGAGTCTCAC | loxF8L | 3 | loxF8R | 4 | TCAAAAAA/TTTTTTGA | 7 |
| chr1 | 146026155 | CAAAGATTCTTTT-TTTTTTGA-GACAGAATCTTAC | loxF8R | 4 | loxF8R | 3 | TCAAAAAA/TTTTTTGA | 7 |
| --- | | | | | | | | 0 |
| chr10 | 9423006 | ATAAATCTATCTA-TCAATAGA-TATACAGATAGAT | loxF8L | 3 | loxF8L | 4 | TCAATAGA/TCTATTGA | 7 |
| chr12 | 117506798 | ATATGTCAGTAGA-TCAATAGA-TAGATAGATTTAT | loxF8L | 4 | loxF8L | 3 | TCAATAGA/TCTATTGA | 7 |
| chr4 | 34431773 | ATAATGCTGTGTG-TCAATAGA-GACACAATATTAT | loxF8L | 4 | loxF8R | 3 | TCAATAGA/TCTATTGA | 7 |
| chr4 | 100890207 | ATTGATCTCTGGA-TCAATAGA-TCCACATCTAGAT | loxF8L | 3 | loxF8L | 4 | TCAATAGA/TCTATTGA | 7 |
| --- | | | | | | | | 0 |
| chr7 | 63663680 | ATAAAACAGATGA-TGAATTGA-TAGACAGATCTAT | loxF8L | 4 | loxF8L | 3 | TCAATTCA/TGAATTGA | 7 |
| chr7 | 56913231 | ATAGATCTGTCTA-TCAATTCA-TCATCTGTTTTAT | loxF8L | 3 | loxF8L | 4 | TCAATTCA/TGAATTGA | 7 |
| chr2 | 138306082 | TTAATTCTGTGAT-TCAATTCA-TCCACTGACTTAC | loxF8L | 4 | loxF8L | 3 | TCAATTCA/TGAATTGA | 7 |
| --- | | | | | | | | 0 |
| chr11 | 25804871 | ATAACTATGTAAA-TTAAGTGA-CTTACAGATTTAT | loxF8L | 4 | loxF8L | 3 | TCACTTAA/TTAAGTGA | 7 |
| chr11 | 25802540 | ATAACTATGTAAA-TTAAGTGA-CTTACAGATTTAT | loxF8L | 4 | loxF8L | 3 | TCACTTAA/TTAAGTGA | 7 |
| --- | | | | | | | | 0 |
| chr11 | 61222214 | CTCAGAATGTGGC-TCAGGGCA-GCCACAATCTTAT | loxF8R | 4 | loxF8R | 3 | TCAGGGCA/TGCCCTGA | 7 |
| chr11 | 61241040 | CTCAGAATGTGGC-TCAGGGCA-GCCACAATCTTAT | loxF8R | 4 | loxF8R | 3 | TCAGGGCA/TGCCCTGA | 7 |
| --- | | | | | | | | 0 |
| chrX | 120161550 | ATAAATTTGTAGG-TCAGTGGA-TCCAAAGGAGTAT | loxF8L | 3 | loxF8L | 4 | TCAGTGGA/TCCACTGA | 7 |
| chrX | 120074718 | ATACTCCTTTGGA-TCCACTGA-CCTACAAATTTAT | loxF8L | 4 | loxF8L | 3 | TCAGTGGA/TCCACTGA | 7 |

| | | | | | | | | |
|---|---|---|---|---|---|---|---|---|
| --- | | | | | | | | 0 |
| chr11 | 103035527 | ATTATTTTTTGGA-TTTTATGA-TTCACACATTTTT | loxF8L | 4 | loxF8L | 3 | TCATAAAA/TTTTATGA | 7 |
| chr14 | 79177489 | ATAAATTTTTGTG-TCATAAAA-TGCACAAATTAAT | loxF8L | 4 | loxF8L | 3 | TCATAAAA/TTTTATGA | 7 |
| --- | | | | | | | | 0 |
| chrX | 119997032 | AGAAATCTTTTGA-TCCCATGA-CCTACAAATTAAT | loxF8L | 3 | loxF8L | 4 | TCATGGGA/TCCCATGA | 7 |
| chr13 | 97101236 | ATTGATCTGTGTA-TCATGGGA-CAAACAGATTTAC | loxF8L | 3 | loxF8L | 4 | TCATGGGA/TCCCATGA | 7 |
| --- | | | | | | | | 0 |
| chr10 | 20950929 | ATAATTCTAAGGA-TGCAATGA-TACAAAGCTTAAT | loxF8L | 3 | loxF8L | 4 | TCATTGCA/TGCAATGA | 7 |
| chr13 | 42573182 | TTTGATCTGTGGT-TCATTGCA-TCCACAGATGTGG | loxF8L | 4 | loxF8L | 3 | TCATTGCA/TGCAATGA | 7 |
| --- | | | | | | | | 0 |
| chr17 | 27681371 | GCCAATCTGTGGA-TCATTTAA-AACTCAATTTTAT | loxF8L | 3 | loxF8R | 4 | TCATTTAA/TTAAATGA | 7 |
| chr17 | 30483957 | TCCAATCTGTGGA-TCATTTAA-ACCTCAATCTTAT | loxF8L | 3 | loxF8R | 4 | TCATTTAA/TTAAATGA | 7 |
| --- | | | | | | | | 0 |
| chr2 | 169928303 | ATGAAATTTTGTG-TTTTTGGA-AAAACAATCTTTG | loxF8R | 4 | loxF8R | 3 | TCCAAAAA/TTTTTGGA | 7 |
| chr6 | 88516424 | CTCAGAGTCTGTT-TCCAAAAA-AACACAACCTTAA | loxF8R | 4 | loxF8R | 3 | TCCAAAAA/TTTTTGGA | 7 |
| --- | | | | | | | | 0 |
| chr12 | 109814268 | ATAGATGTATGGA-TGGTTGGA-TGGACAGATGGAT | loxF8L | 3 | loxF8L | 4 | TCCAACCA/TGGTTGGA | 7 |
| chr6 | 22608931 | CTGCATCTATGGA-TCCAACCA-ACCACAGATTGAA | loxF8L | 4 | loxF8L | 3 | TCCAACCA/TGGTTGGA | 7 |
| chr2 | 101366454 | TTGCATCTGTGGA-TCCAACCA-ACCACAGACTGAA | loxF8L | 3 | loxF8L | 4 | TCCAACCA/TGGTTGGA | 7 |
| --- | | | | | | | | 0 |
| chr16 | 26837341 | ATGAGGTTGTGTG-TGTGTGGA-GACATGGATTTAT | loxF8R | 3 | loxF8L | 4 | TCCACACA/TGTGTGGA | 7 |
| chr10 | 64628166 | CTAAGATCTTATG-TGTGTGGA-CACACATACTGTG | loxF8R | 3 | loxF8R | 4 | TCCACACA/TGTGTGGA | 7 |
| --- | | | | | | | | 0 |
| chr4 | 53770110 | ATAAATGAATGGA-TGGGTGGA-TGGACAGATGAAT | loxF8L | 3 | loxF8L | 4 | TCCACCCA/TGGGTGGA | 7 |
| chr3 | 10368688 | ATCCATCTGTCCA-TCCACCCA-TCCACCCATCTAT | loxF8L | 4 | loxF8L | 3 | TCCACCCA/TGGGTGGA | 7 |
| --- | | | | | | | | 0 |
| chrX | 141805512 | ATGGATAGGTGGA-TTGATGGA-TGGACAGATTAAT | loxF8L | 4 | loxF8L | 3 | TCCATCAA/TTGATGGA | 7 |
| chr9 | 122602306 | TTAAAATTTTATG-TCCATCAA-AAGACAATGTTAG | loxF8R | 4 | loxF8R | 3 | TCCATCAA/TTGATGGA | 7 |
| chr13 | 104057621 | TTAGCATTGTGTG-TCCATCAA-CACACATTTATAA | loxF8R | 3 | loxF8R | 4 | TCCATCAA/TTGATGGA | 7 |
| --- | | | | | | | | 0 |
| chr7 | 158987795 | ATCCATCTGTGCA-TCCATTCA-TCCATCCATTCAT | loxF8L | 3 | loxF8L | 4 | TCCATTCA/TGAATGGA | 7 |
| chr7 | 15630117 | ATGCACCTGTGGG-TCCATTCA-CACACTCTCTCAG | loxF8L | 4 | loxF8R | 3 | TCCATTCA/TGAATGGA | 7 |
| --- | | | | | | | | 0 |
| chr2 | 112453653 | ATATATCTATATA-TCCTAGGA-TACATAGATATAT | loxF8L | 4 | loxF8L | 3 | TCCTAGGA/TCCTAGGA | 7 |
| chr3 | 67353687 | CTAACTTTGTATG-TCCTAGGA-AACACAGATTTGG | loxF8R | 3 | loxF8L | 4 | TCCTAGGA/TCCTAGGA | 7 |
| --- | | | | | | | | 0 |
| chrX | 109009138 | ATTATTTTGTGGA-TCCTGACA-AACTCAATCTAAA | loxF8L | 3 | loxF8R | 4 | TCCTGACA/TGTCAGGA | 7 |
| chr4 | 100943550 | CGATGATTTTGTG-TCCTGACA-CTTACAGTATTAG | loxF8R | 3 | loxF8R | 4 | TCCTGACA/TGTCAGGA | 7 |
| --- | | | | | | | | 0 |
| chr4 | 167738135 | TTGAATCCGTGGG-TGACAGGA-TCCATACATTTCT | loxF8L | 4 | loxF8L | 3 | TCCTGTCA/TGACAGGA | 7 |
| chr22 | 33349786 | ATAACTCTTTGGC-TCCTGTCA-CACAGATTCCTAT | loxF8L | 3 | loxF8R | 4 | TCCTGTCA/TGACAGGA | 7 |
| --- | | | | | | | | 0 |
| chr13 | 51844215 | ATATATGTGTGTA-TGTGTAGA-CACACACACATAT | loxF8L | 3 | loxF8R | 4 | TCTACACA/TGTGTAGA | 7 |
| chrX | 29524319 | ATATATTTGTGTA-TGTGTAGA-TACACACATACAT | loxF8L | 3 | loxF8L | 4 | TCTACACA/TGTGTAGA | 7 |
| --- | | | | | | | | 0 |
| chr2 | 89927877 | ATAAATCTGTCCT-TCTATCCA-TCCATCTATGTAT | loxF8L | 3 | loxF8L | 4 | TCTATCCA/TGGATAGA | 7 |
| chr2 | 89254525 | ATACATAGATGGA-TGGATAGA-AGGACAGATTTAT | loxF8L | 4 | loxF8L | 3 | TCTATCCA/TGGATAGA | 7 |
| chr13 | 97201458 | ATAAATGGATGGA-TGGATAGA-TAGACAAATATAT | loxF8L | 3 | loxF8L | 4 | TCTATCCA/TGGATAGA | 7 |
| chr10 | 78891285 | ATGAATATATGGA-TGGATAGA-TGAACAGATGGAT | loxF8L | 3 | loxF8L | 4 | TCTATCCA/TGGATAGA | 7 |
| chr12 | 113438960 | ATTAGACTGTCTG-TCTATCCA-TCCACATATCCAT | loxF8R | 4 | loxF8L | 3 | TCTATCCA/TGGATAGA | 7 |
| --- | | | | | | | | 0 |
| chr6 | 54595205 | TTCAATCTCTGGA-TGAATAGA-TAAATAGATTTCT | loxF8L | 3 | loxF8L | 4 | TCTATTCA/TGAATAGA | 7 |
| chr1 | 17612435 | ATTTATCTGTGCA-TCTATTCA-TTCATCCATTTAT | loxF8L | 3 | loxF8L | 4 | TCTATTCA/TGAATAGA | 7 |
| chr7 | 158987127 | ATGCATCTGTCCA-TCTATTCA-TCCATAAATTCAT | loxF8L | 4 | loxF8L | 3 | TCTATTCA/TGAATAGA | 7 |
| chr7 | 158987357 | ATGCATCTGTCCA-TCTATTCA-TCCATAAATTCAT | loxF8L | 4 | loxF8L | 3 | TCTATTCA/TGAATAGA | 7 |
| --- | | | | | | | | 0 |
| chrX | 34759910 | ATAAATCTGTTAT-TTCTGAGA-TCCAAAGAAATAA | loxF8L | 3 | loxF8L | 4 | TCTCAGAA/TTCTGAGA | 7 |
| chr7 | 82512731 | TTTAAGGTGTGGA-TTCTGAGA-CAGACAATCCTAA | loxF8L | 4 | loxF8R | 3 | TCTCAGAA/TTCTGAGA | 7 |
| --- | | | | | | | | 0 |
| chr13 | 100706378 | ATAAAACTGTCCA-TGATGAGA-TGCTCAGTTTTTT | loxF8L | 3 | loxF8L | 4 | TCTCATCA/TGATGAGA | 7 |
| chr2 | 194063034 | ATATATCTATCTA-TCTCATCA-CCCATATATTTAT | loxF8L | 4 | loxF8L | 3 | TCTCATCA/TGATGAGA | 7 |
| --- | | | | | | | | 0 |
| chr12 | 102695218 | ACAAGATTCTCTG-TCTGGTAA-TCCAGAGAATTCT | loxF8R | 4 | loxF8L | 3 | TCTGGTAA/TTACCAGA | 7 |
| chr11 | 74213673 | GTAAGAATCTCTG-TCTGGTAA-TCCAGAGAATTCT | loxF8R | 4 | loxF8L | 3 | TCTGGTAA/TTACCAGA | 7 |
| chrY | 6204560 | ATAATGCTCTGGA-TTACCAGA-CAGAGACTCTTAT | loxF8L | 3 | loxF8R | 4 | TCTGGTAA/TTACCAGA | 7 |
| --- | | | | | | | | 0 |
| chr3 | 158138998 | ATAAATCTGTTTC-TTAAAAGA-TCATCATATTTGT | loxF8L | 3 | loxF8L | 4 | TCTTTTAA/TTAAAAGA | 7 |
| chr3 | 191706575 | ATAAATGTGTACA-TTAAAAGA-AAAACAGAATTAT | loxF8L | 3 | loxF8L | 4 | TCTTTTAA/TTAAAAGA | 7 |
| --- | | | | | | | | 0 |
| chr1 | 154982508 | ATCAATCTGCATA-TGAAAAGA-TCCCCAGATTATT | loxF8L | 4 | loxF8L | 3 | TCTTTTCA/TGAAAAGA | 7 |
| chr10 | 109584530 | GTAAATATGTGCA-TGAAAAGA-TACTCAATATTAT | loxF8L | 3 | loxF8R | 4 | TCTTTTCA/TGAAAAGA | 7 |
| chr2 | 236662826 | ATAATTCTCTGGG-TGAAAAGA-TCCATAGAGCTAA | loxF8L | 3 | loxF8L | 4 | TCTTTTCA/TGAAAAGA | 7 |
| chr3 | 17442566 | CTAATTCTGTGTA-TGAAAAGA-CACAAATCCTGAG | loxF8L | 3 | loxF8R | 4 | TCTTTTCA/TGAAAAGA | 7 |
| --- | | | | | | | | 0 |
| chrX | 5344101 | TTAGATCTTTAGA-TTTATTCA-TCTACATATTTGT | loxF8L | 4 | loxF8L | 3 | TGAATAAA/TTTATTCA | 7 |
| chr4 | 96757295 | ACAAATGAATGGA-TTTATTCA-TCCACACATTAGT | loxF8L | 4 | loxF8L | 3 | TGAATAAA/TTTATTCA | 7 |
| chr4 | 53400048 | ATAGTTTTGTGTA-TGAATAAA-GAGACAATCTTAT | loxF8L | 4 | loxF8R | 3 | TGAATAAA/TTTATTCA | 7 |
| chrY | 16181644 | GCAAATATGTGGA-TGAATAAA-TCTAAAGATCTAA | loxF8L | 3 | loxF8L | 4 | TGAATAAA/TTTATTCA | 7 |
| chrY | 16403665 | TTAGATCTTTAGA-TTTATTCA-TCCACATATTTGC | loxF8L | 4 | loxF8L | 3 | TGAATAAA/TTTATTCA | 7 |
| --- | | | | | | | | 0 |
| chr8 | 132413063 | AAAAATGAGTGAA-TGAATGAA-TGCACAAACTTAT | loxF8L | 4 | loxF8L | 3 | TGAATGAA/TTCATTCA | 7 |
| chr16 | 47182501 | CTTAGTTTGTGTA-TTCATTCA-TCTAAACATTTAG | loxF8R | 3 | loxF8L | 4 | TGAATGAA/TTCATTCA | 7 |
| --- | | | | | | | | 0 |
| chr11 | 49369330 | ATAAATCTAGGCA-TGAATGCA-TTCACAAATTTTA | loxF8L | 3 | loxF8L | 4 | TGAATGCA/TGCATTCA | 7 |
| chr11 | 89336872 | TAAAATTTGTGAA-TGCATTCA-TGCCTAGATTTAT | loxF8L | 4 | loxF8L | 3 | TGAATGCA/TGCATTCA | 7 |
| chr18 | 73478639 | ATACATATGTGCG-TGAATGCA-TACACACATGTAT | loxF8L | 4 | loxF8L | 3 | TGAATGCA/TGCATTCA | 7 |
| chr7 | 93771320 | TTAAAAATGTGTT-TGAATGCA-CACACACACTCAG | loxF8R | 4 | loxF8R | 3 | TGAATGCA/TGCATTCA | 7 |

| | | | | | | | | |
|---|---|---|---|---|---|---|---|---|
| --- | | | | | | | | 0 |
| chr21 | 20866536 | GTATGGGTGTGTG-TGAATTCA-TACACAAACTTAA | loxF8R | 4 | loxF8R | 3 | TGAATTCA/TGAATTCA | 7 |
| --- | | | | | | | | 0 |
| chr4 | 13457502 | ATAACATTCTGTG-TGACTGCA-TACACTAATATAT | loxF8R | 3 | loxF8L | 4 | TGACTGCA/TGCAGTCA | 7 |
| chr7 | 112828025 | TTTGATCTGTGGT-TGACTGCA-TCCACAGATGTGG | loxF8L | 4 | loxF8L | 3 | TGACTGCA/TGCAGTCA | 7 |
| --- | | | | | | | | 0 |
| chr5 | 70051745 | GCAAGATTGTGTA-TGAGGGCA-CCCACAGAGTAAA | loxF8R | 3 | loxF8L | 4 | TGAGGGCA/TGCCCTCA | 7 |
| chr5 | 70927163 | GCAAGATTGTGTA-TGAGGGCA-CCCACAGAGTAAA | loxF8R | 3 | loxF8L | 4 | TGAGGGCA/TGCCCTCA | 7 |
| --- | | | | | | | | 0 |
| chr12 | 33393317 | ATAAGCTTCTGGA-TGCAAAAA-ACCAGAGACTTAT | loxF8L | 4 | loxF8L | 3 | TGCAAAAA/TTTTTGCA | 7 |
| chr18 | 74861320 | GTAAATCTGTCCA-TGCAAAAA-TCCAAATGTTTAG | loxF8L | 3 | loxF8L | 4 | TGCAAAAA/TTTTTGCA | 7 |
| --- | | | | | | | | 0 |
| chr4 | 62043408 | CTGTGTCTGTGTG-TGTGTGCA-TGCACACATTTGT | loxF8R | 4 | loxF8L | 3 | TGCACACA/TGTGTGCA | 7 |
| chr6 | 24106226 | CAAATAATGTGTG-TGTGTGCA-TGCACACATGTGT | loxF8R | 3 | loxF8L | 4 | TGCACACA/TGTGTGCA | 7 |
| chrX | 42829939 | ATAAATCTCTTTA-TGCACACA-TGCACATATATAC | loxF8L | 3 | loxF8L | 4 | TGCACACA/TGTGTGCA | 7 |
| chr7 | 44150538 | ATAACTGTGTGTG-TGTGTGCA-TGCACAGATGTAA | loxF8L | 4 | loxF8L | 3 | TGCACACA/TGTGTGCA | 7 |
| chr20 | 26115435 | ATACACATGTGCA-TGTGTGCA-CATACACTCTTAC | loxF8L | 4 | loxF8R | 3 | TGCACACA/TGTGTGCA | 7 |
| chr11 | 38031254 | GTATGTATGTGTG-TGTGTGCA-TCCACACATATAA | loxF8R | 4 | loxF8L | 3 | TGCACACA/TGTGTGCA | 7 |
| --- | | | | | | | | 0 |
| chr4 | 42296143 | ATACATATGTATA-TGCACGCA-TACACATATGTAT | loxF8L | 4 | loxF8L | 3 | TGCACGCA/TGCGTGCA | 7 |
| chr11 | 96291717 | GTAAGTATGTGTG-TGCACGCA-CACACATGCATAC | loxF8R | 3 | loxF8R | 4 | TGCACGCA/TGCGTGCA | 7 |
| --- | | | | | | | | 0 |
| chr3 | 179983128 | ATACATGTATGCA-TGCATAAA-TATACATATTTAT | loxF8L | 4 | loxF8L | 3 | TGCATAAA/TTTATGCA | 7 |
| chr4 | 149695179 | CAGACATTGTGTT-TTTATGCA-TCCATCCATTTAT | loxF8R | 4 | loxF8L | 3 | TGCATAAA/TTTATGCA | 7 |
| chr12 | 21403196 | CTGTATCTGTGCA-TGCATAAA-CAAACAATATTAT | loxF8L | 4 | loxF8R | 3 | TGCATAAA/TTTATGCA | 7 |
| chr11 | 79924771 | ATGAAACTATGTA-TGCATAAA-CACACAAACATAT | loxF8L | 4 | loxF8R | 3 | TGCATAAA/TTTATGCA | 7 |
| --- | | | | | | | | 0 |
| chr6 | 26761928 | ATACATGTGTGCA-TGCATGCA-TACACACATGCAT | loxF8L | 3 | loxF8L | 4 | TGCATGCA/TGCATGCA | 7 |
| --- | | | | | | | | 0 |
| chrY | 16808846 | CAAAATCTGTGTA-TTGTAGCA-TCAGATGATTTAT | loxF8L | 3 | loxF8L | 4 | TGCTACAA/TTGTAGCA | 7 |
| chr8 | 78954883 | ATAACTCTGAGTA-TGCTACAA-ACCATTGAATTAT | loxF8L | 3 | loxF8L | 4 | TGCTACAA/TTGTAGCA | 7 |
| --- | | | | | | | | 0 |
| chr10 | 11438438 | ATGAATGAGTGGA-TGGAGAAA-GACATAAACATAG | loxF8L | 3 | loxF8R | 4 | TGGAGAAA/TTTCTCCA | 7 |
| chr10 | 3157199 | GTAATTCTTTGGA-TGGAGAAA-CAAAAAAACTTAC | loxF8L | 3 | loxF8R | 4 | TGGAGAAA/TTTCTCCA | 7 |
| --- | | | | | | | | 0 |
| chr17 | 77861260 | ATAGATGAGTAGA-TGGATAAA-TGGACAGATTGAT | loxF8L | 4 | loxF8L | 3 | TGGATAAA/TTTATCCA | 7 |
| chr17 | 56892590 | ATCCATCTGTCCA-TTTATCCA-TCCACCCATCTAT | loxF8L | 4 | loxF8L | 3 | TGGATAAA/TTTATCCA | 7 |
| chr7 | 137090569 | CTAGGACTGTCTG-TTTATCCA-TTCACATGTTGAT | loxF8R | 3 | loxF8L | 4 | TGGATAAA/TTTATCCA | 7 |
| --- | | | | | | | | 0 |
| chr8 | 94928266 | ACAAATCTGTTCA-TGTTGCCA-AACAAAATCTCAT | loxF8L | 3 | loxF8R | 4 | TGGCAACA/TGTTGCCA | 7 |
| chr2 | 76978780 | GGAAATCTGGGGA-TGTTGCCA-CCTACAGATTGAG | loxF8L | 3 | loxF8L | 4 | TGGCAACA/TGTTGCCA | 7 |
| --- | | | | | | | | 0 |
| chr4 | 26004667 | ATAGATAGGTGGA-TGGCTGAA-TGCATAGATAGAT | loxF8L | 3 | loxF8L | 4 | TGGCTGAA/TTCAGCCA | 7 |
| chr14 | 65000274 | CTGCATCTGTGGA-TTCAGCCA-ACCACAGATCAAA | loxF8L | 3 | loxF8L | 4 | TGGCTGAA/TTCAGCCA | 7 |
| chr4 | 173674712 | TTGAATCTGTAGT-TGGCTGAA-TCCACAGATGCAG | loxF8L | 4 | loxF8L | 3 | TGGCTGAA/TTCAGCCA | 7 |
| --- | | | | | | | | 0 |
| chr18 | 24438899 | ATACATGTGTGTA-TGTACCCA-CACACAAAGAGAG | loxF8L | 3 | loxF8R | 4 | TGGGTACA/TGTACCCA | 7 |
| chr12 | 13957328 | ATATACATGTGTA-TGGGTACA-CACACAAGCTTGA | loxF8L | 4 | loxF8R | 3 | TGGGTACA/TGTACCCA | 7 |
| --- | | | | | | | | 0 |
| chr5 | 125902207 | CTAAGTTTTTGTT-TGGTTAAA-TCCTTTTATTTAT | loxF8R | 3 | loxF8L | 4 | TGGTTAAA/TTTAACCA | 7 |
| chr22 | 44935025 | ATATCTCTGTGGC-TGGTTAAA-CACACACGCTCAT | loxF8L | 3 | loxF8R | 4 | TGGTTAAA/TTTAACCA | 7 |
| chr5 | 42325769 | CAAAATTTGTGGA-TTTAACCA-AACACAGATTGAA | loxF8L | 3 | loxF8L | 4 | TGGTTAAA/TTTAACCA | 7 |
| --- | | | | | | | | 0 |
| chr5 | 38985375 | TTAGAACTGCGGA-TGGTTCAA-TCCACAGATGTGG | loxF8L | 4 | loxF8L | 3 | TGGTTCAA/TTGAACCA | 7 |
| chr6 | 68969337 | TCACATCTGTGGA-TTGAACCA-ACCACAGATCAAA | loxF8L | 3 | loxF8L | 4 | TGGTTCAA/TTGAACCA | 7 |
| --- | | | | | | | | 0 |
| chr2 | 219980535 | ATTAAATTGTGGA-TTTTTACA-TGGACAGATTATT | loxF8L | 3 | loxF8L | 4 | TGTAAAAA/TTTTTACA | 7 |
| chr11 | 76264666 | ATAAATCTATATA-TGTAAAAA-TATATATATTTAT | loxF8L | 3 | loxF8L | 4 | TGTAAAAA/TTTTTACA | 7 |
| --- | | | | | | | | 0 |
| chr11 | 114062074 | GTAAGTGTGTGTG-TGTAAACA-CACACACACTTTT | loxF8R | 3 | loxF8R | 4 | TGTAAACA/TGTTTACA | 7 |
| chr8 | 13506206 | ATAAATATGTATA-TGTAAACA-CACACATATATAT | loxF8L | 3 | loxF8L | 4 | TGTAAACA/TGTTTACA | 7 |
| chr7 | 131735552 | CTATGCCTGTGTG-TGTTTACA-CACACATTCACAT | loxF8R | 3 | loxF8R | 4 | TGTAAACA/TGTTTACA | 7 |
| --- | | | | | | | | 0 |
| chr3 | 42939047 | ATATATGTGTGTA-TGTGTACA-CACACACATTTCT | loxF8L | 3 | loxF8L | 4 | TGTACACA/TGTGTACA | 7 |
| chr10 | 13607402 | AGAAGTTTGTGTG-TGTGTACA-CACACGCTTTTTG | loxF8R | 3 | loxF8R | 4 | TGTACACA/TGTGTACA | 7 |
| chr11 | 126927337 | TTATGATTTTGTG-TGTGTACA-CACTCATACTTAT | loxF8R | 3 | loxF8R | 4 | TGTACACA/TGTGTACA | 7 |
| chr18 | 53987604 | ATATAATTGTGTG-TGTACACA-CACACACACATAT | loxF8R | 3 | loxF8R | 4 | TGTACACA/TGTGTACA | 7 |
| chr4 | 138050497 | GTAAAGTTGTGTG-TGTACACA-CACACATATATAT | loxF8R | 3 | loxF8L | 4 | TGTACACA/TGTGTACA | 7 |
| chr6 | 23629739 | TTATAATTATGTG-TGTGTACA-TCAATAGATATAT | loxF8R | 4 | loxF8L | 3 | TGTACACA/TGTGTACA | 7 |
| --- | | | | | | | | 0 |
| chr13 | 33044900 | AAAAATCTTAGGA-TTTCTACA-CAGACAATCATGT | loxF8L | 3 | loxF8R | 4 | TGTAGAAA/TTTCTACA | 7 |
| chr14 | 76245387 | ATTCATCTTTGGA-TTTCTACA-CACACACTCACAT | loxF8L | 3 | loxF8R | 4 | TGTAGAAA/TTTCTACA | 7 |
| --- | | | | | | | | 0 |
| chr6 | 166288828 | ATAAATATATAGA-TGTATAAA-TATATAGATATAT | loxF8L | 3 | loxF8L | 4 | TGTATAAA/TTTATACA | 7 |
| chr22 | 47981699 | ATAAATGTATAAA-TTTATACA-TTCATAAATTTAT | loxF8L | 4 | loxF8L | 3 | TGTATAAA/TTTATACA | 7 |
| chr13 | 108677504 | ATACCTGTGTGTA-TTTATACA-TGCACATATATAT | loxF8L | 4 | loxF8L | 3 | TGTATAAA/TTTATACA | 7 |
| chr2 | 142651226 | ATATGAGTGTGTG-TGTATAAA-TTAACATATATAT | loxF8R | 3 | loxF8L | 4 | TGTATAAA/TTTATACA | 7 |
| --- | | | | | | | | 0 |
| chr4 | 66242263 | ATCAGATTTTGTT-TTAATACA-CTCATAATCTAAG | loxF8R | 4 | loxF8R | 3 | TGTATTAA/TTAATACA | 7 |
| chr4 | 153782642 | TTAAAATTGCTTG-TTAATACA-CAAAGAATCTTAA | loxF8R | 4 | loxF8R | 3 | TGTATTAA/TTAATACA | 7 |
| --- | | | | | | | | 0 |
| chr3 | 168118735 | ATCTATGTGTGCA-TGTGAACA-CACAAAATCTTCT | loxF8L | 4 | loxF8R | 3 | TGTGAACA/TGTTCACA | 7 |
| chr2 | 195741133 | CTCAGTGTGTGTG-TGTGAACA-TTATCAAATTTAT | loxF8R | 3 | loxF8L | 4 | TGTGAACA/TGTTCACA | 7 |
| --- | | | | | | | | 0 |
| chr1 | 144383269 | CTAAGAAAGTTTG-TGTGTCAA-TAACCATATTTAT | loxF8R | 3 | loxF8L | 4 | TGTGTCAA/TTGACACA | 7 |
| chr1 | 144583220 | ATAAATATGGTTA-TTGACACA-CAAACTTTCTTAG | loxF8L | 4 | loxF8R | 3 | TGTGTCAA/TTGACACA | 7 |
| chr1 | 149574096 | ATAAATATGGTTA-TTGACACA-CAAACTTTCTTAG | loxF8L | 4 | loxF8R | 3 | TGTGTCAA/TTGACACA | 7 |

| | | | | | | | | |
|---|---|---|---|---|---|---|---|---|
| chr1 | 145251817 | CTAAGAAAGTTTG-TGTGTCAA-TAACCATATTTAT | loxF8R | 3 | loxF8L | 4 | TGTGTCAA/TTGACACA | 7 |
| chr1 | 148492958 | CTAAGAAAGTTTG-TGTGTCAA-TAACCATATTTAT | loxF8R | 3 | loxF8L | 4 | TGTGTCAA/TTGACACA | 7 |
| chr1 | 146354420 | CTAAGAAAGTTTG-TGTGTCAA-TAACCATATTTAT | loxF8R | 3 | loxF8L | 4 | TGTGTCAA/TTGACACA | 7 |
| --- | | | | | | | | 0 |
| chr3 | 8620764 | AAACATGTGTGTA-TTAAAACA-ACCACAGAGGTAT | loxF8L | 4 | loxF8L | 3 | TGTTTTAA/TTAAAACA | 7 |
| chr11 | 57039309 | TAAAATATGTGTA-TTAAAACA-TACACAGATGTGT | loxF8L | 4 | loxF8L | 3 | TGTTTTAA/TTAAAACA | 7 |
| --- | | | | | | | | 0 |
| chr18 | 28571568 | AAAAATTAGTGGC-TTAAAAAA-TCCACCATTTTAT | loxF8L | 4 | loxF8L | 3 | TTAAAAAA/TTTTTTAA | 7 |
| chr1 | 89705786 | ATAAATAAATGGT-TTTTTTAA-AAGACAGATTTAT | loxF8L | 4 | loxF8L | 3 | TTAAAAAA/TTTTTTAA | 7 |
| chrX | 142852303 | ATAAATTTGAAAA-TTAAAAAA-TTCACAGTTTTAA | loxF8L | 4 | loxF8L | 3 | TTAAAAAA/TTTTTTAA | 7 |
| --- | | | | | | | | 0 |
| chr4 | 76525653 | ATTAGATTGTTTT-TTTCTTAA-TGCAAAGATTTTT | loxF8R | 4 | loxF8L | 3 | TTAAGAAA/TTTCTTAA | 7 |
| chr7 | 156436053 | ATTAAATTGTATG-TTAAGAAA-TACATAAATTTAT | loxF8R | 4 | loxF8L | 3 | TTAAGAAA/TTTCTTAA | 7 |
| --- | | | | | | | | 0 |
| chr1 | 90505729 | ATAAATTAATGTA-TTAATAAA-TCAACTGATTTTT | loxF8L | 4 | loxF8L | 3 | TTAATAAA/TTTATTAA | 7 |
| chr7 | 31844218 | ATAAATATTTAGA-TTTATTAA-TCTAAAAATTAAT | loxF8L | 3 | loxF8L | 4 | TTAATAAA/TTTATTAA | 7 |
| chr14 | 71416159 | ATAAATCTATAGT-TTTATTAA-CACAGAAACTGAT | loxF8L | 3 | loxF8R | 4 | TTAATAAA/TTTATTAA | 7 |
| chr4 | 125580575 | CAAAGACTCTGTG-TTAATAAA-TACACACATACAT | loxF8R | 3 | loxF8L | 4 | TTAATAAA/TTTATTAA | 7 |
| --- | | | | | | | | 0 |
| chr17 | 65047940 | GTAAATCTCTGTG-TTAATGAA-TACACTATCTTAT | loxF8L | 4 | loxF8R | 3 | TTAATGAA/TTCATTAA | 7 |
| chr8 | 15308153 | CAACGATGGTGTG-TTAATGAA-TCCATACAAATAT | loxF8R | 3 | loxF8L | 4 | TTAATGAA/TTCATTAA | 7 |
| --- | | | | | | | | 0 |
| chr1 | 115921406 | ATATATATGTATA-TTACATAA-TCCACATGCTTAT | loxF8L | 4 | loxF8L | 3 | TTACATAA/TTATGTAA | 7 |
| chr5 | 165831759 | CTGATGTTCTGTG-TTACATAA-TACTCAGATATAT | loxF8R | 4 | loxF8L | 3 | TTACATAA/TTATGTAA | 7 |
| chr6 | 136931640 | CTCAGATTGTCAG-TTACATAA-AACAAAGTCTTTG | loxF8R | 3 | loxF8R | 4 | TTACATAA/TTATGTAA | 7 |
| --- | | | | | | | | 0 |
| chr11 | 99406336 | AGAATTCTCTGGA-TTACCAAA-CAGACATTCTTGT | loxF8L | 3 | loxF8R | 4 | TTACCAAA/TTTGGTAA | 7 |
| chr16 | 58361080 | AGAATTCTCTGGA-TTACCAAA-CAGAGATTCTTAT | loxF8L | 3 | loxF8R | 4 | TTACCAAA/TTTGGTAA | 7 |
| --- | | | | | | | | 0 |
| chr5 | 30677381 | TTAAATCTTTTGA-TTACTGAA-ACCACTGATCTTT | loxF8L | 3 | loxF8L | 4 | TTACTGAA/TTCAGTAA | 7 |
| chr22 | 40281377 | GGAAATTTGTTGA-TTACTGAA-TGCACAGAATGAT | loxF8L | 4 | loxF8L | 3 | TTACTGAA/TTCAGTAA | 7 |
| --- | | | | | | | | 0 |
| chr2 | 26797662 | ATAAATTTGTGTT-TTATCTAA-GCCACTGAGTTGT | loxF8L | 3 | loxF8L | 4 | TTAGATAA/TTATCTAA | 7 |
| chrX | 16076892 | TTAAATCTAGGGA-TTAGATAA-GGCAAAGGTTTAT | loxF8L | 3 | loxF8L | 4 | TTAGATAA/TTATCTAA | 7 |
| --- | | | | | | | | 0 |
| chrX | 967276 | ATAAATTTGTATA-TTATATAA-TTTACAGAATTTT | loxF8L | 3 | loxF8L | 4 | TTATATAA/TTATATAA | 7 |
| chrY | 967276 | ATAAATTTGTATA-TTATATAA-TTTACAGAATTTT | loxF8L | 3 | loxF8L | 4 | TTATATAA/TTATATAA | 7 |
| --- | | | | | | | | 0 |
| chr10 | 17883585 | ATGAGATTGTTTG-TTCATGAA-TACACAGATAATT | loxF8R | 3 | loxF8L | 4 | TTCATGAA/TTCATGAA | 7 |
| chr4 | 20929887 | ATAATATTCTGTG-TTCATGAA-TCAAAAAATTTAA | loxF8R | 3 | loxF8L | 4 | TTCATGAA/TTCATGAA | 7 |
| --- | | | | | | | | 0 |
| chr6 | 49934985 | ATAAATCTGTCAC-TTCCAAAA-TCCCCACATATCT | loxF8L | 3 | loxF8L | 4 | TTCCAAAA/TTTTGGAA | 7 |
| chr1 | 228698999 | CTCTGATTTTGTT-TTCCAAAA-GACCCAATCTCAG | loxF8R | 4 | loxF8R | 3 | TTCCAAAA/TTTTGGAA | 7 |
| chr15 | 97218722 | TTAAGAATGTGAG-TTTTGGAA-TAGACAGATCTAA | loxF8R | 3 | loxF8L | 4 | TTCCAAAA/TTTTGGAA | 7 |
| --- | | | | | | | | 0 |
| chr11 | 32404895 | CTAACTCTGTGGT-TTCCAGAA-TCCACTGGGTTTT | loxF8L | 3 | loxF8L | 4 | TTCCAGAA/TTCTGGAA | 7 |
| chrY | 14139133 | ATAAATCTCAATA-TTCTGGAA-TCCAAATATTTAC | loxF8L | 4 | loxF8L | 3 | TTCCAGAA/TTCTGGAA | 7 |
| --- | | | | | | | | 0 |
| chr10 | 110967513 | ACAATTCAGTGGT-TTTTAGAA-TATTCAGATTTAT | loxF8L | 4 | loxF8L | 3 | TTCTAAAA/TTTTAGAA | 7 |
| chr4 | 35598847 | CTCTAACTGTGGA-TTCTAAAA-TTCAAAGAATTAT | loxF8L | 4 | loxF8L | 3 | TTCTAAAA/TTTTAGAA | 7 |
| chr12 | 56123364 | CTAACATTGTGGA-TTTTAGAA-ACAACACATTGAT | loxF8R | 3 | loxF8L | 4 | TTCTAAAA/TTTTAGAA | 7 |
| --- | | | | | | | | 0 |
| chr4 | 77071854 | AAAAAGCTGAGTA-TTTGTAAA-ACCAAAGATTTAA | loxF8L | 4 | loxF8L | 3 | TTTACAAA/TTTGTAAA | 7 |
| chr1 | 73584961 | TAAAAAGTGTGGA-TTTACAAA-CACAAAATATGAG | loxF8L | 4 | loxF8R | 3 | TTTACAAA/TTTGTAAA | 7 |
