## Supplementary Table 3 for "Correction of a Factor VIII genomic inversion with designer-recombinases"

| PDF page # | Chrom. | Peak coordinates | | Summit position (center of GB views) | -log10(p-value) |
|---|---|---|---|---|---|
| 1 | chr12 | 48.157.417 | 48.157.690 | 48.157.547 | 5,87 |
| 2 | chr5 | 27.241.681 | 27.242.246 | 27.242.082 | 5,71 |
| 3 | chr5 | 36.369.195 | 36.369.550 | 36.369.421 | 4,69 |
| 4 | chr3 | 179.015.562 | 179.015.925 | 179.015.756 | 4,59 |
| 5 | chr3 | 195.102.719 | 195.103.029 | 195.102.902 | 4,59 |
| 6 | chr14 | 71.674.583 | 71.674.789 | 71.674.656 | 4,48 |
| 7 | chr1 | 36.217.612 | 36.217.941 | 36.217.772 | 4,41 |
| 8 | chr2 | 46.461.481 | 46.461.755 | 46.461.643 | 4,21 |
| 9 | chr9 | 34.771.050 | 34.771.434 | 34.771.259 | 4,17 |
| 10 | chr22 | 33.114.481 | 33.114.871 | 33.114.676 | 4,13 |
| 11 | chr5 | 13.817.511 | 13.817.793 | 13.817.638 | 4,08 |
| 12 | chr7 | 5.856.411 | 5.856.676 | 5.856.537 | 4,08 |
| 13 | chr8 | 143.454.719 | 143.454.963 | 143.454.816 | 4,08 |
| 14 | chr12 | 25.829.046 | 25.829.500 | 25.829.259 | 4,04 |
| 15 | chr3 | 182.794.686 | 182.795.018 | 182.794.815 | 4,04 |
| 16 | chr11 | 102.119.960 | 102.120.257 | 102.120.099 | 3,95 |
| 17 | chr11 | 95.245.011 | 95.245.318 | 95.245.129 | 3,95 |
| 18 | chr5 | 32.642.619 | 32.643.038 | 32.642.847 | 3,95 |
| 19 | chr7 | 22.561.263 | 22.561.577 | 22.561.374 | 3,95 |
| 20 | chr5 | 14.908.143 | 14.908.418 | 14.908.301 | 3,91 |
| 21 | chr3 | 118.531.239 | 118.531.527 | 118.531.357 | 3,91 |
| 22 | chr5 | 27.316.388 | 27.316.633 | 27.316.517 | 3,91 |
| 23 | chr11 | 74.288.108 | 74.288.368 | 74.288.193 | 3,86 |
| 24 | chr12 | 10.182.691 | 10.182.982 | 10.182.827 | 3,86 |
| 25 | chr6 | 10.177.752 | 10.178.063 | 10.177.921 | 3,86 |
| 26 | chr6 | 7.998.080 | 7.998.389 | 7.998.245 | 3,86 |
| 27 | chr10 | 102.963.319 | 102.963.631 | 102.963.488 | 3,81 |
| 28 | chr2 | 240.069.999 | 240.070.289 | 240.070.139 | 3,81 |
| 29 | chr7 | 137.667.744 | 137.668.097 | 137.667.972 | 3,81 |
| 30 | chr11 | 27.331.972 | 27.332.247 | 27.332.093 | 3,76 |
| 31 | chr1 | 32.769.319 | 32.769.614 | 32.769.538 | 3,76 |
| 32 | chr17 | 40.813.853 | 40.814.198 | 40.814.066 | 3,76 |
| 33 | chr2 | 43.426.384 | 43.426.765 | 43.426.607 | 3,72 |
| 34 | chr3 | 261.060 | 261.322 | 261.179 | 3,72 |
| 35 | chr10 | 60.473.598 | 60.473.867 | 60.473.758 | 3,70 |
| 36 | chr5 | 123.454.639 | 123.454.954 | 123.454.759 | 3,66 |
| 37 | chr5 | 143.544.004 | 143.544.290 | 143.544.143 | 3,66 |
| 38 | chr6 | 4.313.227 | 4.313.583 | 4.313.459 | 3,66 |
| 39 | chr12 | 12.361.974 | 12.362.269 | 12.362.104 | 3,61 |
| 40 | chr6 | 9.127.264 | 9.127.600 | 9.127.455 | 3,61 |
| 41 | chr7 | 47.604.997 | 47.605.281 | 47.605.232 | 3,61 |
| 42 | chr2 | 100.381.171 | 100.381.395 | 100.381.315 | 3,56 |
| 43 | chr5 | 33.343.661 | 33.344.150 | 33.343.780 | 3,56 |
| 44 | chr7 | 18.974.778 | 18.975.084 | 18.974.958 | 3,56 |
| 45 | chrX | 17.032.370 | 17.032.737 | 17.032.483 | 3,56 |

| | | | | | |
|---|---|---|---|---|---|
| 46 | chr10 | 60.955.983 | 60.956.383 | 60.956.147 | 3,51 |
| 47 | chr12 | 131.114.803 | 131.115.076 | 131.114.948 | 3,51 |
| 48 | chr15 | 100.873.083 | 100.873.365 | 100.873.188 | 3,51 |
| 49 | chr15 | 99.213.193 | 99.213.418 | 99.213.330 | 3,51 |
| 50 | chr17 | 57.480.917 | 57.481.221 | 57.481.061 | 3,51 |
| 51 | chr3 | 32.045.089 | 32.045.307 | 32.045.205 | 3,51 |
| 52 | chr10 | 4.663.220 | 4.663.520 | 4.663.318 | 3,45 |
| 53 | chr14 | 75.230.535 | 75.230.810 | 75.230.672 | 3,45 |
| 54 | chr3 | 187.017.019 | 187.017.290 | 187.017.123 | 3,45 |
| 55 | chr6 | 2.237.119 | 2.237.442 | 2.237.302 | 3,45 |
| 56 | chr8 | 118.850.186 | 118.850.431 | 118.850.271 | 3,45 |
| 57 | chr12 | 46.473.878 | 46.474.158 | 46.474.011 | 3,39 |
| 58 | chr19 | 14.337.395 | 14.337.729 | 14.337.610 | 3,39 |
| 59 | chr5 | 16.493.901 | 16.494.133 | 16.493.984 | 3,39 |
| 60 | chr6 | 7.730.724 | 7.730.956 | 7.730.831 | 3,39 |
| 61 | chr8 | 118.961.994 | 118.962.381 | 118.962.167 | 3,39 |
| 62 | chr11 | 10.122.793 | 10.123.241 | 10.123.064 | 3,34 |
| 63 | chr1 | 220.426.909 | 220.427.252 | 220.427.003 | 3,34 |
| 64 | chr1 | 87.930.169 | 87.930.421 | 87.930.296 | 3,34 |
| 65 | chr1 | 97.093.875 | 97.094.536 | 97.093.930 | 3,34 |
| 66 | chr22 | 28.712.518 | 28.712.771 | 28.712.700 | 3,34 |
| 67 | chr3 | 22.106.343 | 22.106.839 | 22.106.505 | 3,34 |
| 68 | chr10 | 97.407.314 | 97.407.635 | 97.407.430 | 3,28 |
| 69 | chr5 | 149.113.657 | 149.113.919 | 149.113.819 | 3,28 |
| 70 | chr7 | 22.696.845 | 22.697.116 | 22.696.969 | 3,28 |
| 71 | chr10 | 15.106.196 | 15.106.647 | 15.106.434 | 3,22 |
| 72 | chr10 | 94.963.966 | 94.964.294 | 94.964.056 | 3,22 |
| 73 | chr1 | 100.502.299 | 100.502.713 | 100.502.562 | 3,22 |
| 74 | chr5 | 13.881.729 | 13.882.036 | 13.881.886 | 3,22 |
| 75 | chr9 | 116.232.038 | 116.232.550 | 116.232.235 | 3,22 |
| 76 | chr9 | 125.807.105 | 125.807.823 | 125.807.302 | 3,22 |
| 77 | chrX | 11.447.071 | 11.447.320 | 11.447.213 | 3,22 |
| 78 | chr5 | 17.340.730 | 17.341.211 | 17.340.869 | 3,15 |
| 79 | chr5 | 7.583.616 | 7.584.003 | 7.583.833 | 3,15 |
| 80 | chr10 | 99.435.775 | 99.436.171 | 99.436.031 | 3,09 |
| 81 | chr11 | 94.700.513 | 94.700.994 | 94.700.867 | 3,09 |
| 82 | chr11 | 95.521.062 | 95.521.465 | 95.521.315 | 3,09 |
| 83 | chr12 | 46.430.288 | 46.430.569 | 46.430.369 | 3,09 |
| 84 | chr8 | 137.149.494 | 137.149.914 | 137.149.774 | 3,09 |
| 85 | chr7 | 115.230.957 | 115.231.376 | 115.231.257 | 2,95 |
