## Supplementary Table 4 for "Correction of a Factor VIII genomic inversion with designer-recombinases"

| Primer count | Primer name | Primer seq. | Description |
|---|---|---|---|
| 1 | eGFP_F | GCTAATACGACTCACTATAGGGAGAGCCGCCACCATGCCAAAAAAGAAGAG | Primer to generate DNA template for IVT |
| 2 | eGFP_R | TTTTTTTTTTTTTTTTGGTTTATTCCTAGTACAGCTCGTCCATGCC | Primer to generate DNA template for IVT |
| 3 | mCherry_F | GCTAATACGACTCACTATAGGGAGAGATGGTGAGCAAGGGCG | Primer to generate DNA template for IVT |
| 4 | mCherry_R | TTTTTTTTTTTTTTTTGGTTTATTCTTACTTGTACAGCTCGTCC | Primer to generate DNA template for IVT |
| 5 | tagBFP_F | GCTAATACGACTCACTATAGGGAGAGATGAGCGAGCTGATTAAGGAGA | Primer to generate DNA template for IVT |
| 6 | tagBFP_R | TTTTTTTTTTTTTTTTGGTTTATTCTTAATTAAGCTTGTGCCCCAGTT | Primer to generate DNA template for IVT |
| 7 | D71st_F | GCTAATACGACTCACTATAGGGAGAGCCGCCACCATGCCAAAAAAGAAGAGAAAGGTAATGAGCAACCTGCAGACCC | Primer to generate DNA template for IVT |
| 8 | D71st_R | TTTTTTTTTTTTTTTTGGTTTATTCTCAATCGCCGTCCTCAAGGA | Primer to generate DNA template for IVT |
| 9 | D72nd_F | GCTAATACGACTCACTATAGGGAGAGCCGCCACCATGCCAAAAAAGAAGAGAAAGGTAATGTCCAATATCCAAACACCTCACC | Primer to generate DNA template for IVT |
| 10 | D72nd_R | TTTTTTTTTTTTTTTTGGTTTATTCTCAGTCGGAATCTTCCAGCAGCC | Primer to generate DNA template for IVT |
| 11 | L8_F | GCTAATACGACTCACTATAGGGAGAGCCGCCACCATGCCAAAAAAGAAGAGAAAGGTAATGTCCAATATCCAAACACCTCACC | Primer to generate DNA template for IVT |
| 12 | L8_R | TTTTTTTTTTTTTTTTGGTTTATTCTCAATCGCCGTCCTCAAGGA | Primer to generate DNA template for IVT |
| 13 | qPCR_F8_F | ggaactgtcatgggactatatgc | Primer used for qPCR to detect F8 transcript Ex1-Ex2 boundary |
| 14 | qPCR_F8_R | gcactctaggaggaaatcttgc | Primer used for qPCR to detect F8 transcript Ex1-Ex2 boundary |
| 15 | 1stF8_F | TCCTGGTAAAGGTGGGACTAT | Primer for inversion orientation detection |
| 16 | 1stF8_R | CAGAGGCTAGCTGATGTGAAT | Primer for inversion orientation detection |
| 17 | 2ndF8_F | GCAGGGATCTTGTTGGTAA | Primer for inversion orientation detection |
| 18 | 2ndF8_R | GGATGAGAAGGGTACACAGAAA | Primer for inversion orientation detection |
| 19 | HG2_ch15_F | GCTATTAATGTTCCCCTTAGCC | Detection of HG2 off-target activity |
| 20 | HG2_ch15_R | CTCAAGGACAGACTACATATTAGGC | Detection of HG2 off-target activity |
| 21 | HG2_ch7_F | TCCAGGATAGATCATATTTTAGGCC | Detection of HG2 off-target activity |
| 22 | HG2_ch7_R | GATGCAGCATACCAACATGG | Detection of HG2 off-target activity |
| 23 | peak1_F | ATCGagatctCGTGACCCGGAAGCAGAAGCTGTCGGGGGCAGGCCCTCTGTTTACCGGACTAGAAGACCAACTAAGATGGGCGCCTGCGCACTGATGAGTGgcttgcatgcctgcagatc | Cloning of ChipSeq peak to pEVO vector for rec. assay |
| 24 | peak1_R | ATCGagatctCACTCATCAGTGCGCAGGCGCCCATCTTAGTTGGTCTTCTAGTCCGGTAAACAGAGGGCCTGCCCCCGACAGCTTCTGCTTCCGGGTCACGagtaggtctctgcagtgtcg | Cloning of ChipSeq peak to pEVO vector for rec. assay |
| 25 | peak2_F | ATCGagatctGAAGTTGCAAAATAACAGATGCTAGTGAGGCTCTGGAGAAAAAGAAATGCTTATACACTGTTGGTAGGAATGTAAATTACTTCAGTCACTGgcttgcatgcctgcagatc | Cloning of ChipSeq peak to pEVO vector for rec. assay |
| 26 | peak2_R | ATCGagatctCAGTGACTGAAGTAATTTACATTCCTACCAACAGTGTATAAGCATTTCTTTTTCTCCAGAGCCTCACTAGCATCTGTTATTTTGCAACTTCagtaggtctctgcagtgtcg | Cloning of ChipSeq peak to pEVO vector for rec. assay |
| 27 | peak3_F | ATCGagatctAAAGGGAGTCATCCAAGAATAGACGCTAAAAGCCTTTTGATGGGAATAGGAAATAGGGAATCTGATTTCAGGTAAGCCTATGGGAATTTCCgcttgcatgcctgcagatc | Cloning of ChipSeq peak to pEVO vector for rec. assay |
| 28 | peak3_R | ATCGagatctGGAAATTCCCATAGGCTTACCTGAAATCAGATTCCCTATTTCCTATTCCCATCAAAAGGCTTTTAGCGTCTATTCTTGGATGACTCCCTTTagtaggtctctgcagtgtcg | Cloning of ChipSeq peak to pEVO vector for rec. assay |
| 29 | peak4_F | ATCGagatctGGTGGCACTAGATACCGTTGGTCAAAGTCCATAACCTAGTAAGGATTTCTAGAAAACAGGGTTAGAGAGGTGCCTCTGGCATCAGCAAAGAgcttgcatgcctgcagatc | Cloning of ChipSeq peak to pEVO vector for rec. assay |
| 30 | peak4_R | ATCGagatctTCTTTGCTGATGCCAGAGGCACCTCTCTAACCCTGTTTTCTAGAAATCCTTACTAGGTTATGGACTTTGACCAACGGTATCTAGTGCCACCagtaggtctctgcagtgtcg | Cloning of ChipSeq peak to pEVO vector for rec. assay |
| 31 | peak5_F | ATCGagatctGTTCTAGTTTTATTTCTTTAGGAATTTCCATATTGTTTTCCATCATGGCTGTACCAACCTACATTCCTACCAACGGTGCAGCGTGCTGGGGgcttgcatgcctgcagatc | Cloning of ChipSeq peak to pEVO vector for rec. assay |
| 32 | peak5_R | ATCGagatctCCCCAGCACGCTGCACCGTTGGTAGGAATGTAGGTTGGTACAGCCATGATGGAAAACAATATGGAAATTCCTAAAGAAATAAAACTAGAACagtaggtctctgcagtgtcg | Cloning of ChipSeq peak to pEVO vector for rec. assay |
| 33 | peak6_F | ATCGagatctCTGGAGTGCAGTGGCGGGGTCTCGGCTCACTGCAAGCTCCGCCTCCCGGGTTCACGCCATTCTCCTGCCTCAGCCTCCCAAGTAGCTGGGAgcttgcatgcctgcagatc | Cloning of ChipSeq peak to pEVO vector for rec. assay |
| 34 | peak6_R | ATCGagatctTCCCAGCTACTTGGGAGGCTGAGGCAGGAGAATGGCGTGAACCCGGGAGGCGGAGCTTGCAGTGAGCCGAGACCCCGCCACTGCACTCCAGagtaggtctctgcagtgtcg | Cloning of ChipSeq peak to pEVO vector for rec. assay |
| 35 | peak7_F | ATCGagatctTGTTACTATTGTAATTGTTTGGCAGTGCCACATACTGCACCCATATAAGTGGTAAACTTAGTAAGTGTTGTGTGTGTACTGACTGCTCCACgcttgcatgcctgcagatc | Cloning of ChipSeq peak to pEVO vector for rec. assay |
| 36 | peak7_R | ATCGagatctGTGGAGCAGTCAGTACACACACAACACTTACTAAGTTTACCACTTATATGGGTGCAGTATGTGGCACTGCCAAACAATTACAATAGTAACAagtaggtctctgcagtgtcg | Cloning of ChipSeq peak to pEVO vector for rec. assay |
| 37 | peak8_F | ATCGagatctACGATCTTGGGCCCAGGTTTCTAAAGCAAACTTGAAAATCAACCAGGGGAGGGCCATGAAGAAAATTACTCTTGAATGGCAAGAAATCAATgcttgcatgcctgcagatc | Cloning of ChipSeq peak to pEVO vector for rec. assay |
| 38 | peak8_R | ATCGagatctATTGATTTCTTGCCATTCAAGAGTAATTTTCTTCATGGCCCTCCCCTGGTTGATTTTCAAGTTTGCTTTAGAAACCTGGGCCCAAGATCGTagtaggtctctgcagtgtcg | Cloning of ChipSeq peak to pEVO vector for rec. assay |
| 39 | peak9_F | ATCGagatctTTGTGCCAGCATCATAGACTGTATTTACACAAACCTAGATGGTATAGCCTACTACATGCTGAGGCTATATATTGCGTAGTCTATTGCTGCTgcttgcatgcctgcagatc | Cloning of ChipSeq peak to pEVO vector for rec. assay |
| 40 | peak9_R | ATCGagatctAGCAGCAATAGACTACGCAATATATAGCCTCAGCATGTAGTAGGCTATACCATCTAGGTTTGTGTAAATACAGTCTATGATGCTGGCACAAagtaggtctctgcagtgtcg | Cloning of ChipSeq peak to pEVO vector for rec. assay |
| 41 | peak10_F | ATCGagatctAATGTGTTTCTATGCCAGGTTCACATTTAACTGAGTTAAACATAGTGGGTGTGAACTCTGGCATTGGCAGATAGTGGCTCTGTGGCCTTAGgcttgcatgcctgcagatc | Cloning of ChipSeq peak to pEVO vector for rec. assay |
| 42 | peak10_R | ATCGagatctCTAAGGCCACAGAGCCACTATCTGCCAATGCCAGAGTTCACACCCACTATGTTTAACTCAGTTAAATGTGAACCTGGCATAGAAACACATTagtaggtctctgcagtgtcg | Cloning of ChipSeq peak to pEVO vector for rec. assay |
| 43 | peak20_F | ATCGagatctCCCGGCCTGGTATATTTTTACCCAAGGACTAGATTATTAGTTGCTAGTGGGTAAGGAAAACAACTTAAAATTCCTTTGCAAGTTGTCTTTCgcttgcatgcctgcagatc | Cloning of ChipSeq peak to pEVO vector for rec. assay |
| 44 | peak20_R | ATCGagatctGAAAGACAACTTGCAAAGGAATTTTAAGTTGTTTTCCTTACCCACTAGCAACTAATAATCTAGTCCTTGGGTAAAAATATACCAGGCCGGGagtaggtctctgcagtgtcg | Cloning of ChipSeq peak to pEVO vector for rec. assay |
| 45 | peak68_F | ATCGagatctACCAACTAGTAGACTGGGATTTTGCAAATCAGATGCGGAGATGCAGAGGCCAAGGGTAGTGGCAGATGTCCCATCCTTTTCCTGGGCTACAgcttgcatgcctgcagatc | Cloning of ChipSeq peak to pEVO vector for rec. assay |
| 46 | peak68_R | ATCGagatctTGTAGCCCAGGAAAAGGATGGGACATCTGCCACTACCCTTGGCCTCTGCATCTCCGCATCTGATTTGCAAAATCCCAGTCTACTAGTTGGTagtaggtctctgcagtgtcg | Cloning of ChipSeq peak to pEVO vector for rec. assay |
| 47 | qPCR F8 mRNA F | GGAACTGTCATGGGACTATATGC | Binding in exon1 |
| 48 | qPCR F8 mRNA R | ACAAACAGAGTCTTTTTGTACACG | Binding in exon1 |
| 49 | off_L8_qPCR_F_1 | CTGATGAGTGTTCGTCAGCG | qPCR peak validation |
| 50 | off_L8_qPCR_R_1 | GCGCAAAGCCTCTGAGTAAA | qPCR peak validation |
| 51 | off_L8_qPCR_F_2 | GCAAGTCGAAACCACCATGAG | qPCR peak validation |
| 52 | off_L8_qPCR_R_2 | TCTCCAGAGCCTCACTAGCA | qPCR peak validation |
| 53 | off_L8_qPCR_F_3 | TGGGCTGGTTGCCCAATAAA | qPCR peak validation |
| 54 | off_L8_qPCR_R_3 | GCCCACCATGATGTGTCACT | qPCR peak validation |
| 55 | off_L8_qPCR_F_4 | TAGACAGTCCACAAGGTGGC | qPCR peak validation |
| 56 | off_L8_qPCR_R_4 | ATGCCAGAGGCACCTCTCTA | qPCR peak validation |
| 57 | off_L8_qPCR_F_5 | ATGCCCAGAAGAAGGACTGC | qPCR peak validation |
| 58 | off_L8_qPCR_R_5 | AGATAGCCACTGTTGGCGAG | qPCR peak validation |
| 59 | off_L8_qPCR_F_6 | CTCATAACAACCCTCTCTTGTGA | qPCR peak validation |
| 60 | off_L8_qPCR_R_6 | CAGAGCGAGACTCCGTCTAA | qPCR peak validation |
| 61 | off_L8_qPCR_F_7 | AGTGCCACATACTGCACCCA | qPCR peak validation |
| 62 | off_L8_qPCR_R_7 | ACAGGGAATGGCCAGTTGGT | qPCR peak validation |
| 63 | off_L8_qPCR_F_8 | GTACTGCTGCCAGAGAGACG | qPCR peak validation |
| 64 | off_L8_qPCR_R_8 | GCCCTCCCCTGGTTGATTTT | qPCR peak validation |
| 65 | off_L8_qPCR_F_9 | TGTGCCAGCATCATAGACTGT | qPCR peak validation |
| 66 | off_L8_qPCR_R_9 | CCTAGCAGCAATAGACTACGCA | qPCR peak validation |
| 67 | off_L8_qPCR_F_10 | TGTGTTTCTATGCCAGGTTCACA | qPCR peak validation |
| 68 | off_L8_qPCR_R_10 | CCACAGAGCCACTATCTGCC | qPCR peak validation |
| 69 | off_L8_qPCR_F_20 | TGCTAGTGGGTAAGGAAAACAACT | qPCR peak validation |
| 70 | off_L8_qPCR_R_20 | ACGGTAAGTATGGGCTGCAC | qPCR peak validation |
| 71 | off_L8_qPCR_F_68 | GATGCAGAGGCCAAGGGTAG | qPCR peak validation |
| 72 | off_L8_qPCR_R_68 | CAATGCCCATACCAGGCTGA | qPCR peak validation |
| 73 | off_L8_qPCR_F_HG2 | GCGTTTTTGTTTTCATTCATCTCCA | qPCR peak validation |
| 74 | off_L8_qPCR_R_HG2 | TGGGACATTCTCAAGGACAGAC | qPCR peak validation |
| 75 | off_L8_qPCR_F_offsym2L | CAGATTTGTGCATCTGCTCTTTGA | qPCR peak validation |
| 76 | off_L8_qPCR_R_offsym2L | TTGTGCAATCTCAAACTCCAAACTA | qPCR peak validation |
| 77 | qPCR_F8_F | ATCACCCAAGGAAGCACATAAGTT | qPCR peak validation |
| 78 | qPCR_F8_R | CCCTCAGTTACCACAGCGTT | qPCR peak validation |
