## Supplementary Table 5 for "Correction of a Factor VIII genomic inversion with designer-recombinases"

| Count | Cycling Protocol | Temperature (°C) | Time (s) | Cycles |
|---|---|---|---|---|
| **1** | F8 inversion PCR | 94 | 45 | |
| | | 94 | 20 | 40x |
| | | 52 | 20 | |
| | | 72 | 45 | |
| | | 72 | 300 | |
| | | 8 | infinite | |
| **2** | qPCR to qunatify the inversion | 94 | 180 | |
| | | 94 | 20 | 50x |
| | | 56 | 20 | |
| | | 72 | 180 | |
| **3** | In vitro transcription | 37 | 30 | |
| | | 8 | pause | |
| | | 37 | 15 | add Dnase |
| | | 8 | pause | |
| | | 37 | 30 | add poly(A) polymerase |
| | | 8 | infinite | |
| **4** | Peak insert generation | 94 | 15 | 30x |
| | | 58 | 15 | |
| | | 72 | 30 | |
| | | 72 | 300 | |
| | | 8 | infinite | |
| **5** | HG2 translocation | 94 | 15 | 30x |
| | | 56 | 15 | |
| | | 72 | 20 | |
| | | 72 | 300 | |
| | | 8 | infinite | |
